# ComMet: A method for comparing metabolic states in genome-scale metabolic models

**DOI:** 10.1101/2020.09.14.296145

**Authors:** Chaitra Sarathy, Marian Breuer, Martina Kutmon, Michiel E. Adriaens, Chris T. Evelo, Ilja C.W. Arts

## Abstract

Being comprehensive knowledge bases of cellular metabolism, Genome-scale metabolic models (GEMs) serve as mathematical tools for studying cellular flux states in various organisms. However, analysis of large-scale (human) GEMs, still presents considerable challenges with respect to objective selection and reaction flux constraints. In this study, we introduce a model-based method, ComMet (Comparison of Metabolic states), for comprehensive analysis of large metabolic flux spaces and comparison of various metabolic states. ComMet allows (a) an in-depth characterisation of achievable flux states, (b) comparison of flux spaces from several conditions of interest and (c) identification and visualization of metabolically distinct network modules. As a proof-of-principle, we employed ComMet to extract the biochemical differences in the human adipocyte network (iAdipocytes1809) arising due to unlimited/blocked uptake of branched-chain amino acids. Our study opens avenues for exploring several metabolic conditions of interest in both microbe and human models. ComMet is open-source and is available at https://github.com/macsbio/commet.

## 1 INTRODUCTION

Metabolism plays a central role in maintaining cell functionality as it provides the energy and building blocks for cellular growth. In humans, metabolic dysfunction is associated with a wide range of clinical conditions including obesity, diabetes, neurodegenerative diseases, cancer and inborn errors of metabolism (Hotamisligil, 2006; Coller, 2014; Saudubray et al., 2016). Therefore, systems-level understanding of human metabolism is pivotal to comprehending phenotypic changes (in both normal and diseased states) and to develop prevention and treatment strategies.

Advancements in experimental and computational techniques have enabled the construction of genome-scale metabolic models (GEMs) in the last two decades. GEMs are mathematical formulations of the complete set of metabolic reactions taking place in a cell, tissue, organ or organism (Robinson and Nielsen, 2016). GEMs contain extensive descriptions of molecular relationships between genes, reactions and metabolites. These comprehensive knowledgebases enable prediction of reaction fluxes under varying environmental conditions thus contributing to systems-level understanding of metabolism. GEMs have facilitated investigating various metabolic dysfunctions in cancer (Folger et al., 2011; Agren et al., 2012; Yizhak et al., 2014, 2015), obesity (Mardinoglu et al., 2013) and non-alcoholic fatty liver disease (Mardinoglu et al., 2014, 2017). Despite successful applications and advances in algorithms (Lewis et al., 2012), conducting studies with human GEMs still requires certain assumptions and/or prerequisites which are detailed below.

Flux Balance Analysis (FBA) (Orth et al., 2010), Elementary Flux Modes (EFM) analysis (Schuster and Hilgetag, 1994) and Flux Space Sampling (or Sampling) (Schellenberger and Palsson, 2008) are frequently used methods for analysing GEMs. Given the cellular uptake rates of metabolites, FBA optimises an assumed objective (such as biomass production) and estimates flux values for all reactions. However, the accuracy of FBA estimates predominantly depends on two factors: (a) assumed objective and (b) precise description of media/nutrient levels. Due to the complex nature of human cellular metabolism, selecting the objective function is not as straightforward as biomass production and requires careful consideration of the underlying physiology. In addition, the absence of accurate public data from human cell line studies (describing uptake/release rates of plasma metabolites) limits the applicability of FBA. Alternative to identifying a single optimal flux distribution, EFM analysis and Sampling characterise all the possible flux states in the metabolic network. EFMs are non-decomposable steady-state pathways through a metabolic network. Owing to the combinatorial explosion in the number of EFMs, identification of the complete set of EFMs is computationally demanding (Klamt and Stelling, 2002), making it unsuitable for large GEMs. In addition, estimating the likelihood of observing an EFM in a given phenotype is difficult (Güell et al., 2015) which further limits their practical applicability on human GEMs. Sampling, on the other hand, provides a realistic alternative to EFMs in exploring the properties of network states. Sampling identifies the feasible range as well as a probability distribution for every reaction flux in the model by generating uniformly distributed random points in the flux space (a geometrical polytope containing the set of feasible metabolic states). Most importantly, unlike FBA, Sampling does not require the specification of an objective function. These methodological differences offer great benefits in using Sampling to assess metabolic differences in various physiological states. However, most of the currently available algorithms for Sampling the human GEMs are computationally intense.

In this study, we introduce ComMet (Comparison of Metabolic states), a method for comprehensive analysis and comparison of large metabolic flux spaces. ComMet provides a scalable and model-dependent framework that is computationally feasible and independent of objective specification. The functionalities available in ComMet allow in-depth characterisation of flux states achievable by GEMs followed by identification of metabolic differences between several conditions of interest. Characterisation of metabolic flux spaces involves identifying the key players and predicting flux statistics of the reactions active in that metabolic state. This identification is achieved in ComMet by combining two existing approaches. First, the iterative algorithm developed by Braunstein et al., (Braunstein et al., 2017) gives an analytical approximation of the probability distribution of fluxes instead of describing this probability distribution through sampling random points within the flux space. The flux predictions obtained through their approach are as accurate as conventional Sampling algorithms and involve very little processing times, making it suitable even for large GEMs. Second, in an earlier study Barrett et al. (Barrett et al., 2009) demonstrated that applying Principal Component Analysis (PCA) on a sampled flux space decomposes flux states into reaction sets whose variability accounts for the variation in the flux space. Such a transformation extracts what are called “modules” (or biochemical features characterising a given state) based on network-wide flux interactions and provides useful insights into the underlying physiology.

The novelty of ComMet lies in its ability to investigate differences between various metabolic states such as presence/absence of obesity. Metabolic features distinguishing the different conditions are extracted through rigorous optimisation of comparative strategies. The resulting distinctions are subsequently visualised in two network modes: reaction map and metabolic map. We demonstrate the applicability of ComMet on a large GEM, the metabolic reconstruction of human adipocyte, iAdipocytes1809 (Mardinoglu et al., 2013). We highlight the differences in the flux space of the adipocyte model arising due to presence or absence of branched-chain amino acids (BCAAs: leucine, valine and isoleucine). Elevated BCAAs are considered as strong biomarkers for obesity and diabetes (Felig et al., 1969; Newgard et al., 2009). Although a mechanistic explanation for the observed increased levels is currently unavailable, impaired BCAA catabolism in adipocytes has been hypothesised as a contributing factor (Herman et al., 2010). By extracting biochemically interpretable adipocyte modules, ComMet was able to provide additional insights into adipocyte-specific BCAA metabolism. We validated predictions from ComMet by identifying molecular processes that were functionally related to BCAA metabolism. Apart from this, we also highlight the utility of ComMet as a tool for generating hypotheses that could potentially be tested in a laboratory setting.

## 2 RESULTS

First, this section will briefly explain the workflow underlying ComMet which is followed by the description of modules identified from the adipocyte model (iAdipocytes1809 (Mardinoglu et al., 2013)). To demonstrate the value of ComMet for distinguishing between metabolic conditions, two metabolic states of an adipocyte were simulated: an unconstrained substrate uptake and blocked uptake of BCAAs. The simulated scenario demonstrates a proof of principle of ComMet’s ability to investigate metabolic differences in various metabolic states.

### 2.1 Development of the ComMet workflow

Starting with a GEM, ComMet describes an eight-step pipeline to analyse and compare metabolic flux spaces (Figure 1). All the preprocessing necessary for Sampling was performed in the first step (Figure 1, i). The second step involved specification of constraints necessary for studying metabolic states of interest (Figure 1, ii). To identify the differences between unconstrained and blocked uptake of BCAAs, two metabolic conditions were simulated: (a) Unconstrained substrate uptake (Figure 1, ii, green: where all the exchange metabolites, including the uptake of BCAAs, were kept unlimited) and (b) Constrained substrate uptake, (Figure 1, ii, purple: where only the uptake of leucine, valine and isoleucine were limited to zero). Specifying constraints for the metabolic states under study resulted in two condition-specific flux spaces each of which were decomposed into modules (sets of reactions having key contribution to the flux space) in the following manner. Sampling was carried out in both conditions (Figure 1, iii) using the algorithm developed by (Braunstein et al., 2017). Next, Principal Component Analysis (Figure 1, iv) and basis rotation (Figure 1, v) were applied to each flux space using the approach described by (Barrett et al., 2009). Such a decomposition determined the principal components (PCs or flux vectors) explaining the variation within each flux space. Subsequently, our analysis followed two separate directions to (a) identify condition-specific modules and (b) compare metabolic conditions.

**FIGURE 1.**
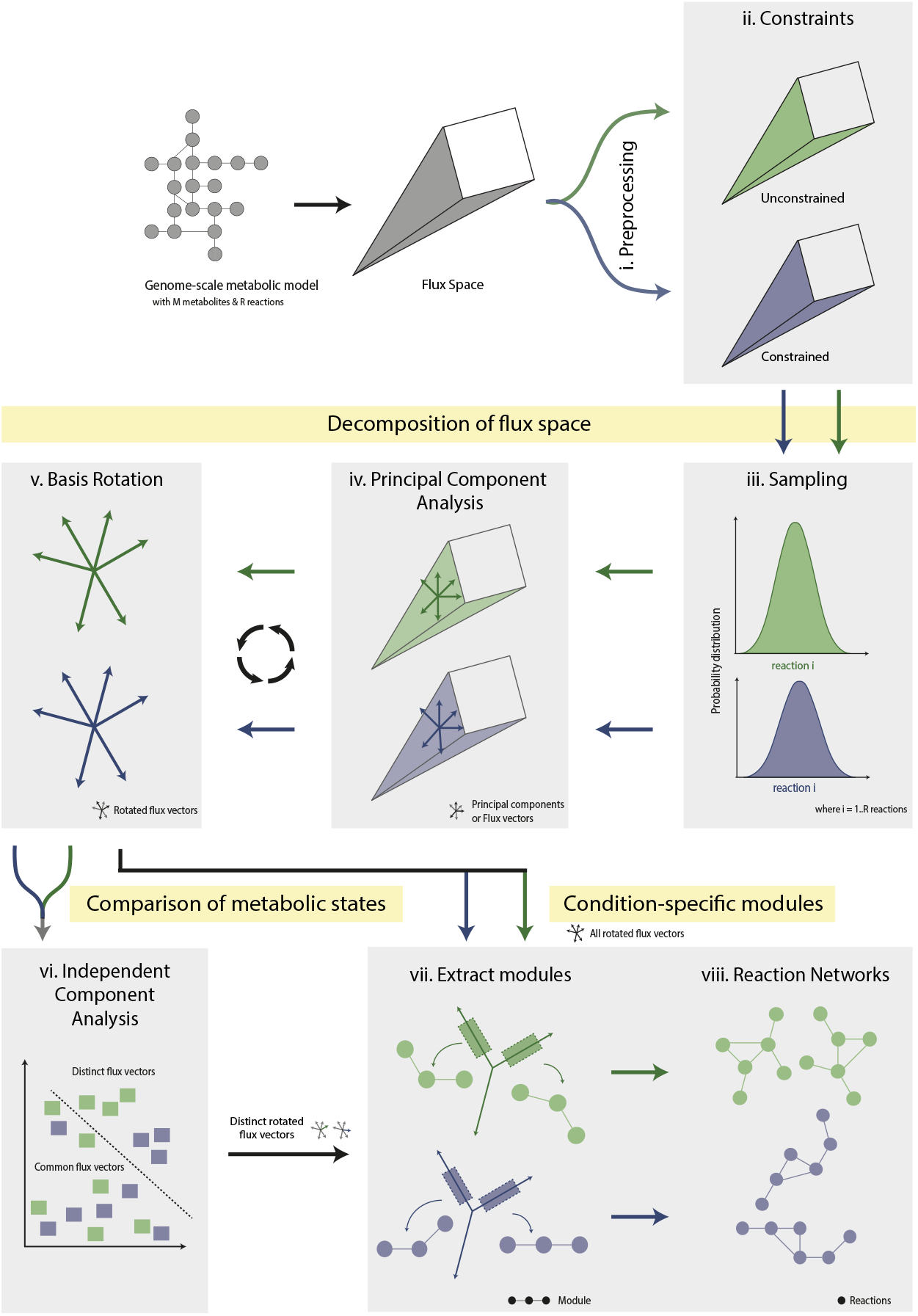
Overview of the eight-step pipeline in ComMet to compare various metabolic states in genome-scale metabolic models (GEMs). ComMet focuses on the flux space of an input GEM with M metabolites and R reactions. Once the GEM is preprocessed (**i**), reaction flux constraints are specified to generate condition-specific flux spaces ((**ii**) green and purple). Both flux spaces are then decomposed into modules through Sampling (**iii**) followed by Principal Component Analysis (**iv**) and basis rotation (**v**). The decomposed flux spaces can be studied individually or compared using Independent Component Analysis (**vi**) by extracting modules from all or distinct flux vectors respectively. Once the modules are extracted, they are visualised as a reaction network (**vii-viii**).

Essentially, the condition-specific modules contained sets of reactions whose fluxes contributed substantially in determining the underlying metabolic state. A module was extracted from each rotated PC which contained the most significant reactions within that vector (Figure 1, vii). The collection of modules from an individual flux space formed the “global modules” of that metabolic condition. To facilitate interpretation of modules and to identify the interplay between individual modules, the global modules were then visualised as reaction networks (Figure 1, viii). For comparing the two simulated metabolic conditions, the rotated PCs obtained previously were then subjected to Independent Component Analysis (ICA) which revealed vectors showing noticeable differences between the two conditions (Figure 1, vi). Finally, modules were extracted from these distinct flux vectors containing metabolic differences (Figure 1, vii) and were also visualised as a reaction map/network (Figure 1, viii). The entire analysis was carried out on a workstation running Windows 10 with E5-1650 6-core 3.5 GHz CPU and 32 GB RAM. The processing time for each step in ComMet’s pipeline is shown in Table S6.

### 2.2 Flux Space Sampling

During preprocessing, blocked reactions were defined as the reactions that are incapable of carrying any flux under the imposed conditions. The preprocessing step removed 2,043 blocked reactions thereby retaining 4,067 reactions. The 4,067 reactions formed a 1608-dimensional flux space. Subsequently, the (Braunstein et al., 2017) algorithm allowed exploring the probability distributions of reaction fluxes in both conditions. Mean and standard deviation of flux distributions were obtained for every reaction. A histogram analysis was first performed to broadly understand the impact of imposing constraints (Figure S1). The histograms of means from both conditions (Figure S1 A and B) were unimodal and centred around 1 mmol/gDW/h (millimoles per gram dry weight per hour, the unit of flux used in GEMs). Both graphs were roughly symmetric with long tails extending till +/− 1000 mmol/gDW/h. The histogram of standard deviations (Figure S1 C and D), on the other hand, were bimodal with a very large peak around 10 mmol/gDW/h and a smaller peak at 550 mmol/gDW/h. The shape and spread of the corresponding histograms (Figure S1 A vs B and Figure S1 C vs D) were notably similar between conditions. However, a closer inspection revealed differences.

In order to quantify the flux differences arising due to constraint imposition, next, a reaction-wise comparison of flux statistics was carried out. Figure 2 shows that a majority of the means and standard deviations lie on or close to, the identity line, suggesting a strong similarity in fluxes between the simulations. Deviations from the identity line seemed to indicate that the flux distributions of only a few reactions were visibly affected by limiting BCAA uptake. However, the mean fluxes were identical in only 412 reactions and the difference in means were between 0.01-10 mmol/gDW/h for 3,388 reactions. As detailed in Table S1, about 267 reactions showed a change in mean flux between 10-700 mmol/gDW/h. It is important to note that, as expected, the mean fluxes of several reactions involved in BCAA metabolism reduced to almost zero upon constraining BCAA uptake (black rectangle in Figure 2).

**FIGURE 2.**
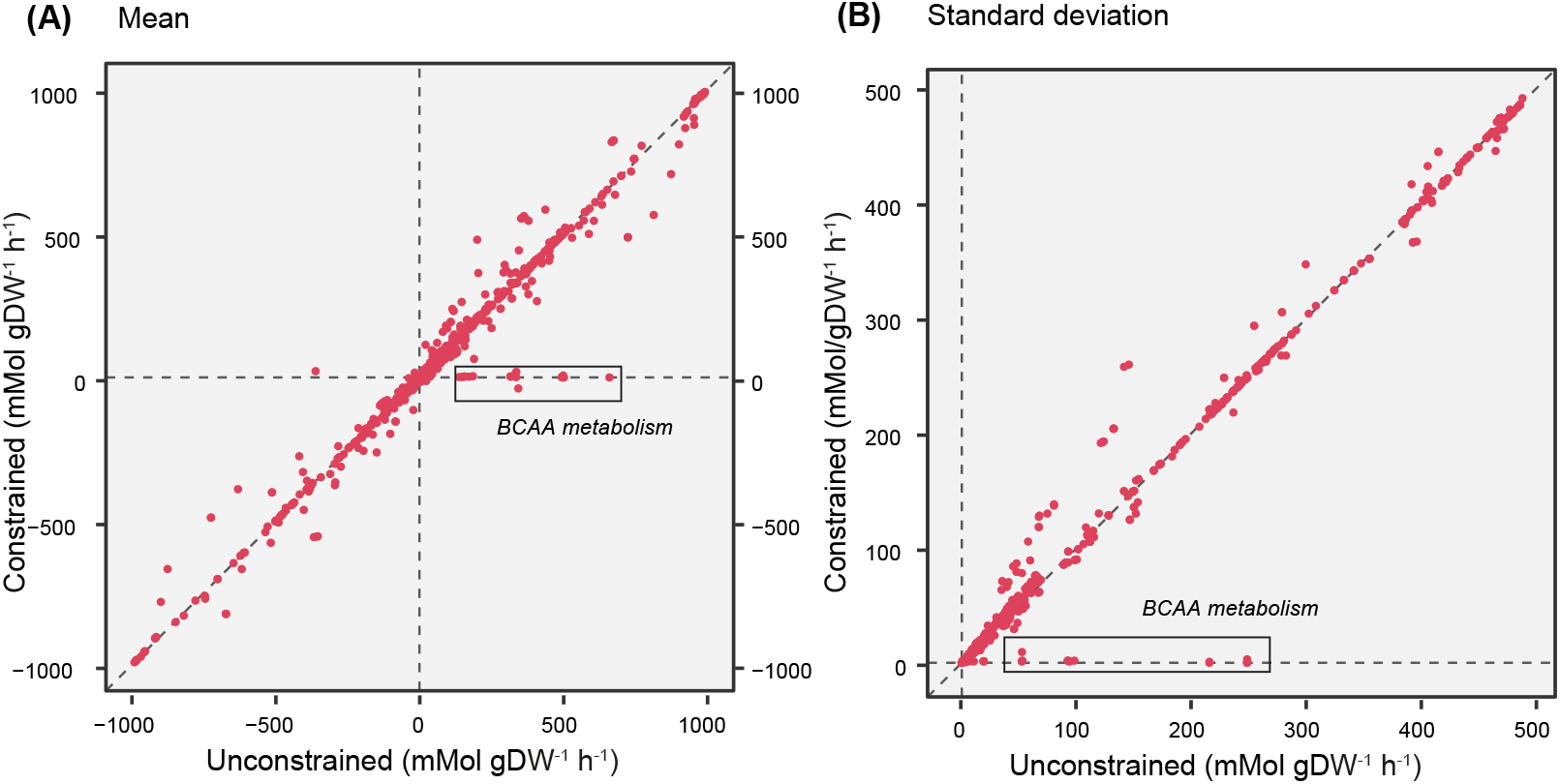
Reaction-wise comparison of (a) means and (b) standard deviations between the unconstrained and constrained simulations. The reactions involved in BCAA metabolism are highlighted in black rectangle.

### 2.3 Reactions with significantly different flux statistics

In order to evaluate the statistical significance of the flux differences, z-scores were computed using the means and standard deviations of reaction fluxes from both the conditions. Introduction of the constraint significantly affected the probability distributions through 21 reactions in the network (p < 0.05). The changes in flux distribution of all these reactions are shown using density plots (Figure 3, Figure S2 & Table S2). The shape of each distribution gives information about the sensitivity of the solution space to the applied constraint. As evident from Figures 3 (i-iii) & S2 (i-ix), in the constrained simulation, the flux distributions shifted to the left and spanned across a narrow range compared to a broad range in their unconstrained counterpart. This shift implied a reduction in the fluxes and as expected, this pattern was observed in several reactions in BCAA breakdown (Figures 3 (i-iii) & S2 (i-vi)). Additionally, the fluxes through the mitochondrial transport of intermediate metabolites involved in this pathway (isoleucine, methylmalonate and 3-methyl-2-oxobutyrate) also diminished (Fig S2, vii-ix). Application of the constraint also resulted in marginal release of BCAAs (not shown) the potential sources of which includes breakdown of lipoproteins that is taken up from the *in silico* media.

**FIGURE 3.**
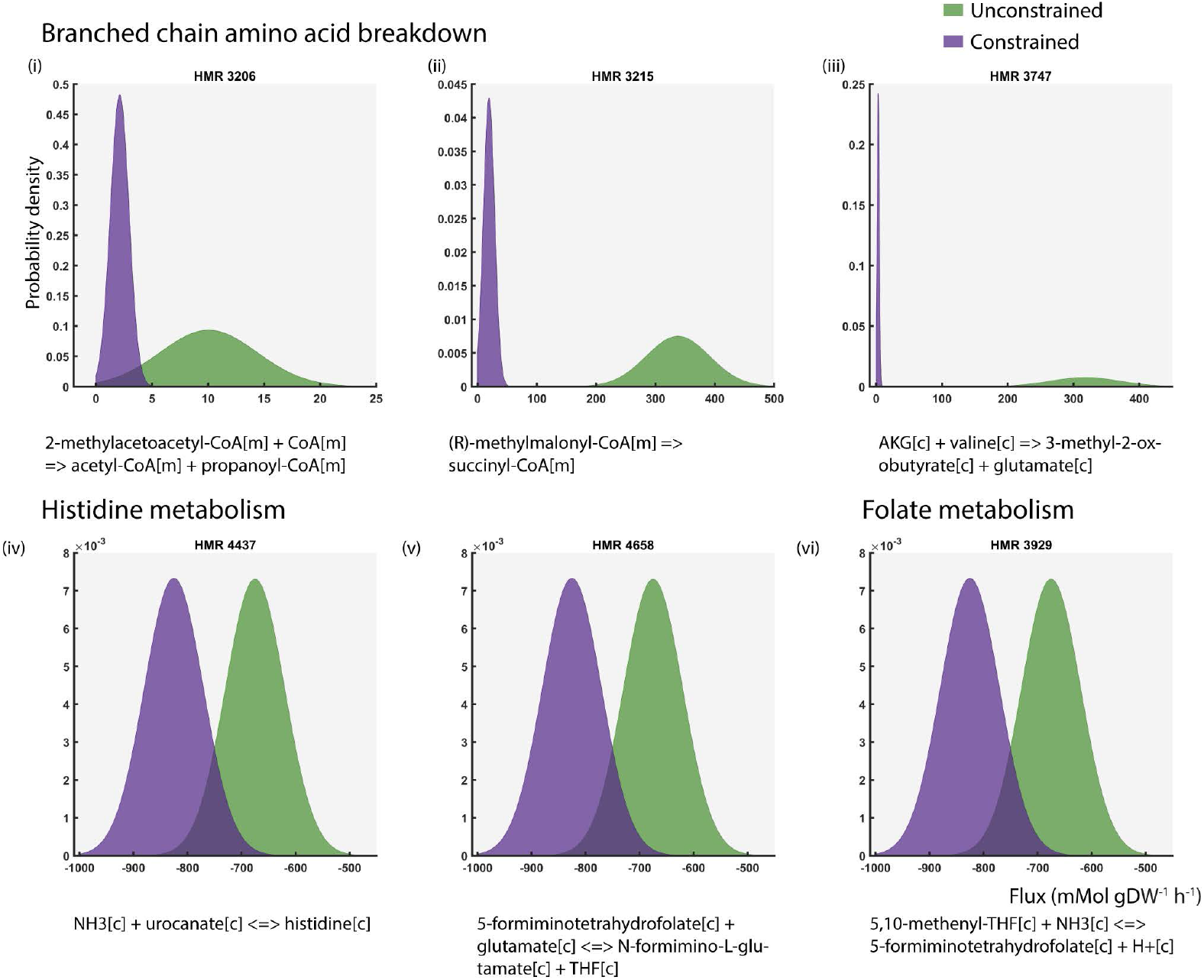
Flux distributions of 6 out of 21 reactions having significant differences in flux statistics (p < 0.05) between the unconstrained and constrained simulations. The reaction IDs and chemical equations have been shown above and below the plots respectively.

On the contrary, the flux distribution through ammonia exchange not only changed in shape, but also reversed in reaction directionality. Under unlimited substrate uptake, our simulations indicated that ammonia is predominantly taken up whereas ammonia is predominantly released on limiting BCAA uptake (Table S2). Interestingly, in the constrained simulation, the flux distribution through the reactions HMR_4437, HMR_4658 and HMR_3929 (Figure 3 iv-vi) retained the same shape and range as the unconstrained curve, but is shifted to the left on the negative axis. This indicated an increase in fluxes and was observed in the reactions releasing ammonia through folate and histidine metabolism, which could potentially contribute to the excess ammonia release.

Overall, the results of Sampling seem to suggest that the restriction of merely three metabolite uptakes can affect the overall behaviour of the network, even in pathways whose connection to these metabolites is not obvious.

However, this test inspected only the differences in individual reaction fluxes caused by the introduced metabolic perturbation without considering any interactions between reactions. Therefore, we followed a PCA-based approach to identify sets of interacting reactions contributing to the underlying metabolic states.

### 2.4 Principal Components

During Sampling, a flux covariance matrix was also computed along with descriptive statistics of individual reaction flux distributions. The rows and columns of the matrix were equal to the number of reactions (4,067×4,067). Essentially, every column was a vector of covariance between the flux distributions of one reaction and all other reactions. Performing PCA on this matrix resulted in PCs explaining variation within each flux space. Each PC was a flux vector containing different values of loadings for all reactions. The graph in Figure 4 reports the percentage of cumulative variance explained by the PCs in both flux spaces. 99.9% of the variation in the metabolic flux spaces were explained by 519 and 514 PCs in the unconstrained and the constrained conditions respectively. The inset in Figure 4 zooms into the first 10 PCs and shows that the highest absolute variance is: 3.03% in the unconstrained and 1.3% in the constrained flux space. Individually, each of the remaining components explained less than 1% of the variation. Nonetheless, these results clearly demonstrate a considerable dimensionality reduction from 1,608 to approximately 500 dimensions in both flux spaces. This reduction also implies that the metabolic state of the adipocyte network can be largely set by regulating these 519 and 514 PCs respectively in the unconstrained and constrained flux spaces.

**FIGURE 4.**
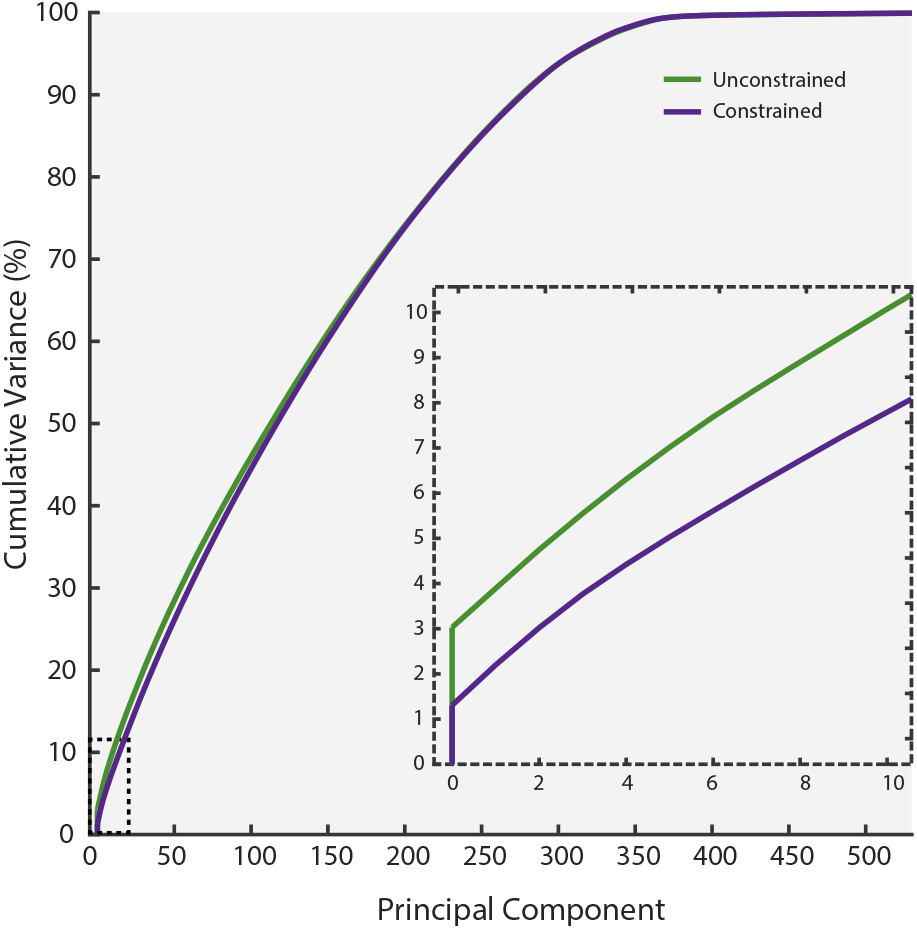
Cumulative variance (%) explained by the principal components of flux spaces from both unconstrained and constrained simulations and the inset zooms into the first 10 principal components.

### 2.5 Basis Rotation

Next, applying a basis rotation on PCs allowed gaining a biochemically meaningful interpretation of the flux vectors by condensing the loadings of 4,067 reactions within each PC into a few high-loading reactions. The reactions whose loadings were at least half of the largest absolute loading value in a given flux vector were considered to be part of the module. In other words, the variance explained by a component was driven by the high-loading reactions. Every module contained distinct sets of reactions which had either positive or negative loading values. We would like to reiterate that the collection of all the modules from an individual flux space formed the global modules of that metabolic condition. Biochemically, it means that the reaction sets forming the global modules operate independently to maintain metabolic homeostasis in the underlying condition.

### 2.6 Global Modules In the adipocyte network: Unconstrained & Constrained

The global modules obtained from all the 519 modules in the unconstrained adipocyte network contained a total of 739 high-loading reactions. Although the reactions within a module were unique, some reactions were present in more than one module. In terms of size, the number of high-loading reactions in each module varied between 1 and 49. Constructing a reaction map from the reactions in the modules resulted in a reaction-reaction network (Figure S3 (A)). Each node represents a reaction and the connections indicate shared metabolites (either a reactant or a product). This network contained 4,212 edges, one highly connected subset (488 reactions) and 142 unconnected reactions. The network structure of the reactions from the global modules of the constrained adipocyte (Figure S3 (B)) was highly comparable to that from the unconstrained model (Figure S3 (A)). The network in Figure S3 (B) had fewer reactions (718) and edges (4,200). Just as seen in Figure S3 (A), this network also contained one highly connected subset (485 reactions) and 135 unconnected nodes. Both networks also contained small subsets of 2-4 reactions.

The connection between reactions and the modules was explored further by associating each reaction with its involved module(s). As highlighted in Figure S3, the color of the nodes were mapped to the number of involved modules. This interlinking revealed that 85-90% of the reactions were present in just one module. Only 12 reactions were present in 3 to (maximum of) 5 modules in both the networks. The dominance of single module reactions implied that very few reactions had a significant loading in multiple modules and most reactions contributed to only one PC of the flux space. In a metabolic context, they were responsible for regulating only one aspect of the network behaviour. Examining the biochemical processes (or subsystems) revealed that nearly half of the reactions in each network were involved in extracellular and mitochondrial transport. These nodes were also responsible for the high network connectivity. A list of all the 67 unique subsystems with the number of reactions in each is given in (Table S3 & S4).

### 2.7 Identification of BCAA-specific modules

ICA originated in the field of signal processing for separating individual sources from a mixture of non-Gaussian signals. In our analysis, ICA enabled identification of the rotated PCs that were distinct between the conditions, thus, contributing to the metabolic differences between the simulations. ICA was performed on the combined set of 1,035 rotated PCs from both conditions (519 and 514). An optimisation was carried out to address the algorithmic stochasticity and to select the number of independent components (ICs or features to be estimated, N).

ICA identified 302 distinct features among which, 159 PCs were from unconstrained and 143 PCs from constrained simulations. Module extraction followed by construction of a combined reaction map resulted in a network of 179 reactions (Figure 5). Colouring the nodes of the network by condition showed that 48 reactions were unique to the unconstrained distinct modules (green nodes) whereas 18 reactions were present only in the distinct constrained modules (violet nodes). 113 reactions were part of the distinct modules from both conditions (orange nodes). Grouping the nodes by subsystem (as described in the model), revealed the involvement of 31 subsystems affected by the introduced perturbation (Table S5). Addition of first neighbouring reactions from the original adipocyte model (that were not part of distinct modules), shown in grey nodes, improved the connectivity between the subsystems and enabled obtaining a more complete picture of the intracellular metabolism. As expected, both cytosolic and mitochondrial BCAA breakdowns were affected by zero BCAA uptake and were observed only in the unconstrained modules. The biochemical relationships between BCAA metabolism and the subsystems predicted in this study were checked for consistency with existing knowledge and are detailed below.

**FIGURE 5.**
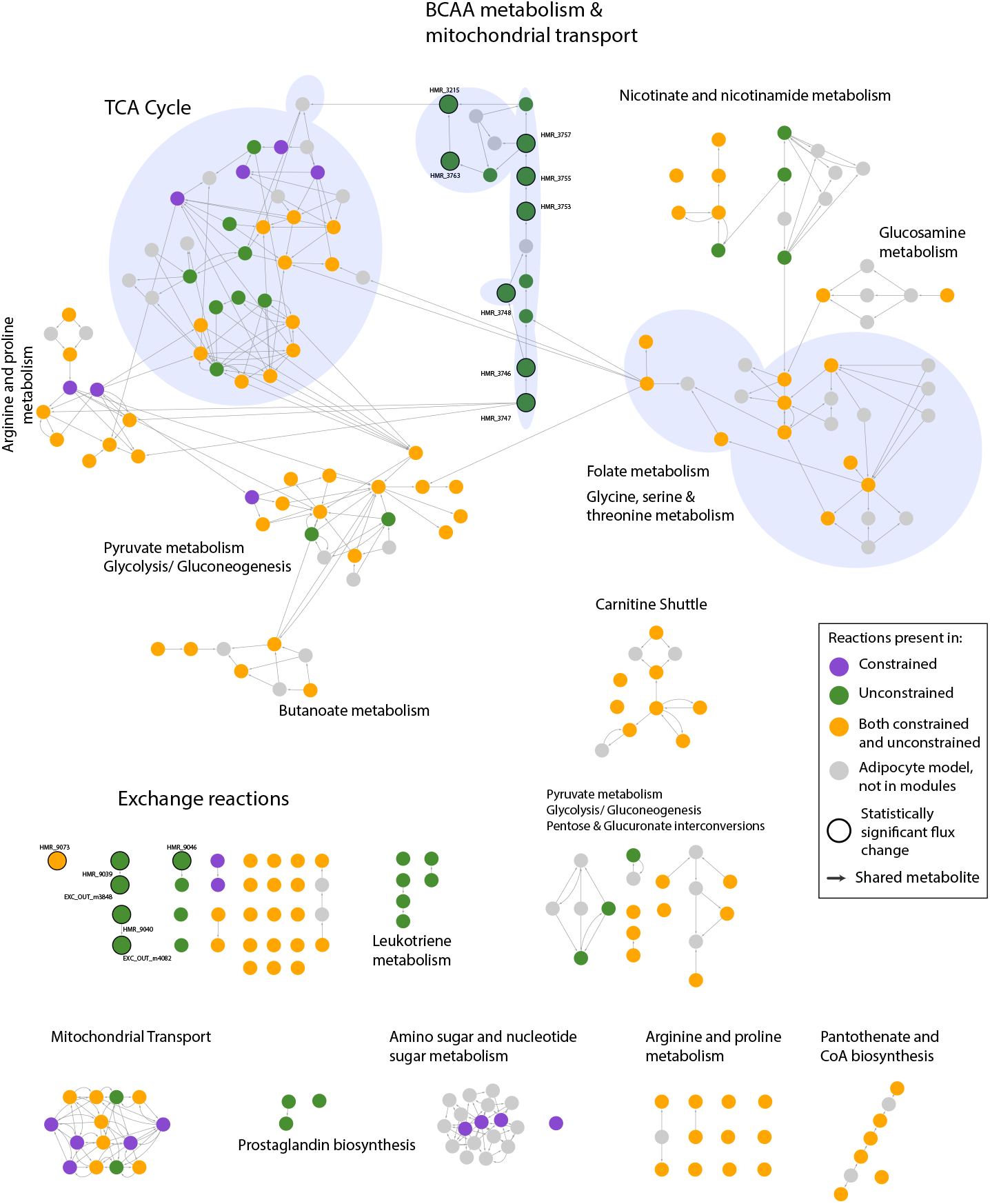
Combined reaction map/network of modules extracted from the 302 PCs that were biochemically distinct between the unconstrained and constrained simulations. Nodes represent reaction and colours indicate the condition: unconstrained (green), constrained (purple) and both (orange). Reactions are grouped by subsystems, some of which are highlighted in blue underlay. Grey nodes are the first neighbours of the reactions in distinct modules extracted from the adipocyte model. The reactions containing significantly changed fluxes are shown as bigger nodes with a black outline. The NDEx link https://bit.ly/2JoxpFf can be used to study the networks interactively.

#### 2.7.1 Comparison of ComMet predictions with existing knowledge

##### Bckdh knockdown

Knockdown of BCKDH (branched-chain alpha-ketoacid dehydrogenase, an enzyme in BCAA degradation) has been shown to reduce BCAA catabolic activity in several studies involving various biological systems (Green et al., 2015). The blocked BCAA uptake simulated here resulted in a similar observation. Unsurprisingly, the biochemically distinct modules showed an active BCAA metabolism only in the presence of BCAA uptake (Figures 5 & 6A).

##### Fatty acid metabolism

Through C-13 labelling studies, (Green et al., 2015) showed that the end products of mitochondrial BCAA catabolism in adipocytes include succinyl-CoA and propionyl-CoA which are then metabolised via TCA cycle into lactate and acetyl-CoA. These metabolites are then channeled towards Fatty acid (FA) metabolism. The distinct modules resulting from our simulations are in line with these experimental observations. Figure 5 reveals the presence of TCA cycle and subsystems related to FA metabolism such as Carnitine shuttle, Leukotriene metabolism, Prostaglandin biosynthesis, Pantothenate metabolism, Beta oxidation and Butanoate metabolism. Notably, Leukotriene metabolism and Prostaglandin biosynthesis were active only in the case of unlimited BCAAs, indicating that these reactions are more significant when BCAAs are available to the cell. On the other hand, the mean reaction fluxes showed an increase in Carnitine shuttle, Beta oxidation and Butanoate metabolism upon limiting BCAA uptake. This increase could be a compensatory response to the decreased availability of acetyl-CoA from BCAA breakdown. Overall, the distinct modules suggest that FA metabolism is affected by BCAA availability in the adipocytes (Figures 5 & 6D).

##### Carbon metabolism

With respect to carbon metabolism, the reactions from TCA cycle were present in both conditions. However, there were notable differences in the mean fluxes of reactions converting alpha-ketoglutarate to succinyl-CoA. In the absence of BCAAs, there was an increased flux in reactions breaking down alpha-ketoglutarate (HMR_6411, HMR_6414, HMR_5297) and a marginal increase in the reactions forming alpha-ketoglutarate (HMR_3957, HMR_3958). Although our analysis did not identify these increases as statistically significant (p-values 0.68, 0.83, 0.78, 0.5, 0.49 respectively), it predicts an increased flux towards succinyl-CoA with no BCAA uptake (Figure 6B). As mentioned above, the end products of BCAA metabolism have been shown to enter TCA cycle as succinyl-CoA (Green et al., 2015). The observed differences in fluxes suggest that, in the absence of BCAAs, metabolic products of alternate sources (such as other amino acids) enter TCA cycle upstream of succinyl-CoA as alpha-ketoglutarate. Overall, the distinct modules indicate that the absence of BCAA uptake affects both TCA cycle and Fatty acid metabolism. Our results emphasise the importance of TCA cycle and the subsequent use of end-products in Fatty acid metabolism in adipocytes.

**FIGURE 6.**
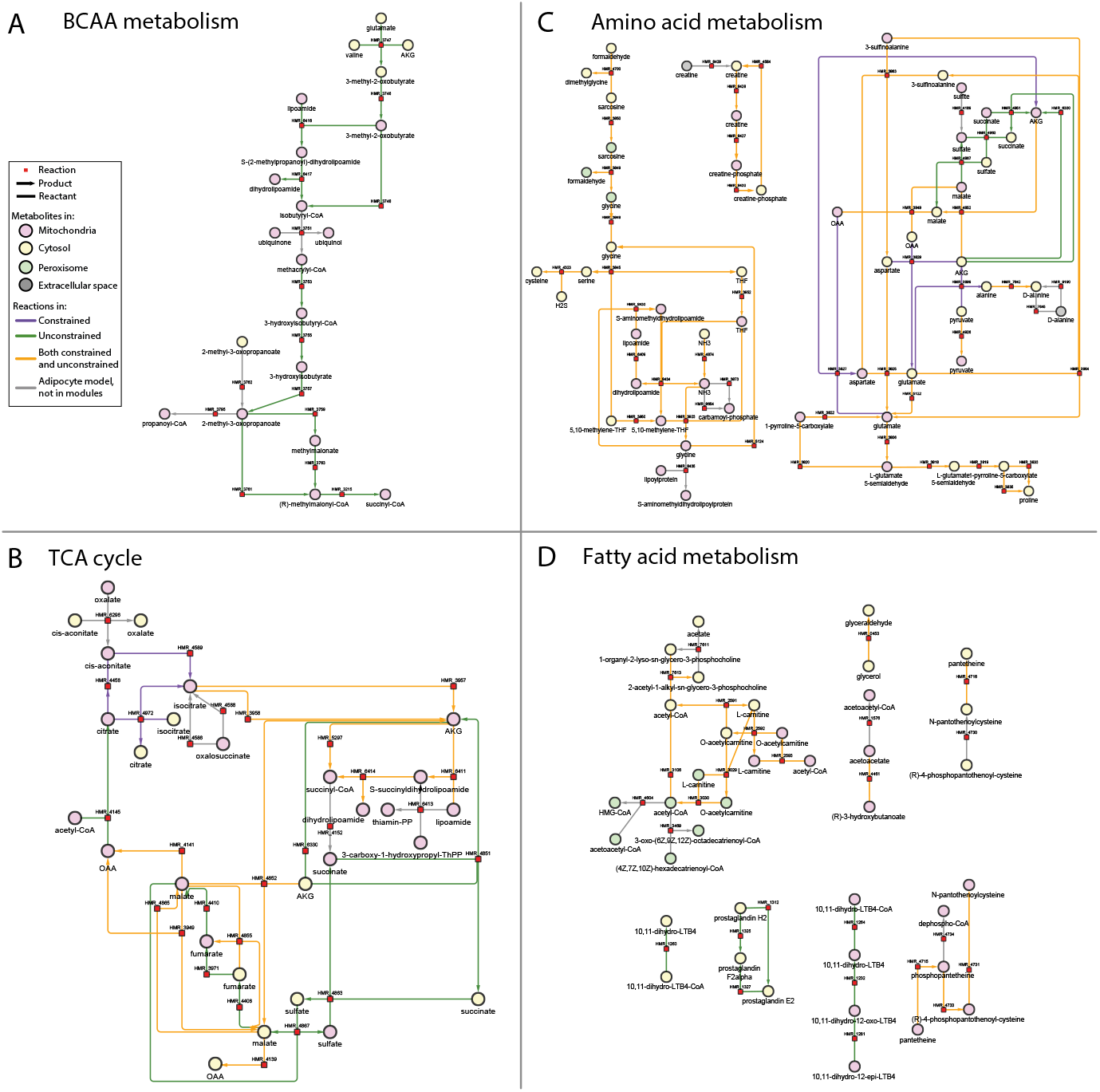
Metabolic map of selected reactions from Figure 5 that were part of distinct modules. Four subsystems are shown:(A) BCAA metabolism, (B) TCA cycle, (C) Amino acid metabolism and (D) Fatty acid metabolism. The nodes in this network represent metabolites (circles) and reactions (squares) while the edges correspond to reactants and products. Metabolite node colours indicate the intracellular location, mitochondria (pink), cytosol (cream) and endoplasmic reticulum (grey). The colour scheme of the edges follows that of the reactions in Figure 5.

#### 2.7.2 Novel mechanisms predicted by ComMet

##### Amino acid metabolism

Amino acids are metabolised by adipocytes and utilised for energy production through the TCA cycle with subsequent usage in fatty acid metabolism (Figures 5 & 6C). The metabolism of Alanine, Aspartate and Glutamate increased and the release of glutamate into the *in silico* media decreased when BCAAs were unavailable. This suggests that adipocytes could be compensating for the lack of BCAAs by increasing the catabolism of alanine, aspartate and glutamate towards TCA cycle via alpha-ketogluterate. In addition, uptake and release of proline from the *in silico* media was exclusive to limited BCAA uptake. On the other hand, decreased glycine secretion was reported in differentiated mice adipocytes cultured on low BCAA levels (Green et al., 2015). In another study, Alves et al. (2019) found that circulating glycine levels were increased in obese rodents and these levels decreased when restricting dietary BCAA intake. In our analysis, increased peroxisomal glycine formation (HMR_3848, HMR_3849) along with increased mitochondrial glycine flux (HMR_3923, HMR_5124) and cytosolic glycine breakdown (HMR_3845, HMR_4700) was observed when BCAAs were unavailable (Figures 5 & 6C and Table S5). Thus, the additional reactions metabolising glycine provide a possible explanation for glycine reduction in absence of BCAAs.

##### Metabolite exchanges with *in silico* media

Several notable differences were observed between the simulations in the exchange profile of the adipocytes. The distinct modules indicate that release of BCAAs were exclusive to the presence of BCAA uptake and ammonia exchange was present in both simulations. Intriguingly, under unlimited BCAA uptake, ammonia was taken up, whereas ammonia was released on limiting BCAA uptake (as discussed under Sampling). Our simulations also indicate that the adipocytes take up nicotinate and release methylnicotinamide in the presence of BCAAs. In particular, the availabiltiy of BCAAs seemed to have affected the profile of ketone bodies (3-hydroxybutanoate and acetoacetate) and non-esterified fatty acids (NEFAs). In the absence of BCAAs, both uptake and release of NEFAs increased, while, the uptake of 3-hydroxybutanoate and release of acetoacetate also increased. Additionally, there were interesting observations in the profile of carbon sources and amino acids. The uptake of glucose and fructose were present in both simulations with marginal differences in their mean fluxes. On the other hand, the uptake of glycerol, glucosamine and xylitol increased while L-arabinose uptake and L-arabitol release reduced. Furthermore, pyruvate uptake and L-lactate release reduced in the absence of BCAAs. As for the profile of amino acids, sarcosine uptake increased while the release of cysteine, glutamate, serine and D-alanine reduced. Taken together, the metabolites describing the changed exchange profile can be used to validate the predictions from our study.

### 2.8 Distinct modules vs flux statistics

Finally, the results from the BCAA-specific modules were compared with those from Sampling alone. The 255 reactions from the distinct modules (Figure 7A) and the 21 reactions with significantly changed fluxes (Figure 7B) have been highlighted on the same plot that compared reaction-wise flux means (Figure 2A). Conversely, about 14 out of these 21 reactions were also found in the distinct modules (Figure 5, bigger nodes with a black outline) which were involved in BCAA metabolism and ammonia exchange. As seen from Figure 7, the number of reactions identified in PCA-based analysis were substantially higher than the reactions with significantly changed flux statistics. The mean fluxes of the reactions from distinct modules appear visually comparable and showed statistically insignificant differences between the simulations. Nonetheless, they revealed biologically meaningful connections between BCAA breakdown and other subsystems (described above) that were not found from Sampling alone.

**FIGURE 7.**
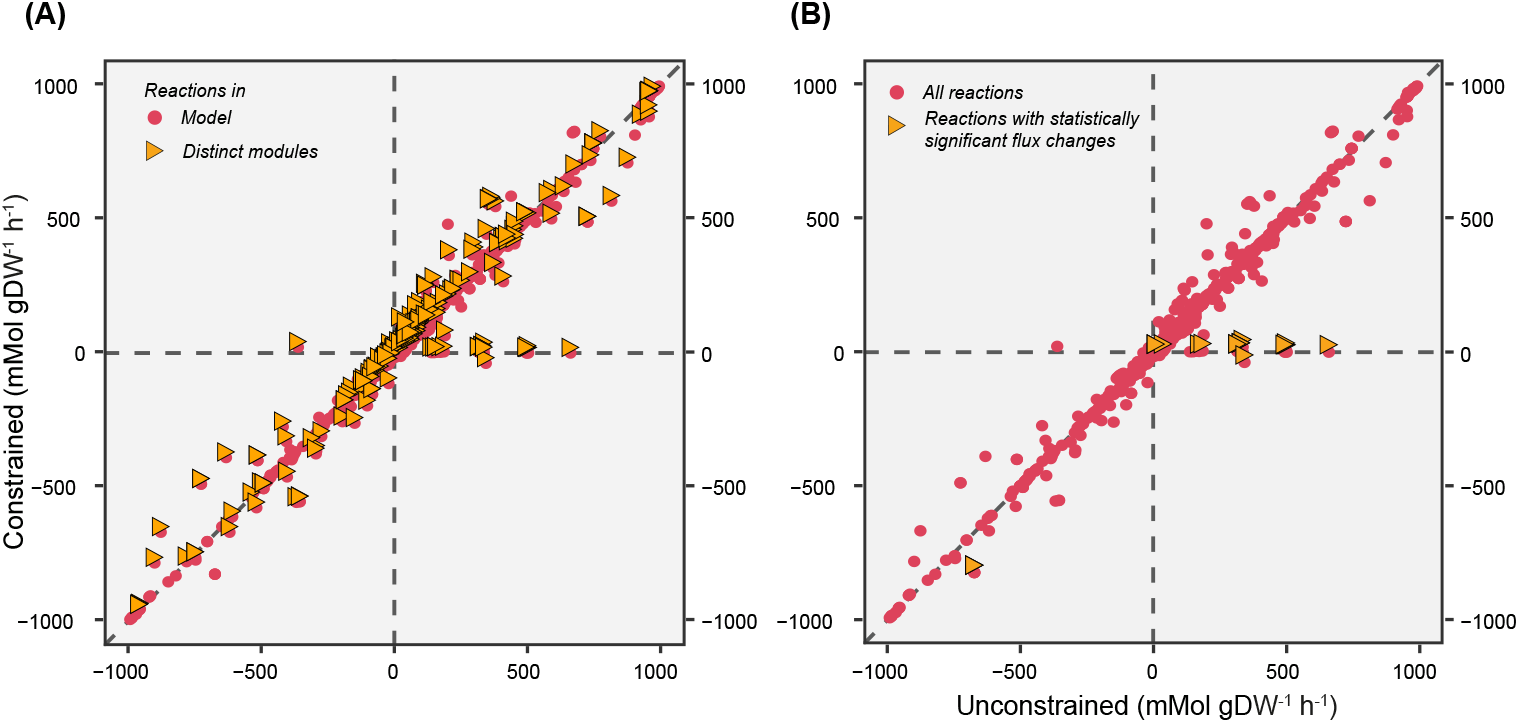
Reaction-wise comparison of means between the unconstrained and constrained simulations (same as Figure 2A) highlighting (A) reactions from the distinct modules and (B) reactions with significantly changed fluxes in yellow triangles.

## 3 DISCUSSION

In this paper we present a novel method, ComMet, for in-depth characterisation of and comparison of distinct metabolic states, which has remained a challenge for large metabolic networks. ComMet facilitates investigating several metabolic states through rigorous optimisation of strategies for comparing metabolic states followed by intuitive network visualisation. Using ComMet, biologically interpretable modules associated with different metabolic states of the adipocyte model (iAdipocytes1809) were extracted.

ComMet was first applied to explore adipocyte flux space and determine condition-specific modules. Principal components of the individual flux spaces uncovered that the metabolic state of an adipocyte network can be set by regulating 500 dimensions. The corresponding modules spanned across a wide array of subsystems thereby controlling various aspects of adipocyte functionality. Using BCAAs as an example, ComMet was then used for identifying differences in adipocyte flux spaces arising from blocked uptake. Histograms of the flux statistics obtained from Sampling provided a broad sense of the flux ranges in each condition. The similarity in shape and spread of the corresponding histograms (Figure S1 A vs B and Figure S1 C vs D) suggested a close correspondence in the overall flux profile between conditions. However, subsequent reaction-wise comparison followed by evaluation of statistical significance indicated otherwise. Identifying reactions with shifts in flux means and/or standard deviations provided a general outline of affected metabolic pathways. Since metabolic reactions seldom act independently, the downstream PCA-based analysis and meticulous comparison were vital in determining network-wide consequences of the introduced perturbation. The observations from only comparing flux statistics failed to highlight metabolic pathways and connections that were indicated in distinct modules and these pathways were even experimentally shown to be affected in other studies. Most notably, the connections between BCAA metabolism and TCA cycle and the differences in metabolite exchanges, resulting from blocked BCAA uptake, were revealed in the distinct modules but not in Sampling alone. Moreover, the changes in glycine breakdown profile predicted by the distinct modules, provide a possible explanation for the reduction in extracellular glycine levels in adipocytes upon limiting BCAA uptake.

The highlight of this study lies not only in comparing metabolic states but also in successful integration of approaches to provide meaningful biological insights. The functionalities included in ComMet offer several advantages. To begin with, the analysis presented here demonstrated an application of ICA in the context of flux spaces. ICA is a useful technique for source separation and has found successful uses in the field of image analysis. Here, ICA enabled distinguishing between flux vectors that are biochemically different in the compared metabolic states. Despite its stochasticity, incorporating ICA optimisation strategies strongly supports reproducibility and ensures data-dependent parameter selection. Automated visualisation of modules as metabolic map and reaction map enabled inferring network-wide consequences of the introduced perturbation thus, contributing to understanding the underlying biological processes. Finally, the computational feasibility offered by ComMet allows application even to human models.

Extracting the biochemical differences from two simulated states of an adipocyte model opens avenues for extensive exploration of metabolic conditions. ComMet can not only be applied on diverse biological systems, ranging from microbe to human models, but it also allows analysing any metabolic states of interest, for example, comparing (a) metabolic shifts between healthy and cancer cells, or (b) metabolic capabilities of different members of gut microbiome, or (c) identifying metabolic differences between cell/tissue types, to name a few. Broadly, ComMet can be employed in two independent scenarios: (1) model-based hypothesis generation or (2) data-driven analysis. As a proof of principle, the first approach was demonstrated here on a human adipocyte GEM. The objective was to showcase comprehensive analysis of large metabolic flux spaces irrespective of the challenges presented by GEMs for multicellular organisms with respect to objective selection and model constraining. The model-based approach can also be extended on a much greater scale by blocking multiple (or combinations of) uptake metabolites. Nevertheless, using experimental data in place of simulated states follows a data-driven approach. When available, omics data can be used to constrain flux spaces prior to the Simulation step in the presented workflow to study physiologically accurate scenarios.

By showcasing the adipocyte modules that are affected by differences in BCAA uptake, we would like to emphasise the utility of ComMet as a tool for generating hypotheses which could be tested in a laboratory setting. Taken together, we demonstrate that ComMet is a powerful tool for holistic understanding of cellular physiology in several metabolic states.

## 4 METHODS

The entire analysis presented in the current study was carried out in MATLAB R2017b (MATLAB, 2017) and all the networks were visualised using Cytoscape v3.7.2 (Shannon, 2003). The genome-scale metabolic reconstruction of human adipocyte, iAdipocytes1809 (Mardinoglu et al., 2013), was used. The model was imported using the RAVEN toolbox (Wang et al., 2018) and it contained 1,809 genes, 6,110 reactions and 4,361 metabolites. Reactions releasing leucine, isoleucine and valine from cytosol to extracellular space were added to the model.

### 4.1 Preprocessing

Prior to Sampling, first, a steady-state flux space was defined by imposing a default set of constraints on all the reactions in the model. The bounds of all the reversible reactions were set to [−1,000 1,000] mmol/gDW/h and the irreversible reactions to [0 1,000] mmol/gDW/h. Same rules were applied to the 151 exchange reactions in the model, which were originally set to no uptake or efflux, depending on directionality. The model was then preprocessed by removing all the blocked reactions (2,043, in this case), which were the reactions incapable of carrying any flux under the imposed conditions. Using Flux Variability Analysis, the minimum and maximum steady-state flux ranges of the remaining 4,067 reactions were then identified and subsequently used for Sampling.

### 4.2 Additional constraints for simulating metabolic states

Starting with the preprocessed model, two metabolic states were simulated: (a) Unconstrained substrate uptake (same as the preprocessed model) and (b) Constrained uptake of BCAAs. Setting both the upper and lower flux bounds of BCAA exchange reactions (HMR_9039: isoleucine uptake, HMR_9040: leucine uptake, HMR_9046: valine uptake) to zero resulted in the constrained model.

### 4.3 Flux Space Sampling

A MATLAB implementation of the Expectation Propagation (EP) algorithm (Braunstein et al., 2017) that was available with the original publication was downloaded and installed. EP Sampling was run with the following parameters individually for each condition: (a) 1,000 as maximum number of iterations (b) 1e^−5^ as the precision accuracy and (c) 1e^8^ as beta. Each EP run resulted in statistics of marginal distributions (means and variances) for each reaction and a matrix describing covariances between all reaction flux distributions. The number of rows and columns in the resulting covariance matrix was equal to the number of reactions (4,067) and this matrix was used for the downstream PCA-based analysis.

### 4.4 Differential flux statistics

The objective here was to obtain a Z-score (Z_i_^flux^) for each reaction, i, using means (E) and variances (Var) of flux distributions (v) in both conditions (denoted by subscripts 1 and 2 in equation (1)). The Z-score was used to quantify the significance of the change in each flux distribution between considered conditions (Bordel et al., 2010). It was calculated as the difference between the means in each of the conditions divided by the square root of the sum of variances in the respective conditions.

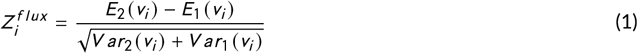

These Z-scores were then transformed into probabilities of change using a cumulative Gaussian distribution. These p-values represented the significance of change in fluxes between the conditions.

### 4.5 Principal Component Analysis and Basis Rotation

Eigenvectors and eigenvalues of the covariance matrix were calculated. The variance explained by each vector was computed by normalising the eigenvalues. The number of vectors explaining 99.9% of the variance was then identified and all the non-zero loadings were rotated using varimax rotation. This step resulted in one matrix of reaction loadings for each condition, where the rows represented reactions and the columns represented PCs.

### 4.6 Module Extraction

The reactions whose loadings were within half of the maximal loading within each principal component were considered as part of the module (same as the criterion used in (Barrett et al., 2009) study). The number of reactions above the defined threshold varied in each component. The set of reactions present in at least one module in a condition, or the global modules, were identified and used for constructing the reaction network.

### 4.7 Independent Component Analysis

For identifying BCAA-specific modules, the matrices containing reaction loadings from both conditions were concatenated. Next, a loading cut-off was calculated by summing the mean absolute deviation and median of the maximum absolute loadings. This cut-off was used to identify and remove reactions with low loadings in all the PCs.

To identify the optimum number of ICs, first, a bootstrapping analysis was performed using the FastICA algorithm implemented In icasso toolbox (Himberg and Hyvarinen, 2003). N ICs were computed in 100 iterations with random initial conditions, where N = [2 to 90, in steps of 1]. pow3 nonlinearity and symmetrical approach were used for the decomposition. Next, the consistency of the estimated ICs across iterations was assessed by plotting a stability profile for each N using the BIODICA toolbox (Kairov et al., 2017a,b). Figure S4 (A) shows that, on average, the stability of the clusters decreases with increasing N (grey lines). Next, 2-means clustering was used to group the stability measures into two lines: one with uniform stability distribution (blue line) and the other with low stability distribution (red line). The point of intersection of these two lines revealed the optimal number of IC as 20 (black vertical line).

Next, ICA was rerun with the identified optimum N (20) for 9000 iterations. To ensure reproducibility of the estimated features, a random number was explicitly set for each iteration. The resulting kurtosis values for all the ICs were examined and the features having kurtosis greater or equal to 1 and lesser or equal to −1 were extracted. The plot of estimation frequency (the number of runs in which each feature was estimated as an IC, Figure S4 (B)) revealed that the top 20 features were estimated in about 70% of the iterations (Figure S4 (B)). These were too low to extract modules describing meaningful biological differences (Figure S5). Then, Kaplan (2020) was used to identify the knee point of this curve, indicated by the cyan vertical line in Figure S4 (B). All the features on the left of the cyan line were selected for further analysis. The selected features corresponded to 302 rotated-PCs that were distinct, and thus, described the metabolic differences between the two simulated conditions. Modules were then extracted from these rotated-PCs and the corresponding reactions were also visualised as a reaction network.

### 4.8 Reaction Networks

To construct a reaction map/network from the global modules, two text files were generated: one graph file and an attributes file. The graph file described the connectivity between reactions and was built based on the connectivities defined in the metabolic network. Two reactions were defined to be connected if they shared either a product or a reactant. The following ubiquitous metabolites were removed from the calculation of connectivity - CoA, ubiquinol, ubiquinone, NH3, O2, H2O, H^+^, ATP, ADP, AMP, dADP, dATP, Pi, PPi, CTP, CDP, CMP, dCTP, dCDP, dCMP, UTP, UDP, UMP, dUTP, dUDP, dUMP, GTP, GDP, GMP, dGTP, dGDP, dGMP, ITP, IDP, IMP, dITP, dIDP, dIMP, TTP, TDP, TMP, dTTP, dTDP, dTMP, NADH, NADPH, NAD^+^, NADP^+^, FADH2, FAD, CO2, Na^+^, HCO3^−^. The attributes file, on the other hand, described features of reactions. For each reaction, the following properties were identified: (1) the number of modules it is involved in, (2) list of all the modules involved, (3) subsystem to which it belongs and (4) its chemical equation. The reaction tables were written into semi-colon separated text files which were imported into Cytoscape for further investigation. In the case of the distinct modules network, the reaction tables from both the conditions were merged prior to import. These combined reactions were visualised as the distinct modules network in Cytoscape (Figure 5). In addition to the reactions shown in Figure 5, the network contained reactions from Extracellular Transport (with 128 nodes) which were hidden from visualisation for the ease of interpretation and have been reported in Table S5. To construct metabolic map, reactions from the distinct modules network belonging to subsystems of interest were extracted as submodels and visualised using EFMviz (Sarathy et al., 2020). All the network operations were automated through an R script (R v3.5.1) using the library, RCy3 (Ono et al., 2015). NDEx (Pratt et al., 2015) links for all the reaction networks have been provided for further interactive exploration.

## Supporting information

ComMet Supplementary Figures and Tables

## Data and Code Availability

All the code used in this study are available at https://github.com/macsbio/commet.

## Conflict of interest

The authors declare no conflict of interest.

## Supplementary Information

This section includes five supplemental figures (S1-S5) and six supplemental tables (Table S1-S6)

### Supplementary Figures

**FIGURE S1** Histogram of means (A,B) and standard deviations (C,D) from both unconstrained and constrained simulations.

**FIGURE S2** Flux distributions of 9 out of 21 reactions having significant differences in flux statistics (p < 0.05) between the unconstrained and constrained simulations.

**FIGURE S3** Reaction map/network of the global Modules from the (A) unconstrained and (B) constrained adipocyte network.

**FIGURE S4** ICA optimization for selecting the number of features.

**FIGURE S5** Combined reaction map/network of modules extracted from the top 17 frequently occurring features.

### Supplementary Tables

**TABLE S1** Table with flux statistics of all the reactions from both simulations

**TABLE S2** Reactions with significant changes in flux statistics between unconstrained and constrained simulations

**TABLE S3** Subsystem distribution: reactions from global modules - unconstrained

**TABLE S4** Subsystem distribution: reactions from global modules - constrained

**TABLE S5** Table with reactions in the modules that were distinct between the simulations

**TABLE S6** Table with processing time for each step in ComMet’s pipeline

## Notes

**Funding information** This research has been made possible with the support of the Dutch Province of Limburg, The Netherlands.

### Competing Interest Statement

The authors have declared no competing interest.

### Summary of Updates

Result section revised

