## Supplementary material for "ComMet: A method for comparing metabolic states in genome-scale metabolic models": ComMet Supplementary Figures and Tables: commet_supp.pdf

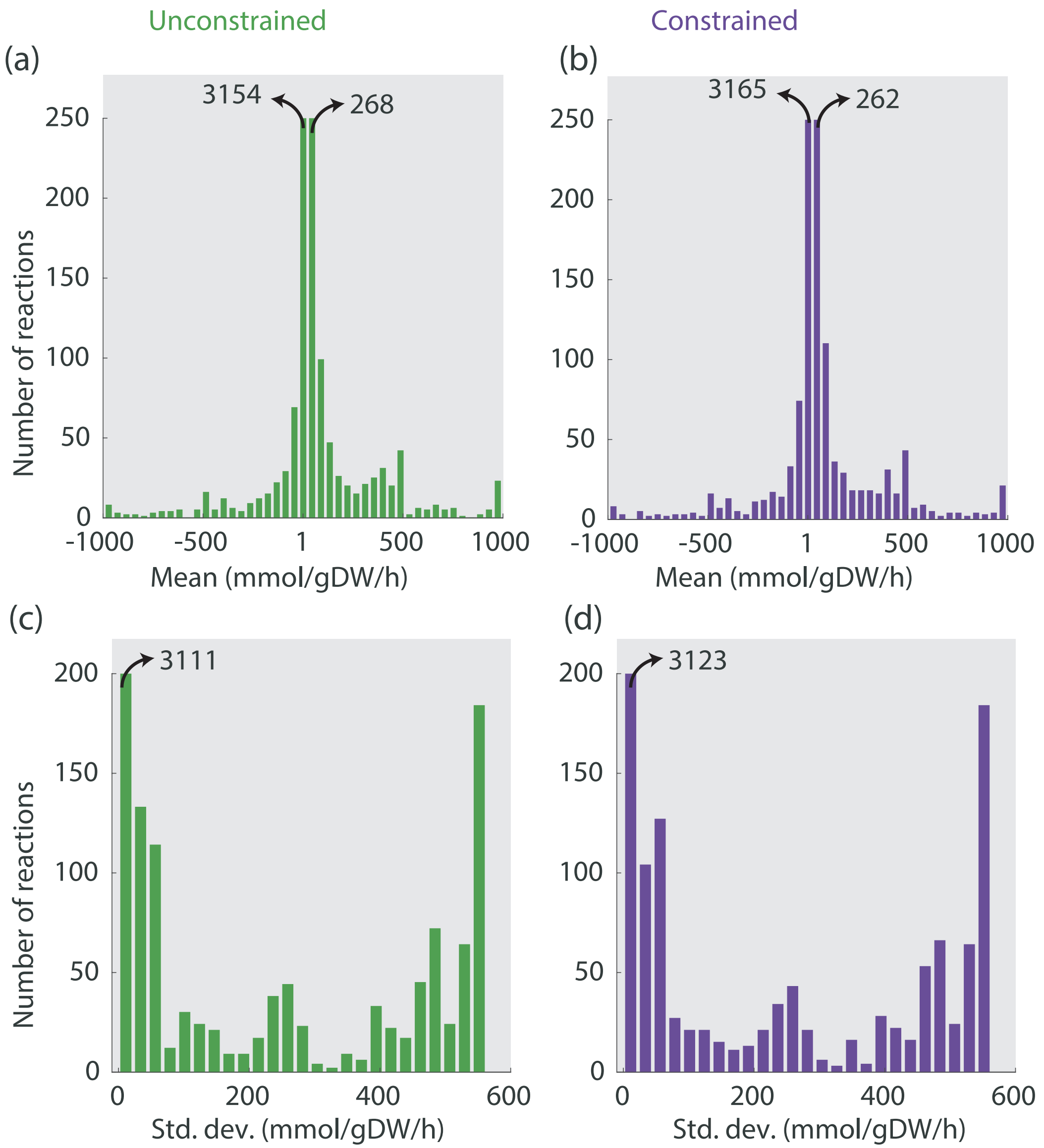


**FIGURE S1** Histogram of means (A,B) and standard deviations (C,D) from both unconstrained and constrained simulations. Due to the very high number of reactions for the lowest values, their number of reactions have been indicated separately with arrows.

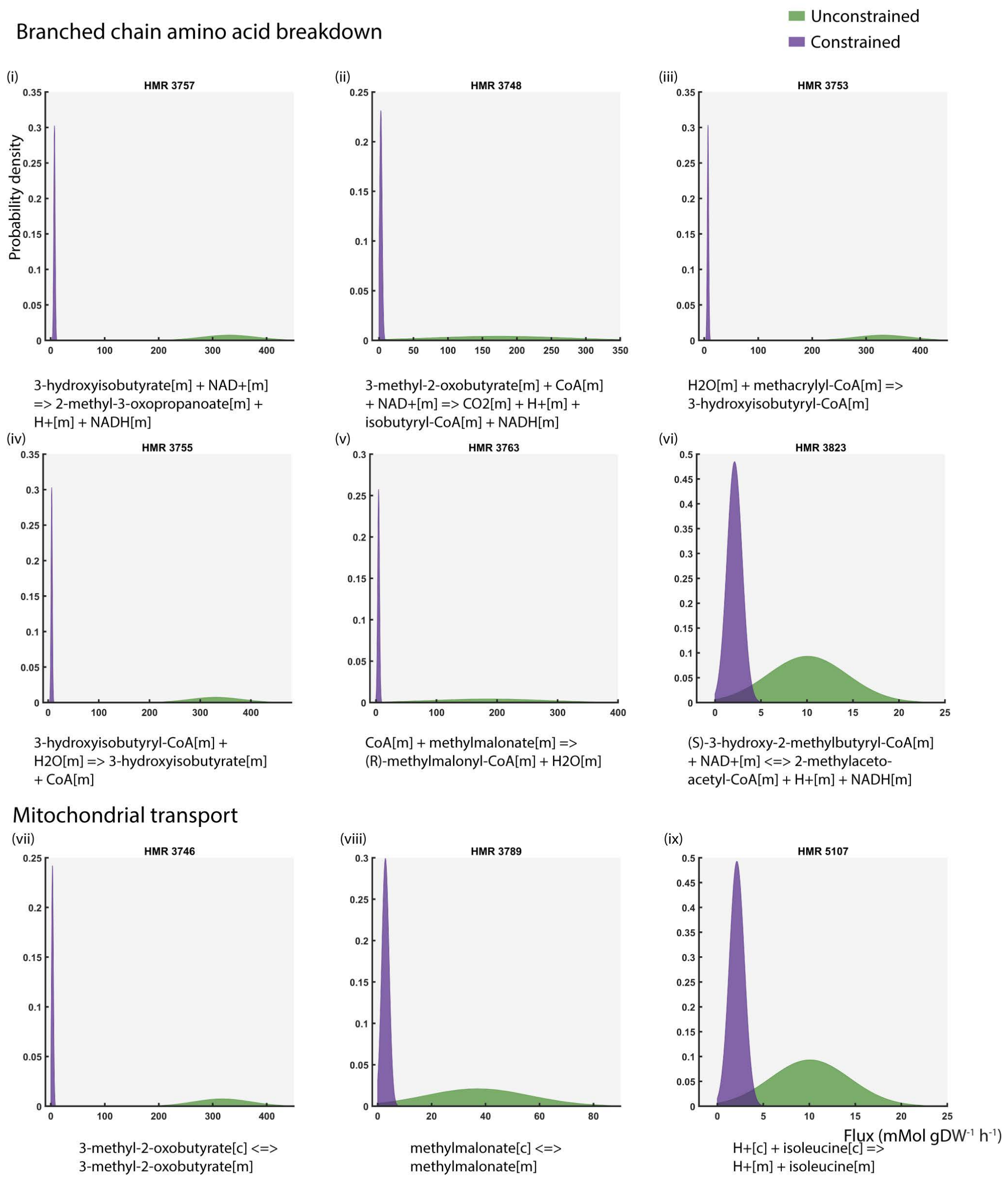


**FIGURE S2** Flux distributions of 9 out of 21 reactions having significant differences in flux statistics ($p < 0.05$) between the unconstrained and constrained simulations. The reaction IDs and chemical equations have been shown above and below the plots respectively.

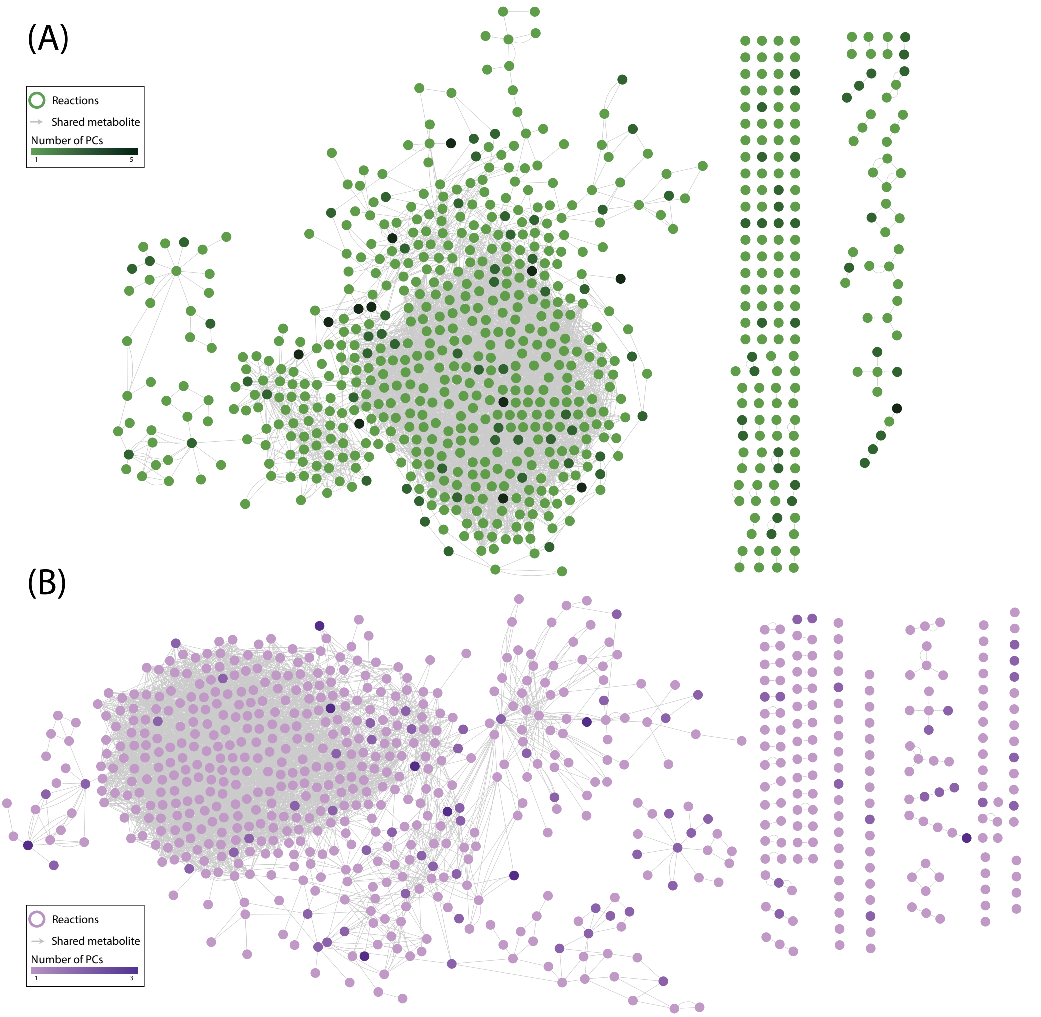


**FIGURE S3** Reaction map/network of the global Modules from the (A) unconstrained and (B) constrained adipocyte network. Nodes represent reactions and the edges indicate shared reactant/product. Node colour is mapped to the number of involved modules. The NDEx links https://bit.ly/38HtnCp and https://bit.ly/35q0sk5 can be used to study the networks interactively.

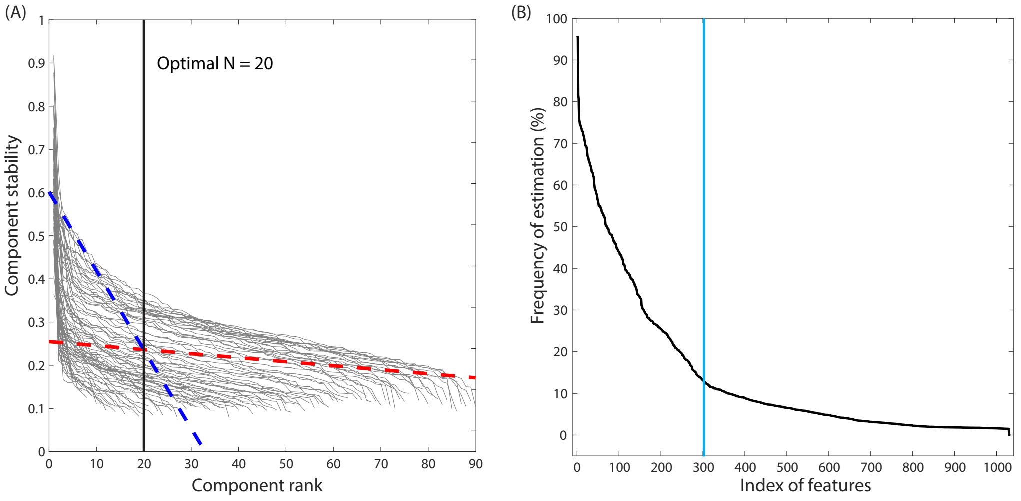


**FIGURE S4** ICA optimisation for selecting the number of features. (A) Grey lines show the stability profiles of the bootstrapped Independent Component Analysis (ICA) runs. Blue and red dashed lines are results of two-line clustering with the optimal N determined as the point of their intersection (black vertical line). (B) Frequency of estimation of a feature as an independent component. Cyan vertical line marks the knee-point cut-off (302) for selecting distinct features.

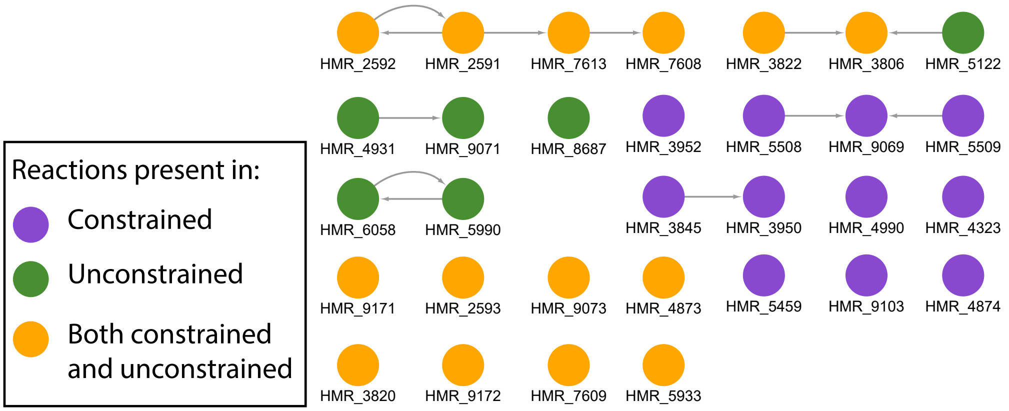


**FIGURE S5** Combined reaction map/network of modules extracted from the top 20 frequently occurring features.

| TableS1: Table with flux statistics of all the reactions from both simulations | | | | | | | | |
|---|---|---|---|---|---|---|---|---|
| Reaction | Subsystem | Mean (unconstrained) | Variance (unconstrained) | Mean (constrained) | Variance (constrained) | AbsoluteDifference in means (unconstrained - constrained) | Z-score | P-value |
| HMR_3905 | Glycolysis / Gluconeogenesis | 12.5517 | 109.8467 | 13.709 | 130.7367 | 1.1573 | -0.0746 | 0.5297 |
| HMR_3907 | Glycolysis / Gluconeogenesis | 12.7381 | 112.087 | 13.9292 | 133.6063 | 1.1911 | -0.076 | 0.5303 |
| HMR_4097 | Glycolysis / Gluconeogenesis | 2.638 | 7.2158 | 2.6991 | 7.5017 | 0.0611 | -0.0159 | 0.5064 |
| HMR_4099 | Glycolysis / Gluconeogenesis | 2.6652 | 7.3593 | 2.7265 | 7.6415 | 0.0613 | -0.0158 | 0.5063 |
| HMR_4108 | Glycolysis / Gluconeogenesis | 2.6446 | 3.267 | 2.7047 | 3.4175 | 0.06 | -0.0232 | 0.5093 |
| HMR_4133 | Glycolysis / Gluconeogenesis | 2.6433 | 3.267 | 2.7033 | 3.4175 | 0.0601 | -0.0232 | 0.5093 |
| HMR_4137 | Glycolysis / Gluconeogenesis | 380.6548 | 23774.0592 | 546.6356 | 26030.4809 | 165.9808 | -0.7437 | 0.7715 |
| HMR_4281 | Glycolysis / Gluconeogenesis | 74.9785 | 112.5135 | 77.9618 | 117.6653 | 2.9834 | -0.1966 | 0.5779 |
| HMR_4283 | Glycolysis / Gluconeogenesis | 16.3557 | 202.626 | 15.8668 | 197.489 | 0.4889 | 0.0244 | 0.4902 |
| HMR_4301 | Glycolysis / Gluconeogenesis | 327.0007 | 53826.1228 | 326.2865 | 53709.3669 | 0.7142 | 0.0022 | 0.4991 |
| HMR_4355 | Glycolysis / Gluconeogenesis | 209.4661 | 61493.4089 | 217.4848 | 61647.9212 | 8.0187 | -0.0229 | 0.5091 |
| HMR_4358 | Glycolysis / Gluconeogenesis | 289.3114 | 53011.4705 | 286.3514 | 52343.2201 | 2.9599 | 0.0091 | 0.4964 |
| HMR_4360 | Glycolysis / Gluconeogenesis | 4.1401 | 18.3704 | 4.1782 | 18.4723 | 0.0381 | -0.0063 | 0.5025 |
| HMR_4362 | Glycolysis / Gluconeogenesis | 2.8684 | 8.3558 | 2.7377 | 7.6121 | 0.1308 | 0.0327 | 0.4869 |
| HMR_4363 | Glycolysis / Gluconeogenesis | 996.144 | 12.3724 | 996.3671 | 10.9577 | 0.223 | -0.0462 | 0.5184 |
| HMR_4365 | Glycolysis / Gluconeogenesis | -988.7502 | 34.2434 | -989.4238 | 30.1957 | 0.6735 | 0.0839 | 0.4666 |
| HMR_4368 | Glycolysis / Gluconeogenesis | 981.0597 | 62.5437 | 982.2938 | 55.8896 | 1.234 | -0.1134 | 0.5451 |
| HMR_4370 | Glycolysis / Gluconeogenesis | 2.8471 | 9.2012 | 2.718 | 8.4671 | 0.1291 | 0.0307 | 0.4878 |
| HMR_4371 | Glycolysis / Gluconeogenesis | 2.8687 | 3.8318 | 2.7407 | 3.4962 | 0.1281 | 0.0473 | 0.4811 |
| HMR_4372 | Glycolysis / Gluconeogenesis | 2.8651 | 3.8317 | 2.7369 | 3.4961 | 0.1282 | 0.0474 | 0.4811 |
| HMR_4373 | Glycolysis / Gluconeogenesis | -986.7775 | 51.1501 | -987.7545 | 45.5148 | 0.9771 | 0.0994 | 0.4604 |
| HMR_4375 | Glycolysis / Gluconeogenesis | -648.2784 | 61021.9362 | -647.8991 | 60958.2022 | 0.3792 | -0.0011 | 0.5004 |
| HMR_4377 | Glycolysis / Gluconeogenesis | 2.8591 | 8.3011 | 2.7287 | 7.5624 | 0.1303 | 0.0327 | 0.4869 |
| HMR_4379 | Glycolysis / Gluconeogenesis | 324.1358 | 53415.3082 | 324.3403 | 53430.2806 | 0.2045 | -0.0006 | 0.5002 |
| HMR_4381 | Glycolysis / Gluconeogenesis | -115.9017 | 215911.4323 | -113.354 | 216123.2405 | 2.5477 | -0.0039 | 0.5015 |
| HMR_4388 | Glycolysis / Gluconeogenesis | 190.4681 | 92659.1446 | 64.6954 | 124469.5954 | 125.7727 | 0.2699 | 0.3936 |
| HMR_4391 | Glycolysis / Gluconeogenesis | 447.612 | 626.9786 | 440.411 | 883.4532 | 7.201 | 0.1853 | 0.4265 |
| HMR_4394 | Glycolysis / Gluconeogenesis | 38.7267 | 1223.1204 | 38.4554 | 1210.8269 | 0.2713 | 0.0055 | 0.4978 |
| HMR_4396 | Glycolysis / Gluconeogenesis | 41.388 | 400.4323 | 37.3071 | 346.4768 | 4.0809 | 0.1493 | 0.4406 |
| HMR_4521 | Glycolysis / Gluconeogenesis | 3.0439 | 4.3327 | 2.8989 | 3.9302 | 0.145 | 0.0505 | 0.4799 |
| HMR_6410 | Glycolysis / Gluconeogenesis | 241.6285 | 20618.9221 | 253.556 | 23090.989 | 11.9275 | -0.0571 | 0.5227 |
| HMR_6412 | Glycolysis / Gluconeogenesis | 241.6286 | 20618.9225 | 253.5549 | 23090.9894 | 11.9263 | -0.057 | 0.5227 |
| HMR_7745 | Glycolysis / Gluconeogenesis | -80.3183 | 1768.0679 | -79.722 | 1749.9121 | 0.5962 | -0.0101 | 0.504 |
| HMR_7746 | Glycolysis / Gluconeogenesis | 39.9506 | 1160.7228 | 39.6525 | 1145.6563 | 0.2981 | 0.0062 | 0.4975 |
| HMR_7747 | Glycolysis / Gluconeogenesis | 40.3677 | 1176.736 | 40.0695 | 1161.6404 | 0.2982 | 0.0062 | 0.4975 |
| HMR_7748 | Glycolysis / Gluconeogenesis | -19.446 | 215542.629 | -18.3327 | 215736.3826 | 1.1133 | -0.0017 | 0.5007 |
| HMR_7749 | Glycolysis / Gluconeogenesis | 99.7644 | 215548.1904 | 98.0552 | 215764.9515 | 1.7092 | 0.0026 | 0.499 |
| HMR_8357 | Glycolysis / Gluconeogenesis | 32.2469 | 353.1375 | 35.0406 | 400.9381 | 2.7936 | -0.1017 | 0.5405 |

| | | | | | | | | |
|---|---|---|---|---|---|---|---|---|
| HMR_4122 | Starch and sucrose metabolism | 11.8356 | 5.4094 | 11.9043 | 5.4722 | 0.0687 | -0.0208 | 0.5083 |
| HMR_5395 | Starch and sucrose metabolism | 0.5343 | 0.0748 | 0.5973 | 0.0934 | 0.0631 | -0.1537 | 0.5611 |
| HMR_5396 | Starch and sucrose metabolism | 0.5416 | 0.0747 | 0.606 | 0.0933 | 0.0645 | -0.1573 | 0.5625 |
| HMR_5397 | Starch and sucrose metabolism | 0.559 | 0.0752 | 0.625 | 0.0937 | 0.0659 | -0.1605 | 0.5637 |
| HMR_5398 | Starch and sucrose metabolism | 0.5737 | 0.0756 | 0.6414 | 0.0942 | 0.0677 | -0.1643 | 0.5653 |
| HMR_5399 | Starch and sucrose metabolism | 0.5768 | 0.0759 | 0.6462 | 0.0945 | 0.0694 | -0.1682 | 0.5668 |
| HMR_5400 | Starch and sucrose metabolism | 0.5775 | 0.0758 | 0.6489 | 0.0946 | 0.0714 | -0.173 | 0.5687 |
| HMR_5401 | Starch and sucrose metabolism | 0.576 | 0.0755 | 0.6498 | 0.0943 | 0.0737 | -0.179 | 0.571 |
| HMR_3944 | Galactose metabolism | 451.7087 | 72154.8911 | 452.2265 | 72257.2041 | 0.5178 | -0.0014 | 0.5005 |
| HMR_4128 | Galactose metabolism | 37.51 | 305.4737 | 34.2591 | 257.9942 | 3.2508 | 0.1369 | 0.4455 |
| HMR_4130 | Galactose metabolism | 20.4138 | 258.9201 | 18.9268 | 221.1151 | 1.487 | 0.0679 | 0.4729 |
| HMR_4131 | Galactose metabolism | -444.2958 | 72347.377 | -442.409 | 72424.9099 | 1.8868 | -0.005 | 0.502 |
| HMR_4132 | Galactose metabolism | 23.0837 | 209.4661 | 21.4191 | 179.7263 | 1.6645 | 0.0844 | 0.4664 |
| HMR_4414 | Galactose metabolism | 23.083 | 209.4651 | 21.4185 | 179.7253 | 1.6646 | 0.0844 | 0.4664 |
| HMR_4415 | Galactose metabolism | 37.4984 | 305.4627 | 34.2475 | 257.9824 | 3.2508 | 0.137 | 0.4455 |
| HMR_8767 | Galactose metabolism | 423.8814 | 72167.8524 | 423.4814 | 72271.3468 | 0.4 | 0.0011 | 0.4996 |
| HMR_0454 | Fructose and Mannose metabolism | 65.3347 | 2008.8572 | 81.7344 | 3060.3358 | 16.3997 | -0.2303 | 0.5911 |
| HMR_4297 | Fructose and mannose metabolism | 2.8486 | 3.7933 | 2.7197 | 3.4577 | 0.129 | 0.0479 | 0.4809 |
| HMR_4319 | Fructose and mannose metabolism | 40.6876 | 1287.4057 | 40.3998 | 1272.2654 | 0.2879 | 0.0057 | 0.4977 |
| HMR_4706 | Fructose and Mannose metabolism | 2.8479 | 3.7933 | 2.7189 | 3.4577 | 0.129 | 0.0479 | 0.4809 |
| HMR_4590 | Pentose and glucuronate interconversions | 408.3837 | 57852.9907 | 409.1603 | 57953.2198 | 0.7767 | -0.0023 | 0.5009 |
| HMR_4591 | Pentose and glucuronate interconversions | 408.3838 | 57852.9909 | 409.1605 | 57953.2201 | 0.7767 | -0.0023 | 0.5009 |
| HMR_4593 | Pentose and glucuronate interconversions | 42.084 | 614.1273 | 47.8576 | 789.897 | 5.7737 | -0.1541 | 0.5612 |
| HMR_4595 | Pentose and glucuronate interconversions | -14.7835 | 800.3764 | -19.5305 | 983.3895 | 4.7471 | 0.1124 | 0.4553 |
| HMR_8341 | Pentose and glucuronate interconversions | 726.8843 | 18062.424 | 489.2168 | 42958.3562 | 237.6675 | 0.9621 | 0.168 |
| HMR_1568 | Pyruvate metabolism | 16.0333 | 196.5293 | 15.5606 | 191.4682 | 0.4727 | 0.024 | 0.4904 |
| HMR_3853 | Pyruvate metabolism | 10.1577 | 83.1787 | 11.8099 | 114.2413 | 1.6523 | -0.1176 | 0.5468 |
| HMR_3854 | Pyruvate metabolism | 10.1541 | 83.0819 | 11.7994 | 113.9638 | 1.6453 | -0.1172 | 0.5467 |
| HMR_3855 | Pyruvate metabolism | 405.8563 | 57868.9341 | 407.7317 | 57975.1657 | 1.8754 | -0.0055 | 0.5022 |
| HMR_3857 | Pyruvate metabolism | 6.1245 | 34.339 | 5.8859 | 32.7043 | 0.2386 | 0.0291 | 0.4884 |
| HMR_3859 | Pyruvate metabolism | 57.9608 | 2139.189 | 33.5178 | 895.0812 | 24.4429 | 0.4437 | 0.3286 |
| HMR_4087 | Pyruvate metabolism | 13.9878 | 257.4397 | 10.4382 | 124.3568 | 3.5496 | 0.1817 | 0.4279 |
| HMR_4089 | Pyruvate metabolism | 67.763 | 2944.0156 | 55.6878 | 2272.5154 | 12.0752 | 0.1672 | 0.4336 |
| HMR_4091 | Pyruvate metabolism | 14.0711 | 261.6656 | 10.4709 | 125.3567 | 3.6002 | 0.183 | 0.4274 |
| HMR_4093 | Pyruvate metabolism | 8.3929 | 74.6354 | 8.2255 | 70.338 | 0.1674 | 0.0139 | 0.4945 |
| HMR_4095 | Pyruvate metabolism | 11.6439 | 152.7193 | 12.1087 | 158.171 | 0.4647 | -0.0264 | 0.5105 |
| HMR_4101 | Pyruvate metabolism | 73.5502 | 3204.5065 | 60.405 | 2534.7521 | 13.1452 | 0.1735 | 0.4311 |
| HMR_4103 | Pyruvate metabolism | 4.4877 | 25.6688 | 4.2157 | 21.8456 | 0.272 | 0.0395 | 0.4843 |
| HMR_4143 | Pyruvate metabolism | 6.1883 | 44.4471 | 7.6682 | 67.686 | 1.4799 | -0.1398 | 0.5556 |
| HMR_4193 | Pyruvate metabolism | 274.6427 | 49050.1399 | 272.722 | 48593.3492 | 1.9207 | 0.0061 | 0.4975 |
| HMR_4280 | Pyruvate metabolism | 409.8723 | 80298.802 | 266.2514 | 96369.1685 | 143.6209 | 0.3417 | 0.3663 |
| HMR_8497 | Pyruvate metabolism | 10.5394 | 49.2269 | 12.1552 | 65.8264 | 1.6158 | -0.1506 | 0.5599 |
| HMR_8500 | Pyruvate metabolism | 10.5401 | 49.2264 | 12.156 | 65.8259 | 1.6158 | -0.1506 | 0.5599 |
| HMR_8501 | Pyruvate metabolism | 10.0282 | 81.4508 | 11.6352 | 111.444 | 1.607 | -0.1157 | 0.5461 |

| | | | | | | | | |
|---|---|---|---|---|---|---|---|---|
| HMR_8502 | Pyruvate metabolism | -19.4295 | 195.1702 | -26.7535 | 291.4194 | 7.3241 | 0.332 | 0.3699 |
| HMR_8503 | Pyruvate metabolism | 6.1252 | 34.3371 | 5.8869 | 32.7044 | 0.2383 | 0.0291 | 0.4884 |
| HMR_8504 | Pyruvate metabolism | 6.1245 | 34.339 | 5.8859 | 32.7043 | 0.2386 | 0.0291 | 0.4884 |
| HMR_8506 | Pyruvate metabolism | 9.2015 | 37.99 | 8.873 | 35.8967 | 0.3285 | 0.0382 | 0.4848 |
| HMR_8507 | Pyruvate metabolism | 411.1352 | 57872.4374 | 413.8824 | 57975.4133 | 2.7472 | -0.0081 | 0.5032 |
| HMR_8508 | Pyruvate metabolism | 411.0285 | 57869.0764 | 410.6923 | 57974.2181 | 0.3363 | 0.001 | 0.4996 |
| HMR_8509 | Pyruvate metabolism | 405.7674 | 57872.3848 | 404.6882 | 57973.1187 | 1.0793 | 0.0032 | 0.4987 |
| HMR_8511 | Pyruvate metabolism | 23.4751 | 156.7447 | 32.6914 | 276.3405 | 9.2163 | -0.4429 | 0.6711 |
| HMR_8512 | Pyruvate metabolism | -16.3396 | 2426.7632 | 16.8103 | 1248.7158 | 33.1499 | -0.5468 | 0.7077 |
| HMR_8514 | Pyruvate metabolism | 57.8271 | 2131.9962 | 33.487 | 893.554 | 24.3401 | 0.4425 | 0.3291 |
| HMR_0153 | Propanoate metabolism | -1.3329 | 220512.8174 | -1.35 | 220512.8061 | 0.0171 | 0.00E+00 | 0.5 |
| HMR_3212 | Propanoate metabolism | 9.8408 | 61.9882 | 8.9893 | 53.6535 | 0.8515 | 0.0792 | 0.4684 |
| HMR_3797 | Propanoate metabolism | -1.4441 | 220512.931 | -1.4817 | 220512.955 | 0.0376 | 1.00E-04 | 0.5 |
| HMR_3800 | Propanoate metabolism | 13.7878 | 97.1058 | 24.487 | 144.8419 | 10.6993 | -0.6878 | 0.7542 |
| HMR_4105 | Propanoate metabolism | 4.4081 | 20.2974 | 4.5321 | 21.2827 | 0.124 | -0.0192 | 0.5077 |
| HMR_4106 | Propanoate metabolism | 4.3508 | 19.6555 | 4.4627 | 20.4827 | 0.1119 | -0.0177 | 0.507 |
| HMR_4107 | Propanoate metabolism | 4.4067 | 6.9116 | 4.5249 | 7.2897 | 0.1181 | -0.0313 | 0.5125 |
| HMR_4282 | Propanoate metabolism | 34.722 | 400.2878 | 27.6738 | 286.6804 | 7.0483 | 0.2689 | 0.394 |
| HMR_4331 | Propanoate metabolism | -0.666 | 220511.4306 | -0.6747 | 220511.3941 | 0.0087 | 0.00E+00 | 0.5 |
| HMR_4459 | Propanoate metabolism | 0.667 | 220511.4272 | 0.6753 | 220511.3908 | 0.0083 | 0.00E+00 | 0.5 |
| HMR_4460 | Propanoate metabolism | 0.7221 | 220511.2964 | 0.7406 | 220511.2524 | 0.0185 | 0.00E+00 | 0.5 |
| HMR_4464 | Propanoate metabolism | 9.5666 | 27.5756 | 10.1706 | 31.1627 | 0.6041 | -0.0788 | 0.5314 |
| HMR_4497 | Propanoate metabolism | 19.702 | 75.0447 | 17.9855 | 70.1721 | 1.7164 | 0.1424 | 0.4434 |
| HMR_4741 | Propanoate metabolism | 19.7026 | 75.0373 | 17.9861 | 70.1657 | 1.7165 | 0.1424 | 0.4434 |
| HMR_1434 | Butanoate metabolism | -7.3964 | 94.1988 | -6.7993 | 86.0405 | 0.5971 | -0.0445 | 0.5177 |
| HMR_1436 | Butanoate metabolism | 2.6381 | 7.2165 | 2.6993 | 7.5025 | 0.0612 | -0.0159 | 0.5064 |
| HMR_3163 | Butanoate metabolism | 17.6448 | 6.4963 | 19.5838 | 7.7163 | 1.939 | -0.5143 | 0.6965 |
| HMR_4461 | Butanoate metabolism | 118.8764 | 15525.5669 | 232.3723 | 38166.3946 | 113.4959 | -0.4898 | 0.6879 |
| HMR_4604 | Butanoate metabolism | 1.8334 | 0.9951 | 1.776 | 0.9419 | 0.0574 | 0.0412 | 0.4835 |
| HMR_5351 | Butanoate metabolism | 1.2266 | 246.5171 | 9.6999 | 284.6678 | 8.4733 | -0.3676 | 0.6434 |
| HMR_4052 | Pentose phosphate pathway | 4.3536 | 15.1371 | 4.7184 | 17.1277 | 0.3649 | -0.0642 | 0.5256 |
| HMR_4304 | Pentose phosphate pathway | 463.8326 | 74701.7097 | 463.6696 | 74671.6352 | 0.163 | 0.0004 | 0.4998 |
| HMR_4306 | Pentose phosphate pathway | -10.9194 | 55.5336 | -10.3376 | 49.8226 | 0.5818 | -0.0567 | 0.5226 |
| HMR_4351 | Pentose phosphate pathway | 231.4187 | 152082.7464 | 230.4352 | 152186.6518 | 0.9836 | 0.0018 | 0.4993 |
| HMR_4352 | Pentose phosphate pathway | -226.673 | 152205.1514 | -227.7727 | 152316.0789 | 1.0997 | 0.002 | 0.4992 |
| HMR_4354 | Pentose phosphate pathway | 14.8343 | 58.6996 | 14.5236 | 55.1182 | 0.3107 | 0.0291 | 0.4884 |
| HMR_4398 | Pentose phosphate pathway | 14.4798 | 47.8666 | 13.8786 | 45.0861 | 0.6012 | 0.0624 | 0.4751 |
| HMR_4404 | Pentose phosphate pathway | 20.5396 | 131.1194 | 21.317 | 147.4359 | 0.7773 | -0.0466 | 0.5186 |
| HMR_4473 | Pentose phosphate pathway | -613.2203 | 30586.9335 | -611.9159 | 30745.432 | 1.3044 | -0.0053 | 0.5021 |
| HMR_4474 | Pentose phosphate pathway | 634.7672 | 30803.5196 | 632.3535 | 30937.2192 | 2.4137 | 0.0097 | 0.4961 |
| HMR_4477 | Pentose phosphate pathway | -26.2935 | 257.0356 | -23.1009 | 262.1191 | 3.1925 | -0.1401 | 0.5557 |
| HMR_4501 | Pentose phosphate pathway | 20.5378 | 131.1134 | 21.315 | 147.43 | 0.7772 | -0.0466 | 0.5186 |
| HMR_4565 | Pentose phosphate pathway | 230.0053 | 61654.9242 | 238.8013 | 61820.5611 | 8.796 | -0.025 | 0.51 |
| HMR_4567 | Pentose phosphate pathway | 114.5344 | 237588.4273 | 115.3496 | 237268.2492 | 0.8151 | -0.0012 | 0.5005 |

| | | | | | | | | |
|---|---|---|---|---|---|---|---|---|
| HMR_4568 | Pentose phosphate pathway | 94.9327 | 238199.6293 | 102.1363 | 237634.277 | 7.2036 | -0.0104 | 0.5042 |
| HMR_4623 | Pentose phosphate pathway | 10.6281 | 134.5923 | 10.1007 | 116.7559 | 0.5274 | 0.0333 | 0.4867 |
| HMR_4625 | Pentose phosphate pathway | 10.9194 | 55.5336 | 10.3375 | 49.8226 | 0.5819 | 0.0567 | 0.4774 |
| HMR_4710 | Pentose phosphate pathway | 14.48 | 47.8578 | 13.8788 | 45.0771 | 0.6012 | 0.0624 | 0.4751 |
| HMR_8653 | Pentose phosphate pathway | 453.2049 | 74784.4493 | 453.5693 | 74743.268 | 0.3644 | -0.0009 | 0.5004 |
| HMR_3998 | Purine metabolism | 2.8475 | 8.2306 | 2.7183 | 7.5014 | 0.1292 | 0.0326 | 0.487 |
| HMR_4000 | Purine metabolism | 2.0134 | 1.8973 | 2.0372 | 1.9426 | 0.0238 | -0.0122 | 0.5049 |
| HMR_4002 | Purine metabolism | 187.703 | 217425.2547 | 189.053 | 217819.5291 | 1.35 | -0.002 | 0.5008 |
| HMR_4004 | Purine metabolism | -382.6905 | 183703.0171 | -390.0134 | 181311.639 | 7.3229 | 0.0121 | 0.4952 |
| HMR_4010 | Purine metabolism | 4.2585 | 20.115 | 4.215 | 19.1139 | 0.0435 | 0.0069 | 0.4972 |
| HMR_4012 | Purine metabolism | 8.2323 | 64.4844 | 7.4361 | 68.0662 | 0.7962 | 0.0692 | 0.4724 |
| HMR_4014 | Purine metabolism | 2.0151 | 1.8974 | 2.0389 | 1.9426 | 0.0238 | -0.0121 | 0.5048 |
| HMR_4016 | Purine metabolism | 234.6171 | 266456.3452 | 231.9121 | 267046.4731 | 2.705 | 0.0037 | 0.4985 |
| HMR_4018 | Purine metabolism | -264.5017 | 253001.7036 | -268.6148 | 251952.8318 | 4.1131 | 0.0058 | 0.4977 |
| HMR_4020 | Purine metabolism | -16.7637 | 25.3927 | -16.5775 | 24.7482 | 0.1861 | -0.0263 | 0.5105 |
| HMR_4022 | Purine metabolism | 4.0284 | 4.0172 | 4.0785 | 4.1179 | 0.05 | -0.0175 | 0.507 |
| HMR_4038 | Purine metabolism | 0.5285 | 0.0696 | 0.5155 | 0.0668 | 0.0131 | 0.0354 | 0.4859 |
| HMR_4040 | Purine metabolism | 5.582 | 10.721 | 5.4885 | 10.2296 | 0.0934 | 0.0204 | 0.4919 |
| HMR_4042 | Purine metabolism | 2.9072 | 3.6289 | 2.8122 | 3.3961 | 0.095 | 0.0359 | 0.4857 |
| HMR_4044 | Purine metabolism | -78.0192 | 299154.7268 | -89.086 | 298058.3728 | 11.0668 | 0.0143 | 0.4943 |
| HMR_4046 | Purine metabolism | 2.064 | 3.5344 | 2.0166 | 3.4039 | 0.0474 | 0.018 | 0.4928 |
| HMR_4048 | Purine metabolism | 1.9335 | 3.0347 | 1.9568 | 3.1099 | 0.0233 | -0.0094 | 0.5037 |
| HMR_4080 | Purine metabolism | 3.9475 | 8.7231 | 3.7558 | 8.0484 | 0.1917 | 0.0468 | 0.4813 |
| HMR_4081 | Purine metabolism | 22.2902 | 44.3891 | 21.711 | 41.7688 | 0.5792 | 0.0624 | 0.4751 |
| HMR_4082 | Purine metabolism | 4.3509 | 20.8578 | 4.462 | 21.2674 | 0.111 | -0.0171 | 0.5068 |
| HMR_4085 | Purine metabolism | 17.6336 | 39.1966 | 16.5379 | 37.8349 | 1.0957 | 0.1248 | 0.4503 |
| HMR_4168 | Purine metabolism | 2.01 | 4.2007 | 2.0344 | 4.2795 | 0.0244 | -0.0084 | 0.5033 |
| HMR_4171 | Purine metabolism | 252.369 | 44235.8753 | 250.2913 | 43720.0938 | 2.0777 | 0.007 | 0.4972 |
| HMR_4291 | Purine metabolism | -115.2181 | 2428.3758 | -113.6286 | 2388.0858 | 1.5895 | -0.0229 | 0.5091 |
| HMR_4406 | Purine metabolism | 0.4979 | 0.0661 | 0.484 | 0.0631 | 0.0139 | 0.0387 | 0.4846 |
| HMR_4412 | Purine metabolism | -2.9065 | 3.6299 | -2.8115 | 3.3971 | 0.0951 | -0.0359 | 0.5143 |
| HMR_4419 | Purine metabolism | 2.2677 | 3.7901 | 2.278 | 3.8148 | 0.0103 | -0.0037 | 0.5015 |
| HMR_4420 | Purine metabolism | 509.2385 | 80565.6762 | 512.2033 | 80021.2185 | 2.9647 | -0.0074 | 0.503 |
| HMR_4421 | Purine metabolism | 300.4233 | 54796.281 | 299.1562 | 54491.6748 | 1.2671 | 0.0038 | 0.4985 |
| HMR_4449 | Purine metabolism | 3.135 | 8.6819 | 2.9951 | 7.9651 | 0.1399 | 0.0343 | 0.4863 |
| HMR_4452 | Purine metabolism | -11.3508 | 29.3959 | -10.8215 | 28.0193 | 0.5293 | -0.0699 | 0.5278 |
| HMR_4453 | Purine metabolism | 4.7648 | 8.9395 | 4.8254 | 9.0181 | 0.0606 | -0.0143 | 0.5057 |
| HMR_4480 | Purine metabolism | -54.1613 | 237287.2542 | -62.5316 | 236628.142 | 8.3703 | 0.0122 | 0.4951 |
| HMR_4486 | Purine metabolism | 38.9293 | 1194.4435 | 38.6448 | 1179.3174 | 0.2846 | 0.0058 | 0.4977 |
| HMR_4487 | Purine metabolism | 38.4408 | 1207.5495 | 38.1341 | 1193.3741 | 0.3068 | 0.0063 | 0.4975 |
| HMR_4488 | Purine metabolism | 40.932 | 1258.6803 | 40.6351 | 1241.1531 | 0.2968 | 0.0059 | 0.4976 |
| HMR_4489 | Purine metabolism | 40.3803 | 1271.4155 | 40.0546 | 1254.42 | 0.3257 | 0.0065 | 0.4974 |
| HMR_4495 | Purine metabolism | 20.6661 | 343.1963 | 19.0772 | 293.3825 | 1.5889 | 0.063 | 0.4749 |
| HMR_4496 | Purine metabolism | 20.4225 | 339.2858 | 18.8488 | 289.7303 | 1.5738 | 0.0627 | 0.475 |

| | | | | | | | | |
|---|---|---|---|---|---|---|---|---|
| HMR_4519 | Purine metabolism | 11.319 | 61.506 | 10.5597 | 55.344 | 0.7594 | 0.0703 | 0.472 |
| HMR_4520 | Purine metabolism | 7.8068 | 42.4878 | 7.5367 | 39.2274 | 0.2701 | 0.0299 | 0.4881 |
| HMR_4570 | Purine metabolism | 7.2267 | 289919.0453 | -6.1 | 289967.5648 | 13.3267 | 0.0175 | 0.493 |
| HMR_4572 | Purine metabolism | 336.4413 | 14713.2099 | 366.5177 | 17494.1914 | 30.0764 | -0.1676 | 0.5665 |
| HMR_4573 | Purine metabolism | 285.0683 | 51245.4992 | 284.0764 | 50992.0295 | 0.9919 | 0.0031 | 0.4988 |
| HMR_4600 | Purine metabolism | 2.8289 | 9.5952 | 2.7014 | 8.7147 | 0.1275 | 0.0298 | 0.4881 |
| HMR_4602 | Purine metabolism | 13.6805 | 28.3676 | 13.2132 | 26.4517 | 0.4673 | 0.0631 | 0.4748 |
| HMR_4603 | Purine metabolism | -7.8626 | 16.4229 | -7.6734 | 15.6356 | 0.1892 | -0.0334 | 0.5133 |
| HMR_4611 | Purine metabolism | 3.8638 | 11.4403 | 3.7717 | 10.8924 | 0.092 | 0.0195 | 0.4922 |
| HMR_4612 | Purine metabolism | 3.8612 | 11.4285 | 3.7689 | 10.8798 | 0.0923 | 0.0195 | 0.4922 |
| HMR_4614 | Purine metabolism | 3.936 | 11.0714 | 3.8422 | 10.5458 | 0.0938 | 0.0202 | 0.492 |
| HMR_4615 | Purine metabolism | 3.9283 | 11.0405 | 3.8346 | 10.5159 | 0.0938 | 0.0202 | 0.4919 |
| HMR_4617 | Purine metabolism | 4.5077 | 12.2329 | 4.3614 | 11.4613 | 0.1463 | 0.0301 | 0.488 |
| HMR_4618 | Purine metabolism | 13.3057 | 121.8506 | 14.2632 | 139.3055 | 0.9574 | -0.0592 | 0.5236 |
| HMR_4619 | Purine metabolism | 4.5031 | 12.2197 | 4.3569 | 11.4488 | 0.1462 | 0.03 | 0.488 |
| HMR_4621 | Purine metabolism | 13.3005 | 121.7917 | 14.2593 | 139.2621 | 0.9587 | -0.0593 | 0.5237 |
| HMR_4632 | Purine metabolism | 10.5517 | 24.636 | 10.2394 | 23.1931 | 0.3123 | 0.0452 | 0.482 |
| HMR_4646 | Purine metabolism | 19.5624 | 118.0773 | 18.3967 | 109.8745 | 1.1657 | 0.0772 | 0.4692 |
| HMR_4649 | Purine metabolism | 11.7649 | 98.2106 | 11.5726 | 92.4759 | 0.1924 | 0.0139 | 0.4944 |
| HMR_4651 | Purine metabolism | 3.4896 | 16.7264 | 3.1499 | 15.8557 | 0.3397 | 0.0595 | 0.4763 |
| HMR_4663 | Purine metabolism | 7.513 | 11.835 | 7.3408 | 11.175 | 0.1721 | 0.0359 | 0.4857 |
| HMR_4664 | Purine metabolism | 4.1522 | 10.2272 | 4.0296 | 9.6405 | 0.1227 | 0.0275 | 0.489 |
| HMR_4694 | Purine metabolism | 17.8541 | 76.4814 | 16.9539 | 70.2907 | 0.9002 | 0.0743 | 0.4704 |
| HMR_4695 | Purine metabolism | 18.9571 | 39.242 | 17.8287 | 37.8638 | 1.1283 | 0.1285 | 0.4489 |
| HMR_4705 | Purine metabolism | 4.5827 | 10.735 | 4.363 | 9.9676 | 0.2197 | 0.0483 | 0.4807 |
| HMR_4709 | Purine metabolism | -27.9812 | 36.6073 | -26.6728 | 34.5236 | 1.3083 | -0.1551 | 0.5616 |
| HMR_4799 | Purine metabolism | 0.5007 | 0.0597 | 0.4868 | 0.0569 | 0.0138 | 0.0405 | 0.4839 |
| HMR_4802 | Purine metabolism | 0.5091 | 0.0531 | 0.4954 | 0.0504 | 0.0137 | 0.0426 | 0.483 |
| HMR_4804 | Purine metabolism | -0.5154 | 0.0536 | -0.5019 | 0.051 | 0.0135 | -0.0417 | 0.5166 |
| HMR_4808 | Purine metabolism | 0.504 | 0.0538 | 0.4901 | 0.0511 | 0.0139 | 0.0428 | 0.4829 |
| HMR_4810 | Purine metabolism | 0.519 | 0.0553 | 0.5056 | 0.0526 | 0.0134 | 0.0408 | 0.4837 |
| HMR_4812 | Purine metabolism | -0.5232 | 0.0582 | -0.5099 | 0.0556 | 0.0133 | -0.0394 | 0.5157 |
| HMR_4814 | Purine metabolism | 0.5269 | 0.0628 | 0.5137 | 0.0601 | 0.0132 | 0.0376 | 0.485 |
| HMR_5301 | Purine metabolism | 14.7235 | 157.4618 | 15.3035 | 170.4375 | 0.58 | -0.032 | 0.5128 |
| HMR_5353 | Purine metabolism | 7.3589 | 16.5207 | 7.1645 | 15.7354 | 0.1944 | 0.0342 | 0.4864 |
| HMR_6605 | Purine metabolism | -6.9663 | 220504.859 | -8.5789 | 220500.8947 | 1.6126 | 0.0024 | 0.499 |
| HMR_6606 | Purine metabolism | 10.4485 | 28.5459 | 12.8686 | 42.6146 | 2.4201 | -0.2869 | 0.6129 |
| HMR_6607 | Purine metabolism | -3.4843 | 220495.3983 | -4.2914 | 220486.8085 | 0.807 | 0.0012 | 0.4995 |
| HMR_7144 | Purine metabolism | 14.8016 | 98.4445 | 14.1711 | 90.6247 | 0.6305 | 0.0459 | 0.4817 |
| HMR_8755 | Purine metabolism | 8.8478 | 25.8453 | 8.5688 | 24.1678 | 0.279 | 0.0394 | 0.4843 |
| HMR_3931 | Pyrimidine metabolism | 7.3428 | 35.9612 | 7.0478 | 33.3547 | 0.2951 | 0.0354 | 0.4859 |
| HMR_3969 | Pyrimidine metabolism | 14.6952 | 156.2506 | 15.2763 | 169.0466 | 0.5811 | -0.0322 | 0.5129 |
| HMR_3970 | Pyrimidine metabolism | 3.3313 | 13.1035 | 3.1423 | 11.329 | 0.189 | 0.0382 | 0.4848 |
| HMR_4006 | Pyrimidine metabolism | 15.152 | 298725.3054 | 10.3107 | 298767.0784 | 4.8413 | 0.0063 | 0.4975 |

| | | | | | | | | |
|---|---|---|---|---|---|---|---|---|
| HMR_4008 | Pyrimidine metabolism | 21.1694 | 305194.5635 | 13.8828 | 305304.449 | 7.2866 | 0.0093 | 0.4963 |
| HMR_4028 | Pyrimidine metabolism | 80.0181 | 285963.6719 | 73.8405 | 286426.9486 | 6.1776 | 0.0082 | 0.4967 |
| HMR_4032 | Pyrimidine metabolism | 8.3998 | 19.465 | 8.095 | 18.0577 | 0.3049 | 0.0498 | 0.4802 |
| HMR_4034 | Pyrimidine metabolism | 0.9626 | 0.2327 | 0.9286 | 0.2181 | 0.0339 | 0.0505 | 0.4798 |
| HMR_4036 | Pyrimidine metabolism | 0.9894 | 0.2279 | 0.956 | 0.2136 | 0.0334 | 0.0503 | 0.4799 |
| HMR_4050 | Pyrimidine metabolism | -0.9686 | 0.2261 | -0.9349 | 0.2116 | 0.0337 | -0.051 | 0.5203 |
| HMR_4056 | Pyrimidine metabolism | 3.7241 | 11.613 | 3.5341 | 10.597 | 0.19 | 0.0403 | 0.4839 |
| HMR_4059 | Pyrimidine metabolism | 8.6492 | 3.8105 | 8.6413 | 3.7912 | 0.0078 | 0.0028 | 0.4989 |
| HMR_4127 | Pyrimidine metabolism | 2.0103 | 4.1564 | 2.0346 | 4.2325 | 0.0244 | -0.0084 | 0.5034 |
| HMR_4177 | Pyrimidine metabolism | 4.5032 | 15.3561 | 4.3154 | 14.024 | 0.1879 | 0.0347 | 0.4862 |
| HMR_4179 | Pyrimidine metabolism | 7.2121 | 13.2884 | 6.6957 | 11.5846 | 0.5165 | 0.1036 | 0.4588 |
| HMR_4182 | Pyrimidine metabolism | 4.8755 | 17.3084 | 4.6134 | 15.5075 | 0.262 | 0.0457 | 0.4818 |
| HMR_4183 | Pyrimidine metabolism | 7.2032 | 13.2852 | 6.6866 | 11.5815 | 0.5166 | 0.1036 | 0.4587 |
| HMR_4192 | Pyrimidine metabolism | 7.1943 | 13.2856 | 6.6775 | 11.5819 | 0.5168 | 0.1036 | 0.4587 |
| HMR_4194 | Pyrimidine metabolism | 2.5197 | 5.745 | 2.4104 | 5.2549 | 0.1093 | 0.033 | 0.4869 |
| HMR_4210 | Pyrimidine metabolism | 284.3413 | 51243.3003 | 282.1577 | 50739.0283 | 2.1836 | 0.0068 | 0.4973 |
| HMR_4345 | Pyrimidine metabolism | -1.4265 | 0.4818 | -1.3954 | 0.4648 | 0.0311 | -0.0319 | 0.5127 |
| HMR_4346 | Pyrimidine metabolism | 1.4272 | 0.4857 | 1.3959 | 0.4685 | 0.0313 | 0.032 | 0.4872 |
| HMR_4484 | Pyrimidine metabolism | 5.7355 | 22.6724 | 5.955 | 23.7391 | 0.2195 | -0.0322 | 0.5129 |
| HMR_4510 | Pyrimidine metabolism | 5.0455 | 8.1053 | 4.8183 | 7.3992 | 0.2272 | 0.0577 | 0.477 |
| HMR_4512 | Pyrimidine metabolism | 30.0656 | 304944.2889 | 23.1413 | 305115.4604 | 6.9243 | 0.0089 | 0.4965 |
| HMR_4513 | Pyrimidine metabolism | 2.7862 | 5.8185 | 2.6605 | 5.3056 | 0.1256 | 0.0377 | 0.485 |
| HMR_4514 | Pyrimidine metabolism | 2.274 | 3.7217 | 2.1727 | 3.4077 | 0.1013 | 0.0379 | 0.4849 |
| HMR_4577 | Pyrimidine metabolism | -0.9837 | 0.2235 | -0.9503 | 0.2091 | 0.0334 | -0.0508 | 0.5203 |
| HMR_4579 | Pyrimidine metabolism | 3.8313 | 42.7451 | 4.0324 | 40.3445 | 0.2011 | -0.0221 | 0.5088 |
| HMR_4580 | Pyrimidine metabolism | 2.1162 | 3.9143 | 1.9966 | 3.4967 | 0.1197 | 0.044 | 0.4825 |
| HMR_4608 | Pyrimidine metabolism | -0.975 | 0.2225 | -0.9415 | 0.208 | 0.0335 | -0.051 | 0.5203 |
| HMR_4637 | Pyrimidine metabolism | -127.6541 | 1739.582 | -101.6279 | 1456.8417 | 26.0262 | -0.4603 | 0.6774 |
| HMR_4642 | Pyrimidine metabolism | 5.0484 | 20.2672 | 4.7317 | 18.069 | 0.3167 | 0.0512 | 0.4796 |
| HMR_4643 | Pyrimidine metabolism | 5.49 | 9.727 | 5.5729 | 10.0362 | 0.0829 | -0.0187 | 0.5074 |
| HMR_4670 | Pyrimidine metabolism | -45.1907 | 286871.5403 | -56.7342 | 286278.4375 | 11.5435 | 0.0152 | 0.4939 |
| HMR_4672 | Pyrimidine metabolism | 3.3903 | 8.2216 | 3.2947 | 7.7548 | 0.0955 | 0.0239 | 0.4905 |
| HMR_4673 | Pyrimidine metabolism | -52.7475 | 408.7965 | -54.6987 | 449.1735 | 1.9512 | 0.0666 | 0.4734 |
| HMR_4675 | Pyrimidine metabolism | 9.1348 | 69.2094 | 9.7285 | 78.3489 | 0.5937 | -0.0489 | 0.5195 |
| HMR_4676 | Pyrimidine metabolism | 4.8631 | 17.2449 | 4.6025 | 15.4542 | 0.2606 | 0.0456 | 0.4818 |
| HMR_5299 | Pyrimidine metabolism | 14.6125 | 155.7553 | 15.185 | 168.5375 | 0.5725 | -0.0318 | 0.5127 |
| HMR_5352 | Pyrimidine metabolism | 14.6125 | 155.7553 | 15.185 | 168.5375 | 0.5725 | -0.0318 | 0.5127 |
| HMR_6612 | Pyrimidine metabolism | 127.1533 | 1739.2724 | 101.1214 | 1456.6706 | 26.0319 | 0.4605 | 0.3226 |
| HMR_6614 | Pyrimidine metabolism | -343.6587 | 193871.4172 | -348.4428 | 191808.9079 | 4.7841 | 0.0077 | 0.4969 |
| HMR_6615 | Pyrimidine metabolism | 48.1721 | 94299.7855 | 49.0535 | 95556.5955 | 0.8814 | -0.002 | 0.5008 |
| HMR_6616 | Pyrimidine metabolism | -88.0855 | 237688.2276 | -92.5731 | 237018.4225 | 4.4877 | 0.0065 | 0.4974 |
| HMR_6621 | Pyrimidine metabolism | 53.6199 | 172.9157 | 56.6319 | 193.2668 | 3.012 | -0.1574 | 0.5625 |
| HMR_6622 | Pyrimidine metabolism | 15.5003 | 68.2041 | 16.6568 | 78.648 | 1.1565 | -0.0954 | 0.538 |
| HMR_6623 | Pyrimidine metabolism | 4.8687 | 19.1135 | 4.7754 | 18.3467 | 0.0933 | 0.0152 | 0.4939 |

| | | | | | | | | |
|---|---|---|---|---|---|---|---|---|
| HMR_6624 | Pyrimidine metabolism | 4.3547 | 14.8398 | 4.4652 | 15.6088 | 0.1105 | -0.02 | 0.508 |
| HMR_6627 | Pyrimidine metabolism | -609.1076 | 87532.3412 | -611.0316 | 86632.4361 | 1.924 | 0.0046 | 0.4982 |
| HMR_3965 | Nucleotide metabolism | 2.8449 | 8.2142 | 2.7158 | 7.487 | 0.129 | 0.0326 | 0.487 |
| HMR_3966 | Nucleotide metabolism | 2.1714 | 4.8951 | 2.227 | 5.1003 | 0.0556 | -0.0176 | 0.507 |
| HMR_3967 | Nucleotide metabolism | -223.8629 | 257801.4375 | -227.6959 | 256895.8175 | 3.833 | 0.0053 | 0.4979 |
| HMR_3968 | Nucleotide metabolism | -15.6957 | 63.7573 | -15.6672 | 60.7268 | 0.0285 | -0.0026 | 0.501 |
| HMR_3999 | Nucleotide metabolism | 16.9177 | 55.9273 | 16.5413 | 53.1666 | 0.3764 | 0.036 | 0.4856 |
| HMR_4024 | Nucleotide metabolism | 25.2246 | 305091.7265 | 18.3332 | 305225.8458 | 6.8914 | 0.0088 | 0.4965 |
| HMR_4054 | Nucleotide metabolism | 6.4644 | 26.4665 | 6.3985 | 25.9877 | 0.0659 | 0.0091 | 0.4964 |
| HMR_4083 | Nucleotide metabolism | 32.7005 | 154.2802 | 35.4357 | 158.2806 | 2.7352 | -0.1547 | 0.5615 |
| HMR_4180 | Nucleotide metabolism | 4.8631 | 17.2449 | 4.6025 | 15.4542 | 0.2606 | 0.0456 | 0.4818 |
| HMR_4185 | Nucleotide metabolism | 13.4158 | 11.4071 | 13.2663 | 11.2216 | 0.1494 | 0.0314 | 0.4875 |
| HMR_4186 | Nucleotide metabolism | 16.2636 | 19.6089 | 15.9852 | 18.6855 | 0.2784 | 0.045 | 0.4821 |
| HMR_4483 | Nucleotide metabolism | 6.9656 | 28.7306 | 6.8412 | 27.9985 | 0.1243 | 0.0165 | 0.4934 |
| HMR_4507 | Nucleotide metabolism | 3.3905 | 8.2225 | 3.2954 | 7.757 | 0.0951 | 0.0238 | 0.4905 |
| HMR_4516 | Nucleotide metabolism | 2.8455 | 8.2183 | 2.7163 | 7.4898 | 0.1292 | 0.0326 | 0.487 |
| HMR_4601 | Nucleotide metabolism | 2.9914 | 3.174 | 2.8404 | 2.8622 | 0.151 | 0.0615 | 0.4755 |
| HMR_4633 | Nucleotide metabolism | 2.8429 | 7.5437 | 2.7154 | 6.8954 | 0.1276 | 0.0336 | 0.4866 |
| HMR_4635 | Nucleotide metabolism | -386.2437 | 192925.0666 | -398.2171 | 188652.952 | 11.9733 | 0.0194 | 0.4923 |
| HMR_4640 | Nucleotide metabolism | 2.7519 | 5.8904 | 2.7932 | 6.0731 | 0.0413 | -0.0119 | 0.5048 |
| HMR_4641 | Nucleotide metabolism | 4.8683 | 19.1112 | 4.7757 | 18.3487 | 0.0927 | 0.0151 | 0.494 |
| HMR_4680 | Nucleotide metabolism | 2.4065 | 42.4246 | 2.6385 | 40.0336 | 0.232 | -0.0256 | 0.5102 |
| HMR_4806 | Nucleotide metabolism | 0.5034 | 0.0559 | 0.4895 | 0.0531 | 0.0139 | 0.042 | 0.4833 |
| HMR_5394 | Nucleotide metabolism | 2.9307 | 5.3794 | 2.9784 | 5.4671 | 0.0476 | -0.0145 | 0.5058 |
| HMR_6625 | Nucleotide metabolism | 4.3561 | 14.8477 | 4.4665 | 15.6166 | 0.1105 | -0.02 | 0.508 |
| HMR_6626 | Nucleotide metabolism | 8.7032 | 24.6571 | 8.9243 | 25.9444 | 0.2211 | -0.0311 | 0.5124 |
| HMR_7160 | Nucleotide metabolism | 2.0107 | 1.8924 | 2.0343 | 1.9372 | 0.0236 | -0.0121 | 0.5048 |
| HMR_7161 | Nucleotide metabolism | 2.0499 | 3.4934 | 2.0681 | 3.524 | 0.0183 | -0.0069 | 0.5027 |
| HMR_7162 | Nucleotide metabolism | 2.7918 | 5.7132 | 2.6737 | 5.3013 | 0.1182 | 0.0356 | 0.4858 |
| HMR_7163 | Nucleotide metabolism | 2.0126 | 1.8925 | 2.0361 | 1.9372 | 0.0235 | -0.012 | 0.5048 |
| HMR_7164 | Nucleotide metabolism | 4.8456 | 7.736 | 4.7456 | 7.394 | 0.0999 | 0.0257 | 0.4898 |
| HMR_7713 | Nucleotide metabolism | 4.2972 | 6.5621 | 4.2456 | 6.4088 | 0.0517 | 0.0143 | 0.4943 |
| HMR_7716 | Nucleotide metabolism | 3.376 | 4.0466 | 3.179 | 3.5902 | 0.197 | 0.0713 | 0.4716 |
| HMR_7717 | Nucleotide metabolism | 53.6304 | 172.9371 | 56.6427 | 193.2887 | 3.0123 | -0.1574 | 0.5625 |
| HMR_7721 | Nucleotide metabolism | 4.9751 | 8.3709 | 4.7759 | 7.703 | 0.1992 | 0.0497 | 0.4802 |
| HMR_7725 | Nucleotide metabolism | 3.9702 | 5.4592 | 3.7504 | 4.8848 | 0.2198 | 0.0683 | 0.4728 |
| HMR_7728 | Nucleotide metabolism | 3.2796 | 3.7631 | 3.1277 | 3.4259 | 0.1519 | 0.0566 | 0.4774 |
| HMR_7800 | Nucleotide metabolism | -168.6584 | 217239.9474 | -162.5157 | 217653.6098 | 6.1427 | -0.0093 | 0.5037 |
| HMR_7801 | Nucleotide metabolism | -148.3964 | 183837.9955 | -158.3124 | 181467.4482 | 9.916 | 0.0164 | 0.4935 |
| HMR_7802 | Nucleotide metabolism | 61.3709 | 237282.3867 | 69.5507 | 236623.6828 | 8.1798 | -0.0119 | 0.5047 |
| HMR_7863 | Nucleotide metabolism | -60.0926 | 278759.3734 | -61.8183 | 278635.593 | 1.7258 | 0.0023 | 0.4991 |
| HMR_7864 | Nucleotide metabolism | -2.3357 | 280297.4643 | -2.4548 | 280296.8611 | 0.119 | 0.0002 | 0.4999 |
| HMR_7865 | Nucleotide metabolism | 7.242 | 285981.1553 | 4.3464 | 285959.6748 | 2.8956 | 0.0038 | 0.4985 |
| HMR_7866 | Nucleotide metabolism | -2.3373 | 280297.467 | -2.4564 | 280296.8594 | 0.1191 | 0.0002 | 0.4999 |

| | | | | | | | | |
|---|---|---|---|---|---|---|---|---|
| HMR_7867 | Nucleotide metabolism | -227.7826 | 256958.0149 | -231.9709 | 255960.9648 | 4.1883 | 0.0058 | 0.4977 |
| HMR_7868 | Nucleotide metabolism | -2.34 | 280297.4571 | -2.4591 | 280296.8545 | 0.1191 | 0.0002 | 0.4999 |
| HMR_7869 | Nucleotide metabolism | 106.644 | 282413.3981 | 110.6633 | 281888.5732 | 4.0192 | -0.0054 | 0.5021 |
| HMR_7870 | Nucleotide metabolism | 20.1306 | 285116.4492 | 21.1559 | 285077.1857 | 1.0253 | -0.0014 | 0.5005 |
| HMR_7871 | Nucleotide metabolism | 71.4678 | 277031.6443 | 75.2202 | 276731.9848 | 3.7524 | -0.005 | 0.502 |
| HMR_7872 | Nucleotide metabolism | -2.34 | 280297.4578 | -2.4592 | 280296.8545 | 0.1191 | 0.0002 | 0.4999 |
| HMR_7873 | Nucleotide metabolism | 16.5423 | 281078.4404 | 22.2176 | 280990.3702 | 5.6753 | -0.0076 | 0.503 |
| HMR_7874 | Nucleotide metabolism | -2.34 | 280297.4574 | -2.4591 | 280296.856 | 0.1191 | 0.0002 | 0.4999 |
| HMR_7875 | Nucleotide metabolism | 53.6274 | 172.9299 | 56.6396 | 193.2813 | 3.0122 | -0.1574 | 0.5625 |
| HMR_7876 | Nucleotide metabolism | -8.4423 | 304274.9608 | -8.8721 | 304267.989 | 0.4298 | 0.0006 | 0.4998 |
| HMR_7878 | Nucleotide metabolism | -186.7361 | 130175.7848 | -195.8193 | 128041.8975 | 9.0833 | 0.0179 | 0.4929 |
| HMR_7879 | Nucleotide metabolism | 1.3235 | 0.538 | 1.3282 | 0.5418 | 0.0047 | -0.0046 | 0.5018 |
| HMR_7880 | Nucleotide metabolism | 1.3208 | 0.5427 | 1.3254 | 0.5465 | 0.0046 | -0.0044 | 0.5018 |
| HMR_7881 | Nucleotide metabolism | 186.7448 | 130175.7701 | 195.8281 | 128041.8828 | 9.0833 | -0.0179 | 0.5071 |
| HMR_7882 | Nucleotide metabolism | -7.0193 | 286654.3512 | -7.3766 | 286649.5785 | 0.3573 | 0.0005 | 0.4998 |
| HMR_7883 | Nucleotide metabolism | 53.6239 | 172.9227 | 56.636 | 193.274 | 3.0121 | -0.1574 | 0.5625 |
| HMR_7884 | Nucleotide metabolism | -7.0218 | 286654.335 | -7.3792 | 286649.5605 | 0.3574 | 0.0005 | 0.4998 |
| HMR_7885 | Nucleotide metabolism | -7.0233 | 286654.3246 | -7.3807 | 286649.5499 | 0.3574 | 0.0005 | 0.4998 |
| HMR_7886 | Nucleotide metabolism | 272.322 | 98236.5036 | 296.4863 | 99867.7183 | 24.1643 | -0.0543 | 0.5216 |
| HMR_7887 | Nucleotide metabolism | -112.115 | 252412.9708 | -117.8432 | 251413.8815 | 5.7282 | 0.0081 | 0.4968 |
| HMR_7888 | Nucleotide metabolism | -7.0193 | 286654.3501 | -7.3766 | 286649.5789 | 0.3573 | 0.0005 | 0.4998 |
| HMR_7889 | Nucleotide metabolism | 13.4929 | 70.1325 | 14.6256 | 80.6135 | 1.1327 | -0.0923 | 0.5368 |
| HMR_7890 | Nucleotide metabolism | 274.867 | 12071.7946 | 298.4248 | 14363.1457 | 23.5579 | -0.1449 | 0.5576 |
| HMR_7891 | Nucleotide metabolism | -7.0192 | 286654.3504 | -7.3765 | 286649.5798 | 0.3573 | 0.0005 | 0.4998 |
| HMR_7892 | Nucleotide metabolism | -27.6929 | 296285.0354 | -29.1048 | 296211.01 | 1.4119 | 0.0018 | 0.4993 |
| HMR_8085 | Nucleotide metabolism | 2.1058 | 1.305 | 2.1319 | 1.338 | 0.0262 | -0.0161 | 0.5064 |
| HMR_8443 | Nucleotide metabolism | 17.9529 | 90.1558 | 17.1973 | 82.8106 | 0.7556 | 0.0575 | 0.4771 |
| HMR_8448 | Nucleotide metabolism | 21.1652 | 280992.355 | 27.282 | 280867.1914 | 6.1169 | -0.0082 | 0.5033 |
| HMR_8449 | Nucleotide metabolism | -2.3404 | 280297.4561 | -2.4596 | 280296.8494 | 0.1191 | 0.0002 | 0.4999 |
| HMR_8450 | Nucleotide metabolism | 26.6826 | 280864.0303 | 32.7503 | 280708.0123 | 6.0677 | -0.0081 | 0.5032 |
| HMR_8451 | Nucleotide metabolism | -2.3392 | 280297.4595 | -2.4583 | 280296.8536 | 0.1191 | 0.0002 | 0.4999 |
| HMR_8452 | Nucleotide metabolism | 81.5241 | 276301.22 | 85.6571 | 275935.0075 | 4.1329 | -0.0056 | 0.5022 |
| HMR_8453 | Nucleotide metabolism | -2.3393 | 280297.453 | -2.4584 | 280296.856 | 0.1191 | 0.0002 | 0.4999 |
| HMR_8454 | Nucleotide metabolism | 116.0595 | 281360.6037 | 120.4303 | 280756.3443 | 4.3708 | -0.0058 | 0.5023 |
| HMR_8455 | Nucleotide metabolism | 20.1314 | 285116.4323 | 21.1567 | 285077.1694 | 1.0253 | -0.0014 | 0.5005 |
| HMR_8456 | Nucleotide metabolism | 17.1203 | 285862.8613 | 14.6218 | 285864.4284 | 2.4985 | 0.0033 | 0.4987 |
| HMR_8457 | Nucleotide metabolism | -2.3365 | 280297.4648 | -2.4556 | 280296.8617 | 0.1191 | 0.0002 | 0.4999 |
| HMR_8458 | Nucleotide metabolism | -8.4427 | 304274.9538 | -8.8726 | 304267.9847 | 0.4298 | 0.0006 | 0.4998 |
| HMR_8459 | Nucleotide metabolism | -8.4417 | 304274.9596 | -8.8715 | 304267.9986 | 0.4298 | 0.0006 | 0.4998 |
| HMR_8460 | Nucleotide metabolism | -2.335 | 280297.4669 | -2.454 | 280296.8637 | 0.119 | 0.0002 | 0.4999 |
| HMR_8461 | Nucleotide metabolism | -219.157 | 258792.1227 | -223.0498 | 257889.8654 | 3.8928 | 0.0054 | 0.4978 |
| HMR_8462 | Nucleotide metabolism | -2.3392 | 280297.4587 | -2.4583 | 280296.8527 | 0.1191 | 0.0002 | 0.4999 |
| HMR_8463 | Nucleotide metabolism | -49.9993 | 279286.1084 | -51.3283 | 279198.1695 | 1.3289 | 0.0018 | 0.4993 |
| HMR_8464 | Nucleotide metabolism | -2.3404 | 280297.4555 | -2.4596 | 280296.8507 | 0.1191 | 0.0002 | 0.4999 |

| | | | | | | | | |
|---|---|---|---|---|---|---|---|---|
| HMR_8465 | Nucleotide metabolism | -55.4981 | 279010.1162 | -56.7825 | 278917.8199 | 1.2844 | 0.0017 | 0.4993 |
| HMR_8466 | Nucleotide metabolism | -2.3362 | 280297.466 | -2.4553 | 280296.8572 | 0.1191 | 0.0002 | 0.4999 |
| HMR_8467 | Nucleotide metabolism | 11.7444 | 285941.5736 | 9.2853 | 285929.4634 | 2.4591 | 0.0033 | 0.4987 |
| HMR_8468 | Nucleotide metabolism | -2.3378 | 280297.4615 | -2.4569 | 280296.861 | 0.1191 | 0.0002 | 0.4999 |
| HMR_8469 | Nucleotide metabolism | 110.9422 | 281943.38 | 115.3655 | 281354.865 | 4.4234 | -0.0059 | 0.5024 |
| HMR_8470 | Nucleotide metabolism | 20.1302 | 285116.4557 | 21.1555 | 285077.1983 | 1.0253 | -0.0014 | 0.5005 |
| HMR_8471 | Nucleotide metabolism | 76.0571 | 276710.0131 | 80.2436 | 276360.9827 | 4.1864 | -0.0056 | 0.5022 |
| HMR_8472 | Nucleotide metabolism | -2.3405 | 280297.4556 | -2.4596 | 280296.853 | 0.1191 | 0.0002 | 0.4999 |
| HMR_8473 | Nucleotide metabolism | 8.8092 | 46.7283 | 8.3964 | 42.4155 | 0.4128 | 0.0437 | 0.4826 |
| HMR_8474 | Nucleotide metabolism | 5.3726 | 19.2839 | 5.1698 | 17.9013 | 0.2028 | 0.0332 | 0.4867 |
| HMR_8482 | Nucleotide metabolism | 7.3742 | 36.1605 | 7.0637 | 33.4547 | 0.3105 | 0.0372 | 0.4852 |
| HMR_8483 | Nucleotide metabolism | 2.0115 | 1.8972 | 2.0352 | 1.9424 | 0.0237 | -0.0121 | 0.5048 |
| HMR_8488 | Nucleotide metabolism | 1.6794 | 0.9944 | 1.689 | 1.0045 | 0.0096 | -0.0068 | 0.5027 |
| HMR_8489 | Nucleotide metabolism | 1.7472 | 1.0773 | 1.7533 | 1.0832 | 0.0061 | -0.0042 | 0.5017 |
| HMR_3802 | Alanine, aspartate and glutamate metabolism | 380.0282 | 177894.7946 | 290.2012 | 204503.407 | 89.827 | 0.1453 | 0.4423 |
| HMR_3804 | Alanine, aspartate and glutamate metabolism | 251.7031 | 178066.2326 | 172.023 | 204872.2922 | 79.6801 | 0.1288 | 0.4488 |
| HMR_3811 | Alanine, aspartate and glutamate metabolism | 3.4758 | 2.6184 | 3.5614 | 2.7302 | 0.0855 | -0.037 | 0.5148 |
| HMR_3813 | Alanine, aspartate and glutamate metabolism | -3.4742 | 2.6259 | -3.5597 | 2.7377 | 0.0855 | 0.0369 | 0.4853 |
| HMR_3822 | Alanine, aspartate and glutamate metabolism | 61.8081 | 2064.2262 | 120.5889 | 7281.4717 | 58.7807 | -0.608 | 0.7284 |
| HMR_3827 | Alanine, aspartate and glutamate metabolism | -120.6201 | 4185.1396 | -147.5874 | 5947.4511 | 26.9673 | 0.2679 | 0.3944 |
| HMR_3829 | Alanine, aspartate and glutamate metabolism | 120.9761 | 4277.772 | 149.8874 | 6030.5463 | 28.9113 | -0.2848 | 0.6121 |
| HMR_3831 | Alanine, aspartate and glutamate metabolism | 8.7327 | 84.3671 | 8.3812 | 73.2813 | 0.3515 | 0.028 | 0.4888 |
| HMR_3865 | Alanine, aspartate and glutamate metabolism | 26.7648 | 267.5957 | 34.154 | 397.8151 | 7.3891 | -0.2865 | 0.6127 |
| HMR_3870 | Alanine, aspartate and glutamate metabolism | 126.031 | 226739.8258 | 86.5307 | 232346.5797 | 39.5003 | 0.0583 | 0.4768 |
| HMR_3873 | Alanine, aspartate and glutamate metabolism | 0.9682 | 0.7068 | 1.0082 | 0.7658 | 0.04 | -0.0329 | 0.5131 |
| HMR_3890 | Alanine, aspartate and glutamate metabolism | 21.766 | 54.0345 | 19.3002 | 48.3811 | 2.4658 | 0.2437 | 0.4037 |
| HMR_3892 | Alanine, aspartate and glutamate metabolism | 3.3565 | 13.7307 | 3.1143 | 11.4637 | 0.2422 | 0.0482 | 0.4808 |
| HMR_3899 | Alanine, aspartate and glutamate metabolism | -878.2522 | 4904.5319 | -668.7194 | 5398.6806 | 209.5329 | -2.0643 | 0.9805 |
| HMR_3903 | Alanine, aspartate and glutamate metabolism | 1.7091 | 1.6725 | 1.5723 | 1.6915 | 0.1368 | 0.0746 | 0.4703 |
| HMR_4109 | Alanine, aspartate and glutamate metabolism | 24.6383 | 238.0914 | 18.5659 | 131.8803 | 6.0724 | 0.3157 | 0.3761 |
| HMR_4114 | Alanine, aspartate and glutamate metabolism | 2.8703 | 3.8474 | 2.7249 | 3.4663 | 0.1454 | 0.0538 | 0.4786 |
| HMR_4115 | Alanine, aspartate and glutamate metabolism | 3.5677 | 5.8928 | 3.3267 | 5.1141 | 0.241 | 0.0726 | 0.471 |
| HMR_4118 | Alanine, aspartate and glutamate metabolism | 371.4012 | 70199.1918 | 372.2912 | 70342.8691 | 0.8901 | -0.0024 | 0.5009 |
| HMR_4172 | Alanine, aspartate and glutamate metabolism | 1.5128 | 1.4818 | 1.8337 | 2.0447 | 0.3209 | -0.1709 | 0.5678 |
| HMR_4196 | Alanine, aspartate and glutamate metabolism | 2.8696 | 3.8474 | 2.7242 | 3.4662 | 0.1454 | 0.0538 | 0.4786 |
| HMR_4197 | Alanine, aspartate and glutamate metabolism | 3.5661 | 5.8927 | 3.3251 | 5.1139 | 0.2411 | 0.0727 | 0.471 |
| HMR_4287 | Alanine, aspartate and glutamate metabolism | 6.2739 | 26.1621 | 6.7237 | 29.7593 | 0.4498 | -0.0601 | 0.524 |
| HMR_4693 | Alanine, aspartate and glutamate metabolism | 12.5343 | 34.2884 | 13.4364 | 38.2715 | 0.9022 | -0.1059 | 0.5422 |
| HMR_6780 | Alanine, aspartate and glutamate metabolism | 1.5492 | 0.8261 | 1.5061 | 0.7815 | 0.0431 | 0.034 | 0.4865 |
| HMR_7642 | Alanine, aspartate and glutamate metabolism | -514.2668 | 66969.5784 | -401.3674 | 89060.9287 | 112.8994 | -0.2858 | 0.6125 |
| HMR_8654 | Alanine, aspartate and glutamate metabolism | 0.9682 | 0.7068 | 1.0082 | 0.7658 | 0.04 | -0.0329 | 0.5131 |
| HMR_3806 | Arginine and proline metabolism | 926.9322 | 2347.641 | 870.223 | 7750.0844 | 56.7092 | 0.5643 | 0.2863 |
| HMR_3807 | Arginine and proline metabolism | 53.3528 | 940.1139 | 62.5909 | 1621.7839 | 9.2381 | -0.1825 | 0.5724 |
| HMR_3809 | Arginine and proline metabolism | -1.9302 | 1.1277 | -2.0102 | 1.2207 | 0.08 | 0.0522 | 0.4792 |

| | | | | | | | | |
|---|---|---|---|---|---|---|---|---|
| HMR_3819 | Arginine and proline metabolism | -83.1241 | 2349.4113 | -153.989 | 6490.2303 | 70.8648 | 0.7537 | 0.2255 |
| HMR_3820 | Arginine and proline metabolism | -901.6278 | 3441.0324 | -783.3651 | 11523.0125 | 118.2627 | -0.9668 | 0.8332 |
| HMR_3832 | Arginine and proline metabolism | 1.5615 | 1.8701 | 1.567 | 1.9029 | 0.0055 | -0.0028 | 0.5011 |
| HMR_3833 | Arginine and proline metabolism | 47.7135 | 1587.5929 | 81.143 | 4502.6581 | 33.4294 | -0.4284 | 0.6658 |
| HMR_3835 | Arginine and proline metabolism | 50.5074 | 1714.1336 | 89.356 | 5099.4791 | 38.8487 | -0.4706 | 0.6811 |
| HMR_3837 | Arginine and proline metabolism | 13.4321 | 171.5739 | 15.0473 | 302.0202 | 1.6152 | -0.0742 | 0.5296 |
| HMR_3838 | Arginine and proline metabolism | 905.3339 | 4732.2124 | 813.3978 | 16984.1828 | 91.9361 | 0.6239 | 0.2664 |
| HMR_3895 | Arginine and proline metabolism | 4.4684 | 9.3193 | 4.5378 | 9.619 | 0.0694 | -0.0159 | 0.5064 |
| HMR_3897 | Arginine and proline metabolism | 4.468 | 9.3193 | 4.5374 | 9.619 | 0.0695 | -0.016 | 0.5064 |
| HMR_3956 | Arginine and proline metabolism | 162.7914 | 292.1401 | 174.2954 | 341.4103 | 11.504 | -0.457 | 0.6762 |
| HMR_4073 | Arginine and proline metabolism | 1.3237 | 0.3216 | 1.3058 | 0.3053 | 0.0179 | 0.0226 | 0.491 |
| HMR_4077 | Arginine and proline metabolism | 1.33 | 0.3132 | 1.3048 | 0.2968 | 0.0252 | 0.0323 | 0.4871 |
| HMR_4212 | Arginine and proline metabolism | 4.4543 | 6.8799 | 4.5595 | 7.2101 | 0.1052 | -0.028 | 0.5112 |
| HMR_4583 | Arginine and proline metabolism | 921.6891 | 986.1053 | 910.148 | 1486.7445 | 11.5411 | 0.2321 | 0.4082 |
| HMR_4605 | Arginine and proline metabolism | 529.341 | 62232.8024 | 520.4039 | 62536.8034 | 8.9371 | 0.0253 | 0.4899 |
| HMR_4606 | Arginine and proline metabolism | 2.0133 | 1.8973 | 2.0371 | 1.9426 | 0.0238 | -0.0121 | 0.5048 |
| HMR_4607 | Arginine and proline metabolism | 2.0152 | 1.8974 | 2.039 | 1.9427 | 0.0238 | -0.0121 | 0.5048 |
| HMR_4777 | Arginine and proline metabolism | 12.0664 | 17.0272 | 11.6065 | 15.8182 | 0.4599 | 0.0802 | 0.468 |
| HMR_4779 | Arginine and proline metabolism | 4.0153 | 11.5721 | 3.8766 | 10.7752 | 0.1387 | 0.0293 | 0.4883 |
| HMR_4781 | Arginine and proline metabolism | 4.0214 | 11.5968 | 3.8805 | 10.7905 | 0.1408 | 0.0298 | 0.4881 |
| HMR_4783 | Arginine and proline metabolism | 4.0335 | 11.4924 | 3.8533 | 10.5413 | 0.1801 | 0.0384 | 0.4847 |
| HMR_4784 | Arginine and proline metabolism | 6.0303 | 20.5862 | 5.8025 | 19.0674 | 0.2278 | 0.0362 | 0.4856 |
| HMR_4785 | Arginine and proline metabolism | 6.0417 | 20.6198 | 5.8098 | 19.0883 | 0.2319 | 0.0368 | 0.4853 |
| HMR_4786 | Arginine and proline metabolism | 6.0363 | 20.6057 | 5.8064 | 19.0801 | 0.2299 | 0.0365 | 0.4854 |
| HMR_4787 | Arginine and proline metabolism | 12.0718 | 17.0288 | 11.6118 | 15.8198 | 0.46 | 0.0803 | 0.468 |
| HMR_5384 | Arginine and proline metabolism | 1.3272 | 0.3152 | 1.2947 | 0.2985 | 0.0325 | 0.0415 | 0.4835 |
| HMR_8098 | Arginine and proline metabolism | 6.0363 | 20.6057 | 5.8064 | 19.0801 | 0.2299 | 0.0365 | 0.4854 |
| HMR_8427 | Arginine and proline metabolism | -39.8074 | 474.0112 | -47.8181 | 683.0071 | 8.0107 | 0.2355 | 0.4069 |
| HMR_8428 | Arginine and proline metabolism | 39.8074 | 474.0163 | 47.8181 | 683.0122 | 8.0107 | -0.2355 | 0.5931 |
| HMR_8430 | Arginine and proline metabolism | 39.8097 | 474.0115 | 47.8204 | 683.0073 | 8.0107 | -0.2355 | 0.5931 |
| HMR_8606 | Arginine and proline metabolism | 4.4544 | 6.8763 | 4.5597 | 7.2065 | 0.1053 | -0.0281 | 0.5112 |
| HMR_8609 | Arginine and proline metabolism | 13.3527 | 169.6251 | 14.9771 | 298.8673 | 1.6244 | -0.0751 | 0.5299 |
| HMR_8610 | Arginine and proline metabolism | 15.0948 | 213.2439 | 16.4989 | 358.8723 | 1.404 | -0.0587 | 0.5234 |
| HMR_8611 | Arginine and proline metabolism | 92.2989 | 4647.0846 | 180.6581 | 16708.7227 | 88.3592 | -0.6046 | 0.7273 |
| HMR_4285 | Glutamate and glutamine metabolism | 6.2599 | 26.0875 | 6.7126 | 29.6973 | 0.4527 | -0.0606 | 0.5242 |
| HMR_0457 | Glycine, serine and threonine metabolism | 7.0423 | 28.671 | 6.613 | 25.2436 | 0.4293 | 0.0585 | 0.4767 |
| HMR_0460 | Glycine, serine and threonine metabolism | 3.3479 | 3.0099 | 3.2863 | 2.8993 | 0.0616 | 0.0253 | 0.4899 |
| HMR_3752 | Glycine, serine and threonine metabolism | 201.4253 | 82353.8474 | 480.6641 | 74005.0451 | 279.2388 | -0.7062 | 0.76 |
| HMR_3770 | Glycine, serine and threonine metabolism | 114.5294 | 227879.6245 | 77.487 | 232678.9235 | 37.0423 | 0.0546 | 0.4782 |
| HMR_3784 | Glycine, serine and threonine metabolism | -112.1537 | 227873.5781 | -78.9788 | 232681.1613 | 33.1749 | -0.0489 | 0.5195 |
| HMR_3839 | Glycine, serine and threonine metabolism | 8.4076 | 23.4862 | 8.2198 | 22.2779 | 0.1879 | 0.0278 | 0.4889 |
| HMR_3841 | Glycine, serine and threonine metabolism | 8.4049 | 23.4843 | 8.2169 | 22.2761 | 0.188 | 0.0278 | 0.4889 |
| HMR_3843 | Glycine, serine and threonine metabolism | 8.4025 | 23.4883 | 8.2145 | 22.2802 | 0.1881 | 0.0278 | 0.4889 |
| HMR_3845 | Glycine, serine and threonine metabolism | -403.3471 | 2614.9089 | -462.3558 | 2485.8236 | 59.0086 | 0.8262 | 0.2043 |

| | | | | | | | | |
|---|---|---|---|---|---|---|---|---|
| HMR_3847 | Glycine, serine and threonine metabolism | -55.1749 | 201.9002 | -63.383 | 295.4768 | 8.2081 | 0.368 | 0.3564 |
| HMR_3849 | Glycine, serine and threonine metabolism | 114.2256 | 1561.1423 | 124.1594 | 1928.7224 | 9.9339 | -0.1682 | 0.5668 |
| HMR_3852 | Glycine, serine and threonine metabolism | 55.1755 | 201.8719 | 63.3834 | 295.448 | 8.2079 | -0.3681 | 0.6436 |
| HMR_3856 | Glycine, serine and threonine metabolism | 10.1577 | 83.1787 | 11.8099 | 114.2413 | 1.6523 | -0.1176 | 0.5468 |
| HMR_3860 | Glycine, serine and threonine metabolism | 55.1747 | 201.8904 | 63.3826 | 295.4669 | 8.208 | -0.368 | 0.6436 |
| HMR_3883 | Glycine, serine and threonine metabolism | 8.9474 | 86.2684 | 8.4876 | 75.6229 | 0.4599 | 0.0361 | 0.4856 |
| HMR_3901 | Glycine, serine and threonine metabolism | 17.6587 | 286.0863 | 18.6702 | 320.4388 | 1.0114 | -0.0411 | 0.5164 |
| HMR_3939 | Glycine, serine and threonine metabolism | -3.7457 | 220493.1892 | -3.3908 | 220496.5824 | 0.3549 | -0.0005 | 0.5002 |
| HMR_3974 | Glycine, serine and threonine metabolism | 8.9782 | 37.6164 | 8.5055 | 33.7829 | 0.4727 | 0.0559 | 0.4777 |
| HMR_4198 | Glycine, serine and threonine metabolism | 10.8123 | 26.3897 | 10.0433 | 22.96 | 0.7691 | 0.1095 | 0.4564 |
| HMR_4200 | Glycine, serine and threonine metabolism | 8.9788 | 37.6164 | 8.5061 | 33.7829 | 0.4727 | 0.0559 | 0.4777 |
| HMR_4284 | Glycine, serine and threonine metabolism | 9.7934 | 85.1598 | 9.2845 | 78.083 | 0.5089 | 0.0398 | 0.4841 |
| HMR_4348 | Glycine, serine and threonine metabolism | 11.2429 | 148.3253 | 11.0512 | 137.9341 | 0.1917 | 0.0113 | 0.4955 |
| HMR_4582 | Glycine, serine and threonine metabolism | -921.6884 | 986.1031 | -910.1472 | 1486.7446 | 11.5412 | -0.2321 | 0.5918 |
| HMR_4584 | Glycine, serine and threonine metabolism | -39.8119 | 474.0166 | -47.8227 | 683.0123 | 8.0108 | 0.2355 | 0.4069 |
| HMR_4696 | Glycine, serine and threonine metabolism | 2.9508 | 2.3013 | 3.0222 | 2.4526 | 0.0714 | -0.0327 | 0.5131 |
| HMR_4697 | Glycine, serine and threonine metabolism | 1.4738 | 1.551 | 1.5095 | 1.6377 | 0.0356 | -0.02 | 0.508 |
| HMR_4698 | Glycine, serine and threonine metabolism | 1.4764 | 1.5549 | 1.5121 | 1.6418 | 0.0357 | -0.02 | 0.508 |
| HMR_4699 | Glycine, serine and threonine metabolism | 2.9453 | 2.3007 | 3.0167 | 2.4524 | 0.0714 | -0.0328 | 0.5131 |
| HMR_4700 | Glycine, serine and threonine metabolism | 93.3976 | 1449.8572 | 101.7879 | 1845.1335 | 8.3903 | -0.1462 | 0.5581 |
| HMR_4742 | Glycine, serine and threonine metabolism | -7.4898 | 220499.8376 | -6.7806 | 220502.1745 | 0.7092 | -0.0011 | 0.5004 |
| HMR_4792 | Glycine, serine and threonine metabolism | -29.6257 | 402.0206 | -25.9246 | 310.8318 | 3.701 | -0.1386 | 0.5551 |
| HMR_5392 | Glycine, serine and threonine metabolism | 0.5743 | 0.1577 | 0.5488 | 0.1442 | 0.0256 | 0.0465 | 0.4815 |
| HMR_5393 | Glycine, serine and threonine metabolism | 0.5813 | 0.1578 | 0.5559 | 0.1444 | 0.0253 | 0.0461 | 0.4816 |
| HMR_6409 | Glycine, serine and threonine metabolism | -781.347 | 24515.5263 | -778.8089 | 26473.4525 | 2.5381 | -0.0112 | 0.5045 |
| HMR_7702 | Glycine, serine and threonine metabolism | -27.3003 | 185.982 | -28.3267 | 199.0992 | 1.0264 | 0.0523 | 0.4791 |
| HMR_8433 | Glycine, serine and threonine metabolism | -160.1704 | 9980.7612 | -144.9535 | 8263.9911 | 15.2169 | -0.1127 | 0.5448 |
| HMR_8434 | Glycine, serine and threonine metabolism | 160.171 | 9980.7612 | 144.9528 | 8263.991 | 15.2182 | 0.1127 | 0.4551 |
| HMR_8435 | Glycine, serine and threonine metabolism | 283.0288 | 22132.9671 | 239.8145 | 16042.2507 | 43.2143 | 0.2212 | 0.4125 |
| HMR_8436 | Glycine, serine and threonine metabolism | 283.029 | 22132.9672 | 239.8147 | 16042.251 | 43.2143 | 0.2212 | 0.4125 |
| HMR_8437 | Glycine, serine and threonine metabolism | -283.0289 | 22132.9682 | -239.8146 | 16042.252 | 43.2143 | -0.2212 | 0.5875 |
| HMR_4428 | Histidine metabolism | -4.3158 | 4.6244 | -4.0753 | 4.1275 | 0.2405 | -0.0813 | 0.5324 |
| HMR_4429 | Histidine metabolism | 4.3158 | 4.6112 | 4.0754 | 4.1143 | 0.2404 | 0.0814 | 0.4676 |
| HMR_4430 | Histidine metabolism | 4.3162 | 4.6079 | 4.0759 | 4.111 | 0.2403 | 0.0814 | 0.4676 |
| HMR_4431 | Histidine metabolism | 4.3171 | 4.6079 | 4.0768 | 4.111 | 0.2402 | 0.0814 | 0.4676 |
| HMR_4437 | Histidine metabolism | -674.8828 | 2979.9278 | -825.1714 | 2964.8204 | 150.2886 | 1.9492 | 0.0256 |
| HMR_4658 | Histidine metabolism | -674.8828 | 2979.9177 | -825.1716 | 2964.8095 | 150.2888 | 1.9492 | 0.0256 |
| HMR_4660 | Histidine metabolism | 674.8825 | 2979.9148 | 825.1712 | 2964.8075 | 150.2887 | -1.9492 | 0.9744 |
| HMR_4712 | Histidine metabolism | 674.8825 | 2979.9182 | 825.1711 | 2964.8111 | 150.2886 | -1.9492 | 0.9744 |
| HMR_5338 | Histidine metabolism | 8.4147 | 33.0536 | 8.2392 | 31.7053 | 0.1755 | 0.0218 | 0.4913 |
| HMR_5339 | Histidine metabolism | 8.4144 | 33.0535 | 8.2389 | 31.7053 | 0.1755 | 0.0218 | 0.4913 |
| HMR_4239 | Lysine metabolism | 8.1114 | 38.5245 | 8.0698 | 35.5974 | 0.0416 | 0.0048 | 0.4981 |
| HMR_4288 | Lysine metabolism | 15.9859 | 41.1623 | 16.0195 | 32.1106 | 0.0336 | -0.0039 | 0.5016 |
| HMR_4597 | Lysine metabolism | -15.9844 | 41.1459 | -16.0176 | 32.1063 | 0.0332 | 0.0039 | 0.4985 |

| | | | | | | | | |
|---|---|---|---|---|---|---|---|---|
| HMR_4599 | Lysine metabolism | 9.0582 | 31.5248 | 9.1163 | 30.7193 | 0.0581 | -0.0074 | 0.5029 |
| HMR_4667 | Lysine metabolism | 121.765 | 226720.1142 | 82.0743 | 232324.4382 | 39.6907 | 0.0586 | 0.4766 |
| HMR_4668 | Lysine metabolism | -137.7517 | 226721.0598 | -98.0936 | 232323.6574 | 39.6581 | -0.0585 | 0.5233 |
| HMR_4737 | Lysine metabolism | 7.6464 | 23.8493 | 7.6357 | 23.1951 | 0.0106 | 0.0015 | 0.4994 |
| HMR_4739 | Lysine metabolism | 8.3361 | 38.9002 | 8.3833 | 35.2278 | 0.0473 | -0.0055 | 0.5022 |
| HMR_6415 | Lysine metabolism | 9.0573 | 31.527 | 9.1143 | 30.7219 | 0.0569 | -0.0072 | 0.5029 |
| HMR_3164 | Tryptophan metabolism | 34.8068 | 47.5012 | 36.7571 | 39.746 | 1.9503 | -0.2088 | 0.5827 |
| HMR_3166 | Tryptophan metabolism | 34.8025 | 47.5084 | 36.7524 | 39.7531 | 1.9499 | -0.2087 | 0.5827 |
| HMR_3885 | Tryptophan metabolism | 28.9456 | 252.1581 | 31.1928 | 246.0714 | 2.2472 | -0.1007 | 0.5401 |
| HMR_4250 | Tryptophan metabolism | 1.5881 | 0.9891 | 1.606 | 1.0024 | 0.0179 | -0.0127 | 0.5051 |
| HMR_8566 | Tryptophan metabolism | 20.8985 | 114.1834 | 25.7389 | 170.457 | 4.8404 | -0.2869 | 0.6129 |
| HMR_3935 | Tyrosine metabolism | 1.6675 | 220507.8773 | 1.5821 | 220508.3022 | 0.0854 | 0.0001 | 0.4999 |
| HMR_4683 | Tyrosine metabolism | 408.3836 | 57852.9897 | 409.1603 | 57953.2154 | 0.7766 | -0.0023 | 0.5009 |
| HMR_4685 | Tyrosine metabolism | 470.6589 | 62232.8031 | 479.596 | 62536.8015 | 8.9371 | -0.0253 | 0.5101 |
| HMR_4686 | Tyrosine metabolism | 408.3838 | 57852.9896 | 409.1604 | 57953.2153 | 0.7766 | -0.0023 | 0.5009 |
| HMR_4687 | Tyrosine metabolism | 470.6589 | 62232.8029 | 479.596 | 62536.8013 | 8.9371 | -0.0253 | 0.5101 |
| HMR_7701 | Tyrosine metabolism | 17.4751 | 285.6216 | 18.3635 | 314.8409 | 0.8884 | -0.0363 | 0.5145 |
| HMR_8530 | Tyrosine metabolism | 3.3416 | 220514.0576 | 3.1712 | 220513.9435 | 0.1704 | 0.0003 | 0.4999 |
| HMR_8534 | Tyrosine metabolism | 1.2259 | 0.5227 | 1.2109 | 0.5102 | 0.015 | 0.0148 | 0.4941 |
| HMR_8537 | Tyrosine metabolism | 4.8316 | 20.0793 | 4.7823 | 19.761 | 0.0493 | 0.0078 | 0.4969 |
| HMR_8541 | Tyrosine metabolism | 3.3394 | 10.3258 | 3.2275 | 9.7073 | 0.1119 | 0.025 | 0.49 |
| HMR_3206 | Valine, leucine, and isoleucine metabolism | 10.051 | 18.6784 | 2.121 | 0.6926 | 7.93 | 1.8017 | 0.0358 |
| HMR_3208 | Valine, leucine, and isoleucine metabolism | 16.7376 | 85.0567 | 18.1254 | 99.3413 | 1.3878 | -0.1022 | 0.5407 |
| HMR_3211 | Valine, leucine, and isoleucine metabolism | 9.8713 | 62.2177 | 9.0072 | 53.7899 | 0.8641 | 0.0802 | 0.468 |
| HMR_3213 | Valine, leucine, and isoleucine metabolism | 12.2728 | 74.8994 | 13.5459 | 88.6097 | 1.2731 | -0.0996 | 0.5397 |
| HMR_3215 | Valine, leucine, and isoleucine metabolism | 337.834 | 2839.3221 | 19.3698 | 90.0565 | 318.4642 | 5.884 | 0.00E+00 |
| HMR_3744 | Valine, leucine, and isoleucine metabolism | 12.4183 | 71.7493 | 2.6795 | 2.5603 | 9.7388 | 1.1298 | 0.1293 |
| HMR_3747 | Valine, leucine, and isoleucine metabolism | 317.6482 | 2872.6209 | 3.1127 | 2.9041 | 314.5355 | 5.8656 | 0.00E+00 |
| HMR_3757 | Valine, leucine, and isoleucine metabolism | 330.9161 | 2827.6229 | 7.0144 | 1.7376 | 323.9017 | 6.0893 | 0.00E+00 |
| HMR_3761 | Valine, leucine, and isoleucine metabolism | 138.8494 | 9135.923 | 1.4655 | 1.5517 | 137.3839 | 1.4372 | 0.0753 |
| HMR_3765 | Valine, leucine, and isoleucine metabolism | -5.3653 | 5.7459 | -5.1181 | 5.0215 | 0.2473 | -0.0754 | 0.53 |
| HMR_3778 | Valine, leucine, and isoleucine metabolism | -10.0704 | 18.6645 | -2.1307 | 0.659 | 7.9397 | -1.8062 | 0.9646 |
| HMR_3795 | Valine, leucine, and isoleucine metabolism | 5.3533 | 25.2377 | 1.2009 | 1.1415 | 4.1523 | 0.8085 | 0.2094 |
| HMR_4797 | Valine, leucine, and isoleucine metabolism | 25.4258 | 360.959 | 1.4864 | 1.4414 | 23.9394 | 1.2575 | 0.1043 |
| HMR_6417 | Valine, leucine, and isoleucine metabolism | 156.8625 | 8788.5715 | 3.1272 | 3.3036 | 153.7353 | 1.6396 | 0.0505 |
| HMR_6419 | Valine, leucine, and isoleucine metabolism | 5.2813 | 11.2871 | 1.1349 | 0.4976 | 4.1464 | 1.2078 | 0.1136 |
| HMR_6420 | Valine, leucine, and isoleucine metabolism | 5.2812 | 11.2867 | 1.1372 | 0.4989 | 4.1441 | 1.2071 | 0.1137 |
| HMR_6421 | Valine, leucine, and isoleucine metabolism | 2.8108 | 3.2504 | 2.6864 | 2.9418 | 0.1244 | 0.05 | 0.4801 |
| HMR_6422 | Valine, leucine, and isoleucine metabolism | 2.8098 | 3.2503 | 2.6844 | 2.942 | 0.1253 | 0.0504 | 0.4799 |
| HMR_8787 | Valine, leucine, and isoleucine metabolism | 4.4616 | 18.3137 | 4.5762 | 19.4308 | 0.1146 | -0.0186 | 0.5074 |
| HMR_3940 | Phenylalanine, tyrosine and tryptophan biosynthesis | 6.6722 | 57.0609 | 6.8934 | 62.0563 | 0.2212 | -0.0203 | 0.5081 |
| HMR_4214 | Phenylalanine, tyrosine and tryptophan biosynthesis | 2.7545 | 1.2436 | 2.755 | 1.2398 | 0.0006 | -0.0004 | 0.5001 |
| HMR_4216 | Phenylalanine, tyrosine and tryptophan biosynthesis | 2.7559 | 1.2352 | 2.7563 | 1.2317 | 0.0004 | -0.0003 | 0.5001 |
| HMR_4220 | Phenylalanine, tyrosine and tryptophan biosynthesis | -2.7575 | 1.2282 | -2.7578 | 1.2249 | 0.0003 | 0.0002 | 0.4999 |

| | | | | | | | | |
|---|---|---|---|---|---|---|---|---|
| HMR_4225 | Phenylalanine, tyrosine and tryptophan biosynthesis | 2.7644 | 1.2103 | 2.7641 | 1.2077 | 0.0003 | 0.0002 | 0.4999 |
| HMR_4231 | Phenylalanine, tyrosine and tryptophan biosynthesis | 1.1843 | 0.4122 | 1.1653 | 0.3996 | 0.019 | 0.021 | 0.4916 |
| HMR_4233 | Phenylalanine, tyrosine and tryptophan biosynthesis | 1.1875 | 0.4085 | 1.1684 | 0.3959 | 0.0191 | 0.0213 | 0.4915 |
| HMR_4235 | Phenylalanine, tyrosine and tryptophan biosynthesis | 1.1891 | 0.4092 | 1.1699 | 0.3966 | 0.0192 | 0.0214 | 0.4915 |
| HMR_4242 | Phenylalanine, tyrosine and tryptophan biosynthesis | -125.7124 | 227853.1601 | -88.5773 | 232645.8992 | 37.1351 | -0.0547 | 0.5218 |
| HMR_4243 | Phenylalanine, tyrosine and tryptophan biosynthesis | 108.5462 | 227854.7766 | 71.3993 | 232648.2035 | 37.1469 | 0.0547 | 0.4782 |
| HMR_4244 | Phenylalanine, tyrosine and tryptophan biosynthesis | 499.998 | 55128.0374 | 499.9982 | 55128.0394 | 0.0002 | 0.00E+00 | 0.5 |
| HMR_4245 | Phenylalanine, tyrosine and tryptophan biosynthesis | 500.0001 | 55128.0341 | 500.0001 | 55128.036 | 0.00E+00 | 0.00E+00 | 0.5 |
| HMR_4246 | Phenylalanine, tyrosine and tryptophan biosynthesis | -500.0019 | 55128.0374 | -500.0017 | 55128.0394 | 0.0002 | 0.00E+00 | 0.5 |
| HMR_6728 | Phenylalanine, tyrosine and tryptophan biosynthesis | 4.8121 | 22.0087 | 4.7553 | 21.5332 | 0.0568 | 0.0086 | 0.4966 |
| HMR_6729 | Phenylalanine, tyrosine and tryptophan biosynthesis | 13.582 | 123.3817 | 14.4372 | 134.3994 | 0.8552 | -0.0533 | 0.5212 |
| HMR_6731 | Phenylalanine, tyrosine and tryptophan biosynthesis | 29.9038 | 175.5344 | 30.4287 | 182.8024 | 0.5249 | -0.0277 | 0.5111 |
| HMR_6743 | Phenylalanine, tyrosine and tryptophan biosynthesis | 3.1942 | 3.0843 | 3.0231 | 2.7636 | 0.171 | 0.0707 | 0.4718 |
| HMR_6768 | Phenylalanine, tyrosine and tryptophan biosynthesis | 1.5574 | 1.9218 | 1.5474 | 1.8898 | 0.01 | 0.0051 | 0.498 |
| HMR_6770 | Phenylalanine, tyrosine and tryptophan biosynthesis | 1.2584 | 1.3456 | 1.2483 | 1.3211 | 0.0101 | 0.0062 | 0.4975 |
| HMR_6772 | Phenylalanine, tyrosine and tryptophan biosynthesis | 4.3544 | 3.2933 | 4.2913 | 3.2068 | 0.0632 | 0.0248 | 0.4901 |
| HMR_6774 | Phenylalanine, tyrosine and tryptophan biosynthesis | 4.3503 | 3.2991 | 4.2871 | 3.2125 | 0.0632 | 0.0248 | 0.4901 |
| HMR_6776 | Phenylalanine, tyrosine and tryptophan biosynthesis | 4.3477 | 3.3063 | 4.2844 | 3.2197 | 0.0633 | 0.0248 | 0.4901 |
| HMR_6778 | Phenylalanine, tyrosine and tryptophan biosynthesis | 4.345 | 3.3148 | 4.2816 | 3.2282 | 0.0634 | 0.0248 | 0.4901 |
| HMR_6826 | Phenylalanine, tyrosine and tryptophan biosynthesis | 1.2327 | 0.5212 | 1.2178 | 0.5087 | 0.0149 | 0.0147 | 0.4941 |
| HMR_6829 | Phenylalanine, tyrosine and tryptophan biosynthesis | 1.1804 | 0.6538 | 1.1655 | 0.6375 | 0.0149 | 0.0132 | 0.4948 |
| HMR_6831 | Phenylalanine, tyrosine and tryptophan biosynthesis | 1.1832 | 0.6539 | 1.1683 | 0.6375 | 0.0149 | 0.0131 | 0.4948 |
| HMR_6834 | Phenylalanine, tyrosine and tryptophan biosynthesis | -1.231 | 0.521 | -1.2161 | 0.5085 | 0.015 | -0.0147 | 0.5059 |
| HMR_6838 | Phenylalanine, tyrosine and tryptophan biosynthesis | 1.136 | 1.0441 | 1.1217 | 1.0182 | 0.0143 | 0.01 | 0.496 |
| HMR_6839 | Phenylalanine, tyrosine and tryptophan biosynthesis | 2.3648 | 1.4306 | 2.3356 | 1.3956 | 0.0292 | 0.0174 | 0.4931 |
| HMR_6844 | Phenylalanine, tyrosine and tryptophan biosynthesis | 1.1802 | 0.6529 | 1.1652 | 0.6365 | 0.0149 | 0.0132 | 0.4947 |
| HMR_6845 | Phenylalanine, tyrosine and tryptophan biosynthesis | 1.1822 | 0.6529 | 1.1672 | 0.6365 | 0.0149 | 0.0132 | 0.4948 |
| HMR_6850 | Phenylalanine, tyrosine and tryptophan biosynthesis | 1.1802 | 0.6529 | 1.1652 | 0.6365 | 0.0149 | 0.0132 | 0.4947 |
| HMR_6851 | Phenylalanine, tyrosine and tryptophan biosynthesis | 1.1822 | 0.6529 | 1.1672 | 0.6365 | 0.0149 | 0.0132 | 0.4948 |
| HMR_8539 | Phenylalanine, tyrosine and tryptophan biosynthesis | 6.5491 | 34.5938 | 6.7608 | 36.5149 | 0.2117 | -0.0251 | 0.51 |
| HMR_3875 | Cysteine and methionine metabolism | 7.0453 | 5.3125 | 9.544 | 6.8268 | 2.4987 | -0.7172 | 0.7634 |
| HMR_3877 | Cysteine and methionine metabolism | 5.7349 | 4.9841 | 8.2368 | 6.5875 | 2.5019 | -0.7355 | 0.769 |
| HMR_3879 | Cysteine and methionine metabolism | 1.3171 | 0.8083 | 3.7382 | 1.8652 | 2.4211 | -1.4807 | 0.9307 |
| HMR_3881 | Cysteine and methionine metabolism | 1.3197 | 0.8084 | 3.7371 | 1.8746 | 2.4174 | -1.4758 | 0.93 |
| HMR_3917 | Cysteine and methionine metabolism | 1.4789 | 2.0813 | 1.4808 | 2.371 | 0.0019 | -0.0009 | 0.5004 |
| HMR_3995 | Cysteine and methionine metabolism | -213.1611 | 170455.8913 | -246.0751 | 169393.5824 | 32.9141 | 0.0565 | 0.4775 |
| HMR_3996 | Cysteine and methionine metabolism | 453.5471 | 166818.2896 | 442.8945 | 166809.9302 | 10.6525 | 0.0184 | 0.4926 |
| HMR_4189 | Cysteine and methionine metabolism | 5.9205 | 39.3082 | 5.9955 | 39.8761 | 0.075 | -0.0084 | 0.5034 |
| HMR_4302 | Cysteine and methionine metabolism | 8.9475 | 86.2673 | 8.4876 | 75.6229 | 0.4599 | 0.0361 | 0.4856 |
| HMR_4323 | Cysteine and methionine metabolism | 302.374 | 4600.7833 | 374.7793 | 3850.407 | 72.4053 | -0.7876 | 0.7845 |
| HMR_5381 | Cysteine and methionine metabolism | 1.6293 | 3.3385 | 1.6281 | 3.3588 | 0.0012 | 0.0005 | 0.4998 |
| HMR_5385 | Cysteine and methionine metabolism | 1.3248 | 0.3186 | 1.2851 | 0.3014 | 0.0397 | 0.0504 | 0.4799 |
| HMR_5386 | Cysteine and methionine metabolism | 1.3219 | 0.3233 | 1.2749 | 0.3058 | 0.047 | 0.0592 | 0.4764 |
| HMR_5387 | Cysteine and methionine metabolism | 1.3181 | 0.3297 | 1.2637 | 0.3117 | 0.0543 | 0.0678 | 0.473 |

| | | | | | | | | |
|---|---|---|---|---|---|---|---|---|
| HMR_5389 | Cysteine and methionine metabolism | 1.3165 | 0.3377 | 1.2548 | 0.3195 | 0.0616 | 0.076 | 0.4697 |
| HMR_5390 | Cysteine and methionine metabolism | 5.5714 | 17.7265 | 5.4938 | 17.1349 | 0.0775 | 0.0131 | 0.4948 |
| HMR_5391 | Cysteine and methionine metabolism | 6.8876 | 17.8163 | 6.741 | 17.2164 | 0.1466 | 0.0248 | 0.4901 |
| HMR_3868 | Glutathione metabolism | 352.9185 | 70390.5673 | 352.7974 | 70552.9972 | 0.1212 | 0.0003 | 0.4999 |
| HMR_3869 | Glutathione metabolism | -133.6016 | 226752.2507 | -93.7198 | 232360.8112 | 39.8818 | -0.0589 | 0.5235 |
| HMR_3889 | Glutathione metabolism | 2.3957 | 1.6358 | 2.2878 | 1.4919 | 0.1079 | 0.061 | 0.4757 |
| HMR_3933 | Glutathione metabolism | -1.6745 | 220507.874 | -1.5894 | 220508.2988 | 0.0851 | -0.0001 | 0.5001 |
| HMR_3937 | Glutathione metabolism | 5.0234 | 18.5457 | 4.7677 | 16.9274 | 0.2557 | 0.0429 | 0.4829 |
| HMR_4116 | Glutathione metabolism | 15.557 | 226.4505 | 16.3391 | 248.9359 | 0.7821 | -0.0359 | 0.5143 |
| HMR_4121 | Glutathione metabolism | 7.5571 | 76.4973 | 7.1751 | 64.8149 | 0.382 | 0.0321 | 0.4872 |
| HMR_4175 | Glutathione metabolism | 1.4276 | 0.9268 | 1.3637 | 0.8455 | 0.0639 | 0.048 | 0.4809 |
| HMR_4176 | Glutathione metabolism | 1.0406 | 0.8296 | 0.993 | 0.7552 | 0.0476 | 0.0378 | 0.4849 |
| HMR_4324 | Glutathione metabolism | 4.7642 | 3.2427 | 4.5414 | 2.9639 | 0.2229 | 0.0895 | 0.4644 |
| HMR_4326 | Glutathione metabolism | 3.4141 | 2.0847 | 3.2525 | 1.9086 | 0.1616 | 0.0808 | 0.4678 |
| HMR_4328 | Glutathione metabolism | 1.4304 | 0.9273 | 1.3666 | 0.8459 | 0.0639 | 0.048 | 0.4809 |
| HMR_4329 | Glutathione metabolism | 2.0849 | 1.1407 | 1.9906 | 1.0395 | 0.0943 | 0.0638 | 0.4745 |
| HMR_4708 | Glutathione metabolism | 1.3533 | 1.3471 | 1.2921 | 1.2278 | 0.0612 | 0.0381 | 0.4848 |
| HMR_8769 | Glutathione metabolism | 1.0439 | 0.4627 | 0.9973 | 0.4224 | 0.0466 | 0.0495 | 0.4803 |
| HMR_8770 | Glutathione metabolism | 1.0452 | 0.463 | 0.9987 | 0.4227 | 0.0465 | 0.0494 | 0.4803 |
| HMR_4078 | beta-Alanine metabolism | 1.3347 | 0.3137 | 1.3095 | 0.2974 | 0.0252 | 0.0322 | 0.4871 |
| HMR_4079 | beta-Alanine metabolism | 0.6724 | 0.2807 | 0.6598 | 0.2692 | 0.0126 | 0.017 | 0.4932 |
| HMR_4330 | beta-Alanine metabolism | -15.0198 | 367.4596 | -9.6886 | 262.2135 | 5.3313 | -0.2125 | 0.5841 |
| HMR_4347 | beta-Alanine metabolism | 1.427 | 0.4925 | 1.3956 | 0.4753 | 0.0314 | 0.032 | 0.4872 |
| HMR_4425 | beta-Alanine metabolism | 2.0134 | 1.8973 | 2.0373 | 1.9426 | 0.0239 | -0.0122 | 0.5049 |
| HMR_4427 | beta-Alanine metabolism | 2.015 | 1.8974 | 2.0387 | 1.9426 | 0.0237 | -0.0121 | 0.5048 |
| HMR_4688 | beta-Alanine metabolism | -529.3412 | 62232.8029 | -520.404 | 62536.8039 | 8.9372 | -0.0253 | 0.5101 |
| HMR_4689 | beta-Alanine metabolism | 4.4547 | 6.8791 | 4.5601 | 7.2093 | 0.1053 | -0.0281 | 0.5112 |
| HMR_4735 | beta-Alanine metabolism | 19.7074 | 75.0325 | 17.9912 | 70.1621 | 1.7162 | 0.1424 | 0.4434 |
| HMR_7993 | beta-Alanine metabolism | 0.6719 | 0.2805 | 0.6594 | 0.269 | 0.0126 | 0.017 | 0.4932 |
| HMR_3748 | Valine, leucine and isoleucine degradation | 173.2033 | 9933.0953 | 2.6856 | 3.47 | 170.5177 | 1.7106 | 0.0436 |
| HMR_3751 | Valine, leucine and isoleucine degradation | 532.3481 | 79919.026 | 487.6683 | 74003.2302 | 44.6798 | 0.1139 | 0.4547 |
| HMR_3753 | Valine, leucine and isoleucine degradation | 330.9193 | 2827.6242 | 7.0063 | 1.7275 | 323.913 | 6.0895 | 0.00E+00 |
| HMR_3755 | Valine, leucine and isoleucine degradation | 330.9156 | 2827.6235 | 7.0081 | 1.7323 | 323.9075 | 6.0894 | 0.00E+00 |
| HMR_3759 | Valine, leucine and isoleucine degradation | 149.7057 | 9003.9901 | 1.473 | 1.5585 | 148.2327 | 1.562 | 0.0591 |
| HMR_3763 | Valine, leucine and isoleucine degradation | 186.7176 | 8960.7653 | 4.3624 | 2.4122 | 182.3552 | 1.9261 | 0.027 |
| HMR_3767 | Valine, leucine and isoleucine degradation | 2.5541 | 4.4007 | 2.4319 | 3.9176 | 0.1222 | 0.0424 | 0.4831 |
| HMR_3769 | Valine, leucine and isoleucine degradation | -119.8863 | 227879.4042 | -82.5958 | 232678.8381 | 37.2905 | -0.0549 | 0.5219 |
| HMR_3773 | Valine, leucine and isoleucine degradation | 5.3508 | 5.7514 | 5.1025 | 5.0293 | 0.2483 | 0.0756 | 0.4699 |
| HMR_3775 | Valine, leucine and isoleucine degradation | 5.3462 | 5.7592 | 5.0974 | 5.0372 | 0.2487 | 0.0757 | 0.4698 |
| HMR_3780 | Valine, leucine and isoleucine degradation | 4.789 | 14.9006 | 1.0016 | 0.598 | 3.7873 | 0.962 | 0.168 |
| HMR_3783 | Valine, leucine and isoleucine degradation | 122.2185 | 227872.306 | 81.1128 | 232681.2067 | 41.1056 | 0.0606 | 0.4759 |
| HMR_3785 | Valine, leucine and isoleucine degradation | -10.0603 | 18.6652 | -2.1299 | 0.6756 | 7.9304 | -1.8032 | 0.9643 |
| HMR_3790 | Valine, leucine and isoleucine degradation | 11.5848 | 113.7258 | 1.4007 | 1.3351 | 10.1841 | 0.9494 | 0.1712 |
| HMR_3796 | Valine, leucine and isoleucine degradation | -0.7221 | 220511.2997 | -0.7411 | 220511.2557 | 0.0191 | 0.00E+00 | 0.5 |

| | | | | | | | | |
|---|---|---|---|---|---|---|---|---|
| HMR_3823 | Valine, leucine and isoleucine degradation | 10.0557 | 18.671 | 2.1256 | 0.6837 | 7.9301 | 1.8025 | 0.0357 |
| HMR_6416 | Valine, leucine and isoleucine degradation | 156.8624 | 8788.5725 | 3.1215 | 3.3016 | 153.741 | 1.6396 | 0.0505 |
| HMR_3986 | Amino sugar and nucleotide sugar metabolism | 0.8646 | 0.2059 | 0.8789 | 0.2131 | 0.0143 | -0.0222 | 0.5088 |
| HMR_3988 | Amino sugar and nucleotide sugar metabolism | 0.853 | 0.2064 | 0.8676 | 0.2136 | 0.0146 | -0.0225 | 0.509 |
| HMR_4158 | Amino sugar and nucleotide sugar metabolism | 5.1858 | 3.2092 | 5.286 | 3.3309 | 0.1001 | -0.0392 | 0.5156 |
| HMR_4299 | Amino sugar and nucleotide sugar metabolism | 62.9198 | 754.8803 | 58.0047 | 648.6672 | 4.9151 | 0.1312 | 0.4478 |
| HMR_4300 | Amino sugar and nucleotide sugar metabolism | 2.8523 | 8.6139 | 2.7231 | 8.0282 | 0.1292 | 0.0317 | 0.4874 |
| HMR_4494 | Amino sugar and nucleotide sugar metabolism | 20.501 | 341.4852 | 18.925 | 291.7041 | 1.576 | 0.0626 | 0.475 |
| HMR_4524 | Amino sugar and nucleotide sugar metabolism | 3.6714 | 2.4608 | 3.725 | 2.5307 | 0.0536 | -0.024 | 0.5096 |
| HMR_4526 | Amino sugar and nucleotide sugar metabolism | 1.9819 | 1.7348 | 2.0059 | 1.7782 | 0.024 | -0.0128 | 0.5051 |
| HMR_4527 | Amino sugar and nucleotide sugar metabolism | 1.9796 | 1.7344 | 2.0036 | 1.7778 | 0.024 | -0.0128 | 0.5051 |
| HMR_4531 | Amino sugar and nucleotide sugar metabolism | 502.1546 | 35483.9956 | 501.9543 | 35496.2343 | 0.2004 | 0.0008 | 0.4997 |
| HMR_4532 | Amino sugar and nucleotide sugar metabolism | -502.1521 | 35483.9932 | -501.9517 | 35496.2319 | 0.2004 | -0.0008 | 0.5003 |
| HMR_4533 | Amino sugar and nucleotide sugar metabolism | 219.5434 | 31069.5885 | 219.7083 | 31095.784 | 0.1649 | -0.0007 | 0.5003 |
| HMR_4534 | Amino sugar and nucleotide sugar metabolism | 282.6189 | 39274.1514 | 282.2536 | 39243.5905 | 0.3652 | 0.0013 | 0.4995 |
| HMR_4536 | Amino sugar and nucleotide sugar metabolism | 703.7806 | 22942.2651 | 703.5683 | 22971.2977 | 0.2122 | 0.001 | 0.4996 |
| HMR_4628 | Amino sugar and nucleotide sugar metabolism | 1.523 | 1.4041 | 1.5692 | 1.4831 | 0.0462 | -0.0272 | 0.5109 |
| HMR_4631 | Amino sugar and nucleotide sugar metabolism | 5.1901 | 3.2085 | 5.2901 | 3.3302 | 0.1 | -0.0391 | 0.5156 |
| HMR_8377 | Amino sugar and nucleotide sugar metabolism | 214.2059 | 28853.9175 | 214.3499 | 28879.6049 | 0.144 | -0.0006 | 0.5002 |
| HMR_5130 | Aminoacyl-tRNA biosynthesis | 0.1936 | 0.0075 | 0.1837 | 0.0071 | 0.0098 | 0.0814 | 0.4676 |
| HMR_5131 | Aminoacyl-tRNA biosynthesis | 0.5828 | 0.0314 | 0.551 | 0.0284 | 0.0318 | 0.1301 | 0.4483 |
| HMR_5132 | Aminoacyl-tRNA biosynthesis | 0.3387 | 0.0141 | 0.3202 | 0.013 | 0.0185 | 0.1121 | 0.4554 |
| HMR_5133 | Aminoacyl-tRNA biosynthesis | 0.1415 | 0.0055 | 0.1354 | 0.0052 | 0.0061 | 0.0584 | 0.4767 |
| HMR_5134 | Aminoacyl-tRNA biosynthesis | 0.3802 | 0.0151 | 0.3588 | 0.0138 | 0.0215 | 0.1262 | 0.4498 |
| HMR_5135 | Aminoacyl-tRNA biosynthesis | 0.1197 | 0.0048 | 0.1147 | 0.0045 | 0.005 | 0.0519 | 0.4793 |
| HMR_5136 | Aminoacyl-tRNA biosynthesis | 0.3335 | 0.0141 | 0.3151 | 0.013 | 0.0184 | 0.1118 | 0.4555 |
| HMR_5137 | Aminoacyl-tRNA biosynthesis | 0.4746 | 0.0223 | 0.4485 | 0.0203 | 0.0262 | 0.1267 | 0.4496 |
| HMR_5138 | Aminoacyl-tRNA biosynthesis | 0.3855 | 0.0155 | 0.3645 | 0.0143 | 0.021 | 0.1219 | 0.4515 |
| HMR_5139 | Aminoacyl-tRNA biosynthesis | 0.1242 | 0.0047 | 0.1193 | 0.0045 | 0.005 | 0.0515 | 0.4795 |
| HMR_5140 | Aminoacyl-tRNA biosynthesis | 0.2161 | 0.0092 | 0.2009 | 0.0085 | 0.0153 | 0.1149 | 0.4542 |
| HMR_5141 | Aminoacyl-tRNA biosynthesis | 0.9152 | 0.0623 | 0.8636 | 0.0554 | 0.0516 | 0.1503 | 0.4403 |
| HMR_5142 | Aminoacyl-tRNA biosynthesis | 0.4449 | 0.0198 | 0.4201 | 0.018 | 0.0248 | 0.1275 | 0.4493 |
| HMR_5143 | Aminoacyl-tRNA biosynthesis | 0.2118 | 0.0078 | 0.2041 | 0.0074 | 0.0077 | 0.0629 | 0.4749 |
| HMR_5145 | Aminoacyl-tRNA biosynthesis | 0.2563 | 0.0096 | 0.242 | 0.0089 | 0.0143 | 0.105 | 0.4582 |
| HMR_5146 | Aminoacyl-tRNA biosynthesis | 0.3848 | 0.0165 | 0.363 | 0.0151 | 0.0218 | 0.1228 | 0.4511 |
| HMR_5147 | Aminoacyl-tRNA biosynthesis | 0.6078 | 0.0308 | 0.5734 | 0.0277 | 0.0345 | 0.1425 | 0.4433 |
| HMR_5148 | Aminoacyl-tRNA biosynthesis | 0.3603 | 0.015 | 0.3401 | 0.0138 | 0.0201 | 0.1186 | 0.4528 |
| HMR_5149 | Aminoacyl-tRNA biosynthesis | 0.1075 | 0.0041 | 0.1037 | 0.0039 | 0.0038 | 0.0428 | 0.4829 |
| HMR_5150 | Aminoacyl-tRNA biosynthesis | 0.5096 | 0.0227 | 0.476 | 0.0206 | 0.0336 | 0.1616 | 0.4358 |
| HMR_5155 | Protein assembly | 0.0055 | 0.00E+00 | 0.0051 | 0.00E+00 | 0.0004 | 0.0498 | 0.4801 |
| HMR_5157 | Protein assembly | 0.0083 | 1.00E-04 | 0.0077 | 1.00E-04 | 0.0006 | 0.051 | 0.4797 |
| HMR_5158 | Protein assembly | 0.0067 | 0.00E+00 | 0.0064 | 0.00E+00 | 0.0003 | 0.0369 | 0.4853 |
| HMR_5159 | Protein assembly | 0.0071 | 0.00E+00 | 0.0067 | 0.00E+00 | 0.0004 | 0.0424 | 0.4831 |
| HMR_5166 | Protein assembly | 0.0026 | 0.00E+00 | 0.0025 | 0.00E+00 | 0.0001 | 0.0342 | 0.4863 |

| | | | | | | | | |
|---|---|---|---|---|---|---|---|---|
| HMR_5169 | Protein assembly | 0.0025 | 0.00E+00 | 0.0024 | 0.00E+00 | 0.0001 | 0.0386 | 0.4846 |
| HMR_5170 | Protein assembly | 0.0038 | 0.00E+00 | 0.0036 | 0.00E+00 | 0.0003 | 0.0517 | 0.4794 |
| HMR_5171 | Protein assembly | 0.0022 | 0.00E+00 | 0.0021 | 0.00E+00 | 1.00E-04 | 0.0309 | 0.4877 |
| HMR_5172 | Protein assembly | 0.0021 | 0.00E+00 | 0.0019 | 0.00E+00 | 0.0001 | 0.0476 | 0.481 |
| HMR_5173 | Protein assembly | 0.0015 | 0.00E+00 | 0.0015 | 0.00E+00 | 1.00E-04 | 0.0419 | 0.4833 |
| HMR_7617 | Protein modification | 2.8476 | 3.7899 | 2.7186 | 3.4543 | 0.129 | 0.0479 | 0.4809 |
| HMR_7618 | Protein modification | 2.8471 | 3.7899 | 2.718 | 3.4543 | 0.129 | 0.0479 | 0.4809 |
| HMR_7621 | Protein modification | 1.9506 | 1.2901 | 1.9493 | 1.2883 | 0.0013 | 0.0008 | 0.4997 |
| HMR_7624 | Protein modification | 1.6178 | 1.1556 | 1.6017 | 1.1339 | 0.0161 | 0.0106 | 0.4958 |
| HMR_7626 | Protein modification | 1.95 | 1.2871 | 1.9487 | 1.2853 | 0.0014 | 0.0009 | 0.4997 |
| HMR_5266 | Protein degradation | 0.0494 | 0.0027 | 0.0764 | 0.0048 | 0.027 | -0.3115 | 0.6223 |
| HMR_5268 | Protein degradation | 0.0475 | 0.0017 | 0.0611 | 0.0024 | 0.0137 | -0.212 | 0.584 |
| HMR_5269 | Protein degradation | 0.0448 | 0.0015 | 0.0519 | 0.0019 | 0.0072 | -0.1216 | 0.5484 |
| HMR_5270 | Protein degradation | 0.0436 | 0.0015 | 0.0526 | 0.0019 | 0.009 | -0.1546 | 0.5614 |
| HMR_5283 | Protein degradation | 0.0195 | 0.0004 | 0.023 | 0.0005 | 0.0034 | -0.1179 | 0.5469 |
| HMR_5284 | Protein degradation | 0.0269 | 0.0006 | 0.0332 | 0.0009 | 0.0062 | -0.1623 | 0.5645 |
| HMR_5285 | Protein degradation | 0.0262 | 0.0006 | 0.0365 | 0.001 | 0.0103 | -0.2609 | 0.6029 |
| HMR_5286 | Protein degradation | 0.0323 | 0.0008 | 0.0489 | 0.0017 | 0.0166 | -0.3329 | 0.6304 |
| HMR_5287 | Protein degradation | 0.0192 | 0.0003 | 0.0225 | 0.0004 | 0.0032 | -0.1174 | 0.5467 |
| HMR_5288 | Protein degradation | 0.0188 | 0.0004 | 0.0311 | 0.0008 | 0.0124 | -0.3548 | 0.6386 |
| HMR_5289 | Protein degradation | 0.0144 | 0.0002 | 0.0208 | 0.0003 | 0.0064 | -0.2841 | 0.6118 |
| HMR_5291 | Protein degradation | 0.015 | 0.0002 | 0.0277 | 0.0005 | 0.0127 | -0.4705 | 0.681 |
| HMR_5407 | Nitrogen metabolism | 6.4691 | 16.7966 | 5.9754 | 14.3556 | 0.4938 | 0.0885 | 0.4648 |
| HMR_3851 | Sulfur metabolism | 11.5679 | 60.5865 | 12.6152 | 72.0109 | 1.0474 | -0.091 | 0.5362 |
| HMR_3861 | Sulfur metabolism | 11.5674 | 60.586 | 12.6148 | 72.0105 | 1.0474 | -0.091 | 0.5362 |
| HMR_3866 | Sulfur metabolism | 26.7662 | 267.5985 | 34.1555 | 397.8187 | 7.3893 | -0.2865 | 0.6127 |
| HMR_3915 | Sulfur metabolism | 15.7134 | 229.3958 | 17.5326 | 283.3851 | 1.8191 | -0.0803 | 0.532 |
| HMR_4084 | Sulfur metabolism | 14.6587 | 12.0925 | 14.4819 | 11.8931 | 0.1768 | 0.0361 | 0.4856 |
| HMR_4187 | Sulfur metabolism | 2.8436 | 8.2841 | 2.7148 | 7.5428 | 0.1288 | 0.0324 | 0.4871 |
| HMR_4701 | Sulfur metabolism | 96.3442 | 1447.5511 | 104.8067 | 1842.6752 | 8.4626 | -0.1475 | 0.5586 |
| HMR_8662 | C5-branched dibasic acid metabolism | -198.6215 | 209135.7445 | -202.1654 | 208333.0264 | 3.5438 | 0.0055 | 0.4978 |
| HMR_8663 | C5-branched dibasic acid metabolism | -205.2863 | 209136.87 | -208.8474 | 208334.3354 | 3.5612 | 0.0055 | 0.4978 |
| HMR_8664 | C5-branched dibasic acid metabolism | -6.6623 | 8.8872 | -6.6791 | 8.9381 | 0.0168 | 0.004 | 0.4984 |
| HMR_8665 | C5-branched dibasic acid metabolism | 3.499 | 5.033 | 3.5077 | 5.0583 | 0.0086 | -0.0027 | 0.5011 |
| HMR_8666 | C5-branched dibasic acid metabolism | 3.1607 | 6.7715 | 3.1683 | 6.8073 | 0.0077 | -0.0021 | 0.5008 |
| HMR_8667 | C5-branched dibasic acid metabolism | -201.9772 | 209136.9335 | -205.5303 | 208334.1407 | 3.5531 | 0.0055 | 0.4978 |
| HMR_8668 | C5-branched dibasic acid metabolism | -201.979 | 209136.9326 | -205.5321 | 208334.1398 | 3.5531 | 0.0055 | 0.4978 |
| HMR_8669 | C5-branched dibasic acid metabolism | 3.4935 | 5.0361 | 3.5017 | 5.0615 | 0.0082 | -0.0026 | 0.501 |
| HMR_0710 | Tricarboxylic acid cycle and glyoxylate/dicarboxylate metabolism | 4.4329 | 25.3019 | 4.2421 | 22.9543 | 0.1908 | 0.0275 | 0.489 |
| HMR_3787 | Tricarboxylic acid cycle and glyoxylate/dicarboxylate metabolism | 6.1176 | 38.9677 | 6.3416 | 41.59 | 0.224 | -0.025 | 0.51 |
| HMR_3957 | Tricarboxylic acid cycle and glyoxylate/dicarboxylate metabolism | 188.8379 | 21834.3084 | 190.8248 | 22148.772 | 1.9869 | -0.0095 | 0.5038 |
| HMR_3958 | Tricarboxylic acid cycle and glyoxylate/dicarboxylate metabolism | 201.4573 | 23331.725 | 199.5893 | 23189.6889 | 1.868 | 0.0087 | 0.4965 |
| HMR_4111 | Tricarboxylic acid cycle and glyoxylate/dicarboxylate metabolism | 4.5011 | 9.4124 | 4.3046 | 8.6113 | 0.1965 | 0.0463 | 0.4815 |
| HMR_4112 | Tricarboxylic acid cycle and glyoxylate/dicarboxylate metabolism | 188.981 | 21664.7952 | 189.3453 | 21766.9372 | 0.3643 | -0.0017 | 0.5007 |

| | | | | | | | | |
|---|---|---|---|---|---|---|---|---|
| HMR_4113 | Tricarboxylic acid cycle and glyoxylate/dicarboxylate metabolism | 4.1108 | 6.1063 | 4.0713 | 5.9876 | 0.0396 | 0.0114 | 0.4955 |
| HMR_4139 | Tricarboxylic acid cycle and glyoxylate/dicarboxylate metabolism | -406.0951 | 72489.5424 | -330.2298 | 71030.3695 | 75.8653 | -0.2003 | 0.5794 |
| HMR_4141 | Tricarboxylic acid cycle and glyoxylate/dicarboxylate metabolism | -195.1828 | 72866.8115 | -256.2166 | 72177.2844 | 61.0338 | 0.1603 | 0.4363 |
| HMR_4145 | Tricarboxylic acid cycle and glyoxylate/dicarboxylate metabolism | -537.2401 | 4590.9263 | -539.8813 | 5044.8112 | 2.6412 | 0.0269 | 0.4893 |
| HMR_4147 | Tricarboxylic acid cycle and glyoxylate/dicarboxylate metabolism | -417.6905 | 169159.6667 | -408.2763 | 173762.3414 | 9.4143 | -0.0161 | 0.5064 |
| HMR_4149 | Tricarboxylic acid cycle and glyoxylate/dicarboxylate metabolism | 1.8537 | 3.6138 | 1.7851 | 3.3458 | 0.0685 | 0.026 | 0.4896 |
| HMR_4152 | Tricarboxylic acid cycle and glyoxylate/dicarboxylate metabolism | 223.1688 | 169586.4594 | 198.0324 | 174217.5094 | 25.1364 | 0.0429 | 0.4829 |
| HMR_4209 | Tricarboxylic acid cycle and glyoxylate/dicarboxylate metabolism | 108.9185 | 4645.498 | 193.4631 | 14443.6438 | 84.5446 | -0.6119 | 0.7297 |
| HMR_4408 | Tricarboxylic acid cycle and glyoxylate/dicarboxylate metabolism | -274.0191 | 1756.9443 | -311.4262 | 1654.172 | 37.4071 | 0.6405 | 0.2609 |
| HMR_4410 | Tricarboxylic acid cycle and glyoxylate/dicarboxylate metabolism | 971.125 | 758.2075 | 974.4708 | 618.7853 | 3.3458 | -0.0902 | 0.5359 |
| HMR_4454 | Tricarboxylic acid cycle and glyoxylate/dicarboxylate metabolism | 196.7746 | 219937.2616 | 196.489 | 219989.9173 | 0.2856 | 0.0004 | 0.4998 |
| HMR_4456 | Tricarboxylic acid cycle and glyoxylate/dicarboxylate metabolism | 231.8781 | 241773.4138 | 232.0445 | 241774.3363 | 0.1663 | -0.0002 | 0.5001 |
| HMR_4458 | Tricarboxylic acid cycle and glyoxylate/dicarboxylate metabolism | -167.2287 | 207823.823 | -167.3838 | 207807.7477 | 0.1551 | 0.0002 | 0.4999 |
| HMR_4465 | Tricarboxylic acid cycle and glyoxylate/dicarboxylate metabolism | 5.3891 | 19.5867 | 5.007 | 16.9541 | 0.3821 | 0.0632 | 0.4748 |
| HMR_4585 | Tricarboxylic acid cycle and glyoxylate/dicarboxylate metabolism | -4.5014 | 9.4124 | -4.305 | 8.6113 | 0.1964 | -0.0463 | 0.5185 |
| HMR_4586 | Tricarboxylic acid cycle and glyoxylate/dicarboxylate metabolism | -212.7393 | 224270.0147 | -176.858 | 229360.2086 | 35.8813 | -0.0533 | 0.5212 |
| HMR_4587 | Tricarboxylic acid cycle and glyoxylate/dicarboxylate metabolism | -4.1114 | 6.1032 | -4.072 | 5.9844 | 0.0395 | -0.0113 | 0.5045 |
| HMR_4588 | Tricarboxylic acid cycle and glyoxylate/dicarboxylate metabolism | 23.7596 | 226704.7656 | -12.4924 | 231150.9447 | 36.252 | 0.0536 | 0.4786 |
| HMR_4589 | Tricarboxylic acid cycle and glyoxylate/dicarboxylate metabolism | 167.2288 | 207823.8182 | 167.3841 | 207807.7446 | 0.1553 | -0.0002 | 0.5001 |
| HMR_4652 | Tricarboxylic acid cycle and glyoxylate/dicarboxylate metabolism | -391.9269 | 170382.0649 | -360.4776 | 177681.8303 | 31.4492 | -0.0533 | 0.5213 |
| HMR_5294 | Tricarboxylic acid cycle and glyoxylate/dicarboxylate metabolism | 11.6439 | 152.7193 | 12.1087 | 158.171 | 0.4647 | -0.0264 | 0.5105 |
| HMR_5297 | Tricarboxylic acid cycle and glyoxylate/dicarboxylate metabolism | 114.8465 | 6661.0282 | 239.2187 | 19702.574 | 124.3721 | -0.766 | 0.7782 |
| HMR_6411 | Tricarboxylic acid cycle and glyoxylate/dicarboxylate metabolism | 96.6179 | 5739.0272 | 170.7694 | 17458.2444 | 74.1515 | -0.4869 | 0.6868 |
| HMR_6413 | Tricarboxylic acid cycle and glyoxylate/dicarboxylate metabolism | 108.9188 | 4645.4979 | 193.4635 | 14443.6429 | 84.5447 | -0.6119 | 0.7297 |
| HMR_6414 | Tricarboxylic acid cycle and glyoxylate/dicarboxylate metabolism | 205.5347 | 6661.693 | 364.2391 | 19361.2244 | 158.7044 | -0.9838 | 0.8374 |
| HMR_8743 | Tricarboxylic acid cycle and glyoxylate/dicarboxylate metabolism | -285.204 | 170456.0782 | -283.021 | 177748.7456 | 2.1831 | -0.0037 | 0.5015 |
| HMR_8772 | Tricarboxylic acid cycle and glyoxylate/dicarboxylate metabolism | 27.3 | 185.9912 | 28.3264 | 199.1087 | 1.0264 | -0.0523 | 0.5209 |
| HMR_8773 | Tricarboxylic acid cycle and glyoxylate/dicarboxylate metabolism | 27.3004 | 185.9847 | 28.3268 | 199.1021 | 1.0264 | -0.0523 | 0.5209 |
| HMR_8775 | Tricarboxylic acid cycle and glyoxylate/dicarboxylate metabolism | 27.3002 | 185.9815 | 28.3266 | 199.0988 | 1.0264 | -0.0523 | 0.5209 |
| HMR_8780 | Tricarboxylic acid cycle and glyoxylate/dicarboxylate metabolism | 39.3725 | 191.9127 | 39.9386 | 204.0312 | 0.5661 | -0.0284 | 0.5113 |
| HMR_8781 | Tricarboxylic acid cycle and glyoxylate/dicarboxylate metabolism | 7.3961 | 27.3782 | 6.9457 | 24.1348 | 0.4504 | 0.0628 | 0.475 |
| HMR_8782 | Tricarboxylic acid cycle and glyoxylate/dicarboxylate metabolism | 5.4225 | 19.7455 | 5.0356 | 17.0817 | 0.3869 | 0.0638 | 0.4746 |
| HMR_3975 | Oxidative phosphorylation | 4.8102 | 26.1917 | 4.0651 | 17.5487 | 0.7451 | 0.1127 | 0.4551 |
| HMR_3977 | Oxidative phosphorylation | 4.8201 | 26.2847 | 4.0721 | 17.5997 | 0.748 | 0.1129 | 0.4551 |
| HMR_3979 | Oxidative phosphorylation | 0.7111 | 0.2195 | 0.7047 | 0.2165 | 0.0065 | 0.0098 | 0.4961 |
| HMR_6911 | Oxidative phosphorylation | 157.7749 | 282365.306 | 108.7005 | 290210.4358 | 49.0744 | 0.0649 | 0.4741 |
| HMR_6912 | Oxidative phosphorylation | 508.2222 | 424.7767 | 522.9237 | 457.5579 | 14.7015 | -0.4949 | 0.6897 |
| HMR_6914 | Oxidative phosphorylation | 556.4294 | 626.1916 | 529.985 | 391.0247 | 26.4444 | 0.8291 | 0.2035 |
| HMR_6916 | Oxidative phosphorylation | 993.4695 | 51.6153 | 993.6592 | 45.2438 | 0.1897 | -0.0193 | 0.5077 |
| HMR_6918 | Oxidative phosphorylation | 990.8335 | 78.8216 | 987.4079 | 141.9929 | 3.4256 | 0.2305 | 0.4088 |
| HMR_6921 | Oxidative phosphorylation | 439.3819 | 4481.0589 | 585.5435 | 5783.3944 | 146.1616 | -1.4427 | 0.9254 |
| HMR_3980 | ROS detoxification | 65.8499 | 2178.8386 | 68.8719 | 2381.5466 | 3.022 | -0.0447 | 0.5178 |
| HMR_3982 | ROS detoxification | 12.727 | 146.6913 | 11.445 | 123.0471 | 1.282 | 0.0781 | 0.4689 |
| HMR_6608 | ROS detoxification | 3.4838 | 220495.4002 | 4.2909 | 220486.8103 | 0.8071 | -0.0012 | 0.5005 |

| | | | | | | | | |
|---|---|---|---|---|---|---|---|---|
| HMR_0188 | Fatty acid activation (cytosolic) | -2.4269 | 5.04 | -2.4775 | 5.2002 | 0.0506 | 0.0158 | 0.4937 |
| HMR_0192 | Fatty acid activation (cytosolic) | -2.4731 | 2.2303 | -2.5279 | 2.3073 | 0.0548 | 0.0257 | 0.4897 |
| HMR_0196 | Fatty acid activation (cytosolic) | -4.075 | 4.6199 | -4.1447 | 4.75 | 0.0696 | 0.0228 | 0.4909 |
| HMR_0200 | Fatty acid activation (cytosolic) | 1.7925 | 3.1582 | 1.7979 | 3.2111 | 0.0055 | -0.0022 | 0.5009 |
| HMR_0204 | Fatty acid activation (cytosolic) | 1.8877 | 1.9769 | 1.9109 | 2.0223 | 0.0231 | -0.0116 | 0.5046 |
| HMR_0209 | Fatty acid activation (cytosolic) | -2.0034 | 1.9078 | -2.0276 | 1.9532 | 0.0243 | 0.0124 | 0.4951 |
| HMR_0213 | Fatty acid activation (cytosolic) | -2.5052 | 3.8487 | -2.5591 | 3.9518 | 0.054 | 0.0193 | 0.4923 |
| HMR_0217 | Fatty acid activation (cytosolic) | -28.6702 | 12.1051 | -29.6504 | 12.5566 | 0.9802 | 0.1974 | 0.4218 |
| HMR_0226 | Fatty acid activation (cytosolic) | -6.7699 | 5.1411 | -7.155 | 5.3739 | 0.3851 | 0.1188 | 0.4527 |
| HMR_0233 | Fatty acid activation (cytosolic) | -2.1248 | 4.7981 | -2.1216 | 5.0001 | 0.0032 | -0.001 | 0.5004 |
| HMR_0237 | Fatty acid activation (cytosolic) | -4.1434 | 3.4751 | -4.2844 | 3.6289 | 0.141 | 0.0529 | 0.4789 |
| HMR_0241 | Fatty acid activation (cytosolic) | 2.3228 | 2.0405 | 2.3859 | 2.1127 | 0.0632 | -0.031 | 0.5124 |
| HMR_0245 | Fatty acid activation (cytosolic) | -1.7625 | 3.7504 | -1.761 | 3.825 | 0.0015 | -0.0005 | 0.5002 |
| HMR_0249 | Fatty acid activation (cytosolic) | -15.9135 | 8.3141 | -16.4262 | 8.6791 | 0.5127 | 0.1244 | 0.4505 |
| HMR_0255 | Fatty acid activation (cytosolic) | 1.7594 | 3.4992 | 1.7621 | 3.5625 | 0.0027 | -0.001 | 0.5004 |
| HMR_0259 | Fatty acid activation (cytosolic) | -1.5192 | 4.801 | -1.4033 | 4.9747 | 0.116 | -0.0371 | 0.5148 |
| HMR_0263 | Fatty acid activation (cytosolic) | 21.2667 | 9.869 | 22.0956 | 10.4052 | 0.8288 | -0.1841 | 0.573 |
| HMR_0267 | Fatty acid activation (cytosolic) | -2.0153 | 1.9078 | -2.0388 | 1.9531 | 0.0235 | 0.012 | 0.4952 |
| HMR_0271 | Fatty acid activation (cytosolic) | -0.6324 | 0.1673 | -0.6874 | 0.1981 | 0.055 | 0.0909 | 0.4638 |
| HMR_0275 | Fatty acid activation (cytosolic) | 2.0109 | 1.9074 | 2.0344 | 1.9527 | 0.0236 | -0.012 | 0.5048 |
| HMR_0279 | Fatty acid activation (cytosolic) | -2.0272 | 1.9097 | -2.0511 | 1.9551 | 0.0239 | 0.0121 | 0.4952 |
| HMR_0283 | Fatty acid activation (cytosolic) | -1.9731 | 2.0161 | -1.9995 | 2.0636 | 0.0264 | 0.0131 | 0.4948 |
| HMR_0289 | Fatty acid activation (cytosolic) | 1.9667 | 1.9298 | 1.989 | 1.9746 | 0.0224 | -0.0113 | 0.5045 |
| HMR_0293 | Fatty acid activation (cytosolic) | -1.7334 | 5.3196 | -1.7213 | 5.5896 | 0.0121 | -0.0037 | 0.5015 |
| HMR_0297 | Fatty acid activation (cytosolic) | 1.8331 | 2.3238 | 1.848 | 2.3671 | 0.0149 | -0.0069 | 0.5028 |
| HMR_0301 | Fatty acid activation (cytosolic) | -1.9595 | 1.9275 | -1.9824 | 1.9725 | 0.0229 | 0.0116 | 0.4954 |
| HMR_0305 | Fatty acid activation (cytosolic) | -2.0484 | 1.9079 | -2.0745 | 1.9537 | 0.0261 | 0.0133 | 0.4947 |
| HMR_0309 | Fatty acid activation (cytosolic) | -2.0305 | 1.9096 | -2.0544 | 1.9549 | 0.0239 | 0.0121 | 0.4952 |
| HMR_0313 | Fatty acid activation (cytosolic) | -2.0544 | 1.9181 | -2.0814 | 1.9644 | 0.027 | 0.0137 | 0.4945 |
| HMR_0319 | Fatty acid activation (cytosolic) | -1.9769 | 1.9234 | -2.0002 | 1.9686 | 0.0233 | 0.0118 | 0.4953 |
| HMR_0323 | Fatty acid activation (cytosolic) | -1.9997 | 1.9105 | -2.0236 | 1.956 | 0.0239 | 0.0122 | 0.4952 |
| HMR_0327 | Fatty acid activation (cytosolic) | -2.0364 | 1.9048 | -2.0612 | 1.9503 | 0.0248 | 0.0126 | 0.495 |
| HMR_0331 | Fatty acid activation (cytosolic) | -2.0969 | 1.92 | -2.1276 | 1.9676 | 0.0307 | 0.0156 | 0.4938 |
| HMR_0337 | Fatty acid activation (cytosolic) | -2.0467 | 1.9075 | -2.0721 | 1.9531 | 0.0253 | 0.0129 | 0.4949 |
| HMR_0341 | Fatty acid activation (cytosolic) | -2.0601 | 1.9097 | -2.0863 | 1.9555 | 0.0262 | 0.0133 | 0.4947 |
| HMR_0345 | Fatty acid activation (cytosolic) | -2.0196 | 1.9061 | -2.0434 | 1.9514 | 0.0238 | 0.0121 | 0.4952 |
| HMR_0349 | Fatty acid activation (cytosolic) | -13.2609 | 7.3197 | -14.373 | 8.032 | 1.1121 | 0.2838 | 0.3883 |
| HMR_0353 | Fatty acid activation (cytosolic) | -2.063 | 1.9111 | -2.0866 | 1.9563 | 0.0236 | 0.012 | 0.4952 |
| HMR_0357 | Fatty acid activation (cytosolic) | -2.3028 | 2.2342 | -2.3203 | 2.2782 | 0.0175 | 0.0082 | 0.4967 |
| HMR_0361 | Fatty acid activation (cytosolic) | -63.0998 | 154240.9638 | -63.6068 | 154195.9752 | 0.507 | 0.0009 | 0.4996 |
| HMR_0365 | Fatty acid activation (cytosolic) | -2.3061 | 2.2688 | -2.3364 | 2.322 | 0.0303 | 0.0141 | 0.4944 |
| HMR_0369 | Fatty acid activation (cytosolic) | 2.0254 | 1.9079 | 2.0488 | 1.9532 | 0.0233 | -0.0119 | 0.5047 |
| HMR_0373 | Fatty acid activation (cytosolic) | 2.0249 | 1.9077 | 2.0483 | 1.9529 | 0.0234 | -0.0119 | 0.5048 |
| HMR_0377 | Fatty acid activation (cytosolic) | -3.5188 | 2.994 | -3.5694 | 3.0706 | 0.0506 | 0.0206 | 0.4918 |

| | | | | | | | | |
|---|---|---|---|---|---|---|---|---|
| HMR_0381 | Fatty acid activation (cytosolic) | 2.0548 | 1.9674 | 2.0884 | 2.018 | 0.0336 | -0.0168 | 0.5067 |
| HMR_0385 | Fatty acid activation (cytosolic) | 2.0181 | 1.9058 | 2.0423 | 1.9512 | 0.0242 | -0.0123 | 0.5049 |
| HMR_0389 | Fatty acid activation (cytosolic) | 2.0209 | 1.9077 | 2.0444 | 1.953 | 0.0235 | -0.0119 | 0.5048 |
| HMR_0393 | Fatty acid activation (cytosolic) | 2.014 | 1.9059 | 2.0377 | 1.9512 | 0.0237 | -0.0121 | 0.5048 |
| HMR_0397 | Fatty acid activation (cytosolic) | -30.0144 | 13.4766 | -31.0792 | 14.1873 | 1.0649 | 0.2025 | 0.4198 |
| HMR_0401 | Fatty acid activation (cytosolic) | -2.3024 | 2.2458 | -2.3663 | 2.3239 | 0.0639 | 0.0299 | 0.4881 |
| HMR_0405 | Fatty acid activation (cytosolic) | -2.7329 | 2.5558 | -2.7622 | 2.6119 | 0.0293 | 0.0129 | 0.4949 |
| HMR_0409 | Fatty acid activation (cytosolic) | -5.2936 | 5.7132 | -5.1646 | 5.8697 | 0.129 | -0.0379 | 0.5151 |
| HMR_0413 | Fatty acid activation (cytosolic) | -2.0486 | 2.0847 | -2.0801 | 2.1362 | 0.0314 | 0.0153 | 0.4939 |
| HMR_0417 | Fatty acid activation (cytosolic) | 2.0186 | 1.9067 | 2.0422 | 1.952 | 0.0237 | -0.0121 | 0.5048 |
| HMR_0421 | Fatty acid activation (cytosolic) | 2.0189 | 1.9067 | 2.0426 | 1.952 | 0.0237 | -0.0121 | 0.5048 |
| HMR_0425 | Fatty acid activation (cytosolic) | 2.1001 | 1.9465 | 2.1237 | 1.992 | 0.0236 | -0.0119 | 0.5047 |
| HMR_0429 | Fatty acid activation (cytosolic) | 1.903 | 2.7522 | 1.9247 | 2.8062 | 0.0217 | -0.0092 | 0.5037 |
| HMR_0433 | Fatty acid activation (cytosolic) | 2.0284 | 1.9047 | 2.0527 | 1.9502 | 0.0244 | -0.0124 | 0.5049 |
| HMR_0437 | Fatty acid activation (cytosolic) | -2.0153 | 1.9064 | -2.0393 | 1.9518 | 0.024 | 0.0122 | 0.4951 |
| HMR_2942 | Fatty acid activation (endoplasmic reticular) | -2.003 | 1.445 | -2.0271 | 1.4799 | 0.0241 | 0.0141 | 0.4944 |
| HMR_2943 | Fatty acid activation (endoplasmic reticular) | -2.003 | 1.445 | -2.0271 | 1.4799 | 0.0241 | 0.0141 | 0.4944 |
| HMR_2944 | Fatty acid activation (endoplasmic reticular) | -1.9603 | 1.4557 | -1.9841 | 1.4906 | 0.0239 | 0.0139 | 0.4945 |
| HMR_2945 | Fatty acid activation (endoplasmic reticular) | -2.003 | 1.445 | -2.0271 | 1.4799 | 0.0241 | 0.0141 | 0.4944 |
| HMR_2946 | Fatty acid activation (endoplasmic reticular) | 2.003 | 1.445 | 2.0271 | 1.4799 | 0.0241 | -0.0141 | 0.5056 |
| HMR_2947 | Fatty acid activation (endoplasmic reticular) | 2.003 | 1.445 | 2.0271 | 1.4799 | 0.0241 | -0.0141 | 0.5056 |
| HMR_2948 | Fatty acid activation (endoplasmic reticular) | -2.003 | 1.445 | -2.0271 | 1.4799 | 0.0241 | 0.0141 | 0.4944 |
| HMR_2949 | Fatty acid activation (endoplasmic reticular) | -1.9693 | 1.7147 | -1.9922 | 1.7546 | 0.0228 | 0.0122 | 0.4951 |
| HMR_2951 | Fatty acid activation (endoplasmic reticular) | -1.9428 | 1.5054 | -1.966 | 1.5408 | 0.0232 | 0.0133 | 0.4947 |
| HMR_2952 | Fatty acid activation (endoplasmic reticular) | -2.003 | 1.445 | -2.0271 | 1.4799 | 0.0241 | 0.0141 | 0.4944 |
| HMR_2954 | Fatty acid activation (endoplasmic reticular) | -2.003 | 1.445 | -2.0271 | 1.4799 | 0.0241 | 0.0141 | 0.4944 |
| HMR_2955 | Fatty acid activation (endoplasmic reticular) | 2.003 | 1.445 | 2.0271 | 1.4799 | 0.0241 | -0.0141 | 0.5056 |
| HMR_2956 | Fatty acid activation (endoplasmic reticular) | -2.003 | 1.445 | -2.0271 | 1.4799 | 0.0241 | 0.0141 | 0.4944 |
| HMR_2957 | Fatty acid activation (endoplasmic reticular) | -1.9581 | 1.4599 | -1.9818 | 1.4949 | 0.0237 | 0.0138 | 0.4945 |
| HMR_2959 | Fatty acid activation (endoplasmic reticular) | 1.9855 | 1.7905 | 2.0087 | 1.8323 | 0.0232 | -0.0122 | 0.5049 |
| HMR_2961 | Fatty acid activation (endoplasmic reticular) | -2.003 | 1.445 | -2.0271 | 1.4799 | 0.0241 | 0.0141 | 0.4944 |
| HMR_2962 | Fatty acid activation (endoplasmic reticular) | 2.003 | 1.445 | 2.0271 | 1.4799 | 0.0241 | -0.0141 | 0.5056 |
| HMR_2963 | Fatty acid activation (endoplasmic reticular) | -2.003 | 1.445 | -2.0271 | 1.4799 | 0.0241 | 0.0141 | 0.4944 |
| HMR_2964 | Fatty acid activation (endoplasmic reticular) | 2.003 | 1.445 | 2.0271 | 1.4799 | 0.0241 | -0.0141 | 0.5056 |
| HMR_2965 | Fatty acid activation (endoplasmic reticular) | -2.003 | 1.445 | -2.0271 | 1.4799 | 0.0241 | 0.0141 | 0.4944 |
| HMR_2966 | Fatty acid activation (endoplasmic reticular) | -2.003 | 1.445 | -2.0271 | 1.4799 | 0.0241 | 0.0141 | 0.4944 |
| HMR_2967 | Fatty acid activation (endoplasmic reticular) | -2.003 | 1.445 | -2.0271 | 1.4799 | 0.0241 | 0.0141 | 0.4944 |
| HMR_2968 | Fatty acid activation (endoplasmic reticular) | 2.003 | 1.445 | 2.0271 | 1.4799 | 0.0241 | -0.0141 | 0.5056 |
| HMR_2969 | Fatty acid activation (endoplasmic reticular) | 2.003 | 1.445 | 2.0271 | 1.4799 | 0.0241 | -0.0141 | 0.5056 |
| HMR_2970 | Fatty acid activation (endoplasmic reticular) | -2.003 | 1.445 | -2.0271 | 1.4799 | 0.0241 | 0.0141 | 0.4944 |
| HMR_2971 | Fatty acid activation (endoplasmic reticular) | -2.003 | 1.445 | -2.0271 | 1.4799 | 0.0241 | 0.0141 | 0.4944 |
| HMR_2972 | Fatty acid activation (endoplasmic reticular) | -2.003 | 1.445 | -2.0271 | 1.4799 | 0.0241 | 0.0141 | 0.4944 |
| HMR_2973 | Fatty acid activation (endoplasmic reticular) | -2.003 | 1.445 | -2.0271 | 1.4799 | 0.0241 | 0.0141 | 0.4944 |
| HMR_2974 | Fatty acid activation (endoplasmic reticular) | -2.003 | 1.445 | -2.0271 | 1.4799 | 0.0241 | 0.0141 | 0.4944 |

| | | | | | | | | |
|---|---|---|---|---|---|---|---|---|
| HMR_2975 | Fatty acid activation (endoplasmic reticular) | -2.003 | 1.445 | -2.0271 | 1.4799 | 0.0241 | 0.0141 | 0.4944 |
| HMR_2976 | Fatty acid activation (endoplasmic reticular) | -2.003 | 1.445 | -2.0271 | 1.4799 | 0.0241 | 0.0141 | 0.4944 |
| HMR_2977 | Fatty acid activation (endoplasmic reticular) | -2.003 | 1.445 | -2.0271 | 1.4799 | 0.0241 | 0.0141 | 0.4944 |
| HMR_2978 | Fatty acid activation (endoplasmic reticular) | -2.003 | 1.445 | -2.0271 | 1.4799 | 0.0241 | 0.0141 | 0.4944 |
| HMR_2979 | Fatty acid activation (endoplasmic reticular) | -2.003 | 1.445 | -2.0271 | 1.4799 | 0.0241 | 0.0141 | 0.4944 |
| HMR_2980 | Fatty acid activation (endoplasmic reticular) | -2.003 | 1.445 | -2.0271 | 1.4799 | 0.0241 | 0.0141 | 0.4944 |
| HMR_2981 | Fatty acid activation (endoplasmic reticular) | -1.961 | 1.4544 | -1.9849 | 1.4893 | 0.0239 | 0.0139 | 0.4944 |
| HMR_2982 | Fatty acid activation (endoplasmic reticular) | -2.003 | 1.445 | -2.0271 | 1.4799 | 0.0241 | 0.0141 | 0.4944 |
| HMR_2983 | Fatty acid activation (endoplasmic reticular) | -2.003 | 1.445 | -2.0271 | 1.4799 | 0.0241 | 0.0141 | 0.4944 |
| HMR_2984 | Fatty acid activation (endoplasmic reticular) | 58.4489 | 154238.1751 | 58.8973 | 154193.1137 | 0.4484 | -0.0008 | 0.5003 |
| HMR_2985 | Fatty acid activation (endoplasmic reticular) | -1.9954 | 1.4455 | -2.0195 | 1.4804 | 0.0241 | 0.0141 | 0.4944 |
| HMR_2986 | Fatty acid activation (endoplasmic reticular) | 2.003 | 1.445 | 2.0271 | 1.4799 | 0.0241 | -0.0141 | 0.5056 |
| HMR_2987 | Fatty acid activation (endoplasmic reticular) | 2.003 | 1.445 | 2.0271 | 1.4799 | 0.0241 | -0.0141 | 0.5056 |
| HMR_2988 | Fatty acid activation (endoplasmic reticular) | -1.9618 | 1.4532 | -1.9858 | 1.4881 | 0.024 | 0.014 | 0.4944 |
| HMR_2989 | Fatty acid activation (endoplasmic reticular) | 2.003 | 1.445 | 2.0271 | 1.4799 | 0.0241 | -0.0141 | 0.5056 |
| HMR_2990 | Fatty acid activation (endoplasmic reticular) | 2.003 | 1.445 | 2.0271 | 1.4799 | 0.0241 | -0.0141 | 0.5056 |
| HMR_2991 | Fatty acid activation (endoplasmic reticular) | -2.0123 | 1.8849 | -2.036 | 1.9297 | 0.0237 | 0.0121 | 0.4952 |
| HMR_2992 | Fatty acid activation (endoplasmic reticular) | -2.003 | 1.445 | -2.0271 | 1.4799 | 0.0241 | 0.0141 | 0.4944 |
| HMR_2994 | Fatty acid activation (endoplasmic reticular) | -1.9627 | 1.4534 | -1.9866 | 1.4883 | 0.0239 | 0.0139 | 0.4944 |
| HMR_2996 | Fatty acid activation (endoplasmic reticular) | -1.9451 | 1.5274 | -1.9679 | 1.5634 | 0.0228 | 0.013 | 0.4948 |
| HMR_2998 | Fatty acid activation (endoplasmic reticular) | -1.9954 | 1.4455 | -2.0195 | 1.4804 | 0.0241 | 0.0141 | 0.4944 |
| HMR_2999 | Fatty acid activation (endoplasmic reticular) | 2.003 | 1.445 | 2.0271 | 1.4799 | 0.0241 | -0.0141 | 0.5056 |
| HMR_3000 | Fatty acid activation (endoplasmic reticular) | 2.003 | 1.445 | 2.0271 | 1.4799 | 0.0241 | -0.0141 | 0.5056 |
| HMR_3001 | Fatty acid activation (endoplasmic reticular) | 2.003 | 1.445 | 2.0271 | 1.4799 | 0.0241 | -0.0141 | 0.5056 |
| HMR_3002 | Fatty acid activation (endoplasmic reticular) | 2.003 | 1.445 | 2.0271 | 1.4799 | 0.0241 | -0.0141 | 0.5056 |
| HMR_3003 | Fatty acid activation (endoplasmic reticular) | 2.003 | 1.445 | 2.0271 | 1.4799 | 0.0241 | -0.0141 | 0.5056 |
| HMR_2152 | Fatty acid biosynthesis (even-chain) | 2.2132 | 0.5925 | 2.2825 | 0.6325 | 0.0693 | -0.0626 | 0.525 |
| HMR_2153 | Fatty acid biosynthesis (even-chain) | 2.2101 | 0.5893 | 2.2797 | 0.6293 | 0.0696 | -0.0631 | 0.5252 |
| HMR_2154 | Fatty acid biosynthesis (even-chain) | 2.2068 | 0.5862 | 2.2768 | 0.6262 | 0.07 | -0.0636 | 0.5253 |
| HMR_2155 | Fatty acid biosynthesis (even-chain) | 2.2032 | 0.5832 | 2.2735 | 0.6231 | 0.0703 | -0.064 | 0.5255 |
| HMR_2156 | Fatty acid biosynthesis (even-chain) | 2.1994 | 0.5802 | 2.2701 | 0.6201 | 0.0707 | -0.0645 | 0.5257 |
| HMR_2157 | Fatty acid biosynthesis (even-chain) | 2.1978 | 0.5773 | 2.2688 | 0.6171 | 0.071 | -0.065 | 0.5259 |
| HMR_2158 | Fatty acid biosynthesis (even-chain) | 2.1961 | 0.5744 | 2.2675 | 0.6142 | 0.0713 | -0.0654 | 0.5261 |
| HMR_2159 | Fatty acid biosynthesis (even-chain) | 2.1941 | 0.5715 | 2.2657 | 0.6113 | 0.0716 | -0.0659 | 0.5263 |
| HMR_2160 | Fatty acid biosynthesis (even-chain) | 2.192 | 0.5687 | 2.2639 | 0.6085 | 0.072 | -0.0663 | 0.5264 |
| HMR_2161 | Fatty acid biosynthesis (even-chain) | 2.192 | 0.5658 | 2.2642 | 0.6057 | 0.0722 | -0.0667 | 0.5266 |
| HMR_2162 | Fatty acid biosynthesis (even-chain) | 2.1919 | 0.563 | 2.2644 | 0.6028 | 0.0725 | -0.0672 | 0.5268 |
| HMR_2163 | Fatty acid biosynthesis (even-chain) | 2.1915 | 0.5602 | 2.2643 | 0.6 | 0.0728 | -0.0676 | 0.5269 |
| HMR_2164 | Fatty acid biosynthesis (even-chain) | 2.1909 | 0.5573 | 2.264 | 0.5972 | 0.0731 | -0.068 | 0.5271 |
| HMR_2165 | Fatty acid biosynthesis (even-chain) | 2.1926 | 0.5545 | 2.2659 | 0.5944 | 0.0733 | -0.0684 | 0.5273 |
| HMR_2166 | Fatty acid biosynthesis (even-chain) | 2.1941 | 0.5516 | 2.2677 | 0.5916 | 0.0736 | -0.0688 | 0.5274 |
| HMR_2167 | Fatty acid biosynthesis (even-chain) | 2.1954 | 0.5487 | 2.2692 | 0.5887 | 0.0738 | -0.0692 | 0.5276 |
| HMR_2168 | Fatty acid biosynthesis (even-chain) | 2.1965 | 0.5457 | 2.2706 | 0.5858 | 0.0741 | -0.0697 | 0.5278 |
| HMR_2169 | Fatty acid biosynthesis (even-chain) | 2.1998 | 0.5427 | 2.2741 | 0.5828 | 0.0743 | -0.07 | 0.5279 |

| | | | | | | | | |
|---|---|---|---|---|---|---|---|---|
| HMR_2170 | Fatty acid biosynthesis (even-chain) | 2.203 | 0.5396 | 2.2775 | 0.5798 | 0.0745 | -0.0705 | 0.5281 |
| HMR_2171 | Fatty acid biosynthesis (even-chain) | 2.2059 | 0.5364 | 2.2806 | 0.5767 | 0.0747 | -0.0708 | 0.5282 |
| HMR_2173 | Fatty acid biosynthesis (even-chain) | 1.6718 | 0.438 | 1.7122 | 0.4647 | 0.0404 | -0.0425 | 0.517 |
| HMR_2174 | Fatty acid biosynthesis (even-chain) | 1.6766 | 0.4339 | 1.7172 | 0.4606 | 0.0406 | -0.0429 | 0.5171 |
| HMR_2175 | Fatty acid biosynthesis (even-chain) | 1.681 | 0.4304 | 1.7218 | 0.4572 | 0.0408 | -0.0433 | 0.5173 |
| HMR_2176 | Fatty acid biosynthesis (even-chain) | 1.685 | 0.4274 | 1.7259 | 0.4542 | 0.0409 | -0.0436 | 0.5174 |
| HMR_2178 | Fatty acid biosynthesis (even-chain) | 0.9165 | 0.2174 | 0.929 | 0.224 | 0.0125 | -0.0188 | 0.5075 |
| HMR_2179 | Fatty acid biosynthesis (even-chain) | 0.9203 | 0.2137 | 0.933 | 0.2203 | 0.0126 | -0.0192 | 0.5077 |
| HMR_2180 | Fatty acid biosynthesis (even-chain) | 0.9223 | 0.213 | 0.9351 | 0.2196 | 0.0128 | -0.0195 | 0.5078 |
| HMR_2181 | Fatty acid biosynthesis (even-chain) | 0.9221 | 0.2152 | 0.9351 | 0.2219 | 0.0129 | -0.0196 | 0.5078 |
| HMR_4156 | Fatty acid biosynthesis (even-chain) | 4.2637 | 14.3366 | 3.8198 | 11.6853 | 0.4439 | 0.087 | 0.4653 |
| HMR_4295 | Fatty acid biosynthesis (even-chain) | 133.5467 | 163.4802 | 142.3921 | 176.2583 | 8.8454 | -0.4799 | 0.6843 |
| HMR_2227 | Fatty acid biosynthesis (odd-chain) | 2.7292 | 0.6991 | 2.8284 | 0.748 | 0.0993 | -0.0825 | 0.5329 |
| HMR_2228 | Fatty acid biosynthesis (odd-chain) | 2.7234 | 0.6976 | 2.8235 | 0.7465 | 0.1 | -0.0833 | 0.5332 |
| HMR_2229 | Fatty acid biosynthesis (odd-chain) | 2.7199 | 0.696 | 2.8207 | 0.7449 | 0.1008 | -0.0839 | 0.5334 |
| HMR_2230 | Fatty acid biosynthesis (odd-chain) | 2.7164 | 0.6941 | 2.8179 | 0.743 | 0.1015 | -0.0847 | 0.5337 |
| HMR_2231 | Fatty acid biosynthesis (odd-chain) | 2.7126 | 0.692 | 2.8148 | 0.7409 | 0.1022 | -0.0854 | 0.534 |
| HMR_2232 | Fatty acid biosynthesis (odd-chain) | 2.7087 | 0.6898 | 2.8116 | 0.7386 | 0.1029 | -0.0861 | 0.5343 |
| HMR_2233 | Fatty acid biosynthesis (odd-chain) | 2.707 | 0.6873 | 2.8107 | 0.7362 | 0.1036 | -0.0868 | 0.5346 |
| HMR_2234 | Fatty acid biosynthesis (odd-chain) | 2.7053 | 0.6847 | 2.8097 | 0.7335 | 0.1043 | -0.0876 | 0.5349 |
| HMR_2235 | Fatty acid biosynthesis (odd-chain) | 2.7034 | 0.6818 | 2.8084 | 0.7306 | 0.105 | -0.0883 | 0.5352 |
| HMR_2236 | Fatty acid biosynthesis (odd-chain) | 2.7013 | 0.6787 | 2.807 | 0.7275 | 0.1057 | -0.0891 | 0.5355 |
| HMR_2237 | Fatty acid biosynthesis (odd-chain) | 2.7015 | 0.6754 | 2.8079 | 0.7241 | 0.1063 | -0.0899 | 0.5358 |
| HMR_2238 | Fatty acid biosynthesis (odd-chain) | 2.7017 | 0.6719 | 2.8087 | 0.7206 | 0.107 | -0.0907 | 0.5361 |
| HMR_2239 | Fatty acid biosynthesis (odd-chain) | 2.7016 | 0.6682 | 2.8092 | 0.7168 | 0.1077 | -0.0915 | 0.5364 |
| HMR_2240 | Fatty acid biosynthesis (odd-chain) | 2.7014 | 0.6642 | 2.8097 | 0.7128 | 0.1083 | -0.0923 | 0.5368 |
| HMR_2241 | Fatty acid biosynthesis (odd-chain) | 2.7035 | 0.6601 | 2.8124 | 0.7086 | 0.109 | -0.0931 | 0.5371 |
| HMR_2242 | Fatty acid biosynthesis (odd-chain) | 2.7055 | 0.6557 | 2.8151 | 0.7041 | 0.1096 | -0.094 | 0.5374 |
| HMR_2243 | Fatty acid biosynthesis (odd-chain) | 2.7072 | 0.651 | 2.8175 | 0.6994 | 0.1102 | -0.0948 | 0.5378 |
| HMR_2244 | Fatty acid biosynthesis (odd-chain) | 2.7089 | 0.6461 | 2.8198 | 0.6945 | 0.1109 | -0.0957 | 0.5381 |
| HMR_2245 | Fatty acid biosynthesis (odd-chain) | 2.7129 | 0.641 | 2.8243 | 0.6893 | 0.1114 | -0.0966 | 0.5385 |
| HMR_2246 | Fatty acid biosynthesis (odd-chain) | 2.7168 | 0.6356 | 2.8288 | 0.6838 | 0.112 | -0.0975 | 0.5389 |
| HMR_2247 | Fatty acid biosynthesis (odd-chain) | 2.7204 | 0.6299 | 2.833 | 0.6781 | 0.1126 | -0.0985 | 0.5392 |
| HMR_2249 | Fatty acid biosynthesis (odd-chain) | 2.1499 | 0.5506 | 2.2316 | 0.5905 | 0.0817 | -0.0765 | 0.5305 |
| HMR_2250 | Fatty acid biosynthesis (odd-chain) | 2.1557 | 0.5445 | 2.238 | 0.5842 | 0.0823 | -0.0774 | 0.5309 |
| HMR_2251 | Fatty acid biosynthesis (odd-chain) | 2.1613 | 0.5383 | 2.2442 | 0.5779 | 0.0828 | -0.0784 | 0.5313 |
| HMR_2252 | Fatty acid biosynthesis (odd-chain) | 2.1667 | 0.5321 | 2.2501 | 0.5716 | 0.0834 | -0.0794 | 0.5316 |
| HMR_2254 | Fatty acid biosynthesis (odd-chain) | 1.6579 | 0.4511 | 1.7159 | 0.4813 | 0.058 | -0.06 | 0.5239 |
| HMR_2255 | Fatty acid biosynthesis (odd-chain) | 1.665 | 0.4451 | 1.7235 | 0.4752 | 0.0585 | -0.061 | 0.5243 |
| HMR_2256 | Fatty acid biosynthesis (odd-chain) | 1.6718 | 0.4399 | 1.7309 | 0.4698 | 0.0591 | -0.062 | 0.5247 |
| HMR_2257 | Fatty acid biosynthesis (odd-chain) | 1.678 | 0.4352 | 1.7377 | 0.465 | 0.0597 | -0.0629 | 0.5251 |
| HMR_2305 | Fatty acid biosynthesis (unsaturated) | 5.5959 | 4.0414 | 6.0148 | 4.7313 | 0.4189 | -0.1414 | 0.5562 |
| HMR_2307 | Fatty acid biosynthesis (unsaturated) | 5.596 | 4.034 | 6.0144 | 4.7239 | 0.4185 | -0.1414 | 0.5562 |
| HMR_2309 | Fatty acid biosynthesis (unsaturated) | 5.5959 | 4.0271 | 6.014 | 4.717 | 0.4181 | -0.1414 | 0.5562 |

| | | | | | | | | |
|---|---|---|---|---|---|---|---|---|
| HMR_2311 | Fatty acid biosynthesis (unsaturated) | 5.5956 | 4.0206 | 6.0133 | 4.7106 | 0.4177 | -0.1414 | 0.5562 |
| HMR_2315 | Fatty acid biosynthesis (unsaturated) | 5.2871 | 3.2826 | 5.6848 | 3.8631 | 0.3976 | -0.1488 | 0.5591 |
| HMR_2317 | Fatty acid biosynthesis (unsaturated) | 5.2889 | 3.2743 | 5.6861 | 3.8548 | 0.3972 | -0.1488 | 0.5591 |
| HMR_2319 | Fatty acid biosynthesis (unsaturated) | 5.2907 | 3.2662 | 5.6875 | 3.8467 | 0.3968 | -0.1488 | 0.5591 |
| HMR_2321 | Fatty acid biosynthesis (unsaturated) | 5.2922 | 3.2585 | 5.6885 | 3.839 | 0.3963 | -0.1488 | 0.5591 |
| HMR_2324 | Fatty acid biosynthesis (unsaturated) | 1.59 | 0.3253 | 1.6462 | 0.3444 | 0.0562 | -0.0687 | 0.5274 |
| HMR_2326 | Fatty acid biosynthesis (unsaturated) | 1.5935 | 0.317 | 1.6492 | 0.336 | 0.0558 | -0.069 | 0.5275 |
| HMR_2328 | Fatty acid biosynthesis (unsaturated) | 1.5968 | 0.309 | 1.6521 | 0.328 | 0.0553 | -0.0693 | 0.5276 |
| HMR_2330 | Fatty acid biosynthesis (unsaturated) | 1.5997 | 0.3014 | 1.6546 | 0.3203 | 0.0549 | -0.0696 | 0.5277 |
| HMR_2332 | Fatty acid biosynthesis (unsaturated) | 0.8849 | 0.1616 | 0.9153 | 0.1715 | 0.0303 | -0.0526 | 0.521 |
| HMR_2334 | Fatty acid biosynthesis (unsaturated) | 0.8888 | 0.1547 | 0.9187 | 0.1645 | 0.0299 | -0.0529 | 0.5211 |
| HMR_2336 | Fatty acid biosynthesis (unsaturated) | 0.8915 | 0.1494 | 0.9211 | 0.1591 | 0.0296 | -0.0532 | 0.5212 |
| HMR_2338 | Fatty acid biosynthesis (unsaturated) | 0.893 | 0.1455 | 0.9222 | 0.1552 | 0.0292 | -0.0533 | 0.5212 |
| HMR_2342 | Fatty acid biosynthesis (unsaturated) | 4.3233 | 2.8778 | 4.6952 | 3.4207 | 0.3719 | -0.1482 | 0.5589 |
| HMR_2343 | Fatty acid biosynthesis (unsaturated) | 4.3304 | 2.8693 | 4.7015 | 3.4122 | 0.3711 | -0.1481 | 0.5589 |
| HMR_2344 | Fatty acid biosynthesis (unsaturated) | 4.3373 | 2.8617 | 4.7077 | 3.4046 | 0.3703 | -0.1479 | 0.5588 |
| HMR_2345 | Fatty acid biosynthesis (unsaturated) | 4.344 | 2.8549 | 4.7134 | 3.3979 | 0.3695 | -0.1478 | 0.5587 |
| HMR_2347 | Fatty acid biosynthesis (unsaturated) | 1.367 | 0.2586 | 1.5003 | 0.3002 | 0.1334 | -0.1784 | 0.5708 |
| HMR_2348 | Fatty acid biosynthesis (unsaturated) | 1.374 | 0.2496 | 1.5066 | 0.2911 | 0.1326 | -0.1803 | 0.5716 |
| HMR_2349 | Fatty acid biosynthesis (unsaturated) | 1.3807 | 0.2411 | 1.5126 | 0.2825 | 0.1319 | -0.1823 | 0.5723 |
| HMR_2350 | Fatty acid biosynthesis (unsaturated) | 1.387 | 0.233 | 1.5182 | 0.2742 | 0.1312 | -0.1842 | 0.5731 |
| HMR_2353 | Fatty acid biosynthesis (unsaturated) | 0.658 | 0.1017 | 0.713 | 0.1172 | 0.055 | -0.1175 | 0.5468 |
| HMR_2354 | Fatty acid biosynthesis (unsaturated) | 0.6642 | 0.0955 | 0.7187 | 0.1108 | 0.0545 | -0.12 | 0.5478 |
| HMR_2355 | Fatty acid biosynthesis (unsaturated) | 0.6685 | 0.0913 | 0.7227 | 0.1065 | 0.0542 | -0.122 | 0.5485 |
| HMR_2356 | Fatty acid biosynthesis (unsaturated) | 0.6709 | 0.0888 | 0.7249 | 0.1038 | 0.0541 | -0.1232 | 0.549 |
| HMR_2361 | Fatty acid biosynthesis (unsaturated) | 2.014 | 0.5296 | 2.2824 | 0.6696 | 0.2684 | -0.2451 | 0.5968 |
| HMR_2362 | Fatty acid biosynthesis (unsaturated) | 2.0151 | 0.5208 | 2.2831 | 0.6608 | 0.2679 | -0.2465 | 0.5973 |
| HMR_2363 | Fatty acid biosynthesis (unsaturated) | 2.0161 | 0.5125 | 2.2836 | 0.6525 | 0.2675 | -0.2478 | 0.5979 |
| HMR_2364 | Fatty acid biosynthesis (unsaturated) | 2.0168 | 0.5046 | 2.2838 | 0.6445 | 0.2671 | -0.2491 | 0.5984 |
| HMR_2190 | Fatty acid elongation (even-chain) | 11.7201 | 6.7272 | 12.4241 | 7.7723 | 0.704 | -0.1849 | 0.5733 |
| HMR_2191 | Fatty acid elongation (even-chain) | 11.7242 | 6.7263 | 12.4275 | 7.7713 | 0.7033 | -0.1847 | 0.5733 |
| HMR_2193 | Fatty acid elongation (even-chain) | 11.7282 | 6.7254 | 12.4309 | 7.7703 | 0.7027 | -0.1846 | 0.5732 |
| HMR_2194 | Fatty acid elongation (even-chain) | 11.732 | 6.7245 | 12.4339 | 7.7693 | 0.7019 | -0.1844 | 0.5731 |
| HMR_2201 | Fatty acid elongation (even-chain) | 4.6848 | 3.6384 | 5.0577 | 4.2701 | 0.3729 | -0.1326 | 0.5528 |
| HMR_2202 | Fatty acid elongation (even-chain) | 4.6907 | 3.6308 | 5.0629 | 4.2624 | 0.3722 | -0.1325 | 0.5527 |
| HMR_2203 | Fatty acid elongation (even-chain) | 4.6964 | 3.6242 | 5.0679 | 4.2559 | 0.3715 | -0.1323 | 0.5526 |
| HMR_2204 | Fatty acid elongation (even-chain) | 4.7018 | 3.6189 | 5.0725 | 4.2506 | 0.3708 | -0.1322 | 0.5526 |
| HMR_2205 | Fatty acid elongation (even-chain) | 4.7932 | 3.1539 | 5.1765 | 3.7206 | 0.3832 | -0.1462 | 0.5581 |
| HMR_2208 | Fatty acid elongation (even-chain) | 4.8006 | 3.1458 | 5.1831 | 3.7125 | 0.3825 | -0.1461 | 0.5581 |
| HMR_2209 | Fatty acid elongation (even-chain) | 4.8079 | 3.1383 | 5.1897 | 3.705 | 0.3818 | -0.1459 | 0.558 |
| HMR_2210 | Fatty acid elongation (even-chain) | 4.8149 | 3.1313 | 5.1959 | 3.6981 | 0.381 | -0.1458 | 0.558 |
| HMR_2211 | Fatty acid elongation (even-chain) | 1.212 | 0.3109 | 1.2555 | 0.3305 | 0.0435 | -0.0543 | 0.5216 |
| HMR_2212 | Fatty acid elongation (even-chain) | 1.2202 | 0.3044 | 1.263 | 0.3239 | 0.0427 | -0.0539 | 0.5215 |
| HMR_2213 | Fatty acid elongation (even-chain) | 1.2275 | 0.2998 | 1.2696 | 0.3192 | 0.0421 | -0.0535 | 0.5213 |

| | | | | | | | | |
|---|---|---|---|---|---|---|---|---|
| HMR_2214 | Fatty acid elongation (even-chain) | 1.2338 | 0.2971 | 1.2752 | 0.3163 | 0.0414 | -0.0528 | 0.5211 |
| HMR_2215 | Fatty acid elongation (even-chain) | 3.5774 | 2.6895 | 3.7018 | 2.8795 | 0.1244 | -0.0527 | 0.521 |
| HMR_2217 | Fatty acid elongation (even-chain) | 3.5848 | 2.6827 | 3.7085 | 2.8726 | 0.1238 | -0.0525 | 0.5209 |
| HMR_2218 | Fatty acid elongation (even-chain) | 3.5918 | 2.6779 | 3.715 | 2.8679 | 0.1231 | -0.0523 | 0.5209 |
| HMR_2219 | Fatty acid elongation (even-chain) | 3.5983 | 2.6752 | 3.7208 | 2.8651 | 0.1225 | -0.052 | 0.5208 |
| HMR_2259 | Fatty acid elongation (odd-chain) | 3.9389 | 3.1305 | 4.2845 | 3.7164 | 0.3456 | -0.1321 | 0.5525 |
| HMR_2260 | Fatty acid elongation (odd-chain) | 3.9522 | 3.1248 | 4.2976 | 3.7107 | 0.3454 | -0.1321 | 0.5525 |
| HMR_2261 | Fatty acid elongation (odd-chain) | 3.9652 | 3.1211 | 4.3104 | 3.7068 | 0.3452 | -0.1321 | 0.5525 |
| HMR_2262 | Fatty acid elongation (odd-chain) | 3.9777 | 3.1191 | 4.3227 | 3.7048 | 0.3449 | -0.132 | 0.5525 |
| HMR_2263 | Fatty acid elongation (odd-chain) | 4.0429 | 2.767 | 4.3975 | 3.2993 | 0.3545 | -0.1439 | 0.5572 |
| HMR_2264 | Fatty acid elongation (odd-chain) | 4.0573 | 2.7599 | 4.4116 | 3.2921 | 0.3543 | -0.144 | 0.5573 |
| HMR_2265 | Fatty acid elongation (odd-chain) | 4.0715 | 2.7539 | 4.4255 | 3.2861 | 0.3541 | -0.1441 | 0.5573 |
| HMR_2266 | Fatty acid elongation (odd-chain) | 4.0852 | 2.7489 | 4.4391 | 3.2811 | 0.3539 | -0.1441 | 0.5573 |
| HMR_2267 | Fatty acid elongation (odd-chain) | 0.4628 | 0.0607 | 0.4695 | 0.0624 | 0.0067 | -0.0191 | 0.5076 |
| HMR_2268 | Fatty acid elongation (odd-chain) | 0.4739 | 0.0557 | 0.4804 | 0.0573 | 0.0065 | -0.0194 | 0.5078 |
| HMR_2269 | Fatty acid elongation (odd-chain) | 0.4814 | 0.0533 | 0.4878 | 0.0548 | 0.0065 | -0.0196 | 0.5078 |
| HMR_2270 | Fatty acid elongation (odd-chain) | 0.4857 | 0.0528 | 0.492 | 0.0543 | 0.0064 | -0.0194 | 0.5078 |
| HMR_2281 | Fatty acid desaturation (even-chain) | 1.1592 | 0.2666 | 1.2825 | 0.3273 | 0.1233 | -0.16 | 0.5636 |
| HMR_2282 | Fatty acid desaturation (even-chain) | 0.7705 | 0.3229 | 0.8662 | 0.393 | 0.0957 | -0.1131 | 0.545 |
| HMR_2284 | Fatty acid desaturation (even-chain) | 0.8236 | 0.2901 | 0.9251 | 0.3571 | 0.1015 | -0.1262 | 0.5502 |
| HMR_2286 | Fatty acid desaturation (even-chain) | 0.804 | 0.613 | 0.9183 | 0.794 | 0.1143 | -0.0964 | 0.5384 |
| HMR_2287 | Fatty acid desaturation (even-chain) | 1.0166 | 0.6514 | 1.1119 | 0.7945 | 0.0953 | -0.0793 | 0.5316 |
| HMR_2292 | Fatty acid desaturation (even-chain) | 0.7062 | 0.2665 | 0.7885 | 0.3291 | 0.0823 | -0.1066 | 0.5425 |
| HMR_2293 | Fatty acid desaturation (even-chain) | 3.4664 | 1.1332 | 3.7311 | 1.2849 | 0.2647 | -0.1702 | 0.5676 |
| HMR_2294 | Fatty acid desaturation (even-chain) | 1.1652 | 1.0278 | 1.2421 | 1.172 | 0.0769 | -0.0519 | 0.5207 |
| HMR_2295 | Fatty acid desaturation (even-chain) | 0.7945 | 0.1647 | 0.8814 | 0.2002 | 0.0868 | -0.1438 | 0.5572 |
| HMR_2296 | Fatty acid desaturation (even-chain) | 2.7842 | 0.5912 | 3.0608 | 0.702 | 0.2767 | -0.2433 | 0.5961 |
| HMR_2359 | Fatty acid desaturation (even-chain) | 3.342 | 0.9884 | 3.8307 | 1.2876 | 0.4887 | -0.324 | 0.627 |
| HMR_2365 | Fatty acid desaturation (even-chain) | 0.9797 | 0.2491 | 1.1027 | 0.3153 | 0.123 | -0.1637 | 0.565 |
| HMR_2288 | Fatty acid desaturation (odd-chain) | 0.4149 | 0.0961 | 0.4366 | 0.109 | 0.0217 | -0.0479 | 0.5191 |
| HMR_2289 | Fatty acid desaturation (odd-chain) | 0.7952 | 0.1153 | 0.8464 | 0.1267 | 0.0511 | -0.1039 | 0.5414 |
| HMR_0709 | Fatty acid transfer reactions | 1.1765 | 0.8656 | 1.1696 | 0.8543 | 0.0069 | 0.0053 | 0.4979 |
| HMR_2150 | Fatty acid transfer reactions | 2.2186 | 0.5957 | 2.2875 | 0.6358 | 0.0689 | -0.0621 | 0.5248 |
| HMR_2151 | Fatty acid transfer reactions | 30.9358 | 44.2356 | 32.0154 | 47.5674 | 1.0796 | -0.1127 | 0.5449 |
| HMR_2172 | Fatty acid transfer reactions | 0.5368 | 0.2121 | 0.5713 | 0.241 | 0.0345 | -0.0512 | 0.5204 |
| HMR_2177 | Fatty acid transfer reactions | 0.7719 | 0.3471 | 0.8004 | 0.3761 | 0.0284 | -0.0334 | 0.5133 |
| HMR_2182 | Fatty acid transfer reactions | 0.92 | 0.2204 | 0.9332 | 0.2271 | 0.0131 | -0.0196 | 0.5078 |
| HMR_2248 | Fatty acid transfer reactions | 0.574 | 0.1563 | 0.6055 | 0.1716 | 0.0315 | -0.055 | 0.5219 |
| HMR_2253 | Fatty acid transfer reactions | 0.5139 | 0.1586 | 0.5399 | 0.1755 | 0.026 | -0.0449 | 0.5179 |
| HMR_2258 | Fatty acid transfer reactions | 1.6839 | 0.4311 | 1.7442 | 0.4608 | 0.0603 | -0.0638 | 0.5255 |
| HMR_2472 | Omega-3 fatty acid metabolism | 7.7491 | 2.3152 | 8.4543 | 2.7196 | 0.7053 | -0.3143 | 0.6234 |
| HMR_2475 | Omega-3 fatty acid metabolism | 6.4983 | 1.8977 | 7.0443 | 2.1893 | 0.546 | -0.2701 | 0.6064 |
| HMR_2478 | Omega-3 fatty acid metabolism | 6.5022 | 1.8886 | 7.0476 | 2.1802 | 0.5453 | -0.2704 | 0.6066 |
| HMR_2480 | Omega-3 fatty acid metabolism | 6.5061 | 1.8795 | 7.0508 | 2.1711 | 0.5447 | -0.2707 | 0.6067 |

| | | | | | | | | |
|---|---|---|---|---|---|---|---|---|
| HMR_2482 | Omega-3 fatty acid metabolism | 6.5097 | 1.8704 | 7.0538 | 2.162 | 0.5441 | -0.2709 | 0.6068 |
| HMR_2484 | Omega-3 fatty acid metabolism | 3.5422 | 1.0636 | 3.8276 | 1.2219 | 0.2855 | -0.1888 | 0.5749 |
| HMR_2486 | Omega-3 fatty acid metabolism | 2.7661 | 0.7098 | 2.9613 | 0.8061 | 0.1952 | -0.1585 | 0.563 |
| HMR_2489 | Omega-3 fatty acid metabolism | 2.7731 | 0.7004 | 2.9675 | 0.7967 | 0.1944 | -0.1589 | 0.5631 |
| HMR_2491 | Omega-3 fatty acid metabolism | 2.7801 | 0.6911 | 2.9738 | 0.7874 | 0.1937 | -0.1593 | 0.5633 |
| HMR_2493 | Omega-3 fatty acid metabolism | 2.7868 | 0.6819 | 2.9797 | 0.7782 | 0.1929 | -0.1597 | 0.5634 |
| HMR_2495 | Omega-3 fatty acid metabolism | 0.6508 | 0.2569 | 0.7068 | 0.3055 | 0.056 | -0.0747 | 0.5298 |
| HMR_2497 | Omega-3 fatty acid metabolism | 1.3829 | 0.2958 | 1.4513 | 0.3197 | 0.0684 | -0.0872 | 0.5348 |
| HMR_2499 | Omega-3 fatty acid metabolism | 1.3915 | 0.2872 | 1.4591 | 0.311 | 0.0677 | -0.0875 | 0.5349 |
| HMR_2501 | Omega-3 fatty acid metabolism | 1.3997 | 0.2794 | 1.4667 | 0.3031 | 0.067 | -0.0877 | 0.535 |
| HMR_2503 | Omega-3 fatty acid metabolism | 1.4075 | 0.2721 | 1.4737 | 0.2957 | 0.0662 | -0.0879 | 0.535 |
| HMR_2505 | Omega-3 fatty acid metabolism | 0.8193 | 0.1498 | 0.8678 | 0.166 | 0.0485 | -0.0863 | 0.5344 |
| HMR_2510 | Omega-3 fatty acid metabolism | 0.7507 | 0.1253 | 0.7974 | 0.1413 | 0.0466 | -0.0903 | 0.536 |
| HMR_2511 | Omega-3 fatty acid metabolism | 0.7547 | 0.1238 | 0.8009 | 0.1397 | 0.0462 | -0.09 | 0.5359 |
| HMR_2512 | Omega-3 fatty acid metabolism | 0.7573 | 0.1243 | 0.8031 | 0.1403 | 0.0459 | -0.0892 | 0.5355 |
| HMR_2513 | Omega-3 fatty acid metabolism | 0.7583 | 0.1269 | 0.8039 | 0.143 | 0.0456 | -0.0878 | 0.535 |
| HMR_2514 | Omega-3 fatty acid metabolism | -0.7576 | 0.1316 | -0.803 | 0.1479 | 0.0454 | 0.0859 | 0.4658 |
| HMR_2515 | Omega-3 fatty acid metabolism | 1.8688 | 0.5339 | 2.0108 | 0.6071 | 0.1419 | -0.1329 | 0.5529 |
| HMR_2516 | Omega-3 fatty acid metabolism | 1.8744 | 0.5251 | 2.0157 | 0.5982 | 0.1413 | -0.1333 | 0.553 |
| HMR_2518 | Omega-3 fatty acid metabolism | 1.8798 | 0.5171 | 2.0205 | 0.5902 | 0.1407 | -0.1337 | 0.5532 |
| HMR_2520 | Omega-3 fatty acid metabolism | 1.8847 | 0.5099 | 2.0248 | 0.5828 | 0.1401 | -0.134 | 0.5533 |
| HMR_2522 | Omega-3 fatty acid metabolism | 1.0787 | 0.3619 | 1.1346 | 0.4024 | 0.0559 | -0.064 | 0.5255 |
| HMR_2524 | Omega-3 fatty acid metabolism | 1.0832 | 0.3594 | 1.1386 | 0.3998 | 0.0554 | -0.0636 | 0.5254 |
| HMR_2526 | Omega-3 fatty acid metabolism | 0.7876 | 0.1603 | 0.807 | 0.1689 | 0.0194 | -0.0338 | 0.5135 |
| HMR_2528 | Omega-3 fatty acid metabolism | 0.7871 | 0.1562 | 0.8061 | 0.1647 | 0.019 | -0.0335 | 0.5134 |
| HMR_2530 | Omega-3 fatty acid metabolism | 1.7365 | 0.4328 | 1.9356 | 0.5289 | 0.1991 | -0.203 | 0.5804 |
| HMR_2533 | Omega-3 fatty acid metabolism | 1.7389 | 0.4238 | 1.9375 | 0.5199 | 0.1986 | -0.2044 | 0.581 |
| HMR_2535 | Omega-3 fatty acid metabolism | 1.7412 | 0.4154 | 1.9393 | 0.5114 | 0.1981 | -0.2058 | 0.5815 |
| HMR_2537 | Omega-3 fatty acid metabolism | 1.743 | 0.4076 | 1.9406 | 0.5035 | 0.1976 | -0.2071 | 0.582 |
| HMR_2540 | Omega-3 fatty acid metabolism | 0.8751 | 0.1655 | 0.9639 | 0.1987 | 0.0889 | -0.1473 | 0.5585 |
| HMR_2541 | Omega-3 fatty acid metabolism | 0.8776 | 0.1589 | 0.9661 | 0.192 | 0.0885 | -0.1495 | 0.5594 |
| HMR_2542 | Omega-3 fatty acid metabolism | 0.8788 | 0.1542 | 0.9671 | 0.1872 | 0.0884 | -0.1512 | 0.5601 |
| HMR_2543 | Omega-3 fatty acid metabolism | 0.8786 | 0.1513 | 0.9668 | 0.1842 | 0.0882 | -0.1523 | 0.5605 |
| HMR_2368 | Omega-6 fatty acid metabolism | 9.2974 | 2.9262 | 9.8464 | 3.2387 | 0.549 | -0.2211 | 0.5875 |
| HMR_2371 | Omega-6 fatty acid metabolism | 14.6856 | 7.6527 | 15.6812 | 8.851 | 0.9956 | -0.2451 | 0.5968 |
| HMR_2374 | Omega-6 fatty acid metabolism | 14.6895 | 7.6439 | 15.6845 | 8.8422 | 0.995 | -0.245 | 0.5968 |
| HMR_2376 | Omega-6 fatty acid metabolism | 14.6934 | 7.6351 | 15.6878 | 8.8334 | 0.9943 | -0.245 | 0.5968 |
| HMR_2378 | Omega-6 fatty acid metabolism | 14.6971 | 7.6262 | 15.6907 | 8.8245 | 0.9937 | -0.245 | 0.5968 |
| HMR_2380 | Omega-6 fatty acid metabolism | 10.0011 | 3.7818 | 10.5872 | 4.2612 | 0.5861 | -0.2067 | 0.5819 |
| HMR_2383 | Omega-6 fatty acid metabolism | 2.2648 | 0.6955 | 2.4505 | 0.8058 | 0.1856 | -0.1515 | 0.5602 |
| HMR_2387 | Omega-6 fatty acid metabolism | 2.2718 | 0.6865 | 2.4566 | 0.7967 | 0.1849 | -0.1518 | 0.5603 |
| HMR_2389 | Omega-6 fatty acid metabolism | 2.2785 | 0.678 | 2.4627 | 0.7881 | 0.1842 | -0.1521 | 0.5604 |
| HMR_2391 | Omega-6 fatty acid metabolism | 2.285 | 0.6699 | 2.4684 | 0.78 | 0.1834 | -0.1523 | 0.5605 |
| HMR_2393 | Omega-6 fatty acid metabolism | 0.5963 | 0.239 | 0.6585 | 0.2926 | 0.0623 | -0.0854 | 0.534 |

| | | | | | | | | |
|---|---|---|---|---|---|---|---|---|
| HMR_2395 | Omega-6 fatty acid metabolism | 0.9858 | 0.3389 | 1.0406 | 0.3776 | 0.0548 | -0.0647 | 0.5258 |
| HMR_2397 | Omega-6 fatty acid metabolism | 0.9911 | 0.3365 | 1.0452 | 0.375 | 0.0542 | -0.0642 | 0.5256 |
| HMR_2399 | Omega-6 fatty acid metabolism | 1.2888 | 0.2248 | 1.3786 | 0.2552 | 0.0899 | -0.1297 | 0.5516 |
| HMR_2401 | Omega-6 fatty acid metabolism | 1.2903 | 0.219 | 1.3798 | 0.2495 | 0.0894 | -0.1307 | 0.552 |
| HMR_2403 | Omega-6 fatty acid metabolism | 0.9932 | 0.1882 | 1.0777 | 0.2189 | 0.0845 | -0.1324 | 0.5527 |
| HMR_2408 | Omega-6 fatty acid metabolism | 0.9015 | 0.1686 | 0.9852 | 0.2 | 0.0838 | -0.138 | 0.5549 |
| HMR_2409 | Omega-6 fatty acid metabolism | 0.9012 | 0.1674 | 0.9848 | 0.199 | 0.0836 | -0.1381 | 0.5549 |
| HMR_2410 | Omega-6 fatty acid metabolism | 0.9 | 0.1681 | 0.9836 | 0.1998 | 0.0836 | -0.1378 | 0.5548 |
| HMR_2411 | Omega-6 fatty acid metabolism | 0.8975 | 0.1707 | 0.9812 | 0.2026 | 0.0837 | -0.137 | 0.5545 |
| HMR_2412 | Omega-6 fatty acid metabolism | -0.8938 | 0.1753 | -0.9777 | 0.2074 | 0.0839 | 0.1356 | 0.4461 |
| HMR_2415 | Omega-6 fatty acid metabolism | 0.8044 | 0.1474 | 0.8816 | 0.1741 | 0.0772 | -0.1361 | 0.5541 |
| HMR_2417 | Omega-6 fatty acid metabolism | 0.8086 | 0.1402 | 0.8853 | 0.1668 | 0.0767 | -0.1385 | 0.5551 |
| HMR_2419 | Omega-6 fatty acid metabolism | 0.8112 | 0.1351 | 0.8876 | 0.1617 | 0.0764 | -0.1403 | 0.5558 |
| HMR_2421 | Omega-6 fatty acid metabolism | 0.8122 | 0.1319 | 0.8885 | 0.1584 | 0.0762 | -0.1415 | 0.5563 |
| HMR_2423 | Omega-6 fatty acid metabolism | 1.7626 | 0.405 | 1.9659 | 0.4941 | 0.2032 | -0.2143 | 0.5849 |
| HMR_2426 | Omega-6 fatty acid metabolism | 1.7652 | 0.3958 | 1.9679 | 0.4848 | 0.2027 | -0.216 | 0.5855 |
| HMR_2428 | Omega-6 fatty acid metabolism | 1.7676 | 0.387 | 1.9698 | 0.476 | 0.2022 | -0.2177 | 0.5862 |
| HMR_2430 | Omega-6 fatty acid metabolism | 1.7696 | 0.3786 | 1.9713 | 0.4675 | 0.2017 | -0.2193 | 0.5868 |
| HMR_2433 | Omega-6 fatty acid metabolism | 0.8318 | 0.1523 | 0.9167 | 0.1826 | 0.0849 | -0.1467 | 0.5583 |
| HMR_2434 | Omega-6 fatty acid metabolism | 0.8343 | 0.1455 | 0.9189 | 0.1757 | 0.0846 | -0.1492 | 0.5593 |
| HMR_2435 | Omega-6 fatty acid metabolism | 0.8355 | 0.1407 | 0.92 | 0.1709 | 0.0844 | -0.1512 | 0.5601 |
| HMR_2436 | Omega-6 fatty acid metabolism | 0.8353 | 0.1377 | 0.9196 | 0.1678 | 0.0843 | -0.1525 | 0.5606 |
| HMR_3444 | Omega-6 fatty acid metabolism | 2.5649 | 1.6725 | 2.7697 | 1.9478 | 0.2049 | -0.1077 | 0.5429 |
| HMR_3446 | Omega-6 fatty acid metabolism | 2.5681 | 1.6748 | 2.7723 | 1.9502 | 0.2042 | -0.1072 | 0.5427 |
| HMR_3447 | Omega-6 fatty acid metabolism | 2.5707 | 1.6807 | 2.7743 | 1.9561 | 0.2036 | -0.1067 | 0.5425 |
| HMR_3448 | Omega-6 fatty acid metabolism | 3.5704 | 3.7917 | 3.9023 | 4.5209 | 0.3319 | -0.1151 | 0.5458 |
| HMR_3449 | Omega-6 fatty acid metabolism | 6.1433 | 5.2318 | 6.6784 | 6.1937 | 0.5351 | -0.1583 | 0.5629 |
| HMR_3450 | Omega-6 fatty acid metabolism | 6.145 | 5.2404 | 6.6796 | 6.2022 | 0.5346 | -0.158 | 0.5628 |
| HMR_3451 | Omega-6 fatty acid metabolism | 6.1467 | 5.2495 | 6.6807 | 6.2114 | 0.534 | -0.1577 | 0.5627 |
| HMR_3452 | Omega-6 fatty acid metabolism | 0.9774 | 0.2232 | 1.1035 | 0.2814 | 0.1262 | -0.1776 | 0.5705 |
| HMR_3453 | Omega-6 fatty acid metabolism | 0.9784 | 0.2162 | 1.104 | 0.2744 | 0.1256 | -0.1794 | 0.5712 |
| HMR_3454 | Omega-6 fatty acid metabolism | 0.9779 | 0.2118 | 1.1031 | 0.2699 | 0.1253 | -0.1805 | 0.5716 |
| HMR_3455 | Omega-6 fatty acid metabolism | -0.9761 | 0.2097 | -1.1011 | 0.2678 | 0.125 | 0.1809 | 0.4282 |
| HMR_3456 | Omega-6 fatty acid metabolism | 0.9731 | 0.2098 | 1.0979 | 0.268 | 0.1249 | -0.1806 | 0.5717 |
| HMR_3457 | Omega-6 fatty acid metabolism | 0.9692 | 0.2123 | 1.094 | 0.2707 | 0.1248 | -0.1796 | 0.5713 |
| HMR_3458 | Omega-6 fatty acid metabolism | 0.964 | 0.2173 | 1.0889 | 0.2758 | 0.125 | -0.1779 | 0.5706 |
| HMR_3459 | Omega-6 fatty acid metabolism | 1.0881 | 0.2277 | 1.1051 | 0.2337 | 0.017 | -0.025 | 0.51 |
| HMR_3461 | Omega-6 fatty acid metabolism | 1.0952 | 0.2217 | 1.1116 | 0.2276 | 0.0164 | -0.0245 | 0.5098 |
| HMR_3462 | Omega-6 fatty acid metabolism | 1.1021 | 0.2173 | 1.1181 | 0.2232 | 0.016 | -0.0241 | 0.5096 |
| HMR_3463 | Omega-6 fatty acid metabolism | 1.1083 | 0.2143 | 1.1239 | 0.2201 | 0.0156 | -0.0237 | 0.5094 |
| HMR_3464 | Omega-6 fatty acid metabolism | 1.1142 | 0.2124 | 1.1294 | 0.2181 | 0.0152 | -0.0232 | 0.5093 |
| HMR_3465 | Omega-6 fatty acid metabolism | 1.1195 | 0.2115 | 1.1343 | 0.2172 | 0.0149 | -0.0227 | 0.5091 |
| HMR_3466 | Omega-6 fatty acid metabolism | 1.1242 | 0.2117 | 1.1387 | 0.2173 | 0.0145 | -0.0221 | 0.5088 |
| HMR_3467 | Omega-6 fatty acid metabolism | 1.129 | 0.2128 | 1.143 | 0.2184 | 0.0141 | -0.0214 | 0.5085 |

| | | | | | | | | |
|---|---|---|---|---|---|---|---|---|
| HMR_3468 | Omega-6 fatty acid metabolism | 1.1334 | 0.2149 | 1.1471 | 0.2205 | 0.0137 | -0.0208 | 0.5083 |
| HMR_3469 | Omega-6 fatty acid metabolism | -1.1374 | 0.2179 | -1.1507 | 0.2235 | 0.0133 | 0.02 | 0.492 |
| HMR_3470 | Omega-6 fatty acid metabolism | 1.1409 | 0.222 | 1.1537 | 0.2276 | 0.0129 | -0.0192 | 0.5077 |
| HMR_3471 | Omega-6 fatty acid metabolism | 1.1443 | 0.2272 | 1.1567 | 0.2328 | 0.0124 | -0.0183 | 0.5073 |
| HMR_3472 | Omega-6 fatty acid metabolism | 1.1477 | 0.2339 | 1.1597 | 0.2395 | 0.0121 | -0.0176 | 0.507 |
| HMR_0958 | Arachidonic acid metabolism | 2.9077 | 0.5285 | 3.1368 | 0.5953 | 0.2291 | -0.2161 | 0.5856 |
| HMR_0960 | Arachidonic acid metabolism | 2.9138 | 0.5186 | 3.1427 | 0.5854 | 0.2289 | -0.2178 | 0.5862 |
| HMR_0973 | Arachidonic acid metabolism | 2.9196 | 0.5087 | 3.1482 | 0.5755 | 0.2286 | -0.2196 | 0.5869 |
| HMR_0976 | Arachidonic acid metabolism | 2.9255 | 0.4988 | 3.1538 | 0.5656 | 0.2283 | -0.2213 | 0.5876 |
| HMR_0979 | Arachidonic acid metabolism | 1.4657 | 0.246 | 1.5796 | 0.2796 | 0.1139 | -0.1571 | 0.5624 |
| HMR_0989 | Arachidonic acid metabolism | 385.6753 | 158501.3985 | 382.12 | 160072.799 | 3.5553 | 0.0063 | 0.4975 |
| HMR_0990 | Arachidonic acid metabolism | -385.6753 | 158501.3985 | -382.12 | 160072.7992 | 3.5554 | -0.0063 | 0.5025 |
| HMR_1131 | Leukotriene metabolism | -8.5321 | 32.4767 | -8.9753 | 35.9181 | 0.4432 | 0.0536 | 0.4786 |
| HMR_1134 | Leukotriene metabolism | 8.6497 | 25.8454 | 9.0999 | 28.5948 | 0.4502 | -0.061 | 0.5243 |
| HMR_1136 | Leukotriene metabolism | -0.1173 | 61.6337 | -0.1242 | 68.2182 | 0.007 | 0.0006 | 0.4998 |
| HMR_1138 | Leukotriene metabolism | 8.5311 | 32.476 | 8.9744 | 35.9174 | 0.4433 | -0.0536 | 0.5214 |
| HMR_1140 | Leukotriene metabolism | 8.6484 | 25.8447 | 9.0986 | 28.594 | 0.4502 | -0.061 | 0.5243 |
| HMR_1142 | Leukotriene metabolism | -8.6494 | 25.8439 | -9.0996 | 28.5932 | 0.4502 | 0.061 | 0.4757 |
| HMR_1150 | Leukotriene metabolism | 0.7511 | 0.1257 | 0.8168 | 0.1474 | 0.0657 | -0.1258 | 0.55 |
| HMR_1152 | Leukotriene metabolism | 0.7277 | 0.1133 | 0.7751 | 0.1273 | 0.0474 | -0.0966 | 0.5385 |
| HMR_1158 | Leukotriene metabolism | -0.7531 | 0.1212 | -0.8188 | 0.1427 | 0.0657 | 0.128 | 0.4491 |
| HMR_1160 | Leukotriene metabolism | -0.7304 | 0.1083 | -0.7778 | 0.1222 | 0.0474 | 0.0988 | 0.4607 |
| HMR_1161 | Leukotriene metabolism | 0.7538 | 0.1187 | 0.8192 | 0.1402 | 0.0654 | -0.1286 | 0.5511 |
| HMR_1162 | Leukotriene metabolism | 0.7321 | 0.1052 | 0.7791 | 0.119 | 0.0471 | -0.0994 | 0.5396 |
| HMR_1163 | Leukotriene metabolism | 0.7536 | 0.1181 | 0.8188 | 0.1396 | 0.0652 | -0.1284 | 0.5511 |
| HMR_1164 | Leukotriene metabolism | 0.733 | 0.1035 | 0.7797 | 0.1172 | 0.0467 | -0.0995 | 0.5396 |
| HMR_1165 | Leukotriene metabolism | 0.7519 | 0.1192 | 0.8169 | 0.1407 | 0.065 | -0.1274 | 0.5507 |
| HMR_1166 | Leukotriene metabolism | 0.7329 | 0.1031 | 0.7794 | 0.1168 | 0.0465 | -0.0992 | 0.5395 |
| HMR_1167 | Leukotriene metabolism | 0.7487 | 0.1223 | 0.8136 | 0.1439 | 0.0649 | -0.1258 | 0.5501 |
| HMR_1168 | Leukotriene metabolism | 0.7317 | 0.104 | 0.778 | 0.1178 | 0.0463 | -0.0983 | 0.5392 |
| HMR_1170 | Leukotriene metabolism | 0.7438 | 0.1274 | 0.8088 | 0.1491 | 0.065 | -0.1235 | 0.5492 |
| HMR_1171 | Leukotriene metabolism | 0.7292 | 0.1063 | 0.7754 | 0.1202 | 0.0462 | -0.0971 | 0.5387 |
| HMR_1186 | Leukotriene metabolism | -1.4657 | 0.246 | -1.5796 | 0.2796 | 0.1139 | 0.1571 | 0.4376 |
| HMR_1194 | Leukotriene metabolism | 0.7511 | 0.1257 | 0.8168 | 0.1474 | 0.0657 | -0.1258 | 0.55 |
| HMR_1196 | Leukotriene metabolism | 0.7277 | 0.1133 | 0.7751 | 0.1273 | 0.0474 | -0.0966 | 0.5385 |
| HMR_1201 | Leukotriene metabolism | -0.7531 | 0.1212 | -0.8188 | 0.1427 | 0.0657 | 0.128 | 0.4491 |
| HMR_1203 | Leukotriene metabolism | -0.7304 | 0.1083 | -0.7778 | 0.1222 | 0.0474 | 0.0988 | 0.4607 |
| HMR_1204 | Leukotriene metabolism | 0.7538 | 0.1187 | 0.8192 | 0.1402 | 0.0654 | -0.1286 | 0.5511 |
| HMR_1205 | Leukotriene metabolism | 0.7321 | 0.1052 | 0.7791 | 0.119 | 0.0471 | -0.0994 | 0.5396 |
| HMR_1206 | Leukotriene metabolism | 0.7536 | 0.1181 | 0.8188 | 0.1396 | 0.0652 | -0.1284 | 0.5511 |
| HMR_1207 | Leukotriene metabolism | 0.733 | 0.1035 | 0.7797 | 0.1172 | 0.0467 | -0.0995 | 0.5396 |
| HMR_1208 | Leukotriene metabolism | 0.7519 | 0.1192 | 0.8169 | 0.1407 | 0.065 | -0.1274 | 0.5507 |
| HMR_1209 | Leukotriene metabolism | 0.7329 | 0.1031 | 0.7794 | 0.1168 | 0.0465 | -0.0992 | 0.5395 |
| HMR_1210 | Leukotriene metabolism | 0.7487 | 0.1223 | 0.8136 | 0.1439 | 0.0649 | -0.1258 | 0.5501 |

| | | | | | | | | |
|---|---|---|---|---|---|---|---|---|
| HMR_1211 | Leukotriene metabolism | 0.7317 | 0.104 | 0.778 | 0.1178 | 0.0463 | -0.0983 | 0.5392 |
| HMR_1212 | Leukotriene metabolism | 0.7438 | 0.1274 | 0.8088 | 0.1491 | 0.065 | -0.1235 | 0.5492 |
| HMR_1213 | Leukotriene metabolism | 0.7292 | 0.1063 | 0.7754 | 0.1202 | 0.0462 | -0.0971 | 0.5387 |
| HMR_1231 | Leukotriene metabolism | 173.3315 | 10623.7403 | 169.3971 | 10124.2926 | 3.9344 | 0.0273 | 0.4891 |
| HMR_1232 | Leukotriene metabolism | 491.6876 | 407.6802 | 505.9255 | 435.6059 | 14.2378 | -0.4903 | 0.688 |
| HMR_1233 | Leukotriene metabolism | 185.3536 | 8138.5588 | 178.5188 | 7577.4078 | 6.8348 | 0.0545 | 0.4783 |
| HMR_1263 | Leukotriene metabolism | 677.045 | 8365.6574 | 684.4435 | 7829.2506 | 7.3984 | -0.0581 | 0.5232 |
| HMR_1264 | Leukotriene metabolism | -491.6876 | 407.6735 | -505.9256 | 435.599 | 14.238 | 0.4903 | 0.312 |
| HMR_1266 | Leukotriene metabolism | -185.3535 | 8138.5585 | -178.5189 | 7577.4075 | 6.8347 | -0.0545 | 0.5217 |
| HMR_1280 | Leukotriene metabolism | 173.3315 | 10623.7414 | 169.397 | 10124.2937 | 3.9345 | 0.0273 | 0.4891 |
| HMR_1281 | Leukotriene metabolism | 491.6877 | 407.6853 | 505.9254 | 435.6111 | 14.2377 | -0.4903 | 0.688 |
| HMR_1282 | Leukotriene metabolism | 185.3542 | 8138.5602 | 178.5194 | 7577.4092 | 6.8348 | 0.0545 | 0.4783 |
| HMR_1285 | Leukotriene metabolism | -850.3883 | 11602.5453 | -853.8383 | 11034.9389 | 3.45 | 0.0229 | 0.4909 |
| HMR_8559 | Eicosanoid metabolism | 393.5255 | 114437.1355 | 392.1597 | 114879.3615 | 1.3658 | 0.0029 | 0.4989 |
| HMR_0154 | Acyl-CoA hydrolysis | 2.0093 | 4.1514 | 2.0338 | 4.228 | 0.0245 | -0.0085 | 0.5034 |
| HMR_0189 | Acyl-CoA hydrolysis | 1.9876 | 4.8133 | 2.0107 | 4.9457 | 0.0231 | -0.0074 | 0.503 |
| HMR_0193 | Acyl-CoA hydrolysis | 1.8751 | 2.172 | 1.8971 | 2.2417 | 0.0219 | -0.0104 | 0.5042 |
| HMR_0197 | Acyl-CoA hydrolysis | 2.0093 | 4.1512 | 2.0338 | 4.2279 | 0.0245 | -0.0085 | 0.5034 |
| HMR_0201 | Acyl-CoA hydrolysis | 1.9173 | 3.1363 | 1.9378 | 3.1883 | 0.0205 | -0.0081 | 0.5032 |
| HMR_0206 | Acyl-CoA hydrolysis | 1.8943 | 1.9727 | 1.9176 | 2.0181 | 0.0233 | -0.0117 | 0.5047 |
| HMR_0210 | Acyl-CoA hydrolysis | 1.9821 | 1.9069 | 2.0054 | 1.9523 | 0.0234 | -0.0119 | 0.5047 |
| HMR_0214 | Acyl-CoA hydrolysis | 1.9759 | 3.6925 | 1.9993 | 3.7796 | 0.0234 | -0.0086 | 0.5034 |
| HMR_0223 | Acyl-CoA hydrolysis | 2.0093 | 4.1512 | 2.0338 | 4.2279 | 0.0245 | -0.0085 | 0.5034 |
| HMR_0230 | Acyl-CoA hydrolysis | 2.0093 | 4.1513 | 2.0338 | 4.2279 | 0.0245 | -0.0085 | 0.5034 |
| HMR_0234 | Acyl-CoA hydrolysis | 1.9899 | 4.739 | 2.0107 | 4.9383 | 0.0208 | -0.0067 | 0.5027 |
| HMR_0238 | Acyl-CoA hydrolysis | 1.9219 | 3.1311 | 1.9494 | 3.2535 | 0.0275 | -0.0109 | 0.5043 |
| HMR_0242 | Acyl-CoA hydrolysis | 1.8799 | 2.0284 | 1.8969 | 2.0986 | 0.0171 | -0.0084 | 0.5034 |
| HMR_0246 | Acyl-CoA hydrolysis | 1.9779 | 3.7146 | 1.9996 | 3.7863 | 0.0218 | -0.0079 | 0.5032 |
| HMR_0250 | Acyl-CoA hydrolysis | 2.0093 | 4.1512 | 2.0338 | 4.2279 | 0.0245 | -0.0085 | 0.5034 |
| HMR_0256 | Acyl-CoA hydrolysis | 1.9525 | 3.4707 | 1.9735 | 3.5323 | 0.021 | -0.0079 | 0.5032 |
| HMR_0260 | Acyl-CoA hydrolysis | 2.0093 | 4.1512 | 2.0338 | 4.2279 | 0.0245 | -0.0085 | 0.5034 |
| HMR_0264 | Acyl-CoA hydrolysis | 2.0093 | 4.1512 | 2.0338 | 4.2279 | 0.0245 | -0.0085 | 0.5034 |
| HMR_0268 | Acyl-CoA hydrolysis | 1.9821 | 1.9069 | 2.006 | 1.9522 | 0.0239 | -0.0122 | 0.5049 |
| HMR_0272 | Acyl-CoA hydrolysis | 0.6054 | 0.1684 | 0.66 | 0.1993 | 0.0546 | -0.09 | 0.5359 |
| HMR_0276 | Acyl-CoA hydrolysis | 1.9849 | 1.9066 | 2.0088 | 1.9519 | 0.0239 | -0.0122 | 0.5048 |
| HMR_0280 | Acyl-CoA hydrolysis | 1.9748 | 1.9085 | 1.9984 | 1.9539 | 0.0236 | -0.012 | 0.5048 |
| HMR_0284 | Acyl-CoA hydrolysis | 1.8772 | 2.0108 | 1.8996 | 2.0584 | 0.0224 | -0.0111 | 0.5044 |
| HMR_0290 | Acyl-CoA hydrolysis | 1.9342 | 1.9272 | 1.9584 | 1.9721 | 0.0242 | -0.0122 | 0.5049 |
| HMR_0294 | Acyl-CoA hydrolysis | 1.9705 | 5.2811 | 1.9872 | 5.5886 | 0.0166 | -0.005 | 0.502 |
| HMR_0298 | Acyl-CoA hydrolysis | 1.8487 | 2.3131 | 1.8705 | 2.3564 | 0.0218 | -0.0101 | 0.504 |
| HMR_0302 | Acyl-CoA hydrolysis | 1.9372 | 1.9251 | 1.9612 | 1.9702 | 0.024 | -0.0121 | 0.5048 |
| HMR_0306 | Acyl-CoA hydrolysis | 1.9824 | 1.907 | 2.0044 | 1.9528 | 0.022 | -0.0112 | 0.5045 |
| HMR_0310 | Acyl-CoA hydrolysis | 1.9753 | 1.9084 | 1.9989 | 1.9538 | 0.0236 | -0.012 | 0.5048 |
| HMR_0314 | Acyl-CoA hydrolysis | 1.9536 | 1.9162 | 1.9753 | 1.9626 | 0.0217 | -0.011 | 0.5044 |

| | | | | | | | | |
|---|---|---|---|---|---|---|---|---|
| HMR_0320 | Acyl-CoA hydrolysis | 1.9435 | 1.9212 | 1.9673 | 1.9665 | 0.0238 | -0.0121 | 0.5048 |
| HMR_0324 | Acyl-CoA hydrolysis | 1.9698 | 1.9093 | 1.9935 | 1.9547 | 0.0237 | -0.012 | 0.5048 |
| HMR_0328 | Acyl-CoA hydrolysis | 1.992 | 1.9042 | 2.0149 | 1.9497 | 0.0229 | -0.0117 | 0.5047 |
| HMR_0332 | Acyl-CoA hydrolysis | 1.9513 | 1.918 | 1.9711 | 1.9656 | 0.0198 | -0.01 | 0.504 |
| HMR_0338 | Acyl-CoA hydrolysis | 1.9847 | 1.9066 | 2.0072 | 1.9522 | 0.0225 | -0.0115 | 0.5046 |
| HMR_0342 | Acyl-CoA hydrolysis | 1.975 | 1.9086 | 1.9969 | 1.9544 | 0.022 | -0.0112 | 0.5045 |
| HMR_0346 | Acyl-CoA hydrolysis | 1.9897 | 1.9054 | 2.0134 | 1.9507 | 0.0237 | -0.0121 | 0.5048 |
| HMR_0350 | Acyl-CoA hydrolysis | 2.0093 | 4.1516 | 2.0338 | 4.228 | 0.0245 | -0.0085 | 0.5034 |
| HMR_0354 | Acyl-CoA hydrolysis | 1.9712 | 1.9099 | 1.995 | 1.9552 | 0.0238 | -0.0121 | 0.5048 |
| HMR_0358 | Acyl-CoA hydrolysis | 1.8495 | 2.2224 | 1.8717 | 2.2664 | 0.0222 | -0.0105 | 0.5042 |
| HMR_0362 | Acyl-CoA hydrolysis | 2.0092 | 4.1506 | 2.0336 | 4.2272 | 0.0245 | -0.0085 | 0.5034 |
| HMR_0366 | Acyl-CoA hydrolysis | 1.8489 | 2.2567 | 1.8708 | 2.3099 | 0.022 | -0.0103 | 0.5041 |
| HMR_0370 | Acyl-CoA hydrolysis | 1.9821 | 1.907 | 2.0061 | 1.9522 | 0.0241 | -0.0122 | 0.5049 |
| HMR_0374 | Acyl-CoA hydrolysis | 1.9824 | 1.9067 | 2.0064 | 1.952 | 0.024 | -0.0122 | 0.5049 |
| HMR_0378 | Acyl-CoA hydrolysis | 1.897 | 2.9259 | 1.9201 | 3.0012 | 0.0231 | -0.0095 | 0.5038 |
| HMR_0382 | Acyl-CoA hydrolysis | 1.9015 | 1.9631 | 1.9218 | 2.0137 | 0.0203 | -0.0102 | 0.5041 |
| HMR_0386 | Acyl-CoA hydrolysis | 1.9915 | 1.9051 | 2.015 | 1.9506 | 0.0235 | -0.012 | 0.5048 |
| HMR_0390 | Acyl-CoA hydrolysis | 1.9833 | 1.9068 | 2.0073 | 1.9521 | 0.024 | -0.0122 | 0.5049 |
| HMR_0394 | Acyl-CoA hydrolysis | 1.991 | 1.9052 | 2.0148 | 1.9506 | 0.0238 | -0.0121 | 0.5048 |
| HMR_0398 | Acyl-CoA hydrolysis | 2.0093 | 4.1512 | 2.0338 | 4.2279 | 0.0245 | -0.0085 | 0.5034 |
| HMR_0402 | Acyl-CoA hydrolysis | 1.8512 | 2.2267 | 1.8732 | 2.3027 | 0.022 | -0.0103 | 0.5041 |
| HMR_0406 | Acyl-CoA hydrolysis | 1.8611 | 2.5325 | 1.8828 | 2.5884 | 0.0218 | -0.0096 | 0.5038 |
| HMR_0410 | Acyl-CoA hydrolysis | 2.0093 | 4.1512 | 2.0338 | 4.2279 | 0.0245 | -0.0085 | 0.5034 |
| HMR_0414 | Acyl-CoA hydrolysis | 1.8607 | 2.0777 | 1.8825 | 2.1293 | 0.0218 | -0.0106 | 0.5042 |
| HMR_0418 | Acyl-CoA hydrolysis | 1.9874 | 1.9059 | 2.0112 | 1.9513 | 0.0238 | -0.0121 | 0.5048 |
| HMR_0422 | Acyl-CoA hydrolysis | 1.9871 | 1.9059 | 2.0109 | 1.9513 | 0.0238 | -0.0121 | 0.5048 |
| HMR_0426 | Acyl-CoA hydrolysis | 1.917 | 1.9434 | 1.9405 | 1.9889 | 0.0234 | -0.0118 | 0.5047 |
| HMR_0430 | Acyl-CoA hydrolysis | 1.8772 | 2.7378 | 1.8985 | 2.7918 | 0.0213 | -0.0091 | 0.5036 |
| HMR_0434 | Acyl-CoA hydrolysis | 1.9991 | 1.9043 | 2.0224 | 1.9497 | 0.0233 | -0.0119 | 0.5047 |
| HMR_0438 | Acyl-CoA hydrolysis | 1.9874 | 1.9057 | 2.011 | 1.9511 | 0.0236 | -0.012 | 0.5048 |
| HMR_3477 | Acyl-CoA hydrolysis | 0.8252 | 0.1184 | 1.0606 | 0.1819 | 0.2354 | -0.4295 | 0.6662 |
| HMR_2591 | Carnitine shuttle (cytosolic) | 346.9992 | 162157.1957 | 443.498 | 139135.5041 | 96.4988 | -0.1758 | 0.5698 |
| HMR_2594 | Carnitine shuttle (cytosolic) | 170.0958 | 229150.6339 | 198.4414 | 223132.3396 | 28.3456 | -0.0421 | 0.5168 |
| HMR_2599 | Carnitine shuttle (cytosolic) | 124.7792 | 138.2814 | 133.3924 | 149.5942 | 8.6131 | -0.5076 | 0.6941 |
| HMR_2602 | Carnitine shuttle (cytosolic) | -0.3219 | 0.2506 | -0.3498 | 0.2821 | 0.0279 | 0.0383 | 0.4847 |
| HMR_2605 | Carnitine shuttle (cytosolic) | -0.4963 | 0.1571 | -0.5298 | 0.1719 | 0.0335 | 0.0584 | 0.4767 |
| HMR_2608 | Carnitine shuttle (cytosolic) | 1.2871 | 1.0817 | 1.5622 | 1.3432 | 0.2751 | -0.1767 | 0.5701 |
| HMR_2611 | Carnitine shuttle (cytosolic) | 0.9333 | 0.2365 | 1.0411 | 0.2947 | 0.1078 | -0.1479 | 0.5588 |
| HMR_2614 | Carnitine shuttle (cytosolic) | 0.6553 | 0.3343 | 0.7518 | 0.402 | 0.0966 | -0.1125 | 0.5448 |
| HMR_2620 | Carnitine shuttle (cytosolic) | 0.7399 | 0.2899 | 0.8428 | 0.3557 | 0.1029 | -0.128 | 0.5509 |
| HMR_2623 | Carnitine shuttle (cytosolic) | -0.4221 | 0.1718 | -0.4533 | 0.1891 | 0.0312 | 0.052 | 0.4793 |
| HMR_2626 | Carnitine shuttle (cytosolic) | -0.0456 | 5.469 | -0.1255 | 6.5741 | 0.08 | 0.023 | 0.4908 |
| HMR_2633 | Carnitine shuttle (cytosolic) | 0.1024 | 0.6373 | 0.1744 | 0.7725 | 0.072 | -0.0606 | 0.5242 |
| HMR_2636 | Carnitine shuttle (cytosolic) | -5.4447 | 1.8871 | -5.9211 | 2.3055 | 0.4764 | 0.2326 | 0.408 |

| | | | | | | | | |
|---|---|---|---|---|---|---|---|---|
| HMR_2642 | Carnitine shuttle (cytosolic) | 3.0132 | 3.1974 | 3.3186 | 3.7947 | 0.3054 | -0.1155 | 0.546 |
| HMR_2648 | Carnitine shuttle (cytosolic) | 0.7538 | 0.1108 | 0.8217 | 0.1285 | 0.0679 | -0.1389 | 0.5552 |
| HMR_2651 | Carnitine shuttle (cytosolic) | 0.4923 | 0.0712 | 0.5201 | 0.0792 | 0.0278 | -0.0716 | 0.5285 |
| HMR_2654 | Carnitine shuttle (cytosolic) | -0.2924 | 7.4165 | -0.3084 | 8.8189 | 0.0159 | 0.0039 | 0.4984 |
| HMR_2657 | Carnitine shuttle (cytosolic) | 1.0375 | 0.2468 | 1.1568 | 0.3063 | 0.1194 | -0.1605 | 0.5638 |
| HMR_2660 | Carnitine shuttle (cytosolic) | 2.9045 | 2.958 | 3.1543 | 3.5393 | 0.2498 | -0.098 | 0.539 |
| HMR_2663 | Carnitine shuttle (cytosolic) | -0.0842 | 4.8196 | -0.0173 | 5.7063 | 0.0669 | -0.0206 | 0.5082 |
| HMR_2666 | Carnitine shuttle (cytosolic) | 0.7306 | 0.1429 | 0.8165 | 0.1783 | 0.0859 | -0.1516 | 0.5602 |
| HMR_2669 | Carnitine shuttle (cytosolic) | 0.7178 | 0.1529 | 0.8056 | 0.192 | 0.0878 | -0.1495 | 0.5594 |
| HMR_2672 | Carnitine shuttle (cytosolic) | 1.2518 | 0.455 | 1.4715 | 0.6285 | 0.2197 | -0.2111 | 0.5836 |
| HMR_2675 | Carnitine shuttle (cytosolic) | 0.0968 | 5.7617 | 0.1063 | 6.8799 | 0.0095 | -0.0027 | 0.5011 |
| HMR_2678 | Carnitine shuttle (cytosolic) | 0.091 | 6.2198 | 0.1002 | 7.3997 | 0.0092 | -0.0025 | 0.501 |
| HMR_2681 | Carnitine shuttle (cytosolic) | 3.6016 | 2.7658 | 3.9459 | 3.3065 | 0.3443 | -0.1397 | 0.5556 |
| HMR_2684 | Carnitine shuttle (cytosolic) | -0.0385 | 6.6033 | -0.0496 | 7.8482 | 0.0112 | 0.0029 | 0.4988 |
| HMR_2687 | Carnitine shuttle (cytosolic) | 0.6197 | 0.1118 | 0.6883 | 0.1375 | 0.0685 | -0.1373 | 0.5546 |
| HMR_2690 | Carnitine shuttle (cytosolic) | 0.9589 | 0.2613 | 1.1016 | 0.345 | 0.1427 | -0.1832 | 0.5727 |
| HMR_2693 | Carnitine shuttle (cytosolic) | 0.9447 | 0.2296 | 1.0715 | 0.2955 | 0.1269 | -0.1751 | 0.5695 |
| HMR_2699 | Carnitine shuttle (cytosolic) | -3.5953 | 2.7036 | -3.9425 | 3.2354 | 0.3472 | 0.1425 | 0.4434 |
| HMR_2702 | Carnitine shuttle (cytosolic) | -3.6117 | 2.8704 | -3.9534 | 3.4254 | 0.3417 | 0.1362 | 0.4458 |
| HMR_2705 | Carnitine shuttle (cytosolic) | 3.6358 | 2.9506 | 3.9749 | 3.5179 | 0.339 | -0.1333 | 0.553 |
| HMR_2708 | Carnitine shuttle (cytosolic) | 0.6014 | 0.0826 | 0.6557 | 0.0978 | 0.0544 | -0.128 | 0.5509 |
| HMR_2711 | Carnitine shuttle (cytosolic) | -1.3442 | 0.4441 | -1.5263 | 0.5848 | 0.1821 | 0.1795 | 0.4288 |
| HMR_2714 | Carnitine shuttle (cytosolic) | 1.1461 | 0.3881 | 1.3041 | 0.5027 | 0.158 | -0.1674 | 0.5665 |
| HMR_2717 | Carnitine shuttle (cytosolic) | 0.8917 | 0.2356 | 1.0013 | 0.2969 | 0.1097 | -0.1503 | 0.5597 |
| HMR_2720 | Carnitine shuttle (cytosolic) | 61.3004 | 154238.0413 | 61.8721 | 154193.0235 | 0.5717 | -0.001 | 0.5004 |
| HMR_2726 | Carnitine shuttle (cytosolic) | 0.7357 | 0.1545 | 0.8111 | 0.1879 | 0.0754 | -0.1289 | 0.5513 |
| HMR_2730 | Carnitine shuttle (cytosolic) | 1.3155 | 0.2978 | 1.4423 | 0.3577 | 0.1267 | -0.1566 | 0.5622 |
| HMR_2733 | Carnitine shuttle (cytosolic) | 0.9224 | 0.2406 | 1.0439 | 0.3083 | 0.1215 | -0.164 | 0.5651 |
| HMR_2736 | Carnitine shuttle (cytosolic) | 0.8027 | 0.1452 | 0.8917 | 0.1787 | 0.089 | -0.1565 | 0.5622 |
| HMR_2739 | Carnitine shuttle (cytosolic) | 0.7492 | 0.1625 | 0.8321 | 0.2 | 0.083 | -0.1378 | 0.5548 |
| HMR_2742 | Carnitine shuttle (cytosolic) | -5.7706 | 3.3177 | -5.9742 | 3.4623 | 0.2035 | 0.0782 | 0.4688 |
| HMR_2746 | Carnitine shuttle (cytosolic) | 5.1503 | 5.5029 | 5.5564 | 6.5238 | 0.4062 | -0.1171 | 0.5466 |
| HMR_2752 | Carnitine shuttle (cytosolic) | -3.7437 | 3.8032 | -4.0774 | 4.5324 | 0.3337 | 0.1156 | 0.454 |
| HMR_2755 | Carnitine shuttle (cytosolic) | -3.0089 | 1.7296 | -3.2206 | 2.0054 | 0.2118 | 0.1096 | 0.4564 |
| HMR_2759 | Carnitine shuttle (cytosolic) | 0.7096 | 0.1421 | 0.7844 | 0.1737 | 0.0748 | -0.1331 | 0.5529 |
| HMR_2762 | Carnitine shuttle (cytosolic) | 1.3369 | 0.3322 | 1.4834 | 0.4046 | 0.1464 | -0.1706 | 0.5677 |
| HMR_2768 | Carnitine shuttle (cytosolic) | 0.8626 | 0.2133 | 0.9793 | 0.2749 | 0.1167 | -0.167 | 0.5663 |
| HMR_2771 | Carnitine shuttle (cytosolic) | 0.7618 | 0.1313 | 0.8473 | 0.1618 | 0.0855 | -0.158 | 0.5628 |
| HMR_2774 | Carnitine shuttle (cytosolic) | 0.7392 | 0.1249 | 0.8159 | 0.1517 | 0.0767 | -0.1458 | 0.5579 |
| HMR_2877 | Carnitine shuttle (cytosolic) | 0.0256 | 0.0002 | 0.0256 | 0.0002 | 0.00E+00 | -0.0002 | 0.5001 |
| HMR_3521 | Carnitine shuttle (cytosolic) | 0.8715 | 0.1015 | 1.1461 | 0.1669 | 0.2746 | -0.5299 | 0.7019 |
| HMR_0162 | Carnitine shuttle (mitochondrial) | 2.82 | 1.789 | 3.0045 | 1.9494 | 0.1845 | -0.0954 | 0.538 |
| HMR_0163 | Carnitine shuttle (mitochondrial) | 2.815 | 1.7961 | 2.9996 | 1.9567 | 0.1846 | -0.0953 | 0.538 |
| HMR_2592 | Carnitine shuttle (mitochondrial) | 368.2881 | 12332.6868 | 557.3149 | 12859.8642 | 189.0268 | -1.1909 | 0.8832 |

| | | | | | | | | |
|---|---|---|---|---|---|---|---|---|
| HMR_2593 | Carnitine shuttle (mitochondrial) | -368.2912 | 12332.7765 | -557.3183 | 12859.9574 | 189.0272 | 1.1909 | 0.1168 |
| HMR_2596 | Carnitine shuttle (mitochondrial) | 119.1906 | 153496.0017 | 118.9546 | 151064.6356 | 0.236 | 0.0004 | 0.4998 |
| HMR_2598 | Carnitine shuttle (mitochondrial) | -119.1941 | 153496.003 | -118.9581 | 151064.6348 | 0.236 | -0.0004 | 0.5002 |
| HMR_2600 | Carnitine shuttle (mitochondrial) | 129.1918 | 144.8165 | 137.9243 | 156.5627 | 8.7325 | -0.503 | 0.6925 |
| HMR_2601 | Carnitine shuttle (mitochondrial) | -129.1932 | 144.8119 | -137.9262 | 156.5577 | 8.733 | 0.5031 | 0.3075 |
| HMR_2603 | Carnitine shuttle (mitochondrial) | -0.2899 | 0.2585 | -0.3183 | 0.2902 | 0.0284 | 0.0383 | 0.4847 |
| HMR_2604 | Carnitine shuttle (mitochondrial) | 0.2847 | 0.2661 | 0.3133 | 0.2982 | 0.0286 | -0.0381 | 0.5152 |
| HMR_2606 | Carnitine shuttle (mitochondrial) | -0.4692 | 0.1587 | -0.5033 | 0.1736 | 0.0341 | 0.0591 | 0.4764 |
| HMR_2607 | Carnitine shuttle (mitochondrial) | 0.4674 | 0.1623 | 0.5019 | 0.1773 | 0.0345 | -0.0591 | 0.5236 |
| HMR_2609 | Carnitine shuttle (mitochondrial) | 1.481 | 1.0907 | 1.7567 | 1.3544 | 0.2757 | -0.1763 | 0.57 |
| HMR_2610 | Carnitine shuttle (mitochondrial) | -1.4786 | 1.0941 | -1.7548 | 1.3586 | 0.2761 | 0.1763 | 0.43 |
| HMR_2612 | Carnitine shuttle (mitochondrial) | -0.9073 | 0.2332 | -1.015 | 0.2914 | 0.1077 | 0.1487 | 0.4409 |
| HMR_2613 | Carnitine shuttle (mitochondrial) | -0.9054 | 0.2341 | -1.0131 | 0.2924 | 0.1077 | 0.1485 | 0.441 |
| HMR_2616 | Carnitine shuttle (mitochondrial) | -0.6235 | 0.3394 | -0.7208 | 0.4067 | 0.0973 | 0.1127 | 0.4551 |
| HMR_2618 | Carnitine shuttle (mitochondrial) | -0.6175 | 0.3466 | -0.7155 | 0.4138 | 0.0979 | 0.1123 | 0.4553 |
| HMR_2621 | Carnitine shuttle (mitochondrial) | -0.7102 | 0.2921 | -0.8135 | 0.3578 | 0.1033 | 0.1281 | 0.449 |
| HMR_2622 | Carnitine shuttle (mitochondrial) | -0.7059 | 0.2972 | -0.8094 | 0.363 | 0.1036 | 0.1274 | 0.4493 |
| HMR_2624 | Carnitine shuttle (mitochondrial) | -0.3947 | 0.1748 | -0.4264 | 0.1923 | 0.0317 | 0.0523 | 0.4791 |
| HMR_2625 | Carnitine shuttle (mitochondrial) | 0.394 | 0.1775 | 0.426 | 0.1952 | 0.032 | -0.0525 | 0.5209 |
| HMR_2629 | Carnitine shuttle (mitochondrial) | 1.1724 | 5.2291 | 1.1062 | 6.332 | 0.0662 | 0.0195 | 0.4922 |
| HMR_2630 | Carnitine shuttle (mitochondrial) | -1.1693 | 5.2201 | -1.1038 | 6.3231 | 0.0655 | -0.0193 | 0.5077 |
| HMR_2634 | Carnitine shuttle (mitochondrial) | 0.2982 | 0.6372 | 0.2289 | 0.7736 | 0.0694 | 0.0584 | 0.4767 |
| HMR_2635 | Carnitine shuttle (mitochondrial) | 0.3009 | 0.6408 | 0.2318 | 0.7777 | 0.0691 | 0.058 | 0.4769 |
| HMR_2638 | Carnitine shuttle (mitochondrial) | -5.4101 | 1.8977 | -5.8875 | 2.3161 | 0.4773 | 0.2325 | 0.4081 |
| HMR_2640 | Carnitine shuttle (mitochondrial) | 5.4023 | 1.908 | 5.8805 | 2.3263 | 0.4781 | -0.2324 | 0.5919 |
| HMR_2644 | Carnitine shuttle (mitochondrial) | 3.027 | 3.1902 | 3.3324 | 3.7874 | 0.3054 | -0.1156 | 0.546 |
| HMR_2646 | Carnitine shuttle (mitochondrial) | -3.0144 | 3.183 | -3.3201 | 3.7802 | 0.3057 | 0.1158 | 0.4539 |
| HMR_2649 | Carnitine shuttle (mitochondrial) | -0.7247 | 0.1131 | -0.793 | 0.1314 | 0.0683 | 0.1382 | 0.445 |
| HMR_2650 | Carnitine shuttle (mitochondrial) | -0.7205 | 0.1174 | -0.7894 | 0.1361 | 0.0688 | 0.1367 | 0.4456 |
| HMR_2652 | Carnitine shuttle (mitochondrial) | -0.4738 | 0.0673 | -0.5018 | 0.0753 | 0.028 | 0.0743 | 0.4704 |
| HMR_2653 | Carnitine shuttle (mitochondrial) | -0.4766 | 0.0676 | -0.505 | 0.0757 | 0.0285 | 0.0751 | 0.4701 |
| HMR_2655 | Carnitine shuttle (mitochondrial) | -0.0864 | 7.39 | -0.1008 | 8.7923 | 0.0144 | 0.0036 | 0.4986 |
| HMR_2656 | Carnitine shuttle (mitochondrial) | 0.0914 | 7.3805 | 0.1051 | 8.7828 | 0.0137 | -0.0034 | 0.5014 |
| HMR_2658 | Carnitine shuttle (mitochondrial) | -1.0073 | 0.2469 | -1.1272 | 0.3066 | 0.1199 | 0.1611 | 0.436 |
| HMR_2659 | Carnitine shuttle (mitochondrial) | -1.0022 | 0.2499 | -1.1226 | 0.3098 | 0.1204 | 0.1609 | 0.4361 |
| HMR_2661 | Carnitine shuttle (mitochondrial) | 2.9248 | 2.9483 | 3.1753 | 3.5295 | 0.2505 | -0.0984 | 0.5392 |
| HMR_2662 | Carnitine shuttle (mitochondrial) | -2.9186 | 2.9388 | -3.1698 | 3.52 | 0.2513 | 0.0989 | 0.4606 |
| HMR_2664 | Carnitine shuttle (mitochondrial) | 1.6701 | 4.3914 | 1.7546 | 5.2708 | 0.0845 | -0.0272 | 0.5108 |
| HMR_2665 | Carnitine shuttle (mitochondrial) | -1.671 | 4.3834 | -1.7559 | 5.2629 | 0.0849 | 0.0273 | 0.4891 |
| HMR_2667 | Carnitine shuttle (mitochondrial) | 0.7052 | 0.1398 | 0.7911 | 0.1753 | 0.0858 | -0.1529 | 0.5608 |
| HMR_2668 | Carnitine shuttle (mitochondrial) | -0.7033 | 0.1408 | -0.7893 | 0.1764 | 0.086 | 0.1527 | 0.4393 |
| HMR_2670 | Carnitine shuttle (mitochondrial) | -0.6941 | 0.1477 | -0.7817 | 0.1868 | 0.0875 | 0.1514 | 0.4398 |
| HMR_2671 | Carnitine shuttle (mitochondrial) | -0.6933 | 0.1474 | -0.7809 | 0.1865 | 0.0876 | 0.1515 | 0.4398 |
| HMR_2673 | Carnitine shuttle (mitochondrial) | 1.2236 | 0.4501 | 1.4433 | 0.6237 | 0.2197 | -0.212 | 0.584 |

| | | | | | | | | |
|---|---|---|---|---|---|---|---|---|
| HMR_2674 | Carnitine shuttle (mitochondrial) | -1.22 | 0.45 | -1.4399 | 0.6237 | 0.2198 | 0.2122 | 0.416 |
| HMR_2676 | Carnitine shuttle (mitochondrial) | 0.1097 | 5.7593 | 0.1192 | 6.8777 | 0.0095 | -0.0027 | 0.5011 |
| HMR_2677 | Carnitine shuttle (mitochondrial) | -0.0962 | 5.757 | -0.1058 | 6.8755 | 0.0097 | 0.0027 | 0.4989 |
| HMR_2679 | Carnitine shuttle (mitochondrial) | 0.111 | 6.2141 | 0.1208 | 7.3941 | 0.0098 | -0.0027 | 0.5011 |
| HMR_2680 | Carnitine shuttle (mitochondrial) | -0.1045 | 6.2085 | -0.115 | 7.3887 | 0.0105 | 0.0028 | 0.4989 |
| HMR_2682 | Carnitine shuttle (mitochondrial) | -3.5826 | 2.7616 | -3.9262 | 3.3025 | 0.3437 | 0.1396 | 0.4445 |
| HMR_2683 | Carnitine shuttle (mitochondrial) | -3.5896 | 2.7599 | -3.9326 | 3.3009 | 0.3429 | 0.1393 | 0.4446 |
| HMR_2685 | Carnitine shuttle (mitochondrial) | 0.0642 | 6.5951 | 0.0756 | 7.8402 | 0.0114 | -0.003 | 0.5012 |
| HMR_2686 | Carnitine shuttle (mitochondrial) | 0.0634 | 6.5872 | 0.0752 | 7.8323 | 0.0118 | -0.0031 | 0.5012 |
| HMR_2688 | Carnitine shuttle (mitochondrial) | -0.5968 | 0.1072 | -0.6652 | 0.1329 | 0.0684 | 0.1396 | 0.4445 |
| HMR_2689 | Carnitine shuttle (mitochondrial) | -0.5963 | 0.1072 | -0.6649 | 0.133 | 0.0685 | 0.1399 | 0.4444 |
| HMR_2691 | Carnitine shuttle (mitochondrial) | 0.9317 | 0.2568 | 1.0744 | 0.3406 | 0.1427 | -0.1846 | 0.5732 |
| HMR_2692 | Carnitine shuttle (mitochondrial) | -0.9286 | 0.257 | -1.0716 | 0.3408 | 0.1429 | 0.1849 | 0.4267 |
| HMR_2695 | Carnitine shuttle (mitochondrial) | 0.9172 | 0.2274 | 1.0441 | 0.2935 | 0.127 | -0.1759 | 0.5698 |
| HMR_2697 | Carnitine shuttle (mitochondrial) | -0.9141 | 0.2291 | -1.0413 | 0.2952 | 0.1272 | 0.1757 | 0.4303 |
| HMR_2700 | Carnitine shuttle (mitochondrial) | -3.5835 | 2.7005 | -3.9307 | 3.2324 | 0.3472 | 0.1425 | 0.4433 |
| HMR_2701 | Carnitine shuttle (mitochondrial) | 3.5977 | 2.6996 | 3.9448 | 3.2316 | 0.3471 | -0.1425 | 0.5567 |
| HMR_2703 | Carnitine shuttle (mitochondrial) | -3.593 | 2.8651 | -3.9342 | 3.4202 | 0.3412 | 0.1361 | 0.4459 |
| HMR_2704 | Carnitine shuttle (mitochondrial) | 3.6004 | 2.8625 | 3.9409 | 3.4177 | 0.3405 | -0.1359 | 0.554 |
| HMR_2706 | Carnitine shuttle (mitochondrial) | -3.6116 | 2.9445 | -3.9503 | 3.5118 | 0.3388 | 0.1333 | 0.447 |
| HMR_2707 | Carnitine shuttle (mitochondrial) | -3.6135 | 2.9411 | -3.9519 | 3.5085 | 0.3384 | 0.1332 | 0.447 |
| HMR_2709 | Carnitine shuttle (mitochondrial) | -0.5746 | 0.0823 | -0.6293 | 0.0977 | 0.0547 | 0.1289 | 0.4487 |
| HMR_2710 | Carnitine shuttle (mitochondrial) | -0.5716 | 0.0847 | -0.6267 | 0.1004 | 0.0551 | 0.1282 | 0.449 |
| HMR_2712 | Carnitine shuttle (mitochondrial) | -1.1522 | 0.4274 | -1.3339 | 0.5686 | 0.1817 | 0.1821 | 0.4278 |
| HMR_2713 | Carnitine shuttle (mitochondrial) | 1.1503 | 0.4261 | 1.3319 | 0.5674 | 0.1817 | -0.1822 | 0.5723 |
| HMR_2715 | Carnitine shuttle (mitochondrial) | 1.1203 | 0.3828 | 1.2781 | 0.4975 | 0.1578 | -0.1682 | 0.5668 |
| HMR_2716 | Carnitine shuttle (mitochondrial) | -1.1189 | 0.3826 | -1.2766 | 0.4973 | 0.1578 | 0.1682 | 0.4332 |
| HMR_2718 | Carnitine shuttle (mitochondrial) | 0.8672 | 0.2304 | 0.9767 | 0.2916 | 0.1095 | -0.1516 | 0.5602 |
| HMR_2719 | Carnitine shuttle (mitochondrial) | -0.8664 | 0.2301 | -0.9759 | 0.2914 | 0.1095 | 0.1517 | 0.4397 |
| HMR_2722 | Carnitine shuttle (mitochondrial) | 0.8351 | 0.2516 | 0.9333 | 0.3131 | 0.0981 | -0.1306 | 0.5519 |
| HMR_2724 | Carnitine shuttle (mitochondrial) | -0.8371 | 0.2484 | -0.935 | 0.3099 | 0.0978 | 0.1309 | 0.4479 |
| HMR_2727 | Carnitine shuttle (mitochondrial) | 0.6899 | 0.1487 | 0.765 | 0.1821 | 0.0751 | -0.1306 | 0.5519 |
| HMR_2729 | Carnitine shuttle (mitochondrial) | -0.6909 | 0.1482 | -0.766 | 0.1816 | 0.075 | 0.1306 | 0.448 |
| HMR_2731 | Carnitine shuttle (mitochondrial) | -1.1246 | 0.2996 | -1.2519 | 0.3608 | 0.1273 | 0.1567 | 0.4377 |
| HMR_2732 | Carnitine shuttle (mitochondrial) | -1.1215 | 0.3032 | -1.249 | 0.3648 | 0.1275 | 0.156 | 0.438 |
| HMR_2734 | Carnitine shuttle (mitochondrial) | 0.8963 | 0.2363 | 1.0178 | 0.3041 | 0.1215 | -0.1653 | 0.5656 |
| HMR_2735 | Carnitine shuttle (mitochondrial) | -0.8941 | 0.2366 | -1.0158 | 0.3044 | 0.1217 | 0.1654 | 0.4343 |
| HMR_2737 | Carnitine shuttle (mitochondrial) | 0.7743 | 0.1454 | 0.8637 | 0.1791 | 0.0894 | -0.1569 | 0.5623 |
| HMR_2738 | Carnitine shuttle (mitochondrial) | -0.7705 | 0.1484 | -0.8603 | 0.1823 | 0.0898 | 0.1561 | 0.438 |
| HMR_2740 | Carnitine shuttle (mitochondrial) | 0.7254 | 0.1579 | 0.8083 | 0.1955 | 0.0829 | -0.1394 | 0.5554 |
| HMR_2741 | Carnitine shuttle (mitochondrial) | -0.7249 | 0.158 | -0.8078 | 0.1956 | 0.083 | 0.1396 | 0.4445 |
| HMR_2744 | Carnitine shuttle (mitochondrial) | -1.0559 | 0.2554 | -1.2104 | 0.3322 | 0.1546 | 0.2017 | 0.4201 |
| HMR_2745 | Carnitine shuttle (mitochondrial) | 1.0562 | 0.2483 | 1.2105 | 0.3251 | 0.1543 | -0.2037 | 0.5807 |
| HMR_2748 | Carnitine shuttle (mitochondrial) | 5.1739 | 5.493 | 5.5805 | 6.5139 | 0.4066 | -0.1173 | 0.5467 |

| | | | | | | | | |
|---|---|---|---|---|---|---|---|---|
| HMR_2750 | Carnitine shuttle (mitochondrial) | -5.171 | 5.4834 | -5.5782 | 6.5043 | 0.4072 | 0.1176 | 0.4532 |
| HMR_2753 | Carnitine shuttle (mitochondrial) | -3.565 | 3.789 | -3.8979 | 4.5182 | 0.3329 | 0.1155 | 0.454 |
| HMR_2754 | Carnitine shuttle (mitochondrial) | 3.5687 | 3.7879 | 3.901 | 4.5171 | 0.3323 | -0.1153 | 0.5459 |
| HMR_2757 | Carnitine shuttle (mitochondrial) | -2.5567 | 1.6781 | -2.7628 | 1.9534 | 0.2061 | 0.1082 | 0.4569 |
| HMR_2758 | Carnitine shuttle (mitochondrial) | 2.5618 | 1.6736 | 2.7672 | 1.9489 | 0.2055 | -0.1079 | 0.543 |
| HMR_2760 | Carnitine shuttle (mitochondrial) | -0.6633 | 0.1364 | -0.7378 | 0.168 | 0.0745 | 0.1351 | 0.4463 |
| HMR_2761 | Carnitine shuttle (mitochondrial) | -0.6638 | 0.136 | -0.7383 | 0.1677 | 0.0745 | 0.1352 | 0.4462 |
| HMR_2764 | Carnitine shuttle (mitochondrial) | 1.3057 | 0.3362 | 1.4528 | 0.4088 | 0.1471 | -0.1704 | 0.5676 |
| HMR_2766 | Carnitine shuttle (mitochondrial) | -1.3005 | 0.3418 | -1.4481 | 0.4147 | 0.1476 | 0.1697 | 0.4326 |
| HMR_2769 | Carnitine shuttle (mitochondrial) | -0.8365 | 0.2086 | -0.9532 | 0.2703 | 0.1167 | 0.1686 | 0.4331 |
| HMR_2770 | Carnitine shuttle (mitochondrial) | -0.8341 | 0.2087 | -0.9509 | 0.2704 | 0.1169 | 0.1689 | 0.433 |
| HMR_2772 | Carnitine shuttle (mitochondrial) | 0.7331 | 0.1313 | 0.819 | 0.1621 | 0.0859 | -0.1586 | 0.563 |
| HMR_2773 | Carnitine shuttle (mitochondrial) | -0.7289 | 0.1341 | -0.8153 | 0.1652 | 0.0864 | 0.1579 | 0.4373 |
| HMR_2775 | Carnitine shuttle (mitochondrial) | 0.7117 | 0.1248 | 0.7887 | 0.1518 | 0.0769 | -0.1463 | 0.5581 |
| HMR_2776 | Carnitine shuttle (mitochondrial) | -0.7086 | 0.1276 | -0.7859 | 0.1548 | 0.0773 | 0.1454 | 0.4422 |
| HMR_3519 | Carnitine shuttle (mitochondrial) | -0.8745 | 0.1034 | -1.1547 | 0.1709 | 0.2802 | 0.535 | 0.2963 |
| HMR_3520 | Carnitine shuttle (mitochondrial) | 0.8725 | 0.1024 | 1.1499 | 0.1688 | 0.2774 | -0.5327 | 0.7029 |
| HMR_3027 | Carnitine shuttle (peroxisomal) | -4.4099 | 6.911 | -4.5287 | 7.2891 | 0.1189 | 0.0315 | 0.4874 |
| HMR_3028 | Carnitine shuttle (peroxisomal) | 4.4109 | 6.9084 | 4.5295 | 7.2864 | 0.1186 | -0.0315 | 0.5126 |
| HMR_3029 | Carnitine shuttle (peroxisomal) | -21.2871 | 159468.2436 | -113.8137 | 138582.983 | 92.5266 | 0.1695 | 0.4327 |
| HMR_3030 | Carnitine shuttle (peroxisomal) | 21.2872 | 159468.2377 | 113.8137 | 138582.9759 | 92.5265 | -0.1695 | 0.5673 |
| HMR_3033 | Carnitine shuttle (peroxisomal) | 2.8257 | 1.7845 | 3.01 | 1.9445 | 0.1843 | -0.0954 | 0.538 |
| HMR_3034 | Carnitine shuttle (peroxisomal) | -2.8266 | 1.783 | -3.0106 | 1.9423 | 0.184 | 0.0954 | 0.462 |
| HMR_3035 | Carnitine shuttle (peroxisomal) | 50.9104 | 172601.9412 | 79.4913 | 167614.5121 | 28.5809 | -0.049 | 0.5195 |
| HMR_3037 | Carnitine shuttle (peroxisomal) | -50.9108 | 172601.9391 | -79.4914 | 167614.51 | 28.5806 | 0.049 | 0.4805 |
| HMR_2787 | Carnitine shuttle (endoplasmic reticular) | -0.0255 | 0.0002 | -0.0254 | 0.0002 | 0.00E+00 | -0.0007 | 0.5003 |
| HMR_2788 | Carnitine shuttle (endoplasmic reticular) | 0.0255 | 0.0002 | 0.0255 | 0.0002 | 0.00E+00 | 1.00E-04 | 0.5 |
| HMR_2789 | Carnitine shuttle (endoplasmic reticular) | -0.0254 | 0.0002 | -0.0253 | 0.0002 | 0.00E+00 | -0.0009 | 0.5003 |
| HMR_2790 | Carnitine shuttle (endoplasmic reticular) | 0.0255 | 0.0002 | 0.0255 | 0.0002 | 0.00E+00 | 1.00E-04 | 0.5 |
| HMR_2791 | Carnitine shuttle (endoplasmic reticular) | -0.195 | 0.0153 | -0.1953 | 0.0153 | 0.0003 | 0.0016 | 0.4994 |
| HMR_2792 | Carnitine shuttle (endoplasmic reticular) | 0.1839 | 0.0122 | 0.1842 | 0.0122 | 0.0003 | -0.002 | 0.5008 |
| HMR_2793 | Carnitine shuttle (endoplasmic reticular) | -0.0253 | 0.0002 | -0.0253 | 0.0002 | 0.00E+00 | -0.0002 | 0.5001 |
| HMR_2794 | Carnitine shuttle (endoplasmic reticular) | -0.0255 | 0.0002 | -0.0255 | 0.0002 | 0.00E+00 | -1.00E-04 | 0.5 |
| HMR_2795 | Carnitine shuttle (endoplasmic reticular) | -0.0255 | 0.0002 | -0.0254 | 0.0002 | 0.00E+00 | -0.0011 | 0.5004 |
| HMR_2796 | Carnitine shuttle (endoplasmic reticular) | -0.0255 | 0.0002 | -0.0255 | 0.0002 | 0.00E+00 | -0.0001 | 0.5 |
| HMR_2797 | Carnitine shuttle (endoplasmic reticular) | -0.0254 | 0.0002 | -0.0254 | 0.0002 | 0.00E+00 | -0.0007 | 0.5003 |
| HMR_2798 | Carnitine shuttle (endoplasmic reticular) | -0.0255 | 0.0002 | -0.0255 | 0.0002 | 0.00E+00 | -1.00E-04 | 0.5 |
| HMR_2799 | Carnitine shuttle (endoplasmic reticular) | -0.0254 | 0.0002 | -0.0253 | 0.0002 | 0.00E+00 | -0.0008 | 0.5003 |
| HMR_2800 | Carnitine shuttle (endoplasmic reticular) | 0.0255 | 0.0002 | 0.0255 | 0.0002 | 0.00E+00 | 1.00E-04 | 0.5 |
| HMR_2801 | Carnitine shuttle (endoplasmic reticular) | -1.2197 | 0.2401 | -1.233 | 0.2449 | 0.0133 | 0.0191 | 0.4924 |
| HMR_2803 | Carnitine shuttle (endoplasmic reticular) | 1.2295 | 0.2305 | 1.2426 | 0.2352 | 0.0131 | -0.0192 | 0.5076 |
| HMR_2805 | Carnitine shuttle (endoplasmic reticular) | -0.3967 | 0.0414 | -0.3992 | 0.0418 | 0.0025 | 0.0088 | 0.4965 |
| HMR_2806 | Carnitine shuttle (endoplasmic reticular) | -0.3965 | 0.0352 | -0.3992 | 0.0356 | 0.0028 | 0.0103 | 0.4959 |
| HMR_2807 | Carnitine shuttle (endoplasmic reticular) | -0.0255 | 0.0002 | -0.0255 | 0.0002 | 0.00E+00 | -0.0013 | 0.5005 |

| | | | | | | | | |
|---|---|---|---|---|---|---|---|---|
| HMR_2809 | Carnitine shuttle (endoplasmic reticular) | 0.0255 | 0.0002 | 0.0255 | 0.0002 | 0.00E+00 | 0.0001 | 0.5 |
| HMR_2811 | Carnitine shuttle (endoplasmic reticular) | -0.0251 | 0.0002 | -0.0251 | 0.0002 | 0.00E+00 | -0.0002 | 0.5001 |
| HMR_2812 | Carnitine shuttle (endoplasmic reticular) | 0.0255 | 0.0002 | 0.0255 | 0.0002 | 0.00E+00 | 1.00E-04 | 0.5 |
| HMR_2813 | Carnitine shuttle (endoplasmic reticular) | -0.0254 | 0.0002 | -0.0254 | 0.0002 | 0.00E+00 | -0.0007 | 0.5003 |
| HMR_2814 | Carnitine shuttle (endoplasmic reticular) | -0.0255 | 0.0002 | -0.0255 | 0.0002 | 0.00E+00 | -1.00E-04 | 0.5 |
| HMR_2815 | Carnitine shuttle (endoplasmic reticular) | -0.0252 | 0.0002 | -0.0252 | 0.0002 | 0.00E+00 | -0.0006 | 0.5002 |
| HMR_2816 | Carnitine shuttle (endoplasmic reticular) | -0.0255 | 0.0002 | -0.0255 | 0.0002 | 0.00E+00 | -1.00E-04 | 0.5 |
| HMR_2817 | Carnitine shuttle (endoplasmic reticular) | -0.2098 | 0.0172 | -0.2108 | 0.0173 | 0.001 | 0.0052 | 0.4979 |
| HMR_2819 | Carnitine shuttle (endoplasmic reticular) | 0.2036 | 0.0139 | 0.2045 | 0.014 | 0.0009 | -0.0052 | 0.5021 |
| HMR_2821 | Carnitine shuttle (endoplasmic reticular) | -0.0254 | 0.0002 | -0.0254 | 0.0002 | 0.00E+00 | -0.0008 | 0.5003 |
| HMR_2822 | Carnitine shuttle (endoplasmic reticular) | -0.0255 | 0.0002 | -0.0255 | 0.0002 | 0.00E+00 | -1.00E-04 | 0.5 |
| HMR_2823 | Carnitine shuttle (endoplasmic reticular) | -0.0252 | 0.0002 | -0.0252 | 0.0002 | 0.00E+00 | 0.0003 | 0.4999 |
| HMR_2824 | Carnitine shuttle (endoplasmic reticular) | 0.0255 | 0.0002 | 0.0255 | 0.0002 | 0.00E+00 | 1.00E-04 | 0.5 |
| HMR_2825 | Carnitine shuttle (endoplasmic reticular) | 1.7521 | 0.4506 | 1.7695 | 0.4601 | 0.0175 | -0.0183 | 0.5073 |
| HMR_2827 | Carnitine shuttle (endoplasmic reticular) | 1.7581 | 0.4402 | 1.7757 | 0.4497 | 0.0175 | -0.0186 | 0.5074 |
| HMR_2829 | Carnitine shuttle (endoplasmic reticular) | 0.0253 | 0.0002 | 0.0253 | 0.0002 | 0.00E+00 | 0.0002 | 0.4999 |
| HMR_2830 | Carnitine shuttle (endoplasmic reticular) | 0.0255 | 0.0002 | 0.0255 | 0.0002 | 0.00E+00 | 1.00E-04 | 0.5 |
| HMR_2831 | Carnitine shuttle (endoplasmic reticular) | -0.0253 | 0.0002 | -0.0253 | 0.0002 | 0.00E+00 | 0.00E+00 | 0.5 |
| HMR_2832 | Carnitine shuttle (endoplasmic reticular) | -0.0254 | 0.0002 | -0.0254 | 0.0002 | 0.00E+00 | -1.00E-04 | 0.5 |
| HMR_2833 | Carnitine shuttle (endoplasmic reticular) | 0.0254 | 0.0002 | 0.0254 | 0.0002 | 0.00E+00 | 0.0003 | 0.4999 |
| HMR_2834 | Carnitine shuttle (endoplasmic reticular) | 0.0255 | 0.0002 | 0.0255 | 0.0002 | 0.00E+00 | 1.00E-04 | 0.5 |
| HMR_2835 | Carnitine shuttle (endoplasmic reticular) | -0.0251 | 0.0002 | -0.025 | 0.0002 | 0.00E+00 | -0.0003 | 0.5001 |
| HMR_2836 | Carnitine shuttle (endoplasmic reticular) | 0.0255 | 0.0002 | 0.0255 | 0.0002 | 0.00E+00 | 1.00E-04 | 0.5 |
| HMR_2837 | Carnitine shuttle (endoplasmic reticular) | -0.0252 | 0.0002 | -0.0252 | 0.0002 | 0.00E+00 | 0.0003 | 0.4999 |
| HMR_2838 | Carnitine shuttle (endoplasmic reticular) | 0.0255 | 0.0002 | 0.0255 | 0.0002 | 0.00E+00 | 1.00E-04 | 0.5 |
| HMR_2839 | Carnitine shuttle (endoplasmic reticular) | -0.0253 | 0.0002 | -0.0253 | 0.0002 | 0.00E+00 | -1.00E-04 | 0.5 |
| HMR_2840 | Carnitine shuttle (endoplasmic reticular) | -0.0255 | 0.0002 | -0.0255 | 0.0002 | 0.00E+00 | -1.00E-04 | 0.5 |
| HMR_2841 | Carnitine shuttle (endoplasmic reticular) | -0.0253 | 0.0002 | -0.0253 | 0.0002 | 0.00E+00 | -0.0002 | 0.5001 |
| HMR_2842 | Carnitine shuttle (endoplasmic reticular) | -0.0255 | 0.0002 | -0.0255 | 0.0002 | 0.00E+00 | -1.00E-04 | 0.5 |
| HMR_2843 | Carnitine shuttle (endoplasmic reticular) | -0.0252 | 0.0002 | -0.0252 | 0.0002 | 0.00E+00 | 0.0003 | 0.4999 |
| HMR_2844 | Carnitine shuttle (endoplasmic reticular) | -0.0255 | 0.0002 | -0.0255 | 0.0002 | 0.00E+00 | -1.00E-04 | 0.5 |
| HMR_2845 | Carnitine shuttle (endoplasmic reticular) | 0.0254 | 0.0002 | 0.0253 | 0.0002 | 0.00E+00 | 0.0003 | 0.4999 |
| HMR_2846 | Carnitine shuttle (endoplasmic reticular) | -0.0255 | 0.0002 | -0.0255 | 0.0002 | 0.00E+00 | -1.00E-04 | 0.5 |
| HMR_2847 | Carnitine shuttle (endoplasmic reticular) | 0.0254 | 0.0002 | 0.0254 | 0.0002 | 0.00E+00 | 0.0004 | 0.4998 |
| HMR_2848 | Carnitine shuttle (endoplasmic reticular) | -0.0255 | 0.0002 | -0.0255 | 0.0002 | 0.00E+00 | -1.00E-04 | 0.5 |
| HMR_2849 | Carnitine shuttle (endoplasmic reticular) | -0.025 | 0.0002 | -0.025 | 0.0002 | 0.00E+00 | -0.0003 | 0.5001 |
| HMR_2850 | Carnitine shuttle (endoplasmic reticular) | 0.0255 | 0.0002 | 0.0255 | 0.0002 | 0.00E+00 | 1.00E-04 | 0.5 |
| HMR_2851 | Carnitine shuttle (endoplasmic reticular) | -0.0252 | 0.0002 | -0.0252 | 0.0002 | 0.00E+00 | 0.0003 | 0.4999 |
| HMR_2852 | Carnitine shuttle (endoplasmic reticular) | 0.0255 | 0.0002 | 0.0255 | 0.0002 | 0.00E+00 | 1.00E-04 | 0.5 |
| HMR_2853 | Carnitine shuttle (endoplasmic reticular) | -0.0253 | 0.0002 | -0.0253 | 0.0002 | 0.00E+00 | 0.00E+00 | 0.5 |
| HMR_2854 | Carnitine shuttle (endoplasmic reticular) | -0.0255 | 0.0002 | -0.0255 | 0.0002 | 0.00E+00 | -1.00E-04 | 0.5 |
| HMR_2855 | Carnitine shuttle (endoplasmic reticular) | -0.0253 | 0.0002 | -0.0253 | 0.0002 | 0.00E+00 | -0.0006 | 0.5002 |
| HMR_2856 | Carnitine shuttle (endoplasmic reticular) | -0.0255 | 0.0002 | -0.0255 | 0.0002 | 0.00E+00 | -1.00E-04 | 0.5 |
| HMR_2857 | Carnitine shuttle (endoplasmic reticular) | -0.1911 | 0.0147 | -0.1913 | 0.0148 | 0.0001 | 0.0007 | 0.4997 |

| | | | | | | | | |
|---|---|---|---|---|---|---|---|---|
| HMR_2859 | Carnitine shuttle (endoplasmic reticular) | 0.1776 | 0.0116 | 0.1777 | 0.0117 | 0.0002 | -0.0011 | 0.5004 |
| HMR_2861 | Carnitine shuttle (endoplasmic reticular) | 0.0253 | 0.0002 | 0.0253 | 0.0002 | 0.00E+00 | 0.0001 | 0.5 |
| HMR_2862 | Carnitine shuttle (endoplasmic reticular) | -0.0255 | 0.0002 | -0.0255 | 0.0002 | 0.00E+00 | -1.00E-04 | 0.5 |
| HMR_2863 | Carnitine shuttle (endoplasmic reticular) | 0.0253 | 0.0002 | 0.0253 | 0.0002 | 0.00E+00 | 0.0001 | 0.5 |
| HMR_2864 | Carnitine shuttle (endoplasmic reticular) | -0.0255 | 0.0002 | -0.0255 | 0.0002 | 0.00E+00 | -1.00E-04 | 0.5 |
| HMR_2865 | Carnitine shuttle (endoplasmic reticular) | 60.47 | 154237.856 | 60.9431 | 154192.7932 | 0.4731 | -0.0009 | 0.5003 |
| HMR_2866 | Carnitine shuttle (endoplasmic reticular) | -60.4833 | 154237.852 | -60.956 | 154192.7891 | 0.4728 | 0.0009 | 0.4997 |
| HMR_2867 | Carnitine shuttle (endoplasmic reticular) | 0.0494 | 0.0008 | 0.0494 | 0.0008 | 0.00E+00 | 0.00E+00 | 0.5 |
| HMR_2868 | Carnitine shuttle (endoplasmic reticular) | -0.0497 | 0.0008 | -0.0497 | 0.0008 | 0.00E+00 | -0.0001 | 0.5001 |
| HMR_2869 | Carnitine shuttle (endoplasmic reticular) | -0.1878 | 0.0142 | -0.1873 | 0.0142 | 0.0005 | -0.0029 | 0.5011 |
| HMR_2870 | Carnitine shuttle (endoplasmic reticular) | -0.1717 | 0.0112 | -0.1714 | 0.0111 | 0.0003 | -0.0018 | 0.5007 |
| HMR_2871 | Carnitine shuttle (endoplasmic reticular) | 0.0253 | 0.0002 | 0.0253 | 0.0002 | 0.00E+00 | 0.0003 | 0.4999 |
| HMR_2872 | Carnitine shuttle (endoplasmic reticular) | -0.0255 | 0.0002 | -0.0255 | 0.0002 | 0.00E+00 | -1.00E-04 | 0.5 |
| HMR_2873 | Carnitine shuttle (endoplasmic reticular) | 0.0254 | 0.0002 | 0.0254 | 0.0002 | 0.00E+00 | 0.0006 | 0.4997 |
| HMR_2874 | Carnitine shuttle (endoplasmic reticular) | -0.0255 | 0.0002 | -0.0255 | 0.0002 | 0.00E+00 | -1.00E-04 | 0.5 |
| HMR_2875 | Carnitine shuttle (endoplasmic reticular) | 0.0253 | 0.0002 | 0.0253 | 0.0002 | 0.00E+00 | 0.0002 | 0.4999 |
| HMR_2876 | Carnitine shuttle (endoplasmic reticular) | -0.0254 | 0.0002 | -0.0254 | 0.0002 | 0.00E+00 | -1.00E-04 | 0.5 |
| HMR_2878 | Carnitine shuttle (endoplasmic reticular) | 0.0253 | 0.0002 | 0.0253 | 0.0002 | 0.00E+00 | 0.0003 | 0.4999 |
| HMR_2879 | Carnitine shuttle (endoplasmic reticular) | -0.0254 | 0.0002 | -0.0254 | 0.0002 | 0.00E+00 | -1.00E-04 | 0.5 |
| HMR_2880 | Carnitine shuttle (endoplasmic reticular) | -4.7151 | 3.0576 | -4.7636 | 3.1264 | 0.0486 | 0.0195 | 0.4922 |
| HMR_2882 | Carnitine shuttle (endoplasmic reticular) | 4.7239 | 3.0436 | 4.7724 | 3.1124 | 0.0485 | -0.0195 | 0.5078 |
| HMR_2884 | Carnitine shuttle (endoplasmic reticular) | -0.0253 | 0.0002 | -0.0253 | 0.0002 | 0.00E+00 | 0.0002 | 0.4999 |
| HMR_2886 | Carnitine shuttle (endoplasmic reticular) | 0.0255 | 0.0002 | 0.0255 | 0.0002 | 0.00E+00 | 1.00E-04 | 0.5 |
| HMR_2888 | Carnitine shuttle (endoplasmic reticular) | -0.1817 | 0.0138 | -0.182 | 0.0138 | 0.0003 | 0.002 | 0.4992 |
| HMR_2890 | Carnitine shuttle (endoplasmic reticular) | 0.1696 | 0.0109 | 0.1699 | 0.011 | 0.0003 | -0.0018 | 0.5007 |
| HMR_2892 | Carnitine shuttle (endoplasmic reticular) | -0.4566 | 0.0528 | -0.4617 | 0.0536 | 0.0051 | 0.0155 | 0.4938 |
| HMR_2894 | Carnitine shuttle (endoplasmic reticular) | 0.4657 | 0.0463 | 0.4706 | 0.047 | 0.0049 | -0.016 | 0.5064 |
| HMR_2896 | Carnitine shuttle (endoplasmic reticular) | -0.0495 | 0.0008 | -0.0495 | 0.0008 | 0.00E+00 | -1.00E-04 | 0.5 |
| HMR_2897 | Carnitine shuttle (endoplasmic reticular) | -0.0497 | 0.0008 | -0.0497 | 0.0008 | 0.00E+00 | -0.0002 | 0.5001 |
| HMR_2898 | Carnitine shuttle (endoplasmic reticular) | 0.0254 | 0.0002 | 0.0254 | 0.0002 | 0.00E+00 | 0.0009 | 0.4996 |
| HMR_2899 | Carnitine shuttle (endoplasmic reticular) | -0.0255 | 0.0002 | -0.0255 | 0.0002 | 0.00E+00 | -1.00E-04 | 0.5 |
| HMR_2900 | Carnitine shuttle (endoplasmic reticular) | -0.0253 | 0.0002 | -0.0253 | 0.0002 | 0.00E+00 | -0.0003 | 0.5001 |
| HMR_2901 | Carnitine shuttle (endoplasmic reticular) | -0.0255 | 0.0002 | -0.0255 | 0.0002 | 0.00E+00 | -1.00E-04 | 0.5 |
| HMR_2902 | Carnitine shuttle (endoplasmic reticular) | 0.0254 | 0.0002 | 0.0254 | 0.0002 | 0.00E+00 | 0.0007 | 0.4997 |
| HMR_2903 | Carnitine shuttle (endoplasmic reticular) | -0.0255 | 0.0002 | -0.0255 | 0.0002 | 0.00E+00 | -1.00E-04 | 0.5 |
| HMR_2904 | Carnitine shuttle (endoplasmic reticular) | 0.0254 | 0.0002 | 0.0254 | 0.0002 | 0.00E+00 | 0.0006 | 0.4998 |
| HMR_2905 | Carnitine shuttle (endoplasmic reticular) | -0.0254 | 0.0002 | -0.0254 | 0.0002 | 0.00E+00 | -1.00E-04 | 0.5 |
| HMR_2906 | Carnitine shuttle (endoplasmic reticular) | -0.0255 | 0.0002 | -0.0255 | 0.0002 | 0.00E+00 | -0.0001 | 0.5 |
| HMR_2907 | Carnitine shuttle (endoplasmic reticular) | -0.0253 | 0.0002 | -0.0253 | 0.0002 | 0.00E+00 | -0.0003 | 0.5001 |
| HMR_2908 | Carnitine shuttle (endoplasmic reticular) | 0.0255 | 0.0002 | 0.0255 | 0.0002 | 0.00E+00 | 1.00E-04 | 0.5 |
| HMR_2909 | Carnitine shuttle (endoplasmic reticular) | -0.0254 | 0.0002 | -0.0254 | 0.0002 | 0.00E+00 | 0.0005 | 0.4998 |
| HMR_2910 | Carnitine shuttle (endoplasmic reticular) | -0.0253 | 0.0002 | -0.0253 | 0.0002 | 0.00E+00 | -0.0003 | 0.5001 |
| HMR_2911 | Carnitine shuttle (endoplasmic reticular) | 0.0255 | 0.0002 | 0.0255 | 0.0002 | 0.00E+00 | 1.00E-04 | 0.5 |
| HMR_2912 | Carnitine shuttle (endoplasmic reticular) | 0.0254 | 0.0002 | 0.0254 | 0.0002 | 0.00E+00 | -0.0003 | 0.5001 |

| | | | | | | | | |
|---|---|---|---|---|---|---|---|---|
| HMR_2913 | Carnitine shuttle (endoplasmic reticular) | 0.0253 | 0.0002 | 0.0253 | 0.0002 | 0.00E+00 | 0.0003 | 0.4999 |
| HMR_2914 | Carnitine shuttle (endoplasmic reticular) | -0.0255 | 0.0002 | -0.0255 | 0.0002 | 0.00E+00 | -1.00E-04 | 0.5 |
| HMR_2915 | Carnitine shuttle (endoplasmic reticular) | -0.0254 | 0.0002 | -0.0254 | 0.0002 | 0.00E+00 | 0.0005 | 0.4998 |
| HMR_2916 | Carnitine shuttle (endoplasmic reticular) | -0.0253 | 0.0002 | -0.0253 | 0.0002 | 0.00E+00 | -0.0003 | 0.5001 |
| HMR_2917 | Carnitine shuttle (endoplasmic reticular) | 0.0255 | 0.0002 | 0.0255 | 0.0002 | 0.00E+00 | 1.00E-04 | 0.5 |
| HMR_2918 | Carnitine shuttle (endoplasmic reticular) | -0.0255 | 0.0002 | -0.0255 | 0.0002 | 0.00E+00 | 0.0001 | 0.4999 |
| HMR_2919 | Carnitine shuttle (endoplasmic reticular) | -0.0253 | 0.0002 | -0.0253 | 0.0002 | 0.00E+00 | -0.0003 | 0.5001 |
| HMR_2920 | Carnitine shuttle (endoplasmic reticular) | 0.0255 | 0.0002 | 0.0255 | 0.0002 | 0.00E+00 | 1.00E-04 | 0.5 |
| HMR_2921 | Carnitine shuttle (endoplasmic reticular) | 0.0253 | 0.0002 | 0.0253 | 0.0002 | 0.00E+00 | -0.0005 | 0.5002 |
| HMR_2922 | Carnitine shuttle (endoplasmic reticular) | 0.0253 | 0.0002 | 0.0253 | 0.0002 | 0.00E+00 | 0.0003 | 0.4999 |
| HMR_2923 | Carnitine shuttle (endoplasmic reticular) | -0.0255 | 0.0002 | -0.0255 | 0.0002 | 0.00E+00 | -1.00E-04 | 0.5 |
| HMR_2924 | Carnitine shuttle (endoplasmic reticular) | 0.0255 | 0.0002 | 0.0255 | 0.0002 | 0.00E+00 | -0.0002 | 0.5001 |
| HMR_2925 | Carnitine shuttle (endoplasmic reticular) | 0.0253 | 0.0002 | 0.0253 | 0.0002 | 0.00E+00 | 0.0003 | 0.4999 |
| HMR_2926 | Carnitine shuttle (endoplasmic reticular) | -0.0255 | 0.0002 | -0.0255 | 0.0002 | 0.00E+00 | -1.00E-04 | 0.5 |
| HMR_2927 | Carnitine shuttle (endoplasmic reticular) | 0.0255 | 0.0002 | 0.0255 | 0.0002 | 0.00E+00 | -0.0002 | 0.5001 |
| HMR_2928 | Carnitine shuttle (endoplasmic reticular) | 0.0253 | 0.0002 | 0.0253 | 0.0002 | 0.00E+00 | 0.0003 | 0.4999 |
| HMR_2929 | Carnitine shuttle (endoplasmic reticular) | -0.0255 | 0.0002 | -0.0255 | 0.0002 | 0.00E+00 | -1.00E-04 | 0.5 |
| HMR_2930 | Carnitine shuttle (endoplasmic reticular) | 0.0253 | 0.0002 | 0.0254 | 0.0002 | 0.00E+00 | -0.0005 | 0.5002 |
| HMR_2931 | Carnitine shuttle (endoplasmic reticular) | 0.0253 | 0.0002 | 0.0253 | 0.0002 | 0.00E+00 | 0.0003 | 0.4999 |
| HMR_2932 | Carnitine shuttle (endoplasmic reticular) | -0.0255 | 0.0002 | -0.0255 | 0.0002 | 0.00E+00 | -1.00E-04 | 0.5 |
| HMR_0001 | Acylglycerides metabolism | 0.00E+00 | 0.00E+00 | 0.00E+00 | 0.00E+00 | 0.00E+00 | 0.0333 | 0.4867 |
| HMR_0002 | Acylglycerides metabolism | 4.9734 | 3.5105 | 4.8234 | 3.3171 | 0.1501 | 0.0574 | 0.4771 |
| HMR_0003 | Acylglycerides metabolism | 4.9723 | 3.5123 | 4.8222 | 3.3188 | 0.1501 | 0.0574 | 0.4771 |
| HMR_0005 | Acylglycerides metabolism | 4.9708 | 3.5288 | 4.8208 | 3.3355 | 0.15 | 0.0573 | 0.4772 |
| HMR_0604 | Acylglycerides metabolism | 2.7628 | 6.5883 | 2.6424 | 6.0658 | 0.1204 | 0.0338 | 0.4865 |
| HMR_0605 | Acylglycerides metabolism | 5.6941 | 3.3337 | 5.7547 | 3.4229 | 0.0606 | -0.0233 | 0.5093 |
| HMR_0665 | Acylglycerides metabolism | 2.0167 | 1.8786 | 2.0481 | 1.9381 | 0.0314 | -0.0161 | 0.5064 |
| HMR_0667 | Acylglycerides metabolism | 2.969 | 2.5525 | 2.9701 | 2.5572 | 0.0011 | -0.0005 | 0.5002 |
| HMR_0668 | Acylglycerides metabolism | 4.6001 | 3.2107 | 4.5661 | 3.1844 | 0.034 | 0.0135 | 0.4946 |
| HMR_0669 | Acylglycerides metabolism | 2.5127 | 1.8791 | 2.4839 | 1.8391 | 0.0288 | 0.0149 | 0.494 |
| HMR_0670 | Acylglycerides metabolism | 2.6077 | 1.8826 | 2.5878 | 1.8551 | 0.0198 | 0.0103 | 0.4959 |
| HMR_0671 | Acylglycerides metabolism | 2.8665 | 1.8896 | 2.8264 | 1.8431 | 0.0401 | 0.0207 | 0.4917 |
| HMR_0672 | Acylglycerides metabolism | 5.3826 | 5.0673 | 5.231 | 4.8156 | 0.1515 | 0.0482 | 0.4808 |
| HMR_0679 | Acylglycerides metabolism | 1.7529 | 1.8888 | 1.7638 | 1.907 | 0.0109 | -0.0056 | 0.5022 |
| HMR_0680 | Acylglycerides metabolism | 1.3212 | 1.1758 | 1.3062 | 1.1513 | 0.015 | 0.0099 | 0.4961 |
| HMR_0681 | Acylglycerides metabolism | 1.434 | 1.223 | 1.4243 | 1.2061 | 0.0097 | 0.0063 | 0.4975 |
| HMR_0682 | Acylglycerides metabolism | 1.6446 | 1.3055 | 1.6188 | 1.272 | 0.0258 | 0.0161 | 0.4936 |
| HMR_0683 | Acylglycerides metabolism | 2.4699 | 3.01 | 2.4524 | 2.976 | 0.0176 | 0.0072 | 0.4971 |
| HMR_0684 | Acylglycerides metabolism | 3.627 | 4.1866 | 3.5191 | 3.9695 | 0.1079 | 0.0378 | 0.4849 |
| HMR_7588 | Acylglycerides metabolism | 1.1243 | 1.0534 | 1.1196 | 1.0465 | 0.0047 | 0.0032 | 0.4987 |
| HMR_3053 | Beta oxidation of even-chain fatty acids (peroxisomal) | 0.6731 | 0.1555 | 0.6914 | 0.1639 | 0.0183 | -0.0324 | 0.5129 |
| HMR_3054 | Beta oxidation of even-chain fatty acids (peroxisomal) | 0.514 | 0.1201 | 0.5297 | 0.1279 | 0.0158 | -0.0317 | 0.5126 |
| HMR_3055 | Beta oxidation of even-chain fatty acids (peroxisomal) | 0.1826 | 0.0165 | 0.1872 | 0.0173 | 0.0046 | -0.0249 | 0.5099 |
| HMR_3057 | Beta oxidation of even-chain fatty acids (peroxisomal) | 3.5715 | 2.6724 | 3.6969 | 2.8622 | 0.1254 | -0.0533 | 0.5213 |

| | | | | | | | | |
|---|---|---|---|---|---|---|---|---|
| HMR_3058 | Beta oxidation of even-chain fatty acids (peroxisomal) | 3.5776 | 2.6753 | 3.7024 | 2.8651 | 0.1247 | -0.053 | 0.5211 |
| HMR_3059 | Beta oxidation of even-chain fatty acids (peroxisomal) | 3.5836 | 2.6801 | 3.7078 | 2.87 | 0.1241 | -0.0527 | 0.521 |
| HMR_3060 | Beta oxidation of even-chain fatty acids (peroxisomal) | 3.5894 | 2.6871 | 3.7129 | 2.877 | 0.1235 | -0.0524 | 0.5209 |
| HMR_3062 | Beta oxidation of even-chain fatty acids (peroxisomal) | 1.2782 | 0.2615 | 1.3282 | 0.2794 | 0.0501 | -0.0681 | 0.5271 |
| HMR_3063 | Beta oxidation of even-chain fatty acids (peroxisomal) | 1.2837 | 0.2586 | 1.3332 | 0.2765 | 0.0494 | -0.0676 | 0.5269 |
| HMR_3064 | Beta oxidation of even-chain fatty acids (peroxisomal) | 1.2892 | 0.2566 | 1.3381 | 0.2744 | 0.0489 | -0.0671 | 0.5267 |
| HMR_3065 | Beta oxidation of even-chain fatty acids (peroxisomal) | 1.2943 | 0.2552 | 1.3426 | 0.273 | 0.0483 | -0.0664 | 0.5265 |
| HMR_3066 | Beta oxidation of even-chain fatty acids (peroxisomal) | 1.2991 | 0.2546 | 1.3468 | 0.2724 | 0.0477 | -0.0657 | 0.5262 |
| HMR_3067 | Beta oxidation of even-chain fatty acids (peroxisomal) | 1.3041 | 0.2547 | 1.3512 | 0.2723 | 0.0471 | -0.0649 | 0.5259 |
| HMR_3068 | Beta oxidation of even-chain fatty acids (peroxisomal) | 1.309 | 0.2554 | 1.3555 | 0.2729 | 0.0465 | -0.064 | 0.5255 |
| HMR_3069 | Beta oxidation of even-chain fatty acids (peroxisomal) | 1.3136 | 0.2567 | 1.3596 | 0.2742 | 0.046 | -0.0631 | 0.5252 |
| HMR_3070 | Beta oxidation of even-chain fatty acids (peroxisomal) | 1.3179 | 0.2586 | 1.3634 | 0.2761 | 0.0454 | -0.0621 | 0.5248 |
| HMR_3071 | Beta oxidation of even-chain fatty acids (peroxisomal) | 1.3225 | 0.2612 | 1.3673 | 0.2786 | 0.0448 | -0.061 | 0.5243 |
| HMR_3072 | Beta oxidation of even-chain fatty acids (peroxisomal) | 1.327 | 0.2645 | 1.3713 | 0.2818 | 0.0442 | -0.0599 | 0.5239 |
| HMR_3073 | Beta oxidation of even-chain fatty acids (peroxisomal) | 1.3312 | 0.2684 | 1.3749 | 0.2857 | 0.0437 | -0.0587 | 0.5234 |
| HMR_3074 | Beta oxidation of even-chain fatty acids (peroxisomal) | 1.3351 | 0.2731 | 1.3782 | 0.2903 | 0.0431 | -0.0574 | 0.5229 |
| HMR_3075 | Beta oxidation of even-chain fatty acids (peroxisomal) | 1.3392 | 0.2785 | 1.3817 | 0.2956 | 0.0425 | -0.0561 | 0.5224 |
| HMR_3076 | Beta oxidation of even-chain fatty acids (peroxisomal) | 1.3432 | 0.2847 | 1.3851 | 0.3018 | 0.042 | -0.0548 | 0.5218 |
| HMR_3077 | Beta oxidation of even-chain fatty acids (peroxisomal) | 1.3468 | 0.2919 | 1.3882 | 0.3089 | 0.0414 | -0.0534 | 0.5213 |
| HMR_3078 | Beta oxidation of even-chain fatty acids (peroxisomal) | 0.894 | 0.1764 | 0.9395 | 0.1926 | 0.0456 | -0.075 | 0.5299 |
| HMR_3079 | Beta oxidation of even-chain fatty acids (peroxisomal) | 0.8964 | 0.1699 | 0.9414 | 0.1861 | 0.0451 | -0.0756 | 0.5301 |
| HMR_3080 | Beta oxidation of even-chain fatty acids (peroxisomal) | 0.8977 | 0.1655 | 0.9424 | 0.1816 | 0.0447 | -0.0759 | 0.5302 |
| HMR_3081 | Beta oxidation of even-chain fatty acids (peroxisomal) | 0.8979 | 0.1629 | 0.9422 | 0.179 | 0.0443 | -0.0758 | 0.5302 |
| HMR_3082 | Beta oxidation of even-chain fatty acids (peroxisomal) | 0.8969 | 0.162 | 0.941 | 0.1782 | 0.0441 | -0.0755 | 0.5301 |
| HMR_3083 | Beta oxidation of even-chain fatty acids (peroxisomal) | 0.8954 | 0.1629 | 0.9391 | 0.1791 | 0.0438 | -0.0748 | 0.5298 |
| HMR_3084 | Beta oxidation of even-chain fatty acids (peroxisomal) | 0.8929 | 0.1655 | 0.9365 | 0.1817 | 0.0436 | -0.0739 | 0.5295 |
| HMR_3085 | Beta oxidation of even-chain fatty acids (peroxisomal) | 0.8893 | 0.1701 | 0.9327 | 0.1863 | 0.0434 | -0.0727 | 0.529 |
| HMR_3086 | Beta oxidation of even-chain fatty acids (peroxisomal) | 0.8843 | 0.1768 | 0.9276 | 0.1931 | 0.0433 | -0.0712 | 0.5284 |
| HMR_3087 | Beta oxidation of even-chain fatty acids (peroxisomal) | 2.9086 | 0.5428 | 3.0214 | 0.5814 | 0.1129 | -0.1065 | 0.5424 |
| HMR_3088 | Beta oxidation of even-chain fatty acids (peroxisomal) | 2.9029 | 0.5502 | 3.0157 | 0.5888 | 0.1129 | -0.1057 | 0.5421 |
| HMR_3089 | Beta oxidation of even-chain fatty acids (peroxisomal) | 2.8972 | 0.5575 | 3.01 | 0.5962 | 0.1128 | -0.105 | 0.5418 |
| HMR_3090 | Beta oxidation of even-chain fatty acids (peroxisomal) | 2.8915 | 0.5648 | 3.0043 | 0.6035 | 0.1128 | -0.1043 | 0.5415 |
| HMR_3091 | Beta oxidation of even-chain fatty acids (peroxisomal) | 5.9818 | 1.0845 | 6.149 | 1.1495 | 0.1672 | -0.1118 | 0.5445 |
| HMR_3092 | Beta oxidation of even-chain fatty acids (peroxisomal) | 5.977 | 1.0894 | 6.1441 | 1.1545 | 0.1671 | -0.1116 | 0.5444 |
| HMR_3093 | Beta oxidation of even-chain fatty acids (peroxisomal) | 5.9722 | 1.0942 | 6.1392 | 1.1595 | 0.167 | -0.1112 | 0.5443 |
| HMR_3094 | Beta oxidation of even-chain fatty acids (peroxisomal) | 3.1406 | 1.616 | 3.1233 | 1.6342 | 0.0173 | 0.0096 | 0.4962 |
| HMR_3095 | Beta oxidation of even-chain fatty acids (peroxisomal) | 3.1362 | 1.6135 | 3.1187 | 1.631 | 0.0175 | 0.0097 | 0.4961 |
| HMR_3096 | Beta oxidation of even-chain fatty acids (peroxisomal) | 3.1319 | 1.6121 | 3.1143 | 1.629 | 0.0176 | 0.0098 | 0.4961 |
| HMR_3097 | Beta oxidation of even-chain fatty acids (peroxisomal) | 3.1274 | 1.6118 | 3.1096 | 1.6282 | 0.0178 | 0.0099 | 0.4961 |
| HMR_3098 | Beta oxidation of even-chain fatty acids (peroxisomal) | 3.1228 | 1.6126 | 3.1048 | 1.6285 | 0.018 | 0.01 | 0.496 |
| HMR_3099 | Beta oxidation of even-chain fatty acids (peroxisomal) | 3.1185 | 1.6145 | 3.1002 | 1.63 | 0.0183 | 0.0102 | 0.4959 |
| HMR_3100 | Beta oxidation of even-chain fatty acids (peroxisomal) | 3.1143 | 1.6176 | 3.0958 | 1.6328 | 0.0185 | 0.0103 | 0.4959 |
| HMR_3101 | Beta oxidation of even-chain fatty acids (peroxisomal) | 3.11 | 1.6218 | 3.0912 | 1.6367 | 0.0188 | 0.0104 | 0.4958 |
| HMR_3102 | Beta oxidation of even-chain fatty acids (peroxisomal) | 3.1055 | 1.6272 | 3.0863 | 1.642 | 0.0191 | 0.0106 | 0.4958 |

| | | | | | | | | |
|---|---|---|---|---|---|---|---|---|
| HMR_3103 | Beta oxidation of even-chain fatty acids (peroxisomal) | 3.1013 | 1.6338 | 3.0818 | 1.6485 | 0.0195 | 0.0108 | 0.4957 |
| HMR_3104 | Beta oxidation of even-chain fatty acids (peroxisomal) | 3.0972 | 1.6416 | 3.0773 | 1.6563 | 0.0199 | 0.0109 | 0.4956 |
| HMR_3106 | Beta oxidation of even-chain fatty acids (peroxisomal) | -101.876 | 159460.7364 | -196.9886 | 138548.0494 | 95.1126 | 0.1742 | 0.4308 |
| HMR_3056 | Beta oxidation of odd-chain fatty acids (peroxisomal) | 0.4011 | 0.0569 | 0.4096 | 0.0588 | 0.0085 | -0.025 | 0.51 |
| HMR_3326 | Beta oxidation of unsaturated fatty acids (n-9) (peroxisomal) | 0.7968 | 0.1332 | 0.8256 | 0.1428 | 0.0288 | -0.0548 | 0.5218 |
| HMR_3327 | Beta oxidation of unsaturated fatty acids (n-9) (peroxisomal) | 0.7951 | 0.1335 | 0.8236 | 0.1431 | 0.0285 | -0.0543 | 0.5216 |
| HMR_3328 | Beta oxidation of unsaturated fatty acids (n-9) (peroxisomal) | 0.7922 | 0.1357 | 0.8206 | 0.1453 | 0.0284 | -0.0535 | 0.5214 |
| HMR_3329 | Beta oxidation of unsaturated fatty acids (n-9) (peroxisomal) | 0.7878 | 0.14 | 0.8161 | 0.1496 | 0.0283 | -0.0525 | 0.5209 |
| HMR_3330 | Beta oxidation of unsaturated fatty acids (n-9) (peroxisomal) | 1.4362 | 0.2317 | 1.4914 | 0.2494 | 0.0551 | -0.0795 | 0.5317 |
| HMR_3331 | Beta oxidation of unsaturated fatty acids (n-9) (peroxisomal) | 1.4306 | 0.2316 | 1.4856 | 0.2493 | 0.055 | -0.0793 | 0.5316 |
| HMR_3332 | Beta oxidation of unsaturated fatty acids (n-9) (peroxisomal) | 1.4251 | 0.2316 | 1.4801 | 0.2492 | 0.055 | -0.0793 | 0.5316 |
| HMR_3333 | Beta oxidation of unsaturated fatty acids (n-9) (peroxisomal) | 1.4195 | 0.2316 | 1.4744 | 0.2492 | 0.0549 | -0.0791 | 0.5315 |
| HMR_3334 | Beta oxidation of unsaturated fatty acids (n-9) (peroxisomal) | 1.4139 | 0.2317 | 1.4687 | 0.2494 | 0.0548 | -0.079 | 0.5315 |
| HMR_3335 | Beta oxidation of unsaturated fatty acids (n-9) (peroxisomal) | 1.4086 | 0.2319 | 1.4633 | 0.2496 | 0.0547 | -0.0788 | 0.5314 |
| HMR_3336 | Beta oxidation of unsaturated fatty acids (n-9) (peroxisomal) | 1.4036 | 0.2323 | 1.4581 | 0.25 | 0.0546 | -0.0786 | 0.5313 |
| HMR_3337 | Beta oxidation of unsaturated fatty acids (n-9) (peroxisomal) | 1.3984 | 0.2328 | 1.4528 | 0.2505 | 0.0545 | -0.0783 | 0.5312 |
| HMR_3338 | Beta oxidation of unsaturated fatty acids (n-9) (peroxisomal) | 1.3931 | 0.2334 | 1.4474 | 0.2512 | 0.0544 | -0.0781 | 0.5311 |
| HMR_3339 | Beta oxidation of unsaturated fatty acids (n-9) (peroxisomal) | 1.3882 | 0.2342 | 1.4424 | 0.252 | 0.0542 | -0.0777 | 0.531 |
| HMR_3340 | Beta oxidation of unsaturated fatty acids (n-9) (peroxisomal) | 1.3834 | 0.2351 | 1.4375 | 0.2531 | 0.0541 | -0.0774 | 0.5309 |
| HMR_3341 | Beta oxidation of unsaturated fatty acids (n-9) (peroxisomal) | 1.3786 | 0.2363 | 1.4325 | 0.2543 | 0.054 | -0.077 | 0.5307 |
| HMR_3342 | Beta oxidation of unsaturated fatty acids (n-9) (peroxisomal) | 1.3735 | 0.2377 | 1.4274 | 0.2558 | 0.0538 | -0.0766 | 0.5305 |
| HMR_3343 | Beta oxidation of unsaturated fatty acids (n-9) (peroxisomal) | 1.3689 | 0.2393 | 1.4226 | 0.2575 | 0.0537 | -0.0761 | 0.5303 |
| HMR_3344 | Beta oxidation of unsaturated fatty acids (n-9) (peroxisomal) | 1.3643 | 0.2412 | 1.4179 | 0.2595 | 0.0535 | -0.0757 | 0.5302 |
| HMR_3345 | Beta oxidation of unsaturated fatty acids (n-9) (peroxisomal) | 1.3596 | 0.2434 | 1.413 | 0.2618 | 0.0534 | -0.0751 | 0.5299 |
| HMR_3346 | Beta oxidation of unsaturated fatty acids (n-9) (peroxisomal) | 1.3547 | 0.2459 | 1.408 | 0.2645 | 0.0533 | -0.0745 | 0.5297 |
| HMR_3347 | Beta oxidation of unsaturated fatty acids (n-9) (peroxisomal) | 1.3502 | 0.2489 | 1.4033 | 0.2675 | 0.0531 | -0.0739 | 0.5294 |
| HMR_3348 | Beta oxidation of unsaturated fatty acids (n-9) (peroxisomal) | 1.3457 | 0.2522 | 1.3986 | 0.271 | 0.0529 | -0.0732 | 0.5292 |
| HMR_3349 | Beta oxidation of unsaturated fatty acids (n-9) (peroxisomal) | 1.3409 | 0.2561 | 1.3937 | 0.275 | 0.0528 | -0.0724 | 0.5289 |
| HMR_3350 | Beta oxidation of unsaturated fatty acids (n-9) (peroxisomal) | 1.3359 | 0.2605 | 1.3886 | 0.2795 | 0.0527 | -0.0717 | 0.5286 |
| HMR_3351 | Beta oxidation of unsaturated fatty acids (n-9) (peroxisomal) | 1.3312 | 0.2655 | 1.3837 | 0.2847 | 0.0525 | -0.0708 | 0.5282 |
| HMR_3352 | Beta oxidation of unsaturated fatty acids (n-9) (peroxisomal) | 1.3264 | 0.2713 | 1.3788 | 0.2905 | 0.0524 | -0.0699 | 0.5279 |
| HMR_3353 | Beta oxidation of unsaturated fatty acids (n-9) (peroxisomal) | 1.3213 | 0.2779 | 1.3735 | 0.2972 | 0.0523 | -0.0689 | 0.5275 |
| HMR_3355 | Beta oxidation of unsaturated fatty acids (n-9) (peroxisomal) | -1.3157 | 0.2854 | -1.3679 | 0.3047 | 0.0522 | 0.0679 | 0.4729 |
| HMR_3364 | Beta oxidation of unsaturated fatty acids (n-9) (peroxisomal) | 0.7225 | 0.0836 | 0.7416 | 0.0879 | 0.0191 | -0.0462 | 0.5184 |
| HMR_3365 | Beta oxidation of unsaturated fatty acids (n-9) (peroxisomal) | 0.7239 | 0.0843 | 0.7427 | 0.0887 | 0.0188 | -0.0453 | 0.5181 |
| HMR_3367 | Beta oxidation of unsaturated fatty acids (n-9) (peroxisomal) | 0.7248 | 0.0856 | 0.7434 | 0.0899 | 0.0186 | -0.0444 | 0.5177 |
| HMR_3368 | Beta oxidation of unsaturated fatty acids (n-9) (peroxisomal) | 0.7249 | 0.0874 | 0.7433 | 0.0917 | 0.0184 | -0.0434 | 0.5173 |
| HMR_3369 | Beta oxidation of unsaturated fatty acids (n-9) (peroxisomal) | 0.7243 | 0.0898 | 0.7424 | 0.0942 | 0.0181 | -0.0423 | 0.5169 |
| HMR_3370 | Beta oxidation of unsaturated fatty acids (n-9) (peroxisomal) | 0.7233 | 0.093 | 0.7411 | 0.0975 | 0.0179 | -0.041 | 0.5163 |
| HMR_3372 | Beta oxidation of unsaturated fatty acids (n-9) (peroxisomal) | 0.7215 | 0.0972 | 0.7392 | 0.1017 | 0.0177 | -0.0397 | 0.5158 |
| HMR_3373 | Beta oxidation of unsaturated fatty acids (n-9) (peroxisomal) | 0.7186 | 0.1027 | 0.7361 | 0.1073 | 0.0175 | -0.0382 | 0.5152 |
| HMR_3375 | Beta oxidation of unsaturated fatty acids (n-9) (peroxisomal) | -0.7143 | 0.1099 | -0.7317 | 0.1145 | 0.0174 | 0.0367 | 0.4854 |
| HMR_3480 | Beta oxidation of phytanic acid (peroxisomal) | -0.8372 | 0.1163 | -1.0723 | 0.1808 | 0.2351 | 0.4314 | 0.3331 |
| HMR_3481 | Beta oxidation of phytanic acid (peroxisomal) | 0.8351 | 0.1104 | 1.0706 | 0.1744 | 0.2355 | -0.4414 | 0.6705 |

| | | | | | | | | |
|---|---|---|---|---|---|---|---|---|
| HMR_3484 | Beta oxidation of phytanic acid (peroxisomal) | 0.8323 | 0.1052 | 1.0685 | 0.1687 | 0.2362 | -0.4513 | 0.6741 |
| HMR_3486 | Beta oxidation of phytanic acid (peroxisomal) | 0.8324 | 0.1016 | 1.0696 | 0.1645 | 0.2371 | -0.4597 | 0.6771 |
| HMR_3488 | Beta oxidation of phytanic acid (peroxisomal) | 0.8353 | 0.1003 | 1.0745 | 0.1624 | 0.2392 | -0.4666 | 0.6796 |
| HMR_3489 | Beta oxidation of phytanic acid (peroxisomal) | 0.8381 | 0.0993 | 1.0794 | 0.1607 | 0.2413 | -0.4733 | 0.682 |
| HMR_3498 | Beta oxidation of phytanic acid (peroxisomal) | 0.8408 | 0.0987 | 1.084 | 0.1596 | 0.2432 | -0.4784 | 0.6838 |
| HMR_3501 | Beta oxidation of phytanic acid (peroxisomal) | 0.8437 | 0.0983 | 1.0888 | 0.1589 | 0.245 | -0.4831 | 0.6855 |
| HMR_3503 | Beta oxidation of phytanic acid (peroxisomal) | 0.8465 | 0.0981 | 1.0936 | 0.1584 | 0.247 | -0.4878 | 0.6872 |
| HMR_3505 | Beta oxidation of phytanic acid (peroxisomal) | 0.849 | 0.098 | 1.0981 | 0.1581 | 0.2492 | -0.4924 | 0.6888 |
| HMR_3506 | Beta oxidation of phytanic acid (peroxisomal) | 0.8515 | 0.0979 | 1.1025 | 0.158 | 0.251 | -0.4962 | 0.6901 |
| HMR_3508 | Beta oxidation of phytanic acid (peroxisomal) | 0.8543 | 0.098 | 1.1072 | 0.1582 | 0.2529 | -0.4997 | 0.6914 |
| HMR_3509 | Beta oxidation of phytanic acid (peroxisomal) | 0.8569 | 0.0981 | 1.1119 | 0.1585 | 0.255 | -0.5034 | 0.6927 |
| HMR_3510 | Beta oxidation of phytanic acid (peroxisomal) | 0.8593 | 0.0983 | 1.1165 | 0.159 | 0.2572 | -0.5069 | 0.6939 |
| HMR_3511 | Beta oxidation of phytanic acid (peroxisomal) | 0.8613 | 0.0986 | 1.1208 | 0.1596 | 0.2594 | -0.5105 | 0.6952 |
| HMR_3512 | Beta oxidation of phytanic acid (peroxisomal) | 0.863 | 0.0989 | 1.1249 | 0.1604 | 0.2618 | -0.5141 | 0.6964 |
| HMR_3513 | Beta oxidation of phytanic acid (peroxisomal) | 0.865 | 0.0993 | 1.1292 | 0.1614 | 0.2643 | -0.5175 | 0.6976 |
| HMR_3514 | Beta oxidation of phytanic acid (peroxisomal) | 0.8668 | 0.0998 | 1.1337 | 0.1625 | 0.2669 | -0.5211 | 0.6988 |
| HMR_3515 | Beta oxidation of phytanic acid (peroxisomal) | 0.8684 | 0.1003 | 1.138 | 0.1638 | 0.2696 | -0.5245 | 0.7 |
| HMR_3517 | Beta oxidation of phytanic acid (peroxisomal) | -0.8701 | 0.1009 | -1.1421 | 0.1653 | 0.272 | 0.5272 | 0.299 |
| HMR_3301 | Beta oxidation of di-unsaturated fatty acids (n-6) (peroxisomal) | 1.902 | 0.6889 | 1.9501 | 0.7203 | 0.0482 | -0.0406 | 0.5162 |
| HMR_3302 | Beta oxidation of di-unsaturated fatty acids (n-6) (peroxisomal) | 1.9056 | 0.683 | 1.9534 | 0.7144 | 0.0478 | -0.0404 | 0.5161 |
| HMR_3304 | Beta oxidation of di-unsaturated fatty acids (n-6) (peroxisomal) | 1.9089 | 0.6787 | 1.9565 | 0.71 | 0.0475 | -0.0403 | 0.5161 |
| HMR_3305 | Beta oxidation of di-unsaturated fatty acids (n-6) (peroxisomal) | 1.9119 | 0.676 | 1.9591 | 0.7072 | 0.0472 | -0.0401 | 0.516 |
| HMR_3306 | Beta oxidation of di-unsaturated fatty acids (n-6) (peroxisomal) | 1.9145 | 0.6748 | 1.9614 | 0.706 | 0.0469 | -0.0399 | 0.5159 |
| HMR_3307 | Beta oxidation of di-unsaturated fatty acids (n-6) (peroxisomal) | 3.0649 | 0.6616 | 3.1236 | 0.6911 | 0.0587 | -0.0505 | 0.5201 |
| HMR_3309 | Beta oxidation of di-unsaturated fatty acids (n-6) (peroxisomal) | 3.0679 | 0.6599 | 3.1263 | 0.6893 | 0.0584 | -0.0503 | 0.52 |
| HMR_3310 | Beta oxidation of di-unsaturated fatty acids (n-6) (peroxisomal) | 3.0709 | 0.6581 | 3.129 | 0.6875 | 0.0581 | -0.0501 | 0.52 |
| HMR_3311 | Beta oxidation of di-unsaturated fatty acids (n-6) (peroxisomal) | 3.0739 | 0.6563 | 3.1317 | 0.6856 | 0.0577 | -0.0498 | 0.5199 |
| HMR_3312 | Beta oxidation of di-unsaturated fatty acids (n-6) (peroxisomal) | 3.0775 | 0.6543 | 3.1349 | 0.6836 | 0.0573 | -0.0496 | 0.5198 |
| HMR_3314 | Beta oxidation of di-unsaturated fatty acids (n-6) (peroxisomal) | 3.0814 | 0.6523 | 3.1384 | 0.6815 | 0.057 | -0.0493 | 0.5197 |
| HMR_3315 | Beta oxidation of di-unsaturated fatty acids (n-6) (peroxisomal) | 3.0853 | 0.6502 | 3.1419 | 0.6794 | 0.0566 | -0.0491 | 0.5196 |
| HMR_3316 | Beta oxidation of di-unsaturated fatty acids (n-6) (peroxisomal) | 3.0892 | 0.648 | 3.1454 | 0.6771 | 0.0562 | -0.0488 | 0.5195 |
| HMR_3317 | Beta oxidation of di-unsaturated fatty acids (n-6) (peroxisomal) | 3.093 | 0.6457 | 3.1489 | 0.6748 | 0.0559 | -0.0486 | 0.5194 |
| HMR_3319 | Beta oxidation of di-unsaturated fatty acids (n-6) (peroxisomal) | 3.0972 | 0.6434 | 3.1527 | 0.6724 | 0.0555 | -0.0484 | 0.5193 |
| HMR_3320 | Beta oxidation of di-unsaturated fatty acids (n-6) (peroxisomal) | 3.1013 | 0.6409 | 3.1565 | 0.6699 | 0.0551 | -0.0482 | 0.5192 |
| HMR_3321 | Beta oxidation of di-unsaturated fatty acids (n-6) (peroxisomal) | 3.1055 | 0.6384 | 3.1603 | 0.6674 | 0.0548 | -0.0479 | 0.5191 |
| HMR_3322 | Beta oxidation of di-unsaturated fatty acids (n-6) (peroxisomal) | 3.1005 | 0.6462 | 3.1551 | 0.6752 | 0.0546 | -0.0475 | 0.519 |
| HMR_3323 | Beta oxidation of di-unsaturated fatty acids (n-6) (peroxisomal) | -3.0954 | 0.6539 | -3.1499 | 0.683 | 0.0545 | 0.0472 | 0.4812 |
| HMR_3107 | Beta oxidation of even-chain fatty acids (mitochondrial) | 3.6062 | 2.8624 | 3.9461 | 3.4176 | 0.3399 | -0.1356 | 0.5539 |
| HMR_3108 | Beta oxidation of even-chain fatty acids (mitochondrial) | 3.6125 | 2.8647 | 3.9517 | 3.42 | 0.3391 | -0.1353 | 0.5538 |
| HMR_3109 | Beta oxidation of even-chain fatty acids (mitochondrial) | 3.6184 | 2.8695 | 3.9568 | 3.4248 | 0.3384 | -0.1349 | 0.5537 |
| HMR_3110 | Beta oxidation of even-chain fatty acids (mitochondrial) | 3.624 | 2.8767 | 3.9617 | 3.4321 | 0.3377 | -0.1344 | 0.5535 |
| HMR_3111 | Beta oxidation of even-chain fatty acids (mitochondrial) | 3.5248 | 3.3498 | 3.8513 | 3.9763 | 0.3265 | -0.1206 | 0.548 |
| HMR_3112 | Beta oxidation of even-chain fatty acids (mitochondrial) | 3.5303 | 3.3478 | 3.856 | 3.9744 | 0.3257 | -0.1204 | 0.5479 |
| HMR_3113 | Beta oxidation of even-chain fatty acids (mitochondrial) | 3.5351 | 3.3496 | 3.8601 | 3.9763 | 0.325 | -0.1201 | 0.5478 |

| | | | | | | | | |
|---|---|---|---|---|---|---|---|---|
| HMR_3114 | Beta oxidation of even-chain fatty acids (mitochondrial) | 3.5394 | 3.3554 | 3.8636 | 3.982 | 0.3242 | -0.1197 | 0.5476 |
| HMR_3115 | Beta oxidation of even-chain fatty acids (mitochondrial) | 3.6349 | 4.0082 | 3.9724 | 4.7824 | 0.3375 | -0.1138 | 0.5453 |
| HMR_3116 | Beta oxidation of even-chain fatty acids (mitochondrial) | 3.6388 | 4.0033 | 3.9755 | 4.7775 | 0.3367 | -0.1136 | 0.5452 |
| HMR_3117 | Beta oxidation of even-chain fatty acids (mitochondrial) | 3.642 | 4.0034 | 3.9779 | 4.7775 | 0.336 | -0.1134 | 0.5451 |
| HMR_3118 | Beta oxidation of even-chain fatty acids (mitochondrial) | 3.6445 | 4.0083 | 3.9797 | 4.7825 | 0.3353 | -0.1131 | 0.545 |
| HMR_3121 | Beta oxidation of even-chain fatty acids (mitochondrial) | 2.4771 | 1.2026 | 2.8773 | 1.5276 | 0.4002 | -0.2422 | 0.5957 |
| HMR_3122 | Beta oxidation of even-chain fatty acids (mitochondrial) | 2.4793 | 1.1951 | 2.8788 | 1.5198 | 0.3995 | -0.2425 | 0.5958 |
| HMR_3123 | Beta oxidation of even-chain fatty acids (mitochondrial) | 2.4812 | 1.1894 | 2.88 | 1.5136 | 0.3989 | -0.2426 | 0.5959 |
| HMR_3125 | Beta oxidation of even-chain fatty acids (mitochondrial) | 2.4826 | 1.1855 | 2.881 | 1.5089 | 0.3983 | -0.2427 | 0.5959 |
| HMR_3128 | Beta oxidation of even-chain fatty acids (mitochondrial) | 1.0052 | 0.2808 | 1.1269 | 0.3523 | 0.1218 | -0.153 | 0.5608 |
| HMR_3129 | Beta oxidation of even-chain fatty acids (mitochondrial) | 1.005 | 0.2763 | 1.1264 | 0.3477 | 0.1214 | -0.1537 | 0.5611 |
| HMR_3130 | Beta oxidation of even-chain fatty acids (mitochondrial) | 1.0027 | 0.2757 | 1.1239 | 0.3471 | 0.1212 | -0.1536 | 0.561 |
| HMR_3132 | Beta oxidation of even-chain fatty acids (mitochondrial) | 0.9984 | 0.2791 | 1.1196 | 0.3505 | 0.1212 | -0.1528 | 0.5607 |
| HMR_3135 | Beta oxidation of even-chain fatty acids (mitochondrial) | 1.2766 | 0.4744 | 1.4267 | 0.5699 | 0.1501 | -0.1469 | 0.5584 |
| HMR_3136 | Beta oxidation of even-chain fatty acids (mitochondrial) | 13.0152 | 4.1613 | 14.4858 | 5.1174 | 1.4706 | -0.4828 | 0.6854 |
| HMR_3137 | Beta oxidation of even-chain fatty acids (mitochondrial) | 13.0077 | 4.1705 | 14.4787 | 5.1265 | 1.471 | -0.4824 | 0.6853 |
| HMR_3139 | Beta oxidation of even-chain fatty acids (mitochondrial) | 13.0002 | 4.1796 | 14.4716 | 5.1355 | 1.4714 | -0.4821 | 0.6851 |
| HMR_3142 | Beta oxidation of even-chain fatty acids (mitochondrial) | 12.9927 | 4.1886 | 14.4645 | 5.1445 | 1.4717 | -0.4817 | 0.685 |
| HMR_3143 | Beta oxidation of even-chain fatty acids (mitochondrial) | 14.8936 | 4.6632 | 16.6481 | 5.7363 | 1.7545 | -0.5441 | 0.7068 |
| HMR_3144 | Beta oxidation of even-chain fatty acids (mitochondrial) | 14.8868 | 4.6716 | 16.6415 | 5.7446 | 1.7547 | -0.5437 | 0.7067 |
| HMR_3146 | Beta oxidation of even-chain fatty acids (mitochondrial) | 14.8801 | 4.6799 | 16.635 | 5.753 | 1.7549 | -0.5433 | 0.7065 |
| HMR_3149 | Beta oxidation of even-chain fatty acids (mitochondrial) | 17.6887 | 6.4381 | 19.6284 | 7.659 | 1.9397 | -0.5166 | 0.6973 |
| HMR_3150 | Beta oxidation of even-chain fatty acids (mitochondrial) | 17.6828 | 6.4455 | 19.6226 | 7.6662 | 1.9398 | -0.5164 | 0.6972 |
| HMR_3151 | Beta oxidation of even-chain fatty acids (mitochondrial) | 17.6769 | 6.4528 | 19.6167 | 7.6734 | 1.9398 | -0.5161 | 0.6971 |
| HMR_3153 | Beta oxidation of even-chain fatty acids (mitochondrial) | 17.671 | 6.4601 | 19.6108 | 7.6806 | 1.9398 | -0.5158 | 0.697 |
| HMR_3156 | Beta oxidation of even-chain fatty acids (mitochondrial) | 17.6651 | 6.4674 | 19.6049 | 7.6878 | 1.9398 | -0.5156 | 0.6969 |
| HMR_3157 | Beta oxidation of even-chain fatty acids (mitochondrial) | 17.6601 | 6.4747 | 19.5997 | 7.695 | 1.9396 | -0.5153 | 0.6968 |
| HMR_3158 | Beta oxidation of even-chain fatty acids (mitochondrial) | 17.6549 | 6.4819 | 19.5944 | 7.7021 | 1.9394 | -0.515 | 0.6967 |
| HMR_3160 | Beta oxidation of even-chain fatty acids (mitochondrial) | 17.6498 | 6.4891 | 19.5891 | 7.7092 | 1.9392 | -0.5146 | 0.6966 |
| HMR_3398 | Beta oxidation of even-chain fatty acids (mitochondrial) | 0.8998 | 0.2393 | 1.0078 | 0.2977 | 0.108 | -0.1474 | 0.5586 |
| HMR_3406 | Beta oxidation of even-chain fatty acids (mitochondrial) | 0.9942 | 0.2562 | 1.1153 | 0.3163 | 0.1211 | -0.1601 | 0.5636 |
| HMR_3407 | Beta oxidation of even-chain fatty acids (mitochondrial) | 0.697 | 0.146 | 0.7835 | 0.1817 | 0.0865 | -0.1511 | 0.5601 |
| HMR_3408 | Beta oxidation of even-chain fatty acids (mitochondrial) | 0.6875 | 0.1519 | 0.7756 | 0.1911 | 0.088 | -0.1503 | 0.5597 |
| HMR_3409 | Beta oxidation of even-chain fatty acids (mitochondrial) | 1.2132 | 0.4548 | 1.4335 | 0.6286 | 0.2203 | -0.2117 | 0.5838 |
| HMR_3411 | Beta oxidation of even-chain fatty acids (mitochondrial) | 0.5904 | 0.1118 | 0.6595 | 0.1377 | 0.0692 | -0.1385 | 0.5551 |
| HMR_3413 | Beta oxidation of even-chain fatty acids (mitochondrial) | 0.9216 | 0.2619 | 1.0652 | 0.3458 | 0.1436 | -0.1842 | 0.5731 |
| HMR_3414 | Beta oxidation of even-chain fatty acids (mitochondrial) | 0.9076 | 0.2346 | 1.0353 | 0.3009 | 0.1277 | -0.1746 | 0.5693 |
| HMR_3416 | Beta oxidation of even-chain fatty acids (mitochondrial) | 0.5641 | 0.0904 | 0.6201 | 0.1063 | 0.056 | -0.1263 | 0.5502 |
| HMR_3421 | Beta oxidation of even-chain fatty acids (mitochondrial) | 0.6591 | 0.1405 | 0.7341 | 0.1722 | 0.075 | -0.1341 | 0.5533 |
| HMR_3422 | Beta oxidation of even-chain fatty acids (mitochondrial) | 1.2932 | 0.3494 | 1.4415 | 0.4225 | 0.1482 | -0.1687 | 0.567 |
| HMR_3423 | Beta oxidation of even-chain fatty acids (mitochondrial) | 0.8274 | 0.2134 | 0.9449 | 0.2753 | 0.1175 | -0.168 | 0.5667 |
| HMR_3424 | Beta oxidation of even-chain fatty acids (mitochondrial) | 0.7211 | 0.1402 | 0.8083 | 0.1714 | 0.0872 | -0.1562 | 0.5621 |
| HMR_3425 | Beta oxidation of even-chain fatty acids (mitochondrial) | 0.7017 | 0.1336 | 0.7796 | 0.161 | 0.078 | -0.1437 | 0.5571 |
| HMR_3426 | Beta oxidation of even-chain fatty acids (mitochondrial) | 1.1446 | 0.4303 | 1.3266 | 0.5717 | 0.182 | -0.1818 | 0.5721 |

| | | | | | | | | |
|---|---|---|---|---|---|---|---|---|
| HMR_9719 | Beta oxidation of even-chain fatty acids (mitochondrial) | 1.1139 | 0.3873 | 1.2719 | 0.502 | 0.158 | -0.1676 | 0.5665 |
| HMR_3427 | Beta oxidation of even-chain fatty acids (mitochondrial) | 0.8614 | 0.2347 | 0.9713 | 0.296 | 0.1099 | -0.1508 | 0.5599 |
| HMR_3428 | Beta oxidation of even-chain fatty acids (mitochondrial) | 0.8339 | 0.2516 | 0.9319 | 0.3132 | 0.098 | -0.1304 | 0.5519 |
| HMR_3429 | Beta oxidation of even-chain fatty acids (mitochondrial) | 0.6869 | 0.1526 | 0.7623 | 0.1861 | 0.0753 | -0.1295 | 0.5515 |
| HMR_3430 | Beta oxidation of even-chain fatty acids (mitochondrial) | 1.1157 | 0.3097 | 1.2436 | 0.3716 | 0.1279 | -0.1549 | 0.5616 |
| HMR_3431 | Beta oxidation of even-chain fatty acids (mitochondrial) | 0.888 | 0.2415 | 1.0102 | 0.3094 | 0.1222 | -0.1646 | 0.5654 |
| HMR_3432 | Beta oxidation of even-chain fatty acids (mitochondrial) | 0.7632 | 0.1546 | 0.8537 | 0.1886 | 0.0905 | -0.1545 | 0.5614 |
| HMR_3433 | Beta oxidation of even-chain fatty acids (mitochondrial) | 0.7196 | 0.1627 | 0.8031 | 0.2004 | 0.0835 | -0.1386 | 0.5551 |
| HMR_3170 | Beta oxidation of odd-chain fatty acids (mitochondrial) | 3.6104 | 2.7008 | 3.9574 | 3.2329 | 0.347 | -0.1424 | 0.5566 |
| HMR_3171 | Beta oxidation of odd-chain fatty acids (mitochondrial) | 3.6235 | 2.704 | 3.9703 | 3.2361 | 0.3468 | -0.1423 | 0.5566 |
| HMR_3172 | Beta oxidation of odd-chain fatty acids (mitochondrial) | 3.6363 | 2.7091 | 3.9829 | 3.2414 | 0.3466 | -0.1421 | 0.5565 |
| HMR_3173 | Beta oxidation of odd-chain fatty acids (mitochondrial) | 3.6489 | 2.7163 | 3.9952 | 3.2486 | 0.3464 | -0.1418 | 0.5564 |
| HMR_3174 | Beta oxidation of odd-chain fatty acids (mitochondrial) | 3.565 | 3.0743 | 3.9016 | 3.6609 | 0.3366 | -0.1297 | 0.5516 |
| HMR_3175 | Beta oxidation of odd-chain fatty acids (mitochondrial) | 3.5777 | 3.074 | 3.914 | 3.6607 | 0.3363 | -0.1296 | 0.5516 |
| HMR_3176 | Beta oxidation of odd-chain fatty acids (mitochondrial) | 3.5898 | 3.0766 | 3.9258 | 3.6634 | 0.336 | -0.1294 | 0.5515 |
| HMR_3177 | Beta oxidation of odd-chain fatty acids (mitochondrial) | 3.6015 | 3.0822 | 3.9373 | 3.6691 | 0.3358 | -0.1292 | 0.5514 |
| HMR_3178 | Beta oxidation of odd-chain fatty acids (mitochondrial) | 0.5989 | 0.1057 | 0.629 | 0.1158 | 0.0301 | -0.064 | 0.5255 |
| HMR_3179 | Beta oxidation of odd-chain fatty acids (mitochondrial) | 0.607 | 0.1014 | 0.637 | 0.1115 | 0.03 | -0.0651 | 0.5259 |
| HMR_3180 | Beta oxidation of odd-chain fatty acids (mitochondrial) | 0.6117 | 0.1008 | 0.6417 | 0.1109 | 0.0301 | -0.0653 | 0.526 |
| HMR_3181 | Beta oxidation of odd-chain fatty acids (mitochondrial) | 0.6131 | 0.1036 | 0.6433 | 0.1138 | 0.0302 | -0.0648 | 0.5258 |
| HMR_3182 | Beta oxidation of odd-chain fatty acids (mitochondrial) | 1.0051 | 0.1844 | 1.0676 | 0.2057 | 0.0626 | -0.1002 | 0.5399 |
| HMR_3183 | Beta oxidation of odd-chain fatty acids (mitochondrial) | 1.0031 | 0.185 | 1.066 | 0.2065 | 0.0628 | -0.1004 | 0.54 |
| HMR_3184 | Beta oxidation of odd-chain fatty acids (mitochondrial) | 1.0003 | 0.1869 | 1.0634 | 0.2086 | 0.0631 | -0.1004 | 0.54 |
| HMR_3185 | Beta oxidation of odd-chain fatty acids (mitochondrial) | 0.9967 | 0.1902 | 1.0602 | 0.2121 | 0.0635 | -0.1001 | 0.5399 |
| HMR_3186 | Beta oxidation of odd-chain fatty acids (mitochondrial) | 1.4596 | 0.2458 | 1.558 | 0.2767 | 0.0984 | -0.1361 | 0.5541 |
| HMR_3187 | Beta oxidation of odd-chain fatty acids (mitochondrial) | 1.4558 | 0.2469 | 1.5545 | 0.2781 | 0.0987 | -0.1362 | 0.5542 |
| HMR_3188 | Beta oxidation of odd-chain fatty acids (mitochondrial) | 1.4519 | 0.2481 | 1.5509 | 0.2796 | 0.099 | -0.1363 | 0.5542 |
| HMR_3189 | Beta oxidation of odd-chain fatty acids (mitochondrial) | 1.4478 | 0.2494 | 1.5471 | 0.2813 | 0.0993 | -0.1363 | 0.5542 |
| HMR_3190 | Beta oxidation of odd-chain fatty acids (mitochondrial) | 1.4436 | 0.2509 | 1.5432 | 0.2831 | 0.0996 | -0.1363 | 0.5542 |
| HMR_3191 | Beta oxidation of odd-chain fatty acids (mitochondrial) | 1.4401 | 0.2525 | 1.5398 | 0.2851 | 0.0997 | -0.136 | 0.5541 |
| HMR_3192 | Beta oxidation of odd-chain fatty acids (mitochondrial) | 1.4363 | 0.2544 | 1.5362 | 0.2874 | 0.0999 | -0.1357 | 0.554 |
| HMR_3193 | Beta oxidation of odd-chain fatty acids (mitochondrial) | 1.4325 | 0.2565 | 1.5325 | 0.2898 | 0.1 | -0.1354 | 0.5538 |
| HMR_3194 | Beta oxidation of odd-chain fatty acids (mitochondrial) | 1.4285 | 0.2588 | 1.5287 | 0.2925 | 0.1002 | -0.135 | 0.5537 |
| HMR_3195 | Beta oxidation of odd-chain fatty acids (mitochondrial) | 1.4252 | 0.2615 | 1.5254 | 0.2955 | 0.1002 | -0.1343 | 0.5534 |
| HMR_3196 | Beta oxidation of odd-chain fatty acids (mitochondrial) | 1.4216 | 0.2644 | 1.5219 | 0.2987 | 0.1003 | -0.1336 | 0.5532 |
| HMR_3197 | Beta oxidation of odd-chain fatty acids (mitochondrial) | 1.4178 | 0.2677 | 1.5181 | 0.3024 | 0.1003 | -0.1329 | 0.5528 |
| HMR_3198 | Beta oxidation of odd-chain fatty acids (mitochondrial) | 1.4139 | 0.2714 | 1.5143 | 0.3064 | 0.1004 | -0.132 | 0.5525 |
| HMR_3199 | Beta oxidation of odd-chain fatty acids (mitochondrial) | 1.4106 | 0.2755 | 1.5109 | 0.3108 | 0.1003 | -0.131 | 0.5521 |
| HMR_3200 | Beta oxidation of odd-chain fatty acids (mitochondrial) | 1.4069 | 0.2801 | 1.5072 | 0.3157 | 0.1002 | -0.1299 | 0.5517 |
| HMR_3201 | Beta oxidation of odd-chain fatty acids (mitochondrial) | 1.4031 | 0.2853 | 1.5032 | 0.3212 | 0.1002 | -0.1286 | 0.5512 |
| HMR_3202 | Beta oxidation of odd-chain fatty acids (mitochondrial) | 1.3989 | 0.2911 | 1.4991 | 0.3272 | 0.1001 | -0.1274 | 0.5507 |
| HMR_3203 | Beta oxidation of odd-chain fatty acids (mitochondrial) | 1.3953 | 0.2976 | 1.4953 | 0.334 | 0.1 | -0.1259 | 0.5501 |
| HMR_3204 | Beta oxidation of odd-chain fatty acids (mitochondrial) | 1.3913 | 0.3049 | 1.4912 | 0.3415 | 0.0999 | -0.1243 | 0.5494 |
| HMR_3205 | Beta oxidation of odd-chain fatty acids (mitochondrial) | 1.387 | 0.3132 | 1.4868 | 0.3498 | 0.0998 | -0.1226 | 0.5488 |

| | | | | | | | | |
|---|---|---|---|---|---|---|---|---|
| HMR_3396 | Beta oxidation of odd-chain fatty acids (mitochondrial) | 0.7133 | 0.1242 | 0.7829 | 0.1432 | 0.0696 | -0.1346 | 0.5535 |
| HMR_3397 | Beta oxidation of odd-chain fatty acids (mitochondrial) | 0.473 | 0.0722 | 0.5022 | 0.0804 | 0.0292 | -0.0747 | 0.5298 |
| HMR_3240 | Beta oxidation of unsaturated fatty acids (n-7) (mitochondrial) | 3.5951 | 2.7605 | 3.9374 | 3.3015 | 0.3422 | -0.139 | 0.5553 |
| HMR_3241 | Beta oxidation of unsaturated fatty acids (n-7) (mitochondrial) | 3.6011 | 2.7634 | 3.9425 | 3.3045 | 0.3414 | -0.1386 | 0.5551 |
| HMR_3242 | Beta oxidation of unsaturated fatty acids (n-7) (mitochondrial) | 3.6067 | 2.7686 | 3.9473 | 3.3097 | 0.3406 | -0.1382 | 0.5549 |
| HMR_3243 | Beta oxidation of unsaturated fatty acids (n-7) (mitochondrial) | 3.612 | 2.7761 | 3.9518 | 3.3172 | 0.3398 | -0.1377 | 0.5547 |
| HMR_3244 | Beta oxidation of unsaturated fatty acids (n-7) (mitochondrial) | 0.699 | 0.1525 | 0.7871 | 0.1916 | 0.0881 | -0.1502 | 0.5597 |
| HMR_3245 | Beta oxidation of unsaturated fatty acids (n-7) (mitochondrial) | 0.701 | 0.1478 | 0.7886 | 0.1869 | 0.0877 | -0.1515 | 0.5602 |
| HMR_3246 | Beta oxidation of unsaturated fatty acids (n-7) (mitochondrial) | 0.6993 | 0.1478 | 0.7869 | 0.1869 | 0.0876 | -0.1514 | 0.5602 |
| HMR_3247 | Beta oxidation of unsaturated fatty acids (n-7) (mitochondrial) | 0.6942 | 0.1524 | 0.7821 | 0.1916 | 0.0879 | -0.1499 | 0.5596 |
| HMR_3250 | Beta oxidation of unsaturated fatty acids (n-7) (mitochondrial) | 6.0875 | 2.1189 | 6.6544 | 2.589 | 0.5669 | -0.2613 | 0.6031 |
| HMR_3252 | Beta oxidation of unsaturated fatty acids (n-7) (mitochondrial) | 6.0793 | 2.1292 | 6.6468 | 2.5992 | 0.5675 | -0.261 | 0.603 |
| HMR_3254 | Beta oxidation of unsaturated fatty acids (n-7) (mitochondrial) | 6.071 | 2.1394 | 6.6392 | 2.6095 | 0.5682 | -0.2607 | 0.6028 |
| HMR_3256 | Beta oxidation of unsaturated fatty acids (n-7) (mitochondrial) | 6.0627 | 2.1496 | 6.6315 | 2.6197 | 0.5688 | -0.2605 | 0.6027 |
| HMR_3258 | Beta oxidation of unsaturated fatty acids (n-7) (mitochondrial) | 6.672 | 2.5177 | 7.3393 | 3.0714 | 0.6673 | -0.2823 | 0.6111 |
| HMR_3260 | Beta oxidation of unsaturated fatty acids (n-7) (mitochondrial) | 6.6646 | 2.5278 | 7.3324 | 3.0815 | 0.6678 | -0.282 | 0.611 |
| HMR_3262 | Beta oxidation of unsaturated fatty acids (n-7) (mitochondrial) | 6.6571 | 2.5379 | 7.3253 | 3.0916 | 0.6683 | -0.2817 | 0.6109 |
| HMR_3264 | Beta oxidation of unsaturated fatty acids (n-7) (mitochondrial) | 6.6496 | 2.5479 | 7.3183 | 3.1016 | 0.6687 | -0.2813 | 0.6108 |
| HMR_3272 | Beta oxidation of unsaturated fatty acids (n-7) (mitochondrial) | -6.6422 | 2.5579 | -7.3113 | 3.1116 | 0.6691 | 0.281 | 0.3894 |
| HMR_3356 | Beta oxidation of unsaturated fatty acids (n-7) (mitochondrial) | 0.6904 | 0.0986 | 0.7121 | 0.1039 | 0.0218 | -0.0484 | 0.5193 |
| HMR_3357 | Beta oxidation of unsaturated fatty acids (n-7) (mitochondrial) | 0.6973 | 0.0936 | 0.7187 | 0.0986 | 0.0213 | -0.0486 | 0.5194 |
| HMR_3358 | Beta oxidation of unsaturated fatty acids (n-7) (mitochondrial) | 0.7033 | 0.0898 | 0.7242 | 0.0948 | 0.021 | -0.0488 | 0.5194 |
| HMR_3359 | Beta oxidation of unsaturated fatty acids (n-7) (mitochondrial) | 0.7081 | 0.0872 | 0.7287 | 0.0919 | 0.0206 | -0.0487 | 0.5194 |
| HMR_3360 | Beta oxidation of unsaturated fatty acids (n-7) (mitochondrial) | 0.712 | 0.0853 | 0.7323 | 0.09 | 0.0203 | -0.0485 | 0.5193 |
| HMR_3361 | Beta oxidation of unsaturated fatty acids (n-7) (mitochondrial) | 0.7156 | 0.0841 | 0.7356 | 0.0886 | 0.02 | -0.048 | 0.5191 |
| HMR_3362 | Beta oxidation of unsaturated fatty acids (n-7) (mitochondrial) | 0.7186 | 0.0834 | 0.7383 | 0.0879 | 0.0197 | -0.0475 | 0.519 |
| HMR_3363 | Beta oxidation of unsaturated fatty acids (n-7) (mitochondrial) | 0.7209 | 0.0833 | 0.7403 | 0.0877 | 0.0194 | -0.0469 | 0.5187 |
| HMR_3218 | Beta oxidation of unsaturated fatty acids (n-9) (mitochondrial) | 3.6138 | 2.9404 | 3.9519 | 3.5078 | 0.3381 | -0.1331 | 0.553 |
| HMR_3219 | Beta oxidation of unsaturated fatty acids (n-9) (mitochondrial) | 3.6145 | 2.9422 | 3.9522 | 3.5097 | 0.3376 | -0.1329 | 0.5529 |
| HMR_3220 | Beta oxidation of unsaturated fatty acids (n-9) (mitochondrial) | 3.6148 | 2.9467 | 3.9521 | 3.5142 | 0.3372 | -0.1327 | 0.5528 |
| HMR_3221 | Beta oxidation of unsaturated fatty acids (n-9) (mitochondrial) | 3.6148 | 2.9538 | 3.9516 | 3.5213 | 0.3368 | -0.1324 | 0.5526 |
| HMR_3222 | Beta oxidation of unsaturated fatty acids (n-9) (mitochondrial) | 3.5513 | 3.6279 | 3.876 | 4.3053 | 0.3247 | -0.1153 | 0.5459 |
| HMR_3223 | Beta oxidation of unsaturated fatty acids (n-9) (mitochondrial) | 3.5511 | 3.6241 | 3.8753 | 4.3016 | 0.3242 | -0.1152 | 0.5458 |
| HMR_3224 | Beta oxidation of unsaturated fatty acids (n-9) (mitochondrial) | 3.5502 | 3.6248 | 3.8739 | 4.3024 | 0.3238 | -0.115 | 0.5458 |
| HMR_3225 | Beta oxidation of unsaturated fatty acids (n-9) (mitochondrial) | 3.5486 | 3.6299 | 3.872 | 4.3076 | 0.3234 | -0.1148 | 0.5457 |
| HMR_3226 | Beta oxidation of unsaturated fatty acids (n-9) (mitochondrial) | 1.8757 | 0.7572 | 2.114 | 0.9606 | 0.2383 | -0.1818 | 0.5721 |
| HMR_3227 | Beta oxidation of unsaturated fatty acids (n-9) (mitochondrial) | 1.8738 | 0.7511 | 2.1116 | 0.9546 | 0.2379 | -0.1821 | 0.5723 |
| HMR_3228 | Beta oxidation of unsaturated fatty acids (n-9) (mitochondrial) | 1.8711 | 0.7474 | 2.1086 | 0.951 | 0.2375 | -0.1823 | 0.5723 |
| HMR_3229 | Beta oxidation of unsaturated fatty acids (n-9) (mitochondrial) | 1.8678 | 0.7459 | 2.105 | 0.9496 | 0.2372 | -0.1821 | 0.5723 |
| HMR_3230 | Beta oxidation of unsaturated fatty acids (n-9) (mitochondrial) | 1.563 | 0.5805 | 1.869 | 0.8133 | 0.306 | -0.2592 | 0.6023 |
| HMR_3231 | Beta oxidation of unsaturated fatty acids (n-9) (mitochondrial) | 1.5589 | 0.578 | 1.8647 | 0.8112 | 0.3058 | -0.2594 | 0.6024 |
| HMR_3232 | Beta oxidation of unsaturated fatty acids (n-9) (mitochondrial) | 1.5538 | 0.5783 | 1.8596 | 0.8119 | 0.3057 | -0.2593 | 0.6023 |
| HMR_3233 | Beta oxidation of unsaturated fatty acids (n-9) (mitochondrial) | 1.5477 | 0.5816 | 1.8536 | 0.8155 | 0.3059 | -0.2588 | 0.6021 |
| HMR_3234 | Beta oxidation of unsaturated fatty acids (n-9) (mitochondrial) | 2.2465 | 0.7751 | 2.6563 | 1.0584 | 0.4098 | -0.3026 | 0.6189 |

| | | | | | | | | |
|---|---|---|---|---|---|---|---|---|
| HMR_3235 | Beta oxidation of unsaturated fatty acids (n-9) (mitochondrial) | 2.2401 | 0.7808 | 2.6501 | 1.0644 | 0.41 | -0.3019 | 0.6186 |
| HMR_3236 | Beta oxidation of unsaturated fatty acids (n-9) (mitochondrial) | 2.2333 | 0.7874 | 2.6436 | 1.0713 | 0.4103 | -0.301 | 0.6183 |
| HMR_3237 | Beta oxidation of unsaturated fatty acids (n-9) (mitochondrial) | 2.2263 | 0.7949 | 2.6369 | 1.079 | 0.4106 | -0.3 | 0.6179 |
| HMR_3239 | Beta oxidation of unsaturated fatty acids (n-9) (mitochondrial) | -2.2191 | 0.8035 | -2.6301 | 1.0877 | 0.411 | 0.2988 | 0.3825 |
| HMR_3275 | Beta oxidation of di-unsaturated fatty acids (n-6) (mitochondrial) | 1.0541 | 0.2437 | 1.2082 | 0.3204 | 0.1541 | -0.2052 | 0.5813 |
| HMR_3277 | Beta oxidation of di-unsaturated fatty acids (n-6) (mitochondrial) | 1.0515 | 0.2414 | 1.2056 | 0.3181 | 0.154 | -0.2059 | 0.5816 |
| HMR_3278 | Beta oxidation of di-unsaturated fatty acids (n-6) (mitochondrial) | 1.0478 | 0.2414 | 1.2018 | 0.3181 | 0.154 | -0.2059 | 0.5816 |
| HMR_3279 | Beta oxidation of di-unsaturated fatty acids (n-6) (mitochondrial) | 1.0428 | 0.2436 | 1.1969 | 0.3205 | 0.1541 | -0.2052 | 0.5813 |
| HMR_3280 | Beta oxidation of di-unsaturated fatty acids (n-6) (mitochondrial) | 1.0365 | 0.2484 | 1.191 | 0.3254 | 0.1544 | -0.2039 | 0.5808 |
| HMR_3281 | Beta oxidation of di-unsaturated fatty acids (n-6) (mitochondrial) | 1.9939 | 0.4104 | 2.2737 | 0.5342 | 0.2798 | -0.2879 | 0.6133 |
| HMR_3282 | Beta oxidation of di-unsaturated fatty acids (n-6) (mitochondrial) | 1.987 | 0.4157 | 2.2671 | 0.5399 | 0.2801 | -0.2866 | 0.6128 |
| HMR_3283 | Beta oxidation of di-unsaturated fatty acids (n-6) (mitochondrial) | 1.98 | 0.4211 | 2.2605 | 0.5456 | 0.2804 | -0.2852 | 0.6123 |
| HMR_3284 | Beta oxidation of di-unsaturated fatty acids (n-6) (mitochondrial) | 1.973 | 0.4267 | 2.2538 | 0.5516 | 0.2808 | -0.2839 | 0.6117 |
| HMR_3285 | Beta oxidation of di-unsaturated fatty acids (n-6) (mitochondrial) | 1.9668 | 0.4324 | 2.2477 | 0.5576 | 0.2809 | -0.2823 | 0.6112 |
| HMR_3286 | Beta oxidation of di-unsaturated fatty acids (n-6) (mitochondrial) | 1.9604 | 0.4383 | 2.2415 | 0.5639 | 0.2811 | -0.2808 | 0.6106 |
| HMR_3287 | Beta oxidation of di-unsaturated fatty acids (n-6) (mitochondrial) | 1.954 | 0.4445 | 2.2352 | 0.5704 | 0.2813 | -0.2792 | 0.6099 |
| HMR_3288 | Beta oxidation of di-unsaturated fatty acids (n-6) (mitochondrial) | 1.9475 | 0.4508 | 2.2289 | 0.5772 | 0.2814 | -0.2776 | 0.6093 |
| HMR_3290 | Beta oxidation of di-unsaturated fatty acids (n-6) (mitochondrial) | 1.9409 | 0.4575 | 2.2225 | 0.5842 | 0.2816 | -0.2759 | 0.6087 |
| HMR_3292 | Beta oxidation of di-unsaturated fatty acids (n-6) (mitochondrial) | 1.9341 | 0.4645 | 2.2159 | 0.5914 | 0.2818 | -0.2743 | 0.6081 |
| HMR_3293 | Beta oxidation of di-unsaturated fatty acids (n-6) (mitochondrial) | 1.9272 | 0.4718 | 2.2092 | 0.5991 | 0.282 | -0.2725 | 0.6074 |
| HMR_3294 | Beta oxidation of di-unsaturated fatty acids (n-6) (mitochondrial) | 1.9202 | 0.4796 | 2.2025 | 0.6071 | 0.2822 | -0.2707 | 0.6067 |
| HMR_3296 | Beta oxidation of di-unsaturated fatty acids (n-6) (mitochondrial) | 1.9139 | 0.4878 | 2.1963 | 0.6155 | 0.2823 | -0.2688 | 0.606 |
| HMR_3298 | Beta oxidation of di-unsaturated fatty acids (n-6) (mitochondrial) | -1.9075 | 0.4966 | -2.19 | 0.6243 | 0.2825 | 0.2669 | 0.3948 |
| HMR_1174 | Beta oxidation of poly-unsaturated fatty acids (mitochondrial) | 1.4702 | 0.221 | 1.583 | 0.2553 | 0.1128 | -0.1634 | 0.5649 |
| HMR_1175 | Beta oxidation of poly-unsaturated fatty acids (mitochondrial) | 1.4676 | 0.227 | 1.581 | 0.2616 | 0.1134 | -0.1622 | 0.5644 |
| HMR_1176 | Beta oxidation of poly-unsaturated fatty acids (mitochondrial) | 1.4654 | 0.2331 | 1.5794 | 0.268 | 0.114 | -0.1611 | 0.564 |
| HMR_1177 | Beta oxidation of poly-unsaturated fatty acids (mitochondrial) | 1.4632 | 0.2393 | 1.5778 | 0.2746 | 0.1147 | -0.1599 | 0.5635 |
| HMR_1178 | Beta oxidation of poly-unsaturated fatty acids (mitochondrial) | 1.4607 | 0.2457 | 1.576 | 0.2813 | 0.1153 | -0.1588 | 0.5631 |
| HMR_1179 | Beta oxidation of poly-unsaturated fatty acids (mitochondrial) | 1.4582 | 0.2523 | 1.5741 | 0.2882 | 0.1159 | -0.1577 | 0.5626 |
| HMR_1180 | Beta oxidation of poly-unsaturated fatty acids (mitochondrial) | 1.4555 | 0.2591 | 1.5721 | 0.2954 | 0.1166 | -0.1565 | 0.5622 |
| HMR_1181 | Beta oxidation of poly-unsaturated fatty acids (mitochondrial) | 1.4533 | 0.2663 | 1.5705 | 0.3029 | 0.1172 | -0.1553 | 0.5617 |
| HMR_1182 | Beta oxidation of poly-unsaturated fatty acids (mitochondrial) | 1.4509 | 0.2739 | 1.5688 | 0.3108 | 0.1178 | -0.1541 | 0.5612 |
| HMR_1183 | Beta oxidation of poly-unsaturated fatty acids (mitochondrial) | 1.4482 | 0.282 | 1.5668 | 0.3191 | 0.1185 | -0.1529 | 0.5608 |
| HMR_1184 | Beta oxidation of poly-unsaturated fatty acids (mitochondrial) | 1.4454 | 0.2906 | 1.5646 | 0.3278 | 0.1192 | -0.1516 | 0.5603 |
| HMR_1216 | Beta oxidation of poly-unsaturated fatty acids (mitochondrial) | 1.4702 | 0.221 | 1.583 | 0.2553 | 0.1128 | -0.1634 | 0.5649 |
| HMR_1217 | Beta oxidation of poly-unsaturated fatty acids (mitochondrial) | 1.4676 | 0.227 | 1.581 | 0.2616 | 0.1134 | -0.1622 | 0.5644 |
| HMR_1218 | Beta oxidation of poly-unsaturated fatty acids (mitochondrial) | 1.4654 | 0.2331 | 1.5794 | 0.268 | 0.114 | -0.1611 | 0.564 |
| HMR_1219 | Beta oxidation of poly-unsaturated fatty acids (mitochondrial) | -1.4632 | 0.2393 | -1.5778 | 0.2746 | 0.1147 | 0.1599 | 0.4365 |
| HMR_1220 | Beta oxidation of poly-unsaturated fatty acids (mitochondrial) | 1.4607 | 0.2457 | 1.576 | 0.2813 | 0.1153 | -0.1588 | 0.5631 |
| HMR_1221 | Beta oxidation of poly-unsaturated fatty acids (mitochondrial) | 1.4582 | 0.2523 | 1.5741 | 0.2882 | 0.1159 | -0.1577 | 0.5626 |
| HMR_1222 | Beta oxidation of poly-unsaturated fatty acids (mitochondrial) | 1.4533 | 0.2663 | 1.5705 | 0.3029 | 0.1172 | -0.1553 | 0.5617 |
| HMR_1223 | Beta oxidation of poly-unsaturated fatty acids (mitochondrial) | 1.4555 | 0.2591 | 1.5721 | 0.2954 | 0.1166 | -0.1565 | 0.5622 |
| HMR_1224 | Beta oxidation of poly-unsaturated fatty acids (mitochondrial) | 1.4509 | 0.2739 | 1.5688 | 0.3108 | 0.1178 | -0.1541 | 0.5612 |
| HMR_1225 | Beta oxidation of poly-unsaturated fatty acids (mitochondrial) | 1.4482 | 0.282 | 1.5668 | 0.3191 | 0.1185 | -0.1529 | 0.5608 |

| | | | | | | | | |
|---|---|---|---|---|---|---|---|---|
| HMR_1226 | Beta oxidation of poly-unsaturated fatty acids (mitochondrial) | 1.4454 | 0.2906 | 1.5646 | 0.3278 | 0.1192 | -0.1516 | 0.5603 |
| HMR_3522 | Beta oxidation of branched-chain fatty acids (mitochondrial) | 0.8748 | 0.1044 | 1.158 | 0.1732 | 0.2831 | -0.5374 | 0.7045 |
| HMR_3523 | Beta oxidation of branched-chain fatty acids (mitochondrial) | 0.8756 | 0.1056 | 1.1617 | 0.1757 | 0.2861 | -0.5394 | 0.7052 |
| HMR_3524 | Beta oxidation of branched-chain fatty acids (mitochondrial) | 0.876 | 0.107 | 1.1651 | 0.1784 | 0.2891 | -0.5412 | 0.7058 |
| HMR_3525 | Beta oxidation of branched-chain fatty acids (mitochondrial) | 0.876 | 0.1086 | 1.1683 | 0.1815 | 0.2923 | -0.5428 | 0.7064 |
| HMR_3526 | Beta oxidation of branched-chain fatty acids (mitochondrial) | 0.8756 | 0.1104 | 1.1713 | 0.1848 | 0.2957 | -0.5442 | 0.7068 |
| HMR_3527 | Beta oxidation of branched-chain fatty acids (mitochondrial) | 0.8748 | 0.1126 | 1.174 | 0.1885 | 0.2992 | -0.5453 | 0.7072 |
| HMR_3528 | Beta oxidation of branched-chain fatty acids (mitochondrial) | 0.8743 | 0.1151 | 1.1771 | 0.1925 | 0.3027 | -0.5458 | 0.7074 |
| HMR_3529 | Beta oxidation of branched-chain fatty acids (mitochondrial) | 0.8733 | 0.1181 | 1.1798 | 0.197 | 0.3065 | -0.546 | 0.7075 |
| HMR_3530 | Beta oxidation of branched-chain fatty acids (mitochondrial) | 0.8717 | 0.1217 | 1.1822 | 0.2021 | 0.3105 | -0.5457 | 0.7074 |
| HMR_3531 | Beta oxidation of branched-chain fatty acids (mitochondrial) | 0.87 | 0.1261 | 1.1844 | 0.2077 | 0.3144 | -0.5442 | 0.7069 |
| HMR_3532 | Beta oxidation of branched-chain fatty acids (mitochondrial) | 0.8684 | 0.1313 | 1.1869 | 0.214 | 0.3185 | -0.5421 | 0.7061 |
| HMR_3533 | Beta oxidation of branched-chain fatty acids (mitochondrial) | 0.866 | 0.1376 | 1.1889 | 0.221 | 0.323 | -0.5393 | 0.7052 |
| HMR_3534 | Beta oxidation of branched-chain fatty acids (mitochondrial) | 0.8626 | 0.1454 | 1.1906 | 0.2289 | 0.3279 | -0.536 | 0.704 |
| HMR_1573 | Terpenoid backbone biosynthesis | 14.5075 | 159.34 | 14.5105 | 158.0427 | 0.003 | -0.0002 | 0.5001 |
| HMR_1920 | Steroid metabolism | 1.1791 | 0.6563 | 1.1643 | 0.6399 | 0.0148 | 0.013 | 0.4948 |
| HMR_1924 | Steroid metabolism | 472.0928 | 48381.2228 | 472.0879 | 48381.0757 | 0.0049 | 0.00E+00 | 0.5 |
| HMR_1926 | Steroid metabolism | 472.1179 | 48381.2207 | 472.1151 | 48381.0751 | 0.0028 | 0.00E+00 | 0.5 |
| HMR_7648 | Steroid metabolism | 7.752 | 3.6163 | 7.7564 | 3.6018 | 0.0044 | -0.0016 | 0.5007 |
| HMR_7968 | Steroid metabolism | 2.0603 | 2.7708 | 2.0534 | 2.753 | 0.0069 | 0.0029 | 0.4988 |
| HMR_7984 | Steroid metabolism | 0.8951 | 0.1866 | 0.8829 | 0.1817 | 0.0121 | 0.02 | 0.492 |
| HMR_1952 | Androgen metabolism | 1.1802 | 0.6529 | 1.1652 | 0.6365 | 0.0149 | 0.0132 | 0.4947 |
| HMR_1953 | Androgen metabolism | 1.1822 | 0.6529 | 1.1672 | 0.6365 | 0.0149 | 0.0132 | 0.4948 |
| HMR_1962 | Androgen metabolism | 1.1802 | 0.6529 | 1.1652 | 0.6365 | 0.0149 | 0.0132 | 0.4947 |
| HMR_1963 | Androgen metabolism | 1.1822 | 0.6529 | 1.1672 | 0.6365 | 0.0149 | 0.0132 | 0.4948 |
| HMR_1967 | Androgen metabolism | 4.5535 | 2.1855 | 4.6592 | 2.2788 | 0.1057 | -0.05 | 0.52 |
| HMR_1969 | Androgen metabolism | 1.1743 | 1.5138 | 1.16 | 1.4623 | 0.0143 | 0.0083 | 0.4967 |
| HMR_1970 | Androgen metabolism | 4.5488 | 2.183 | 4.6547 | 2.2761 | 0.1059 | -0.0501 | 0.52 |
| HMR_1973 | Androgen metabolism | 1.6943 | 0.803 | 1.7193 | 0.8275 | 0.0249 | -0.0195 | 0.5078 |
| HMR_2016 | Androgen metabolism | 385.6753 | 158501.3963 | 382.12 | 160072.8037 | 3.5553 | 0.0063 | 0.4975 |
| HMR_2017 | Androgen metabolism | -107.3987 | 108734.1463 | -108.3595 | 108772.0383 | 0.9608 | 0.0021 | 0.4992 |
| HMR_2018 | Androgen metabolism | -385.6753 | 158501.3962 | -382.12 | 160072.8036 | 3.5553 | -0.0063 | 0.5025 |
| HMR_2019 | Androgen metabolism | 107.3988 | 108734.1475 | 108.3595 | 108772.0396 | 0.9607 | -0.0021 | 0.5008 |
| HMR_1440 | Cholesterol biosynthesis 1 (Bloch pathway) | 2.393 | 2.5733 | 2.3818 | 2.5221 | 0.0112 | 0.005 | 0.498 |
| HMR_1445 | Cholesterol biosynthesis 1 (Bloch pathway) | 4.2173 | 2.394 | 4.1488 | 2.3437 | 0.0685 | 0.0315 | 0.4874 |
| HMR_1448 | Cholesterol biosynthesis 1 (Bloch pathway) | -4.2152 | 2.3919 | -4.1468 | 2.3416 | 0.0684 | -0.0314 | 0.5125 |
| HMR_1454 | Cholesterol biosynthesis 1 (Bloch pathway) | 1.4015 | 0.275 | 1.3785 | 0.2694 | 0.023 | 0.0312 | 0.4876 |
| HMR_1465 | Cholesterol biosynthesis 1 (Bloch pathway) | -0.7119 | 0.0667 | -0.7011 | 0.0654 | 0.0108 | -0.0298 | 0.5119 |
| HMR_1467 | Cholesterol biosynthesis 1 (Bloch pathway) | 0.7408 | 0.0677 | 0.731 | 0.0664 | 0.0099 | 0.0269 | 0.4893 |
| HMR_1470 | Cholesterol biosynthesis 1 (Bloch pathway) | 0.7729 | 0.0683 | 0.7639 | 0.0671 | 0.009 | 0.0245 | 0.4902 |
| HMR_1473 | Cholesterol biosynthesis 1 (Bloch pathway) | 0.8062 | 0.0686 | 0.798 | 0.0675 | 0.0082 | 0.0223 | 0.4911 |
| HMR_1477 | Cholesterol biosynthesis 1 (Bloch pathway) | 0.5391 | 0.0626 | 0.5344 | 0.0618 | 0.0048 | 0.0135 | 0.4946 |
| HMR_1478 | Cholesterol biosynthesis 1 (Bloch pathway) | 0.5715 | 0.0615 | 0.5675 | 0.0608 | 0.0041 | 0.0116 | 0.4954 |
| HMR_1479 | Cholesterol biosynthesis 1 (Bloch pathway) | 0.6036 | 0.0614 | 0.6001 | 0.0608 | 0.0035 | 0.01 | 0.496 |

| | | | | | | | | |
|---|---|---|---|---|---|---|---|---|
| HMR_1484 | Cholesterol biosynthesis 1 (Bloch pathway) | 0.6364 | 0.0617 | 0.6334 | 0.0612 | 0.003 | 0.0085 | 0.4966 |
| HMR_1490 | Cholesterol biosynthesis 1 (Bloch pathway) | 0.6684 | 0.0622 | 0.666 | 0.0617 | 0.0024 | 0.0068 | 0.4973 |
| HMR_1493 | Cholesterol biosynthesis 1 (Bloch pathway) | 0.7014 | 0.0625 | 0.6995 | 0.0621 | 0.0019 | 0.0054 | 0.4978 |
| HMR_1494 | Cholesterol biosynthesis 1 (Bloch pathway) | 0.7353 | 0.0625 | 0.7338 | 0.0622 | 0.0015 | 0.0042 | 0.4983 |
| HMR_1495 | Cholesterol biosynthesis 1 (Bloch pathway) | 0.3852 | 0.0823 | 0.3846 | 0.082 | 0.0006 | 0.0015 | 0.4994 |
| HMR_1496 | Cholesterol biosynthesis 1 (Bloch pathway) | 0.3851 | 0.0822 | 0.3845 | 0.082 | 0.0006 | 0.0015 | 0.4994 |
| HMR_1500 | Cholesterol biosynthesis 1 (Bloch pathway) | 0.7901 | 0.0655 | 0.7894 | 0.0652 | 0.0007 | 0.002 | 0.4992 |
| HMR_1502 | Cholesterol biosynthesis 1 (Bloch pathway) | 0.8067 | 0.0685 | 0.8063 | 0.0682 | 0.0005 | 0.0013 | 0.4995 |
| HMR_1503 | Cholesterol biosynthesis 1 (Bloch pathway) | 0.8247 | 0.071 | 0.8244 | 0.0708 | 0.0003 | 0.0007 | 0.4997 |
| HMR_1504 | Cholesterol biosynthesis 1 (Bloch pathway) | 0.8437 | 0.0731 | 0.8436 | 0.0729 | 0.0001 | 0.0003 | 0.4999 |
| HMR_1505 | Cholesterol biosynthesis 1 (Bloch pathway) | 0.8641 | 0.0747 | 0.8641 | 0.0746 | 1.00E-04 | 0.0001 | 0.4999 |
| HMR_1509 | Cholesterol biosynthesis 1 (Bloch pathway) | 0.8851 | 0.0758 | 0.8852 | 0.0757 | 1.00E-04 | -0.0002 | 0.5001 |
| HMR_1512 | Cholesterol biosynthesis 1 (Bloch pathway) | 0.5962 | 0.0746 | 0.5964 | 0.0746 | 0.0002 | -0.0004 | 0.5002 |
| HMR_1516 | Cholesterol biosynthesis 1 (Bloch pathway) | 0.2998 | 0.0328 | 0.3 | 0.0328 | 0.0002 | -0.0008 | 0.5003 |
| HMR_1519 | Cholesterol biosynthesis 1 (Bloch pathway) | 0.3084 | 0.0316 | 0.3086 | 0.0317 | 0.0003 | -0.001 | 0.5004 |
| HMR_1526 | Cholesterol biosynthesis 1 (Bloch pathway) | 0.3073 | 0.035 | 0.3076 | 0.0351 | 0.0004 | -0.0014 | 0.5005 |
| HMR_1533 | Cholesterol biosynthesis 2 | 0.3154 | 0.0596 | 0.3154 | 0.0596 | 1.00E-04 | -0.0003 | 0.5001 |
| HMR_1557 | Cholesterol biosynthesis 2 | 0.5354 | 0.1579 | 0.5362 | 0.1583 | 0.0008 | -0.0014 | 0.5006 |
| HMR_1558 | Cholesterol biosynthesis 2 | 0.5351 | 0.1578 | 0.5359 | 0.1583 | 0.0008 | -0.0014 | 0.5006 |
| HMR_1565 | Cholesterol biosynthesis 2 | 1.058 | 0.1243 | 1.0598 | 0.1246 | 0.0018 | -0.0036 | 0.5014 |
| HMR_1535 | Cholesterol biosynthesis 3 (Kandustch-Russell pathway) | 0.3002 | 0.0297 | 0.2975 | 0.0294 | 0.0027 | 0.011 | 0.4956 |
| HMR_1536 | Cholesterol biosynthesis 3 (Kandustch-Russell pathway) | 0.3236 | 0.0272 | 0.3215 | 0.0269 | 0.0021 | 0.0088 | 0.4965 |
| HMR_1538 | Cholesterol biosynthesis 3 (Kandustch-Russell pathway) | 0.3427 | 0.0267 | 0.3411 | 0.0264 | 0.0016 | 0.0068 | 0.4973 |
| HMR_1539 | Cholesterol biosynthesis 3 (Kandustch-Russell pathway) | 0.359 | 0.027 | 0.3578 | 0.0268 | 0.0012 | 0.0051 | 0.498 |
| HMR_1540 | Cholesterol biosynthesis 3 (Kandustch-Russell pathway) | 0.3738 | 0.0278 | 0.373 | 0.0277 | 0.0008 | 0.0036 | 0.4986 |
| HMR_1543 | Cholesterol biosynthesis 3 (Kandustch-Russell pathway) | 0.386 | 0.0288 | 0.3855 | 0.0287 | 0.0005 | 0.0021 | 0.4992 |
| HMR_1544 | Cholesterol biosynthesis 3 (Kandustch-Russell pathway) | 0.3973 | 0.0298 | 0.3971 | 0.0298 | 0.0002 | 0.0008 | 0.4997 |
| HMR_1545 | Cholesterol biosynthesis 3 (Kandustch-Russell pathway) | 0.4076 | 0.031 | 0.4076 | 0.031 | 0.00E+00 | 0.00E+00 | 0.5 |
| HMR_1546 | Cholesterol biosynthesis 3 (Kandustch-Russell pathway) | 0.4173 | 0.0323 | 0.4175 | 0.0323 | 0.0002 | -0.0006 | 0.5003 |
| HMR_1547 | Cholesterol biosynthesis 3 (Kandustch-Russell pathway) | 0.4258 | 0.0337 | 0.4261 | 0.0337 | 0.0004 | -0.0014 | 0.5006 |
| HMR_1548 | Cholesterol biosynthesis 3 (Kandustch-Russell pathway) | 0.4322 | 0.0353 | 0.4328 | 0.0354 | 0.0006 | -0.0023 | 0.5009 |
| HMR_1549 | Cholesterol biosynthesis 3 (Kandustch-Russell pathway) | 0.4381 | 0.0372 | 0.4389 | 0.0373 | 0.0008 | -0.0029 | 0.5012 |
| HMR_1550 | Cholesterol biosynthesis 3 (Kandustch-Russell pathway) | 0.4431 | 0.0398 | 0.444 | 0.0399 | 0.0009 | -0.0032 | 0.5013 |
| HMR_1551 | Cholesterol biosynthesis 3 (Kandustch-Russell pathway) | 0.4472 | 0.0433 | 0.4482 | 0.0434 | 0.001 | -0.0033 | 0.5013 |
| HMR_1552 | Cholesterol biosynthesis 3 (Kandustch-Russell pathway) | 0.4493 | 0.0483 | 0.4504 | 0.0485 | 0.0011 | -0.0036 | 0.5014 |
| HMR_1553 | Cholesterol biosynthesis 3 (Kandustch-Russell pathway) | 0.7576 | 0.1004 | 0.7589 | 0.1006 | 0.0014 | -0.0031 | 0.5012 |
| HMR_1437 | Cholesterol metabolism | 10.0304 | 86.9525 | 9.4943 | 78.5243 | 0.5361 | 0.0417 | 0.4834 |
| HMR_1451 | Cholesterol metabolism | 4.2136 | 2.3905 | 4.1453 | 2.3401 | 0.0683 | 0.0314 | 0.4875 |
| HMR_1457 | Cholesterol metabolism | 1.4035 | 0.2687 | 1.3807 | 0.2631 | 0.0227 | 0.0311 | 0.4876 |
| HMR_1460 | Cholesterol metabolism | 1.4109 | 0.2661 | 1.3886 | 0.2606 | 0.0223 | 0.0307 | 0.4878 |
| HMR_1570 | Cholesterol metabolism | 0.3097 | 0.059 | 0.3098 | 0.059 | 0.00E+00 | -0.0001 | 0.5001 |
| HMR_1576 | Cholesterol metabolism | 2.5368 | 6.6399 | 2.6129 | 6.9998 | 0.076 | -0.0206 | 0.5082 |
| HMR_1577 | Cholesterol metabolism | 27.4809 | 220.6615 | 26.7085 | 212.0409 | 0.7724 | 0.0371 | 0.4852 |
| HMR_3105 | Cholesterol metabolism | 1.2594 | 2.6332 | 1.2965 | 2.5966 | 0.037 | -0.0162 | 0.5065 |

| | | | | | | | | |
|---|---|---|---|---|---|---|---|---|
| HMR_4630 | Cholesterol metabolism | 1.8281 | 0.9899 | 1.7702 | 0.9367 | 0.0578 | 0.0417 | 0.4834 |
| HMR_2029 | Estrogen metabolism | 1.6869 | 0.799 | 1.7119 | 0.8234 | 0.025 | -0.0197 | 0.5078 |
| HMR_2030 | Estrogen metabolism | 1.6781 | 0.7993 | 1.7032 | 0.8237 | 0.0251 | -0.0197 | 0.5078 |
| HMR_2031 | Estrogen metabolism | 1.6678 | 0.8041 | 1.6927 | 0.8285 | 0.0249 | -0.0195 | 0.5078 |
| HMR_2032 | Estrogen metabolism | 2.8483 | 2.0063 | 2.9294 | 2.0978 | 0.081 | -0.04 | 0.516 |
| HMR_2033 | Estrogen metabolism | 2.8416 | 2.0057 | 2.9226 | 2.0972 | 0.081 | -0.04 | 0.516 |
| HMR_2034 | Estrogen metabolism | 2.8339 | 2.0086 | 2.9148 | 2.0999 | 0.0809 | -0.0399 | 0.5159 |
| HMR_2036 | Estrogen metabolism | 1.9844 | 2.424 | 1.978 | 2.4157 | 0.0064 | 0.0029 | 0.4988 |
| HMR_2037 | Estrogen metabolism | 1.1702 | 1.304 | 1.1551 | 1.2684 | 0.0151 | 0.0094 | 0.4962 |
| HMR_2038 | Estrogen metabolism | 4.4601 | 3.4157 | 4.3445 | 3.2726 | 0.1156 | 0.0447 | 0.4822 |
| HMR_2088 | Estrogen metabolism | -190.828 | 16891.4807 | -188.0886 | 17039.6951 | 2.7394 | -0.0149 | 0.5059 |
| HMR_2089 | Estrogen metabolism | 190.8287 | 16891.4736 | 188.0892 | 17039.6879 | 2.7395 | 0.0149 | 0.4941 |
| HMR_3537 | Formation and hydrolysis of cholesterol esters | 9.8556 | 11.5616 | 9.9868 | 11.8421 | 0.1311 | -0.0271 | 0.5108 |
| HMR_3538 | Formation and hydrolysis of cholesterol esters | 0.0242 | 0.0002 | 0.0242 | 0.0002 | 0.00E+00 | -0.0001 | 0.5001 |
| HMR_3539 | Formation and hydrolysis of cholesterol esters | 0.0242 | 0.0002 | 0.0242 | 0.0002 | 0.00E+00 | -0.0001 | 0.5001 |
| HMR_3540 | Formation and hydrolysis of cholesterol esters | 0.0942 | 0.0035 | 0.095 | 0.0035 | 0.0008 | -0.009 | 0.5036 |
| HMR_3541 | Formation and hydrolysis of cholesterol esters | 0.0242 | 0.0002 | 0.0242 | 0.0002 | 0.00E+00 | -0.0001 | 0.5001 |
| HMR_3542 | Formation and hydrolysis of cholesterol esters | 0.0242 | 0.0002 | 0.0242 | 0.0002 | 0.00E+00 | -0.0001 | 0.5001 |
| HMR_3543 | Formation and hydrolysis of cholesterol esters | 0.0242 | 0.0002 | 0.0242 | 0.0002 | 0.00E+00 | -0.0001 | 0.5001 |
| HMR_3544 | Formation and hydrolysis of cholesterol esters | 0.0242 | 0.0002 | 0.0242 | 0.0002 | 0.00E+00 | -0.0001 | 0.5001 |
| HMR_3545 | Formation and hydrolysis of cholesterol esters | 1.3515 | 0.2242 | 1.3697 | 0.2295 | 0.0182 | -0.027 | 0.5108 |
| HMR_3546 | Formation and hydrolysis of cholesterol esters | 0.3935 | 0.0234 | 0.3988 | 0.0239 | 0.0054 | -0.0246 | 0.5098 |
| HMR_3547 | Formation and hydrolysis of cholesterol esters | 0.0242 | 0.0002 | 0.0242 | 0.0002 | 0.00E+00 | -0.0001 | 0.5001 |
| HMR_3548 | Formation and hydrolysis of cholesterol esters | 0.0242 | 0.0002 | 0.0242 | 0.0002 | 0.00E+00 | -0.0001 | 0.5001 |
| HMR_3549 | Formation and hydrolysis of cholesterol esters | 0.0242 | 0.0002 | 0.0242 | 0.0002 | 0.00E+00 | -0.0001 | 0.5001 |
| HMR_3550 | Formation and hydrolysis of cholesterol esters | 0.0242 | 0.0002 | 0.0242 | 0.0002 | 0.00E+00 | -0.0001 | 0.5001 |
| HMR_3551 | Formation and hydrolysis of cholesterol esters | 0.1327 | 0.0053 | 0.1342 | 0.0054 | 0.0015 | -0.0146 | 0.5058 |
| HMR_3552 | Formation and hydrolysis of cholesterol esters | 1.9201 | 0.4467 | 1.9459 | 0.4575 | 0.0258 | -0.0271 | 0.5108 |
| HMR_3553 | Formation and hydrolysis of cholesterol esters | 0.0242 | 0.0002 | 0.0242 | 0.0002 | 0.00E+00 | -0.0001 | 0.5001 |
| HMR_3554 | Formation and hydrolysis of cholesterol esters | 0.0242 | 0.0002 | 0.0242 | 0.0002 | 0.00E+00 | -0.0001 | 0.5001 |
| HMR_3555 | Formation and hydrolysis of cholesterol esters | 0.0242 | 0.0002 | 0.0242 | 0.0002 | 0.00E+00 | -0.0001 | 0.5001 |
| HMR_3556 | Formation and hydrolysis of cholesterol esters | 0.0242 | 0.0002 | 0.0242 | 0.0002 | 0.00E+00 | -0.0001 | 0.5001 |
| HMR_3557 | Formation and hydrolysis of cholesterol esters | 0.0242 | 0.0002 | 0.0242 | 0.0002 | 0.00E+00 | -0.0001 | 0.5001 |
| HMR_3558 | Formation and hydrolysis of cholesterol esters | 0.0242 | 0.0002 | 0.0242 | 0.0002 | 0.00E+00 | -0.0001 | 0.5001 |
| HMR_3559 | Formation and hydrolysis of cholesterol esters | 0.0242 | 0.0002 | 0.0242 | 0.0002 | 0.00E+00 | -0.0001 | 0.5001 |
| HMR_3560 | Formation and hydrolysis of cholesterol esters | 0.0242 | 0.0002 | 0.0242 | 0.0002 | 0.00E+00 | -0.0001 | 0.5001 |
| HMR_3561 | Formation and hydrolysis of cholesterol esters | 0.0242 | 0.0002 | 0.0242 | 0.0002 | 0.00E+00 | -0.0001 | 0.5001 |
| HMR_3562 | Formation and hydrolysis of cholesterol esters | 0.0242 | 0.0002 | 0.0242 | 0.0002 | 0.00E+00 | -0.0001 | 0.5001 |
| HMR_3563 | Formation and hydrolysis of cholesterol esters | 0.0242 | 0.0002 | 0.0242 | 0.0002 | 0.00E+00 | -0.0001 | 0.5001 |
| HMR_3564 | Formation and hydrolysis of cholesterol esters | 0.0242 | 0.0002 | 0.0242 | 0.0002 | 0.00E+00 | -0.0001 | 0.5001 |
| HMR_3565 | Formation and hydrolysis of cholesterol esters | 0.0242 | 0.0002 | 0.0242 | 0.0002 | 0.00E+00 | -0.0001 | 0.5001 |
| HMR_3566 | Formation and hydrolysis of cholesterol esters | 0.0242 | 0.0002 | 0.0242 | 0.0002 | 0.00E+00 | -0.0001 | 0.5001 |
| HMR_3567 | Formation and hydrolysis of cholesterol esters | 0.0242 | 0.0002 | 0.0242 | 0.0002 | 0.00E+00 | -0.0001 | 0.5001 |
| HMR_3568 | Formation and hydrolysis of cholesterol esters | 0.0242 | 0.0002 | 0.0242 | 0.0002 | 0.00E+00 | -0.0001 | 0.5001 |

| | | | | | | | | |
|---|---|---|---|---|---|---|---|---|
| HMR_3569 | Formation and hydrolysis of cholesterol esters | 0.0242 | 0.0002 | 0.0242 | 0.0002 | 0.00E+00 | -0.0001 | 0.5001 |
| HMR_3570 | Formation and hydrolysis of cholesterol esters | 0.0242 | 0.0002 | 0.0242 | 0.0002 | 0.00E+00 | -0.0001 | 0.5001 |
| HMR_3571 | Formation and hydrolysis of cholesterol esters | 0.0242 | 0.0002 | 0.0242 | 0.0002 | 0.00E+00 | -0.0001 | 0.5001 |
| HMR_3572 | Formation and hydrolysis of cholesterol esters | 0.0242 | 0.0002 | 0.0242 | 0.0002 | 0.00E+00 | -0.0001 | 0.5001 |
| HMR_3573 | Formation and hydrolysis of cholesterol esters | 0.0242 | 0.0002 | 0.0242 | 0.0002 | 0.00E+00 | -0.0001 | 0.5001 |
| HMR_3574 | Formation and hydrolysis of cholesterol esters | 0.0816 | 0.0029 | 0.0821 | 0.0029 | 0.0005 | -0.0065 | 0.5026 |
| HMR_3575 | Formation and hydrolysis of cholesterol esters | 0.0242 | 0.0002 | 0.0242 | 0.0002 | 0.00E+00 | -0.0001 | 0.5001 |
| HMR_3576 | Formation and hydrolysis of cholesterol esters | 0.0242 | 0.0002 | 0.0242 | 0.0002 | 0.00E+00 | -0.0001 | 0.5001 |
| HMR_3577 | Formation and hydrolysis of cholesterol esters | 0.0877 | 0.0032 | 0.0883 | 0.0032 | 0.0006 | -0.0078 | 0.5031 |
| HMR_3578 | Formation and hydrolysis of cholesterol esters | 0.0418 | 0.0008 | 0.0418 | 0.0008 | 0.00E+00 | -0.0003 | 0.5001 |
| HMR_3579 | Formation and hydrolysis of cholesterol esters | 0.0242 | 0.0002 | 0.0242 | 0.0002 | 0.00E+00 | -0.0001 | 0.5001 |
| HMR_3580 | Formation and hydrolysis of cholesterol esters | 0.0242 | 0.0002 | 0.0242 | 0.0002 | 0.00E+00 | -0.0001 | 0.5001 |
| HMR_3581 | Formation and hydrolysis of cholesterol esters | 0.0707 | 0.0024 | 0.071 | 0.0025 | 0.0003 | -0.0042 | 0.5017 |
| HMR_3582 | Formation and hydrolysis of cholesterol esters | 0.0242 | 0.0002 | 0.0242 | 0.0002 | 0.00E+00 | -0.0001 | 0.5001 |
| HMR_3583 | Formation and hydrolysis of cholesterol esters | 0.0242 | 0.0002 | 0.0242 | 0.0002 | 0.00E+00 | -0.0001 | 0.5001 |
| HMR_3584 | Formation and hydrolysis of cholesterol esters | 0.0242 | 0.0002 | 0.0242 | 0.0002 | 0.00E+00 | -0.0001 | 0.5001 |
| HMR_3585 | Formation and hydrolysis of cholesterol esters | 0.0242 | 0.0002 | 0.0242 | 0.0002 | 0.00E+00 | -0.0001 | 0.5001 |
| HMR_3586 | Formation and hydrolysis of cholesterol esters | 5.1855 | 3.2182 | 5.2548 | 3.2963 | 0.0693 | -0.0272 | 0.5108 |
| HMR_3587 | Formation and hydrolysis of cholesterol esters | 0.0242 | 0.0002 | 0.0242 | 0.0002 | 0.00E+00 | -0.0001 | 0.5001 |
| HMR_3588 | Formation and hydrolysis of cholesterol esters | 0.0759 | 0.0027 | 0.0763 | 0.0027 | 0.0004 | -0.0053 | 0.5021 |
| HMR_3589 | Formation and hydrolysis of cholesterol esters | 0.5031 | 0.0354 | 0.5099 | 0.0361 | 0.0068 | -0.0256 | 0.5102 |
| HMR_3590 | Formation and hydrolysis of cholesterol esters | 0.0418 | 0.0008 | 0.0418 | 0.0008 | 0.00E+00 | -0.0003 | 0.5001 |
| HMR_3591 | Formation and hydrolysis of cholesterol esters | 0.0242 | 0.0002 | 0.0242 | 0.0002 | 0.00E+00 | -0.0001 | 0.5001 |
| HMR_3592 | Formation and hydrolysis of cholesterol esters | 0.0242 | 0.0002 | 0.0242 | 0.0002 | 0.00E+00 | -0.0001 | 0.5001 |
| HMR_3593 | Formation and hydrolysis of cholesterol esters | 0.0242 | 0.0002 | 0.0242 | 0.0002 | 0.00E+00 | -0.0001 | 0.5001 |
| HMR_3594 | Formation and hydrolysis of cholesterol esters | 0.0242 | 0.0002 | 0.0242 | 0.0002 | 0.00E+00 | -0.0001 | 0.5001 |
| HMR_3595 | Formation and hydrolysis of cholesterol esters | 0.0242 | 0.0002 | 0.0242 | 0.0002 | 0.00E+00 | -0.0001 | 0.5001 |
| HMR_3596 | Formation and hydrolysis of cholesterol esters | 0.0242 | 0.0002 | 0.0242 | 0.0002 | 0.00E+00 | -0.0001 | 0.5001 |
| HMR_3622 | Formation and hydrolysis of cholesterol esters | -8.9631 | 10.8406 | -9.0551 | 11.0872 | 0.092 | 0.0196 | 0.4922 |
| HMR_3625 | Formation and hydrolysis of cholesterol esters | 2.0306 | 1.4451 | 2.0546 | 1.48 | 0.0241 | -0.0141 | 0.5056 |
| HMR_3626 | Formation and hydrolysis of cholesterol esters | 2.0306 | 1.4451 | 2.0546 | 1.48 | 0.0241 | -0.0141 | 0.5056 |
| HMR_3627 | Formation and hydrolysis of cholesterol esters | 2.1174 | 1.4568 | 2.1417 | 1.4917 | 0.0243 | -0.0141 | 0.5056 |
| HMR_3628 | Formation and hydrolysis of cholesterol esters | 2.0306 | 1.4451 | 2.0546 | 1.48 | 0.0241 | -0.0141 | 0.5056 |
| HMR_3629 | Formation and hydrolysis of cholesterol esters | 2.0306 | 1.4451 | 2.0546 | 1.48 | 0.0241 | -0.0141 | 0.5056 |
| HMR_3630 | Formation and hydrolysis of cholesterol esters | 2.0306 | 1.4451 | 2.0546 | 1.48 | 0.0241 | -0.0141 | 0.5056 |
| HMR_3631 | Formation and hydrolysis of cholesterol esters | 2.0306 | 1.4451 | 2.0546 | 1.48 | 0.0241 | -0.0141 | 0.5056 |
| HMR_3632 | Formation and hydrolysis of cholesterol esters | 3.2111 | 1.8937 | 3.2468 | 1.9379 | 0.0357 | -0.0182 | 0.5073 |
| HMR_3633 | Formation and hydrolysis of cholesterol esters | 2.3385 | 1.5167 | 2.3648 | 1.5524 | 0.0262 | -0.015 | 0.506 |
| HMR_3634 | Formation and hydrolysis of cholesterol esters | 2.0306 | 1.4451 | 2.0546 | 1.48 | 0.0241 | -0.0141 | 0.5056 |
| HMR_3635 | Formation and hydrolysis of cholesterol esters | 2.0306 | 1.4451 | 2.0546 | 1.48 | 0.0241 | -0.0141 | 0.5056 |
| HMR_3636 | Formation and hydrolysis of cholesterol esters | 2.0306 | 1.4451 | 2.0546 | 1.48 | 0.0241 | -0.0141 | 0.5056 |
| HMR_3637 | Formation and hydrolysis of cholesterol esters | 2.0306 | 1.4451 | 2.0546 | 1.48 | 0.0241 | -0.0141 | 0.5056 |
| HMR_3638 | Formation and hydrolysis of cholesterol esters | 2.1435 | 1.4615 | 2.1682 | 1.4965 | 0.0247 | -0.0143 | 0.5057 |
| HMR_3639 | Formation and hydrolysis of cholesterol esters | 2.0306 | 1.4451 | 2.0546 | 1.48 | 0.0241 | -0.0141 | 0.5056 |

| | | | | | | | | |
|---|---|---|---|---|---|---|---|---|
| HMR_3640 | Formation and hydrolysis of cholesterol esters | 2.0306 | 1.4451 | 2.0546 | 1.48 | 0.0241 | -0.0141 | 0.5056 |
| HMR_3641 | Formation and hydrolysis of cholesterol esters | 3.7523 | 2.1752 | 3.7932 | 2.2261 | 0.0409 | -0.0195 | 0.5078 |
| HMR_3642 | Formation and hydrolysis of cholesterol esters | 2.0306 | 1.4451 | 2.0546 | 1.48 | 0.0241 | -0.0141 | 0.5056 |
| HMR_3643 | Formation and hydrolysis of cholesterol esters | 0.0254 | 0.0002 | 0.0254 | 0.0002 | 0.00E+00 | 1.00E-04 | 0.5 |
| HMR_3644 | Formation and hydrolysis of cholesterol esters | 2.0306 | 1.4451 | 2.0546 | 1.48 | 0.0241 | -0.0141 | 0.5056 |
| HMR_3645 | Formation and hydrolysis of cholesterol esters | 2.0306 | 1.4451 | 2.0546 | 1.48 | 0.0241 | -0.0141 | 0.5056 |
| HMR_3646 | Formation and hydrolysis of cholesterol esters | 2.0306 | 1.4451 | 2.0546 | 1.48 | 0.0241 | -0.0141 | 0.5056 |
| HMR_3647 | Formation and hydrolysis of cholesterol esters | 2.0306 | 1.4451 | 2.0546 | 1.48 | 0.0241 | -0.0141 | 0.5056 |
| HMR_3648 | Formation and hydrolysis of cholesterol esters | 2.0306 | 1.4451 | 2.0546 | 1.48 | 0.0241 | -0.0141 | 0.5056 |
| HMR_3649 | Formation and hydrolysis of cholesterol esters | 2.0306 | 1.4451 | 2.0546 | 1.48 | 0.0241 | -0.0141 | 0.5056 |
| HMR_3650 | Formation and hydrolysis of cholesterol esters | 2.0306 | 1.4451 | 2.0546 | 1.48 | 0.0241 | -0.0141 | 0.5056 |
| HMR_3651 | Formation and hydrolysis of cholesterol esters | 2.0306 | 1.4451 | 2.0546 | 1.48 | 0.0241 | -0.0141 | 0.5056 |
| HMR_3652 | Formation and hydrolysis of cholesterol esters | 2.0306 | 1.4451 | 2.0546 | 1.48 | 0.0241 | -0.0141 | 0.5056 |
| HMR_3653 | Formation and hydrolysis of cholesterol esters | 2.0306 | 1.4451 | 2.0546 | 1.48 | 0.0241 | -0.0141 | 0.5056 |
| HMR_3654 | Formation and hydrolysis of cholesterol esters | 2.0306 | 1.4451 | 2.0546 | 1.48 | 0.0241 | -0.0141 | 0.5056 |
| HMR_3655 | Formation and hydrolysis of cholesterol esters | 2.0306 | 1.4451 | 2.0546 | 1.48 | 0.0241 | -0.0141 | 0.5056 |
| HMR_3656 | Formation and hydrolysis of cholesterol esters | 2.0306 | 1.4451 | 2.0546 | 1.48 | 0.0241 | -0.0141 | 0.5056 |
| HMR_3657 | Formation and hydrolysis of cholesterol esters | 2.0306 | 1.4451 | 2.0546 | 1.48 | 0.0241 | -0.0141 | 0.5056 |
| HMR_3658 | Formation and hydrolysis of cholesterol esters | 2.0306 | 1.4451 | 2.0546 | 1.48 | 0.0241 | -0.0141 | 0.5056 |
| HMR_3659 | Formation and hydrolysis of cholesterol esters | 2.0306 | 1.4451 | 2.0546 | 1.48 | 0.0241 | -0.0141 | 0.5056 |
| HMR_3660 | Formation and hydrolysis of cholesterol esters | 2.0306 | 1.4451 | 2.0546 | 1.48 | 0.0241 | -0.0141 | 0.5056 |
| HMR_3661 | Formation and hydrolysis of cholesterol esters | 2.1079 | 1.4554 | 2.1321 | 1.4903 | 0.0242 | -0.0141 | 0.5056 |
| HMR_3662 | Formation and hydrolysis of cholesterol esters | 2.0306 | 1.4451 | 2.0546 | 1.48 | 0.0241 | -0.0141 | 0.5056 |
| HMR_3663 | Formation and hydrolysis of cholesterol esters | 2.0306 | 1.4451 | 2.0546 | 1.48 | 0.0241 | -0.0141 | 0.5056 |
| HMR_3664 | Formation and hydrolysis of cholesterol esters | 2.0505 | 1.8992 | 2.0746 | 1.9444 | 0.0242 | -0.0123 | 0.5049 |
| HMR_3665 | Formation and hydrolysis of cholesterol esters | 2.0393 | 1.4456 | 2.0633 | 1.4805 | 0.024 | -0.0141 | 0.5056 |
| HMR_3666 | Formation and hydrolysis of cholesterol esters | 2.0306 | 1.4451 | 2.0546 | 1.48 | 0.0241 | -0.0141 | 0.5056 |
| HMR_3667 | Formation and hydrolysis of cholesterol esters | 2.0306 | 1.4451 | 2.0546 | 1.48 | 0.0241 | -0.0141 | 0.5056 |
| HMR_3668 | Formation and hydrolysis of cholesterol esters | 2.0987 | 1.4541 | 2.1227 | 1.489 | 0.024 | -0.014 | 0.5056 |
| HMR_3669 | Formation and hydrolysis of cholesterol esters | 2.0306 | 1.4451 | 2.0546 | 1.48 | 0.0241 | -0.0141 | 0.5056 |
| HMR_3670 | Formation and hydrolysis of cholesterol esters | 2.0306 | 1.4451 | 2.0546 | 1.48 | 0.0241 | -0.0141 | 0.5056 |
| HMR_3671 | Formation and hydrolysis of cholesterol esters | 0.0254 | 0.0002 | 0.0254 | 0.0002 | 0.00E+00 | 1.00E-04 | 0.5 |
| HMR_3672 | Formation and hydrolysis of cholesterol esters | 0.0254 | 0.0002 | 0.0254 | 0.0002 | 0.00E+00 | 1.00E-04 | 0.5 |
| HMR_3673 | Formation and hydrolysis of cholesterol esters | 6.7477 | 4.8697 | 6.8198 | 4.9841 | 0.0721 | -0.023 | 0.5092 |
| HMR_3674 | Formation and hydrolysis of cholesterol esters | 2.0306 | 1.4451 | 2.0546 | 1.48 | 0.0241 | -0.0141 | 0.5056 |
| HMR_3675 | Formation and hydrolysis of cholesterol esters | 2.1013 | 1.4543 | 2.1255 | 1.4892 | 0.0242 | -0.0141 | 0.5056 |
| HMR_3676 | Formation and hydrolysis of cholesterol esters | 2.4202 | 1.5461 | 2.4479 | 1.5826 | 0.0277 | -0.0156 | 0.5062 |
| HMR_3677 | Formation and hydrolysis of cholesterol esters | 2.0393 | 1.4456 | 2.0633 | 1.4805 | 0.024 | -0.0141 | 0.5056 |
| HMR_3678 | Formation and hydrolysis of cholesterol esters | 2.0306 | 1.4451 | 2.0546 | 1.48 | 0.0241 | -0.0141 | 0.5056 |
| HMR_3679 | Formation and hydrolysis of cholesterol esters | 2.0306 | 1.4451 | 2.0546 | 1.48 | 0.0241 | -0.0141 | 0.5056 |
| HMR_3680 | Formation and hydrolysis of cholesterol esters | 2.0306 | 1.4451 | 2.0546 | 1.48 | 0.0241 | -0.0141 | 0.5056 |
| HMR_3681 | Formation and hydrolysis of cholesterol esters | 2.0306 | 1.4451 | 2.0546 | 1.48 | 0.0241 | -0.0141 | 0.5056 |
| HMR_3682 | Formation and hydrolysis of cholesterol esters | 2.0306 | 1.4451 | 2.0546 | 1.48 | 0.0241 | -0.0141 | 0.5056 |
| HMR_3683 | Formation and hydrolysis of cholesterol esters | 0.0254 | 0.0002 | 0.0254 | 0.0002 | 0.00E+00 | 1.00E-04 | 0.5 |

| | | | | | | | | |
|---|---|---|---|---|---|---|---|---|
| HMR_3684 | Formation and hydrolysis of cholesterol esters | 2.0109 | 1.445 | 2.035 | 1.4799 | 0.0241 | -0.0141 | 0.5056 |
| HMR_3685 | Formation and hydrolysis of cholesterol esters | 2.0109 | 1.445 | 2.035 | 1.4799 | 0.0241 | -0.0141 | 0.5056 |
| HMR_3686 | Formation and hydrolysis of cholesterol esters | 1.9969 | 1.4548 | 2.0206 | 1.4897 | 0.0238 | -0.0138 | 0.5055 |
| HMR_3687 | Formation and hydrolysis of cholesterol esters | 2.0109 | 1.445 | 2.035 | 1.4799 | 0.0241 | -0.0141 | 0.5056 |
| HMR_3688 | Formation and hydrolysis of cholesterol esters | 2.0109 | 1.445 | 2.035 | 1.4799 | 0.0241 | -0.0141 | 0.5056 |
| HMR_3689 | Formation and hydrolysis of cholesterol esters | 2.0109 | 1.445 | 2.035 | 1.4799 | 0.0241 | -0.0141 | 0.5056 |
| HMR_3690 | Formation and hydrolysis of cholesterol esters | 2.0109 | 1.445 | 2.035 | 1.4799 | 0.0241 | -0.0141 | 0.5056 |
| HMR_3691 | Formation and hydrolysis of cholesterol esters | 1.9664 | 1.7141 | 1.9894 | 1.754 | 0.023 | -0.0123 | 0.5049 |
| HMR_3692 | Formation and hydrolysis of cholesterol esters | 1.9533 | 1.5043 | 1.9762 | 1.5397 | 0.0229 | -0.0131 | 0.5052 |
| HMR_3693 | Formation and hydrolysis of cholesterol esters | 2.0109 | 1.445 | 2.035 | 1.4799 | 0.0241 | -0.0141 | 0.5056 |
| HMR_3694 | Formation and hydrolysis of cholesterol esters | 2.0109 | 1.445 | 2.035 | 1.4799 | 0.0241 | -0.0141 | 0.5056 |
| HMR_3695 | Formation and hydrolysis of cholesterol esters | 2.0109 | 1.445 | 2.035 | 1.4799 | 0.0241 | -0.0141 | 0.5056 |
| HMR_3696 | Formation and hydrolysis of cholesterol esters | 2.0109 | 1.445 | 2.035 | 1.4799 | 0.0241 | -0.0141 | 0.5056 |
| HMR_3697 | Formation and hydrolysis of cholesterol esters | 1.9861 | 1.459 | 2.0097 | 1.4939 | 0.0236 | -0.0137 | 0.5055 |
| HMR_3698 | Formation and hydrolysis of cholesterol esters | 2.0109 | 1.445 | 2.035 | 1.4799 | 0.0241 | -0.0141 | 0.5056 |
| HMR_3699 | Formation and hydrolysis of cholesterol esters | 2.0109 | 1.445 | 2.035 | 1.4799 | 0.0241 | -0.0141 | 0.5056 |
| HMR_3700 | Formation and hydrolysis of cholesterol esters | 1.9862 | 1.7901 | 2.0092 | 1.832 | 0.0231 | -0.0121 | 0.5048 |
| HMR_3701 | Formation and hydrolysis of cholesterol esters | 2.0109 | 1.445 | 2.035 | 1.4799 | 0.0241 | -0.0141 | 0.5056 |
| HMR_3703 | Formation and hydrolysis of cholesterol esters | 2.0109 | 1.445 | 2.035 | 1.4799 | 0.0241 | -0.0141 | 0.5056 |
| HMR_3704 | Formation and hydrolysis of cholesterol esters | 2.0109 | 1.445 | 2.035 | 1.4799 | 0.0241 | -0.0141 | 0.5056 |
| HMR_3705 | Formation and hydrolysis of cholesterol esters | 2.0109 | 1.445 | 2.035 | 1.4799 | 0.0241 | -0.0141 | 0.5056 |
| HMR_3706 | Formation and hydrolysis of cholesterol esters | 2.0109 | 1.445 | 2.035 | 1.4799 | 0.0241 | -0.0141 | 0.5056 |
| HMR_3707 | Formation and hydrolysis of cholesterol esters | 2.0109 | 1.445 | 2.035 | 1.4799 | 0.0241 | -0.0141 | 0.5056 |
| HMR_3708 | Formation and hydrolysis of cholesterol esters | 2.0109 | 1.445 | 2.035 | 1.4799 | 0.0241 | -0.0141 | 0.5056 |
| HMR_3709 | Formation and hydrolysis of cholesterol esters | 2.0109 | 1.445 | 2.035 | 1.4799 | 0.0241 | -0.0141 | 0.5056 |
| HMR_3710 | Formation and hydrolysis of cholesterol esters | 2.0109 | 1.445 | 2.035 | 1.4799 | 0.0241 | -0.0141 | 0.5056 |
| HMR_3711 | Formation and hydrolysis of cholesterol esters | 2.0109 | 1.445 | 2.035 | 1.4799 | 0.0241 | -0.0141 | 0.5056 |
| HMR_3712 | Formation and hydrolysis of cholesterol esters | 2.0109 | 1.445 | 2.035 | 1.4799 | 0.0241 | -0.0141 | 0.5056 |
| HMR_3713 | Formation and hydrolysis of cholesterol esters | 2.0109 | 1.445 | 2.035 | 1.4799 | 0.0241 | -0.0141 | 0.5056 |
| HMR_3714 | Formation and hydrolysis of cholesterol esters | 2.0109 | 1.445 | 2.035 | 1.4799 | 0.0241 | -0.0141 | 0.5056 |
| HMR_3715 | Formation and hydrolysis of cholesterol esters | 2.0109 | 1.445 | 2.035 | 1.4799 | 0.0241 | -0.0141 | 0.5056 |
| HMR_3716 | Formation and hydrolysis of cholesterol esters | 2.0109 | 1.445 | 2.035 | 1.4799 | 0.0241 | -0.0141 | 0.5056 |
| HMR_3717 | Formation and hydrolysis of cholesterol esters | 2.0109 | 1.445 | 2.035 | 1.4799 | 0.0241 | -0.0141 | 0.5056 |
| HMR_3718 | Formation and hydrolysis of cholesterol esters | 2.0109 | 1.445 | 2.035 | 1.4799 | 0.0241 | -0.0141 | 0.5056 |
| HMR_3719 | Formation and hydrolysis of cholesterol esters | 2.0109 | 1.445 | 2.035 | 1.4799 | 0.0241 | -0.0141 | 0.5056 |
| HMR_3720 | Formation and hydrolysis of cholesterol esters | 2.0016 | 1.4535 | 2.0254 | 1.4884 | 0.0238 | -0.0139 | 0.5055 |
| HMR_3721 | Formation and hydrolysis of cholesterol esters | 2.0109 | 1.445 | 2.035 | 1.4799 | 0.0241 | -0.0141 | 0.5056 |
| HMR_3722 | Formation and hydrolysis of cholesterol esters | 2.0109 | 1.445 | 2.035 | 1.4799 | 0.0241 | -0.0141 | 0.5056 |
| HMR_3723 | Formation and hydrolysis of cholesterol esters | 1.9808 | 1.8988 | 2.0042 | 1.9441 | 0.0234 | -0.0119 | 0.5048 |
| HMR_3724 | Formation and hydrolysis of cholesterol esters | 2.0112 | 1.4455 | 2.0353 | 1.4804 | 0.0241 | -0.0141 | 0.5056 |
| HMR_3725 | Formation and hydrolysis of cholesterol esters | 2.0109 | 1.445 | 2.035 | 1.4799 | 0.0241 | -0.0141 | 0.5056 |
| HMR_3726 | Formation and hydrolysis of cholesterol esters | 2.0109 | 1.445 | 2.035 | 1.4799 | 0.0241 | -0.0141 | 0.5056 |
| HMR_3727 | Formation and hydrolysis of cholesterol esters | 2.0064 | 1.4524 | 2.0303 | 1.4872 | 0.0239 | -0.0139 | 0.5056 |
| HMR_3728 | Formation and hydrolysis of cholesterol esters | 2.0109 | 1.445 | 2.035 | 1.4799 | 0.0241 | -0.0141 | 0.5056 |

| | | | | | | | | |
|---|---|---|---|---|---|---|---|---|
| HMR_3729 | Formation and hydrolysis of cholesterol esters | 2.0109 | 1.445 | 2.035 | 1.4799 | 0.0241 | -0.0141 | 0.5056 |
| HMR_3732 | Formation and hydrolysis of cholesterol esters | 2.01 | 1.8849 | 2.0337 | 1.9296 | 0.0237 | -0.0121 | 0.5048 |
| HMR_3733 | Formation and hydrolysis of cholesterol esters | 2.0109 | 1.445 | 2.035 | 1.4799 | 0.0241 | -0.0141 | 0.5056 |
| HMR_3734 | Formation and hydrolysis of cholesterol esters | 2.0036 | 1.4526 | 2.0275 | 1.4875 | 0.0239 | -0.0139 | 0.5056 |
| HMR_3735 | Formation and hydrolysis of cholesterol esters | 1.9453 | 1.5264 | 1.9682 | 1.5623 | 0.0229 | -0.013 | 0.5052 |
| HMR_3736 | Formation and hydrolysis of cholesterol esters | 2.0112 | 1.4455 | 2.0353 | 1.4804 | 0.0241 | -0.0141 | 0.5056 |
| HMR_3737 | Formation and hydrolysis of cholesterol esters | 2.0109 | 1.445 | 2.035 | 1.4799 | 0.0241 | -0.0141 | 0.5056 |
| HMR_3738 | Formation and hydrolysis of cholesterol esters | 2.0109 | 1.445 | 2.035 | 1.4799 | 0.0241 | -0.0141 | 0.5056 |
| HMR_3739 | Formation and hydrolysis of cholesterol esters | 2.0109 | 1.445 | 2.035 | 1.4799 | 0.0241 | -0.0141 | 0.5056 |
| HMR_3740 | Formation and hydrolysis of cholesterol esters | 2.0109 | 1.445 | 2.035 | 1.4799 | 0.0241 | -0.0141 | 0.5056 |
| HMR_3741 | Formation and hydrolysis of cholesterol esters | 2.0109 | 1.445 | 2.035 | 1.4799 | 0.0241 | -0.0141 | 0.5056 |
| HMR_0736 | Sphingolipid metabolism | 1.4736 | 1.4327 | 1.4926 | 1.4624 | 0.019 | -0.0112 | 0.5045 |
| HMR_0738 | Sphingolipid metabolism | 0.1317 | 0.0054 | 0.1318 | 0.0054 | 0.0001 | -0.0011 | 0.5004 |
| HMR_0741 | Sphingolipid metabolism | 0.131 | 0.0054 | 0.1311 | 0.0054 | 0.0001 | -0.0013 | 0.5005 |
| HMR_0744 | Sphingolipid metabolism | 2.8098 | 7.4309 | 2.6836 | 6.7868 | 0.1262 | 0.0335 | 0.4866 |
| HMR_0746 | Sphingolipid metabolism | 9.1906 | 14.069 | 8.875 | 13.1091 | 0.3156 | 0.0605 | 0.4759 |
| HMR_0748 | Sphingolipid metabolism | 6.3926 | 7.1094 | 6.2031 | 6.76 | 0.1895 | 0.0509 | 0.4797 |
| HMR_0750 | Sphingolipid metabolism | 3.6062 | 7.6738 | 3.6007 | 7.5562 | 0.0055 | 0.0014 | 0.4994 |
| HMR_0753 | Sphingolipid metabolism | -2.8965 | 7.9448 | -2.713 | 7.7529 | 0.1835 | -0.0463 | 0.5185 |
| HMR_0754 | Sphingolipid metabolism | 3.0923 | 6.1174 | 2.9841 | 5.7367 | 0.1083 | 0.0315 | 0.4875 |
| HMR_0758 | Sphingolipid metabolism | 6.4408 | 7.106 | 6.2511 | 6.7571 | 0.1897 | 0.051 | 0.4797 |
| HMR_0760 | Sphingolipid metabolism | 2.8532 | 3.793 | 2.7242 | 3.4573 | 0.1289 | 0.0479 | 0.4809 |
| HMR_0761 | Sphingolipid metabolism | 2.1143 | 2.083 | 2.1414 | 2.1371 | 0.0272 | -0.0132 | 0.5053 |
| HMR_0762 | Sphingolipid metabolism | 2.1075 | 2.083 | 2.1346 | 2.1371 | 0.027 | -0.0132 | 0.5052 |
| HMR_0763 | Sphingolipid metabolism | 0.5213 | 4.7506 | 0.524 | 4.8356 | 0.0028 | -0.0009 | 0.5004 |
| HMR_0767 | Sphingolipid metabolism | 2.8432 | 3.7929 | 2.7141 | 3.4573 | 0.129 | 0.0479 | 0.4809 |
| HMR_0773 | Sphingolipid metabolism | 9.2391 | 14.092 | 8.9232 | 13.1308 | 0.3159 | 0.0605 | 0.4759 |
| HMR_0775 | Sphingolipid metabolism | 2.8104 | 7.4445 | 2.6841 | 6.7992 | 0.1263 | 0.0335 | 0.4867 |
| HMR_0779 | Sphingolipid metabolism | 6.4172 | 7.1081 | 6.2276 | 6.7589 | 0.1896 | 0.0509 | 0.4797 |
| HMR_0781 | Sphingolipid metabolism | 6.405 | 7.1087 | 6.2154 | 6.7595 | 0.1895 | 0.0509 | 0.4797 |
| HMR_0795 | Sphingolipid metabolism | 1.9752 | 3.38 | 1.9965 | 3.4391 | 0.0213 | -0.0081 | 0.5032 |
| HMR_8220 | Sphingolipid metabolism | 3.4026 | 6.8101 | 3.322 | 6.4733 | 0.0806 | 0.0221 | 0.4912 |
| HMR_0453 | Glycerolipid metabolism | -68.9636 | 2009.0675 | -85.2528 | 3060.6255 | 16.2892 | 0.2288 | 0.4095 |
| HMR_0456 | Glycerolipid metabolism | 3.629 | 10.9075 | 3.5183 | 10.144 | 0.1106 | 0.0241 | 0.4904 |
| HMR_0458 | Glycerolipid metabolism | 3.3508 | 3.0102 | 3.2894 | 2.8996 | 0.0614 | 0.0253 | 0.4899 |
| HMR_0459 | Glycerolipid metabolism | 3.3496 | 3.0081 | 3.2882 | 2.8975 | 0.0615 | 0.0253 | 0.4899 |
| HMR_0486 | Glycerolipid metabolism | 0.0938 | 0.0045 | 0.0934 | 0.0045 | 0.0005 | 0.0051 | 0.498 |
| HMR_0487 | Glycerolipid metabolism | 0.0839 | 0.0039 | 0.0835 | 0.0038 | 0.0004 | 0.0045 | 0.4982 |
| HMR_0488 | Glycerolipid metabolism | 0.2645 | 0.0141 | 0.2622 | 0.014 | 0.0022 | 0.0133 | 0.4947 |
| HMR_0489 | Glycerolipid metabolism | 0.0855 | 0.004 | 0.0855 | 0.004 | 1.00E-04 | 0.001 | 0.4996 |
| HMR_0490 | Glycerolipid metabolism | 0.0862 | 0.004 | 0.0857 | 0.004 | 0.0005 | 0.0056 | 0.4978 |
| HMR_0491 | Glycerolipid metabolism | 0.0849 | 0.0039 | 0.0846 | 0.0039 | 0.0003 | 0.0036 | 0.4986 |
| HMR_0492 | Glycerolipid metabolism | 0.0882 | 0.0041 | 0.0878 | 0.0041 | 0.0004 | 0.0047 | 0.4981 |
| HMR_0493 | Glycerolipid metabolism | 4.4721 | 1.6635 | 4.4019 | 1.5953 | 0.0702 | 0.0389 | 0.4845 |

| | | | | | | | | |
|---|---|---|---|---|---|---|---|---|
| HMR_0494 | Glycerolipid metabolism | 0.0938 | 0.0045 | 0.0931 | 0.0044 | 0.0007 | 0.0076 | 0.497 |
| HMR_0495 | Glycerolipid metabolism | 0.3572 | 0.0177 | 0.3522 | 0.0175 | 0.005 | 0.0268 | 0.4893 |
| HMR_0496 | Glycerolipid metabolism | 0.0795 | 0.0036 | 0.0794 | 0.0036 | 0.0002 | 0.0018 | 0.4993 |
| HMR_0497 | Glycerolipid metabolism | 0.0845 | 0.0039 | 0.0843 | 0.0039 | 0.0002 | 0.0027 | 0.4989 |
| HMR_0498 | Glycerolipid metabolism | 0.0821 | 0.0038 | 0.0819 | 0.0038 | 0.0002 | 0.0023 | 0.4991 |
| HMR_0499 | Glycerolipid metabolism | 4.4203 | 0.8068 | 4.4136 | 0.8041 | 0.0067 | 0.0053 | 0.4979 |
| HMR_0500 | Glycerolipid metabolism | 0.0877 | 0.0041 | 0.0873 | 0.0041 | 0.0003 | 0.0038 | 0.4985 |
| HMR_0501 | Glycerolipid metabolism | 0.5308 | 0.0218 | 0.5263 | 0.0214 | 0.0046 | 0.022 | 0.4912 |
| HMR_0502 | Glycerolipid metabolism | 2.5469 | 0.5028 | 2.5325 | 0.4975 | 0.0144 | 0.0144 | 0.4943 |
| HMR_0503 | Glycerolipid metabolism | 0.0827 | 0.0038 | 0.0827 | 0.0038 | 0.00E+00 | 0.0004 | 0.4998 |
| HMR_0504 | Glycerolipid metabolism | 0.0819 | 0.0038 | 0.082 | 0.0038 | 1.00E-04 | -0.0006 | 0.5002 |
| HMR_0505 | Glycerolipid metabolism | 0.0842 | 0.0039 | 0.0841 | 0.0039 | 1.00E-04 | 0.0009 | 0.4996 |
| HMR_0506 | Glycerolipid metabolism | 0.0792 | 0.0036 | 0.079 | 0.0036 | 0.0002 | 0.0019 | 0.4992 |
| HMR_0507 | Glycerolipid metabolism | 0.0835 | 0.0039 | 0.0836 | 0.0039 | 0.00E+00 | 0.00E+00 | 0.5 |
| HMR_0508 | Glycerolipid metabolism | 0.0815 | 0.0037 | 0.0816 | 0.0037 | 1.00E-04 | -0.0006 | 0.5003 |
| HMR_0509 | Glycerolipid metabolism | 0.0919 | 0.0044 | 0.0917 | 0.0044 | 0.0002 | 0.0022 | 0.4991 |
| HMR_0510 | Glycerolipid metabolism | 0.0828 | 0.0038 | 0.0828 | 0.0038 | 0.00E+00 | -1.00E-04 | 0.5 |
| HMR_0511 | Glycerolipid metabolism | 0.0839 | 0.0039 | 0.0838 | 0.0039 | 1.00E-04 | 0.001 | 0.4996 |
| HMR_0512 | Glycerolipid metabolism | 0.0838 | 0.0039 | 0.0837 | 0.0039 | 0.0001 | 0.0014 | 0.4994 |
| HMR_0513 | Glycerolipid metabolism | 0.0787 | 0.0036 | 0.0785 | 0.0036 | 0.0002 | 0.0019 | 0.4993 |
| HMR_0514 | Glycerolipid metabolism | 0.0814 | 0.0037 | 0.0814 | 0.0037 | 0.00E+00 | -0.0004 | 0.5001 |
| HMR_0515 | Glycerolipid metabolism | 0.0837 | 0.0039 | 0.0837 | 0.0039 | 1.00E-04 | 0.001 | 0.4996 |
| HMR_0516 | Glycerolipid metabolism | 0.0833 | 0.0038 | 0.0831 | 0.0038 | 0.0002 | 0.0021 | 0.4992 |
| HMR_0517 | Glycerolipid metabolism | 0.0827 | 0.0037 | 0.0825 | 0.0037 | 0.0002 | 0.0024 | 0.499 |
| HMR_0518 | Glycerolipid metabolism | 0.0816 | 0.0037 | 0.0816 | 0.0037 | 0.00E+00 | 1.00E-04 | 0.5 |
| HMR_0519 | Glycerolipid metabolism | 0.0827 | 0.0038 | 0.0826 | 0.0038 | 1.00E-04 | 0.0009 | 0.4996 |
| HMR_0520 | Glycerolipid metabolism | 0.0814 | 0.0037 | 0.0814 | 0.0037 | 0.00E+00 | 1.00E-04 | 0.5 |
| HMR_0521 | Glycerolipid metabolism | 0.0834 | 0.0038 | 0.0833 | 0.0038 | 0.0001 | 0.0014 | 0.4994 |
| HMR_0522 | Glycerolipid metabolism | 0.1345 | 0.0068 | 0.1338 | 0.0068 | 0.0007 | 0.006 | 0.4976 |
| HMR_0523 | Glycerolipid metabolism | 0.0946 | 0.0045 | 0.0946 | 0.0045 | 0.00E+00 | 1.00E-04 | 0.5 |
| HMR_0524 | Glycerolipid metabolism | 0.2351 | 0.011 | 0.2354 | 0.011 | 0.0003 | -0.002 | 0.5008 |
| HMR_0525 | Glycerolipid metabolism | 0.1979 | 0.0099 | 0.1973 | 0.0098 | 0.0006 | 0.0045 | 0.4982 |
| HMR_0526 | Glycerolipid metabolism | 0.1733 | 0.0091 | 0.1735 | 0.0091 | 0.0002 | -0.0012 | 0.5005 |
| HMR_0527 | Glycerolipid metabolism | 0.0809 | 0.0037 | 0.0809 | 0.0037 | 0.00E+00 | -0.0002 | 0.5001 |
| HMR_0528 | Glycerolipid metabolism | 0.081 | 0.0037 | 0.081 | 0.0037 | 0.00E+00 | -0.0001 | 0.5001 |
| HMR_0529 | Glycerolipid metabolism | 0.5462 | 0.0265 | 0.5456 | 0.0264 | 0.0006 | 0.0028 | 0.4989 |
| HMR_0530 | Glycerolipid metabolism | 0.0834 | 0.0038 | 0.0833 | 0.0038 | 1.00E-04 | 0.0009 | 0.4997 |
| HMR_0531 | Glycerolipid metabolism | 0.0841 | 0.0039 | 0.0839 | 0.0039 | 0.0002 | 0.0024 | 0.499 |
| HMR_0532 | Glycerolipid metabolism | 0.082 | 0.0038 | 0.082 | 0.0038 | 0.00E+00 | 0.0003 | 0.4999 |
| HMR_0533 | Glycerolipid metabolism | 0.0847 | 0.0039 | 0.0846 | 0.0039 | 0.0001 | 0.0012 | 0.4995 |
| HMR_0534 | Glycerolipid metabolism | 1.9952 | 0.2406 | 1.9874 | 0.2387 | 0.0078 | 0.0113 | 0.4955 |
| HMR_0535 | Glycerolipid metabolism | 0.0904 | 0.0043 | 0.0903 | 0.0043 | 0.00E+00 | 0.0004 | 0.4999 |
| HMR_0536 | Glycerolipid metabolism | 0.3166 | 0.0145 | 0.3168 | 0.0145 | 0.0003 | -0.0015 | 0.5006 |
| HMR_0537 | Glycerolipid metabolism | 2.3211 | 0.2753 | 2.3252 | 0.2748 | 0.0041 | -0.0055 | 0.5022 |

| | | | | | | | | |
|---|---|---|---|---|---|---|---|---|
| HMR_0538 | Glycerolipid metabolism | 0.0926 | 0.0044 | 0.0927 | 0.0044 | 0.00E+00 | -0.0005 | 0.5002 |
| HMR_0539 | Glycerolipid metabolism | 0.0832 | 0.0038 | 0.0831 | 0.0038 | 1.00E-04 | 0.001 | 0.4996 |
| HMR_0540 | Glycerolipid metabolism | 0.0831 | 0.0038 | 0.083 | 0.0038 | 1.00E-04 | 0.001 | 0.4996 |
| HMR_0541 | Glycerolipid metabolism | 0.1369 | 0.0069 | 0.1366 | 0.0069 | 0.0004 | 0.0033 | 0.4987 |
| HMR_0542 | Glycerolipid metabolism | 0.0858 | 0.004 | 0.0857 | 0.004 | 1.00E-04 | 0.0011 | 0.4996 |
| HMR_0543 | Glycerolipid metabolism | 0.0842 | 0.0039 | 0.084 | 0.0039 | 0.0002 | 0.0025 | 0.499 |
| HMR_0544 | Glycerolipid metabolism | 0.0837 | 0.0038 | 0.0835 | 0.0038 | 0.0002 | 0.0021 | 0.4992 |
| HMR_0588 | Glycerolipid metabolism | 3.8833 | 3.7576 | 3.8536 | 3.7411 | 0.0297 | 0.0108 | 0.4957 |
| HMR_0596 | Glycerolipid metabolism | 6.6464 | 9.6899 | 6.4962 | 9.1829 | 0.1502 | 0.0346 | 0.4862 |
| HMR_0673 | Glycerolipid metabolism | 1.2192 | 1.1713 | 1.2095 | 1.1547 | 0.0097 | 0.0063 | 0.4975 |
| HMR_0674 | Glycerolipid metabolism | 1.1982 | 1.1178 | 1.1844 | 1.0928 | 0.0137 | 0.0092 | 0.4963 |
| HMR_0675 | Glycerolipid metabolism | 1.1798 | 1.0567 | 1.1697 | 1.0399 | 0.0101 | 0.007 | 0.4972 |
| HMR_0676 | Glycerolipid metabolism | 1.2279 | 1.1359 | 1.2137 | 1.1104 | 0.0143 | 0.0095 | 0.4962 |
| HMR_0677 | Glycerolipid metabolism | 2.1345 | 2.888 | 2.1181 | 2.8493 | 0.0165 | 0.0069 | 0.4973 |
| HMR_0678 | Glycerolipid metabolism | 1.7577 | 2.4743 | 1.714 | 2.3535 | 0.0436 | 0.0199 | 0.4921 |
| HMR_4296 | Glycerolipid metabolism | 2.3487 | 6.6312 | 2.2797 | 6.1579 | 0.069 | 0.0193 | 0.4923 |
| HMR_4390 | Glycerolipid metabolism | 6.1777 | 15.5506 | 5.9391 | 14.4003 | 0.2386 | 0.0436 | 0.4826 |
| HMR_7599 | Glycerolipid metabolism | 0.1521 | 0.01 | 0.152 | 0.01 | 1.00E-04 | 0.0004 | 0.4998 |
| HMR_0450 | Glycerophospholipid metabolism | 2.9672 | 7.1975 | 3.0117 | 7.3518 | 0.0445 | -0.0117 | 0.5047 |
| HMR_0478 | Glycerophospholipid metabolism | 3.083 | 7.6652 | 3.1045 | 7.7257 | 0.0215 | -0.0055 | 0.5022 |
| HMR_0479 | Glycerophospholipid metabolism | 3.0951 | 7.7143 | 3.1167 | 7.7747 | 0.0215 | -0.0055 | 0.5022 |
| HMR_0481 | Glycerophospholipid metabolism | 3.3387 | 3.8266 | 3.3674 | 3.8868 | 0.0287 | -0.0103 | 0.5041 |
| HMR_0482 | Glycerophospholipid metabolism | 3.9328 | 11.1578 | 3.8131 | 10.6166 | 0.1197 | 0.0257 | 0.4898 |
| HMR_0589 | Glycerophospholipid metabolism | 4.7792 | 4.3758 | 4.7556 | 4.3185 | 0.0235 | 0.008 | 0.4968 |
| HMR_0590 | Glycerophospholipid metabolism | 3.3343 | 2.5564 | 3.3167 | 2.5233 | 0.0176 | 0.0078 | 0.4969 |
| HMR_0591 | Glycerophospholipid metabolism | 3.3628 | 2.3676 | 3.3445 | 2.3393 | 0.0183 | 0.0084 | 0.4966 |
| HMR_0592 | Glycerophospholipid metabolism | 7.4056 | 3.5761 | 7.4292 | 3.5812 | 0.0236 | -0.0088 | 0.5035 |
| HMR_0593 | Glycerophospholipid metabolism | 5.3785 | 5.0654 | 5.2274 | 4.8139 | 0.1511 | 0.0481 | 0.4808 |
| HMR_0597 | Glycerophospholipid metabolism | 6.227 | 5.0511 | 6.1799 | 4.9669 | 0.0471 | 0.0149 | 0.4941 |
| HMR_0598 | Glycerophospholipid metabolism | 4.7054 | 3.0387 | 4.6683 | 2.9937 | 0.0371 | 0.0151 | 0.494 |
| HMR_0599 | Glycerophospholipid metabolism | 4.969 | 2.8033 | 4.9412 | 2.7751 | 0.0278 | 0.0118 | 0.4953 |
| HMR_0600 | Glycerophospholipid metabolism | 1.6587 | 1.9836 | 1.6407 | 1.9472 | 0.018 | 0.0091 | 0.4964 |
| HMR_0601 | Glycerophospholipid metabolism | 5.3791 | 5.0657 | 5.2278 | 4.8141 | 0.1514 | 0.0482 | 0.4808 |
| HMR_0607 | Glycerophospholipid metabolism | -5.7553 | 2.5372 | -5.7967 | 2.566 | 0.0414 | 0.0183 | 0.4927 |
| HMR_0610 | Glycerophospholipid metabolism | 5.7763 | 2.5287 | 5.8174 | 2.5574 | 0.0411 | -0.0182 | 0.5073 |
| HMR_0614 | Glycerophospholipid metabolism | 2.1063 | 2.0979 | 2.0889 | 2.0611 | 0.0174 | 0.0085 | 0.4966 |
| HMR_0615 | Glycerophospholipid metabolism | 2.112 | 1.5895 | 2.1241 | 1.5989 | 0.0121 | -0.0068 | 0.5027 |
| HMR_0616 | Glycerophospholipid metabolism | 2.125 | 1.5912 | 2.1369 | 1.6006 | 0.0119 | -0.0066 | 0.5026 |
| HMR_0622 | Glycerophospholipid metabolism | 10.844 | 76.6858 | 9.5298 | 59.9699 | 1.3141 | 0.1124 | 0.4552 |
| HMR_0623 | Glycerophospholipid metabolism | 4.3935 | 7.3944 | 3.9528 | 6.2037 | 0.4407 | 0.1195 | 0.4524 |
| HMR_0625 | Glycerophospholipid metabolism | 3.723 | 3.8634 | 3.7046 | 3.7993 | 0.0184 | 0.0066 | 0.4974 |
| HMR_0627 | Glycerophospholipid metabolism | 15.7721 | 80.0808 | 14.0382 | 62.9603 | 1.7339 | 0.145 | 0.4424 |
| HMR_0629 | Glycerophospholipid metabolism | 1.4421 | 1.6397 | 1.4188 | 1.5884 | 0.0233 | 0.013 | 0.4948 |
| HMR_0630 | Glycerophospholipid metabolism | 2.1678 | 3.3005 | 2.194 | 3.3907 | 0.0262 | -0.0101 | 0.504 |

| | | | | | | | | |
|---|---|---|---|---|---|---|---|---|
| HMR_0632 | Glycerophospholipid metabolism | 3.5606 | 5.2552 | 3.6364 | 5.4498 | 0.0758 | -0.0232 | 0.5092 |
| HMR_0633 | Glycerophospholipid metabolism | 1.0824 | 0.5173 | 1.0825 | 0.5173 | 1.00E-04 | -1.00E-04 | 0.5 |
| HMR_0634 | Glycerophospholipid metabolism | 2.4734 | 3.6641 | 2.5232 | 3.8122 | 0.0498 | -0.0182 | 0.5073 |
| HMR_0635 | Glycerophospholipid metabolism | 1.0841 | 0.518 | 1.0843 | 0.518 | 0.0001 | -0.0001 | 0.5001 |
| HMR_0636 | Glycerophospholipid metabolism | 3.6957 | 7.2175 | 3.6081 | 6.7698 | 0.0876 | 0.0234 | 0.4907 |
| HMR_0638 | Glycerophospholipid metabolism | 5.0914 | 6.3486 | 5.0621 | 6.3053 | 0.0293 | 0.0082 | 0.4967 |
| HMR_0640 | Glycerophospholipid metabolism | 46.5711 | 612.9435 | 41.4583 | 478.2979 | 5.1128 | 0.1548 | 0.4385 |
| HMR_0641 | Glycerophospholipid metabolism | 8.5677 | 70.0576 | 8.3842 | 66.7722 | 0.1835 | 0.0157 | 0.4937 |
| HMR_0642 | Glycerophospholipid metabolism | 1.3654 | 1.336 | 1.3462 | 1.3022 | 0.0192 | 0.0118 | 0.4953 |
| HMR_0643 | Glycerophospholipid metabolism | 2.8007 | 2.4382 | 2.8407 | 2.5132 | 0.0401 | -0.018 | 0.5072 |
| HMR_0644 | Glycerophospholipid metabolism | 1.8577 | 2.2475 | 1.891 | 2.3214 | 0.0333 | -0.0156 | 0.5062 |
| HMR_0645 | Glycerophospholipid metabolism | 0.9401 | 0.392 | 0.9466 | 0.397 | 0.0065 | -0.0073 | 0.5029 |
| HMR_0646 | Glycerophospholipid metabolism | 0.9424 | 0.3927 | 0.9489 | 0.3977 | 0.0065 | -0.0073 | 0.5029 |
| HMR_0647 | Glycerophospholipid metabolism | 12.7253 | 135.358 | 13.388 | 149.7562 | 0.6626 | -0.0392 | 0.5157 |
| HMR_0648 | Glycerophospholipid metabolism | 12.3111 | 21.2659 | 11.9608 | 19.6114 | 0.3504 | 0.0548 | 0.4781 |
| HMR_0649 | Glycerophospholipid metabolism | 2.8232 | 9.6198 | 2.6974 | 8.647 | 0.1258 | 0.0294 | 0.4883 |
| HMR_0651 | Glycerophospholipid metabolism | 4.3573 | 3.1925 | 4.3518 | 3.158 | 0.0055 | 0.0022 | 0.4991 |
| HMR_0653 | Glycerophospholipid metabolism | 4.7088 | 6.4519 | 4.2835 | 5.3385 | 0.4253 | 0.1239 | 0.4507 |
| HMR_0654 | Glycerophospholipid metabolism | 4.7204 | 6.4503 | 4.295 | 5.3369 | 0.4255 | 0.1239 | 0.4507 |
| HMR_0657 | Glycerophospholipid metabolism | 4.7323 | 6.4534 | 4.3067 | 5.3399 | 0.4256 | 0.1239 | 0.4507 |
| HMR_0660 | Glycerophospholipid metabolism | 1.5996 | 1.5701 | 1.5904 | 1.5567 | 0.0092 | 0.0052 | 0.4979 |
| HMR_4627 | Glycerophospholipid metabolism | 2.807 | 2.6308 | 2.6568 | 2.3645 | 0.1502 | 0.0672 | 0.4732 |
| HMR_4838 | Glycerophospholipid metabolism | 2.8073 | 2.6269 | 2.6571 | 2.3606 | 0.1502 | 0.0672 | 0.4732 |
| HMR_4839 | Glycerophospholipid metabolism | 2.8074 | 2.6289 | 2.6572 | 2.3627 | 0.1501 | 0.0672 | 0.4732 |
| HMR_7591 | Glycerophospholipid metabolism | -0.088 | 1.2452 | -0.097 | 1.2337 | 0.009 | 0.0057 | 0.4977 |
| HMR_7594 | Glycerophospholipid metabolism | 1.0284 | 0.2183 | 1.0144 | 0.2126 | 0.014 | 0.0213 | 0.4915 |
| HMR_7597 | Glycerophospholipid metabolism | 1.0274 | 0.2176 | 1.0135 | 0.2119 | 0.0139 | 0.0212 | 0.4915 |
| HMR_7601 | Glycerophospholipid metabolism | 0.1265 | 0.0068 | 0.1265 | 0.0068 | 0.00E+00 | -0.0001 | 0.5001 |
| HMR_7611 | Glycerophospholipid metabolism | 8.3938 | 74.6908 | 8.2253 | 70.3391 | 0.1685 | 0.014 | 0.4944 |
| HMR_7613 | Glycerophospholipid metabolism | 364.2057 | 12495.0048 | 563.5075 | 12705.2491 | 199.3018 | -1.2555 | 0.8953 |
| HMR_7755 | Glycerophospholipid metabolism | 6.1775 | 15.5497 | 5.9389 | 14.3994 | 0.2386 | 0.0436 | 0.4826 |
| HMR_8421 | Glycerophospholipid metabolism | 38.003 | 571.4463 | 33.0739 | 437.3316 | 4.9292 | 0.1552 | 0.4383 |
| HMR_8424 | Glycerophospholipid metabolism | 3.3859 | 6.715 | 3.199 | 6.1212 | 0.187 | 0.0522 | 0.4792 |
| HMR_0816 | Glycosphingolipid biosynthesis-ganglio series | 3.9416 | 4.1105 | 4.005 | 4.2474 | 0.0634 | -0.0219 | 0.5088 |
| HMR_0817 | Glycosphingolipid biosynthesis-ganglio series | 0.7984 | 0.2871 | 0.8092 | 0.295 | 0.0109 | -0.0143 | 0.5057 |
| HMR_0819 | Glycosphingolipid biosynthesis-ganglio series | 2.0565 | 3.2891 | 2.0927 | 3.4024 | 0.0362 | -0.014 | 0.5056 |
| HMR_0820 | Glycosphingolipid biosynthesis-ganglio series | 1.0926 | 0.9772 | 1.1089 | 1.0068 | 0.0163 | -0.0116 | 0.5046 |
| HMR_0823 | Glycosphingolipid biosynthesis-ganglio series | 0.7989 | 0.2871 | 0.8098 | 0.2949 | 0.0108 | -0.0142 | 0.5057 |
| HMR_0824 | Glycosphingolipid biosynthesis-ganglio series | 2.0265 | 1.9185 | 2.0504 | 1.9644 | 0.024 | -0.0122 | 0.5049 |
| HMR_0825 | Glycosphingolipid biosynthesis-ganglio series | 2.0256 | 1.9185 | 2.0495 | 1.9644 | 0.0239 | -0.0122 | 0.5048 |
| HMR_0829 | Glycosphingolipid biosynthesis-ganglio series | 2.0566 | 3.2893 | 2.0927 | 3.4025 | 0.0362 | -0.014 | 0.5056 |
| HMR_0830 | Glycosphingolipid biosynthesis-ganglio series | 3.9418 | 4.1117 | 4.0052 | 4.2485 | 0.0634 | -0.0219 | 0.5087 |
| HMR_0837 | Glycosphingolipid biosynthesis-ganglio series | 1.8953 | 2.3121 | 1.9179 | 2.3687 | 0.0225 | -0.0104 | 0.5042 |
| HMR_0838 | Glycosphingolipid biosynthesis-ganglio series | 2.6896 | 2.4357 | 2.7229 | 2.496 | 0.0334 | -0.015 | 0.506 |

| | | | | | | | | |
|---|---|---|---|---|---|---|---|---|
| HMR_0844 | Glycosphingolipid biosynthesis-ganglio series | 2.0272 | 1.9192 | 2.0511 | 1.9651 | 0.024 | -0.0122 | 0.5049 |
| HMR_0845 | Glycosphingolipid biosynthesis-ganglio series | 2.0254 | 1.9192 | 2.0493 | 1.9651 | 0.0239 | -0.0121 | 0.5048 |
| HMR_0846 | Glycosphingolipid biosynthesis-ganglio series | 2.0272 | 1.9192 | 2.0511 | 1.9651 | 0.024 | -0.0122 | 0.5049 |
| HMR_0847 | Glycosphingolipid biosynthesis-ganglio series | 2.0254 | 1.9192 | 2.0493 | 1.9651 | 0.0239 | -0.0121 | 0.5048 |
| HMR_0849 | Glycosphingolipid biosynthesis-ganglio series | 1.9388 | 2.9706 | 1.9622 | 3.0451 | 0.0234 | -0.0096 | 0.5038 |
| HMR_0850 | Glycosphingolipid biosynthesis-ganglio series | 3.8221 | 3.8178 | 3.8728 | 3.9194 | 0.0506 | -0.0182 | 0.5073 |
| HMR_0794 | Glycosphingolipid metabolism | 5.7325 | 3.6702 | 5.8238 | 3.7145 | 0.0912 | -0.0336 | 0.5134 |
| HMR_1312 | Prostaglandin biosynthesis | 412.6848 | 57406.7293 | 413.0757 | 57453.2657 | 0.391 | -0.0012 | 0.5005 |
| HMR_1321 | Prostaglandin biosynthesis | -0.001 | 199915.4842 | -0.0011 | 199915.5417 | 0.0001 | 0.00E+00 | 0.5 |
| HMR_1322 | Prostaglandin biosynthesis | 0.0003 | 199915.4792 | 0.0004 | 199915.5367 | 0.00E+00 | 0.00E+00 | 0.5 |
| HMR_1323 | Prostaglandin biosynthesis | 0.0003 | 199915.4792 | 0.0004 | 199915.5367 | 0.00E+00 | 0.00E+00 | 0.5 |
| HMR_1324 | Prostaglandin biosynthesis | -0.001 | 199915.4842 | -0.0011 | 199915.5417 | 0.0001 | 0.00E+00 | 0.5 |
| HMR_1325 | Prostaglandin biosynthesis | 60.1156 | 232074.3635 | 56.4775 | 232826.8788 | 3.6381 | 0.0053 | 0.4979 |
| HMR_1326 | Prostaglandin biosynthesis | -465.6435 | 173815.5295 | -464.137 | 174504.1362 | 1.5065 | -0.0026 | 0.501 |
| HMR_1327 | Prostaglandin biosynthesis | 412.6855 | 57406.7296 | 413.0764 | 57453.266 | 0.3909 | -0.0012 | 0.5005 |
| HMR_1329 | Prostaglandin biosynthesis | -7.1573 | 153545.7402 | -5.4165 | 153908.684 | 1.7408 | -0.0031 | 0.5013 |
| HMR_1330 | Prostaglandin biosynthesis | 386.3684 | 71717.1681 | 386.7446 | 71770.2597 | 0.3762 | -0.001 | 0.5004 |
| HMR_1373 | Prostaglandin biosynthesis | 0.00E+00 | 220512.0877 | 0.00E+00 | 220512.1485 | 0.00E+00 | 0.00E+00 | 0.5 |
| HMR_1375 | Prostaglandin biosynthesis | -1.00E-04 | 220512.0877 | -1.00E-04 | 220512.1485 | 0.00E+00 | 0.00E+00 | 0.5 |
| HMR_1376 | Prostaglandin biosynthesis | -1.00E-04 | 220512.0877 | -1.00E-04 | 220512.1485 | 0.00E+00 | 0.00E+00 | 0.5 |
| HMR_0703 | Ether lipid metabolism | 1.0211 | 0.2244 | 1.007 | 0.2187 | 0.0141 | 0.0211 | 0.4916 |
| HMR_7598 | Ether lipid metabolism | 1.0245 | 0.2202 | 1.0107 | 0.2144 | 0.0138 | 0.021 | 0.4916 |
| HMR_7600 | Ether lipid metabolism | 0.8713 | 0.2293 | 0.8577 | 0.2234 | 0.0136 | 0.0203 | 0.4919 |
| HMR_7602 | Ether lipid metabolism | 1.9589 | 1.8244 | 1.9946 | 1.8907 | 0.0357 | -0.0185 | 0.5074 |
| HMR_7603 | Ether lipid metabolism | 2.0196 | 1.8247 | 2.0552 | 1.891 | 0.0356 | -0.0185 | 0.5074 |
| HMR_7614 | Ether lipid metabolism | 1.8453 | 2.1997 | 1.8789 | 2.2638 | 0.0336 | -0.0159 | 0.5063 |
| HMR_7615 | Ether lipid metabolism | 2.697 | 2.3154 | 2.717 | 2.3737 | 0.0201 | -0.0093 | 0.5037 |
| HMR_0025 | Regulation of lipid metabolism | 500 | 63776.8521 | 500 | 63776.8521 | 0.00E+00 | 0.00E+00 | 0.5 |
| HMR_0030 | Regulation of lipid metabolism | 500 | 63776.8521 | 500 | 63776.8521 | 0.00E+00 | 0.00E+00 | 0.5 |
| HMR_3799 | Bile acid biosynthesis | 2.1713 | 4.8953 | 2.2269 | 5.1005 | 0.0556 | -0.0176 | 0.507 |
| HMR_3908 | Bile acid biosynthesis | 38.3312 | 299.7639 | 46.766 | 420.5475 | 8.4348 | -0.3143 | 0.6233 |
| HMR_3910 | Bile acid biosynthesis | 0.0025 | 0.00E+00 | 0.0025 | 0.00E+00 | 0.00E+00 | 0.00E+00 | 0.5 |
| HMR_4679 | Bile acid biosynthesis | -0.0025 | 0.00E+00 | -0.0025 | 0.00E+00 | 0.00E+00 | 0.00E+00 | 0.5 |
| HMR_1896 | Bile acid recycling | 2.8469 | 3.7899 | 2.7179 | 3.4543 | 0.129 | 0.0479 | 0.4809 |
| HMR_0712 | Nicotinate and nicotinamide metabolism | 3.563 | 6.802 | 3.5272 | 6.6626 | 0.0358 | 0.0098 | 0.4961 |
| HMR_4228 | Nicotinate and nicotinamide metabolism | 2.7674 | 1.2053 | 2.767 | 1.2029 | 0.0004 | 0.0003 | 0.4999 |
| HMR_4251 | Nicotinate and nicotinamide metabolism | -1.59 | 0.9913 | -1.6078 | 1.0047 | 0.0178 | 0.0126 | 0.495 |
| HMR_4252 | Nicotinate and nicotinamide metabolism | 2.8465 | 3.7933 | 2.7174 | 3.4577 | 0.1291 | 0.0479 | 0.4809 |
| HMR_4253 | Nicotinate and nicotinamide metabolism | 2.8501 | 3.7934 | 2.7212 | 3.4578 | 0.1289 | 0.0479 | 0.4809 |
| HMR_4254 | Nicotinate and nicotinamide metabolism | 958.6225 | 356.2125 | 957.3413 | 376.0925 | 1.2812 | 0.0473 | 0.4811 |
| HMR_4255 | Nicotinate and nicotinamide metabolism | 2.0099 | 4.154 | 2.0344 | 4.2306 | 0.0245 | -0.0085 | 0.5034 |
| HMR_4257 | Nicotinate and nicotinamide metabolism | -243.5083 | 120421.7284 | -242.9313 | 120583.4513 | 0.577 | -0.0012 | 0.5005 |
| HMR_4259 | Nicotinate and nicotinamide metabolism | -956.5513 | 357.6418 | -955.2804 | 377.4571 | 1.2709 | -0.0469 | 0.5187 |
| HMR_4260 | Nicotinate and nicotinamide metabolism | 0.9231 | 0.7813 | 0.914 | 0.7672 | 0.0092 | 0.0074 | 0.4971 |

| | | | | | | | | |
|---|---|---|---|---|---|---|---|---|
| HMR_4261 | Nicotinate and nicotinamide metabolism | 1.3296 | 1.5004 | 1.3379 | 1.5179 | 0.0083 | -0.0048 | 0.5019 |
| HMR_4262 | Nicotinate and nicotinamide metabolism | 955.21 | 358.2901 | 953.9786 | 378.0735 | 1.2314 | 0.0454 | 0.4819 |
| HMR_4263 | Nicotinate and nicotinamide metabolism | 2.3539 | 3.5963 | 2.3859 | 3.661 | 0.0319 | -0.0118 | 0.5047 |
| HMR_4269 | Nicotinate and nicotinamide metabolism | 2.8463 | 3.8273 | 2.718 | 3.4916 | 0.1283 | 0.0474 | 0.4811 |
| HMR_4270 | Nicotinate and nicotinamide metabolism | 2.869 | 3.8279 | 2.7402 | 3.4923 | 0.1288 | 0.0476 | 0.481 |
| HMR_4271 | Nicotinate and nicotinamide metabolism | 50.7003 | 3377.6735 | 58.7489 | 4603.9485 | 8.0486 | -0.0901 | 0.5359 |
| HMR_4276 | Nicotinate and nicotinamide metabolism | 249.1878 | 120417.1241 | 248.6437 | 120578.7486 | 0.5441 | 0.0011 | 0.4996 |
| HMR_4278 | Nicotinate and nicotinamide metabolism | 1.4301 | 0.7099 | 1.4394 | 0.7191 | 0.0093 | -0.0078 | 0.5031 |
| HMR_7143 | Nicotinate and nicotinamide metabolism | 1.5202 | 1.8503 | 1.5062 | 1.8214 | 0.014 | 0.0073 | 0.4971 |
| HMR_7623 | Nicotinate and nicotinamide metabolism | 1.6173 | 1.1554 | 1.6012 | 1.1337 | 0.016 | 0.0106 | 0.4958 |
| HMR_7625 | Nicotinate and nicotinamide metabolism | 1.949 | 1.2832 | 1.9477 | 1.2814 | 0.0013 | 0.0008 | 0.4997 |
| HMR_7676 | Nicotinate and nicotinamide metabolism | 960.2963 | 355.7223 | 959.0332 | 375.5877 | 1.2631 | 0.0467 | 0.4814 |
| HMR_8790 | Nicotinate and nicotinamide metabolism | 3.7006 | 3.863 | 3.6933 | 3.8558 | 0.0073 | 0.0026 | 0.499 |
| HMR_3871 | Pantothenate and CoA biosynthesis | 4.0978 | 3.323 | 3.9709 | 3.0864 | 0.1268 | 0.0501 | 0.48 |
| HMR_4058 | Pantothenate and CoA biosynthesis | 0.0025 | 0.00E+00 | 0.0025 | 0.00E+00 | 0.00E+00 | 0.00E+00 | 0.5 |
| HMR_4498 | Pantothenate and CoA biosynthesis | 1.2816 | 0.7075 | 1.281 | 0.6926 | 0.0006 | 0.0005 | 0.4998 |
| HMR_4500 | Pantothenate and CoA biosynthesis | 1.2373 | 0.7078 | 1.2102 | 0.6925 | 0.027 | 0.0229 | 0.4909 |
| HMR_4714 | Pantothenate and CoA biosynthesis | 0.0026 | 0.00E+00 | 0.0026 | 0.00E+00 | 0.00E+00 | 0.00E+00 | 0.5 |
| HMR_4715 | Pantothenate and CoA biosynthesis | 748.653 | 12874.1377 | 762.7367 | 11521.6999 | 14.0837 | -0.0902 | 0.5359 |
| HMR_4716 | Pantothenate and CoA biosynthesis | 748.6519 | 12874.134 | 762.7355 | 11521.6952 | 14.0836 | -0.0902 | 0.5359 |
| HMR_4717 | Pantothenate and CoA biosynthesis | -0.0025 | 0.00E+00 | -0.0025 | 0.00E+00 | 0.00E+00 | 0.00E+00 | 0.5 |
| HMR_4718 | Pantothenate and CoA biosynthesis | 0.0026 | 0.00E+00 | 0.0026 | 0.00E+00 | 0.00E+00 | 0.00E+00 | 0.5 |
| HMR_4723 | Pantothenate and CoA biosynthesis | 0.0026 | 0.00E+00 | 0.0026 | 0.00E+00 | 0.00E+00 | 0.00E+00 | 0.5 |
| HMR_4725 | Pantothenate and CoA biosynthesis | 0.0026 | 0.00E+00 | 0.0026 | 0.00E+00 | 0.00E+00 | 0.00E+00 | 0.5 |
| HMR_4727 | Pantothenate and CoA biosynthesis | 3.1838 | 3.0299 | 3.2037 | 3.0701 | 0.0199 | -0.008 | 0.5032 |
| HMR_4730 | Pantothenate and CoA biosynthesis | -1.00E-04 | 0.00E+00 | -1.00E-04 | 0.00E+00 | 0.00E+00 | 0.00E+00 | 0.5 |
| HMR_4731 | Pantothenate and CoA biosynthesis | 748.652 | 12874.1389 | 762.7358 | 11521.7005 | 14.0837 | -0.0902 | 0.5359 |
| HMR_4732 | Pantothenate and CoA biosynthesis | 1.9052 | 2.6155 | 1.9259 | 2.6664 | 0.0207 | -0.009 | 0.5036 |
| HMR_4733 | Pantothenate and CoA biosynthesis | -748.6516 | 12874.1386 | -762.7353 | 11521.701 | 14.0837 | 0.0902 | 0.4641 |
| HMR_4734 | Pantothenate and CoA biosynthesis | 0.0025 | 0.00E+00 | 0.0025 | 0.00E+00 | 0.00E+00 | 0.00E+00 | 0.5 |
| HMR_7731 | Pantothenate and CoA biosynthesis | 2.8273 | 2.7118 | 2.7021 | 2.4826 | 0.1252 | 0.0549 | 0.4781 |
| HMR_0663 | Inositol phosphate metabolism | 2.9351 | 2.589 | 2.9194 | 2.5984 | 0.0157 | 0.0069 | 0.4972 |
| HMR_4308 | Inositol phosphate metabolism | 3.5259 | 1.9041 | 3.556 | 1.9344 | 0.0302 | -0.0154 | 0.5061 |
| HMR_6540 | Inositol phosphate metabolism | 1.0102 | 0.8607 | 1.0125 | 0.865 | 0.0023 | -0.0017 | 0.5007 |
| HMR_6542 | Inositol phosphate metabolism | 4.8985 | 10.823 | 4.6848 | 9.8836 | 0.2137 | 0.047 | 0.4813 |
| HMR_6543 | Inositol phosphate metabolism | 3.1347 | 7.0409 | 2.9905 | 6.4147 | 0.1441 | 0.0393 | 0.4843 |
| HMR_6544 | Inositol phosphate metabolism | 2.8004 | 6.2054 | 2.6733 | 5.654 | 0.127 | 0.0369 | 0.4853 |
| HMR_6545 | Inositol phosphate metabolism | 5.6989 | 9.7323 | 5.4377 | 8.8635 | 0.2612 | 0.0606 | 0.4758 |
| HMR_6546 | Inositol phosphate metabolism | 3.6177 | 8.6516 | 3.4682 | 7.9256 | 0.1495 | 0.0367 | 0.4853 |
| HMR_6547 | Inositol phosphate metabolism | 4.3018 | 10.4738 | 4.0878 | 9.5041 | 0.214 | 0.0479 | 0.4809 |
| HMR_6548 | Inositol phosphate metabolism | 3.2877 | 7.6799 | 3.1378 | 6.9983 | 0.1499 | 0.0391 | 0.4844 |
| HMR_6550 | Inositol phosphate metabolism | 4.8341 | 11.5273 | 4.6181 | 10.5153 | 0.216 | 0.046 | 0.4816 |
| HMR_6551 | Inositol phosphate metabolism | 4.1482 | 10.6018 | 3.9622 | 9.6689 | 0.186 | 0.0413 | 0.4835 |
| HMR_6552 | Inositol phosphate metabolism | 3.918 | 9.916 | 3.7388 | 9.0333 | 0.1792 | 0.0412 | 0.4836 |

| | | | | | | | | |
|---|---|---|---|---|---|---|---|---|
| HMR_6553 | Inositol phosphate metabolism | 4.4048 | 11.4306 | 4.1939 | 10.3861 | 0.2109 | 0.0452 | 0.482 |
| HMR_6554 | Inositol phosphate metabolism | 3.7401 | 9.4334 | 3.5783 | 8.6201 | 0.1617 | 0.0381 | 0.4848 |
| HMR_6555 | Inositol phosphate metabolism | 3.5748 | 8.8996 | 3.4101 | 8.1037 | 0.1647 | 0.0399 | 0.4841 |
| HMR_6556 | Inositol phosphate metabolism | 4.6427 | 12.0523 | 4.437 | 10.9975 | 0.2058 | 0.0429 | 0.4829 |
| HMR_6557 | Inositol phosphate metabolism | 3.5437 | 8.2631 | 3.3833 | 7.5326 | 0.1605 | 0.0404 | 0.4839 |
| HMR_6558 | Inositol phosphate metabolism | 3.7301 | 8.7961 | 3.5577 | 8.0083 | 0.1724 | 0.0421 | 0.4832 |
| HMR_6559 | Inositol phosphate metabolism | 0.6385 | 0.1746 | 0.6397 | 0.1752 | 0.0012 | -0.002 | 0.5008 |
| HMR_6560 | Inositol phosphate metabolism | 0.6339 | 0.1736 | 0.6348 | 0.1742 | 0.0009 | -0.0016 | 0.5006 |
| HMR_6561 | Inositol phosphate metabolism | 1.2633 | 0.3749 | 1.2688 | 0.3785 | 0.0056 | -0.0064 | 0.5026 |
| HMR_6562 | Inositol phosphate metabolism | 1.2561 | 0.3816 | 1.2617 | 0.3851 | 0.0056 | -0.0064 | 0.5026 |
| HMR_6565 | Inositol phosphate metabolism | 2.8471 | 3.7899 | 2.718 | 3.4543 | 0.129 | 0.0479 | 0.4809 |
| HMR_6566 | Inositol phosphate metabolism | 2.8476 | 3.7899 | 2.7186 | 3.4543 | 0.129 | 0.0479 | 0.4809 |
| HMR_6571 | Inositol phosphate metabolism | 1.069 | 0.5132 | 1.0711 | 0.5153 | 0.0021 | -0.0021 | 0.5008 |
| HMR_6572 | Inositol phosphate metabolism | 1.073 | 0.5138 | 1.0751 | 0.516 | 0.0021 | -0.0021 | 0.5008 |
| HMR_6576 | Inositol phosphate metabolism | 1.2643 | 0.5619 | 1.2481 | 0.5474 | 0.0162 | 0.0154 | 0.4939 |
| HMR_6579 | Inositol phosphate metabolism | 4.1025 | 10.4361 | 3.9273 | 9.5422 | 0.1752 | 0.0392 | 0.4844 |
| HMR_6581 | Inositol phosphate metabolism | 2.922 | 3.7776 | 2.7908 | 3.4462 | 0.1312 | 0.0488 | 0.4805 |
| HMR_6583 | Inositol phosphate metabolism | 3.9475 | 9.9767 | 3.7588 | 9.0653 | 0.1887 | 0.0432 | 0.4828 |
| HMR_6585 | Inositol phosphate metabolism | 2.9214 | 3.778 | 2.7901 | 3.4465 | 0.1313 | 0.0488 | 0.4805 |
| HMR_6587 | Inositol phosphate metabolism | 2.8939 | 5.1021 | 2.76 | 4.6449 | 0.1338 | 0.0429 | 0.4829 |
| HMR_6591 | Inositol phosphate metabolism | 5.2507 | 12.729 | 5.0095 | 11.5955 | 0.2412 | 0.0489 | 0.4805 |
| HMR_6592 | Inositol phosphate metabolism | 4.2797 | 10.1087 | 4.0862 | 9.2162 | 0.1935 | 0.044 | 0.4824 |
| HMR_6595 | Inositol phosphate metabolism | 4.0585 | 9.639 | 3.8784 | 8.7942 | 0.1801 | 0.0419 | 0.4833 |
| HMR_7652 | Inositol phosphate metabolism | 3.9202 | 4.9455 | 3.7851 | 4.5691 | 0.1351 | 0.0438 | 0.4825 |
| HMR_7655 | Inositol phosphate metabolism | 2.6558 | 4.6802 | 2.5368 | 4.3046 | 0.1189 | 0.0397 | 0.4842 |
| HMR_7656 | Inositol phosphate metabolism | 1.2643 | 0.5619 | 1.2481 | 0.5474 | 0.0162 | 0.0154 | 0.4939 |
| HMR_8813 | Inositol phosphate metabolism | 2.278 | 3.6963 | 2.1792 | 3.3799 | 0.0988 | 0.0372 | 0.4852 |
| HMR_8815 | Inositol phosphate metabolism | 2.3916 | 4.0539 | 2.2864 | 3.7033 | 0.1052 | 0.0378 | 0.4849 |
| HMR_8822 | Inositol phosphate metabolism | 0.6383 | 0.2843 | 0.6432 | 0.288 | 0.0049 | -0.0064 | 0.5026 |
| HMR_8824 | Inositol phosphate metabolism | 3.183 | 7.5176 | 3.0397 | 6.8539 | 0.1433 | 0.0378 | 0.4849 |
| HMR_8832 | Inositol phosphate metabolism | 1.6005 | 1.5405 | 1.6487 | 1.6075 | 0.0482 | -0.0272 | 0.5108 |
| HMR_8835 | Inositol phosphate metabolism | 1.2623 | 0.5611 | 1.2461 | 0.5466 | 0.0162 | 0.0154 | 0.4939 |
| HMR_3923 | Folate metabolism | 774.7948 | 23750.1754 | 808.6485 | 17442.137 | 33.8536 | -0.1668 | 0.5662 |
| HMR_3925 | Folate metabolism | 397.6947 | 158683.16 | 394.2775 | 160253.5505 | 3.4171 | 0.0061 | 0.4976 |
| HMR_3929 | Folate metabolism | -674.8826 | 2979.9277 | -825.1713 | 2964.8195 | 150.2888 | 1.9492 | 0.0256 |
| HMR_3972 | Folate metabolism | 15.8216 | 233.8344 | 17.621 | 287.3806 | 1.7994 | -0.0788 | 0.5314 |
| HMR_4332 | Folate metabolism | 143.4194 | 180625.6613 | 179.8669 | 178472.0306 | 36.4476 | -0.0608 | 0.5242 |
| HMR_4333 | Folate metabolism | -465.821 | 166559.8042 | -455.871 | 167805.0373 | 9.95 | -0.0172 | 0.5069 |
| HMR_4335 | Folate metabolism | 322.3999 | 23217.1299 | 276.0014 | 18973.7184 | 46.3985 | 0.2259 | 0.4106 |
| HMR_4336 | Folate metabolism | 6.9602 | 53.978 | 6.1324 | 40.2977 | 0.8278 | 0.0853 | 0.466 |
| HMR_4338 | Folate metabolism | 3.583 | 10.7986 | 3.572 | 10.6618 | 0.0111 | 0.0024 | 0.499 |
| HMR_4340 | Folate metabolism | 4.7862 | 7.8121 | 5.0358 | 8.5867 | 0.2496 | -0.0616 | 0.5246 |
| HMR_4440 | Folate metabolism | 132.2267 | 226741.7111 | 92.6522 | 232351.7938 | 39.5744 | 0.0584 | 0.4767 |
| HMR_4442 | Folate metabolism | 670.474 | 3075.1338 | 821.6068 | 3031.2708 | 151.1328 | -1.934 | 0.9734 |

| | | | | | | | | |
|---|---|---|---|---|---|---|---|---|
| HMR_4444 | Folate metabolism | -127.4427 | 226740.3631 | -87.6183 | 232349.4339 | 39.8244 | -0.0588 | 0.5234 |
| HMR_4446 | Folate metabolism | -392.4549 | 158628.5655 | -389.7283 | 160223.7824 | 2.7266 | -0.0048 | 0.5019 |
| HMR_4448 | Folate metabolism | 378.2211 | 158626.5516 | 373.6774 | 160229.5112 | 4.5437 | 0.008 | 0.4968 |
| HMR_4503 | Folate metabolism | -4.4075 | 64.8765 | -3.5635 | 51.1161 | 0.844 | -0.0784 | 0.5312 |
| HMR_4505 | Folate metabolism | 4.7865 | 7.8157 | 5.0361 | 8.5903 | 0.2496 | -0.0616 | 0.5246 |
| HMR_4654 | Folate metabolism | 143.4198 | 180625.6551 | 179.8674 | 178472.0189 | 36.4477 | -0.0608 | 0.5242 |
| HMR_4655 | Folate metabolism | -465.821 | 166559.8026 | -455.871 | 167805.0321 | 9.95 | -0.0172 | 0.5069 |
| HMR_4656 | Folate metabolism | 322.4003 | 23217.1274 | 276.0019 | 18973.7164 | 46.3984 | 0.2259 | 0.4106 |
| HMR_4665 | Folate metabolism | 4.3594 | 8.875 | 4.4691 | 9.3293 | 0.1097 | -0.0257 | 0.5103 |
| HMR_4666 | Folate metabolism | 4.358 | 8.875 | 4.4677 | 9.3293 | 0.1097 | -0.0257 | 0.5103 |
| HMR_8758 | Folate metabolism | 8.6721 | 29.234 | 8.4046 | 27.9703 | 0.2674 | 0.0354 | 0.4859 |
| HMR_4160 | Biopterin metabolism | 0.7611 | 0.4093 | 0.7381 | 0.3851 | 0.023 | 0.0258 | 0.4897 |
| HMR_4165 | Biopterin metabolism | -500.0001 | 55128.0359 | -500.0001 | 55128.0376 | 1.00E-04 | 0.00E+00 | 0.5 |
| HMR_4166 | Biopterin metabolism | 500 | 55128.034 | 500 | 55128.0357 | 0.00E+00 | 0.00E+00 | 0.5 |
| HMR_4167 | Biopterin metabolism | 499.9999 | 55128.0359 | 499.9999 | 55128.0376 | 0.00E+00 | 0.00E+00 | 0.5 |
| HMR_4169 | Biopterin metabolism | 0.8946 | 0.2347 | 0.8694 | 0.2219 | 0.0252 | 0.0373 | 0.4851 |
| HMR_4523 | Biopterin metabolism | -372.124 | 158693.4446 | -368.4157 | 160265.678 | 3.7083 | -0.0066 | 0.5026 |
| HMR_4539 | Biopterin metabolism | 0.00E+00 | 220512.1554 | 0.00E+00 | 220512.174 | 0.00E+00 | 0.00E+00 | 0.5 |
| HMR_4540 | Biopterin metabolism | 1.00E-04 | 220512.1434 | 1.00E-04 | 220512.1395 | 0.00E+00 | 0.00E+00 | 0.5 |
| HMR_4541 | Biopterin metabolism | 0.00E+00 | 220512.1535 | 0.00E+00 | 220512.172 | 0.00E+00 | 0.00E+00 | 0.5 |
| HMR_4542 | Biopterin metabolism | 0.00E+00 | 220512.1434 | 0.00E+00 | 220512.1395 | 0.00E+00 | 0.00E+00 | 0.5 |
| HMR_4543 | Biopterin metabolism | 0.00E+00 | 220512.1554 | 0.00E+00 | 220512.174 | 0.00E+00 | 0.00E+00 | 0.5 |
| HMR_4544 | Biopterin metabolism | 1.00E-04 | 220512.1434 | 1.00E-04 | 220512.1395 | 0.00E+00 | 0.00E+00 | 0.5 |
| HMR_4816 | Biopterin metabolism | 1.6444 | 0.5348 | 1.5955 | 0.5041 | 0.049 | 0.048 | 0.4808 |
| HMR_4817 | Biopterin metabolism | 0.89 | 0.232 | 0.8643 | 0.2192 | 0.0257 | 0.0382 | 0.4848 |
| HMR_4833 | Biopterin metabolism | -0.8953 | 0.2295 | -0.87 | 0.2167 | 0.0253 | -0.0379 | 0.5151 |
| HMR_4835 | Biopterin metabolism | -0.8939 | 0.2286 | -0.8684 | 0.2158 | 0.0255 | -0.0382 | 0.5152 |
| HMR_8538 | Biopterin metabolism | 14.0862 | 50.9888 | 14.2133 | 52.3314 | 0.1271 | -0.0125 | 0.505 |
| HMR_6393 | Ascorbate and aldarate metabolism | -10.4485 | 28.5459 | -12.8686 | 42.6146 | 2.4201 | 0.2869 | 0.3871 |
| HMR_6396 | Ascorbate and aldarate metabolism | 10.4486 | 28.5473 | 12.8686 | 42.6161 | 2.42 | -0.2869 | 0.6129 |
| HMR_3991 | Porphyrin metabolism | 8.0283 | 22.461 | 8.8545 | 27.1095 | 0.8262 | -0.1174 | 0.5467 |
| HMR_4657 | Porphyrin metabolism | 3.746 | 220493.1859 | 3.3913 | 220496.5791 | 0.3547 | 0.0005 | 0.4998 |
| HMR_4744 | Porphyrin metabolism | 5.6188 | 4.9712 | 5.0869 | 4.1619 | 0.5318 | 0.176 | 0.4302 |
| HMR_4746 | Porphyrin metabolism | 1.405 | 0.3102 | 1.2721 | 0.2596 | 0.1329 | 0.176 | 0.4301 |
| HMR_4748 | Porphyrin metabolism | 1.4099 | 0.3033 | 1.2782 | 0.2529 | 0.1317 | 0.1766 | 0.4299 |
| HMR_4750 | Porphyrin metabolism | 1.4142 | 0.2972 | 1.2836 | 0.247 | 0.1306 | 0.177 | 0.4297 |
| HMR_4752 | Porphyrin metabolism | 1.4208 | 0.2919 | 1.2915 | 0.2418 | 0.1292 | 0.1769 | 0.4298 |
| HMR_4755 | Porphyrin metabolism | 0.7184 | 0.2026 | 0.6552 | 0.1682 | 0.0632 | 0.1039 | 0.4586 |
| HMR_4757 | Porphyrin metabolism | 0.7143 | 0.2022 | 0.6503 | 0.1678 | 0.064 | 0.1051 | 0.4581 |
| HMR_4758 | Porphyrin metabolism | 1.4373 | 0.2947 | 1.311 | 0.2448 | 0.1263 | 0.172 | 0.4317 |
| HMR_6638 | Retinol metabolism | 385.6753 | 158501.3966 | 382.12 | 160072.8035 | 3.5553 | 0.0063 | 0.4975 |
| HMR_6646 | Retinol metabolism | 499.9995 | 49978.8813 | 499.9995 | 49978.8827 | 0.00E+00 | 0.00E+00 | 0.5 |
| HMR_6647 | Retinol metabolism | 298.6448 | 46589.1573 | 298.6448 | 46589.1627 | 0.00E+00 | 0.00E+00 | 0.5 |
| HMR_6650 | Retinol metabolism | 356.5483 | 37273.3461 | 356.5483 | 37273.3487 | 0.00E+00 | 0.00E+00 | 0.5 |

| | | | | | | | | |
|---|---|---|---|---|---|---|---|---|
| HMR_6652 | Retinol metabolism | 356.5481 | 37273.3458 | 356.5482 | 37273.3484 | 0.00E+00 | 0.00E+00 | 0.5 |
| HMR_6660 | Retinol metabolism | 356.5483 | 37273.3473 | 356.5483 | 37273.3499 | 0.00E+00 | 0.00E+00 | 0.5 |
| HMR_6661 | Retinol metabolism | -500.0003 | 49978.8813 | -500.0003 | 49978.8827 | 0.00E+00 | 0.00E+00 | 0.5 |
| HMR_6662 | Retinol metabolism | -278.4904 | 162696.014 | -278.4904 | 162696.0487 | 0.00E+00 | 0.00E+00 | 0.5 |
| HMR_6672 | Retinol metabolism | 499.9999 | 49978.8803 | 499.9999 | 49978.8817 | 0.00E+00 | 0.00E+00 | 0.5 |
| HMR_6673 | Retinol metabolism | 655.1932 | 49546.2769 | 655.1933 | 49546.2746 | 0.00E+00 | 0.00E+00 | 0.5 |
| HMR_6674 | Retinol metabolism | -376.7029 | 168196.6065 | -376.7029 | 168196.6352 | 0.00E+00 | 0.00E+00 | 0.5 |
| HMR_6675 | Retinol metabolism | -500.0002 | 49978.8803 | -500.0003 | 49978.8817 | 0.00E+00 | 0.00E+00 | 0.5 |
| HMR_6676 | Retinol metabolism | -278.4906 | 162696.0131 | -278.4905 | 162696.0478 | 0.00E+00 | 0.00E+00 | 0.5 |
| HMR_8709 | Retinol metabolism | -385.6753 | 158501.3965 | -382.12 | 160072.8033 | 3.5553 | -0.0063 | 0.5025 |
| HMR_6506 | Riboflavin metabolism | 2.8479 | 3.7916 | 2.7189 | 3.456 | 0.129 | 0.0479 | 0.4809 |
| HMR_6507 | Riboflavin metabolism | 2.8477 | 3.7916 | 2.7187 | 3.456 | 0.129 | 0.0479 | 0.4809 |
| HMR_6508 | Riboflavin metabolism | 2.0139 | 1.8973 | 2.0377 | 1.9426 | 0.0238 | -0.0121 | 0.5048 |
| HMR_6509 | Riboflavin metabolism | 2.0146 | 1.8974 | 2.0384 | 1.9426 | 0.0238 | -0.0121 | 0.5048 |
| HMR_6510 | Riboflavin metabolism | 12.4112 | 195.5239 | 14.5658 | 273.3731 | 2.1546 | -0.0995 | 0.5396 |
| HMR_6511 | Riboflavin metabolism | 12.2037 | 187.5121 | 14.2841 | 260.2498 | 2.0804 | -0.0983 | 0.5392 |
| HMR_4537 | Serotonin and melatonin biosynthesis | 2.7039 | 2.5171 | 2.6695 | 2.4142 | 0.0344 | 0.0155 | 0.4938 |
| HMR_4548 | Serotonin and melatonin biosynthesis | 20.8992 | 114.1807 | 25.7396 | 170.4545 | 4.8404 | -0.2869 | 0.6129 |
| HMR_4206 | Thiamine metabolism | 2.8476 | 3.7899 | 2.7186 | 3.4543 | 0.129 | 0.0479 | 0.4809 |
| HMR_4207 | Thiamine metabolism | 2.8471 | 3.7899 | 2.718 | 3.4543 | 0.129 | 0.0479 | 0.4809 |
| HMR_4064 | Vitamin B6 metabolism | 8.5466 | 12.5793 | 8.1621 | 11.4716 | 0.3845 | 0.0784 | 0.4688 |
| HMR_4065 | Vitamin B6 metabolism | 2.8008 | 7.2974 | 2.6739 | 6.6513 | 0.1269 | 0.034 | 0.4864 |
| HMR_4066 | Vitamin B6 metabolism | 2.8738 | 2.9126 | 2.7449 | 2.6576 | 0.1289 | 0.0546 | 0.4782 |
| HMR_4067 | Vitamin B6 metabolism | 2.8734 | 2.9143 | 2.7446 | 2.6593 | 0.1288 | 0.0546 | 0.4782 |
| HMR_4068 | Vitamin B6 metabolism | 2.8738 | 2.9126 | 2.7449 | 2.6576 | 0.1289 | 0.0546 | 0.4782 |
| HMR_4069 | Vitamin B6 metabolism | 2.8731 | 2.9143 | 2.7443 | 2.6593 | 0.1289 | 0.0546 | 0.4782 |
| HMR_4070 | Vitamin B6 metabolism | 2.8739 | 2.9143 | 2.745 | 2.6593 | 0.1289 | 0.0546 | 0.4782 |
| HMR_4071 | Vitamin B6 metabolism | 2.8736 | 2.9143 | 2.7447 | 2.6593 | 0.1289 | 0.0546 | 0.4782 |
| HMR_8032 | Metabolism of xenobiotics by cytochrome P450 | 6.7672 | 12.3241 | 7.0051 | 13.2041 | 0.2379 | -0.0471 | 0.5188 |
| HMR_3955 | Miscellaneous | 992.5025 | 29.368 | 992.941 | 25.6586 | 0.4384 | -0.0591 | 0.5236 |
| HMR_3984 | Miscellaneous | 65.8499 | 2178.8386 | 68.8719 | 2381.5466 | 3.022 | -0.0447 | 0.5178 |
| HMR_4201 | Miscellaneous | 15.8164 | 115.4051 | 16.6251 | 127.4062 | 0.8087 | -0.0519 | 0.5207 |
| HMR_4202 | Miscellaneous | 15.8157 | 115.4051 | 16.6242 | 127.4062 | 0.8086 | -0.0519 | 0.5207 |
| HMR_5409 | Miscellaneous | 6.4686 | 16.7952 | 5.9748 | 14.3542 | 0.4938 | 0.0885 | 0.4647 |
| HMR_7794 | Miscellaneous | 4.832 | 10.8782 | 4.0713 | 7.7363 | 0.7606 | 0.1763 | 0.43 |
| HMR_0010 | Pool reactions | 4.9708 | 3.5288 | 4.8208 | 3.3355 | 0.15 | 0.0573 | 0.4772 |
| HMR_0011 | Pool reactions | 19.8909 | 56.1393 | 19.2907 | 53.0436 | 0.6002 | 0.0574 | 0.4771 |
| HMR_0012 | Pool reactions | 19.8907 | 56.1484 | 19.2904 | 53.0527 | 0.6002 | 0.0574 | 0.4771 |
| HMR_0013 | Pool reactions | 0.0047 | 0.00E+00 | 0.0048 | 0.00E+00 | 0.00E+00 | -0.0078 | 0.5031 |
| HMR_0014 | Pool reactions | 0.0004 | 0.00E+00 | 0.0004 | 0.00E+00 | 0.00E+00 | -0.008 | 0.5032 |
| HMR_0031 | Pool reactions | 504.1869 | 63766.5281 | 504.2203 | 63766.3646 | 0.0333 | -1.00E-04 | 0.5 |
| HMR_0032 | Pool reactions | 495.8131 | 63766.5281 | 495.7797 | 63766.3646 | 0.0333 | 1.00E-04 | 0.5 |
| HMR_0546 | Pool reactions | 8.8122 | 9.4156 | 8.8477 | 9.5616 | 0.0355 | -0.0082 | 0.5033 |
| HMR_0547 | Pool reactions | 5.6943 | 3.343 | 5.7554 | 3.4323 | 0.0611 | -0.0235 | 0.5094 |

| | | | | | | | | |
|---|---|---|---|---|---|---|---|---|
| HMR_0548 | Pool reactions | 8.3382 | 9.1255 | 8.3909 | 9.2587 | 0.0528 | -0.0123 | 0.5049 |
| HMR_0549 | Pool reactions | 5.1898 | 4.584 | 5.2061 | 4.6218 | 0.0163 | -0.0054 | 0.5021 |
| HMR_0550 | Pool reactions | 3.3601 | 2.3751 | 3.3424 | 2.3467 | 0.0177 | 0.0081 | 0.4967 |
| HMR_0551 | Pool reactions | 7.4031 | 3.5859 | 7.4273 | 3.591 | 0.0242 | -0.009 | 0.5036 |
| HMR_0552 | Pool reactions | 2.545 | 4.3925 | 2.5786 | 4.4442 | 0.0336 | -0.0113 | 0.5045 |
| HMR_0555 | Pool reactions | 2.5796 | 3.1295 | 2.5507 | 3.0961 | 0.0289 | 0.0116 | 0.4954 |
| HMR_0556 | Pool reactions | 3.5808 | 3.7551 | 3.5711 | 3.7114 | 0.0097 | 0.0035 | 0.4986 |
| HMR_0557 | Pool reactions | 2.1548 | 1.9565 | 2.1474 | 1.9325 | 0.0074 | 0.0038 | 0.4985 |
| HMR_0558 | Pool reactions | 2.135 | 1.8308 | 2.131 | 1.8161 | 0.0039 | 0.002 | 0.4992 |
| HMR_0559 | Pool reactions | 5.2697 | 3.9015 | 5.3099 | 3.9128 | 0.0402 | -0.0144 | 0.5057 |
| HMR_0560 | Pool reactions | 3.6231 | 4.1853 | 3.5159 | 3.9684 | 0.1073 | 0.0376 | 0.485 |
| HMR_0685 | Pool reactions | 11.6521 | 15.6532 | 11.3627 | 14.9297 | 0.2894 | 0.0523 | 0.4791 |
| HMR_0686 | Pool reactions | 6.6574 | 6.3769 | 6.7282 | 6.5112 | 0.0707 | -0.0197 | 0.5079 |
| HMR_0687 | Pool reactions | 2.0185 | 1.8788 | 2.0499 | 1.9383 | 0.0314 | -0.0161 | 0.5064 |
| HMR_0688 | Pool reactions | -7.0853 | 8.5096 | -7.0676 | 8.4934 | 0.0177 | -0.0043 | 0.5017 |
| HMR_0689 | Pool reactions | 7.7842 | 8.0901 | 7.8011 | 8.0982 | 0.0169 | -0.0042 | 0.5017 |
| HMR_0690 | Pool reactions | 4.5142 | 5.2519 | 4.4482 | 5.1176 | 0.066 | 0.0205 | 0.4918 |
| HMR_0691 | Pool reactions | 7.0724 | 9.556 | 7.0207 | 9.4734 | 0.0517 | 0.0118 | 0.4953 |
| HMR_0692 | Pool reactions | -6.1145 | 9.4118 | -6.0385 | 9.2417 | 0.076 | -0.0176 | 0.507 |
| HMR_1185 | Pool reactions | 1.4423 | 0.2998 | 1.5623 | 0.3372 | 0.12 | -0.1503 | 0.5597 |
| HMR_1227 | Pool reactions | 1.4423 | 0.2998 | 1.5623 | 0.3372 | 0.12 | -0.1503 | 0.5597 |
| HMR_3598 | Pool reactions | 9.8233 | 11.5655 | 9.9547 | 11.8461 | 0.1314 | -0.0272 | 0.5108 |
| HMR_3602 | Pool reactions | 9.823 | 11.5841 | 9.9544 | 11.8648 | 0.1314 | -0.0271 | 0.5108 |
| HMR_9022 | Pool reactions | 4.97 | 9.7247 | 4.8245 | 9.1793 | 0.1455 | 0.0335 | 0.4867 |
| HMR_0004 | Transport, extracellular | 4.9716 | 3.5202 | 4.8215 | 3.3268 | 0.15 | 0.0573 | 0.4771 |
| HMR_0006 | Transport, extracellular | 0.0002 | 0.00E+00 | 0.0002 | 0.00E+00 | 0.00E+00 | 0.0386 | 0.4846 |
| HMR_0007 | Transport, extracellular | 4.9734 | 3.5105 | 4.8234 | 3.3169 | 0.15 | 0.0574 | 0.4771 |
| HMR_0008 | Transport, extracellular | 4.9723 | 3.5123 | 4.8222 | 3.3188 | 0.1501 | 0.0574 | 0.4771 |
| HMR_0009 | Transport, extracellular | 4.9716 | 3.5202 | 4.8215 | 3.3268 | 0.15 | 0.0573 | 0.4771 |
| HMR_0015 | Transport, extracellular | 87.989 | 1244.87 | 97.4843 | 1451.3484 | 9.4953 | -0.1829 | 0.5725 |
| HMR_0016 | Transport, extracellular | 88.8631 | 1436.8183 | 95.2818 | 1658.3849 | 6.4187 | -0.1154 | 0.5459 |
| HMR_0018 | Transport, extracellular | -4.4653 | 6.1181 | -4.2797 | 5.594 | 0.1856 | -0.0542 | 0.5216 |
| HMR_0019 | Transport, extracellular | 1.2989 | 0.4522 | 1.3106 | 0.4607 | 0.0117 | -0.0122 | 0.5049 |
| HMR_0187 | Transport, extracellular | 2.7449 | 3.8412 | 2.6149 | 3.5074 | 0.13 | 0.048 | 0.4809 |
| HMR_0190 | Transport, extracellular | 2.9968 | 3.8437 | 2.8731 | 3.5107 | 0.1237 | 0.0456 | 0.4818 |
| HMR_0191 | Transport, extracellular | 2.8473 | 3.7932 | 2.7187 | 3.4576 | 0.1287 | 0.0478 | 0.4809 |
| HMR_0194 | Transport, extracellular | 2.8492 | 3.7932 | 2.7199 | 3.4576 | 0.1293 | 0.048 | 0.4809 |
| HMR_0195 | Transport, extracellular | 2.8975 | 3.8852 | 2.7854 | 3.5629 | 0.1121 | 0.0411 | 0.4836 |
| HMR_0198 | Transport, extracellular | 2.8485 | 3.882 | 2.7127 | 3.5579 | 0.1358 | 0.0498 | 0.4802 |
| HMR_0199 | Transport, extracellular | 2.763 | 3.8218 | 2.6295 | 3.4901 | 0.1335 | 0.0494 | 0.4803 |
| HMR_0202 | Transport, extracellular | 2.9609 | 3.8231 | 2.8418 | 3.4918 | 0.1191 | 0.044 | 0.4824 |
| HMR_0203 | Transport, extracellular | 2.8323 | 3.794 | 2.7028 | 3.4585 | 0.1295 | 0.0481 | 0.4808 |
| HMR_0207 | Transport, extracellular | 2.865 | 3.7941 | 2.7366 | 3.4586 | 0.1284 | 0.0477 | 0.481 |
| HMR_0208 | Transport, extracellular | 2.8468 | 3.7933 | 2.718 | 3.4577 | 0.1288 | 0.0478 | 0.4809 |

| | | | | | | | | |
|---|---|---|---|---|---|---|---|---|
| HMR_0211 | Transport, extracellular | 2.8497 | 3.7933 | 2.7205 | 3.4577 | 0.1292 | 0.048 | 0.4809 |
| HMR_0212 | Transport, extracellular | 2.8106 | 3.7993 | 2.6839 | 3.463 | 0.1268 | 0.047 | 0.4812 |
| HMR_0215 | Transport, extracellular | 2.8912 | 3.7996 | 2.7597 | 3.4637 | 0.1315 | 0.0488 | 0.4805 |
| HMR_0216 | Transport, extracellular | 8.0405 | 22.9599 | 8.7368 | 25.2493 | 0.6963 | -0.1003 | 0.5399 |
| HMR_0224 | Transport, extracellular | 2.9668 | 7.6171 | 2.8058 | 7.024 | 0.161 | 0.0421 | 0.4832 |
| HMR_0225 | Transport, extracellular | 6.2963 | 7.4117 | 6.5623 | 7.3126 | 0.266 | -0.0693 | 0.5276 |
| HMR_0231 | Transport, extracellular | 2.7402 | 6.5671 | 2.6383 | 6.2864 | 0.1019 | 0.0284 | 0.4887 |
| HMR_0232 | Transport, extracellular | 2.7024 | 3.9047 | 2.5693 | 3.5831 | 0.1331 | 0.0487 | 0.4806 |
| HMR_0235 | Transport, extracellular | 3.1019 | 3.9109 | 2.9971 | 3.5912 | 0.1048 | 0.0383 | 0.4847 |
| HMR_0236 | Transport, extracellular | 3.1812 | 3.9712 | 3.0996 | 3.6709 | 0.0816 | 0.0295 | 0.4882 |
| HMR_0239 | Transport, extracellular | 2.6797 | 3.9613 | 2.545 | 3.6577 | 0.1347 | 0.0488 | 0.4805 |
| HMR_0240 | Transport, extracellular | 3.1163 | 3.9193 | 3.0259 | 3.6102 | 0.0905 | 0.033 | 0.4868 |
| HMR_0243 | Transport, extracellular | 2.6971 | 3.9131 | 2.5608 | 3.6019 | 0.1363 | 0.0497 | 0.4802 |
| HMR_0244 | Transport, extracellular | 2.7267 | 3.8623 | 2.5928 | 3.536 | 0.1339 | 0.0492 | 0.4804 |
| HMR_0247 | Transport, extracellular | 3.036 | 3.8658 | 2.9249 | 3.5406 | 0.1111 | 0.0408 | 0.4837 |
| HMR_0248 | Transport, extracellular | 4.5389 | 10.0606 | 4.7825 | 10.6454 | 0.2436 | -0.0535 | 0.5213 |
| HMR_0253 | Transport, extracellular | 3.2576 | 7.2632 | 3.0438 | 6.7103 | 0.2138 | 0.0572 | 0.4772 |
| HMR_0254 | Transport, extracellular | 2.7399 | 3.8437 | 2.6064 | 3.5145 | 0.1335 | 0.0492 | 0.4804 |
| HMR_0257 | Transport, extracellular | 3.0052 | 3.846 | 2.8898 | 3.5176 | 0.1153 | 0.0425 | 0.4831 |
| HMR_0258 | Transport, extracellular | 2.6567 | 5.1436 | 2.5474 | 4.873 | 0.1092 | 0.0345 | 0.4862 |
| HMR_0261 | Transport, extracellular | 4.4621 | 5.511 | 4.4839 | 5.3245 | 0.0218 | -0.0066 | 0.5026 |
| HMR_0262 | Transport, extracellular | 5.8584 | 15.6332 | 6.3395 | 17.1679 | 0.4811 | -0.084 | 0.5335 |
| HMR_0265 | Transport, extracellular | 3.1722 | 7.8419 | 2.9609 | 7.1581 | 0.2113 | 0.0546 | 0.4782 |
| HMR_0266 | Transport, extracellular | 2.8502 | 3.7933 | 2.7211 | 3.4577 | 0.1291 | 0.0479 | 0.4809 |
| HMR_0269 | Transport, extracellular | 2.8464 | 3.7933 | 2.7175 | 3.4577 | 0.1289 | 0.0479 | 0.4809 |
| HMR_0270 | Transport, extracellular | 2.8486 | 3.7932 | 2.7197 | 3.4577 | 0.1289 | 0.0479 | 0.4809 |
| HMR_0273 | Transport, extracellular | 2.8479 | 3.7932 | 2.7188 | 3.4576 | 0.1291 | 0.0479 | 0.4809 |
| HMR_0274 | Transport, extracellular | 2.8483 | 3.7933 | 2.7193 | 3.4577 | 0.1291 | 0.0479 | 0.4809 |
| HMR_0277 | Transport, extracellular | 2.8482 | 3.7933 | 2.7193 | 3.4577 | 0.1289 | 0.0479 | 0.4809 |
| HMR_0278 | Transport, extracellular | 2.8551 | 3.7935 | 2.7262 | 3.4579 | 0.1289 | 0.0479 | 0.4809 |
| HMR_0281 | Transport, extracellular | 2.8416 | 3.7934 | 2.7125 | 3.4578 | 0.129 | 0.0479 | 0.4809 |
| HMR_0282 | Transport, extracellular | 2.8633 | 3.794 | 2.7368 | 3.4586 | 0.1265 | 0.047 | 0.4813 |
| HMR_0287 | Transport, extracellular | 2.8339 | 3.7939 | 2.7026 | 3.4585 | 0.1313 | 0.0487 | 0.4806 |
| HMR_0288 | Transport, extracellular | 2.8463 | 3.7934 | 2.7167 | 3.4578 | 0.1296 | 0.0481 | 0.4808 |
| HMR_0291 | Transport, extracellular | 2.8502 | 3.7934 | 2.7219 | 3.4578 | 0.1283 | 0.0477 | 0.481 |
| HMR_0292 | Transport, extracellular | 2.7001 | 3.9102 | 2.567 | 3.5894 | 0.1331 | 0.0486 | 0.4806 |
| HMR_0295 | Transport, extracellular | 3.1096 | 3.9168 | 3.0058 | 3.5981 | 0.1037 | 0.0378 | 0.4849 |
| HMR_0296 | Transport, extracellular | 2.8126 | 3.7977 | 2.681 | 3.4627 | 0.1316 | 0.0488 | 0.4805 |
| HMR_0299 | Transport, extracellular | 2.888 | 3.7979 | 2.7626 | 3.4631 | 0.1254 | 0.0466 | 0.4814 |
| HMR_0300 | Transport, extracellular | 2.8433 | 3.7934 | 2.7139 | 3.4578 | 0.1294 | 0.0481 | 0.4808 |
| HMR_0303 | Transport, extracellular | 2.8533 | 3.7934 | 2.7248 | 3.4578 | 0.1285 | 0.0477 | 0.481 |
| HMR_0304 | Transport, extracellular | 2.8686 | 3.7945 | 2.7418 | 3.4591 | 0.1268 | 0.0471 | 0.4812 |
| HMR_0307 | Transport, extracellular | 2.829 | 3.7943 | 2.6981 | 3.459 | 0.1309 | 0.0486 | 0.4806 |
| HMR_0308 | Transport, extracellular | 2.856 | 3.7935 | 2.7271 | 3.4579 | 0.1289 | 0.0479 | 0.4809 |

| | | | | | | | | |
|---|---|---|---|---|---|---|---|---|
| HMR_0311 | Transport, extracellular | 2.8407 | 3.7934 | 2.7117 | 3.4578 | 0.1291 | 0.0479 | 0.4809 |
| HMR_0312 | Transport, extracellular | 2.881 | 3.7961 | 2.7552 | 3.4611 | 0.1257 | 0.0467 | 0.4814 |
| HMR_0317 | Transport, extracellular | 2.8182 | 3.7959 | 2.6867 | 3.4608 | 0.1316 | 0.0488 | 0.4805 |
| HMR_0318 | Transport, extracellular | 2.8463 | 3.7933 | 2.7172 | 3.4577 | 0.1291 | 0.0479 | 0.4809 |
| HMR_0321 | Transport, extracellular | 2.8503 | 3.7933 | 2.7214 | 3.4577 | 0.1289 | 0.0479 | 0.4809 |
| HMR_0322 | Transport, extracellular | 2.8478 | 3.7932 | 2.7189 | 3.4576 | 0.1289 | 0.0479 | 0.4809 |
| HMR_0325 | Transport, extracellular | 2.8486 | 3.7931 | 2.7196 | 3.4576 | 0.129 | 0.0479 | 0.4809 |
| HMR_0326 | Transport, extracellular | 2.8565 | 3.7932 | 2.7284 | 3.4577 | 0.1281 | 0.0476 | 0.481 |
| HMR_0329 | Transport, extracellular | 2.8401 | 3.7932 | 2.7103 | 3.4576 | 0.1298 | 0.0482 | 0.4808 |
| HMR_0330 | Transport, extracellular | 2.9097 | 3.8027 | 2.7875 | 3.469 | 0.1222 | 0.0453 | 0.4819 |
| HMR_0335 | Transport, extracellular | 2.7957 | 3.8022 | 2.6622 | 3.4685 | 0.1334 | 0.0495 | 0.4803 |
| HMR_0336 | Transport, extracellular | 2.8651 | 3.7941 | 2.7377 | 3.4587 | 0.1274 | 0.0473 | 0.4811 |
| HMR_0339 | Transport, extracellular | 2.8322 | 3.794 | 2.7018 | 3.4585 | 0.1304 | 0.0484 | 0.4807 |
| HMR_0340 | Transport, extracellular | 2.8756 | 3.7953 | 2.7491 | 3.46 | 0.1265 | 0.047 | 0.4813 |
| HMR_0343 | Transport, extracellular | 2.8228 | 3.7951 | 2.6917 | 3.4598 | 0.131 | 0.0486 | 0.4806 |
| HMR_0344 | Transport, extracellular | 2.8498 | 3.7932 | 2.7208 | 3.4576 | 0.1289 | 0.0479 | 0.4809 |
| HMR_0347 | Transport, extracellular | 2.8467 | 3.7932 | 2.7177 | 3.4576 | 0.129 | 0.0479 | 0.4809 |
| HMR_0348 | Transport, extracellular | 13.6975 | 11.4239 | 14.6573 | 11.3142 | 0.9598 | -0.2013 | 0.5798 |
| HMR_0351 | Transport, extracellular | 2.8442 | 8.2438 | 2.7157 | 7.4943 | 0.1286 | 0.0324 | 0.4871 |
| HMR_0352 | Transport, extracellular | 2.8525 | 3.7934 | 2.7237 | 3.4578 | 0.1288 | 0.0478 | 0.4809 |
| HMR_0355 | Transport, extracellular | 2.8441 | 3.7934 | 2.7149 | 3.4577 | 0.1291 | 0.048 | 0.4809 |
| HMR_0356 | Transport, extracellular | 2.8365 | 3.7938 | 2.7065 | 3.4582 | 0.1301 | 0.0483 | 0.4807 |
| HMR_0359 | Transport, extracellular | 2.8605 | 3.7938 | 2.7327 | 3.4583 | 0.1278 | 0.0475 | 0.4811 |
| HMR_0360 | Transport, extracellular | 2.916 | 3.8043 | 2.7948 | 3.471 | 0.1212 | 0.0449 | 0.4821 |
| HMR_0363 | Transport, extracellular | 2.7911 | 3.8039 | 2.6571 | 3.4704 | 0.1339 | 0.0497 | 0.4802 |
| HMR_0364 | Transport, extracellular | 2.8986 | 3.8 | 2.7765 | 3.466 | 0.1222 | 0.0453 | 0.4819 |
| HMR_0367 | Transport, extracellular | 2.8041 | 3.7996 | 2.6702 | 3.4656 | 0.1338 | 0.0497 | 0.4802 |
| HMR_0368 | Transport, extracellular | 2.8531 | 3.7934 | 2.7239 | 3.4578 | 0.1291 | 0.048 | 0.4809 |
| HMR_0371 | Transport, extracellular | 2.8435 | 3.7933 | 2.7147 | 3.4577 | 0.1288 | 0.0478 | 0.4809 |
| HMR_0372 | Transport, extracellular | 2.8528 | 3.7933 | 2.7237 | 3.4577 | 0.1291 | 0.0479 | 0.4809 |
| HMR_0375 | Transport, extracellular | 2.8437 | 3.7933 | 2.7149 | 3.4577 | 0.1288 | 0.0478 | 0.4809 |
| HMR_0376 | Transport, extracellular | 3.0878 | 3.8932 | 2.9869 | 3.5749 | 0.1009 | 0.0369 | 0.4853 |
| HMR_0379 | Transport, extracellular | 2.7044 | 3.8896 | 2.5691 | 3.5702 | 0.1353 | 0.0495 | 0.4803 |
| HMR_0380 | Transport, extracellular | 2.9144 | 3.8043 | 2.7936 | 3.4712 | 0.1208 | 0.0448 | 0.4821 |
| HMR_0383 | Transport, extracellular | 2.7924 | 3.8038 | 2.6582 | 3.4706 | 0.1342 | 0.0498 | 0.4802 |
| HMR_0384 | Transport, extracellular | 2.849 | 3.7932 | 2.7202 | 3.4576 | 0.1288 | 0.0478 | 0.4809 |
| HMR_0387 | Transport, extracellular | 2.8475 | 3.7932 | 2.7183 | 3.4576 | 0.1292 | 0.048 | 0.4809 |
| HMR_0388 | Transport, extracellular | 2.8515 | 3.7934 | 2.7224 | 3.4578 | 0.1291 | 0.0479 | 0.4809 |
| HMR_0391 | Transport, extracellular | 2.8451 | 3.7933 | 2.7162 | 3.4577 | 0.1289 | 0.0479 | 0.4809 |
| HMR_0392 | Transport, extracellular | 2.8479 | 3.7932 | 2.719 | 3.4576 | 0.129 | 0.0479 | 0.4809 |
| HMR_0395 | Transport, extracellular | 2.8486 | 3.7932 | 2.7196 | 3.4576 | 0.129 | 0.0479 | 0.4809 |
| HMR_0396 | Transport, extracellular | 19.1014 | 15.217 | 19.9814 | 15.0363 | 0.88 | -0.16 | 0.5636 |
| HMR_0399 | Transport, extracellular | 2.8449 | 8.2148 | 2.7158 | 7.4871 | 0.129 | 0.0326 | 0.487 |
| HMR_0400 | Transport, extracellular | 3.0898 | 3.9002 | 2.9954 | 3.5875 | 0.0944 | 0.0345 | 0.4862 |

| | | | | | | | | |
|---|---|---|---|---|---|---|---|---|
| HMR_0403 | Transport, extracellular | 2.7058 | 3.8951 | 2.5691 | 3.5807 | 0.1367 | 0.05 | 0.4801 |
| HMR_0404 | Transport, extracellular | 2.89 | 3.7976 | 2.7662 | 3.463 | 0.1238 | 0.046 | 0.4817 |
| HMR_0407 | Transport, extracellular | 2.8107 | 3.7974 | 2.6778 | 3.4627 | 0.1329 | 0.0493 | 0.4803 |
| HMR_0408 | Transport, extracellular | 440.0046 | 75327.9727 | 439.9849 | 75326.1146 | 0.0197 | 1.00E-04 | 0.5 |
| HMR_0411 | Transport, extracellular | 2.8501 | 8.2464 | 2.7206 | 7.515 | 0.1295 | 0.0326 | 0.487 |
| HMR_0412 | Transport, extracellular | 2.8947 | 3.799 | 2.7719 | 3.4648 | 0.1228 | 0.0456 | 0.4818 |
| HMR_0415 | Transport, extracellular | 2.8071 | 3.7987 | 2.6736 | 3.4644 | 0.1335 | 0.0495 | 0.4802 |
| HMR_0416 | Transport, extracellular | 2.85 | 3.7933 | 2.721 | 3.4577 | 0.129 | 0.0479 | 0.4809 |
| HMR_0419 | Transport, extracellular | 2.8466 | 3.7933 | 2.7176 | 3.4577 | 0.129 | 0.0479 | 0.4809 |
| HMR_0420 | Transport, extracellular | 2.8501 | 3.7933 | 2.7212 | 3.4577 | 0.129 | 0.0479 | 0.4809 |
| HMR_0423 | Transport, extracellular | 2.8464 | 3.7933 | 2.7174 | 3.4577 | 0.129 | 0.0479 | 0.4809 |
| HMR_0424 | Transport, extracellular | 2.8446 | 3.7933 | 2.7159 | 3.4577 | 0.1288 | 0.0478 | 0.4809 |
| HMR_0427 | Transport, extracellular | 2.852 | 3.7933 | 2.7227 | 3.4577 | 0.1292 | 0.048 | 0.4809 |
| HMR_0428 | Transport, extracellular | 2.8222 | 3.796 | 2.6937 | 3.4602 | 0.1284 | 0.0477 | 0.481 |
| HMR_0431 | Transport, extracellular | 2.8768 | 3.7962 | 2.7473 | 3.4606 | 0.1295 | 0.0481 | 0.4808 |
| HMR_0432 | Transport, extracellular | 2.8503 | 3.7932 | 2.7218 | 3.4577 | 0.1286 | 0.0478 | 0.481 |
| HMR_0435 | Transport, extracellular | 2.8462 | 3.7932 | 2.7168 | 3.4576 | 0.1294 | 0.048 | 0.4808 |
| HMR_0436 | Transport, extracellular | 2.849 | 3.7932 | 2.7202 | 3.4576 | 0.1288 | 0.0478 | 0.4809 |
| HMR_0439 | Transport, extracellular | 2.8474 | 3.7932 | 2.7183 | 3.4576 | 0.1291 | 0.0479 | 0.4809 |
| HMR_0440 | Transport, extracellular | 0.839 | 0.1234 | 1.0735 | 0.1885 | 0.2344 | -0.4198 | 0.6627 |
| HMR_0444 | Transport, extracellular | -53.0912 | 2039.1093 | -69.829 | 3088.7337 | 16.7378 | 0.2337 | 0.4076 |
| HMR_1910 | Transport, extracellular | 1.4297 | 1.5314 | 1.3641 | 1.3953 | 0.0655 | 0.0383 | 0.4847 |
| HMR_1911 | Transport, extracellular | 2.705 | 5.2297 | 2.5817 | 4.7765 | 0.1233 | 0.039 | 0.4845 |
| HMR_1913 | Transport, extracellular | 1.4209 | 2.0523 | 1.3566 | 1.871 | 0.0643 | 0.0325 | 0.4871 |
| HMR_1914 | Transport, extracellular | 4.342 | 22.753 | 4.156 | 20.8578 | 0.186 | 0.0282 | 0.4888 |
| HMR_1919 | Transport, extracellular | 1.1878 | 0.6564 | 1.1728 | 0.64 | 0.015 | 0.0132 | 0.4947 |
| HMR_3858 | Transport, extracellular | 5.8987 | 16.2208 | 5.8672 | 16.0474 | 0.0315 | 0.0056 | 0.4978 |
| HMR_3951 | Transport, extracellular | 27.8085 | 396.3998 | 36.8448 | 514.695 | 9.0363 | -0.2994 | 0.6177 |
| HMR_3964 | Transport, extracellular | 2.8449 | 8.2142 | 2.7158 | 7.487 | 0.129 | 0.0326 | 0.487 |
| HMR_4321 | Transport, extracellular | -121.9996 | 2544.3018 | -121.0896 | 2528.2778 | 0.91 | -0.0128 | 0.5051 |
| HMR_4432 | Transport, extracellular | 4.3179 | 4.6112 | 4.0777 | 4.1144 | 0.2402 | 0.0813 | 0.4676 |
| HMR_4433 | Transport, extracellular | -7.9212 | 28.9049 | -7.522 | 26.0694 | 0.3992 | -0.0538 | 0.5215 |
| HMR_4434 | Transport, extracellular | -7.9212 | 28.9049 | -7.522 | 26.0694 | 0.3992 | -0.0538 | 0.5215 |
| HMR_4435 | Transport, extracellular | -7.9212 | 28.9049 | -7.522 | 26.0694 | 0.3992 | -0.0538 | 0.5215 |
| HMR_4436 | Transport, extracellular | 114.3498 | 243627.9905 | 113.7198 | 243631.314 | 0.63 | 0.0009 | 0.4996 |
| HMR_4691 | Transport, extracellular | 3.7287 | 12.1682 | 3.708 | 12.0997 | 0.0207 | 0.0042 | 0.4983 |
| HMR_4721 | Transport, extracellular | 0.0025 | 0.00E+00 | 0.0025 | 0.00E+00 | 0.00E+00 | 0.00E+00 | 0.5 |
| HMR_4844 | Transport, extracellular | -7.43 | 220485.3653 | -7.237 | 220487.2085 | 0.1931 | -0.0003 | 0.5001 |
| HMR_4845 | Transport, extracellular | 8.5076 | 73.5203 | 8.147 | 67.4296 | 0.3605 | 0.0304 | 0.4879 |
| HMR_4849 | Transport, extracellular | 959.6994 | 1862.6423 | 958.4935 | 1978.6574 | 1.2059 | 0.0195 | 0.4922 |
| HMR_4858 | Transport, extracellular | -37.4986 | 305.4577 | -34.2478 | 257.9776 | 3.2508 | -0.137 | 0.5545 |
| HMR_4873 | Transport, extracellular | -362.0261 | 13517.994 | 21.8153 | 13677.2501 | 383.8414 | -2.3276 | 0.99 |
| HMR_4885 | Transport, extracellular | -747.243 | 3760.8001 | -772.4471 | 3802.8496 | 25.2041 | 0.2898 | 0.386 |
| HMR_4896 | Transport, extracellular | -986.4477 | 43.8769 | -985.4725 | 49.7098 | 0.9753 | -0.1008 | 0.5402 |

| | | | | | | | | |
|---|---|---|---|---|---|---|---|---|
| HMR_4910 | Transport, extracellular | 1.6209 | 2.7591 | 1.5561 | 2.5415 | 0.0648 | 0.0282 | 0.4888 |
| HMR_4911 | Transport, extracellular | 1.1327 | 1.3939 | 1.0885 | 1.2858 | 0.0441 | 0.027 | 0.4892 |
| HMR_4912 | Transport, extracellular | 2.0005 | 4.2816 | 1.9217 | 3.9473 | 0.0788 | 0.0275 | 0.489 |
| HMR_4919 | Transport, extracellular | 985.4518 | 136.236 | 986.9298 | 111.2521 | 1.478 | -0.0939 | 0.5374 |
| HMR_4928 | Transport, extracellular | 5.8832 | 38.9795 | 5.8468 | 38.4976 | 0.0364 | 0.0041 | 0.4983 |
| HMR_4931 | Transport, extracellular | 79.647 | 2516.0942 | 65.6381 | 2067.3142 | 14.0089 | 0.2069 | 0.418 |
| HMR_4932 | Transport, extracellular | 2.3432 | 5.6678 | 2.2534 | 5.2429 | 0.0899 | 0.0272 | 0.4891 |
| HMR_4933 | Transport, extracellular | 4.7383 | 23.219 | 4.6535 | 22.3108 | 0.0848 | 0.0126 | 0.495 |
| HMR_4934 | Transport, extracellular | 4.7377 | 23.2121 | 4.6529 | 22.3043 | 0.0848 | 0.0126 | 0.495 |
| HMR_4935 | Transport, extracellular | 5.8755 | 38.8644 | 5.8411 | 38.416 | 0.0343 | 0.0039 | 0.4984 |
| HMR_4939 | Transport, extracellular | 2.0769 | 3.545 | 2.0711 | 3.5332 | 0.0057 | 0.0021 | 0.4991 |
| HMR_4946 | Transport, extracellular | 2.1703 | 5.6183 | 2.0777 | 5.1534 | 0.0927 | 0.0282 | 0.4887 |
| HMR_4952 | Transport, extracellular | 20.4175 | 294.8366 | 15.9492 | 185.9551 | 4.4683 | 0.2038 | 0.4193 |
| HMR_4954 | Transport, extracellular | -34.3449 | 582.3123 | -30.0495 | 447.0295 | 4.2954 | -0.1339 | 0.5533 |
| HMR_4956 | Transport, extracellular | 8.4041 | 72.6048 | 8.0544 | 66.6269 | 0.3497 | 0.0296 | 0.4882 |
| HMR_4976 | Transport, extracellular | 5.8752 | 38.861 | 5.84 | 38.3991 | 0.0352 | 0.004 | 0.4984 |
| HMR_4982 | Transport, extracellular | -93.6207 | 1474.2873 | -102.4714 | 1885.0974 | 8.8507 | 0.1527 | 0.4393 |
| HMR_4983 | Transport, extracellular | 8.2059 | 45.9446 | 7.8581 | 42.1318 | 0.3479 | 0.0371 | 0.4852 |
| HMR_4989 | Transport, extracellular | -958.6225 | 356.218 | -957.3413 | 376.0982 | 1.2812 | -0.0473 | 0.5189 |
| HMR_4990 | Transport, extracellular | -293.4265 | 4691.2983 | -366.2917 | 3928.5556 | 72.8652 | 0.7848 | 0.2163 |
| HMR_4992 | Transport, extracellular | -917.425 | 1002.9748 | -906.0614 | 1501.6082 | 11.3635 | -0.2271 | 0.5898 |
| HMR_4995 | Transport, extracellular | 12.8813 | 157.0492 | 12.5096 | 148.2408 | 0.3717 | 0.0213 | 0.4915 |
| HMR_4996 | Transport, extracellular | 61.5898 | 747.108 | 56.8507 | 641.2907 | 4.7391 | 0.1272 | 0.4494 |
| HMR_4999 | Transport, extracellular | 86.3164 | 243634.914 | 87.0777 | 243637.5231 | 0.7614 | -0.0011 | 0.5004 |
| HMR_5000 | Transport, extracellular | -25.2897 | 157.0377 | -27.6379 | 186.2104 | 2.3482 | 0.1267 | 0.4496 |
| HMR_5003 | Transport, extracellular | 5.5409 | 105.1117 | 5.5163 | 95.7993 | 0.0247 | 0.0017 | 0.4993 |
| HMR_5008 | Transport, extracellular | 12.6108 | 54.1943 | 12.0761 | 49.6957 | 0.5347 | 0.0525 | 0.4791 |
| HMR_5018 | Transport, extracellular | 5.8774 | 38.8938 | 5.844 | 38.4587 | 0.0334 | 0.0038 | 0.4985 |
| HMR_5023 | Transport, extracellular | 8.453 | 33.3819 | 8.0633 | 30.3749 | 0.3897 | 0.0488 | 0.4805 |
| HMR_5029 | Transport, extracellular | -177.9222 | 3084.421 | -177.0513 | 3108.16 | 0.8709 | -0.0111 | 0.5044 |
| HMR_5032 | Transport, extracellular | -2.0986 | 220530.8376 | -2.2534 | 220529.2734 | 0.1548 | 0.0002 | 0.4999 |
| HMR_5034 | Transport, extracellular | 12.6144 | 54.2051 | 12.0798 | 49.706 | 0.5347 | 0.0524 | 0.4791 |
| HMR_5035 | Transport, extracellular | 12.6102 | 54.1926 | 12.0756 | 49.6942 | 0.5346 | 0.0525 | 0.4791 |
| HMR_5037 | Transport, extracellular | 12.6129 | 54.2009 | 12.0783 | 49.7023 | 0.5346 | 0.0524 | 0.4791 |
| HMR_5038 | Transport, extracellular | 8.508 | 73.4574 | 8.1474 | 67.3763 | 0.3606 | 0.0304 | 0.4879 |
| HMR_5039 | Transport, extracellular | 8.3993 | 60.8548 | 8.0422 | 55.7614 | 0.3571 | 0.0331 | 0.4868 |
| HMR_5040 | Transport, extracellular | 8.2066 | 45.935 | 7.8588 | 42.1224 | 0.3478 | 0.0371 | 0.4852 |
| HMR_5041 | Transport, extracellular | 8.2061 | 45.9429 | 7.8582 | 42.1303 | 0.3479 | 0.0371 | 0.4852 |
| HMR_5042 | Transport, extracellular | 5.3305 | 220515.9192 | 4.9828 | 220515.2384 | 0.3477 | 0.0005 | 0.4998 |
| HMR_5068 | Transport, extracellular | 38.908 | 1718.5002 | 40.032 | 1820.8708 | 1.1239 | -0.0189 | 0.5075 |
| HMR_5070 | Transport, extracellular | 959.1877 | 1910.2481 | 957.8291 | 2044.5086 | 1.3587 | 0.0216 | 0.4914 |
| HMR_5071 | Transport, extracellular | 39.6217 | 1786.1775 | 40.7824 | 1894.1069 | 1.1607 | -0.0191 | 0.5076 |
| HMR_5073 | Transport, extracellular | 580.6287 | 78072.1792 | 577.7247 | 78351.0741 | 2.904 | 0.0073 | 0.4971 |
| HMR_5074 | Transport, extracellular | 339.4099 | 246948.5298 | 329.4927 | 250580.342 | 9.9172 | 0.0141 | 0.4944 |

| | | | | | | | | |
|---|---|---|---|---|---|---|---|---|
| HMR_5076 | Transport, extracellular | 385.8077 | 74328.3626 | 389.0218 | 74757.829 | 3.2141 | -0.0083 | 0.5033 |
| HMR_5077 | Transport, extracellular | 400.3186 | 76083.07 | 404.3704 | 76555.0354 | 4.0518 | -0.0104 | 0.5041 |
| HMR_5078 | Transport, extracellular | 372.3495 | 235412.4457 | 360.8396 | 239949.0244 | 11.5099 | 0.0167 | 0.4933 |
| HMR_5079 | Transport, extracellular | 406.7152 | 76791.2136 | 409.2566 | 77073.5242 | 2.5414 | -0.0065 | 0.5026 |
| HMR_5080 | Transport, extracellular | 340.1032 | 246701.6561 | 330.1778 | 250342.4787 | 9.9254 | 0.0141 | 0.4944 |
| HMR_5082 | Transport, extracellular | 339.8038 | 246807.9367 | 329.8857 | 250443.5927 | 9.9181 | 0.0141 | 0.4944 |
| HMR_5084 | Transport, extracellular | 387.0167 | 74414.6794 | 390.067 | 74816.4536 | 3.0503 | -0.0079 | 0.5032 |
| HMR_5085 | Transport, extracellular | 310.8784 | 256335.7871 | 300.7553 | 259775.781 | 10.1231 | 0.0141 | 0.4944 |
| HMR_5087 | Transport, extracellular | 593.5668 | 76762.3406 | 589.2851 | 77228.2354 | 4.2817 | 0.0109 | 0.4956 |
| HMR_5088 | Transport, extracellular | 405.204 | 76627.4152 | 408.0501 | 76946.509 | 2.8461 | -0.0073 | 0.5029 |
| HMR_5089 | Transport, extracellular | 324.9725 | 251953.6753 | 330.6119 | 250192.34 | 5.6394 | -0.008 | 0.5032 |
| HMR_5091 | Transport, extracellular | 405.8485 | 76698.1359 | 408.6506 | 77010.5123 | 2.8021 | -0.0071 | 0.5029 |
| HMR_5092 | Transport, extracellular | 340.1367 | 246689.9524 | 330.6182 | 250190.1499 | 9.5185 | 0.0135 | 0.4946 |
| HMR_5094 | Transport, extracellular | 40.31 | 1863.6269 | 41.5128 | 1979.2681 | 1.2028 | -0.0194 | 0.5077 |
| HMR_5129 | Transport, extracellular | 8.2061 | 45.9424 | 7.8583 | 42.1299 | 0.3479 | 0.0371 | 0.4852 |
| HMR_5206 | Transport, extracellular | 0.0711 | 0.0026 | 0.0708 | 0.0026 | 0.0003 | 0.0043 | 0.4983 |
| HMR_5231 | Transport, extracellular | -0.1465 | 0.0074 | -0.1506 | 0.0077 | 0.0042 | 0.0339 | 0.4865 |
| HMR_5295 | Transport, extracellular | 4.2641 | 18.4469 | 4.0738 | 16.8413 | 0.1903 | 0.032 | 0.4872 |
| HMR_5303 | Transport, extracellular | 11.5494 | 26.2877 | 11.108 | 24.4579 | 0.4414 | 0.062 | 0.4753 |
| HMR_5305 | Transport, extracellular | 8.4221 | 71.992 | 8.0719 | 66.1398 | 0.3502 | 0.0298 | 0.4881 |
| HMR_5307 | Transport, extracellular | 6.1711 | 42.312 | 5.8781 | 38.2575 | 0.293 | 0.0326 | 0.487 |
| HMR_5308 | Transport, extracellular | 8.4276 | 72.0843 | 8.0764 | 66.2115 | 0.3512 | 0.0299 | 0.4881 |
| HMR_5310 | Transport, extracellular | 6.174 | 42.3548 | 5.8805 | 38.2899 | 0.2935 | 0.0327 | 0.487 |
| HMR_5311 | Transport, extracellular | 8.4265 | 72.0678 | 8.0749 | 66.1904 | 0.3515 | 0.0299 | 0.4881 |
| HMR_5313 | Transport, extracellular | 6.1734 | 42.3466 | 5.8797 | 38.2798 | 0.2937 | 0.0327 | 0.487 |
| HMR_5314 | Transport, extracellular | 8.4399 | 72.2978 | 8.0836 | 66.3331 | 0.3563 | 0.0303 | 0.4879 |
| HMR_5315 | Transport, extracellular | 8.4434 | 72.3602 | 8.0903 | 66.4452 | 0.3531 | 0.03 | 0.488 |
| HMR_5316 | Transport, extracellular | 8.4361 | 72.2335 | 8.0836 | 66.3331 | 0.3525 | 0.0299 | 0.4881 |
| HMR_5317 | Transport, extracellular | 8.4361 | 72.2336 | 8.0837 | 66.3347 | 0.3524 | 0.0299 | 0.4881 |
| HMR_5318 | Transport, extracellular | 8.4363 | 72.2366 | 8.0839 | 66.3373 | 0.3525 | 0.0299 | 0.4881 |
| HMR_5319 | Transport, extracellular | 8.4362 | 72.2349 | 8.0838 | 66.3358 | 0.3524 | 0.0299 | 0.4881 |
| HMR_5320 | Transport, extracellular | 8.4089 | 71.7628 | 8.0592 | 65.928 | 0.3497 | 0.0298 | 0.4881 |
| HMR_5322 | Transport, extracellular | 8.4276 | 72.0869 | 8.0771 | 66.2264 | 0.3505 | 0.0298 | 0.4881 |
| HMR_5324 | Transport, extracellular | 8.439 | 72.2848 | 8.0864 | 66.3814 | 0.3526 | 0.0299 | 0.4881 |
| HMR_5326 | Transport, extracellular | 8.4271 | 72.0779 | 8.0754 | 66.1986 | 0.3516 | 0.0299 | 0.4881 |
| HMR_5328 | Transport, extracellular | 5.4604 | 18.9389 | 5.281 | 17.704 | 0.1794 | 0.0296 | 0.4882 |
| HMR_5330 | Transport, extracellular | 8.5411 | 80.9982 | 8.2665 | 76.6103 | 0.2746 | 0.0219 | 0.4913 |
| HMR_5332 | Transport, extracellular | 2.8317 | 8.2235 | 2.6929 | 7.4452 | 0.1387 | 0.035 | 0.486 |
| HMR_5333 | Transport, extracellular | 7.0529 | 50.5072 | 6.8402 | 47.4885 | 0.2127 | 0.0215 | 0.4914 |
| HMR_5335 | Transport, extracellular | 5.4082 | 31.7918 | 5.2016 | 29.3602 | 0.2066 | 0.0264 | 0.4895 |
| HMR_5413 | Transport, extracellular | 2.9227 | 9.9222 | 2.8205 | 9.219 | 0.1022 | 0.0234 | 0.4907 |
| HMR_5429 | Transport, extracellular | 608.581 | 1839.4039 | 546.5837 | 1625.8973 | 61.9974 | 1.0532 | 0.1461 |
| HMR_5430 | Transport, extracellular | 21.9037 | 416.1064 | 20.6012 | 371.3317 | 1.3025 | 0.0464 | 0.4815 |
| HMR_5433 | Transport, extracellular | 591.0727 | 50385.0033 | 500.722 | 52932.1563 | 90.3507 | 0.2811 | 0.3893 |

| | | | | | | | | |
|---|---|---|---|---|---|---|---|---|
| HMR_5441 | Transport, extracellular | 341.3263 | 246269.525 | 331.3391 | 249942.1107 | 9.9871 | 0.0142 | 0.4943 |
| HMR_5442 | Transport, extracellular | 7.7811 | 56.341 | 7.3898 | 50.8603 | 0.3913 | 0.0378 | 0.4849 |
| HMR_5443 | Transport, extracellular | 7.9198 | 28.905 | 7.5205 | 26.0695 | 0.3993 | 0.0539 | 0.4785 |
| HMR_5444 | Transport, extracellular | 7.9198 | 28.905 | 7.5205 | 26.0695 | 0.3993 | 0.0539 | 0.4785 |
| HMR_5445 | Transport, extracellular | 7.9198 | 28.905 | 7.5205 | 26.0695 | 0.3993 | 0.0539 | 0.4785 |
| HMR_5446 | Transport, extracellular | 34.1657 | 43.169 | 32.6779 | 40.4606 | 1.4877 | 0.1627 | 0.4354 |
| HMR_5447 | Transport, extracellular | 2.1595 | 4.2525 | 2.0775 | 3.9384 | 0.082 | 0.0287 | 0.4886 |
| HMR_5448 | Transport, extracellular | 968.5189 | 294.1614 | 969.9505 | 267.1794 | 1.4317 | -0.0604 | 0.5241 |
| HMR_5450 | Transport, extracellular | 8.508 | 73.4574 | 8.1474 | 67.3763 | 0.3606 | 0.0304 | 0.4879 |
| HMR_5455 | Transport, extracellular | 4.2221 | 18.094 | 4.0409 | 16.5772 | 0.1812 | 0.0308 | 0.4877 |
| HMR_5457 | Transport, extracellular | 2.8218 | 8.0799 | 2.7012 | 7.4053 | 0.1206 | 0.0307 | 0.4878 |
| HMR_5458 | Transport, extracellular | -66.8819 | 314334.4641 | -70.2319 | 314197.9852 | 3.35 | 0.0042 | 0.4983 |
| HMR_5459 | Transport, extracellular | -93.4371 | 313435.3412 | -93.5168 | 313532.643 | 0.0798 | 0.0001 | 0.5 |
| HMR_5460 | Transport, extracellular | 95.5337 | 311521.8008 | 92.0856 | 311934.6668 | 3.4481 | 0.0044 | 0.4983 |
| HMR_5461 | Transport, extracellular | 150.2506 | 303846.6157 | 148.2388 | 304232.8963 | 2.0119 | 0.0026 | 0.499 |
| HMR_5462 | Transport, extracellular | 136.1885 | 306438.1392 | 133.6008 | 306887.588 | 2.5877 | 0.0033 | 0.4987 |
| HMR_5463 | Transport, extracellular | 136.5527 | 306381.6878 | 133.951 | 306834.3544 | 2.6017 | 0.0033 | 0.4987 |
| HMR_5464 | Transport, extracellular | 137.0299 | 306307.7961 | 134.4204 | 306763.0347 | 2.6095 | 0.0033 | 0.4987 |
| HMR_5465 | Transport, extracellular | -6.3204 | 316091.5118 | -5.5313 | 316152.5423 | 0.7891 | -0.001 | 0.5004 |
| HMR_5466 | Transport, extracellular | 171.4689 | 299862.2093 | 168.6213 | 300460.8914 | 2.8476 | 0.0037 | 0.4985 |
| HMR_5467 | Transport, extracellular | 94.2824 | 311639.8817 | 98.4846 | 311236.0685 | 4.2023 | -0.0053 | 0.5021 |
| HMR_5468 | Transport, extracellular | -88.8432 | 312200.544 | -86.7587 | 312458.4975 | 2.0845 | -0.0026 | 0.5011 |
| HMR_5469 | Transport, extracellular | 154.3134 | 303446.6616 | 133.0791 | 306966.7875 | 21.2343 | 0.0272 | 0.4892 |
| HMR_5470 | Transport, extracellular | 91.7 | 311908.7609 | 89.4118 | 312193.9859 | 2.2882 | 0.0029 | 0.4988 |
| HMR_5471 | Transport, extracellular | 134.6916 | 306669.1353 | 132.2009 | 307099.7527 | 2.4907 | 0.0032 | 0.4987 |
| HMR_5472 | Transport, extracellular | 136.1472 | 306444.5465 | 133.0715 | 306967.9601 | 3.0757 | 0.0039 | 0.4984 |
| HMR_5473 | Transport, extracellular | 27.5516 | 316815.4099 | 24.2531 | 317024.154 | 3.2985 | 0.0041 | 0.4983 |
| HMR_5474 | Transport, extracellular | 29.0342 | 315896.1768 | 22.1063 | 316144.1884 | 6.9279 | 0.0087 | 0.4965 |
| HMR_5475 | Transport, extracellular | 85.2132 | 312188.6123 | 79.76 | 312748.619 | 5.4533 | 0.0069 | 0.4972 |
| HMR_5476 | Transport, extracellular | 70.7937 | 313778.1382 | 64.747 | 314300.4394 | 6.0467 | 0.0076 | 0.497 |
| HMR_5477 | Transport, extracellular | 71.1691 | 313747.5812 | 65.108 | 314273.5812 | 6.0612 | 0.0076 | 0.4969 |
| HMR_5478 | Transport, extracellular | 71.6611 | 313707.5747 | 65.592 | 314237.6633 | 6.0691 | 0.0077 | 0.4969 |
| HMR_5479 | Transport, extracellular | -60.9665 | 313646.299 | -65.1471 | 313402.5781 | 4.1805 | 0.0053 | 0.4979 |
| HMR_5480 | Transport, extracellular | 107.1977 | 309631.0679 | 100.8902 | 310431.2611 | 6.3075 | 0.008 | 0.4968 |
| HMR_5481 | Transport, extracellular | 27.7534 | 315920.3439 | 28.6288 | 315951.4877 | 0.8754 | -0.0011 | 0.5004 |
| HMR_5482 | Transport, extracellular | 22.2136 | 316072.5218 | 16.6781 | 316245.6729 | 5.5355 | 0.007 | 0.4972 |
| HMR_5483 | Transport, extracellular | 89.5167 | 312059.5882 | 64.2092 | 314340.3113 | 25.3075 | 0.032 | 0.4872 |
| HMR_5484 | Transport, extracellular | 25.1217 | 315995.6712 | 19.378 | 316191.7165 | 5.7438 | 0.0072 | 0.4971 |
| HMR_5485 | Transport, extracellular | 69.2516 | 313902.4689 | 63.3043 | 314406.9337 | 5.9473 | 0.0075 | 0.497 |
| HMR_5486 | Transport, extracellular | 70.7512 | 313781.6099 | 64.2013 | 314340.9135 | 6.5499 | 0.0083 | 0.4967 |
| HMR_5487 | Transport, extracellular | 1.3641 | 317103.6091 | -2.2513 | 317140.4013 | 3.6154 | 0.0045 | 0.4982 |
| HMR_5488 | Transport, extracellular | 57.6225 | 315156.5229 | 55.4257 | 315338.8839 | 2.1968 | 0.0028 | 0.4989 |
| HMR_5489 | Transport, extracellular | 43.1775 | 316343.2554 | 40.4004 | 316524.2919 | 2.7771 | 0.0035 | 0.4986 |
| HMR_5490 | Transport, extracellular | 43.554 | 316324.4874 | 40.762 | 316507.4553 | 2.792 | 0.0035 | 0.4986 |

| | | | | | | | | |
|---|---|---|---|---|---|---|---|---|
| HMR_5491 | Transport, extracellular | 44.0475 | 316299.7262 | 41.2469 | 316484.6753 | 2.8006 | 0.0035 | 0.4986 |
| HMR_5492 | Transport, extracellular | 88.0936 | 312076.6018 | 88.9752 | 312034.4823 | 0.8816 | -0.0011 | 0.5004 |
| HMR_5493 | Transport, extracellular | 79.7023 | 313322.2657 | 76.6168 | 313646.0045 | 3.0855 | 0.0039 | 0.4984 |
| HMR_5494 | Transport, extracellular | 0.0834 | 317090.7833 | 4.2568 | 317130.1072 | 4.1734 | -0.0052 | 0.5021 |
| HMR_5495 | Transport, extracellular | 5.4448 | 317068.1012 | 7.6668 | 317093.514 | 2.222 | -0.0028 | 0.5011 |
| HMR_5496 | Transport, extracellular | 61.9749 | 315202.3012 | 39.8618 | 316549.307 | 22.1131 | 0.0278 | 0.4889 |
| HMR_5497 | Transport, extracellular | -2.5434 | 317082.7524 | -4.9746 | 317114.2729 | 2.4311 | 0.0031 | 0.4988 |
| HMR_5498 | Transport, extracellular | 41.6312 | 316419.7349 | 38.9555 | 316591.2473 | 2.6757 | 0.0034 | 0.4987 |
| HMR_5499 | Transport, extracellular | 43.1349 | 316345.4654 | 39.8539 | 316549.6589 | 3.281 | 0.0041 | 0.4984 |
| HMR_5500 | Transport, extracellular | -56.4806 | 314445.3567 | -57.8912 | 314357.4393 | 1.4106 | 0.0018 | 0.4993 |
| HMR_5501 | Transport, extracellular | 41.9887 | 315567.3265 | 42.8195 | 315537.6855 | 0.8308 | -0.001 | 0.5004 |
| HMR_5502 | Transport, extracellular | -42.3669 | 315548.9854 | -43.1825 | 315519.7948 | 0.8157 | 0.001 | 0.4996 |
| HMR_5503 | Transport, extracellular | 42.8624 | 315525.0116 | 43.6693 | 315495.724 | 0.8069 | -0.001 | 0.5004 |
| HMR_5504 | Transport, extracellular | 89.7556 | 311173.5591 | 87.0788 | 311446.4883 | 2.6768 | 0.0034 | 0.4986 |
| HMR_5505 | Transport, extracellular | -78.6714 | 312576.3298 | -79.1741 | 312542.6866 | 0.5027 | 0.0006 | 0.4997 |
| HMR_5506 | Transport, extracellular | 1.2857 | 316298.2246 | -6.5342 | 316286.5225 | 7.82 | 0.0098 | 0.4961 |
| HMR_5507 | Transport, extracellular | 6.835 | 316266.9655 | 5.4372 | 316282.0029 | 1.3979 | 0.0018 | 0.4993 |
| HMR_5508 | Transport, extracellular | 60.8669 | 314446.1504 | 42.2789 | 315564.288 | 18.588 | 0.0234 | 0.4907 |
| HMR_5509 | Transport, extracellular | -3.9226 | 316286.5093 | -2.734 | 316296.0907 | 1.1886 | -0.0015 | 0.5006 |
| HMR_5510 | Transport, extracellular | -40.436 | 315641.8127 | -41.3692 | 315608.6634 | 0.9332 | 0.0012 | 0.4995 |
| HMR_5511 | Transport, extracellular | -41.946 | 315569.4093 | -42.271 | 315564.6473 | 0.325 | 0.0004 | 0.4998 |
| HMR_5512 | Transport, extracellular | -14.4832 | 316405.2059 | -15.0653 | 316408.0008 | 0.5822 | 0.0007 | 0.4997 |
| HMR_5513 | Transport, extracellular | -14.1038 | 316411.2767 | -14.7011 | 316414.1233 | 0.5973 | 0.0008 | 0.4997 |
| HMR_5514 | Transport, extracellular | -13.6064 | 316419.0921 | -14.2126 | 316422.1672 | 0.6062 | 0.0008 | 0.4997 |
| HMR_5515 | Transport, extracellular | 144.9041 | 303755.5521 | 143.6571 | 303970.0668 | 1.247 | 0.0016 | 0.4994 |
| HMR_5516 | Transport, extracellular | -22.3599 | 315769.6645 | -21.4503 | 315805.7668 | 0.9096 | -0.0011 | 0.5005 |
| HMR_5517 | Transport, extracellular | -57.7619 | 314347.2652 | -51.3976 | 314757.1673 | 6.3643 | -0.008 | 0.5032 |
| HMR_5518 | Transport, extracellular | 63.2717 | 313961.6403 | 63.2927 | 313967.6073 | 0.021 | 0.00E+00 | 0.5 |
| HMR_5519 | Transport, extracellular | 4.4997 | 316500.8161 | -15.6079 | 316398.8484 | 20.1076 | 0.0253 | 0.4899 |
| HMR_5520 | Transport, extracellular | -60.3813 | 314166.7245 | -60.6102 | 314158.2518 | 0.2289 | 0.0003 | 0.4999 |
| HMR_5521 | Transport, extracellular | 16.0407 | 316379.4358 | 16.5203 | 316383.0745 | 0.4796 | -0.0006 | 0.5002 |
| HMR_5522 | Transport, extracellular | 14.526 | 316404.5309 | 15.6158 | 316398.723 | 1.0898 | -0.0014 | 0.5005 |
| HMR_5523 | Transport, extracellular | -0.379 | 316877.4112 | -0.3638 | 316897.5919 | 0.0152 | 0.00E+00 | 0.5 |
| HMR_5524 | Transport, extracellular | 0.8757 | 316877.0668 | 0.8518 | 316897.3072 | 0.0239 | 0.00E+00 | 0.5 |
| HMR_5525 | Transport, extracellular | 130.7464 | 306260.671 | 128.9232 | 306546.0641 | 1.8232 | 0.0023 | 0.4991 |
| HMR_5526 | Transport, extracellular | -36.787 | 315641.9995 | -36.4605 | 315676.6065 | 0.3266 | -0.0004 | 0.5002 |
| HMR_5527 | Transport, extracellular | 43.2706 | 315491.8921 | 36.3157 | 315828.4307 | 6.9549 | 0.0088 | 0.4965 |
| HMR_5528 | Transport, extracellular | 48.7898 | 315195.9847 | 48.2294 | 315240.9103 | 0.5604 | 0.0007 | 0.4997 |
| HMR_5529 | Transport, extracellular | 18.9488 | 316661.0868 | -0.5418 | 316897.8854 | 19.4906 | 0.0245 | 0.4902 |
| HMR_5530 | Transport, extracellular | 45.8942 | 315353.8868 | 45.5418 | 315386.0572 | 0.3524 | 0.0004 | 0.4998 |
| HMR_5531 | Transport, extracellular | 1.5558 | 316877.4649 | 1.4533 | 316897.6914 | 0.1025 | 0.0001 | 0.4999 |
| HMR_5532 | Transport, extracellular | 0.0428 | 316877.6214 | 0.5497 | 316897.8516 | 0.5069 | -0.0006 | 0.5003 |
| HMR_5533 | Transport, extracellular | 0.4967 | 316877.5908 | 0.4879 | 316897.8833 | 0.0088 | 0.00E+00 | 0.5 |
| HMR_5534 | Transport, extracellular | 131.1135 | 306206.2202 | 129.2761 | 306494.5378 | 1.8375 | 0.0023 | 0.4991 |

| | | | | | | | | |
|---|---|---|---|---|---|---|---|---|
| HMR_5535 | Transport, extracellular | -36.4083 | 315657.7664 | -36.0968 | 315691.5539 | 0.3115 | -0.0004 | 0.5002 |
| HMR_5536 | Transport, extracellular | -43.6487 | 315472.9818 | -36.6791 | 315813.1261 | 6.9696 | -0.0088 | 0.5035 |
| HMR_5537 | Transport, extracellular | 49.1675 | 315174.8109 | 48.5921 | 315220.7697 | 0.5754 | 0.0007 | 0.4997 |
| HMR_5538 | Transport, extracellular | 18.5701 | 316669.2064 | -0.9057 | 316897.5011 | 19.4757 | 0.0245 | 0.4902 |
| HMR_5539 | Transport, extracellular | -46.2721 | 315333.9646 | -45.9047 | 315367.0342 | 0.3674 | -0.0005 | 0.5002 |
| HMR_5540 | Transport, extracellular | 1.9347 | 316876.6602 | 1.8171 | 316896.982 | 0.1176 | 0.0001 | 0.4999 |
| HMR_5541 | Transport, extracellular | 0.4218 | 316877.451 | 0.9135 | 316897.489 | 0.4918 | -0.0006 | 0.5002 |
| HMR_5542 | Transport, extracellular | 131.5946 | 306134.7608 | 129.749 | 306425.3626 | 1.8456 | 0.0024 | 0.4991 |
| HMR_5543 | Transport, extracellular | -35.9118 | 315678.2867 | -35.6091 | 315711.5509 | 0.3027 | -0.0004 | 0.5002 |
| HMR_5544 | Transport, extracellular | 44.1441 | 315448.1767 | 37.1663 | 315792.6188 | 6.9778 | 0.0088 | 0.4965 |
| HMR_5545 | Transport, extracellular | 49.6624 | 315146.7778 | 49.0783 | 315193.6191 | 0.5841 | 0.0007 | 0.4997 |
| HMR_5546 | Transport, extracellular | 18.0736 | 316679.595 | -1.3936 | 316896.9122 | 19.4672 | 0.0245 | 0.4902 |
| HMR_5547 | Transport, extracellular | 46.7673 | 315307.633 | 46.3912 | 315341.4385 | 0.3761 | 0.0005 | 0.4998 |
| HMR_5548 | Transport, extracellular | 2.4314 | 316875.3925 | 2.305 | 316895.7999 | 0.1264 | 0.0002 | 0.4999 |
| HMR_5549 | Transport, extracellular | 0.9185 | 316877.0648 | 1.4015 | 316896.8704 | 0.483 | -0.0006 | 0.5002 |
| HMR_5550 | Transport, extracellular | 460.3752 | 81334.0927 | 460.8877 | 81359.0529 | 0.5126 | -0.0013 | 0.5005 |
| HMR_5551 | Transport, extracellular | 88.5168 | 311231.4169 | 93.5416 | 310732.8533 | 5.0248 | -0.0064 | 0.5025 |
| HMR_5552 | Transport, extracellular | -83.0396 | 311758.289 | -81.7345 | 311893.6556 | 1.3051 | -0.0017 | 0.5007 |
| HMR_5553 | Transport, extracellular | 149.0114 | 303368.4709 | 128.3977 | 306622.8237 | 20.6137 | 0.0264 | 0.4895 |
| HMR_5554 | Transport, extracellular | 85.9163 | 311484.1958 | 84.4057 | 311643.3922 | 1.5106 | 0.0019 | 0.4992 |
| HMR_5555 | Transport, extracellular | 129.2382 | 306483.6361 | 127.5129 | 306751.8246 | 1.7253 | 0.0022 | 0.4991 |
| HMR_5556 | Transport, extracellular | 130.705 | 306266.8689 | 128.39 | 306623.9634 | 2.315 | 0.003 | 0.4988 |
| HMR_5557 | Transport, extracellular | -79.9454 | 312451.0345 | -72.7127 | 313100.1469 | 7.2328 | -0.0091 | 0.5036 |
| HMR_5558 | Transport, extracellular | 85.4232 | 311930.1739 | 84.5452 | 312029.4397 | 0.878 | 0.0011 | 0.4996 |
| HMR_5559 | Transport, extracellular | -17.8239 | 316222.0279 | -37.0018 | 315654.1027 | 19.1779 | 0.0241 | 0.4904 |
| HMR_5560 | Transport, extracellular | -82.5502 | 312205.9106 | -81.8781 | 312283.1014 | 0.672 | -0.0009 | 0.5003 |
| HMR_5561 | Transport, extracellular | 38.3415 | 315576.6156 | 37.9125 | 315616.1119 | 0.4289 | 0.0005 | 0.4998 |
| HMR_5562 | Transport, extracellular | 36.8299 | 315640.2743 | 37.0098 | 315653.8245 | 0.1799 | -0.0002 | 0.5001 |
| HMR_5563 | Transport, extracellular | 5.5498 | 316260.9118 | 11.9711 | 316212.3222 | 6.4213 | -0.0081 | 0.5032 |
| HMR_5564 | Transport, extracellular | 62.1437 | 314347.0571 | 35.7745 | 315851.0092 | 26.3693 | 0.0332 | 0.4868 |
| HMR_5565 | Transport, extracellular | -2.6371 | 316276.1956 | -9.2684 | 316246.5814 | 6.6314 | 0.0083 | 0.4967 |
| HMR_5566 | Transport, extracellular | -41.7181 | 315568.4455 | -34.8639 | 315888.5692 | 6.8542 | -0.0086 | 0.5034 |
| HMR_5567 | Transport, extracellular | -43.2278 | 315494.0669 | -35.7665 | 315851.3148 | 7.4613 | -0.0094 | 0.5037 |
| HMR_5568 | Transport, extracellular | 67.6401 | 313932.5443 | 47.6893 | 315270.7618 | 19.9508 | 0.0252 | 0.49 |
| HMR_5569 | Transport, extracellular | 2.9128 | 316269.3058 | 2.7034 | 316278.7109 | 0.2094 | 0.0003 | 0.4999 |
| HMR_5570 | Transport, extracellular | 47.2388 | 315282.4464 | 46.7804 | 315320.7483 | 0.4584 | 0.0006 | 0.4998 |
| HMR_5571 | Transport, extracellular | 48.7471 | 315198.4301 | 47.6814 | 315271.181 | 1.0657 | 0.0013 | 0.4995 |
| HMR_5572 | Transport, extracellular | 64.7572 | 314151.5395 | 45.0014 | 315414.2569 | 19.7558 | 0.0249 | 0.4901 |
| HMR_5573 | Transport, extracellular | 20.5032 | 316627.3925 | 0.9114 | 316898.7561 | 19.5917 | 0.0246 | 0.4902 |
| HMR_5574 | Transport, extracellular | 18.9915 | 316660.2388 | 0.0079 | 316898.2777 | 18.9836 | 0.0238 | 0.4905 |
| HMR_5575 | Transport, extracellular | -44.3423 | 315435.0988 | -44.0921 | 315461.4219 | 0.2502 | -0.0003 | 0.5001 |
| HMR_5576 | Transport, extracellular | -45.8514 | 315356.2104 | -44.9935 | 315414.7237 | 0.8579 | -0.0011 | 0.5004 |
| HMR_5577 | Transport, extracellular | 1.5129 | 316877.6228 | 0.9035 | 316898.8424 | 0.6094 | 0.0008 | 0.4997 |
| HMR_5578 | Transport, extracellular | 220.9805 | 234139.7307 | 220.4437 | 233559.4151 | 0.5368 | 0.0008 | 0.4997 |

| | | | | | | | | |
|---|---|---|---|---|---|---|---|---|
| HMR_5579 | Transport, extracellular | 221.3369 | 227913.6841 | 223.5967 | 226161.7675 | 2.2597 | -0.0034 | 0.5013 |
| HMR_5580 | Transport, extracellular | -311.0263 | 224449.1999 | -337.0135 | 215801.4389 | 25.9871 | 0.0392 | 0.4844 |
| HMR_5581 | Transport, extracellular | -218.7998 | 228350.6971 | -222.0411 | 226419.1973 | 3.2413 | 0.0048 | 0.4981 |
| HMR_5582 | Transport, extracellular | 391.2473 | 184764.9792 | 392.0532 | 183748.2863 | 0.8059 | -0.0013 | 0.5005 |
| HMR_5583 | Transport, extracellular | 453.0517 | 173205.0829 | 470.8976 | 165828.2687 | 17.8459 | -0.0306 | 0.5122 |
| HMR_5584 | Transport, extracellular | 389.541 | 185264.6688 | 391.0137 | 184044.3226 | 1.4727 | -0.0024 | 0.501 |
| HMR_5585 | Transport, extracellular | -162.8882 | 230300.9086 | -199.6718 | 223590.5831 | 36.7836 | 0.0546 | 0.4782 |
| HMR_5586 | Transport, extracellular | 165.6528 | 230085.9383 | 201.2869 | 223416.6342 | 35.6341 | -0.0529 | 0.5211 |
| HMR_5587 | Transport, extracellular | 8.4943 | 73.2413 | 8.1342 | 67.1753 | 0.3601 | 0.0304 | 0.4879 |
| HMR_5588 | Transport, extracellular | 8.4888 | 73.143 | 8.1298 | 67.0988 | 0.359 | 0.0303 | 0.4879 |
| HMR_5589 | Transport, extracellular | 8.4885 | 73.1406 | 8.1302 | 67.1074 | 0.3583 | 0.0303 | 0.4879 |
| HMR_5590 | Transport, extracellular | 8.4772 | 72.9431 | 8.1196 | 66.9298 | 0.3577 | 0.0302 | 0.4879 |
| HMR_5591 | Transport, extracellular | 8.4801 | 72.9931 | 8.1223 | 66.9754 | 0.3578 | 0.0302 | 0.4879 |
| HMR_5592 | Transport, extracellular | 8.4801 | 72.9918 | 8.1223 | 66.9743 | 0.3578 | 0.0302 | 0.4879 |
| HMR_5593 | Transport, extracellular | 8.48 | 72.9901 | 8.1222 | 66.9728 | 0.3578 | 0.0302 | 0.4879 |
| HMR_5594 | Transport, extracellular | 8.5065 | 73.4553 | 8.1462 | 67.3763 | 0.3603 | 0.0304 | 0.4879 |
| HMR_5595 | Transport, extracellular | 8.4728 | 72.8657 | 8.1157 | 66.8644 | 0.3571 | 0.0302 | 0.4879 |
| HMR_5596 | Transport, extracellular | 8.4888 | 73.1445 | 8.129 | 67.0872 | 0.3598 | 0.0304 | 0.4879 |
| HMR_5597 | Transport, extracellular | 8.4899 | 73.164 | 8.1312 | 67.124 | 0.3587 | 0.0303 | 0.4879 |
| HMR_5598 | Transport, extracellular | 8.4763 | 72.928 | 8.1224 | 66.9771 | 0.3539 | 0.0299 | 0.4881 |
| HMR_5599 | Transport, extracellular | 8.4893 | 73.1537 | 8.1307 | 67.1156 | 0.3586 | 0.0303 | 0.4879 |
| HMR_5600 | Transport, extracellular | 8.4804 | 72.9986 | 8.1226 | 66.9799 | 0.3579 | 0.0302 | 0.4879 |
| HMR_5601 | Transport, extracellular | 8.4801 | 72.9933 | 8.1224 | 66.9771 | 0.3577 | 0.0302 | 0.4879 |
| HMR_5602 | Transport, extracellular | 8.5213 | 73.7135 | 8.1602 | 67.6108 | 0.3611 | 0.0304 | 0.4879 |
| HMR_5603 | Transport, extracellular | 8.5022 | 73.3793 | 8.1427 | 67.3167 | 0.3595 | 0.0303 | 0.4879 |
| HMR_5604 | Transport, extracellular | 8.5019 | 73.376 | 8.1431 | 67.3246 | 0.3588 | 0.0303 | 0.4879 |
| HMR_5605 | Transport, extracellular | 8.4906 | 73.1776 | 8.1325 | 67.1461 | 0.3582 | 0.0302 | 0.4879 |
| HMR_5606 | Transport, extracellular | 8.4935 | 73.2278 | 8.1353 | 67.192 | 0.3583 | 0.0302 | 0.4879 |
| HMR_5607 | Transport, extracellular | 8.4935 | 73.2265 | 8.1352 | 67.1909 | 0.3583 | 0.0302 | 0.4879 |
| HMR_5608 | Transport, extracellular | 8.4934 | 73.2247 | 8.1351 | 67.1894 | 0.3583 | 0.0302 | 0.4879 |
| HMR_5609 | Transport, extracellular | 8.52 | 73.6923 | 8.1592 | 67.5949 | 0.3608 | 0.0304 | 0.4879 |
| HMR_5610 | Transport, extracellular | 8.4862 | 73.0998 | 8.1286 | 67.0804 | 0.3576 | 0.0302 | 0.488 |
| HMR_5611 | Transport, extracellular | 8.5022 | 73.3799 | 8.1419 | 67.3043 | 0.3603 | 0.0304 | 0.4879 |
| HMR_5612 | Transport, extracellular | 8.5033 | 73.3995 | 8.1441 | 67.3413 | 0.3592 | 0.0303 | 0.4879 |
| HMR_5613 | Transport, extracellular | 8.4897 | 73.1623 | 8.1354 | 67.1937 | 0.3544 | 0.0299 | 0.4881 |
| HMR_5614 | Transport, extracellular | 8.5027 | 73.3892 | 8.1436 | 67.3328 | 0.3591 | 0.0303 | 0.4879 |
| HMR_5615 | Transport, extracellular | 8.4939 | 73.2333 | 8.1355 | 67.1965 | 0.3583 | 0.0302 | 0.4879 |
| HMR_5616 | Transport, extracellular | 8.4936 | 73.228 | 8.1354 | 67.1937 | 0.3582 | 0.0302 | 0.4879 |
| HMR_5617 | Transport, extracellular | 8.5269 | 73.8097 | 8.1647 | 67.6854 | 0.3621 | 0.0304 | 0.4879 |
| HMR_5618 | Transport, extracellular | 8.5133 | 73.5738 | 8.1517 | 67.4678 | 0.3617 | 0.0305 | 0.4879 |
| HMR_5619 | Transport, extracellular | 8.5075 | 73.4724 | 8.1476 | 67.3995 | 0.3599 | 0.0303 | 0.4879 |
| HMR_5620 | Transport, extracellular | 8.4962 | 73.2735 | 8.137 | 67.2206 | 0.3592 | 0.0303 | 0.4879 |
| HMR_5621 | Transport, extracellular | 8.4991 | 73.3238 | 8.1397 | 67.2666 | 0.3594 | 0.0303 | 0.4879 |
| HMR_5622 | Transport, extracellular | 8.499 | 73.3225 | 8.1397 | 67.2655 | 0.3593 | 0.0303 | 0.4879 |

| | | | | | | | | |
|---|---|---|---|---|---|---|---|---|
| HMR_5623 | Transport, extracellular | 8.4989 | 73.3208 | 8.1396 | 67.264 | 0.3593 | 0.0303 | 0.4879 |
| HMR_5624 | Transport, extracellular | 8.5256 | 73.7893 | 8.1637 | 67.6702 | 0.3619 | 0.0304 | 0.4879 |
| HMR_5625 | Transport, extracellular | 8.4917 | 73.1955 | 8.133 | 67.1548 | 0.3587 | 0.0303 | 0.4879 |
| HMR_5626 | Transport, extracellular | 8.5078 | 73.4763 | 8.1464 | 67.3791 | 0.3614 | 0.0304 | 0.4879 |
| HMR_5627 | Transport, extracellular | 8.5089 | 73.4959 | 8.1486 | 67.4161 | 0.3603 | 0.0303 | 0.4879 |
| HMR_5628 | Transport, extracellular | 8.4953 | 73.2582 | 8.1398 | 67.2683 | 0.3554 | 0.03 | 0.488 |
| HMR_5629 | Transport, extracellular | 8.5083 | 73.4856 | 8.1481 | 67.4077 | 0.3602 | 0.0303 | 0.4879 |
| HMR_5630 | Transport, extracellular | 8.4994 | 73.3293 | 8.14 | 67.2711 | 0.3594 | 0.0303 | 0.4879 |
| HMR_5631 | Transport, extracellular | 8.4991 | 73.324 | 8.1398 | 67.2683 | 0.3593 | 0.0303 | 0.4879 |
| HMR_5632 | Transport, extracellular | 8.5271 | 73.8158 | 8.1643 | 67.6799 | 0.3628 | 0.0305 | 0.4878 |
| HMR_5633 | Transport, extracellular | 8.5136 | 73.5789 | 8.1513 | 67.4615 | 0.3624 | 0.0305 | 0.4878 |
| HMR_5634 | Transport, extracellular | 8.5081 | 73.4808 | 8.1468 | 67.3853 | 0.3613 | 0.0304 | 0.4879 |
| HMR_5635 | Transport, extracellular | 8.4965 | 73.2786 | 8.1365 | 67.2143 | 0.3599 | 0.0304 | 0.4879 |
| HMR_5636 | Transport, extracellular | 8.4994 | 73.3289 | 8.1393 | 67.2603 | 0.36 | 0.0304 | 0.4879 |
| HMR_5637 | Transport, extracellular | 8.4993 | 73.3276 | 8.1392 | 67.2591 | 0.36 | 0.0304 | 0.4879 |
| HMR_5638 | Transport, extracellular | 8.4992 | 73.3259 | 8.1392 | 67.2576 | 0.36 | 0.0304 | 0.4879 |
| HMR_5639 | Transport, extracellular | 8.5258 | 73.7943 | 8.1633 | 67.6637 | 0.3626 | 0.0305 | 0.4878 |
| HMR_5640 | Transport, extracellular | 8.492 | 73.2006 | 8.1326 | 67.1485 | 0.3594 | 0.0303 | 0.4879 |
| HMR_5641 | Transport, extracellular | 8.508 | 73.4813 | 8.146 | 67.3727 | 0.362 | 0.0305 | 0.4878 |
| HMR_5642 | Transport, extracellular | 8.5091 | 73.501 | 8.1482 | 67.4098 | 0.361 | 0.0304 | 0.4879 |
| HMR_5643 | Transport, extracellular | 8.4955 | 73.2633 | 8.1394 | 67.262 | 0.3561 | 0.03 | 0.488 |
| HMR_5644 | Transport, extracellular | 8.5086 | 73.4906 | 8.1477 | 67.4013 | 0.3609 | 0.0304 | 0.4879 |
| HMR_5645 | Transport, extracellular | 8.4997 | 73.3344 | 8.1396 | 67.2648 | 0.3601 | 0.0304 | 0.4879 |
| HMR_5646 | Transport, extracellular | 8.4994 | 73.3291 | 8.1394 | 67.262 | 0.36 | 0.0304 | 0.4879 |
| HMR_5647 | Transport, extracellular | 8.5385 | 74.017 | 8.175 | 67.8607 | 0.3635 | 0.0305 | 0.4878 |
| HMR_5648 | Transport, extracellular | 8.525 | 73.7792 | 8.1619 | 67.6415 | 0.363 | 0.0305 | 0.4878 |
| HMR_5649 | Transport, extracellular | 8.5194 | 73.6806 | 8.1574 | 67.5649 | 0.362 | 0.0305 | 0.4879 |
| HMR_5650 | Transport, extracellular | 8.5191 | 73.6772 | 8.1579 | 67.5728 | 0.3613 | 0.0304 | 0.4879 |
| HMR_5651 | Transport, extracellular | 8.5107 | 73.5281 | 8.15 | 67.4394 | 0.3607 | 0.0304 | 0.4879 |
| HMR_5652 | Transport, extracellular | 8.5106 | 73.5267 | 8.1499 | 67.4382 | 0.3607 | 0.0304 | 0.4879 |
| HMR_5653 | Transport, extracellular | 8.5105 | 73.525 | 8.1498 | 67.4367 | 0.3607 | 0.0304 | 0.4879 |
| HMR_5654 | Transport, extracellular | 8.5372 | 73.9955 | 8.174 | 67.8445 | 0.3633 | 0.0305 | 0.4878 |
| HMR_5655 | Transport, extracellular | 8.5033 | 73.3992 | 8.1432 | 67.3271 | 0.36 | 0.0304 | 0.4879 |
| HMR_5656 | Transport, extracellular | 8.5194 | 73.6811 | 8.1567 | 67.5523 | 0.3627 | 0.0305 | 0.4878 |
| HMR_5657 | Transport, extracellular | 8.5205 | 73.7009 | 8.1589 | 67.5895 | 0.3616 | 0.0304 | 0.4879 |
| HMR_5658 | Transport, extracellular | 8.5069 | 73.4622 | 8.1501 | 67.4411 | 0.3568 | 0.0301 | 0.488 |
| HMR_5659 | Transport, extracellular | 8.5199 | 73.6905 | 8.1584 | 67.581 | 0.3616 | 0.0304 | 0.4879 |
| HMR_5660 | Transport, extracellular | 8.511 | 73.5336 | 8.1502 | 67.4439 | 0.3608 | 0.0304 | 0.4879 |
| HMR_5661 | Transport, extracellular | 8.5107 | 73.5282 | 8.1501 | 67.4411 | 0.3606 | 0.0304 | 0.4879 |
| HMR_5662 | Transport, extracellular | 8.5356 | 73.9653 | 8.1722 | 67.8135 | 0.3634 | 0.0305 | 0.4878 |
| HMR_5663 | Transport, extracellular | 8.5221 | 73.7277 | 8.1592 | 67.5945 | 0.3629 | 0.0305 | 0.4878 |
| HMR_5664 | Transport, extracellular | 8.5165 | 73.6292 | 8.1547 | 67.518 | 0.3618 | 0.0305 | 0.4879 |
| HMR_5665 | Transport, extracellular | 8.5162 | 73.6259 | 8.1551 | 67.5259 | 0.3611 | 0.0304 | 0.4879 |
| HMR_5666 | Transport, extracellular | 8.5049 | 73.4264 | 8.1444 | 67.3465 | 0.3605 | 0.0304 | 0.4879 |

| | | | | | | | | |
|---|---|---|---|---|---|---|---|---|
| HMR_5667 | Transport, extracellular | 8.5077 | 73.4755 | 8.1471 | 67.3915 | 0.3606 | 0.0304 | 0.4879 |
| HMR_5668 | Transport, extracellular | 8.5076 | 73.4738 | 8.147 | 67.39 | 0.3606 | 0.0304 | 0.4879 |
| HMR_5669 | Transport, extracellular | 8.5343 | 73.9438 | 8.1712 | 67.7974 | 0.3631 | 0.0305 | 0.4878 |
| HMR_5670 | Transport, extracellular | 8.5004 | 73.3481 | 8.1405 | 67.2805 | 0.3599 | 0.0304 | 0.4879 |
| HMR_5671 | Transport, extracellular | 8.5165 | 73.6298 | 8.1539 | 67.5055 | 0.3626 | 0.0305 | 0.4878 |
| HMR_5672 | Transport, extracellular | 8.5176 | 73.6495 | 8.1561 | 67.5426 | 0.3615 | 0.0304 | 0.4879 |
| HMR_5673 | Transport, extracellular | 8.504 | 73.411 | 8.1473 | 67.3943 | 0.3567 | 0.0301 | 0.488 |
| HMR_5674 | Transport, extracellular | 8.517 | 73.6391 | 8.1556 | 67.5341 | 0.3614 | 0.0304 | 0.4879 |
| HMR_5675 | Transport, extracellular | 8.5081 | 73.4824 | 8.1475 | 67.3972 | 0.3606 | 0.0304 | 0.4879 |
| HMR_5676 | Transport, extracellular | 8.5078 | 73.477 | 8.1473 | 67.3943 | 0.3605 | 0.0304 | 0.4879 |
| HMR_5677 | Transport, extracellular | 8.5357 | 73.9666 | 8.1723 | 67.8147 | 0.3634 | 0.0305 | 0.4878 |
| HMR_5678 | Transport, extracellular | 8.5221 | 73.729 | 8.1592 | 67.5956 | 0.3629 | 0.0305 | 0.4878 |
| HMR_5679 | Transport, extracellular | 8.5166 | 73.6306 | 8.1547 | 67.5191 | 0.3618 | 0.0305 | 0.4879 |
| HMR_5680 | Transport, extracellular | 8.5163 | 73.6272 | 8.1551 | 67.527 | 0.3611 | 0.0304 | 0.4879 |
| HMR_5681 | Transport, extracellular | 8.5049 | 73.4277 | 8.1445 | 67.3477 | 0.3605 | 0.0304 | 0.4879 |
| HMR_5682 | Transport, extracellular | 8.5079 | 73.4782 | 8.1473 | 67.3937 | 0.3606 | 0.0304 | 0.4879 |
| HMR_5683 | Transport, extracellular | 8.5077 | 73.4751 | 8.1471 | 67.3911 | 0.3606 | 0.0304 | 0.4879 |
| HMR_5684 | Transport, extracellular | 8.5344 | 73.9452 | 8.1712 | 67.7985 | 0.3632 | 0.0305 | 0.4878 |
| HMR_5685 | Transport, extracellular | 8.5005 | 73.3495 | 8.1406 | 67.2816 | 0.3599 | 0.0304 | 0.4879 |
| HMR_5686 | Transport, extracellular | 8.5165 | 73.6311 | 8.1539 | 67.5066 | 0.3626 | 0.0305 | 0.4878 |
| HMR_5687 | Transport, extracellular | 8.5177 | 73.6508 | 8.1561 | 67.5437 | 0.3615 | 0.0304 | 0.4879 |
| HMR_5688 | Transport, extracellular | 8.504 | 73.4124 | 8.1474 | 67.3954 | 0.3567 | 0.0301 | 0.488 |
| HMR_5689 | Transport, extracellular | 8.5171 | 73.6405 | 8.1557 | 67.5352 | 0.3614 | 0.0304 | 0.4879 |
| HMR_5690 | Transport, extracellular | 8.5082 | 73.4837 | 8.1475 | 67.3983 | 0.3606 | 0.0304 | 0.4879 |
| HMR_5691 | Transport, extracellular | 8.5079 | 73.4783 | 8.1474 | 67.3955 | 0.3605 | 0.0304 | 0.4879 |
| HMR_5692 | Transport, extracellular | 8.5358 | 73.9684 | 8.1724 | 67.8162 | 0.3634 | 0.0305 | 0.4878 |
| HMR_5693 | Transport, extracellular | 8.5222 | 73.7308 | 8.1593 | 67.5972 | 0.3629 | 0.0305 | 0.4878 |
| HMR_5694 | Transport, extracellular | 8.5167 | 73.6323 | 8.1548 | 67.5206 | 0.3618 | 0.0305 | 0.4879 |
| HMR_5695 | Transport, extracellular | 8.5164 | 73.6289 | 8.1552 | 67.5286 | 0.3611 | 0.0304 | 0.4879 |
| HMR_5696 | Transport, extracellular | 8.505 | 73.4295 | 8.1446 | 67.3492 | 0.3605 | 0.0304 | 0.4879 |
| HMR_5697 | Transport, extracellular | 8.508 | 73.4799 | 8.1473 | 67.3952 | 0.3606 | 0.0304 | 0.4879 |
| HMR_5698 | Transport, extracellular | 8.5079 | 73.4786 | 8.1473 | 67.3941 | 0.3606 | 0.0304 | 0.4879 |
| HMR_5699 | Transport, extracellular | 8.5345 | 73.9469 | 8.1713 | 67.8 | 0.3632 | 0.0305 | 0.4878 |
| HMR_5700 | Transport, extracellular | 8.5006 | 73.3512 | 8.1406 | 67.2831 | 0.3599 | 0.0304 | 0.4879 |
| HMR_5701 | Transport, extracellular | 8.5166 | 73.6329 | 8.154 | 67.5081 | 0.3626 | 0.0305 | 0.4878 |
| HMR_5702 | Transport, extracellular | 8.5178 | 73.6526 | 8.1562 | 67.5452 | 0.3615 | 0.0304 | 0.4879 |
| HMR_5703 | Transport, extracellular | 8.5041 | 73.4141 | 8.1474 | 67.3969 | 0.3567 | 0.0301 | 0.488 |
| HMR_5704 | Transport, extracellular | 8.5172 | 73.6422 | 8.1557 | 67.5367 | 0.3614 | 0.0304 | 0.4879 |
| HMR_5705 | Transport, extracellular | 8.5083 | 73.4855 | 8.1476 | 67.3998 | 0.3607 | 0.0304 | 0.4879 |
| HMR_5706 | Transport, extracellular | 8.508 | 73.4801 | 8.1474 | 67.397 | 0.3605 | 0.0304 | 0.4879 |
| HMR_5707 | Transport, extracellular | 8.509 | 73.5 | 8.1482 | 67.4102 | 0.3608 | 0.0304 | 0.4879 |
| HMR_5708 | Transport, extracellular | 8.4956 | 73.2647 | 8.1352 | 67.1932 | 0.3603 | 0.0304 | 0.4879 |
| HMR_5709 | Transport, extracellular | 8.49 | 73.1672 | 8.1308 | 67.1174 | 0.3593 | 0.0303 | 0.4879 |
| HMR_5710 | Transport, extracellular | 8.4898 | 73.1638 | 8.1312 | 67.1252 | 0.3586 | 0.0303 | 0.4879 |

| | | | | | | | | |
|---|---|---|---|---|---|---|---|---|
| HMR_5711 | Transport, extracellular | 8.4785 | 72.9663 | 8.1206 | 66.9475 | 0.3579 | 0.0303 | 0.4879 |
| HMR_5712 | Transport, extracellular | 8.4814 | 73.0163 | 8.1233 | 66.9932 | 0.3581 | 0.0303 | 0.4879 |
| HMR_5713 | Transport, extracellular | 8.4813 | 73.0149 | 8.1233 | 66.992 | 0.358 | 0.0303 | 0.4879 |
| HMR_5714 | Transport, extracellular | 8.4812 | 73.0132 | 8.1232 | 66.9906 | 0.358 | 0.0303 | 0.4879 |
| HMR_5715 | Transport, extracellular | 8.474 | 72.8898 | 8.1166 | 66.8829 | 0.3574 | 0.0302 | 0.4879 |
| HMR_5716 | Transport, extracellular | 8.49 | 73.1677 | 8.13 | 67.1049 | 0.36 | 0.0304 | 0.4879 |
| HMR_5717 | Transport, extracellular | 8.4911 | 73.1872 | 8.1322 | 67.1417 | 0.359 | 0.0303 | 0.4879 |
| HMR_5718 | Transport, extracellular | 8.4776 | 72.9511 | 8.1234 | 66.9948 | 0.3542 | 0.0299 | 0.4881 |
| HMR_5719 | Transport, extracellular | 8.4905 | 73.1769 | 8.1317 | 67.1333 | 0.3589 | 0.0303 | 0.4879 |
| HMR_5720 | Transport, extracellular | 8.4817 | 73.0218 | 8.1236 | 66.9977 | 0.3581 | 0.0303 | 0.4879 |
| HMR_5721 | Transport, extracellular | 8.4814 | 73.0164 | 8.1234 | 66.9949 | 0.358 | 0.0303 | 0.4879 |
| HMR_5722 | Transport, extracellular | 8.543 | 74.0966 | 8.179 | 67.9278 | 0.3641 | 0.0305 | 0.4878 |
| HMR_5723 | Transport, extracellular | 8.5295 | 73.8584 | 8.1659 | 67.7083 | 0.3636 | 0.0306 | 0.4878 |
| HMR_5724 | Transport, extracellular | 8.5239 | 73.7597 | 8.1614 | 67.6316 | 0.3625 | 0.0305 | 0.4878 |
| HMR_5725 | Transport, extracellular | 8.5236 | 73.7563 | 8.1618 | 67.6395 | 0.3618 | 0.0304 | 0.4879 |
| HMR_5726 | Transport, extracellular | 8.5122 | 73.5563 | 8.1511 | 67.4596 | 0.3611 | 0.0304 | 0.4879 |
| HMR_5727 | Transport, extracellular | 8.5152 | 73.6069 | 8.1539 | 67.5058 | 0.3613 | 0.0304 | 0.4879 |
| HMR_5728 | Transport, extracellular | 8.5151 | 73.6055 | 8.1538 | 67.5047 | 0.3613 | 0.0304 | 0.4879 |
| HMR_5729 | Transport, extracellular | 8.515 | 73.6038 | 8.1537 | 67.5032 | 0.3612 | 0.0304 | 0.4879 |
| HMR_5730 | Transport, extracellular | 8.5417 | 74.0761 | 8.1779 | 67.9125 | 0.3638 | 0.0305 | 0.4878 |
| HMR_5731 | Transport, extracellular | 8.5239 | 73.7602 | 8.1606 | 67.619 | 0.3633 | 0.0306 | 0.4878 |
| HMR_5732 | Transport, extracellular | 8.525 | 73.78 | 8.1628 | 67.6562 | 0.3622 | 0.0305 | 0.4879 |
| HMR_5733 | Transport, extracellular | 8.5113 | 73.5409 | 8.154 | 67.5075 | 0.3573 | 0.0301 | 0.488 |
| HMR_5734 | Transport, extracellular | 8.5244 | 73.7696 | 8.1623 | 67.6477 | 0.3621 | 0.0304 | 0.4879 |
| HMR_5735 | Transport, extracellular | 8.5155 | 73.6124 | 8.1542 | 67.5104 | 0.3613 | 0.0304 | 0.4879 |
| HMR_5736 | Transport, extracellular | 8.5152 | 73.607 | 8.154 | 67.5076 | 0.3612 | 0.0304 | 0.4879 |
| HMR_5737 | Transport, extracellular | 8.5269 | 73.8119 | 8.1655 | 67.7004 | 0.3614 | 0.0304 | 0.4879 |
| HMR_5738 | Transport, extracellular | 8.5133 | 73.5751 | 8.1524 | 67.482 | 0.3609 | 0.0304 | 0.4879 |
| HMR_5739 | Transport, extracellular | 8.5078 | 73.477 | 8.148 | 67.4057 | 0.3598 | 0.0303 | 0.4879 |
| HMR_5740 | Transport, extracellular | 8.5075 | 73.4736 | 8.1484 | 67.4135 | 0.3591 | 0.0303 | 0.4879 |
| HMR_5741 | Transport, extracellular | 8.4962 | 73.2747 | 8.1377 | 67.2346 | 0.3585 | 0.0302 | 0.4879 |
| HMR_5742 | Transport, extracellular | 8.4991 | 73.3251 | 8.1405 | 67.2806 | 0.3586 | 0.0302 | 0.4879 |
| HMR_5743 | Transport, extracellular | 8.499 | 73.3237 | 8.1404 | 67.2795 | 0.3586 | 0.0302 | 0.4879 |
| HMR_5744 | Transport, extracellular | 8.4989 | 73.322 | 8.1404 | 67.278 | 0.3586 | 0.0302 | 0.4879 |
| HMR_5745 | Transport, extracellular | 8.5256 | 73.7905 | 8.1645 | 67.6843 | 0.3611 | 0.0304 | 0.4879 |
| HMR_5746 | Transport, extracellular | 8.4917 | 73.1968 | 8.1338 | 67.1688 | 0.3579 | 0.0302 | 0.4879 |
| HMR_5747 | Transport, extracellular | 8.5089 | 73.4971 | 8.1494 | 67.4302 | 0.3595 | 0.0303 | 0.4879 |
| HMR_5748 | Transport, extracellular | 8.4953 | 73.2595 | 8.1406 | 67.2823 | 0.3547 | 0.0299 | 0.4881 |
| HMR_5749 | Transport, extracellular | 8.5083 | 73.4868 | 8.1489 | 67.4217 | 0.3594 | 0.0303 | 0.4879 |
| HMR_5750 | Transport, extracellular | 8.4994 | 73.3306 | 8.1408 | 67.2851 | 0.3586 | 0.0302 | 0.4879 |
| HMR_5751 | Transport, extracellular | 8.4991 | 73.3252 | 8.1406 | 67.2823 | 0.3585 | 0.0302 | 0.4879 |
| HMR_5752 | Transport, extracellular | 8.5257 | 73.7921 | 8.1633 | 67.6632 | 0.3625 | 0.0305 | 0.4878 |
| HMR_5753 | Transport, extracellular | 8.5122 | 73.5554 | 8.1502 | 67.4449 | 0.362 | 0.0305 | 0.4878 |
| HMR_5754 | Transport, extracellular | 8.5067 | 73.4573 | 8.1458 | 67.3687 | 0.3609 | 0.0304 | 0.4879 |

| | | | | | | | | |
|---|---|---|---|---|---|---|---|---|
| HMR_5755 | Transport, extracellular | 8.5064 | 73.4539 | 8.1462 | 67.3765 | 0.3602 | 0.0304 | 0.4879 |
| HMR_5756 | Transport, extracellular | 8.4951 | 73.2552 | 8.1355 | 67.1978 | 0.3595 | 0.0303 | 0.4879 |
| HMR_5757 | Transport, extracellular | 8.498 | 73.3055 | 8.1383 | 67.2437 | 0.3597 | 0.0303 | 0.4879 |
| HMR_5758 | Transport, extracellular | 8.4979 | 73.3041 | 8.1382 | 67.2426 | 0.3597 | 0.0303 | 0.4879 |
| HMR_5759 | Transport, extracellular | 8.4978 | 73.3024 | 8.1382 | 67.2411 | 0.3597 | 0.0303 | 0.4879 |
| HMR_5760 | Transport, extracellular | 8.5245 | 73.7707 | 8.1622 | 67.6471 | 0.3622 | 0.0305 | 0.4879 |
| HMR_5761 | Transport, extracellular | 8.4906 | 73.1772 | 8.1316 | 67.132 | 0.359 | 0.0303 | 0.4879 |
| HMR_5762 | Transport, extracellular | 8.5067 | 73.4578 | 8.145 | 67.3562 | 0.3617 | 0.0305 | 0.4878 |
| HMR_5763 | Transport, extracellular | 8.4942 | 73.2399 | 8.1384 | 67.2454 | 0.3558 | 0.03 | 0.488 |
| HMR_5764 | Transport, extracellular | 8.5072 | 73.4671 | 8.1467 | 67.3847 | 0.3605 | 0.0304 | 0.4879 |
| HMR_5765 | Transport, extracellular | 8.4983 | 73.311 | 8.1386 | 67.2483 | 0.3597 | 0.0303 | 0.4879 |
| HMR_5766 | Transport, extracellular | 8.498 | 73.3056 | 8.1384 | 67.2454 | 0.3596 | 0.0303 | 0.4879 |
| HMR_5767 | Transport, extracellular | 8.5394 | 74.0318 | 8.1721 | 67.8118 | 0.3673 | 0.0308 | 0.4877 |
| HMR_5768 | Transport, extracellular | 8.5259 | 73.7939 | 8.1591 | 67.5928 | 0.3668 | 0.0309 | 0.4877 |
| HMR_5769 | Transport, extracellular | 8.5203 | 73.6953 | 8.1546 | 67.5163 | 0.3658 | 0.0308 | 0.4877 |
| HMR_5770 | Transport, extracellular | 8.52 | 73.692 | 8.155 | 67.5242 | 0.3651 | 0.0307 | 0.4877 |
| HMR_5771 | Transport, extracellular | 8.5087 | 73.4922 | 8.1443 | 67.3448 | 0.3644 | 0.0307 | 0.4878 |
| HMR_5772 | Transport, extracellular | 8.5116 | 73.5428 | 8.1471 | 67.3909 | 0.3645 | 0.0307 | 0.4878 |
| HMR_5773 | Transport, extracellular | 8.5115 | 73.5414 | 8.147 | 67.3898 | 0.3645 | 0.0307 | 0.4878 |
| HMR_5774 | Transport, extracellular | 8.5114 | 73.5397 | 8.1469 | 67.3883 | 0.3645 | 0.0307 | 0.4878 |
| HMR_5775 | Transport, extracellular | 8.5382 | 74.0104 | 8.1711 | 67.7957 | 0.3671 | 0.0308 | 0.4877 |
| HMR_5776 | Transport, extracellular | 8.5042 | 73.4139 | 8.1404 | 67.2788 | 0.3638 | 0.0307 | 0.4878 |
| HMR_5777 | Transport, extracellular | 8.5203 | 73.6959 | 8.1538 | 67.5038 | 0.3665 | 0.0308 | 0.4877 |
| HMR_5778 | Transport, extracellular | 8.5214 | 73.7157 | 8.156 | 67.5409 | 0.3654 | 0.0307 | 0.4877 |
| HMR_5779 | Transport, extracellular | 8.5208 | 73.7052 | 8.1555 | 67.5324 | 0.3653 | 0.0307 | 0.4877 |
| HMR_5780 | Transport, extracellular | 8.5119 | 73.5483 | 8.1474 | 67.3955 | 0.3646 | 0.0307 | 0.4878 |
| HMR_5781 | Transport, extracellular | 8.5116 | 73.5429 | 8.1472 | 67.3926 | 0.3644 | 0.0307 | 0.4878 |
| HMR_5782 | Transport, extracellular | 8.5263 | 73.8025 | 8.1638 | 67.6717 | 0.3626 | 0.0305 | 0.4878 |
| HMR_5783 | Transport, extracellular | 8.5128 | 73.5658 | 8.1507 | 67.4534 | 0.3621 | 0.0305 | 0.4878 |
| HMR_5784 | Transport, extracellular | 8.5073 | 73.4677 | 8.1463 | 67.3771 | 0.361 | 0.0304 | 0.4879 |
| HMR_5785 | Transport, extracellular | 8.507 | 73.4643 | 8.1467 | 67.385 | 0.3603 | 0.0304 | 0.4879 |
| HMR_5786 | Transport, extracellular | 8.4957 | 73.2655 | 8.136 | 67.2062 | 0.3596 | 0.0303 | 0.4879 |
| HMR_5787 | Transport, extracellular | 8.4986 | 73.3158 | 8.1388 | 67.2522 | 0.3598 | 0.0303 | 0.4879 |
| HMR_5788 | Transport, extracellular | 8.4985 | 73.3145 | 8.1387 | 67.251 | 0.3597 | 0.0303 | 0.4879 |
| HMR_5789 | Transport, extracellular | 8.4984 | 73.3127 | 8.1387 | 67.2495 | 0.3597 | 0.0303 | 0.4879 |
| HMR_5790 | Transport, extracellular | 8.5251 | 73.7811 | 8.1627 | 67.6556 | 0.3623 | 0.0305 | 0.4878 |
| HMR_5791 | Transport, extracellular | 8.4912 | 73.1875 | 8.1321 | 67.1404 | 0.3591 | 0.0303 | 0.4879 |
| HMR_5792 | Transport, extracellular | 8.5072 | 73.4682 | 8.1455 | 67.3646 | 0.3618 | 0.0305 | 0.4878 |
| HMR_5793 | Transport, extracellular | 8.5084 | 73.4878 | 8.1477 | 67.4016 | 0.3607 | 0.0304 | 0.4879 |
| HMR_5794 | Transport, extracellular | 8.4948 | 73.2502 | 8.1389 | 67.2538 | 0.3559 | 0.03 | 0.488 |
| HMR_5795 | Transport, extracellular | 8.4989 | 73.3213 | 8.1391 | 67.2567 | 0.3598 | 0.0303 | 0.4879 |
| HMR_5796 | Transport, extracellular | 8.4986 | 73.3159 | 8.1389 | 67.2539 | 0.3597 | 0.0303 | 0.4879 |
| HMR_5797 | Transport, extracellular | 8.5353 | 73.9597 | 8.1719 | 67.8089 | 0.3633 | 0.0305 | 0.4878 |
| HMR_5798 | Transport, extracellular | 8.5217 | 73.7221 | 8.1589 | 67.5899 | 0.3629 | 0.0305 | 0.4878 |

| | | | | | | | | |
|---|---|---|---|---|---|---|---|---|
| HMR_5799 | Transport, extracellular | 8.5162 | 73.6237 | 8.1544 | 67.5134 | 0.3618 | 0.0305 | 0.4879 |
| HMR_5800 | Transport, extracellular | 8.5159 | 73.6203 | 8.1548 | 67.5213 | 0.3611 | 0.0304 | 0.4879 |
| HMR_5801 | Transport, extracellular | 8.5046 | 73.4209 | 8.1441 | 67.342 | 0.3604 | 0.0304 | 0.4879 |
| HMR_5802 | Transport, extracellular | 8.5075 | 73.4713 | 8.1469 | 67.3881 | 0.3605 | 0.0304 | 0.4879 |
| HMR_5803 | Transport, extracellular | 8.5074 | 73.47 | 8.1469 | 67.3869 | 0.3605 | 0.0304 | 0.4879 |
| HMR_5804 | Transport, extracellular | 8.5073 | 73.4682 | 8.1468 | 67.3854 | 0.3605 | 0.0304 | 0.4879 |
| HMR_5805 | Transport, extracellular | 8.534 | 73.9382 | 8.1709 | 67.7928 | 0.3631 | 0.0305 | 0.4878 |
| HMR_5806 | Transport, extracellular | 8.5001 | 73.3426 | 8.1402 | 67.276 | 0.3599 | 0.0303 | 0.4879 |
| HMR_5807 | Transport, extracellular | 8.5162 | 73.6242 | 8.1536 | 67.5009 | 0.3626 | 0.0305 | 0.4878 |
| HMR_5808 | Transport, extracellular | 8.5173 | 73.6439 | 8.1558 | 67.538 | 0.3615 | 0.0304 | 0.4879 |
| HMR_5809 | Transport, extracellular | 8.5037 | 73.4055 | 8.147 | 67.3897 | 0.3566 | 0.0301 | 0.488 |
| HMR_5810 | Transport, extracellular | 8.5167 | 73.6335 | 8.1553 | 67.5295 | 0.3614 | 0.0304 | 0.4879 |
| HMR_5811 | Transport, extracellular | 8.5075 | 73.4715 | 8.147 | 67.3898 | 0.3604 | 0.0304 | 0.4879 |
| HMR_5812 | Transport, extracellular | 8.5356 | 73.9651 | 8.1721 | 67.8118 | 0.3635 | 0.0305 | 0.4878 |
| HMR_5813 | Transport, extracellular | 8.522 | 73.7275 | 8.1591 | 67.5928 | 0.363 | 0.0305 | 0.4878 |
| HMR_5814 | Transport, extracellular | 8.5165 | 73.6291 | 8.1546 | 67.5163 | 0.3619 | 0.0305 | 0.4878 |
| HMR_5815 | Transport, extracellular | 8.5162 | 73.6257 | 8.155 | 67.5242 | 0.3612 | 0.0304 | 0.4879 |
| HMR_5816 | Transport, extracellular | 8.5049 | 73.4262 | 8.1443 | 67.3448 | 0.3605 | 0.0304 | 0.4879 |
| HMR_5817 | Transport, extracellular | 8.5078 | 73.4767 | 8.1471 | 67.3909 | 0.3607 | 0.0304 | 0.4879 |
| HMR_5818 | Transport, extracellular | 8.5077 | 73.4753 | 8.147 | 67.3898 | 0.3607 | 0.0304 | 0.4879 |
| HMR_5819 | Transport, extracellular | 8.5076 | 73.4736 | 8.1469 | 67.3883 | 0.3607 | 0.0304 | 0.4879 |
| HMR_5820 | Transport, extracellular | 8.5343 | 73.9437 | 8.1711 | 67.7956 | 0.3632 | 0.0305 | 0.4878 |
| HMR_5821 | Transport, extracellular | 8.5004 | 73.348 | 8.1404 | 67.2788 | 0.36 | 0.0304 | 0.4879 |
| HMR_5822 | Transport, extracellular | 8.5165 | 73.6296 | 8.1538 | 67.5037 | 0.3627 | 0.0305 | 0.4878 |
| HMR_5823 | Transport, extracellular | 8.5176 | 73.6493 | 8.156 | 67.5409 | 0.3616 | 0.0304 | 0.4879 |
| HMR_5824 | Transport, extracellular | 8.504 | 73.4109 | 8.1472 | 67.3926 | 0.3568 | 0.0301 | 0.488 |
| HMR_5825 | Transport, extracellular | 8.517 | 73.6389 | 8.1555 | 67.5324 | 0.3615 | 0.0304 | 0.4879 |
| HMR_5826 | Transport, extracellular | 8.5081 | 73.4822 | 8.1474 | 67.3955 | 0.3607 | 0.0304 | 0.4879 |
| HMR_5827 | Transport, extracellular | 10.6339 | 116.4249 | 10.009 | 103.0391 | 0.6249 | 0.0422 | 0.4832 |
| HMR_5828 | Transport, extracellular | 10.6862 | 117.6291 | 10.055 | 104.0328 | 0.6312 | 0.0424 | 0.4831 |
| HMR_5829 | Transport, extracellular | 10.5837 | 115.3218 | 9.9645 | 102.1182 | 0.6193 | 0.042 | 0.4833 |
| HMR_5830 | Transport, extracellular | 10.5504 | 114.5457 | 9.9277 | 101.3276 | 0.6226 | 0.0424 | 0.4831 |
| HMR_5831 | Transport, extracellular | 10.5844 | 115.3372 | 9.9648 | 102.1265 | 0.6196 | 0.042 | 0.4832 |
| HMR_5832 | Transport, extracellular | 10.6549 | 116.9009 | 10.0286 | 103.4549 | 0.6263 | 0.0422 | 0.4832 |
| HMR_5833 | Transport, extracellular | 10.7075 | 118.1126 | 10.0749 | 104.4547 | 0.6327 | 0.0424 | 0.4831 |
| HMR_5834 | Transport, extracellular | 10.6046 | 115.791 | 9.9839 | 102.5284 | 0.6207 | 0.042 | 0.4832 |
| HMR_5835 | Transport, extracellular | 10.5711 | 115.0101 | 9.947 | 101.7329 | 0.6241 | 0.0424 | 0.4831 |
| HMR_5836 | Transport, extracellular | 10.6053 | 115.8065 | 9.9843 | 102.5367 | 0.621 | 0.042 | 0.4832 |
| HMR_5837 | Transport, extracellular | 10.6637 | 117.0954 | 10.0354 | 103.598 | 0.6282 | 0.0423 | 0.4831 |
| HMR_5838 | Transport, extracellular | 10.7163 | 118.3102 | 10.0817 | 104.5999 | 0.6346 | 0.0425 | 0.483 |
| HMR_5839 | Transport, extracellular | 10.6132 | 115.9827 | 9.9907 | 102.6696 | 0.6226 | 0.0421 | 0.4832 |
| HMR_5840 | Transport, extracellular | 10.5797 | 115.1998 | 9.9537 | 101.8725 | 0.626 | 0.0425 | 0.4831 |
| HMR_5841 | Transport, extracellular | 10.6139 | 115.9983 | 9.991 | 102.6779 | 0.6229 | 0.0421 | 0.4832 |
| HMR_5842 | Transport, extracellular | 10.6641 | 117.1056 | 10.0348 | 103.586 | 0.6293 | 0.0424 | 0.4831 |

| | | | | | | | | |
|---|---|---|---|---|---|---|---|---|
| HMR_5843 | Transport, extracellular | 10.7167 | 118.3206 | 10.0811 | 104.5878 | 0.6357 | 0.0426 | 0.483 |
| HMR_5844 | Transport, extracellular | 10.6137 | 115.9928 | 9.99 | 102.6578 | 0.6236 | 0.0422 | 0.4832 |
| HMR_5845 | Transport, extracellular | 10.5801 | 115.2098 | 9.9531 | 101.8608 | 0.627 | 0.0426 | 0.483 |
| HMR_5846 | Transport, extracellular | 10.6143 | 116.0084 | 9.9904 | 102.6661 | 0.6239 | 0.0422 | 0.4832 |
| HMR_5847 | Transport, extracellular | 10.6819 | 117.5107 | 10.0509 | 103.9306 | 0.6309 | 0.0424 | 0.4831 |
| HMR_5848 | Transport, extracellular | 10.7347 | 118.7321 | 10.0974 | 104.9374 | 0.6374 | 0.0426 | 0.483 |
| HMR_5849 | Transport, extracellular | 10.6313 | 116.3921 | 10.006 | 102.9977 | 0.6253 | 0.0422 | 0.4832 |
| HMR_5850 | Transport, extracellular | 10.5976 | 115.6051 | 9.969 | 102.1967 | 0.6286 | 0.0426 | 0.483 |
| HMR_5851 | Transport, extracellular | 10.632 | 116.4078 | 10.0064 | 103.0061 | 0.6256 | 0.0422 | 0.4832 |
| HMR_5852 | Transport, extracellular | 10.6773 | 117.4064 | 10.0467 | 103.8406 | 0.6306 | 0.0424 | 0.4831 |
| HMR_5853 | Transport, extracellular | 10.7301 | 118.6262 | 10.0931 | 104.8461 | 0.637 | 0.0426 | 0.483 |
| HMR_5854 | Transport, extracellular | 10.6268 | 116.2893 | 10.0019 | 102.9089 | 0.6249 | 0.0422 | 0.4832 |
| HMR_5855 | Transport, extracellular | 10.5931 | 115.5033 | 9.9649 | 102.109 | 0.6283 | 0.0426 | 0.483 |
| HMR_5856 | Transport, extracellular | 10.6274 | 116.305 | 10.0022 | 102.9172 | 0.6252 | 0.0422 | 0.4832 |
| HMR_5857 | Transport, extracellular | 10.6774 | 117.4092 | 10.0468 | 103.8427 | 0.6306 | 0.0424 | 0.4831 |
| HMR_5858 | Transport, extracellular | 10.7302 | 118.6289 | 10.0932 | 104.8483 | 0.637 | 0.0426 | 0.483 |
| HMR_5859 | Transport, extracellular | 10.6269 | 116.292 | 10.002 | 102.911 | 0.6249 | 0.0422 | 0.4832 |
| HMR_5860 | Transport, extracellular | 10.5932 | 115.506 | 9.965 | 102.1111 | 0.6283 | 0.0426 | 0.483 |
| HMR_5861 | Transport, extracellular | 10.6276 | 116.3077 | 10.0023 | 102.9194 | 0.6252 | 0.0422 | 0.4832 |
| HMR_5862 | Transport, extracellular | 10.6776 | 117.4127 | 10.047 | 103.8456 | 0.6306 | 0.0424 | 0.4831 |
| HMR_5863 | Transport, extracellular | 10.7304 | 118.6325 | 10.0934 | 104.8512 | 0.637 | 0.0426 | 0.483 |
| HMR_5864 | Transport, extracellular | 10.6271 | 116.2955 | 10.0021 | 102.9139 | 0.6249 | 0.0422 | 0.4832 |
| HMR_5865 | Transport, extracellular | 10.5934 | 115.5094 | 9.9651 | 102.1139 | 0.6283 | 0.0426 | 0.483 |
| HMR_5866 | Transport, extracellular | 10.6277 | 116.3111 | 10.0025 | 102.9222 | 0.6252 | 0.0422 | 0.4832 |
| HMR_5867 | Transport, extracellular | 10.6358 | 116.4724 | 10.0105 | 103.0732 | 0.6253 | 0.0422 | 0.4832 |
| HMR_5868 | Transport, extracellular | 10.6882 | 117.6774 | 10.0566 | 104.0674 | 0.6316 | 0.0424 | 0.4831 |
| HMR_5869 | Transport, extracellular | 10.5857 | 115.3686 | 9.966 | 102.1518 | 0.6197 | 0.042 | 0.4832 |
| HMR_5870 | Transport, extracellular | 10.5523 | 114.592 | 9.9292 | 101.3607 | 0.6231 | 0.0424 | 0.4831 |
| HMR_5871 | Transport, extracellular | 10.5863 | 115.3841 | 9.9663 | 102.16 | 0.62 | 0.042 | 0.4832 |
| HMR_5872 | Transport, extracellular | 10.6889 | 117.6709 | 10.0569 | 104.0585 | 0.632 | 0.0424 | 0.4831 |
| HMR_5873 | Transport, extracellular | 10.7418 | 118.8949 | 10.1034 | 105.0673 | 0.6384 | 0.0427 | 0.483 |
| HMR_5874 | Transport, extracellular | 10.6383 | 116.5501 | 10.012 | 103.1239 | 0.6263 | 0.0423 | 0.4831 |
| HMR_5875 | Transport, extracellular | 10.6046 | 115.7614 | 9.9749 | 102.3215 | 0.6297 | 0.0426 | 0.483 |
| HMR_5876 | Transport, extracellular | 10.6389 | 116.5658 | 10.0123 | 103.1323 | 0.6266 | 0.0423 | 0.4831 |
| HMR_5877 | Transport, extracellular | 10.6637 | 117.0982 | 10.0366 | 103.6247 | 0.6271 | 0.0422 | 0.4832 |
| HMR_5878 | Transport, extracellular | 10.7163 | 118.313 | 10.0829 | 104.627 | 0.6334 | 0.0424 | 0.4831 |
| HMR_5879 | Transport, extracellular | 10.6132 | 115.9854 | 9.9918 | 102.6959 | 0.6214 | 0.042 | 0.4832 |
| HMR_5880 | Transport, extracellular | 10.5797 | 115.2026 | 9.9549 | 101.8985 | 0.6248 | 0.0424 | 0.4831 |
| HMR_5881 | Transport, extracellular | 10.6139 | 116.001 | 9.9922 | 102.7043 | 0.6217 | 0.042 | 0.4832 |
| HMR_5882 | Transport, extracellular | 10.6619 | 117.0586 | 10.0333 | 103.5544 | 0.6287 | 0.0423 | 0.4831 |
| HMR_5883 | Transport, extracellular | 10.7146 | 118.2728 | 10.0795 | 104.5557 | 0.635 | 0.0425 | 0.483 |
| HMR_5884 | Transport, extracellular | 10.6115 | 115.9464 | 9.9885 | 102.6266 | 0.623 | 0.0421 | 0.4832 |
| HMR_5885 | Transport, extracellular | 10.578 | 115.1639 | 9.9516 | 101.83 | 0.6264 | 0.0425 | 0.483 |
| HMR_5886 | Transport, extracellular | 10.6122 | 115.962 | 9.9889 | 102.6349 | 0.6233 | 0.0422 | 0.4832 |

| | | | | | | | | |
|---|---|---|---|---|---|---|---|---|
| HMR_5887 | Transport, extracellular | 10.6833 | 117.5397 | 10.0466 | 103.8373 | 0.6368 | 0.0428 | 0.4829 |
| HMR_5888 | Transport, extracellular | 10.7362 | 118.7615 | 10.093 | 104.8428 | 0.6432 | 0.043 | 0.4828 |
| HMR_5889 | Transport, extracellular | 10.6327 | 116.4207 | 10.0017 | 102.9057 | 0.631 | 0.0426 | 0.483 |
| HMR_5890 | Transport, extracellular | 10.599 | 115.6333 | 9.9647 | 102.1058 | 0.6343 | 0.043 | 0.4829 |
| HMR_5891 | Transport, extracellular | 10.6334 | 116.4364 | 10.0021 | 102.914 | 0.6313 | 0.0426 | 0.483 |
| HMR_5892 | Transport, extracellular | 10.6628 | 117.0796 | 10.034 | 103.5707 | 0.6288 | 0.0423 | 0.4831 |
| HMR_5893 | Transport, extracellular | 10.7155 | 118.2941 | 10.0803 | 104.5722 | 0.6352 | 0.0425 | 0.483 |
| HMR_5894 | Transport, extracellular | 10.6124 | 115.9671 | 9.9893 | 102.6426 | 0.6232 | 0.0421 | 0.4832 |
| HMR_5895 | Transport, extracellular | 10.5789 | 115.1844 | 9.9523 | 101.8458 | 0.6265 | 0.0425 | 0.483 |
| HMR_5896 | Transport, extracellular | 10.6131 | 115.9827 | 9.9896 | 102.6509 | 0.6235 | 0.0422 | 0.4832 |
| HMR_5897 | Transport, extracellular | 10.6768 | 117.3951 | 10.0463 | 103.8318 | 0.6305 | 0.0424 | 0.4831 |
| HMR_5898 | Transport, extracellular | 10.7296 | 118.6147 | 10.0927 | 104.8372 | 0.6369 | 0.0426 | 0.483 |
| HMR_5899 | Transport, extracellular | 10.6263 | 116.2782 | 10.0015 | 102.9002 | 0.6248 | 0.0422 | 0.4832 |
| HMR_5900 | Transport, extracellular | 10.5926 | 115.4923 | 9.9645 | 102.1004 | 0.6282 | 0.0426 | 0.483 |
| HMR_5901 | Transport, extracellular | 10.6269 | 116.2938 | 10.0018 | 102.9086 | 0.6251 | 0.0422 | 0.4832 |
| HMR_5902 | Transport, extracellular | 10.6773 | 117.4061 | 10.0466 | 103.8373 | 0.6307 | 0.0424 | 0.4831 |
| HMR_5903 | Transport, extracellular | 10.7301 | 118.6258 | 10.093 | 104.8427 | 0.6371 | 0.0426 | 0.483 |
| HMR_5904 | Transport, extracellular | 10.6268 | 116.289 | 10.0017 | 102.9056 | 0.625 | 0.0422 | 0.4832 |
| HMR_5905 | Transport, extracellular | 10.5931 | 115.503 | 9.9647 | 102.1057 | 0.6284 | 0.0426 | 0.483 |
| HMR_5906 | Transport, extracellular | 10.6274 | 116.3047 | 10.0021 | 102.914 | 0.6253 | 0.0422 | 0.4832 |
| HMR_5907 | Transport, extracellular | 41.8935 | 2022.1535 | 43.1179 | 2143.8143 | 1.2244 | -0.019 | 0.5076 |
| HMR_5908 | Transport, extracellular | 41.5733 | 1987.1857 | 42.7617 | 2103.7201 | 1.1884 | -0.0186 | 0.5074 |
| HMR_5909 | Transport, extracellular | 41.4445 | 1972.4319 | 42.6418 | 2089.473 | 1.1973 | -0.0188 | 0.5075 |
| HMR_5910 | Transport, extracellular | 41.4368 | 1972.2378 | 42.6515 | 2091.3652 | 1.2147 | -0.0191 | 0.5076 |
| HMR_5911 | Transport, extracellular | 41.173 | 1943.7821 | 42.3653 | 2059.4769 | 1.1922 | -0.0188 | 0.5075 |
| HMR_5912 | Transport, extracellular | 41.2408 | 1950.7963 | 42.44 | 2067.4407 | 1.1991 | -0.0189 | 0.5075 |
| HMR_5913 | Transport, extracellular | 41.2391 | 1950.6047 | 42.4382 | 2067.2387 | 1.1991 | -0.0189 | 0.5075 |
| HMR_5914 | Transport, extracellular | 41.2368 | 1950.3592 | 42.4357 | 2066.9752 | 1.199 | -0.0189 | 0.5075 |
| HMR_5915 | Transport, extracellular | 41.8615 | 2019.5384 | 43.0882 | 2141.3371 | 1.2267 | -0.019 | 0.5076 |
| HMR_5916 | Transport, extracellular | 41.0698 | 1932.7441 | 42.2604 | 2047.9343 | 1.1906 | -0.0189 | 0.5075 |
| HMR_5917 | Transport, extracellular | 41.4427 | 1972.9274 | 42.6195 | 2087.5489 | 1.1768 | -0.0185 | 0.5074 |
| HMR_5918 | Transport, extracellular | 41.4686 | 1975.8271 | 42.6784 | 2094.4561 | 1.2098 | -0.019 | 0.5076 |
| HMR_5919 | Transport, extracellular | 41.153 | 1941.2382 | 42.4426 | 2067.7387 | 1.2897 | -0.0204 | 0.5081 |
| HMR_5920 | Transport, extracellular | 41.4549 | 1974.338 | 42.6649 | 2092.9471 | 1.21 | -0.019 | 0.5076 |
| HMR_5921 | Transport, extracellular | 41.2481 | 1951.5798 | 42.4471 | 2068.2388 | 1.199 | -0.0189 | 0.5075 |
| HMR_5922 | Transport, extracellular | 41.241 | 1950.8183 | 42.4427 | 2067.7431 | 1.2016 | -0.019 | 0.5076 |
| HMR_5923 | Transport, extracellular | 41.0971 | 1940.6654 | 42.2805 | 2055.6823 | 1.1834 | -0.0187 | 0.5075 |
| HMR_5924 | Transport, extracellular | 40.7888 | 1907.8051 | 41.9378 | 2018.0511 | 1.149 | -0.0183 | 0.5073 |
| HMR_5925 | Transport, extracellular | 40.6647 | 1893.9455 | 41.8224 | 2004.6847 | 1.1578 | -0.0185 | 0.5074 |
| HMR_5926 | Transport, extracellular | 40.6573 | 1893.7551 | 41.8318 | 2006.4511 | 1.1745 | -0.0188 | 0.5075 |
| HMR_5927 | Transport, extracellular | 40.4032 | 1866.9951 | 41.5563 | 1976.5012 | 1.1531 | -0.0186 | 0.5074 |
| HMR_5928 | Transport, extracellular | 40.4685 | 1873.5961 | 41.6282 | 1983.9866 | 1.1596 | -0.0187 | 0.5074 |
| HMR_5929 | Transport, extracellular | 40.4669 | 1873.4157 | 41.6265 | 1983.7967 | 1.1596 | -0.0187 | 0.5074 |
| HMR_5930 | Transport, extracellular | 40.4646 | 1873.185 | 41.6241 | 1983.5492 | 1.1595 | -0.0187 | 0.5074 |

| | | | | | | | | |
|---|---|---|---|---|---|---|---|---|
| HMR_5931 | Transport, extracellular | 41.0663 | 1938.1997 | 42.252 | 2053.3475 | 1.1856 | -0.0188 | 0.5075 |
| HMR_5932 | Transport, extracellular | 40.3038 | 1856.6111 | 41.4553 | 1965.6555 | 1.1516 | -0.0186 | 0.5074 |
| HMR_5933 | Transport, extracellular | 40.663 | 1894.3997 | 41.801 | 2002.8626 | 1.138 | -0.0182 | 0.5073 |
| HMR_5934 | Transport, extracellular | 40.6879 | 1897.1279 | 41.8576 | 2009.3524 | 1.1697 | -0.0187 | 0.5075 |
| HMR_5935 | Transport, extracellular | 40.3839 | 1864.6065 | 41.6307 | 1984.2665 | 1.2469 | -0.0201 | 0.508 |
| HMR_5936 | Transport, extracellular | 40.6748 | 1895.7282 | 41.8447 | 2007.9357 | 1.1699 | -0.0187 | 0.5075 |
| HMR_5937 | Transport, extracellular | 40.4755 | 1874.3328 | 41.6351 | 1984.7361 | 1.1595 | -0.0187 | 0.5074 |
| HMR_5938 | Transport, extracellular | 40.4687 | 1873.6167 | 41.6308 | 1984.2706 | 1.162 | -0.0187 | 0.5075 |
| HMR_5939 | Transport, extracellular | 42.6611 | 2116.3212 | 43.9322 | 2247.3393 | 1.2711 | -0.0192 | 0.5077 |
| HMR_5940 | Transport, extracellular | 42.3295 | 2078.7921 | 43.563 | 2204.1969 | 1.2335 | -0.0188 | 0.5075 |
| HMR_5941 | Transport, extracellular | 42.1962 | 2062.9576 | 43.4388 | 2188.8661 | 1.2427 | -0.0191 | 0.5076 |
| HMR_5942 | Transport, extracellular | 42.1882 | 2062.7552 | 43.4488 | 2190.909 | 1.2606 | -0.0193 | 0.5077 |
| HMR_5943 | Transport, extracellular | 41.9151 | 2032.2444 | 43.1522 | 2156.6288 | 1.2371 | -0.0191 | 0.5076 |
| HMR_5944 | Transport, extracellular | 41.9853 | 2039.7609 | 43.2296 | 2165.1848 | 1.2443 | -0.0192 | 0.5077 |
| HMR_5945 | Transport, extracellular | 41.9835 | 2039.5555 | 43.2278 | 2164.9676 | 1.2442 | -0.0192 | 0.5077 |
| HMR_5946 | Transport, extracellular | 41.9811 | 2039.2925 | 43.2253 | 2164.6845 | 1.2441 | -0.0192 | 0.5077 |
| HMR_5947 | Transport, extracellular | 42.6278 | 2113.5225 | 43.9013 | 2244.6811 | 1.2735 | -0.0193 | 0.5077 |
| HMR_5948 | Transport, extracellular | 41.8083 | 2020.4141 | 43.0435 | 2144.2265 | 1.2353 | -0.0191 | 0.5076 |
| HMR_5949 | Transport, extracellular | 42.1943 | 2063.4957 | 43.4157 | 2186.8011 | 1.2214 | -0.0187 | 0.5075 |
| HMR_5950 | Transport, extracellular | 42.2211 | 2066.6072 | 43.4766 | 2194.2342 | 1.2555 | -0.0192 | 0.5077 |
| HMR_5951 | Transport, extracellular | 41.8944 | 2029.5132 | 43.2324 | 2165.5051 | 1.338 | -0.0207 | 0.5082 |
| HMR_5952 | Transport, extracellular | 42.2069 | 2065.01 | 43.4627 | 2192.6114 | 1.2558 | -0.0192 | 0.5077 |
| HMR_5953 | Transport, extracellular | 41.9928 | 2040.6008 | 43.237 | 2166.0426 | 1.2442 | -0.0192 | 0.5077 |
| HMR_5954 | Transport, extracellular | 41.9856 | 2039.7845 | 43.2324 | 2165.5098 | 1.2469 | -0.0192 | 0.5077 |
| HMR_5955 | Transport, extracellular | 43.2242 | 2172.7528 | 44.6698 | 2326.4 | 1.4457 | -0.0216 | 0.5086 |
| HMR_5956 | Transport, extracellular | 42.8838 | 2133.7209 | 44.2882 | 2280.958 | 1.4044 | -0.0211 | 0.5084 |
| HMR_5957 | Transport, extracellular | 42.747 | 2117.2469 | 44.1599 | 2264.8048 | 1.4129 | -0.0213 | 0.5085 |
| HMR_5958 | Transport, extracellular | 42.7388 | 2117.0346 | 44.1702 | 2266.9616 | 1.4314 | -0.0216 | 0.5086 |
| HMR_5959 | Transport, extracellular | 42.4586 | 2085.3268 | 43.8638 | 2230.8823 | 1.4052 | -0.0214 | 0.5085 |
| HMR_5960 | Transport, extracellular | 42.5306 | 2093.136 | 43.9438 | 2239.8827 | 1.4131 | -0.0215 | 0.5086 |
| HMR_5961 | Transport, extracellular | 42.5288 | 2092.9224 | 43.9418 | 2239.6541 | 1.4131 | -0.0215 | 0.5086 |
| HMR_5962 | Transport, extracellular | 42.5263 | 2092.6492 | 43.9393 | 2239.3564 | 1.413 | -0.0215 | 0.5086 |
| HMR_5963 | Transport, extracellular | 43.19 | 2169.8496 | 44.6379 | 2323.5963 | 1.4478 | -0.0216 | 0.5086 |
| HMR_5964 | Transport, extracellular | 42.3489 | 2073.0312 | 43.7515 | 2217.8326 | 1.4026 | -0.0214 | 0.5085 |
| HMR_5965 | Transport, extracellular | 42.7451 | 2117.8139 | 44.136 | 2262.6277 | 1.3909 | -0.021 | 0.5084 |
| HMR_5966 | Transport, extracellular | 42.7725 | 2121.0507 | 44.1989 | 2270.4692 | 1.4264 | -0.0215 | 0.5086 |
| HMR_5967 | Transport, extracellular | 42.4373 | 2082.4751 | 43.9466 | 2240.2199 | 1.5093 | -0.023 | 0.5092 |
| HMR_5968 | Transport, extracellular | 42.758 | 2119.3902 | 44.1846 | 2268.7608 | 1.4265 | -0.0215 | 0.5086 |
| HMR_5969 | Transport, extracellular | 42.5383 | 2094.0092 | 43.9514 | 2240.7856 | 1.4131 | -0.0215 | 0.5086 |
| HMR_5970 | Transport, extracellular | 42.5308 | 2093.1604 | 43.9467 | 2240.2248 | 1.4158 | -0.0215 | 0.5086 |
| HMR_5971 | Transport, extracellular | 42.6505 | 2115.1282 | 43.9251 | 2246.603 | 1.2746 | -0.0193 | 0.5077 |
| HMR_5972 | Transport, extracellular | 42.3191 | 2077.6314 | 43.556 | 2203.4818 | 1.2369 | -0.0189 | 0.5075 |
| HMR_5973 | Transport, extracellular | 42.1858 | 2061.8105 | 43.4319 | 2188.1586 | 1.2461 | -0.0191 | 0.5076 |
| HMR_5974 | Transport, extracellular | 42.1778 | 2061.608 | 43.4418 | 2190.2002 | 1.264 | -0.0194 | 0.5077 |

| | | | | | | | | |
|---|---|---|---|---|---|---|---|---|
| HMR_5975 | Transport, extracellular | 41.9049 | 2031.123 | 43.1454 | 2155.9365 | 1.2404 | -0.0192 | 0.5076 |
| HMR_5976 | Transport, extracellular | 41.9751 | 2038.6332 | 43.2227 | 2164.4883 | 1.2476 | -0.0192 | 0.5077 |
| HMR_5977 | Transport, extracellular | 41.9733 | 2038.4278 | 43.2209 | 2164.2713 | 1.2476 | -0.0192 | 0.5077 |
| HMR_5978 | Transport, extracellular | 41.9709 | 2038.1649 | 43.2184 | 2163.9882 | 1.2475 | -0.0192 | 0.5077 |
| HMR_5979 | Transport, extracellular | 42.6173 | 2112.3317 | 43.8942 | 2243.9459 | 1.2769 | -0.0193 | 0.5077 |
| HMR_5980 | Transport, extracellular | 41.7981 | 2019.3027 | 43.0367 | 2143.5401 | 1.2386 | -0.0192 | 0.5077 |
| HMR_5981 | Transport, extracellular | 42.184 | 2062.3485 | 43.4087 | 2186.0968 | 1.2248 | -0.0188 | 0.5075 |
| HMR_5982 | Transport, extracellular | 42.2107 | 2065.4568 | 43.4697 | 2193.5239 | 1.2589 | -0.0193 | 0.5077 |
| HMR_5983 | Transport, extracellular | 41.8842 | 2028.3941 | 43.2255 | 2164.8085 | 1.3413 | -0.0207 | 0.5083 |
| HMR_5984 | Transport, extracellular | 42.1966 | 2063.8607 | 43.4557 | 2191.9017 | 1.2592 | -0.0193 | 0.5077 |
| HMR_5985 | Transport, extracellular | 41.9826 | 2039.4726 | 43.2302 | 2165.3459 | 1.2476 | -0.0192 | 0.5077 |
| HMR_5986 | Transport, extracellular | 41.9753 | 2038.6567 | 43.2255 | 2164.8132 | 1.2502 | -0.0193 | 0.5077 |
| HMR_5987 | Transport, extracellular | 13.9445 | 80.4864 | 13.4789 | 75.7148 | 0.4656 | 0.0373 | 0.4851 |
| HMR_5990 | Transport, extracellular | 637.095 | 47946.7152 | 602.9554 | 50294.386 | 34.1396 | 0.1089 | 0.4566 |
| HMR_5993 | Transport, extracellular | 43.6794 | 471.0294 | 42.8095 | 439.2942 | 0.8699 | 0.0288 | 0.4885 |
| HMR_5994 | Transport, extracellular | 18.9161 | 285.0199 | 18.0082 | 261.468 | 0.9079 | 0.0388 | 0.4845 |
| HMR_5995 | Transport, extracellular | 6.0041 | 44.892 | 5.7325 | 40.4617 | 0.2716 | 0.0294 | 0.4883 |
| HMR_5996 | Transport, extracellular | 3.5391 | 13.6131 | 3.3818 | 12.392 | 0.1574 | 0.0309 | 0.4877 |
| HMR_5998 | Transport, extracellular | 5.8713 | 38.796 | 5.839 | 38.3791 | 0.0323 | 0.0037 | 0.4985 |
| HMR_6000 | Transport, extracellular | 87.7712 | 303526.8831 | 75.1515 | 305138.0537 | 12.6197 | 0.0162 | 0.4935 |
| HMR_6001 | Transport, extracellular | -2.7845 | 307841.6831 | -12.917 | 307834.6896 | 10.1325 | 0.0129 | 0.4948 |
| HMR_6002 | Transport, extracellular | 14.1076 | 307736.6218 | 24.3828 | 307599.1019 | 10.2752 | -0.0131 | 0.5052 |
| HMR_6003 | Transport, extracellular | 14.1072 | 307736.6223 | 24.3824 | 307599.082 | 10.2753 | -0.0131 | 0.5052 |
| HMR_6004 | Transport, extracellular | 44.148 | 306706.8164 | 11.7271 | 307876.7665 | 32.421 | 0.0414 | 0.4835 |
| HMR_6005 | Transport, extracellular | 14.1077 | 307736.6331 | 24.3829 | 307599.0941 | 10.2752 | -0.0131 | 0.5052 |
| HMR_6006 | Transport, extracellular | -13.6844 | 307743.94 | -24.7333 | 307590.3495 | 11.0489 | 0.0141 | 0.4944 |
| HMR_6007 | Transport, extracellular | 74.3167 | 304648.3095 | 50.7479 | 306547.2187 | 23.5688 | 0.0301 | 0.488 |
| HMR_6008 | Transport, extracellular | -10.9035 | 307732.7343 | -11.832 | 307736.6708 | 0.9284 | 0.0012 | 0.4995 |
| HMR_6009 | Transport, extracellular | 0.4234 | 307799.1335 | -0.3512 | 307814.8341 | 0.7747 | 0.001 | 0.4996 |
| HMR_6010 | Transport, extracellular | 0.423 | 307799.1265 | -0.3517 | 307814.7797 | 0.7747 | 0.001 | 0.4996 |
| HMR_6011 | Transport, extracellular | -57.7604 | 305927.1847 | -36.4211 | 307284.9415 | 21.3393 | -0.0273 | 0.5109 |
| HMR_6012 | Transport, extracellular | 0.4234 | 307799.1443 | -0.3513 | 307814.7878 | 0.7747 | 0.001 | 0.4996 |
| HMR_6013 | Transport, extracellular | 0.4235 | 307799.0992 | -0.3513 | 307814.8129 | 0.7747 | 0.001 | 0.4996 |
| HMR_6014 | Transport, extracellular | 74.052 | 304669.1598 | 51.2423 | 306518.9884 | 22.8097 | 0.0292 | 0.4884 |
| HMR_6015 | Transport, extracellular | 85.0674 | 303704.1389 | 62.484 | 305814.8044 | 22.5835 | 0.0289 | 0.4885 |
| HMR_6016 | Transport, extracellular | 73.8987 | 304681.5715 | 51.0969 | 306527.0861 | 22.8018 | 0.0292 | 0.4884 |
| HMR_6017 | Transport, extracellular | 73.8991 | 304681.5362 | 51.0974 | 306527.0351 | 22.8018 | 0.0292 | 0.4884 |
| HMR_6018 | Transport, extracellular | 130.7377 | 298408.7769 | 86.5096 | 304534.3594 | 44.228 | 0.057 | 0.4773 |
| HMR_6019 | Transport, extracellular | 73.8987 | 304681.5759 | 51.097 | 306527.0295 | 22.8018 | 0.0292 | 0.4884 |
| HMR_6020 | Transport, extracellular | -73.8987 | 304681.6191 | -51.0969 | 306527.0137 | 22.8018 | -0.0292 | 0.5116 |
| HMR_6021 | Transport, extracellular | 63.7013 | 305471.7525 | 30.5505 | 307394.333 | 33.1508 | 0.0423 | 0.4831 |
| HMR_6022 | Transport, extracellular | -11.1718 | 307728.5242 | -11.3345 | 307741.9585 | 0.1628 | 0.0002 | 0.4999 |
| HMR_6023 | Transport, extracellular | 0.1552 | 307798.3256 | 0.1464 | 307813.8605 | 0.0088 | 0.00E+00 | 0.5 |
| HMR_6024 | Transport, extracellular | 0.1547 | 307798.3438 | 0.146 | 307813.8412 | 0.0087 | 0.00E+00 | 0.5 |

| | | | | | | | | |
|---|---|---|---|---|---|---|---|---|
| HMR_6025 | Transport, extracellular | -13.9525 | 307739.1063 | -24.2367 | 307603.0961 | 10.2842 | 0.0131 | 0.4948 |
| HMR_6026 | Transport, extracellular | -58.0274 | 305907.3346 | -35.9261 | 307300.9948 | 22.1013 | -0.0282 | 0.5113 |
| HMR_6027 | Transport, extracellular | 0.1552 | 307798.3549 | 0.1464 | 307813.8015 | 0.0088 | 0.00E+00 | 0.5 |
| HMR_6028 | Transport, extracellular | 0.1552 | 307798.3595 | 0.1464 | 307813.7993 | 0.0088 | 0.00E+00 | 0.5 |
| HMR_6029 | Transport, extracellular | -10.451 | 307788.2658 | -20.7798 | 307693.4816 | 10.3289 | 0.0132 | 0.4947 |
| HMR_6030 | Transport, extracellular | 0.2682 | 307799.2689 | -0.4976 | 307814.8371 | 0.7658 | 0.001 | 0.4996 |
| HMR_6031 | Transport, extracellular | 11.3269 | 307726.4579 | 11.4809 | 307739.9958 | 0.154 | -0.0002 | 0.5001 |
| HMR_6032 | Transport, extracellular | 11.3265 | 307726.5124 | 11.4805 | 307740.0866 | 0.154 | -0.0002 | 0.5001 |
| HMR_6033 | Transport, extracellular | 46.923 | 306554.7382 | 24.6257 | 307682.4027 | 22.2973 | 0.0285 | 0.4887 |
| HMR_6034 | Transport, extracellular | 11.3269 | 307726.4419 | 11.4809 | 307740.0858 | 0.1539 | -0.0002 | 0.5001 |
| HMR_6035 | Transport, extracellular | 11.327 | 307726.4663 | 11.4809 | 307740.0748 | 0.154 | -0.0002 | 0.5001 |
| HMR_6036 | Transport, extracellular | -21.6141 | 307589.2602 | -32.0903 | 307362.0424 | 10.4761 | 0.0134 | 0.4947 |
| HMR_6037 | Transport, extracellular | -0.0004 | 307798.2718 | -0.0004 | 307813.8021 | 0.00E+00 | 0.00E+00 | 0.5 |
| HMR_6038 | Transport, extracellular | 58.1816 | 305897.2234 | 36.0719 | 307294.9333 | 22.1096 | 0.0282 | 0.4887 |
| HMR_6039 | Transport, extracellular | 0.00E+00 | 307798.2677 | 0.00E+00 | 307813.7551 | 0.00E+00 | 0.00E+00 | 0.5 |
| HMR_6040 | Transport, extracellular | 0.00E+00 | 307798.2814 | 0.00E+00 | 307813.7569 | 0.00E+00 | 0.00E+00 | 0.5 |
| HMR_6041 | Transport, extracellular | -10.2958 | 307789.9346 | -20.6336 | 307696.7676 | 10.3378 | 0.0132 | 0.4947 |
| HMR_6042 | Transport, extracellular | 58.1811 | 305897.2346 | 36.0715 | 307294.98 | 22.1096 | 0.0282 | 0.4887 |
| HMR_6043 | Transport, extracellular | -0.0004 | 307798.275 | -0.0004 | 307813.7742 | 1.00E-04 | 0.00E+00 | 0.5 |
| HMR_6044 | Transport, extracellular | 0.0005 | 307798.2352 | 0.0004 | 307813.7498 | 1.00E-04 | 0.00E+00 | 0.5 |
| HMR_6045 | Transport, extracellular | -10.2963 | 307789.9266 | -20.634 | 307696.7508 | 10.3377 | 0.0132 | 0.4947 |
| HMR_6046 | Transport, extracellular | 14.1076 | 307736.6105 | 24.3828 | 307599.0951 | 10.2752 | -0.0131 | 0.5052 |
| HMR_6047 | Transport, extracellular | -24.3752 | 307683.3413 | -44.875 | 307213.6418 | 20.4999 | 0.0261 | 0.4896 |
| HMR_6048 | Transport, extracellular | 58.1815 | 305897.0891 | 36.0719 | 307294.9623 | 22.1096 | 0.0282 | 0.4887 |
| HMR_6049 | Transport, extracellular | 58.1815 | 305897.0907 | 36.072 | 307294.9846 | 22.1095 | 0.0282 | 0.4887 |
| HMR_6050 | Transport, extracellular | -68.3589 | 305269.5922 | -56.5175 | 306541.3606 | 11.8414 | -0.0151 | 0.506 |
| HMR_6051 | Transport, extracellular | 0.00E+00 | 307798.2789 | 0.00E+00 | 307813.7722 | 0.00E+00 | 0.00E+00 | 0.5 |
| HMR_6052 | Transport, extracellular | -10.2958 | 307789.9332 | -20.6337 | 307696.7613 | 10.3378 | 0.0132 | 0.4947 |
| HMR_6053 | Transport, extracellular | -10.2958 | 307789.9401 | -20.6336 | 307696.7784 | 10.3378 | 0.0132 | 0.4947 |
| HMR_6054 | Transport, extracellular | -10.719 | 307786.1689 | -20.2828 | 307706.1811 | 9.5638 | 0.0122 | 0.4951 |
| HMR_6055 | Transport, extracellular | 4.3768 | 8.9155 | 4.191 | 8.1745 | 0.1858 | 0.0449 | 0.4821 |
| HMR_6056 | Transport, extracellular | 4.3789 | 8.9175 | 4.1931 | 8.1764 | 0.1858 | 0.0449 | 0.4821 |
| HMR_6057 | Transport, extracellular | 2.61 | 7.5763 | 2.5097 | 6.9933 | 0.1003 | 0.0263 | 0.4895 |
| HMR_6058 | Transport, extracellular | 426.6178 | 48422.5293 | 398.0496 | 50511.8806 | 28.5682 | 0.0908 | 0.4638 |
| HMR_6096 | Transport, extracellular | 4.3763 | 8.9155 | 4.1904 | 8.1744 | 0.1858 | 0.045 | 0.4821 |
| HMR_6098 | Transport, extracellular | 4.3749 | 8.9174 | 4.1891 | 8.1764 | 0.1858 | 0.0449 | 0.4821 |
| HMR_6101 | Transport, extracellular | 1.7729 | 2.6727 | 1.6944 | 2.4408 | 0.0785 | 0.0347 | 0.4862 |
| HMR_6129 | Transport, extracellular | 0.9155 | 0.7255 | 0.887 | 0.683 | 0.0285 | 0.024 | 0.4904 |
| HMR_6218 | Transport, extracellular | -436.2331 | 75333.533 | -436.2505 | 75331.1993 | 0.0174 | 0.00E+00 | 0.5 |
| HMR_6219 | Transport, extracellular | 100.5154 | 243724.119 | 100.5638 | 243717.9869 | 0.0483 | -1.00E-04 | 0.5 |
| HMR_6220 | Transport, extracellular | -100.5154 | 243724.1139 | -100.5638 | 243717.9867 | 0.0483 | 1.00E-04 | 0.5 |
| HMR_6221 | Transport, extracellular | 100.5154 | 243724.117 | 100.5638 | 243717.9933 | 0.0483 | -1.00E-04 | 0.5 |
| HMR_6222 | Transport, extracellular | 100.5154 | 243724.1172 | 100.5638 | 243717.9941 | 0.0483 | -1.00E-04 | 0.5 |
| HMR_6223 | Transport, extracellular | 100.5154 | 243724.1177 | 100.5638 | 243718.0009 | 0.0483 | -1.00E-04 | 0.5 |

| | | | | | | | | |
|---|---|---|---|---|---|---|---|---|
| HMR_6224 | Transport, extracellular | 100.5154 | 243724.1185 | 100.5638 | 243718.0007 | 0.0483 | -1.00E-04 | 0.5 |
| HMR_6225 | Transport, extracellular | 100.5153 | 243724.1094 | 100.5635 | 243717.968 | 0.0482 | -1.00E-04 | 0.5 |
| HMR_6226 | Transport, extracellular | 100.5154 | 243724.1188 | 100.5638 | 243717.9999 | 0.0483 | -1.00E-04 | 0.5 |
| HMR_6227 | Transport, extracellular | 100.5153 | 243724.1158 | 100.5635 | 243717.9654 | 0.0482 | -1.00E-04 | 0.5 |
| HMR_6228 | Transport, extracellular | 100.5155 | 243724.1187 | 100.5638 | 243717.9966 | 0.0484 | -1.00E-04 | 0.5 |
| HMR_6229 | Transport, extracellular | 100.5153 | 243724.1034 | 100.5636 | 243717.9531 | 0.0482 | -1.00E-04 | 0.5 |
| HMR_6230 | Transport, extracellular | 100.5155 | 243724.1168 | 100.5637 | 243718.0084 | 0.0483 | -1.00E-04 | 0.5 |
| HMR_6231 | Transport, extracellular | -477.6838 | 63089.9751 | -477.5795 | 63089.6723 | 0.1043 | -0.0003 | 0.5001 |
| HMR_6232 | Transport, extracellular | 100.5155 | 243724.1097 | 100.5637 | 243718.0164 | 0.0481 | -1.00E-04 | 0.5 |
| HMR_6233 | Transport, extracellular | -440.4316 | 75120.9333 | -440.3769 | 75121.421 | 0.0547 | -0.0001 | 0.5001 |
| HMR_6234 | Transport, extracellular | 100.5155 | 243724.1099 | 100.5637 | 243718.0163 | 0.0481 | -1.00E-04 | 0.5 |
| HMR_6236 | Transport, extracellular | -475.5127 | 63090.7687 | -475.5009 | 63090.4004 | 0.0118 | 0.00E+00 | 0.5 |
| HMR_6237 | Transport, extracellular | 100.5155 | 243724.1103 | 100.5637 | 243718.0161 | 0.0481 | -1.00E-04 | 0.5 |
| HMR_6238 | Transport, extracellular | 100.5155 | 243724.111 | 100.5637 | 243718.0176 | 0.0481 | -1.00E-04 | 0.5 |
| HMR_6239 | Transport, extracellular | 100.5155 | 243724.1115 | 100.5635 | 243718.0214 | 0.0481 | -1.00E-04 | 0.5 |
| HMR_6241 | Transport, extracellular | 5.2995 | 113.4318 | 5.1229 | 103.8713 | 0.1765 | 0.012 | 0.4952 |
| HMR_6242 | Transport, extracellular | 4.254 | 18.3643 | 4.0737 | 16.8441 | 0.1803 | 0.0304 | 0.4879 |
| HMR_6245 | Transport, extracellular | 38.0032 | 571.4444 | 33.074 | 437.3298 | 4.9291 | 0.1552 | 0.4383 |
| HMR_6246 | Transport, extracellular | 2.7688 | 5.4436 | 2.6434 | 4.9627 | 0.1254 | 0.0389 | 0.4845 |
| HMR_6247 | Transport, extracellular | 475.5125 | 63090.7688 | 475.5008 | 63090.4004 | 0.0117 | 0.00E+00 | 0.5 |
| HMR_6248 | Transport, extracellular | 1.1323 | 1.042 | 1.1006 | 0.9873 | 0.0317 | 0.0223 | 0.4911 |
| HMR_6249 | Transport, extracellular | 1.575 | 1.7156 | 1.6178 | 1.805 | 0.0428 | -0.0228 | 0.5091 |
| HMR_6250 | Transport, extracellular | -2.8478 | 3.7899 | -2.7188 | 3.4543 | 0.129 | -0.0479 | 0.5191 |
| HMR_6251 | Transport, extracellular | 100.5154 | 243724.1193 | 100.5638 | 243717.9872 | 0.0483 | -1.00E-04 | 0.5 |
| HMR_6252 | Transport, extracellular | 100.5154 | 243724.1142 | 100.5638 | 243717.9869 | 0.0483 | -1.00E-04 | 0.5 |
| HMR_6254 | Transport, extracellular | 100.5154 | 243724.1172 | 100.5638 | 243717.9936 | 0.0483 | -1.00E-04 | 0.5 |
| HMR_6255 | Transport, extracellular | 100.5154 | 243724.1175 | 100.5638 | 243717.9943 | 0.0483 | -1.00E-04 | 0.5 |
| HMR_6256 | Transport, extracellular | 100.5154 | 243724.1179 | 100.5638 | 243718.0012 | 0.0483 | -1.00E-04 | 0.5 |
| HMR_6259 | Transport, extracellular | 100.5154 | 243724.1188 | 100.5638 | 243718.001 | 0.0483 | -1.00E-04 | 0.5 |
| HMR_6260 | Transport, extracellular | 100.5156 | 243724.1097 | 100.5639 | 243717.9683 | 0.0483 | -1.00E-04 | 0.5 |
| HMR_6261 | Transport, extracellular | 100.5154 | 243724.1191 | 100.5638 | 243718.0002 | 0.0483 | -1.00E-04 | 0.5 |
| HMR_6262 | Transport, extracellular | 100.5156 | 243724.1161 | 100.5639 | 243717.9657 | 0.0483 | -1.00E-04 | 0.5 |
| HMR_6263 | Transport, extracellular | 100.5155 | 243724.119 | 100.5638 | 243717.9969 | 0.0484 | -1.00E-04 | 0.5 |
| HMR_6264 | Transport, extracellular | 100.5157 | 243724.1037 | 100.564 | 243717.9534 | 0.0483 | -1.00E-04 | 0.5 |
| HMR_6265 | Transport, extracellular | 100.5155 | 243724.1171 | 100.5637 | 243718.0087 | 0.0483 | -1.00E-04 | 0.5 |
| HMR_6266 | Transport, extracellular | 473.3522 | 63090.1499 | 473.432 | 63089.8322 | 0.0799 | -0.0002 | 0.5001 |
| HMR_6267 | Transport, extracellular | 100.5155 | 243724.1099 | 100.5637 | 243718.0167 | 0.0481 | -1.00E-04 | 0.5 |
| HMR_6268 | Transport, extracellular | 438.9583 | 75100.8507 | 438.9575 | 75103.0251 | 0.0007 | 0.00E+00 | 0.5 |
| HMR_6269 | Transport, extracellular | 100.5155 | 243724.1102 | 100.5637 | 243718.0166 | 0.0481 | -1.00E-04 | 0.5 |
| HMR_6271 | Transport, extracellular | 100.5155 | 243724.1106 | 100.5637 | 243718.0163 | 0.0481 | -1.00E-04 | 0.5 |
| HMR_6272 | Transport, extracellular | 100.5155 | 243724.1112 | 100.5637 | 243718.0178 | 0.0481 | -1.00E-04 | 0.5 |
| HMR_6273 | Transport, extracellular | 100.5156 | 243724.1118 | 100.5636 | 243718.0217 | 0.0481 | -1.00E-04 | 0.5 |
| HMR_6351 | Transport, extracellular | 8.2064 | 45.9384 | 7.8586 | 42.1259 | 0.3479 | 0.0371 | 0.4852 |
| HMR_6352 | Transport, extracellular | 8.2063 | 45.9392 | 7.8585 | 42.1266 | 0.3478 | 0.0371 | 0.4852 |

| | | | | | | | | |
|---|---|---|---|---|---|---|---|---|
| HMR_6353 | Transport, extracellular | 8.5068 | 73.506 | 8.1463 | 67.4173 | 0.3605 | 0.0304 | 0.4879 |
| HMR_6354 | Transport, extracellular | 12.8191 | 80.3988 | 12.2429 | 73.3662 | 0.5762 | 0.0465 | 0.4815 |
| HMR_6355 | Transport, extracellular | 4.254 | 18.3685 | 4.0737 | 16.8476 | 0.1803 | 0.0304 | 0.4879 |
| HMR_6356 | Transport, extracellular | 4.4081 | 17.0322 | 4.2213 | 15.6187 | 0.1868 | 0.0327 | 0.487 |
| HMR_6357 | Transport, extracellular | 4.2538 | 18.3667 | 4.0735 | 16.8461 | 0.1803 | 0.0304 | 0.4879 |
| HMR_6358 | Transport, extracellular | 4.4084 | 17.033 | 4.2216 | 15.6194 | 0.1868 | 0.0327 | 0.487 |
| HMR_6359 | Transport, extracellular | 4.4082 | 17.0325 | 4.2213 | 15.6189 | 0.1869 | 0.0327 | 0.487 |
| HMR_6360 | Transport, extracellular | 4.4082 | 17.0325 | 4.2214 | 15.6189 | 0.1869 | 0.0327 | 0.487 |
| HMR_6361 | Transport, extracellular | 4.254 | 18.3643 | 4.0737 | 16.8441 | 0.1803 | 0.0304 | 0.4879 |
| HMR_6362 | Transport, extracellular | 4.4084 | 17.0329 | 4.2215 | 15.6193 | 0.1868 | 0.0327 | 0.487 |
| HMR_6363 | Transport, extracellular | 4.2624 | 25.6041 | 4.0813 | 23.515 | 0.1811 | 0.0258 | 0.4897 |
| HMR_6364 | Transport, extracellular | 4.4086 | 17.0334 | 4.2217 | 15.6198 | 0.1868 | 0.0327 | 0.487 |
| HMR_6365 | Transport, extracellular | 5.7007 | 27.4941 | 5.453 | 25.1471 | 0.2477 | 0.0341 | 0.4864 |
| HMR_6366 | Transport, extracellular | 12.6132 | 54.2019 | 12.0786 | 49.7032 | 0.5346 | 0.0524 | 0.4791 |
| HMR_6367 | Transport, extracellular | 21.4724 | 117.1507 | 20.5214 | 107.0281 | 0.9511 | 0.0635 | 0.4747 |
| HMR_6368 | Transport, extracellular | 12.608 | 54.1858 | 12.0732 | 49.6872 | 0.5348 | 0.0525 | 0.4791 |
| HMR_6369 | Transport, extracellular | 9.7123 | 62.2385 | 9.3012 | 56.9768 | 0.4111 | 0.0377 | 0.485 |
| HMR_6371 | Transport, extracellular | 12.6645 | 152.1068 | 12.3107 | 143.833 | 0.3538 | 0.0206 | 0.4918 |
| HMR_6372 | Transport, extracellular | 12.6937 | 152.7723 | 12.3397 | 144.4762 | 0.354 | 0.0205 | 0.4918 |
| HMR_6373 | Transport, extracellular | 12.7058 | 153.0485 | 12.3515 | 144.7391 | 0.3543 | 0.0205 | 0.4918 |
| HMR_6374 | Transport, extracellular | 12.7063 | 153.0596 | 12.3487 | 144.6772 | 0.3575 | 0.0207 | 0.4917 |
| HMR_6375 | Transport, extracellular | 8.4278 | 72.0907 | 8.0759 | 66.2066 | 0.3519 | 0.0299 | 0.4881 |
| HMR_6377 | Transport, extracellular | 8.4102 | 71.7858 | 8.0602 | 65.9457 | 0.35 | 0.0298 | 0.4881 |
| HMR_6379 | Transport, extracellular | 8.4358 | 72.2282 | 8.0834 | 66.3303 | 0.3524 | 0.0299 | 0.4881 |
| HMR_6380 | Transport, extracellular | 21.4113 | 399.7501 | 20.1793 | 357.9245 | 1.232 | 0.0448 | 0.4822 |
| HMR_6381 | Transport, extracellular | 21.4126 | 399.7849 | 20.1766 | 357.8337 | 1.236 | 0.0449 | 0.4821 |
| HMR_6382 | Transport, extracellular | 21.4045 | 399.5219 | 20.1709 | 357.6558 | 1.2336 | 0.0448 | 0.4821 |
| HMR_6383 | Transport, extracellular | 21.4611 | 401.3833 | 20.2208 | 359.2282 | 1.2402 | 0.045 | 0.4821 |
| HMR_6384 | Transport, extracellular | 14.5346 | 196.9382 | 14.0904 | 185.276 | 0.4442 | 0.0227 | 0.4909 |
| HMR_6524 | Transport, extracellular | 8.4782 | 73.132 | 8.1029 | 66.8155 | 0.3753 | 0.0317 | 0.4873 |
| HMR_6525 | Transport, extracellular | 4.3284 | 22.2612 | 4.1402 | 20.3808 | 0.1883 | 0.0288 | 0.4885 |
| HMR_6526 | Transport, extracellular | 8.4527 | 33.3818 | 8.063 | 30.3749 | 0.3897 | 0.0488 | 0.4805 |
| HMR_6530 | Transport, extracellular | 39.5091 | 272.2601 | 39.1389 | 274.5711 | 0.3703 | 0.0158 | 0.4937 |
| HMR_6531 | Transport, extracellular | 20.6771 | 237.8771 | 20.8584 | 244.0577 | 0.1812 | -0.0083 | 0.5033 |
| HMR_6532 | Transport, extracellular | 4.3202 | 22.2218 | 4.1282 | 20.2985 | 0.192 | 0.0294 | 0.4883 |
| HMR_6534 | Transport, extracellular | 2.1661 | 5.5739 | 2.0729 | 5.1089 | 0.0932 | 0.0285 | 0.4886 |
| HMR_6732 | Transport, extracellular | 38.3985 | 285.8664 | 38.5761 | 283.8813 | 0.1776 | -0.0074 | 0.503 |
| HMR_6733 | Transport, extracellular | 4.254 | 18.3643 | 4.0737 | 16.8441 | 0.1803 | 0.0304 | 0.4879 |
| HMR_7253 | Transport, extracellular | -42.0842 | 614.1216 | -47.8579 | 789.8917 | 5.7737 | 0.1541 | 0.4388 |
| HMR_7568 | Transport, extracellular | 1.5074 | 1.3933 | 1.5433 | 1.4615 | 0.0359 | -0.0212 | 0.5085 |
| HMR_7569 | Transport, extracellular | 1.7017 | 1.1338 | 1.742 | 1.1885 | 0.0403 | -0.0264 | 0.5105 |
| HMR_7608 | Transport, extracellular | 355.812 | 12420.3153 | 555.2822 | 12637.3217 | 199.4702 | -1.2601 | 0.8962 |
| HMR_7609 | Transport, extracellular | -354.9605 | 12420.2303 | -554.4444 | 12637.4558 | 199.4839 | 1.2602 | 0.1038 |
| HMR_7633 | Transport, extracellular | 1.4234 | 2.0568 | 1.3588 | 1.8746 | 0.0646 | 0.0326 | 0.487 |

| | | | | | | | | |
|---|---|---|---|---|---|---|---|---|
| HMR_7634 | Transport, extracellular | 0.9013 | 0.2794 | 0.8898 | 0.268 | 0.0115 | 0.0156 | 0.4938 |
| HMR_7635 | Transport, extracellular | 1.0645 | 0.3427 | 1.0074 | 0.3071 | 0.057 | 0.0707 | 0.4718 |
| HMR_7636 | Transport, extracellular | 1.4224 | 2.0536 | 1.3579 | 1.8717 | 0.0645 | 0.0326 | 0.487 |
| HMR_7637 | Transport, extracellular | 1.4224 | 2.0536 | 1.3579 | 1.8717 | 0.0645 | 0.0326 | 0.487 |
| HMR_7643 | Transport, extracellular | 452.2186 | 158480.5776 | 407.2401 | 179538.459 | 44.9785 | 0.0774 | 0.4692 |
| HMR_7644 | Transport, extracellular | 392.2823 | 170052.0934 | 336.3279 | 193450.5203 | 55.9543 | 0.0928 | 0.463 |
| HMR_7645 | Transport, extracellular | 12.5363 | 149.1461 | 12.2069 | 141.4999 | 0.3294 | 0.0193 | 0.4923 |
| HMR_7650 | Transport, extracellular | 9.4682 | 6.7296 | 9.3963 | 6.4426 | 0.072 | 0.0198 | 0.4921 |
| HMR_7651 | Transport, extracellular | 1.7233 | 3.1177 | 1.6472 | 2.8431 | 0.076 | 0.0311 | 0.4876 |
| HMR_7679 | Transport, extracellular | 2.8581 | 8.513 | 2.7279 | 7.7284 | 0.1302 | 0.0323 | 0.4871 |
| HMR_7699 | Transport, extracellular | 4.2617 | 18.6859 | 4.0799 | 17.1234 | 0.1819 | 0.0304 | 0.4879 |
| HMR_7700 | Transport, extracellular | 2.8402 | 8.2347 | 2.7192 | 7.5489 | 0.1209 | 0.0304 | 0.4879 |
| HMR_7735 | Transport, extracellular | 1.4389 | 0.3104 | 1.3128 | 0.2604 | 0.1261 | 0.167 | 0.4337 |
| HMR_7736 | Transport, extracellular | 3.929 | 13.4971 | 4.1022 | 13.7726 | 0.1732 | -0.0332 | 0.5132 |
| HMR_7737 | Transport, extracellular | 2.8787 | 8.4178 | 2.7457 | 7.6588 | 0.133 | 0.0332 | 0.4868 |
| HMR_7738 | Transport, extracellular | 2.3919 | 3.7985 | 2.2289 | 3.4978 | 0.163 | 0.0603 | 0.4759 |
| HMR_7781 | Transport, extracellular | 10.7406 | 59.8476 | 10.6028 | 57.3349 | 0.1378 | 0.0127 | 0.4949 |
| HMR_7787 | Transport, extracellular | 7.1854 | 13.2895 | 6.6684 | 11.5859 | 0.5169 | 0.1036 | 0.4587 |
| HMR_7896 | Transport, extracellular | -9.0613 | 222.4859 | -10.8261 | 219.2931 | 1.7648 | 0.084 | 0.4665 |
| HMR_7898 | Transport, extracellular | -1.7551 | 1.0717 | -1.7612 | 1.0776 | 0.0061 | 0.0042 | 0.4983 |
| HMR_7900 | Transport, extracellular | -1.3396 | 0.5392 | -1.3444 | 0.543 | 0.0047 | 0.0046 | 0.4982 |
| HMR_7902 | Transport, extracellular | 2.2926 | 6.8743 | 2.1886 | 6.2103 | 0.104 | 0.0287 | 0.4885 |
| HMR_7905 | Transport, extracellular | -9.7136 | 96.5652 | -9.1144 | 88.8623 | 0.5992 | -0.044 | 0.5175 |
| HMR_7906 | Transport, extracellular | 1.6457 | 0.5408 | 1.5967 | 0.5101 | 0.0489 | 0.0477 | 0.481 |
| HMR_7986 | Transport, extracellular | 0.8968 | 0.1917 | 0.8845 | 0.1868 | 0.0123 | 0.02 | 0.492 |
| HMR_8031 | Transport, extracellular | -6.7678 | 12.3261 | -7.0057 | 13.2062 | 0.2379 | 0.0471 | 0.4812 |
| HMR_8033 | Transport, extracellular | 6.7678 | 12.3261 | 7.0057 | 13.2062 | 0.2379 | -0.0471 | 0.5188 |
| HMR_8075 | Transport, extracellular | 5.8765 | 38.8794 | 5.8422 | 38.4318 | 0.0343 | 0.0039 | 0.4984 |
| HMR_8340 | Transport, extracellular | -726.8843 | 18062.4276 | -489.2168 | 42958.3599 | 237.6675 | -0.9621 | 0.832 |
| HMR_8343 | Transport, extracellular | 726.8843 | 18062.4276 | 489.2168 | 42958.3599 | 237.6675 | 0.9621 | 0.168 |
| HMR_8354 | Transport, extracellular | 7.8016 | 37.6673 | 8.154 | 40.213 | 0.3524 | -0.0399 | 0.5159 |
| HMR_8364 | Transport, extracellular | 9.5662 | 27.5705 | 10.1701 | 31.1577 | 0.604 | -0.0788 | 0.5314 |
| HMR_8429 | Transport, extracellular | 4.2599 | 18.5858 | 4.0822 | 17.0648 | 0.1776 | 0.0297 | 0.4881 |
| HMR_8629 | Transport, extracellular | 6.9933 | 49.6476 | 6.7809 | 46.66 | 0.2124 | 0.0216 | 0.4914 |
| HMR_8676 | Transport, extracellular | 8.5156 | 33.8725 | 8.1551 | 31.0635 | 0.3605 | 0.0447 | 0.4822 |
| HMR_8677 | Transport, extracellular | 8.5107 | 33.8725 | 8.1499 | 31.0635 | 0.3608 | 0.0448 | 0.4821 |
| HMR_8687 | Transport, extracellular | 160.7384 | 10376.519 | 131.1897 | 8359.5839 | 29.5487 | 0.2159 | 0.4145 |
| HMR_8733 | Transport, extracellular | 236.4323 | 229669.9998 | 237.7932 | 229428.5165 | 1.3608 | -0.002 | 0.5008 |
| HMR_8734 | Transport, extracellular | 126.1888 | 236160.061 | 127.7654 | 235993.9766 | 1.5766 | -0.0023 | 0.5009 |
| HMR_8736 | Transport, extracellular | 20.9882 | 232.8285 | 21.27 | 238.867 | 0.2818 | -0.013 | 0.5052 |
| HMR_8847 | Transport, extracellular | 1.2034 | 1.1463 | 1.2149 | 1.1741 | 0.0115 | -0.0075 | 0.503 |
| HMR_8884 | Transport, extracellular | 4.254 | 18.3643 | 4.0737 | 16.8441 | 0.1803 | 0.0304 | 0.4879 |
| HMR_8892 | Transport, extracellular | 2.8405 | 8.1848 | 2.7159 | 7.4842 | 0.1246 | 0.0315 | 0.4874 |
| HMR_8911 | Transport, extracellular | 1.2968 | 1.1555 | 1.3174 | 1.1881 | 0.0206 | -0.0135 | 0.5054 |

| | | | | | | | | |
|---|---|---|---|---|---|---|---|---|
| HMR_8912 | Transport, extracellular | 1.1498 | 0.9742 | 1.1602 | 0.9924 | 0.0104 | -0.0074 | 0.503 |
| HMR_8917 | Transport, extracellular | 7.0161 | 49.9721 | 6.8021 | 46.9535 | 0.2139 | 0.0217 | 0.4913 |
| HMR_8925 | Transport, extracellular | 8.508 | 73.4574 | 8.1474 | 67.3763 | 0.3606 | 0.0304 | 0.4879 |
| HMR_8930 | Transport, extracellular | 0.0025 | 0.00E+00 | 0.0025 | 0.00E+00 | 0.00E+00 | 0.00E+00 | 0.5 |
| HMR_8932 | Transport, extracellular | 4.3315 | 22.3111 | 4.1474 | 20.4741 | 0.1841 | 0.0282 | 0.4888 |
| HMR_8933 | Transport, extracellular | 4.3235 | 22.2009 | 4.14 | 20.3774 | 0.1835 | 0.0281 | 0.4888 |
| HMR_8934 | Transport, extracellular | 2.8502 | 8.2469 | 2.7207 | 7.5154 | 0.1295 | 0.0326 | 0.487 |
| HMR_8935 | Transport, extracellular | 4.5389 | 6.881 | 4.3349 | 6.2762 | 0.204 | 0.0562 | 0.4776 |
| HMR_8938 | Transport, extracellular | 4.2338 | 18.19 | 4.0561 | 16.6979 | 0.1778 | 0.0301 | 0.488 |
| HMR_8939 | Transport, extracellular | 4.9535 | 17.3419 | 5.1398 | 19.1412 | 0.1863 | -0.0308 | 0.5123 |
| HMR_8940 | Transport, extracellular | 3.9065 | 10.6969 | 4.0804 | 11.7372 | 0.1739 | -0.0367 | 0.5146 |
| HMR_8941 | Transport, extracellular | 3.263 | 7.2804 | 3.3629 | 7.7146 | 0.0999 | -0.0258 | 0.5103 |
| HMR_9173 | Transport, extracellular | -1.6832 | 0.9914 | -1.6928 | 1.0017 | 0.0096 | 0.0068 | 0.4973 |
| HMR_9175 | Transport, extracellular | -6.6654 | 8.9011 | -6.6827 | 8.952 | 0.0173 | 0.0041 | 0.4984 |
| HMR_9176 | Transport, extracellular | -0.9035 | 0.1921 | -0.8915 | 0.1872 | 0.012 | -0.0195 | 0.5078 |
| HMR_9183 | Transport, extracellular | -17.9532 | 90.1572 | -17.1975 | 82.812 | 0.7556 | -0.0575 | 0.5229 |
| HMR_9190 | Transport, extracellular | 317.7012 | 59952.9733 | 329.9959 | 62697.1969 | 12.2947 | -0.0351 | 0.514 |
| HMR_9191 | Transport, extracellular | 362.6211 | 68676.1299 | 365.5586 | 69121.875 | 2.9376 | -0.0079 | 0.5032 |
| HMR_9197 | Transport, extracellular | 12.5073 | 80.4782 | 12.1678 | 75.7166 | 0.3395 | 0.0272 | 0.4892 |
| HMR_9198 | Transport, extracellular | 2.8449 | 8.2143 | 2.7158 | 7.487 | 0.129 | 0.0326 | 0.487 |
| HMR_9590 | Transport, extracellular | 6.4564 | 39.3033 | 6.267 | 37.0634 | 0.1893 | 0.0217 | 0.4914 |
| HMR_9591 | Transport, extracellular | 4.2254 | 18.123 | 4.044 | 16.6038 | 0.1814 | 0.0308 | 0.4877 |
| HMR_9592 | Transport, extracellular | 3.8328 | 14.8991 | 3.6921 | 13.8251 | 0.1407 | 0.0263 | 0.4895 |
| HMR_9593 | Transport, extracellular | 3.8507 | 15.0398 | 3.7097 | 13.9581 | 0.141 | 0.0262 | 0.4896 |
| HMR_9594 | Transport, extracellular | 4.2289 | 18.1535 | 4.047 | 16.6282 | 0.1819 | 0.0309 | 0.4877 |
| HMR_9595 | Transport, extracellular | 4.2307 | 18.1695 | 4.0487 | 16.6422 | 0.1821 | 0.0309 | 0.4877 |
| HMR_9596 | Transport, extracellular | 3.8397 | 14.9525 | 3.6984 | 13.8725 | 0.1412 | 0.0263 | 0.4895 |
| HMR_9597 | Transport, extracellular | 4.2296 | 18.1599 | 4.0477 | 16.6341 | 0.182 | 0.0308 | 0.4877 |
| HMR_9598 | Transport, extracellular | 4.2299 | 18.1616 | 4.047 | 16.6282 | 0.1829 | 0.031 | 0.4876 |
| HMR_9599 | Transport, extracellular | 4.2224 | 18.0969 | 4.0411 | 16.5793 | 0.1813 | 0.0308 | 0.4877 |
| HMR_9600 | Transport, extracellular | 4.2268 | 18.1354 | 4.0451 | 16.6122 | 0.1818 | 0.0308 | 0.4877 |
| HMR_9601 | Transport, extracellular | 3.6174 | 14.7017 | 3.3307 | 12.777 | 0.2867 | 0.0547 | 0.4782 |
| HMR_9607 | Transport, extracellular | 0.0026 | 0.00E+00 | 0.0026 | 0.00E+00 | 0.00E+00 | 0.00E+00 | 0.5 |
| HMR_9610 | Transport, extracellular | 2.6499 | 5.2172 | 2.6732 | 5.2961 | 0.0234 | -0.0072 | 0.5029 |
| HMR_9613 | Transport, extracellular | 3.7236 | 12.138 | 3.6995 | 12.0503 | 0.024 | 0.0049 | 0.4981 |
| HMR_9614 | Transport, extracellular | 8.508 | 73.4574 | 8.1474 | 67.3763 | 0.3606 | 0.0304 | 0.4879 |
| HMR_9615 | Transport, extracellular | 8.508 | 73.4574 | 8.1474 | 67.3763 | 0.3606 | 0.0304 | 0.4879 |
| HMR_9616 | Transport, extracellular | 8.3799 | 71.4468 | 8.0157 | 65.3836 | 0.3641 | 0.0311 | 0.4876 |
| HMR_9617 | Transport, extracellular | 8.3984 | 71.769 | 8.0334 | 65.6786 | 0.3649 | 0.0311 | 0.4876 |
| HMR_9618 | Transport, extracellular | 957.4382 | 362.4168 | 956.3055 | 381.6555 | 1.1328 | 0.0415 | 0.4834 |
| HMR_9623 | Transport, extracellular | 1.7847 | 2.012 | 1.8358 | 2.1184 | 0.0511 | -0.0251 | 0.51 |
| HMR_9624 | Transport, extracellular | 1.2501 | 1.2224 | 1.2101 | 1.1512 | 0.04 | 0.026 | 0.4896 |
| HMR_0446 | Transport, mitochondrial | 3.3514 | 3.0162 | 3.29 | 2.9057 | 0.0614 | 0.0252 | 0.4899 |
| HMR_0483 | Transport, mitochondrial | 3.9276 | 11.1364 | 3.8097 | 10.6028 | 0.1179 | 0.0253 | 0.4899 |

| | | | | | | | | |
|---|---|---|---|---|---|---|---|---|
| HMR_1145 | Transport, mitochondrial | 0.7474 | 0.1326 | 0.8133 | 0.1543 | 0.0659 | -0.123 | 0.5489 |
| HMR_1173 | Transport, mitochondrial | 0.7289 | 0.116 | 0.776 | 0.1302 | 0.0471 | -0.0949 | 0.5378 |
| HMR_1187 | Transport, mitochondrial | 0.7474 | 0.1326 | 0.8133 | 0.1543 | 0.0659 | -0.123 | 0.5489 |
| HMR_1215 | Transport, mitochondrial | 0.7289 | 0.116 | 0.776 | 0.1302 | 0.0471 | -0.0949 | 0.5378 |
| HMR_1267 | Transport, mitochondrial | -491.6879 | 407.676 | -505.9255 | 435.6018 | 14.2376 | 0.4903 | 0.312 |
| HMR_1284 | Transport, mitochondrial | 491.6877 | 407.6887 | 505.9253 | 435.6146 | 14.2376 | -0.4903 | 0.688 |
| HMR_1572 | Transport, mitochondrial | 7.6321 | 89.1301 | 7.1063 | 80.6827 | 0.5258 | 0.0403 | 0.4839 |
| HMR_1915 | Transport, mitochondrial | -279.9101 | 202471.2402 | -279.9333 | 202468.1567 | 0.0232 | 0.00E+00 | 0.5 |
| HMR_1925 | Transport, mitochondrial | 472.1054 | 48381.2217 | 472.1015 | 48381.0754 | 0.0039 | 0.00E+00 | 0.5 |
| HMR_3746 | Transport, mitochondrial | 317.6473 | 2872.6132 | 3.1205 | 2.9051 | 314.5268 | 5.8654 | 0.00E+00 |
| HMR_3762 | Transport, mitochondrial | -37.01 | 381.3218 | -2.8813 | 1.8327 | 34.1288 | -1.7435 | 0.9594 |
| HMR_3789 | Transport, mitochondrial | 37.0111 | 381.3205 | 2.8885 | 1.8372 | 34.1227 | 1.7432 | 0.0406 |
| HMR_3818 | Transport, mitochondrial | -83.1239 | 2349.4168 | -153.9887 | 6490.2349 | 70.8648 | 0.7537 | 0.2255 |
| HMR_3825 | Transport, mitochondrial | 51.2964 | 2496.698 | 61.1877 | 3519.8656 | 9.8913 | -0.1275 | 0.5507 |
| HMR_3863 | Transport, mitochondrial | 75.3605 | 1744.6765 | 92.2088 | 2540.5405 | 16.8483 | -0.2574 | 0.6016 |
| HMR_3864 | Transport, mitochondrial | 48.5968 | 1595.7538 | 58.059 | 2306.7484 | 9.4622 | -0.1515 | 0.5602 |
| HMR_3946 | Transport, mitochondrial | 16.6073 | 688.662 | 13.0597 | 591.9757 | 3.5476 | 0.0991 | 0.4605 |
| HMR_3949 | Transport, mitochondrial | 455.3691 | 69093.5599 | 421.4984 | 67265.9145 | 33.8707 | 0.0917 | 0.4635 |
| HMR_3950 | Transport, mitochondrial | 297.1823 | 1803.0401 | 392.9233 | 1608.4393 | 95.741 | -1.6392 | 0.9494 |
| HMR_3952 | Transport, mitochondrial | -619.5795 | 24279.5093 | -668.9211 | 20215.1917 | 49.3416 | 0.2339 | 0.4075 |
| HMR_3954 | Transport, mitochondrial | 55.1751 | 201.8811 | 63.383 | 295.4575 | 8.2079 | -0.3681 | 0.6436 |
| HMR_3971 | Transport, mitochondrial | -88.0795 | 280725.0211 | -108.3228 | 278531.3248 | 20.2433 | 0.0271 | 0.4892 |
| HMR_4184 | Transport, mitochondrial | 6.2375 | 43.5009 | 5.559 | 34.0989 | 0.6785 | 0.077 | 0.4693 |
| HMR_4237 | Transport, mitochondrial | 1.189 | 0.4145 | 1.1696 | 0.4019 | 0.0194 | 0.0214 | 0.4915 |
| HMR_4367 | Transport, mitochondrial | -3.346 | 3.0157 | -3.2843 | 2.905 | 0.0617 | -0.0254 | 0.5101 |
| HMR_4575 | Transport, mitochondrial | 0.9797 | 0.2217 | 0.9463 | 0.2072 | 0.0334 | 0.0509 | 0.4797 |
| HMR_4692 | Transport, mitochondrial | 12.5347 | 34.2931 | 13.4366 | 38.2753 | 0.9019 | -0.1059 | 0.5422 |
| HMR_4720 | Transport, mitochondrial | 748.6524 | 12874.1365 | 762.7361 | 11521.6982 | 14.0836 | -0.0902 | 0.5359 |
| HMR_4729 | Transport, mitochondrial | -748.652 | 12874.137 | -762.7357 | 11521.6985 | 14.0837 | 0.0902 | 0.4641 |
| HMR_4738 | Transport, mitochondrial | -7.6462 | 23.8484 | -7.6356 | 23.1941 | 0.0106 | -0.0015 | 0.5006 |
| HMR_4743 | Transport, mitochondrial | -11.2366 | 19.8939 | -10.1728 | 16.6565 | 1.0639 | -0.176 | 0.5698 |
| HMR_4754 | Transport, mitochondrial | 0.7151 | 0.2022 | 0.6511 | 0.1678 | 0.064 | 0.1053 | 0.4581 |
| HMR_4756 | Transport, mitochondrial | 0.7193 | 0.2026 | 0.656 | 0.1682 | 0.0633 | 0.104 | 0.4586 |
| HMR_4760 | Transport, mitochondrial | -1.4383 | 0.3022 | -1.312 | 0.2523 | 0.1262 | -0.1695 | 0.5673 |
| HMR_4843 | Transport, mitochondrial | 929.9073 | 4399.4961 | 922.0636 | 5398.4689 | 7.8438 | 0.0792 | 0.4684 |
| HMR_4850 | Transport, mitochondrial | 2.7518 | 5.8902 | 2.7932 | 6.0731 | 0.0414 | -0.012 | 0.5048 |
| HMR_4851 | Transport, mitochondrial | 82.135 | 262343.3183 | 95.3082 | 260813.9518 | 13.1732 | -0.0182 | 0.5073 |
| HMR_4852 | Transport, mitochondrial | 133.6729 | 259450.2847 | 142.7941 | 258019.5148 | 9.1213 | -0.0127 | 0.5051 |
| HMR_4854 | Transport, mitochondrial | 17.1273 | 285346.6957 | 21.5202 | 285139.7513 | 4.3929 | -0.0058 | 0.5023 |
| HMR_4855 | Transport, mitochondrial | 62.8735 | 293243.6613 | 63.9827 | 293110.5476 | 1.1092 | -0.0014 | 0.5006 |
| HMR_4862 | Transport, mitochondrial | -70.94 | 282708.6457 | -87.0144 | 281171.2981 | 16.0744 | 0.0214 | 0.4915 |
| HMR_4863 | Transport, mitochondrial | 21.2456 | 271967.7577 | 21.0544 | 271904.2567 | 0.1912 | 0.0003 | 0.4999 |
| HMR_4864 | Transport, mitochondrial | 24.9066 | 271893.2532 | 26.4934 | 271788.5021 | 1.5867 | -0.0022 | 0.5009 |
| HMR_4865 | Transport, mitochondrial | -21.4964 | 287221.7343 | -40.7272 | 286371.1456 | 19.2308 | 0.0254 | 0.4899 |

| | | | | | | | | |
|---|---|---|---|---|---|---|---|---|
| HMR_4867 | Transport, mitochondrial | -31.423 | 275491.9485 | -28.231 | 275546.4173 | 3.192 | -0.0043 | 0.5017 |
| HMR_4868 | Transport, mitochondrial | -27.8459 | 275600.0679 | -22.9136 | 275686.1767 | 4.9322 | -0.0066 | 0.5027 |
| HMR_4870 | Transport, mitochondrial | -40.1206 | 255059.7761 | -52.6665 | 254347.5837 | 12.546 | 0.0176 | 0.493 |
| HMR_4871 | Transport, mitochondrial | -22.1482 | 251481.295 | -29.4317 | 251219.474 | 7.2836 | 0.0103 | 0.4959 |
| HMR_4872 | Transport, mitochondrial | -17.9728 | 251543.2365 | -23.2352 | 251340.3727 | 5.2625 | 0.0074 | 0.497 |
| HMR_4874 | Transport, mitochondrial | 958.335 | 1234.0723 | 881.6842 | 4162.2979 | 76.6508 | 1.0434 | 0.1484 |
| HMR_4888 | Transport, mitochondrial | -971.235 | 753.6263 | -974.5624 | 615.0006 | 3.3274 | 0.0899 | 0.4642 |
| HMR_4898 | Transport, mitochondrial | 573.3187 | 842.8721 | 547.8776 | 585.9691 | 25.4411 | 0.673 | 0.2505 |
| HMR_4905 | Transport, mitochondrial | 155.151 | 1665.0732 | 129.5473 | 1382.6358 | 25.6036 | 0.4638 | 0.3214 |
| HMR_4914 | Transport, mitochondrial | 366.0316 | 153535.7767 | 380.5436 | 151047.9825 | 14.512 | -0.0263 | 0.5105 |
| HMR_4922 | Transport, mitochondrial | -821.9209 | 13700.6522 | -831.4615 | 12391.1729 | 9.5406 | 0.0591 | 0.4765 |
| HMR_4926 | Transport, mitochondrial | 425.2217 | 67850.8365 | 408.1788 | 66618.1158 | 17.0429 | 0.0465 | 0.4815 |
| HMR_4940 | Transport, mitochondrial | 91.6267 | 224730.0636 | 106.985 | 221810.784 | 15.3583 | -0.023 | 0.5092 |
| HMR_4944 | Transport, mitochondrial | 4.7871 | 7.8146 | 5.0367 | 8.589 | 0.2496 | -0.0616 | 0.5246 |
| HMR_4953 | Transport, mitochondrial | 15.0202 | 367.4555 | 9.6891 | 262.2098 | 5.3312 | 0.2125 | 0.4159 |
| HMR_4963 | Transport, mitochondrial | 25.9615 | 578.8992 | 29.5796 | 821.8522 | 3.6181 | -0.0967 | 0.5385 |
| HMR_4964 | Transport, mitochondrial | 4.0924 | 20.5579 | 3.9756 | 19.3522 | 0.1168 | 0.0185 | 0.4926 |
| HMR_4971 | Transport, mitochondrial | 36.9856 | 259224.3183 | 37.4179 | 259207.4578 | 0.4323 | -0.0006 | 0.5002 |
| HMR_4972 | Transport, mitochondrial | -71.8081 | 277486.6992 | -72.2804 | 277455.9192 | 0.4723 | 0.0006 | 0.4997 |
| HMR_4977 | Transport, mitochondrial | 371.2317 | 57781.7122 | 317.0525 | 49111.2593 | 54.1792 | 0.1657 | 0.4342 |
| HMR_4997 | Transport, mitochondrial | 2.9912 | 3.1773 | 2.8403 | 2.8656 | 0.1509 | 0.0614 | 0.4755 |
| HMR_4998 | Transport, mitochondrial | 2.991 | 3.1773 | 2.8398 | 2.8655 | 0.1512 | 0.0615 | 0.4755 |
| HMR_5006 | Transport, mitochondrial | 322.4007 | 23217.1301 | 276.0024 | 18973.7199 | 46.3983 | 0.2259 | 0.4106 |
| HMR_5015 | Transport, mitochondrial | 28.4445 | 761.9951 | 31.0702 | 921.1859 | 2.6258 | -0.064 | 0.5255 |
| HMR_5016 | Transport, mitochondrial | 147.3204 | 15138.7626 | 263.4443 | 37875.3464 | 116.1238 | -0.5043 | 0.693 |
| HMR_5031 | Transport, mitochondrial | 28.3494 | 135.3312 | 30.9735 | 138.3095 | 2.6241 | -0.1586 | 0.563 |
| HMR_5033 | Transport, mitochondrial | 1.2301 | 31.6204 | 0.8863 | 30.8929 | 0.3439 | 0.0435 | 0.4827 |
| HMR_5043 | Transport, mitochondrial | 982.2308 | 298.1853 | 986.9795 | 197.7808 | 4.7487 | -0.2132 | 0.5844 |
| HMR_5046 | Transport, mitochondrial | -192.195 | 196813.9838 | -192.1692 | 196813.3419 | 0.0259 | 0.00E+00 | 0.5 |
| HMR_5099 | Transport, mitochondrial | 1.551 | 0.8298 | 1.5081 | 0.7852 | 0.043 | 0.0338 | 0.4865 |
| HMR_5101 | Transport, mitochondrial | 30.4862 | 335.4011 | 34.02 | 436.7109 | 3.5339 | -0.1272 | 0.5506 |
| HMR_5102 | Transport, mitochondrial | 23.5652 | 329.0749 | 27.5836 | 432.152 | 4.0184 | -0.1456 | 0.5579 |
| HMR_5105 | Transport, mitochondrial | 21.6674 | 420.5663 | 25.3654 | 575.0644 | 3.6981 | -0.1172 | 0.5466 |
| HMR_5107 | Transport, mitochondrial | 10.0702 | 18.6729 | 2.1245 | 0.6608 | 7.9457 | 1.8071 | 0.0354 |
| HMR_5109 | Transport, mitochondrial | 5.3653 | 5.7539 | 5.1179 | 5.0284 | 0.2474 | 0.0754 | 0.47 |
| HMR_5113 | Transport, mitochondrial | 21.0726 | 237.1716 | 15.2414 | 130.6553 | 5.8311 | 0.304 | 0.3805 |
| HMR_5114 | Transport, mitochondrial | 29.6256 | 402.0208 | 25.9246 | 310.8319 | 3.7011 | 0.1386 | 0.4449 |
| HMR_5115 | Transport, mitochondrial | 25.2239 | 295.6533 | 29.3951 | 401.3994 | 4.1712 | -0.158 | 0.5628 |
| HMR_5116 | Transport, mitochondrial | 25.2239 | 295.6533 | 29.3951 | 401.3994 | 4.1712 | -0.158 | 0.5628 |
| HMR_5118 | Transport, mitochondrial | 12.4185 | 71.7494 | 2.6711 | 2.5596 | 9.7474 | 1.1308 | 0.1291 |
| HMR_5122 | Transport, mitochondrial | 81.4758 | 2833.1352 | 159.0627 | 6307.8511 | 77.5869 | -0.8115 | 0.7915 |
| HMR_5124 | Transport, mitochondrial | -294.8093 | 1631.2119 | -376.252 | 1434.0944 | 81.4427 | 1.471 | 0.0706 |
| HMR_5125 | Transport, mitochondrial | 23.7521 | 528.2807 | 28.9089 | 1081.9637 | 5.1568 | -0.1285 | 0.5511 |
| HMR_5126 | Transport, mitochondrial | 877.4688 | 3658.2459 | 709.675 | 8219.0351 | 167.7938 | 1.5396 | 0.0618 |

| | | | | | | | | |
|---|---|---|---|---|---|---|---|---|
| HMR_5225 | Transport, mitochondrial | 472.1432 | 48381.2206 | 472.1413 | 48381.0758 | 0.0019 | 0.00E+00 | 0.5 |
| HMR_5292 | Transport, mitochondrial | -52.8684 | 653.8259 | -56.5298 | 716.2558 | 3.6615 | 0.0989 | 0.4606 |
| HMR_5304 | Transport, mitochondrial | -128.3827 | 1768.2048 | -102.0069 | 1486.7677 | 26.3758 | -0.4623 | 0.6781 |
| HMR_5348 | Transport, mitochondrial | 37.6073 | 442.123 | 52.2534 | 691.6961 | 14.6462 | -0.435 | 0.6682 |
| HMR_5349 | Transport, mitochondrial | 23.8187 | 390.5346 | 27.7668 | 592.734 | 3.9481 | -0.1259 | 0.5501 |
| HMR_5361 | Transport, mitochondrial | 118.5549 | 153495.9614 | 118.7485 | 151064.2733 | 0.1936 | -0.0004 | 0.5001 |
| HMR_5411 | Transport, mitochondrial | -419.0729 | 80299.3969 | -275.1234 | 96376.3014 | 143.9494 | -0.3425 | 0.634 |
| HMR_5420 | Transport, mitochondrial | 1.4373 | 0.3038 | 1.3109 | 0.254 | 0.1263 | 0.1691 | 0.4328 |
| HMR_5426 | Transport, mitochondrial | -518.5006 | 63542.669 | -577.3467 | 64848.7007 | 58.8461 | 0.1642 | 0.4348 |
| HMR_6276 | Transport, mitochondrial | 34.0877 | 259290.6999 | 34.568 | 259273.7104 | 0.4803 | -0.0007 | 0.5003 |
| HMR_6277 | Transport, mitochondrial | -47.4205 | 258825.2552 | -47.4831 | 258820.4844 | 0.0625 | 1.00E-04 | 0.5 |
| HMR_6286 | Transport, mitochondrial | 4.0876 | 20.501 | 3.9708 | 19.2975 | 0.1168 | 0.0185 | 0.4926 |
| HMR_6287 | Transport, mitochondrial | 4.0904 | 20.5369 | 3.9739 | 19.3344 | 0.1165 | 0.0185 | 0.4926 |
| HMR_6288 | Transport, mitochondrial | 4.1216 | 14.998 | 4.0042 | 14.1582 | 0.1173 | 0.0217 | 0.4913 |
| HMR_6289 | Transport, mitochondrial | 4.0903 | 20.5363 | 3.9733 | 19.3298 | 0.117 | 0.0185 | 0.4926 |
| HMR_6290 | Transport, mitochondrial | 4.093 | 20.5723 | 3.9764 | 19.3668 | 0.1167 | 0.0185 | 0.4926 |
| HMR_6291 | Transport, mitochondrial | 4.095 | 20.5933 | 3.9781 | 19.3847 | 0.1169 | 0.0185 | 0.4926 |
| HMR_6292 | Transport, mitochondrial | 4.1242 | 15.0147 | 4.0067 | 14.1736 | 0.1175 | 0.0217 | 0.4913 |
| HMR_6293 | Transport, mitochondrial | 4.0887 | 20.5176 | 3.9718 | 19.3128 | 0.1169 | 0.0185 | 0.4926 |
| HMR_6294 | Transport, mitochondrial | 4.0915 | 20.5535 | 3.9749 | 19.3497 | 0.1166 | 0.0185 | 0.4926 |
| HMR_6295 | Transport, mitochondrial | 4.0934 | 20.5745 | 3.9766 | 19.3675 | 0.1168 | 0.0185 | 0.4926 |
| HMR_6296 | Transport, mitochondrial | 4.1227 | 15.0046 | 4.0053 | 14.1644 | 0.1174 | 0.0217 | 0.4913 |
| HMR_6297 | Transport, mitochondrial | -44.5347 | 258914.6127 | -44.6448 | 258908.2242 | 0.1101 | 0.0002 | 0.4999 |
| HMR_6298 | Transport, mitochondrial | 3.0629 | 252518.9448 | 3.0127 | 252514.7209 | 0.0502 | 1.00E-04 | 0.5 |
| HMR_6299 | Transport, mitochondrial | 33.9823 | 945.3418 | 39.7912 | 1291.625 | 5.809 | -0.1228 | 0.5489 |
| HMR_6300 | Transport, mitochondrial | 54.9432 | 1969.6448 | 63.4717 | 2639.777 | 8.5285 | -0.1256 | 0.55 |
| HMR_6301 | Transport, mitochondrial | 45.2389 | 1461.5702 | 52.9946 | 2004.8941 | 7.7558 | -0.1317 | 0.5524 |
| HMR_6302 | Transport, mitochondrial | 38.6579 | 1146.7297 | 44.8758 | 1544.7438 | 6.218 | -0.1199 | 0.5477 |
| HMR_6303 | Transport, mitochondrial | 33.9823 | 945.3418 | 39.7912 | 1291.6249 | 5.809 | -0.1228 | 0.5489 |
| HMR_6304 | Transport, mitochondrial | 54.9432 | 1969.6449 | 63.4717 | 2639.7771 | 8.5285 | -0.1256 | 0.55 |
| HMR_6305 | Transport, mitochondrial | 39.7204 | 1176.7045 | 46.502 | 1607.4476 | 6.7816 | -0.1285 | 0.5511 |
| HMR_6306 | Transport, mitochondrial | 42.9083 | 1323.9191 | 49.7144 | 1779.9987 | 6.8061 | -0.1222 | 0.5486 |
| HMR_6307 | Transport, mitochondrial | 62.3693 | 2419.7065 | 72.7146 | 3314.9462 | 10.3454 | -0.1366 | 0.5543 |
| HMR_6308 | Transport, mitochondrial | 32.1251 | 871.7823 | 37.5194 | 1186.662 | 5.3944 | -0.1189 | 0.5473 |
| HMR_6309 | Transport, mitochondrial | 43.9209 | 1529.1661 | 51.1335 | 2068.6802 | 7.2126 | -0.1202 | 0.5479 |
| HMR_6310 | Transport, mitochondrial | 43.9209 | 1529.1663 | 51.1335 | 2068.6804 | 7.2126 | -0.1202 | 0.5479 |
| HMR_6311 | Transport, mitochondrial | 52.198 | 1826.1003 | 60.7404 | 2474.1868 | 8.5424 | -0.1303 | 0.5518 |
| HMR_6312 | Transport, mitochondrial | 35.0958 | 995.5219 | 40.9243 | 1351.2844 | 5.8286 | -0.1203 | 0.5479 |
| HMR_6313 | Transport, mitochondrial | 32.1251 | 871.7822 | 37.5194 | 1186.6619 | 5.3944 | -0.1189 | 0.5473 |
| HMR_6314 | Transport, mitochondrial | 62.3693 | 2419.7066 | 72.7146 | 3314.9463 | 10.3454 | -0.1366 | 0.5543 |
| HMR_6315 | Transport, mitochondrial | 52.198 | 1826.1004 | 60.7404 | 2474.187 | 8.5424 | -0.1303 | 0.5518 |
| HMR_6316 | Transport, mitochondrial | 35.0958 | 995.5219 | 40.9243 | 1351.2843 | 5.8286 | -0.1203 | 0.5479 |
| HMR_6317 | Transport, mitochondrial | 47.105 | 1545.6716 | 54.8981 | 2106.0865 | 7.7931 | -0.129 | 0.5513 |
| HMR_6318 | Transport, mitochondrial | 37.2227 | 1076.3281 | 43.4364 | 1462.9471 | 6.2137 | -0.1233 | 0.5491 |

| | | | | | | | | |
|---|---|---|---|---|---|---|---|---|
| HMR_6321 | Transport, mitochondrial | 22.5411 | 447.6277 | 26.433 | 613.4642 | 3.8919 | -0.1195 | 0.5476 |
| HMR_6323 | Transport, mitochondrial | 95.2981 | 3761.8924 | 110.2334 | 5121.9152 | 14.9353 | -0.1585 | 0.563 |
| HMR_6324 | Transport, mitochondrial | 26.9906 | 654.7154 | 31.5421 | 891.3456 | 4.5515 | -0.1158 | 0.5461 |
| HMR_6325 | Transport, mitochondrial | 95.2981 | 3761.8924 | 110.2334 | 5121.9151 | 14.9353 | -0.1585 | 0.563 |
| HMR_6326 | Transport, mitochondrial | 26.9906 | 654.7154 | 31.5421 | 891.3456 | 4.5515 | -0.1158 | 0.5461 |
| HMR_6327 | Transport, mitochondrial | 23.0588 | 466.0448 | 26.962 | 635.4711 | 3.9032 | -0.1176 | 0.5468 |
| HMR_6328 | Transport, mitochondrial | 935.2027 | 3842.8467 | 928.5278 | 4658.275 | 6.675 | 0.0724 | 0.4711 |
| HMR_6330 | Transport, mitochondrial | 142.9513 | 273835.1905 | 169.5932 | 269290.4797 | 26.6419 | -0.0362 | 0.5144 |
| HMR_6331 | Transport, mitochondrial | -12.3676 | 38.3557 | -12.0152 | 36.2184 | 0.3524 | -0.0408 | 0.5163 |
| HMR_6332 | Transport, mitochondrial | 11.0702 | 143.8298 | 9.3708 | 94.6211 | 1.6994 | 0.11 | 0.4562 |
| HMR_6333 | Transport, mitochondrial | 11.0698 | 143.8207 | 9.3705 | 94.6149 | 1.6993 | 0.1101 | 0.4562 |
| HMR_6334 | Transport, mitochondrial | 11.088 | 144.3981 | 9.3848 | 94.9417 | 1.7032 | 0.1101 | 0.4562 |
| HMR_6335 | Transport, mitochondrial | 9.5182 | 104.5445 | 8.3244 | 74.9491 | 1.1938 | 0.0891 | 0.4645 |
| HMR_6336 | Transport, mitochondrial | 9.5179 | 104.538 | 8.3241 | 74.944 | 1.1938 | 0.0891 | 0.4645 |
| HMR_6337 | Transport, mitochondrial | 9.5312 | 104.9133 | 8.3353 | 75.1854 | 1.1958 | 0.0891 | 0.4645 |
| HMR_6338 | Transport, mitochondrial | 9.4895 | 103.9678 | 8.2981 | 74.5332 | 1.1914 | 0.0892 | 0.4645 |
| HMR_6339 | Transport, mitochondrial | 9.5259 | 104.8079 | 8.328 | 75.0342 | 1.1979 | 0.0893 | 0.4644 |
| HMR_6340 | Transport, mitochondrial | 4.8034 | 21.6792 | 4.6138 | 20.2068 | 0.1896 | 0.0293 | 0.4883 |
| HMR_6341 | Transport, mitochondrial | 8.3094 | 80.3018 | 7.459 | 61.1224 | 0.8504 | 0.0715 | 0.4715 |
| HMR_6342 | Transport, mitochondrial | 8.3373 | 80.8727 | 7.483 | 61.5067 | 0.8544 | 0.0716 | 0.4715 |
| HMR_6343 | Transport, mitochondrial | 4.492 | 19.0738 | 4.3505 | 18.0433 | 0.1414 | 0.0232 | 0.4907 |
| HMR_6391 | Transport, mitochondrial | 115.1352 | 224725.037 | 139.7096 | 221831.4485 | 24.5744 | -0.0368 | 0.5147 |
| HMR_6513 | Transport, mitochondrial | 39.1797 | 589.9231 | 44.8579 | 871.6252 | 5.6783 | -0.1485 | 0.559 |
| HMR_6521 | Transport, mitochondrial | 45.9123 | 234730.2505 | 46.7224 | 234658.7576 | 0.8101 | -0.0012 | 0.5005 |
| HMR_6522 | Transport, mitochondrial | -45.912 | 234730.251 | -46.7219 | 234658.7583 | 0.81 | 0.0012 | 0.4995 |
| HMR_6618 | Transport, mitochondrial | -60.9937 | 190.4716 | -63.695 | 209.2007 | 2.7013 | 0.1351 | 0.4463 |
| HMR_6620 | Transport, mitochondrial | 60.9934 | 190.4716 | 63.6948 | 209.2007 | 2.7014 | -0.1351 | 0.5537 |
| HMR_6781 | Transport, mitochondrial | -1.5451 | 0.8286 | -1.502 | 0.7841 | 0.0431 | -0.0339 | 0.5135 |
| HMR_7638 | Transport, mitochondrial | 49.7581 | 3132.9094 | 57.95 | 4291.3052 | 8.1919 | -0.0951 | 0.5379 |
| HMR_7708 | Transport, mitochondrial | -12.0719 | 17.0381 | -11.6118 | 15.8291 | 0.4601 | -0.0802 | 0.532 |
| HMR_7719 | Transport, mitochondrial | -53.6353 | 172.9451 | -56.6475 | 193.2966 | 3.0122 | 0.1574 | 0.4375 |
| HMR_7804 | Transport, mitochondrial | 11.7607 | 273909.2312 | 12.655 | 274069.5795 | 0.8943 | -0.0012 | 0.5005 |
| HMR_7806 | Transport, mitochondrial | -1.3705 | 274001.0784 | -1.781 | 274183.2763 | 0.4105 | 0.0006 | 0.4998 |
| HMR_7808 | Transport, mitochondrial | -52.5057 | 272048.6286 | -63.9528 | 271439.8524 | 11.4471 | 0.0155 | 0.4938 |
| HMR_7810 | Transport, mitochondrial | -68.8887 | 271195.65 | -74.8831 | 270835.6062 | 5.9944 | 0.0081 | 0.4968 |
| HMR_7812 | Transport, mitochondrial | -67.5354 | 271282.658 | -73.129 | 270955.66 | 5.5935 | 0.0076 | 0.497 |
| HMR_7814 | Transport, mitochondrial | -55.9049 | 271912.5452 | -60.6371 | 271684.9022 | 4.7322 | 0.0064 | 0.4974 |
| HMR_7815 | Transport, mitochondrial | 11.0756 | 143.8142 | 9.3734 | 94.587 | 1.7022 | 0.1102 | 0.4561 |
| HMR_7816 | Transport, mitochondrial | 9.522 | 104.546 | 8.3263 | 74.9306 | 1.1956 | 0.0892 | 0.4644 |
| HMR_7818 | Transport, mitochondrial | -16.5597 | 273864.7153 | -11.1115 | 274073.975 | 5.4482 | -0.0074 | 0.5029 |
| HMR_7820 | Transport, mitochondrial | -15.1893 | 273896.0184 | -9.3298 | 274104.696 | 5.8595 | -0.0079 | 0.5032 |
| HMR_7822 | Transport, mitochondrial | -3.4373 | 273787.5405 | 3.3316 | 273885.0954 | 6.7689 | -0.0091 | 0.5036 |
| HMR_7824 | Transport, mitochondrial | -13.1307 | 273893.0996 | -14.4349 | 274051.5232 | 1.3043 | 0.0018 | 0.4993 |
| HMR_7825 | Transport, mitochondrial | 9.5204 | 104.6101 | 8.3263 | 74.991 | 1.1941 | 0.0891 | 0.4645 |

| | | | | | | | | |
|---|---|---|---|---|---|---|---|---|
| HMR_7826 | Transport, mitochondrial | 11.0733 | 143.9303 | 9.3732 | 94.6779 | 1.7 | 0.1101 | 0.4562 |
| HMR_7827 | Transport, mitochondrial | 8.136 | 56.7836 | 8.7035 | 64.8048 | 0.5675 | -0.0515 | 0.5205 |
| HMR_7828 | Transport, mitochondrial | 8.1288 | 56.6961 | 8.6936 | 64.675 | 0.5647 | -0.0513 | 0.5204 |
| HMR_7829 | Transport, mitochondrial | 8.1381 | 56.8065 | 8.705 | 64.8212 | 0.567 | -0.0514 | 0.5205 |
| HMR_7830 | Transport, mitochondrial | 63.2419 | 3649.6345 | 69.2646 | 4361.3283 | 6.0226 | -0.0673 | 0.5268 |
| HMR_7831 | Transport, mitochondrial | 8.1379 | 56.8044 | 8.7048 | 64.8179 | 0.5669 | -0.0514 | 0.5205 |
| HMR_7832 | Transport, mitochondrial | 8.1363 | 56.7853 | 8.7028 | 64.7928 | 0.5665 | -0.0514 | 0.5205 |
| HMR_7833 | Transport, mitochondrial | 9.4282 | 102.3396 | 8.2492 | 73.5308 | 1.179 | 0.0889 | 0.4646 |
| HMR_7834 | Transport, mitochondrial | 8.2624 | 79.1918 | 7.4195 | 60.3724 | 0.8429 | 0.0714 | 0.4716 |
| HMR_7835 | Transport, mitochondrial | 60.1753 | 3326.2047 | 65.4659 | 3926.4006 | 5.2906 | -0.0621 | 0.5248 |
| HMR_7836 | Transport, mitochondrial | 60.7143 | 3395.047 | 66.2216 | 4029.0936 | 5.5073 | -0.0639 | 0.5255 |
| HMR_7837 | Transport, mitochondrial | 60.6732 | 3373.4817 | 66.1074 | 3992.3858 | 5.4343 | -0.0633 | 0.5252 |
| HMR_7838 | Transport, mitochondrial | 60.6608 | 3372.045 | 66.0883 | 3989.9675 | 5.4275 | -0.0633 | 0.5252 |
| HMR_7839 | Transport, mitochondrial | 62.8037 | 3606.5363 | 68.6958 | 4300.5404 | 5.8921 | -0.0663 | 0.5264 |
| HMR_7840 | Transport, mitochondrial | 60.5794 | 3364.7337 | 65.9852 | 3980.0251 | 5.4057 | -0.0631 | 0.5251 |
| HMR_7841 | Transport, mitochondrial | 64.4528 | 3733.3421 | 70.9184 | 4502.4863 | 6.4656 | -0.0712 | 0.5284 |
| HMR_7842 | Transport, mitochondrial | 63.8165 | 3651.164 | 70.0851 | 4385.0464 | 6.2686 | -0.0699 | 0.5279 |
| HMR_7843 | Transport, mitochondrial | 64.369 | 3704.142 | 70.8089 | 4460.5059 | 6.4398 | -0.0713 | 0.5284 |
| HMR_7844 | Transport, mitochondrial | 64.3587 | 3703.2371 | 70.7934 | 4459.1224 | 6.4347 | -0.0712 | 0.5284 |
| HMR_7845 | Transport, mitochondrial | 8.3379 | 80.9547 | 7.4838 | 61.5544 | 0.8541 | 0.0715 | 0.4715 |
| HMR_7846 | Transport, mitochondrial | 9.5258 | 105.0063 | 8.3286 | 75.1339 | 1.1972 | 0.0892 | 0.4645 |
| HMR_7847 | Transport, mitochondrial | 64.7001 | 3782.136 | 71.1669 | 4556.0093 | 6.4668 | -0.0708 | 0.5282 |
| HMR_7848 | Transport, mitochondrial | 63.2022 | 3644.2933 | 69.2817 | 4361.4262 | 6.0796 | -0.0679 | 0.5271 |
| HMR_7849 | Transport, mitochondrial | 64.1507 | 3728.1733 | 70.4492 | 4479.1696 | 6.2985 | -0.0695 | 0.5277 |
| HMR_7850 | Transport, mitochondrial | 64.567 | 3768.0235 | 71.0664 | 4544.1838 | 6.4994 | -0.0713 | 0.5284 |
| HMR_7851 | Transport, mitochondrial | 64.91 | 3825.5292 | 71.4458 | 4618.6851 | 6.5359 | -0.0711 | 0.5284 |
| HMR_7852 | Transport, mitochondrial | 64.6054 | 3772.9905 | 71.0433 | 4542.9129 | 6.4378 | -0.0706 | 0.5281 |
| HMR_7853 | Transport, mitochondrial | 63.3432 | 3659.3335 | 69.3979 | 4375.1934 | 6.0547 | -0.0675 | 0.5269 |
| HMR_7854 | Transport, mitochondrial | 63.526 | 3699.3194 | 69.6332 | 4431.6024 | 6.1072 | -0.0677 | 0.527 |
| HMR_8356 | Transport, mitochondrial | 32.2477 | 353.1339 | 35.0413 | 400.9341 | 2.7936 | -0.1017 | 0.5405 |
| HMR_8365 | Transport, mitochondrial | 9.5664 | 27.5706 | 10.1704 | 31.1577 | 0.604 | -0.0788 | 0.5314 |
| HMR_8505 | Transport, mitochondrial | 9.2017 | 37.9895 | 8.8731 | 35.8964 | 0.3286 | 0.0382 | 0.4848 |
| HMR_8510 | Transport, mitochondrial | 23.4689 | 156.7442 | 32.685 | 276.3384 | 9.2161 | -0.4429 | 0.6711 |
| HMR_8513 | Transport, mitochondrial | -23.4745 | 156.75 | -32.6907 | 276.3466 | 9.2163 | 0.4429 | 0.3289 |
| HMR_8741 | Transport, mitochondrial | 41.7596 | 277660.2292 | 47.8155 | 277353.9806 | 6.0559 | -0.0081 | 0.5032 |
| HMR_8742 | Transport, mitochondrial | 38.2468 | 277785.753 | 42.6048 | 277567.625 | 4.358 | -0.0058 | 0.5023 |
| HMR_9174 | Transport, mitochondrial | 6.6651 | 8.8951 | 6.6824 | 8.946 | 0.0173 | -0.0041 | 0.5016 |
| HMR_0713 | Transport, peroxisomal | 479.0845 | 59496.2314 | 479.1713 | 59502.3439 | 0.0869 | -0.0003 | 0.5001 |
| HMR_1144 | Transport, peroxisomal | 0.7239 | 0.1204 | 0.7715 | 0.1345 | 0.0476 | -0.0943 | 0.5376 |
| HMR_1172 | Transport, peroxisomal | -0.7298 | 0.1102 | -0.7764 | 0.1243 | 0.0466 | 0.0962 | 0.4617 |
| HMR_1188 | Transport, peroxisomal | 0.7239 | 0.1204 | 0.7715 | 0.1345 | 0.0476 | -0.0943 | 0.5376 |
| HMR_1214 | Transport, peroxisomal | -0.7298 | 0.1102 | -0.7764 | 0.1243 | 0.0466 | 0.0962 | 0.4617 |
| HMR_1268 | Transport, peroxisomal | -185.3539 | 8138.5623 | -178.5189 | 7577.4113 | 6.835 | -0.0545 | 0.5217 |
| HMR_1283 | Transport, peroxisomal | 185.3543 | 8138.5632 | 178.5193 | 7577.4122 | 6.835 | 0.0545 | 0.4783 |

| | | | | | | | | |
|---|---|---|---|---|---|---|---|---|
| HMR_1443 | Transport, peroxisomal | -1.8267 | 0.9929 | -1.769 | 0.9397 | 0.0577 | -0.0415 | 0.5166 |
| HMR_2407 | Transport, peroxisomal | 0.9011 | 0.1716 | 0.9851 | 0.2028 | 0.084 | -0.1373 | 0.5546 |
| HMR_2509 | Transport, peroxisomal | 0.7456 | 0.1291 | 0.7927 | 0.1451 | 0.0471 | -0.09 | 0.5359 |
| HMR_3007 | Transport, peroxisomal | -74.9722 | 112.541 | -77.9557 | 117.6923 | 2.9835 | 0.1966 | 0.4221 |
| HMR_3008 | Transport, peroxisomal | 2.8452 | 8.2168 | 2.7163 | 7.4902 | 0.1289 | 0.0325 | 0.487 |
| HMR_3011 | Transport, peroxisomal | 1.7415 | 1.136 | 1.7253 | 1.1319 | 0.0162 | 0.0107 | 0.4957 |
| HMR_3013 | Transport, peroxisomal | 1.8985 | 0.6964 | 1.947 | 0.728 | 0.0485 | -0.0406 | 0.5162 |
| HMR_3014 | Transport, peroxisomal | 1.0807 | 0.2354 | 1.0982 | 0.2414 | 0.0175 | -0.0253 | 0.5101 |
| HMR_3017 | Transport, peroxisomal | 0.6824 | 0.1056 | 0.7047 | 0.1109 | 0.0223 | -0.0479 | 0.5191 |
| HMR_3018 | Transport, peroxisomal | 0.4064 | 0.0532 | 0.4148 | 0.055 | 0.0083 | -0.0254 | 0.5101 |
| HMR_3019 | Transport, peroxisomal | -2.3171 | 2.9222 | -2.3901 | 3.1305 | 0.0731 | 0.0297 | 0.4882 |
| HMR_3020 | Transport, peroxisomal | 0.6544 | 0.2266 | 0.6811 | 0.2448 | 0.0268 | -0.039 | 0.5156 |
| HMR_3021 | Transport, peroxisomal | 0.6801 | 0.1518 | 0.6987 | 0.1602 | 0.0186 | -0.0332 | 0.5133 |
| HMR_3022 | Transport, peroxisomal | 3.5657 | 2.6716 | 3.6917 | 2.8614 | 0.126 | -0.0536 | 0.5214 |
| HMR_3023 | Transport, peroxisomal | 0.7977 | 0.1348 | 0.8267 | 0.1443 | 0.029 | -0.0549 | 0.5219 |
| HMR_3024 | Transport, peroxisomal | 0.5157 | 0.1195 | 0.5319 | 0.1274 | 0.0162 | -0.0326 | 0.513 |
| HMR_3025 | Transport, peroxisomal | 0.2039 | 0.0183 | 0.2083 | 0.0191 | 0.0044 | -0.0228 | 0.5091 |
| HMR_3473 | Transport, peroxisomal | 483.1466 | 78986.3961 | 482.4985 | 78830.7628 | 0.6481 | 0.0016 | 0.4993 |
| HMR_3848 | Transport, peroxisomal | -114.2253 | 1561.1508 | -124.1591 | 1928.7314 | 9.9338 | 0.1682 | 0.4332 |
| HMR_3850 | Transport, peroxisomal | 114.2256 | 1561.1519 | 124.1594 | 1928.7319 | 9.9339 | -0.1682 | 0.5668 |
| HMR_3867 | Transport, peroxisomal | -114.2267 | 1561.139 | -124.1606 | 1928.7196 | 9.9339 | 0.1682 | 0.4332 |
| HMR_3947 | Transport, peroxisomal | -56.7948 | 8860.205 | -62.8748 | 9705.4111 | 6.08 | 0.0446 | 0.4822 |
| HMR_4393 | Transport, peroxisomal | 3.3392 | 3.8303 | 3.3678 | 3.8905 | 0.0287 | -0.0103 | 0.5041 |
| HMR_4647 | Transport, peroxisomal | -8.8445 | 25.8493 | -8.5653 | 24.1717 | 0.2791 | -0.0395 | 0.5157 |
| HMR_4890 | Transport, peroxisomal | 76.3262 | 8884.8955 | 82.9935 | 9729.5258 | 6.6673 | -0.0489 | 0.5195 |
| HMR_4900 | Transport, peroxisomal | 123.4754 | 3060.499 | 132.8153 | 3458.6311 | 9.3399 | -0.1157 | 0.546 |
| HMR_4915 | Transport, peroxisomal | 229.4103 | 223201.1674 | 289.6853 | 205416.621 | 60.2749 | -0.0921 | 0.5367 |
| HMR_4924 | Transport, peroxisomal | -12.9133 | 12.6226 | -13.1932 | 12.9391 | 0.2799 | 0.0554 | 0.4779 |
| HMR_4930 | Transport, peroxisomal | 74.9847 | 112.5059 | 77.968 | 117.6582 | 2.9833 | -0.1966 | 0.5779 |
| HMR_4945 | Transport, peroxisomal | -0.8292 | 0.1096 | -1.0653 | 0.1728 | 0.2361 | 0.4444 | 0.3284 |
| HMR_5344 | Transport, peroxisomal | -1.4175 | 0.8768 | -1.4046 | 0.8651 | 0.0129 | -0.0098 | 0.5039 |
| HMR_5346 | Transport, peroxisomal | -186.9406 | 8139.1353 | -180.4451 | 7577.7246 | 6.4955 | -0.0518 | 0.5207 |
| HMR_5362 | Transport, peroxisomal | 45.9518 | 172607.1192 | 74.1341 | 167618.9187 | 28.1822 | -0.0483 | 0.5193 |
| HMR_5414 | Transport, peroxisomal | -3.3373 | 3.8291 | -3.3661 | 3.8893 | 0.0288 | 0.0104 | 0.4959 |
| HMR_6986 | Transport, peroxisomal | 486.7591 | 59495.2008 | 486.7717 | 59501.217 | 0.0125 | 0.00E+00 | 0.5 |
| HMR_6987 | Transport, peroxisomal | -0.8305 | 0.1141 | -1.0665 | 0.1777 | 0.236 | 0.4369 | 0.3311 |
| HMR_7590 | Transport, peroxisomal | 1.0329 | 0.2257 | 1.019 | 0.2199 | 0.0139 | 0.0208 | 0.4917 |
| HMR_7596 | Transport, peroxisomal | -1.0285 | 0.217 | -1.0145 | 0.2113 | 0.0139 | -0.0213 | 0.5085 |
| HMR_7771 | Transport, peroxisomal | 1.6391 | 1.1341 | 1.6696 | 1.1777 | 0.0305 | -0.0201 | 0.508 |
| HMR_7775 | Transport, peroxisomal | 1.6293 | 1.1339 | 1.6599 | 1.1775 | 0.0306 | -0.0201 | 0.508 |
| HMR_7777 | Transport, peroxisomal | 1.7235 | 1.2093 | 1.7778 | 1.2812 | 0.0543 | -0.0344 | 0.5137 |
| HMR_7779 | Transport, peroxisomal | 1.7159 | 1.2091 | 1.7704 | 1.2811 | 0.0545 | -0.0345 | 0.5138 |
| HMR_7783 | Transport, peroxisomal | 295.4924 | 85128.2564 | 301.3457 | 84490.354 | 5.8532 | -0.0142 | 0.5057 |
| HMR_7785 | Transport, peroxisomal | -295.4927 | 85128.2562 | -301.346 | 84490.3538 | 5.8533 | 0.0142 | 0.4943 |

| | | | | | | | | |
|---|---|---|---|---|---|---|---|---|
| HMR_7797 | Transport, peroxisomal | -485.9261 | 59495.4219 | -485.7023 | 59501.5977 | 0.2238 | -0.0006 | 0.5003 |
| HMR_8754 | Transport, peroxisomal | 8.8517 | 25.8473 | 8.5728 | 24.1697 | 0.2789 | 0.0394 | 0.4843 |
| HMR_9678 | Transport, peroxisomal | 2.2069 | 0.9548 | 2.186 | 0.9411 | 0.0209 | 0.0152 | 0.4939 |
| HMR_1095 | Transport, nuclear | 3.702 | 3.8687 | 3.6947 | 3.8614 | 0.0073 | 0.0026 | 0.499 |
| HMR_4274 | Transport, nuclear | -247.7643 | 120417.9756 | -247.2109 | 120579.607 | 0.5534 | -0.0011 | 0.5004 |
| HMR_4275 | Transport, nuclear | 249.1853 | 120417.1317 | 248.6411 | 120578.7562 | 0.5442 | 0.0011 | 0.4996 |
| HMR_4277 | Transport, nuclear | 1.4323 | 0.7135 | 1.4417 | 0.7227 | 0.0093 | -0.0078 | 0.5031 |
| HMR_4535 | Transport, nuclear | 703.772 | 22942.268 | 703.5597 | 22971.3007 | 0.2123 | 0.001 | 0.4996 |
| HMR_4893 | Transport, nuclear | -8.6572 | 80.9674 | -10.5504 | 88.3556 | 1.8932 | 0.1455 | 0.4422 |
| HMR_5422 | Transport, nuclear | 452.2859 | 124850.0251 | 451.8313 | 125022.0456 | 0.4546 | 0.0009 | 0.4996 |
| HMR_5423 | Transport, nuclear | -703.8062 | 22942.2833 | -703.5944 | 22971.3159 | 0.2118 | -0.001 | 0.5004 |
| HMR_5424 | Transport, nuclear | -703.7766 | 22942.2688 | -703.5644 | 22971.3015 | 0.2122 | -0.001 | 0.5004 |
| HMR_6617 | Transport, nuclear | 15.501 | 68.2078 | 16.6575 | 78.6516 | 1.1565 | -0.0954 | 0.538 |
| HMR_6619 | Transport, nuclear | -15.5009 | 68.2078 | -16.6574 | 78.6516 | 1.1565 | 0.0954 | 0.462 |
| HMR_7712 | Transport, nuclear | -1.4318 | 0.7135 | -1.4411 | 0.7227 | 0.0093 | 0.0078 | 0.4969 |
| HMR_7764 | Transport, nuclear | -9.6618 | 43.513 | -8.1405 | 30.9451 | 1.5213 | -0.1763 | 0.57 |
| HMR_7773 | Transport, nuclear | -5.1354 | 4.0651 | -5.1374 | 4.0647 | 0.002 | 0.0007 | 0.4997 |
| HMR_7786 | Transport, nuclear | 254.3177 | 120417.0834 | 253.7756 | 120578.7253 | 0.5421 | 0.0011 | 0.4996 |
| HMR_7796 | Transport, nuclear | 15.4841 | 68.2187 | 16.6405 | 78.6624 | 1.1564 | -0.0954 | 0.538 |
| HMR_7855 | Transport, nuclear | 186.7485 | 130175.7765 | 195.8318 | 128041.889 | 9.0833 | -0.0179 | 0.5071 |
| HMR_7856 | Transport, nuclear | 200.2005 | 130162.5056 | 210.4161 | 128027.0761 | 10.2156 | -0.0201 | 0.508 |
| HMR_7857 | Transport, nuclear | -200.2013 | 130162.5051 | -210.4169 | 128027.0757 | 10.2156 | 0.0201 | 0.492 |
| HMR_7858 | Transport, nuclear | -13.4966 | 70.1411 | -14.6293 | 80.622 | 1.1327 | 0.0923 | 0.4632 |
| HMR_7859 | Transport, nuclear | -2.017 | 1.8973 | -2.0406 | 1.9424 | 0.0237 | 0.0121 | 0.4952 |
| HMR_7877 | Transport, nuclear | -186.7404 | 130175.7764 | -195.8237 | 128041.8891 | 9.0833 | 0.0179 | 0.4929 |
| HMR_8789 | Transport, nuclear | 3.6982 | 3.8684 | 3.6908 | 3.8611 | 0.0074 | 0.0027 | 0.4989 |
| HMR_0796 | Transport, lysosomal | 482.437 | 59953.8267 | 482.4366 | 59953.7632 | 0.0004 | 0.00E+00 | 0.5 |
| HMR_1917 | Transport, lysosomal | -117.7298 | 256844.4721 | -117.7845 | 256837.9873 | 0.0546 | 1.00E-04 | 0.5 |
| HMR_2087 | Transport, lysosomal | -190.8284 | 16891.4789 | -188.0889 | 17039.6933 | 2.7395 | -0.0149 | 0.5059 |
| HMR_2090 | Transport, lysosomal | 190.8284 | 16891.4789 | 188.0889 | 17039.6933 | 2.7395 | 0.0149 | 0.4941 |
| HMR_3535 | Transport, lysosomal | 503.595 | 55115.5049 | 503.6517 | 55115.2134 | 0.0567 | -0.0002 | 0.5001 |
| HMR_3601 | Transport, lysosomal | 9.8232 | 11.5746 | 9.9546 | 11.8553 | 0.1314 | -0.0271 | 0.5108 |
| HMR_3987 | Transport, lysosomal | 0.86 | 0.2042 | 0.8744 | 0.2113 | 0.0144 | -0.0224 | 0.5089 |
| HMR_4887 | Transport, lysosomal | 436.7697 | 67553.1161 | 439.4165 | 68127.2553 | 2.6468 | -0.0072 | 0.5029 |
| HMR_4907 | Transport, lysosomal | 20.4533 | 40.7772 | 29.3679 | 65.4088 | 8.9146 | -0.8651 | 0.8065 |
| HMR_4962 | Transport, lysosomal | 447.1913 | 71211.1964 | 447.2004 | 71213.0303 | 0.0091 | 0.00E+00 | 0.5 |
| HMR_5047 | Transport, lysosomal | 43.178 | 71.3476 | 51.1468 | 93.4979 | 7.9688 | -0.6207 | 0.7326 |
| HMR_5048 | Transport, lysosomal | 2.158 | 0.5526 | 3.0117 | 0.8661 | 0.8537 | -0.7167 | 0.7632 |
| HMR_5049 | Transport, lysosomal | 1.194 | 0.1848 | 1.7788 | 0.3251 | 0.5847 | -0.8188 | 0.7936 |
| HMR_5050 | Transport, lysosomal | 1.9719 | 0.4104 | 2.8212 | 0.6529 | 0.8493 | -0.8236 | 0.7949 |
| HMR_5051 | Transport, lysosomal | 0.9287 | 0.1355 | 1.3809 | 0.2154 | 0.4522 | -0.7634 | 0.7774 |
| HMR_5052 | Transport, lysosomal | 1.6503 | 0.3834 | 2.3006 | 0.599 | 0.6503 | -0.6561 | 0.7441 |
| HMR_5053 | Transport, lysosomal | 2.6893 | 0.7522 | 3.866 | 1.1867 | 1.1766 | -0.845 | 0.8009 |
| HMR_5054 | Transport, lysosomal | 1.0357 | 0.1243 | 1.5031 | 0.2088 | 0.4674 | -0.8098 | 0.791 |

| | | | | | | | | |
|---|---|---|---|---|---|---|---|---|
| HMR_5055 | Transport, lysosomal | 4.7814 | 2.3988 | 6.7242 | 3.714 | 1.9428 | -0.7858 | 0.784 |
| HMR_5056 | Transport, lysosomal | 2.7162 | 0.7784 | 3.9318 | 1.2819 | 1.2156 | -0.8469 | 0.8015 |
| HMR_5057 | Transport, lysosomal | 1.1421 | 0.1722 | 1.5799 | 0.2599 | 0.4379 | -0.6662 | 0.7474 |
| HMR_5058 | Transport, lysosomal | 1.3588 | 0.213 | 1.993 | 0.367 | 0.6342 | -0.8327 | 0.7975 |
| HMR_5059 | Transport, lysosomal | 2.2865 | 0.6137 | 3.2541 | 0.9645 | 0.9676 | -0.7703 | 0.7794 |
| HMR_5060 | Transport, lysosomal | 3.2615 | 1.0704 | 4.7626 | 1.7677 | 1.5012 | -0.8911 | 0.8136 |
| HMR_5061 | Transport, lysosomal | 0.5566 | 0.0514 | 0.8122 | 0.0867 | 0.2556 | -0.6878 | 0.7542 |
| HMR_5062 | Transport, lysosomal | 1.1491 | 0.1608 | 1.6508 | 0.2638 | 0.5017 | -0.7699 | 0.7793 |
| HMR_5063 | Transport, lysosomal | 2.6418 | 0.7049 | 3.842 | 1.1595 | 1.2002 | -0.879 | 0.8103 |
| HMR_5064 | Transport, lysosomal | 3.1817 | 1.0966 | 4.4601 | 1.6876 | 1.2784 | -0.7662 | 0.7782 |
| HMR_5065 | Transport, lysosomal | 1.5578 | 0.3613 | 2.3888 | 0.6034 | 0.831 | -0.8461 | 0.8013 |
| HMR_5066 | Transport, lysosomal | 1.7882 | 0.3339 | 2.6173 | 0.56 | 0.8291 | -0.8769 | 0.8097 |
| HMR_5067 | Transport, lysosomal | 2.984 | 1.0111 | 4.2397 | 1.5971 | 1.2557 | -0.7776 | 0.7816 |
| HMR_5204 | Transport, lysosomal | 0.08 | 0.003 | 0.0818 | 0.0031 | 0.0018 | -0.023 | 0.5092 |
| HMR_5224 | Transport, lysosomal | 0.0781 | 0.0033 | 0.082 | 0.0036 | 0.0039 | -0.0466 | 0.5186 |
| HMR_5226 | Transport, lysosomal | 0.0639 | 0.0022 | 0.0679 | 0.0024 | 0.004 | -0.0589 | 0.5235 |
| HMR_5228 | Transport, lysosomal | 0.0606 | 0.002 | 0.0613 | 0.002 | 0.0007 | -0.0107 | 0.5043 |
| HMR_5229 | Transport, lysosomal | 0.0595 | 0.002 | 0.0608 | 0.002 | 0.0013 | -0.0209 | 0.5083 |
| HMR_5230 | Transport, lysosomal | 0.1013 | 0.0042 | 0.1074 | 0.0045 | 0.0061 | -0.0652 | 0.526 |
| HMR_7199 | Transport, lysosomal | 0 | 220512.1421 | 0 | 220512.1403 | 0 | 0 | 0.5 |
| HMR_7711 | Transport, lysosomal | 4.2985 | 6.5654 | 4.2469 | 6.4122 | 0.0517 | 0.0143 | 0.4943 |
| HMR_7714 | Transport, lysosomal | -4.298 | 6.5654 | -4.2465 | 6.4121 | 0.0516 | -0.0143 | 0.5057 |
| HMR_7715 | Transport, lysosomal | 3.3721 | 4.0498 | 3.1751 | 3.5934 | 0.197 | 0.0712 | 0.4716 |
| HMR_7718 | Transport, lysosomal | -3.3818 | 4.0499 | -3.1848 | 3.5936 | 0.197 | -0.0713 | 0.5284 |
| HMR_7720 | Transport, lysosomal | 4.9678 | 8.3747 | 4.7685 | 7.7067 | 0.1993 | 0.0497 | 0.4802 |
| HMR_7722 | Transport, lysosomal | -4.9847 | 8.3733 | -4.7857 | 7.7054 | 0.199 | -0.0496 | 0.5198 |
| HMR_7724 | Transport, lysosomal | 3.9715 | 5.4618 | 3.7518 | 4.8874 | 0.2197 | 0.0683 | 0.4728 |
| HMR_7726 | Transport, lysosomal | -3.9709 | 5.4615 | -3.7511 | 4.8871 | 0.2198 | -0.0683 | 0.5272 |
| HMR_7727 | Transport, lysosomal | 3.2788 | 3.7663 | 3.1271 | 3.4292 | 0.1517 | 0.0565 | 0.4775 |
| HMR_7729 | Transport, lysosomal | -3.2823 | 3.7663 | -3.1303 | 3.4292 | 0.1519 | -0.0566 | 0.5226 |
| HMR_7730 | Transport, lysosomal | 2.8326 | 2.7156 | 2.7079 | 2.4865 | 0.1247 | 0.0547 | 0.4782 |
| HMR_7732 | Transport, lysosomal | -2.8237 | 2.714 | -2.6981 | 2.4849 | 0.1256 | -0.0551 | 0.522 |
| HMR_7762 | Transport, lysosomal | -190.8299 | 16891.4821 | -188.0904 | 17039.6965 | 2.7395 | -0.0149 | 0.5059 |
| HMR_3597 | Transport, lysosome to ER | 493.7285 | 55117.714 | 493.6542 | 55117.4482 | 0.0743 | 0.0002 | 0.4999 |
| HMR_4961 | Transport, lysosome to ER | -445.4901 | 71211.5355 | -445.4699 | 71213.3667 | 0.0202 | -1.00E-04 | 0.5 |
| HMR_5045 | Transport, lysosome to ER | 85.544 | 219824.4168 | 85.5933 | 219820.2522 | 0.0493 | -1.00E-04 | 0.5 |
| HMR_5249 | Transport, lysosome to ER | 482.4316 | 59953.8277 | 482.4311 | 59953.7641 | 0.0005 | 0.00E+00 | 0.5 |
| HMR_4892 | Transport, Golgi apparatus | 2.1066 | 1.3103 | 2.1327 | 1.3432 | 0.0261 | -0.016 | 0.5064 |
| HMR_7331 | Transport, Golgi apparatus | -2.1087 | 1.3103 | -2.1347 | 1.3432 | 0.026 | 0.016 | 0.4936 |
| HMR_7431 | Transport, Golgi apparatus | 0 | 220512.1421 | 0 | 220512.1403 | 0 | 0 | 0.5 |
| HMR_8084 | Transport, Golgi apparatus | 2.1008 | 1.3092 | 2.1272 | 1.3422 | 0.0264 | -0.0162 | 0.5065 |
| HMR_7432 | Transport, Golgi to lysosome | 0 | 220512.1421 | 0 | 220512.1403 | 0 | 0 | 0.5 |
| HMR_0020 | Transport, endoplasmic reticular | -502.6807 | 55115.266 | -502.6984 | 55114.9637 | 0.0177 | 1.00E-04 | 0.5 |
| HMR_0730 | Transport, endoplasmic reticular | 0.5301 | 4.7565 | 0.533 | 4.8414 | 0.0028 | -0.0009 | 0.5004 |

| | | | | | | | | |
|---|---|---|---|---|---|---|---|---|
| HMR_0771 | Transport, endoplasmic reticular | -482.9474 | 59957.1585 | -482.9495 | 59957.1521 | 0.0021 | 0.00E+00 | 0.5 |
| HMR_1311 | Transport, endoplasmic reticular | -393.5254 | 114437.1379 | -392.1596 | 114879.3638 | 1.3658 | -0.0029 | 0.5011 |
| HMR_1331 | Transport, endoplasmic reticular | 393.5256 | 114437.1372 | 392.1598 | 114879.3631 | 1.3658 | 0.0029 | 0.4989 |
| HMR_1530 | Transport, endoplasmic reticular | -73.864 | 219822.7952 | -73.891 | 219818.6623 | 0.027 | 0.00E+00 | 0.5 |
| HMR_2039 | Transport, endoplasmic reticular | 4.1315 | 3.5007 | 4.118 | 3.48 | 0.0135 | 0.0051 | 0.498 |
| HMR_2041 | Transport, endoplasmic reticular | 2.063 | 2.7753 | 2.0561 | 2.7576 | 0.0068 | 0.0029 | 0.4988 |
| HMR_2569 | Transport, endoplasmic reticular | -60.4341 | 154237.8608 | -60.9056 | 154192.7978 | 0.4716 | 0.0008 | 0.4997 |
| HMR_3953 | Transport, endoplasmic reticular | -623.8482 | 30768.3839 | -622.0163 | 30904.957 | 1.8318 | -0.0074 | 0.5029 |
| HMR_4123 | Transport, endoplasmic reticular | 2.1484 | 199905.915 | 2.1466 | 199906.0017 | 0.0018 | 0.00E+00 | 0.5 |
| HMR_4765 | Transport, endoplasmic reticular | -97.5825 | 181988.5521 | -97.679 | 181988.5578 | 0.0965 | 0.0002 | 0.4999 |
| HMR_4846 | Transport, endoplasmic reticular | -49.3272 | 154266.489 | -50.174 | 154222.5669 | 0.8468 | 0.0015 | 0.4994 |
| HMR_4856 | Transport, endoplasmic reticular | -480.0599 | 72467.5725 | -480.6003 | 72428.0157 | 0.5403 | 0.0014 | 0.4994 |
| HMR_4876 | Transport, endoplasmic reticular | 231.4183 | 152082.7499 | 230.4347 | 152186.6553 | 0.9836 | 0.0018 | 0.4993 |
| HMR_4877 | Transport, endoplasmic reticular | 381.8009 | 157287.4288 | 381.4798 | 157425.5649 | 0.3211 | 0.0006 | 0.4998 |
| HMR_4878 | Transport, endoplasmic reticular | -2.1757 | 199905.915 | -2.1737 | 199906.0018 | 0.0021 | 0.00E+00 | 0.5 |
| HMR_4879 | Transport, endoplasmic reticular | 2.1657 | 199905.9102 | 2.1637 | 199905.9969 | 0.002 | 0.00E+00 | 0.5 |
| HMR_4880 | Transport, endoplasmic reticular | 6.4884 | 199907.8138 | 6.4826 | 199907.8929 | 0.0058 | 0.00E+00 | 0.5 |
| HMR_4883 | Transport, endoplasmic reticular | 108.8698 | 470.9811 | 106.7154 | 469.9374 | 2.1543 | 0.0702 | 0.472 |
| HMR_4894 | Transport, endoplasmic reticular | 17.1773 | 55.0061 | 18.0706 | 60.7974 | 0.8933 | -0.083 | 0.5331 |
| HMR_4906 | Transport, endoplasmic reticular | 5.7817 | 8.9 | 5.5194 | 8.1129 | 0.2623 | 0.0636 | 0.4747 |
| HMR_4917 | Transport, endoplasmic reticular | 613.2209 | 30586.9414 | 611.9166 | 30745.4399 | 1.3043 | 0.0053 | 0.4979 |
| HMR_4941 | Transport, endoplasmic reticular | -0.5292 | 4.7606 | -0.5321 | 4.8456 | 0.0029 | 0.0009 | 0.4996 |
| HMR_4960 | Transport, endoplasmic reticular | 445.4948 | 71211.5401 | 445.4744 | 71213.3714 | 0.0203 | 1.00E-04 | 0.5 |
| HMR_5004 | Transport, endoplasmic reticular | 97.5825 | 181988.5492 | 97.679 | 181988.5549 | 0.0965 | -0.0002 | 0.5001 |
| HMR_5027 | Transport, endoplasmic reticular | -3.0455 | 4.3328 | -2.9006 | 3.9303 | 0.1449 | -0.0504 | 0.5201 |
| HMR_5342 | Transport, endoplasmic reticular | 639.5348 | 30695.622 | 640.5398 | 30841.0208 | 1.0049 | -0.0041 | 0.5016 |
| HMR_5347 | Transport, endoplasmic reticular | -119.8281 | 241499.3159 | -120.5556 | 241433.1807 | 0.7276 | 0.001 | 0.4996 |
| HMR_5354 | Transport, endoplasmic reticular | 0.5091 | 4.7586 | 0.5118 | 4.8435 | 0.0027 | -0.0009 | 0.5003 |
| HMR_6386 | Transport, endoplasmic reticular | 493.7311 | 72360.059 | 493.5989 | 72334.8777 | 0.1322 | 0.0003 | 0.4999 |
| HMR_6387 | Transport, endoplasmic reticular | -97.5821 | 181988.5481 | -97.6786 | 181988.5538 | 0.0965 | 0.0002 | 0.4999 |
| HMR_6388 | Transport, endoplasmic reticular | 75.3138 | 194036.1849 | 74.4492 | 194055.4558 | 0.8647 | 0.0014 | 0.4994 |
| HMR_7649 | Transport, endoplasmic reticular | -7.7485 | 3.6262 | -7.7527 | 3.6117 | 0.0042 | 0.0016 | 0.4994 |
| HMR_7772 | Transport, endoplasmic reticular | 365.0527 | 38037.9111 | 365.3425 | 38135.1439 | 0.2898 | -0.0011 | 0.5004 |
| HMR_7776 | Transport, endoplasmic reticular | 365.0428 | 38037.9113 | 365.3327 | 38135.1441 | 0.2899 | -0.0011 | 0.5004 |
| HMR_7778 | Transport, endoplasmic reticular | 276.0319 | 28920.4969 | 276.7378 | 29048.8388 | 0.7059 | -0.0029 | 0.5012 |
| HMR_7780 | Transport, endoplasmic reticular | 276.0243 | 28920.497 | 276.7304 | 29048.8389 | 0.706 | -0.0029 | 0.5012 |
| HMR_7969 | Transport, endoplasmic reticular | 3.6406 | 2.9169 | 3.659 | 2.9244 | 0.0184 | -0.0076 | 0.503 |
| HMR_7983 | Transport, endoplasmic reticular | 0.9001 | 0.188 | 0.888 | 0.1831 | 0.0121 | 0.0198 | 0.4921 |
| HMR_7985 | Transport, endoplasmic reticular | -0.8967 | 0.1878 | -0.8845 | 0.1829 | 0.0122 | -0.0201 | 0.508 |
| HMR_9023 | Exchange reactions | 4.9708 | 9.7269 | 4.8253 | 9.1815 | 0.1455 | 0.0335 | 0.4867 |
| HMR_9024 | Exchange reactions | 0.0016 | 0.00E+00 | 0.0016 | 0.00E+00 | 0.00E+00 | 0.00E+00 | 0.5 |
| HMR_9025 | Exchange reactions | 0.012 | 0.00E+00 | 0.012 | 0.00E+00 | 0.00E+00 | 0.00E+00 | 0.5 |
| HMR_9026 | Exchange reactions | 0.0815 | 0.0037 | 0.0816 | 0.0037 | 0.00E+00 | -0.0002 | 0.5001 |
| HMR_9027 | Exchange reactions | 0.0479 | 0.0009 | 0.0479 | 0.0009 | 0.00E+00 | 0.00E+00 | 0.5 |

| | | | | | | | | |
|---|---|---|---|---|---|---|---|---|
| HMR_9033 | Exchange reactions | 87.989 | 1244.8789 | 97.4843 | 1451.3575 | 9.4953 | -0.1829 | 0.5725 |
| HMR_9034 | Exchange reactions | 576.5929 | 60465.3626 | 577.5123 | 60370.8353 | 0.9194 | -0.0026 | 0.5011 |
| HMR_9035 | Exchange reactions | 20.9393 | 12.9589 | 22.3765 | 14.7978 | 1.4373 | -0.2728 | 0.6075 |
| HMR_9036 | Exchange reactions | 11.2741 | 3.2197 | 12.41 | 3.8535 | 1.1359 | -0.4271 | 0.6653 |
| HMR_9037 | Exchange reactions | 0.84 | 0.1318 | 1.0741 | 0.1972 | 0.234 | -0.408 | 0.6584 |
| HMR_9038 | Exchange reactions | 677.9054 | 2984.6591 | 827.3137 | 2968.6134 | 149.4083 | -1.9364 | 0.9736 |
| HMR_9039 | Exchange reactions | 504.0355 | 63768.3726 | 0.00E+00 | 0 | 504.0355 | 1.996 | 0.023 |
| HMR_9040 | Exchange reactions | 498.6294 | 63776.7019 | 0 | 0 | 498.6294 | 1.9745 | 0.0242 |
| HMR_9041 | Exchange reactions | 12.3094 | 36.249 | 9.9612 | 32.7443 | 2.3482 | 0.2827 | 0.3887 |
| HMR_9042 | Exchange reactions | 0.7396 | 0.3501 | 1.2799 | 1.4916 | 0.5403 | -0.3981 | 0.6547 |
| HMR_9043 | Exchange reactions | 505.4483 | 63766.5441 | 505.1028 | 63769.4521 | 0.3455 | 0.001 | 0.4996 |
| HMR_9044 | Exchange reactions | 508.9006 | 63748.552 | 507.7905 | 63758.6084 | 1.1102 | 0.0031 | 0.4988 |
| HMR_9045 | Exchange reactions | 4.5594 | 3.7317 | 4.1132 | 3.5674 | 0.4462 | 0.1651 | 0.4344 |
| HMR_9046 | Exchange reactions | 662.7702 | 48045.6149 | 0.00E+00 | 0 | 662.7702 | 3.0237 | 0.0012 |
| HMR_9047 | Exchange reactions | 961.7954 | 1385.699 | 967.0895 | 1513.2784 | 5.2941 | -0.0983 | 0.5392 |
| HMR_9048 | Exchange reactions | 994.4741 | 25.9026 | 994.3251 | 27.6264 | 0.1491 | 0.0204 | 0.4919 |
| HMR_9052 | Exchange reactions | 0.0016 | 0.00E+00 | 0.0016 | 0.00E+00 | 0.00E+00 | 0.00E+00 | 0.5 |
| HMR_9053 | Exchange reactions | 0.012 | 0.00E+00 | 0.012 | 0.00E+00 | 0.00E+00 | 0.00E+00 | 0.5 |
| HMR_9054 | Exchange reactions | 0.0815 | 0.0037 | 0.0816 | 0.0037 | 0.00E+00 | -0.0002 | 0.5001 |
| HMR_9055 | Exchange reactions | 0.0479 | 0.0009 | 0.0479 | 0.0009 | 0.00E+00 | 0.00E+00 | 0.5 |
| HMR_9056 | Exchange reactions | 88.8631 | 1436.8267 | 95.2818 | 1658.3934 | 6.4187 | -0.1154 | 0.5459 |
| HMR_9057 | Exchange reactions | 38.2046 | 1385.6982 | 32.9105 | 1513.2784 | 5.2941 | 0.0983 | 0.4608 |
| HMR_9058 | Exchange reactions | 978.9824 | 143.6766 | 980.9541 | 117.7508 | 1.9717 | -0.1219 | 0.5485 |
| HMR_9059 | Exchange reactions | 494.5517 | 63766.5441 | 494.8972 | 63769.4521 | 0.3455 | -0.001 | 0.5004 |
| HMR_9060 | Exchange reactions | 491.0994 | 63748.552 | 492.2095 | 63758.6084 | 1.1102 | -0.0031 | 0.5012 |
| HMR_9061 | Exchange reactions | 682.2768 | 53903.9081 | 637.5296 | 63703.9654 | 44.7472 | 0.1305 | 0.4481 |
| HMR_9062 | Exchange reactions | 1.672 | 1.6621 | 2.0654 | 2.2787 | 0.3934 | -0.1982 | 0.5786 |
| HMR_9063 | Exchange reactions | 3.0407 | 9.7786 | 2.914 | 9.2079 | 0.1267 | 0.0291 | 0.4884 |
| HMR_9064 | Exchange reactions | 490.3975 | 63744.859 | 490.8119 | 63751.2174 | 0.4144 | -0.0012 | 0.5005 |
| HMR_9065 | Exchange reactions | 738.2995 | 34630.7581 | 718.6153 | 33379.6144 | 19.6842 | 0.0755 | 0.4699 |
| HMR_9066 | Exchange reactions | 926.8753 | 986.5268 | 916.8396 | 1487.5362 | 10.0357 | 0.2018 | 0.4201 |
| HMR_9067 | Exchange reactions | 961.3804 | 1356.7079 | 971.9034 | 1058.102 | 10.523 | -0.2141 | 0.5848 |
| HMR_9068 | Exchange reactions | 957.1225 | 1286.7978 | 905.0507 | 5209.8204 | 52.0719 | 0.646 | 0.2591 |
| HMR_9069 | Exchange reactions | 41.22 | 1553.4974 | 34.849 | 1130.3351 | 6.371 | 0.123 | 0.4511 |
| HMR_9070 | Exchange reactions | -7.6204 | 20.6087 | -7.359 | 19.2475 | 0.2614 | -0.0414 | 0.5165 |
| HMR_9071 | Exchange reactions | 72.6339 | 2512.0576 | 58.803 | 2084.1028 | 13.8309 | 0.204 | 0.4192 |
| HMR_9072 | Exchange reactions | -66.321 | 147.9018 | -63.6056 | 140.0008 | 2.7153 | -0.16 | 0.5636 |
| HMR_9073 | Exchange reactions | 344.7089 | 13368.3228 | -38.359 | 13560.6515 | 383.0679 | 2.3343 | 0.0098 |
| HMR_9074 | Exchange reactions | -18.3915 | 89.2078 | -17.7945 | 80.4128 | 0.597 | -0.0458 | 0.5183 |
| HMR_9075 | Exchange reactions | -31.3208 | 35.0447 | -29.9621 | 33.0553 | 1.3587 | -0.1646 | 0.5654 |
| HMR_9076 | Exchange reactions | 33.5501 | 359.4624 | 36.7288 | 433.8086 | 3.1787 | -0.1129 | 0.5449 |
| HMR_9078 | Exchange reactions | -993.7448 | 24.6245 | -994.1672 | 21.4008 | 0.4223 | 0.0623 | 0.4752 |
| HMR_9079 | Exchange reactions | 92.2644 | 1239.1014 | 100.4476 | 1326.0613 | 8.1832 | -0.1616 | 0.5642 |
| HMR_9083 | Exchange reactions | -3.659 | 11.4113 | -3.0244 | 10.203 | 0.6346 | -0.1365 | 0.5543 |

| | | | | | | | | |
|---|---|---|---|---|---|---|---|---|
| HMR_9084 | Exchange reactions | 9.0613 | 222.4959 | 10.8261 | 219.3031 | 1.7648 | -0.084 | 0.5335 |
| HMR_9085 | Exchange reactions | 53.0912 | 2039.1193 | 69.829 | 3088.7437 | 16.7378 | -0.2337 | 0.5924 |
| HMR_9086 | Exchange reactions | -151.7474 | 1415.799 | -148.7011 | 1321.3382 | 3.0463 | -0.0582 | 0.5232 |
| HMR_9087 | Exchange reactions | 981.4245 | 297.9846 | 979.3077 | 383.4358 | 2.1168 | 0.0811 | 0.4677 |
| HMR_9088 | Exchange reactions | -27.8085 | 396.4098 | -36.8448 | 514.705 | 9.0363 | 0.2994 | 0.3823 |
| HMR_9090 | Exchange reactions | -4.3182 | 4.6179 | -4.0781 | 4.121 | 0.2402 | -0.0812 | 0.5324 |
| HMR_9091 | Exchange reactions | 8.0782 | 37.4908 | 8.8747 | 41.1594 | 0.7965 | -0.0898 | 0.5358 |
| HMR_9092 | Exchange reactions | -29.9039 | 175.5506 | -30.4288 | 182.8185 | 0.5249 | 0.0277 | 0.4889 |
| HMR_9093 | Exchange reactions | 3.1957 | 3.0942 | 3.0248 | 2.7736 | 0.1709 | 0.0706 | 0.4719 |
| HMR_9094 | Exchange reactions | -2.7051 | 2.5182 | -2.6705 | 2.4159 | 0.0346 | -0.0156 | 0.5062 |
| HMR_9095 | Exchange reactions | -3.1942 | 3.0892 | -3.0232 | 2.7686 | 0.171 | -0.0707 | 0.5282 |
| HMR_9096 | Exchange reactions | -32.1131 | 359.3788 | -35.4181 | 433.7557 | 3.305 | 0.1174 | 0.4533 |
| HMR_9098 | Exchange reactions | -514.2668 | 66969.5874 | -401.3674 | 89060.9394 | 112.8994 | -0.2858 | 0.6125 |
| HMR_9099 | Exchange reactions | 25.2898 | 157.0444 | 27.638 | 186.2172 | 2.3482 | -0.1267 | 0.5504 |
| HMR_9102 | Exchange reactions | 37.4984 | 305.4702 | 34.2476 | 257.9899 | 3.2508 | 0.1369 | 0.4455 |
| HMR_9103 | Exchange reactions | 293.4265 | 4691.3083 | 366.2917 | 3928.5656 | 72.8652 | -0.7848 | 0.7837 |
| HMR_9104 | Exchange reactions | -960.2962 | 355.7349 | -959.0331 | 375.6005 | 1.2631 | -0.0467 | 0.5186 |
| HMR_9107 | Exchange reactions | -1.4392 | 0.3194 | -1.3131 | 0.2694 | 0.1261 | -0.1643 | 0.5653 |
| HMR_9108 | Exchange reactions | -1.6463 | 0.5487 | -1.5974 | 0.518 | 0.0489 | -0.0474 | 0.5189 |
| HMR_9122 | Exchange reactions | -1.704 | 1.1361 | -1.7442 | 1.1908 | 0.0402 | 0.0264 | 0.4895 |
| HMR_9123 | Exchange reactions | 0.1943 | 3.4544 | 0.1987 | 3.6196 | 0.0044 | -0.0017 | 0.5007 |
| HMR_9129 | Exchange reactions | -7.745 | 3.6462 | -7.749 | 3.6317 | 0.0041 | 0.0015 | 0.4994 |
| HMR_9130 | Exchange reactions | -0.17 | 0.0115 | -0.1733 | 0.0119 | 0.0034 | 0.0222 | 0.4912 |
| HMR_9131 | Exchange reactions | 93.6207 | 1474.2973 | 102.4714 | 1885.1074 | 8.8507 | -0.1527 | 0.5607 |
| HMR_9132 | Exchange reactions | -148.9745 | 15798.4998 | -261.8774 | 38397.3651 | 112.9029 | 0.485 | 0.3138 |
| HMR_9133 | Exchange reactions | 817.1257 | 20602.2992 | 566.9194 | 68672.1263 | 250.2063 | 0.8374 | 0.2012 |
| HMR_9134 | Exchange reactions | 118.8781 | 15525.5815 | 232.374 | 38166.4115 | 113.4959 | -0.4898 | 0.6879 |
| HMR_9135 | Exchange reactions | -632.8099 | 21866.2032 | -390.1747 | 69705.6106 | 242.6352 | -0.8018 | 0.7887 |
| HMR_9136 | Exchange reactions | 5.8996 | 16.2208 | 5.868 | 16.0474 | 0.0315 | 0.0056 | 0.4978 |
| HMR_9138 | Exchange reactions | 42.0843 | 614.1252 | 47.858 | 789.8953 | 5.7737 | -0.1541 | 0.5612 |
| HMR_9139 | Exchange reactions | 121.9996 | 2544.3105 | 121.0896 | 2528.2865 | 0.91 | 0.0128 | 0.4949 |
| HMR_9141 | Exchange reactions | 9.7136 | 96.5752 | 9.1144 | 88.8723 | 0.5992 | 0.044 | 0.4825 |
| HMR_9142 | Exchange reactions | 958.6224 | 356.2258 | 957.3413 | 376.106 | 1.2812 | 0.0473 | 0.4811 |
| HMR_9145 | Exchange reactions | 0.0026 | 0.00E+00 | 0.0026 | 0.00E+00 | 0.00E+00 | 0.00E+00 | 0.5 |
| HMR_9155 | Exchange reactions | 1.757 | 1.0753 | 1.763 | 1.0812 | 0.0061 | -0.0041 | 0.5017 |
| HMR_9156 | Exchange reactions | 1.3415 | 0.5442 | 1.3462 | 0.548 | 0.0047 | -0.0045 | 0.5018 |
| HMR_9165 | Exchange reactions | -39.3737 | 191.9313 | -39.94 | 204.0491 | 0.5663 | 0.0285 | 0.4886 |
| HMR_9166 | Exchange reactions | 1.7304 | 0.8295 | 1.7591 | 0.8581 | 0.0287 | -0.0221 | 0.5088 |
| HMR_9168 | Exchange reactions | 61.5899 | 747.1161 | 56.8507 | 641.2987 | 4.7391 | 0.1272 | 0.4494 |
| HMR_9171 | Exchange reactions | -355.812 | 12420.3253 | -555.2822 | 12637.3317 | 199.4702 | 1.2601 | 0.1038 |
| HMR_9172 | Exchange reactions | 354.9605 | 12420.2403 | 554.4444 | 12637.4658 | 199.4839 | -1.2602 | 0.8962 |
| HMR_9223 | Exchange reactions | -6.7681 | 12.3322 | -7.006 | 13.2122 | 0.2379 | 0.0471 | 0.4812 |
| HMR_9241 | Exchange reactions | -726.8843 | 18062.4345 | -489.2168 | 42958.3668 | 237.6675 | -0.9621 | 0.832 |
| HMR_9243 | Exchange reactions | 9.5664 | 27.5756 | 10.1704 | 31.1627 | 0.604 | -0.0788 | 0.5314 |

| | | | | | | | | |
|---|---|---|---|---|---|---|---|---|
| HMR_9260 | Exchange reactions | 12.2476 | 367.1221 | 6.9733 | 261.9294 | 5.2743 | 0.2103 | 0.4167 |
| HMR_9270 | Exchange reactions | 726.8843 | 18062.4345 | 489.2168 | 42958.3668 | 237.6675 | 0.9621 | 0.168 |
| HMR_9286 | Exchange reactions | 72.1383 | 220.479 | 69.9322 | 207.0845 | 2.2061 | 0.1067 | 0.4575 |
| HMR_9299 | Exchange reactions | 6.7681 | 12.3322 | 7.006 | 13.2122 | 0.2379 | -0.0471 | 0.5188 |
| HMR_9302 | Exchange reactions | 4.5632 | 2.201 | 4.6685 | 2.2943 | 0.1054 | -0.0497 | 0.5198 |
| HMR_9315 | Exchange reactions | -0.8986 | 0.1985 | -0.8863 | 0.1935 | 0.0123 | -0.0196 | 0.5078 |
| HMR_9317 | Exchange reactions | 3.2983 | 2.2338 | 3.1981 | 2.1247 | 0.1002 | 0.048 | 0.4809 |
| HMR_9361 | Exchange reactions | 2.4473 | 1.5781 | 2.4783 | 1.6123 | 0.031 | -0.0174 | 0.5069 |
| HMR_9363 | Exchange reactions | 240.387 | 8858.2734 | 196.8231 | 7888.6419 | 43.5638 | 0.3366 | 0.3682 |
| HMR_9415 | Exchange reactions | 4.7274 | 7.3288 | 4.745 | 7.3891 | 0.0177 | -0.0046 | 0.5018 |
| HMR_9450 | Exchange reactions | 1.6851 | 0.995 | 1.6947 | 1.0052 | 0.0096 | -0.0068 | 0.5027 |
| HMR_9451 | Exchange reactions | 6.6656 | 8.9091 | 6.6828 | 8.9599 | 0.0173 | -0.0041 | 0.5016 |
| HMR_9452 | Exchange reactions | 0.9053 | 0.199 | 0.8933 | 0.194 | 0.012 | 0.0191 | 0.4924 |
| HMR_9456 | Exchange reactions | 17.9533 | 90.1629 | 17.1976 | 82.8176 | 0.7556 | 0.0575 | 0.4771 |
| HMR_obj | Objective function | 8.3739 | 7.8136 | 8.4406 | 7.9528 | 0.0667 | -0.0168 | 0.5067 |
| EXC_OUT_m4082 | | 501.3706 | 63776.7018 | 6.5167 | 9.2994 | 494.8538 | 1.9594 | 0.025 |
| EXC_OUT_m3848 | | 495.9645 | 63768.3726 | 1.0512 | 0.6424 | 494.9133 | 1.9599 | 0.025 |
| EXC_OUT_m5559 | | 337.2298 | 48045.6149 | 0.922 | 0.6866 | 336.3078 | 1.5343 | 0.0625 |
| EXC_OUT_m3413 | | 423.4071 | 60465.3629 | 422.4877 | 60370.8354 | 0.9194 | 0.0026 | 0.4989 |
| EXC_IN_m5479 | | 509.6025 | 63744.859 | 509.1881 | 63751.2173 | 0.4144 | 0.0012 | 0.4995 |
| EXC_IN_m3462 | | 38.6196 | 1356.7079 | 28.0966 | 1058.102 | 10.523 | 0.2141 | 0.4152 |
| EXC_IN_m2118 | | 317.7232 | 53903.9085 | 362.4704 | 63703.9664 | 44.7472 | -0.1305 | 0.5519 |
| EXC_IN_m4849 | | 42.8775 | 1286.7978 | 94.9493 | 5209.8204 | 52.0719 | -0.646 | 0.7409 |
| EXC_BOTH_m2731 | | -921.6847 | 986.1509 | -910.1434 | 1486.7866 | 11.5413 | -0.2321 | 0.5918 |
| EXC_BOTH_m1103 | | -1.2275 | 246.5368 | -9.7008 | 284.6876 | 8.4733 | 0.3676 | 0.3566 |

| TableS2: Reactions with significant changes in flux statistics between unconstrained and constrained (bcaa=0) simulations | | | | | | |
|---|---|---|---|---|---|---|
| **Reaction** | **Chemical Equation** | **Subsystem** | **Mean (unconstrained)** | **Mean (constrained)** | **Std dev (unconstrained)** | **Std dev (constrained)** |
| EXC_OUT_m3848 | isoleucine[s] => | Exchange reactions | 495.96 | 1.05 | 252.52 | 0.8 |
| EXC_OUT_m4082 | leucine[s] => | Exchange reactions | 501.37 | 6.52 | 252.54 | 3.05 |
| HMR_3206 | 2-methylacetoacetyl-CoA[m] + CoA[m] => acetyl-CoA[m] + propanoyl-CoA[m] | Valine, leucine, and isoleucine metabolism | 10.05 | 2.12 | 4.32 | 0.83 |
| HMR_3215 | (R)-methylmalonyl-CoA[m] => succinyl-CoA[m] | Valine, leucine, and isoleucine metabolism | 337.83 | 19.37 | 53.29 | 9.49 |
| HMR_3746 | 3-methyl-2-oxobutyrate[c] <=> 3-methyl-2-oxobutyrate[m] | Transport, mitochondrial | 317.65 | 3.12 | 53.6 | 1.7 |
| HMR_3747 | AKG[c] + valine[c] => 3-methyl-2-oxobutyrate[c] + glutamate[c] | Valine, leucine, and isoleucine metabolism | 317.65 | 3.11 | 53.6 | 1.7 |
| HMR_3748 | 3-methyl-2-oxobutyrate[m] + CoA[m] + NAD+[m] => CO2[m] + H+[m] + isobutyryl-CoA[m] + NADH[m] | Valine, leucine and isoleucine degradation | 173.2 | 2.69 | 99.66 | 1.86 |
| HMR_3753 | H2O[m] + methacrylyl-CoA[m] => 3-hydroxyisobutyryl-CoA[m] | Valine, leucine and isoleucine degradation | 330.92 | 7.01 | 53.18 | 1.31 |
| HMR_3755 | 3-hydroxyisobutyryl-CoA[m] + H2O[m] => 3-hydroxyisobutyrate[m] + CoA[m] | Valine, leucine and isoleucine degradation | 330.92 | 7.01 | 53.18 | 1.32 |
| HMR_3757 | 3-hydroxyisobutyrate[m] + NAD+[m] => 2-methyl-3-oxopropanoate[m] + H+[m] + NADH[m] | Valine, leucine, and isoleucine metabolism | 330.92 | 7.01 | 53.18 | 1.32 |
| HMR_3763 | CoA[m] + methylmalonate[m] => (R)-methylmalonyl-CoA[m] + H2O[m] | Valine, leucine and isoleucine degradation | 186.72 | 4.36 | 94.66 | 1.55 |
| HMR_3789 | methylmalonate[c] <=> methylmalonate[m] | Transport, mitochondrial | 37.01 | 2.89 | 19.53 | 1.36 |
| HMR_3823 | (S)-3-hydroxy-2-methylbutyryl-CoA[m] + NAD+[m] <=> 2-methylacetoacetyl-CoA[m] + H+[m] + NADH[m] | Valine, leucine and isoleucine degradation | 10.06 | 2.13 | 4.32 | 0.83 |
| HMR_3929 | 5,10-methenyl-THF[c] + NH3[c] <=> 5-formiminotetrahydrofolate[c] + H+[c] | Folate metabolism | -674.88 | -825.17 | 54.59 | 54.45 |
| HMR_4437 | NH3[c] + urocanate[c] <=> histidine[c] | Histidine metabolism | -674.88 | -825.17 | 54.59 | 54.45 |
| HMR_4658 | 5-formiminotetrahydrofolate[c] + glutamate[c] <=> N-formimino-L-glutamate[c] + THF[c] | Histidine metabolism | -674.88 | -825.17 | 54.59 | 54.45 |
| HMR_5107 | H+[c] + isoleucine[c] => H+[m] + isoleucine[m] | Transport, mitochondrial | 10.07 | 2.12 | 4.32 | 0.81 |
| HMR_9039 | => isoleucine[s] | Exchange reactions | 504.04 | 0 | 252.52 | 0 |
| HMR_9040 | => leucine[s] | Exchange reactions | 498.63 | 0 | 252.54 | 0 |
| HMR_9046 | => valine[s] | Exchange reactions | 662.77 | 0 | 219.19 | 0 |
| HMR_9073 | <=> NH3[s] | Exchange reactions | 344.71 | -38.36 | 115.62 | 116.45 |

| Table S3: Subsystem distribution: Reactions from global modules (unconstrained) | | |
|---|---|---|
| Subsystem | Num Reactions | % Reactions |
| Transport, extracellular | 331 | 44.97 |
| Transport, mitochondrial | 62 | 8.42 |
| Exchange reactions | 33 | 4.48 |
| Tricarboxylic acid cycle and glyoxylate/dicarboxylate metabolism | 23 | 3.13 |
| Transport, endoplasmic reticular | 17 | 2.31 |
| Glycolysis / Gluconeogenesis | 15 | 2.04 |
| Glycine, serine and threonine metabolism | 14 | 1.90 |
| Retinol metabolism | 14 | 1.90 |
| Prostaglandin biosynthesis | 13 | 1.77 |
| Arginine and proline metabolism | 11 | 1.49 |
| Pyruvate metabolism | 10 | 1.36 |
| Leukotriene metabolism | 10 | 1.36 |
| Folate metabolism | 10 | 1.36 |
| Biopterin metabolism | 10 | 1.36 |
| Transport, peroxisomal | 10 | 1.36 |
| Pentose phosphate pathway | 9 | 1.22 |
| Valine, leucine and isoleucine degradation | 9 | 1.22 |
| Alanine, aspartate and glutamate metabolism | 8 | 1.09 |
| Purine metabolism | 7 | 0.95 |
| Amino sugar and nucleotide sugar metabolism | 7 | 0.95 |
| Transport, lysosomal | 7 | 0.95 |
| Tyrosine metabolism | 6 | 0.82 |
| Nicotinate and nicotinamide metabolism | 6 | 0.82 |
| Transport, nuclear | 6 | 0.82 |
| Pentose and glucuronate interconversions | 5 | 0.68 |
| Propanoate metabolism | 5 | 0.68 |
| Valine, leucine, and isoleucine metabolism | 5 | 0.68 |
| Phenylalanine, tyrosine and tryptophan biosynthesis | 5 | 0.68 |
| C5-branched dibasic acid metabolism | 4 | 0.54 |
| Carnitine shuttle (mitochondrial) | 4 | 0.54 |
| Carnitine shuttle (peroxisomal) | 4 | 0.54 |
| Androgen metabolism | 4 | 0.54 |
| Pantothenate and CoA biosynthesis | 4 | 0.54 |
| Transport, lysosome to ER | 4 | 0.54 |
| Galactose metabolism | 3 | 0.41 |
| Nucleotide metabolism | 3 | 0.41 |
| Cysteine and methionine metabolism | 3 | 0.41 |
| Glutathione metabolism | 3 | 0.41 |
| Carnitine shuttle (cytosolic) | 3 | 0.41 |
| Pyrimidine metabolism | 2 | 0.27 |
| Lysine metabolism | 2 | 0.27 |
| Arachidonic acid metabolism | 2 | 0.27 |
| Carnitine shuttle (endoplasmic reticular) | 2 | 0.27 |
| Steroid metabolism | 2 | 0.27 |
| Estrogen metabolism | 2 | 0.27 |

| | | |
|---|---|---|
| Ether lipid metabolism | 2 | 0.27 |
| Pool reactions | 2 | 0.27 |
| Fructose and Mannose metabolism | 1 | 0.14 |
| Butanoate metabolism | 1 | 0.14 |
| beta-Alanine metabolism | 1 | 0.14 |
| Sulfur metabolism | 1 | 0.14 |
| ROS detoxification | 1 | 0.14 |
| Fatty acid activation (cytosolic) | 1 | 0.14 |
| Fatty acid activation (endoplasmic reticular) | 1 | 0.14 |
| Eicosanoid metabolism | 1 | 0.14 |
| Beta oxidation of even-chain fatty acids (peroxisomal) | 1 | 0.14 |
| Glycerolipid metabolism | 1 | 0.14 |
| Glycerophospholipid metabolism | 1 | 0.14 |
| Porphyrin metabolism | 1 | 0.14 |
| Transport, Golgi apparatus | 1 | 0.14 |

| Table S4: Subsystem distribution: Reactions from global modules (constrained) | | |
|---|---|---|
| **Subsystem** | **Num Reactions** | **% Reactions** |
| Transport, extracellular | 328 | 45.62 |
| Transport, mitochondrial | 68 | 9.46 |
| Exchange reactions | 29 | 4.03 |
| Tricarboxylic acid cycle and glyoxylate/dicarboxylate metabolism | 21 | 2.92 |
| Glycine, serine and threonine metabolism | 17 | 2.36 |
| Transport, endoplasmic reticular | 17 | 2.36 |
| Glycolysis / Gluconeogenesis | 15 | 2.09 |
| Retinol metabolism | 14 | 1.95 |
| Prostaglandin biosynthesis | 13 | 1.81 |
| Folate metabolism | 12 | 1.67 |
| Arginine and proline metabolism | 11 | 1.53 |
| Biopterin metabolism | 10 | 1.39 |
| Transport, peroxisomal | 10 | 1.39 |
| Pyruvate metabolism | 9 | 1.25 |
| Pentose phosphate pathway | 9 | 1.25 |
| Alanine, aspartate and glutamate metabolism | 9 | 1.25 |
| Transport, lysosomal | 8 | 1.11 |
| Purine metabolism | 7 | 0.97 |
| Amino sugar and nucleotide sugar metabolism | 7 | 0.97 |
| Leukotriene metabolism | 7 | 0.97 |
| Tyrosine metabolism | 6 | 0.83 |
| Transport, nuclear | 6 | 0.83 |
| Pentose and glucuronate interconversions | 5 | 0.70 |
| Propanoate metabolism | 5 | 0.70 |
| Phenylalanine, tyrosine and tryptophan biosynthesis | 5 | 0.70 |
| C5-branched dibasic acid metabolism | 4 | 0.56 |
| Carnitine shuttle (mitochondrial) | 4 | 0.56 |
| Carnitine shuttle (peroxisomal) | 4 | 0.56 |
| Androgen metabolism | 4 | 0.56 |
| Pantothenate and CoA biosynthesis | 4 | 0.56 |
| Transport, lysosome to ER | 4 | 0.56 |
| Galactose metabolism | 3 | 0.42 |
| Nucleotide metabolism | 3 | 0.42 |
| Cysteine and methionine metabolism | 3 | 0.42 |
| Glutathione metabolism | 3 | 0.42 |
| Valine, leucine and isoleucine degradation | 3 | 0.42 |
| Carnitine shuttle (cytosolic) | 3 | 0.42 |
| Pyrimidine metabolism | 2 | 0.28 |
| Lysine metabolism | 2 | 0.28 |
| Arachidonic acid metabolism | 2 | 0.28 |
| Carnitine shuttle (endoplasmic reticular) | 2 | 0.28 |
| Steroid metabolism | 2 | 0.28 |
| Estrogen metabolism | 2 | 0.28 |
| Nicotinate and nicotinamide metabolism | 2 | 0.28 |
| Pool reactions | 2 | 0.28 |

| | | |
|---|---|---|
| Fructose and Mannose metabolism | 1 | 0.14 |
| Butanoate metabolism | 1 | 0.14 |
| beta-Alanine metabolism | 1 | 0.14 |
| Sulfur metabolism | 1 | 0.14 |
| ROS detoxification | 1 | 0.14 |
| Fatty acid activation (cytosolic) | 1 | 0.14 |
| Fatty acid activation (endoplasmic reticular) | 1 | 0.14 |
| Eicosanoid metabolism | 1 | 0.14 |
| Beta oxidation of even-chain fatty acids (peroxisomal) | 1 | 0.14 |
| Glycerolipid metabolism | 1 | 0.14 |
| Glycerophospholipid metabolism | 1 | 0.14 |
| Porphyrin metabolism | 1 | 0.14 |
| Transport, Golgi apparatus | 1 | 0.14 |

**TableS5: Table with reactions in the modules that were distinct between the simulations**

| Reaction id | Equation | Subsystem | Type | Mean (Constrained) | Mean (Unconstrained) | NumberOfPCs (Constrained) | NumberOfPCs (Unconstrained) | PC Indices (Constrained) | PC Indices (Unconstrained) | Stddev (Constrained) | Stddev (Unconstrained) |
|---|---|---|---|---|---|---|---|---|---|---|---|
| HMR_9618 | 1-methylnicotinamide[c] <=> 1-methylnicotinamide[s] | Transport, extracellular | un | 0 | 957.4382 | 0 | 1 | null | 472 | 0 | 19.03725 |
| HMR_9270 | <=> L-arabinose[s] | Exchange reactions | both | 489.2168 | 726.8843 | 1 | 1 | 296 | 341 | 207.264 | 134.3966 |
| HMR_9241 | <=> L-arabitol[s] | Exchange reactions | both | -489.2168 | -726.8843 | 1 | 1 | 296 | 341 | 207.264 | 134.3966 |
| HMR_9172 | <=> 1-organyl-2-lyso-sn-glycero-3-phosphocholine[s] | Exchange reactions | both | 554.4444 | 354.9605 | 1 | 1 | 331 | 319 | 112.4165 | 111.4461 |
| HMR_9171 | <=> 2-acetyl-1-alkyl-sn-glycero-3-phosphocholine[s] | Exchange reactions | both | -555.2822 | -355.812 | 1 | 1 | 331 | 319 | 112.4159 | 111.4465 |
| HMR_9168 | <=> glucosamine[s] | Exchange reactions | both | 56.85073 | 61.58986 | 1 | 1 | 426 | 415 | 25.32388 | 27.33342 |
| HMR_9142 | <=> nicotinate[s] | Exchange reactions | un | 0 | 958.6224 | 0 | 1 | null | 472 | 0 | 18.87394 |
| HMR_9139 | <=> fructose[s] | Exchange reactions | both | 121.0896 | 121.9996 | 1 | 1 | 385 | 386 | 50.28207 | 50.44116 |
| HMR_9138 | <=> xylitol[s] | Exchange reactions | both | 47.85797 | 42.08431 | 1 | 1 | 413 | 421 | 28.10508 | 24.78155 |
| HMR_9135 | <=> L-lactate[s] | Exchange reactions | both | -390.1747 | -632.8099 | 1 | 1 | 228 | 366 | 264.0182 | 147.8723 |
| HMR_9134 | <=> (R)-3-hydroxybutanoate[s] | Exchange reactions | both | 232.374 | 118.8781 | 1 | 1 | 350 | 350 | 195.3623 | 124.6017 |
| HMR_9133 | <=> pyruvate[s] | Exchange reactions | both | 566.9194 | 817.1257 | 2 | 1 | 228,350 | 366 | 262.0537 | 143.535 |
| HMR_9132 | <=> acetoacetate[s] | Exchange reactions | both | -261.8774 | -148.9745 | 1 | 1 | 350 | 350 | 195.9525 | 125.6921 |
| HMR_9131 | <=> sarcosine[s] | Exchange reactions | both | 102.4714 | 93.62069 | 1 | 1 | 453 | 385 | 43.41782 | 38.39658 |
| HMR_9104 | <=> 1-methylnicotinamide[s] | Exchange reactions | un | 0 | -960.2962 | 0 | 1 | null | 472 | 0 | 18.86094 |
| HMR_9103 | <=> H2S[s] | Exchange reactions | both | 366.2917 | 293.4265 | 1 | 1 | 379 | 384 | 62.67827 | 68.49313 |
| HMR_9098 | <=> D-alanine[s] | Exchange reactions | both | -401.3674 | -514.2668 | 1 | 1 | 271 | 296 | 298.4308 | 258.7848 |
| HMR_9085 | <=> glycerol[s] | Exchange reactions | both | 69.82899 | 53.09124 | 1 | 1 | 383 | 388 | 55.57647 | 45.15661 |
| HMR_9073 | <=> NH3[s] | Exchange reactions | both | -38.35902 | 344.7089 | 1 | 1 | 398 | 403 | 116.4502 | 115.6215 |
| HMR_9071 | glutamate[s] <=> | Exchange reactions | both | 58.80301 | 72.63386 | 1 | 1 | 391 | 398 | 45.65197 | 50.12043 |
| HMR_9069 | serine[s] => | Exchange reactions | both | 34.84896 | 41.21996 | 1 | 1 | 379 | 384 | 33.62046 | 39.41443 |
| HMR_9068 | proline[s] => | Exchange reactions | con | 905.0507 | null | 1 | 0 | 392 | null | 72.17909 | null |
| HMR_9065 | cysteine[s] => | Exchange reactions | both | 718.6153 | 738.2995 | 1 | 1 | 365 | 363 | 182.7009 | 186.0934 |
| HMR_9056 | NEFA blood pool out[s] => | Exchange reactions | both | 95.28183 | 88.86313 | 1 | 1 | 402 | 409 | 40.72338 | 37.9055 |
| HMR_9046 | => valine[s] | Exchange reactions | un | 0 | 662.7702 | 0 | 1 | null | 343 | 0 | 219.1931 |
| HMR_9040 | =>leucine[s] | Exchange reactions | un | 0 | 498.6294 | 0 | 1 | null | 323 | 0 | 252.5405 |
| HMR_9039 | => isoleucine[s] | Exchange reactions | un | 0 | 504.0355 | 0 | 1 | null | 329 | 0 | 252.524 |
| HMR_9034 | => glucose[s] | Exchange reactions | both | 577.5123 | 576.5929 | 1 | 1 | 426 | 454 | 245.7048 | 245.8971 |
| HMR_9033 | => NEFA blood pool in[s] | Exchange reactions | both | 97.48427 | 87.989 | 1 | 1 | 402 | 409 | 38.09669 | 35.28284 |
| HMR_8687 | cystine[s] + serine[c] => cystine[c] + serine[s] | Transport, extracellular | both | 131.1897 | 160.7384 | 1 | 1 | 391 | 398 | 91.43076 | 101.8652 |
| HMR_8653 | glucono-1,5-lactone-6-phosphate[r] + H+[r] + NADH[r] <=> glucose-6-phosphate[r] + NAD+[r] | Pentose phosphate pathway | un | 0 | 453.2049 | 0 | 1 | null | 338 | 0 | 273.4675 |
| HMR_8514 | 2 ferricytochrome C[m] + L-lactate[c] => 2 ferrocytochrome C[m] + 2 H+[c] + pyruvate[c] | Pyruvate metabolism | un | 0 | 57.82712 | 0 | 2 | null | 503,402 | 0 | 46.17354 |
| HMR_8512 | D-lactate[c] + NAD+[c] <=> H+[c] + NADH[c] + pyruvate[c] | Pyruvate metabolism | both | 16.81031 | -16.33962 | 1 | 1 | 376 | 402 | 35.33717 | 49.26219 |
| HMR_8509 | D-lactaldehyde[c] + H+[c] + NADPH[c] => NADP+[c] + propane-1,2-diol[c] | Pyruvate metabolism | un | 0 | 405.7674 | 0 | 1 | null | 334 | 0 | 240.5668 |
| HMR_8507 | NAD+[c] + propane-1,2-diol[c] => D-lactaldehyde[c] + H+[c] + NADH[c] | Pyruvate metabolism | un | 0 | 411.1352 | 0 | 1 | null | 334 | 0 | 240.5669 |
| HMR_8434 | S-aminomethyldihydrolipoamide[m] + THF[m] => 5,10-methylene-THF[m] + dihydrolipoamide[m] + NH3[m] | Glycine, serine and threonine metabolism | both | 144.9528 | 160.171 | 1 | 1 | 374 | 370 | 90.9065 | 99.90376 |
| HMR_8433 | CO2[m] + S-aminomethyldihydrolipoamide[m] <=> glycine[m] + lipoamide[m] | Glycine, serine and threonine metabolism | both | -144.9535 | -160.1704 | 1 | 1 | 374 | 370 | 90.9065 | 99.90376 |
| HMR_8430 | creatine-phosphate[m] => creatine-phosphate[c] | Arginine and proline metabolism | both | 47.8204 | 39.80966 | 1 | 1 | 432 | 489 | 26.13441 | 21.7718 |
| HMR_8428 | creatine[c] => creatine[m] | Arginine and proline metabolism | both | 47.81811 | 39.80744 | 1 | 1 | 432 | 489 | 26.1345 | 21.77192 |
| HMR_8427 | ADP[m] + creatine-phosphate[m] + 2 H+[m] <=> ATP[m] + creatine[m] | Arginine and proline metabolism | both | -47.81807 | -39.80739 | 1 | 1 | 432 | 489 | 26.1344 | 21.7718 |
| HMR_8377 | CTP[c] + N-acetylneuraminate[c] => CMP-N-acetylneuraminate[c] + PPi[c] | Amino sugar and nucleotide sugar metabolism | con | 214.3499 | null | 1 | 0 | 354 | null | 169.94 | null |
| HMR_8343 | L-arabitol[c] => L-arabitol[s] | Transport, extracellular | both | 489.2168 | 726.8843 | 1 | 1 | 296 | 341 | 207.264 | 134.3965 |
| HMR_8341 | H+[c] + L-arabinose[c] + NADPH[c] => L-arabitol[c] + NADP+[c] | Pentose and glucuronate interconversions | both | 489.2168 | 726.8843 | 1 | 1 | 296 | 341 | 207.264 | 134.3965 |
| HMR_8340 | L-arabinose[c] <=> L-arabinose[s] | Transport, extracellular | both | -489.2168 | -726.8843 | 1 | 1 | 296 | 341 | 207.264 | 134.3965 |
| HMR_7881 | ADP[n] + dITP[n] <=> ATP[n] + dIDP[n] | Nucleotide metabolism | both | 195.8281 | 186.7448 | 1 | 1 | 5 | 11 | 357.8294 | 360.7988 |
| HMR_7878 | ADP[c] + dITP[c] <=> ATP[c] + dIDP[c] | Nucleotide metabolism | both | -195.8193 | -186.7361 | 1 | 1 | 5 | 11 | 357.8294 | 360.7988 |
| HMR_7877 | dIDP[c] <=> dIDP[n] | Transport, nuclear | both | -195.8237 | -186.7404 | 1 | 1 | 5 | 11 | 357.8294 | 360.7988 |
| HMR_7855 | dITP[c] <=> dITP[n] | Transport, nuclear | both | 195.8318 | 186.7485 | 1 | 1 | 5 | 11 | 357.8294 | 360.7988 |
| HMR_7748 | beta-D-glucose-6-phosphate[c] <=> glucose-6-phosphate[c] | Glycolysis / Gluconeogenesis | both | -18.33267 | -19.44595 | 1 | 1 | 422 | 412 | 464.4743 | 464.2657 |
| HMR_7747 | ADP[c] + beta-D-glucose[c] => AMP[c] + beta-D-glucose-6-phosphate[c] | Glycolysis / Gluconeogenesis | both | 40.06949 | 40.36772 | 1 | 1 | 422 | 412 | 34.08285 | 34.30359 |
| HMR_7745 | beta-D-glucose[c] <=> glucose[c] | Glycolysis / Gluconeogenesis | both | -79.72204 | -80.31828 | 1 | 1 | 422 | 412 | 41.83195 | 42.0484 |
| HMR_7676 | H+[c] + nicotinamide[c] + SAM[c] => 1-methylnicotinamide[c] + SAH[c] | Nicotinate and nicotinamide metabolism | un | 0 | 960.2963 | 0 | 1 | null | 472 | 0 | 18.8606 |
| HMR_7643 | D-alanine[c] + glutamine[s] <=> D-alanine[s] + glutamine[c] | Transport, extracellular | both | 407.2401 | 452.2186 | 1 | 1 | 271 | 296 | 423.7198 | 398.0962 |
| HMR_7642 | D-alanine[c] <=> alanine[c] | Alanine, aspartate and glutamate metabolism | both | -401.3674 | -514.2668 | 1 | 1 | 271 | 296 | 298.4308 | 258.7848 |
| HMR_7613 | 1-organyl-2-lyso-sn-glycero-3-phosphocholine[c] + acetyl-CoA[c] => 2-acetyl-1-alkyl-sn-glycero-3-phosphocholine[c] + CoA[c] | Glycerophospholipid metabolism | both | 563.5075 | 364.2057 | 1 | 1 | 331 | 319 | 112.7176 | 111.7811 |
| HMR_7609 | 1-organyl-2-lyso-sn-glycero-3-phosphocholine[c] <=> 1-organyl-2-lyso-sn-glycero-3-phosphocholine[s] | Transport, extracellular | both | -554.4444 | -354.9605 | 1 | 1 | 331 | 319 | 112.4164 | 111.4461 |

| | | | | | | | | | | | |
|---|---|---|---|---|---|---|---|---|---|---|---|
| HMR_7608 | 2-acetyl-1-alkyl-sn-glycero-3-phosphocholine[c] <=> 2-acetyl-1-alkyl-sn-glycero-3-phosphocholine[s] | Transport, extracellular | both | 555.2822 | 355.812 | 1 | 1 | 331 | 319 | 112.4158 | 111.4465 |
| HMR_7253 | xylitol[c] <=> xylitol[s] | Transport, extracellular | both | -47.8579 | -42.08423 | 1 | 1 | 413 | 421 | 28.10501 | 24.78148 |
| HMR_6627 | dTDP[c] + PEP[c] <=> dTTP[c] + pyruvate[c] | Pyrimidine metabolism | both | -611.0316 | -609.1076 | 1 | 2 | 317 | 11,311 | 294.3339 | 295.8587 |
| HMR_6417 | CoA[m] + S-(2-methylpropanoyl)-dihydrolipoamide[m] <=> dihydrolipoamide[m] + isobutyryl-CoA[m] | Valine, leucine, and isoleucine metabolism | un | 0 | 156.8625 | 0 | 1 | null | 372 | 0 | 93.74738 |
| HMR_6416 | 3-methyl-2-oxobutyrate[m] + lipoamide[m] => CO2[m] + S-(2-methylpropanoyl)-dihydrolipoamide[m] | Valine, leucine and isoleucine degradation | un | 0 | 156.8624 | 0 | 1 | null | 372 | 0 | 93.74739 |
| HMR_6414 | CoA[m] + S-succinyldihydrolipoamide[m] <=> dihydrolipoamide[m] + succinyl-CoA[m] | Tricarboxylic acid cycle and glyoxylate/dicarboxylate metabolism | both | 364.2391 | 205.5347 | 1 | 1 | 363 | 379 | 139.1446 | 81.6192 |
| HMR_6412 | CoA[m] + S-acetyldihydrolipoamide[m] => acetyl-CoA[m] + dihydrolipoamide[m] | Glycolysis / Gluconeogenesis | both | 253.5549 | 241.6286 | 1 | 1 | 344 | 359 | 151.9572 | 143.5929 |
| HMR_6411 | AKG[m] + lipoamide[m] => CO2[m] + S-succinyldihydrolipoamide[m] | Tricarboxylic acid cycle and glyoxylate/dicarboxylate metabolism | both | 170.7694 | 96.61792 | 1 | 1 | 363 | 379 | 132.1297 | 75.75637 |
| HMR_6410 | lipoamide[m] + pyruvate[m] => CO2[m] + S-acetyldihydrolipoamide[m] | Glycolysis / Gluconeogenesis | both | 253.556 | 241.6285 | 1 | 1 | 344 | 359 | 151.9572 | 143.5929 |
| HMR_6409 | H+[m] + lipoamide[m] + NADH[m] <=> dihydrolipoamide[m] + NAD+[m] | Glycine, serine and threonine metabolism | both | -778.8089 | -781.347 | 3 | 4 | 374,363,344 | 370,379,372,359 | 162.7066 | 156.5743 |
| HMR_6330 | AKG[c] + Pi[m] <=> AKG[m] + Pi[c] | Transport, mitochondrial | un | 0 | 142.9513 | 0 | 1 | null | 132 | 0 | 523.2926 |
| HMR_6325 | H+[c] + histidine[c] => H+[m] + histidine[m] | Transport, mitochondrial | un | 0 | 95.29811 | 0 | 1 | null | 489 | 0 | 61.33427 |
| HMR_6323 | arginine[c] + H+[c] => arginine[m] + H+[m] | Transport, mitochondrial | un | 0 | 95.29812 | 0 | 1 | null | 489 | 0 | 61.33427 |
| HMR_6317 | citrulline[m] + H+[c] + ornithine[c] => citrulline[c] + H+[m] + ornithine[m] | Transport, mitochondrial | both | 54.89813 | 47.10501 | 1 | 1 | 450 | 489 | 45.89212 | 39.31503 |
| HMR_6315 | citrulline[c] + H+[c] + histidine[m] => citrulline[m] + H+[m] + histidine[c] | Transport, mitochondrial | both | 60.74043 | 52.19804 | 1 | 1 | 429 | 456 | 49.7412 | 42.7329 |
| HMR_6314 | H+[c] + histidine[m] + ornithine[c] => H+[m] + histidine[c] + ornithine[m] | Transport, mitochondrial | both | 72.71464 | 62.36928 | 1 | 1 | 415 | 447 | 57.57557 | 49.19051 |
| HMR_6311 | arginine[m] + citrulline[c] + H+[c] => arginine[c] + citrulline[m] + H+[m] | Transport, mitochondrial | both | 60.74042 | 52.19804 | 1 | 2 | 431 | 456,489 | 49.7412 | 42.73289 |
| HMR_6310 | arginine[c] + H+[c] + histidine[m] => arginine[m] + H+[m] + histidine[c] | Transport, mitochondrial | con | 51.13353 | null | 1 | 0 | 452 | null | 45.48275 | null |
| HMR_6309 | arginine[m] + H+[c] + histidine[c] => arginine[c] + H+[m] + histidine[m] | Transport, mitochondrial | con | 51.13353 | null | 1 | 0 | 452 | null | 45.48275 | null |
| HMR_6307 | arginine[m] + H+[c] + ornithine[c] => arginine[c] + H+[m] + ornithine[m] | Transport, mitochondrial | both | 72.71463 | 62.36928 | 1 | 1 | 424 | 427 | 57.57557 | 49.19051 |
| HMR_6306 | citrulline[m] + H+[c] + lysine[c] => citrulline[c] + H+[m] + lysine[m] | Transport, mitochondrial | con | 49.71444 | null | 1 | 0 | 504 | null | 42.19003 | null |
| HMR_6305 | citrulline[c] + H+[c] + lysine[m] => citrulline[m] + H+[m] + lysine[c] | Transport, mitochondrial | con | 46.50197 | null | 1 | 0 | 504 | null | 40.09299 | null |
| HMR_6304 | H+[c] + histidine[m] + lysine[c] => H+[m] + histidine[c] + lysine[m] | Transport, mitochondrial | both | 63.47167 | 54.94319 | 1 | 1 | 367 | 449 | 51.37876 | 44.38068 |
| HMR_6301 | H+[c] + lysine[m] + ornithine[c] => H+[m] + lysine[c] + ornithine[m] | Transport, mitochondrial | con | 52.99465 | null | 1 | 0 | 488 | null | 44.77604 | null |
| HMR_6300 | arginine[m] + H+[c] + lysine[c] => arginine[c] + H+[m] + lysine[m] | Transport, mitochondrial | both | 63.47167 | 54.94319 | 1 | 1 | 425 | 453 | 51.37876 | 44.38068 |
| HMR_6058 | chloride[s] => chloride[c] | Transport, extracellular | both | 398.0496 | 426.6178 | 1 | 1 | 316 | 315 | 224.7485 | 220.0512 |
| HMR_6054 | (R)-3-hydroxybutanoate[c] + 2-hydroxybutyrate[s] <=> (R)-3-hydroxybutanoate[s] + 2-hydroxybutyrate[c] | Transport, extracellular | both | -20.2828 | -10.71901 | 1 | 1 | 350 | 297 | 554.7127 | 554.7848 |
| HMR_6049 | L-lactate[c] + retinoate[s] <=> L-lactate[s] + retinoate[c] | Transport, extracellular | con | 36.07198 | null | 1 | 0 | 42 | null | 554.3419 | null |
| HMR_6043 | butyrate[c] + propanoate[s] <=> butyrate[s] + propanoate[c] | Transport, extracellular | un | 0 | -4.46E-04 | 0 | 1 | null | 292 | 0 | 554.7957 |
| HMR_6039 | butyrate[c] + formate[s] <=> butyrate[s] + formate[c] | Transport, extracellular | con | 3.29E-05 | null | 1 | 0 | 303 | null | 554.8097 | null |
| HMR_6037 | formate[c] + propanoate[s] <=> formate[s] + propanoate[c] | Transport, extracellular | un | 0 | -4.49E-04 | 0 | 1 | null | 252 | 0 | 554.7957 |
| HMR_6033 | acetate[s] + L-lactate[c] <=> acetate[c] + L-lactate[s] | Transport, extracellular | un | 0 | 46.92303 | 0 | 1 | null | 32 | 0 | 553.6739 |
| HMR_6026 | AKG[c] + L-lactate[s] <=> AKG[s] + L-lactate[c] | Transport, extracellular | un | 0 | -58.02741 | 0 | 1 | null | 135 | 0 | 553.0889 |
| HMR_6025 | acetoacetate[s] + AKG[c] <=> acetoacetate[c] + AKG[s] | Transport, extracellular | un | 0 | -13.9525 | 0 | 1 | null | 141 | 0 | 554.7424 |
| HMR_6021 | (R)-3-hydroxybutanoate[c] + pyruvate[s] <=> (R)-3-hydroxybutanoate[s] + pyruvate[c] | Transport, extracellular | both | 30.55049 | 63.70129 | 1 | 1 | 24 | 350 | 554.4315 | 552.695 |
| HMR_6015 | acetate[c] + pyruvate[s] <=> acetate[s] + pyruvate[c] | Transport, extracellular | con | 62.48395 | null | 1 | 0 | 203 | null | 553.0052 | null |
| HMR_6013 | 2-hydroxybutyrate[c] + retinoate[s] <=> 2-hydroxybutyrate[s] + retinoate[c] | Transport, extracellular | un | 0 | 0.4234824 | 0 | 1 | null | 302 | 0 | 554.7964 |
| HMR_6011 | 2-hydroxybutyrate[c] + L-lactate[s] <=> 2-hydroxybutyrate[s] + L-lactate[c] | Transport, extracellular | con | -36.42105 | null | 1 | 0 | 294 | null | 554.3329 | null |
| HMR_6010 | 2-hydroxybutyrate[c] + propanoate[s] <=> 2-hydroxybutyrate[s] + propanoate[c] | Transport, extracellular | un | 0 | 0.423004 | 0 | 1 | null | 65 | 0 | 554.7965 |
| HMR_6007 | 2-hydroxybutyrate[c] + pyruvate[s] <=> 2-hydroxybutyrate[s] + pyruvate[c] | Transport, extracellular | un | 0 | 74.31668 | 0 | 1 | null | 167 | 0 | 551.9496 |
| HMR_6002 | acetoacetate[c] + formate[s] <=> acetoacetate[s] + formate[c] | Transport, extracellular | con | 24.38282 | null | 1 | 0 | 93 | null | 554.6162 | null |
| HMR_6000 | acetoacetate[c] + pyruvate[s] <=> acetoacetate[s] + pyruvate[c] | Transport, extracellular | both | 75.15149 | 87.77119 | 1 | 1 | 350 | 350 | 552.393 | 550.9327 |
| HMR_5990 | chloride[c] + K+[c] => chloride[s] + K+[s] | Transport, extracellular | both | 602.9554 | 637.095 | 1 | 1 | 316 | 315 | 224.2641 | 218.9674 |
| HMR_5986 | citrulline[s] + isoleucine[c] => citrulline[c] + isoleucine[s] | Transport, extracellular | both | 43.22554 | 41.9753 | 1 | 1 | 482 | 486 | 46.52755 | 45.15149 |
| HMR_5983 | citrulline[s] + valine[c] => citrulline[c] + valine[s] | Transport, extracellular | un | 0 | 41.88418 | 0 | 1 | null | 444 | 0 | 45.0377 |
| HMR_5978 | citrulline[s] + tyrosine[c] => citrulline[c] + tyrosine[s] | Transport, extracellular | un | 0 | 41.97087 | 0 | 1 | null | 439 | 0 | 45.14604 |
| HMR_5976 | citrulline[s] + tryptophan[c] => citrulline[c] + tryptophan[s] | Transport, extracellular | un | 0 | 41.9751 | 0 | 1 | null | 441 | 0 | 45.15123 |
| HMR_5975 | citrulline[s] + methionine[c] => citrulline[c] + methionine[s] | Transport, extracellular | both | 43.14536 | 41.90492 | 1 | 1 | 444 | 443 | 46.43206 | 45.06798 |
| HMR_5974 | citrulline[s] + serine[c] => citrulline[c] + serine[s] | Transport, extracellular | un | 0 | 42.17782 | 0 | 1 | null | 438 | 0 | 45.40493 |
| HMR_5970 | histidine[s] + isoleucine[c] => histidine[c] + isoleucine[s] | Transport, extracellular | un | 0 | 42.53084 | 0 | 1 | null | 471 | 0 | 45.75107 |
| HMR_5969 | histidine[s] + homoserine[c] => histidine[c] + homoserine[s] | Transport, extracellular | both | 43.95142 | 42.53832 | 1 | 1 | 469 | 468 | 47.33694 | 45.76034 |
| HMR_5968 | histidine[s] + threonine[c] => histidine[c] + threonine[s] | Transport, extracellular | un | 0 | 42.75802 | 0 | 1 | null | 466 | 0 | 46.03684 |
| HMR_5967 | histidine[s] + valine[c] => histidine[c] + valine[s] | Transport, extracellular | both | 43.94661 | 42.43732 | 1 | 1 | 467 | 474 | 47.33096 | 45.63414 |
| HMR_5964 | histidine[s] + leucine[c] => histidine[c] + leucine[s] | Transport, extracellular | both | 43.7515 | 42.34891 | 1 | 1 | 472 | 479 | 47.09387 | 45.53055 |
| HMR_5961 | histidine[s] + phenylalanine[c] => histidine[c] + phenylalanine[s] | Transport, extracellular | un | 0 | 42.52876 | 0 | 1 | null | 470 | 0 | 45.74847 |
| HMR_5960 | histidine[s] + tryptophan[c] => histidine[c] + tryptophan[s] | Transport, extracellular | con | 43.94375 | null | 1 | 0 | 468 | null | 47.3274 | null |
| HMR_5959 | histidine[s] + methionine[c] => histidine[c] + methionine[s] | Transport, extracellular | un | 0 | 42.45858 | 0 | 1 | null | 473 | 0 | 45.66538 |
| HMR_5958 | histidine[s] + serine[c] => histidine[c] + serine[s] | Transport, extracellular | both | 44.1702 | 42.73879 | 1 | 1 | 458 | 465 | 47.61262 | 46.01124 |
| HMR_5955 | glycine[c] + histidine[s] => glycine[s] + histidine[c] | Transport, extracellular | both | 44.66982 | 43.22415 | 1 | 1 | 454 | 457 | 48.23277 | 46.6128 |
| HMR_5954 | isoleucine[c] + ornithine[s] => isoleucine[s] + ornithine[c] | Transport, extracellular | un | 0 | 41.98555 | 0 | 1 | null | 442 | 0 | 45.16397 |

| | | | | | | | | | | | |
|---|---|---|---|---|---|---|---|---|---|---|---|
| HMR_5952 | ornithine[s] + threonine[c] => ornithine[c] + threonine[s] | Transport, extracellular | un | 0 | 42.20694 | 0 | 1 | null | 434 | 0 | 45.44238 |
| HMR_5951 | ornithine[s] + valine[c] => ornithine[c] + valine[s] | Transport, extracellular | both | 43.23237 | 41.89439 | 1 | 1 | 442 | 488 | 46.53499 | 45.05012 |
| HMR_5947 | cysteine[c] + ornithine[s] => cysteine[s] + ornithine[c] | Transport, extracellular | un | 0 | 42.62784 | 0 | 1 | null | 432 | 0 | 45.97306 |
| HMR_5945 | ornithine[s] + phenylalanine[c] => ornithine[c] + phenylalanine[s] | Transport, extracellular | both | 43.22776 | 41.98352 | 1 | 1 | 441 | 485 | 46.52921 | 45.16144 |
| HMR_5944 | ornithine[s] + tryptophan[c] => ornithine[c] + tryptophan[s] | Transport, extracellular | con | 43.22961 | null | 1 | 0 | 438 | null | 46.53155 | null |
| HMR_5943 | methionine[c] + ornithine[s] => methionine[s] + ornithine[c] | Transport, extracellular | both | 43.15221 | 41.91514 | 1 | 1 | 485 | 487 | 46.43952 | 45.08042 |
| HMR_5942 | ornithine[s] + serine[c] => ornithine[c] + serine[s] | Transport, extracellular | both | 43.44878 | 42.18817 | 1 | 1 | 435 | 478 | 46.80715 | 45.41757 |
| HMR_5939 | glycine[c] + ornithine[s] => glycine[s] + ornithine[c] | Transport, extracellular | both | 43.93221 | 42.66106 | 1 | 1 | 430 | 433 | 47.40611 | 46.00349 |
| HMR_5938 | arginine[s] + isoleucine[c] => arginine[c] + isoleucine[s] | Transport, extracellular | con | 41.63078 | null | 1 | 0 | 514 | null | 44.54515 | null |
| HMR_5937 | arginine[s] + homoserine[c] => arginine[c] + homoserine[s] | Transport, extracellular | un | 0 | 40.47552 | 0 | 1 | null | 514 | 0 | 43.29356 |
| HMR_5936 | arginine[s] + threonine[c] => arginine[c] + threonine[s] | Transport, extracellular | both | 41.84471 | 40.67478 | 1 | 1 | 509 | 508 | 44.81 | 43.53996 |
| HMR_5934 | arginine[s] + asparagine[c] => arginine[c] + asparagine[s] | Transport, extracellular | both | 41.85763 | 40.68794 | 1 | 1 | 508 | 509 | 44.8258 | 43.55603 |
| HMR_5933 | arginine[s] + proline[c] => arginine[c] + proline[s] | Transport, extracellular | both | 41.80102 | 40.66303 | 1 | 1 | 510 | 513 | 44.75335 | 43.5247 |
| HMR_5931 | arginine[s] + cysteine[c] => arginine[c] + cysteine[s] | Transport, extracellular | both | 42.25196 | 41.06631 | 1 | 1 | 503 | 506 | 45.31388 | 44.02499 |
| HMR_5930 | arginine[s] + tyrosine[c] => arginine[c] + tyrosine[s] | Transport, extracellular | both | 41.62413 | 40.46461 | 1 | 1 | 513 | 516 | 44.53705 | 43.28031 |
| HMR_5929 | arginine[s] + phenylalanine[c] => arginine[c] + phenylalanine[s] | Transport, extracellular | un | 0 | 40.46685 | 0 | 1 | null | 517 | 0 | 43.28297 |
| HMR_5928 | arginine[s] + tryptophan[c] => arginine[c] + tryptophan[s] | Transport, extracellular | un | 0 | 40.46855 | 0 | 1 | null | 515 | 0 | 43.28506 |
| HMR_5927 | arginine[s] + methionine[c] => arginine[c] + methionine[s] | Transport, extracellular | un | 0 | 40.40324 | 0 | 1 | null | 518 | 0 | 43.20874 |
| HMR_5926 | arginine[s] + serine[c] => arginine[c] + serine[s] | Transport, extracellular | both | 41.83178 | 40.6573 | 1 | 1 | 512 | 511 | 44.79343 | 43.5173 |
| HMR_5925 | arginine[s] + glutamine[c] => arginine[c] + glutamine[s] | Transport, extracellular | both | 41.82243 | 40.66467 | 1 | 1 | 511 | 510 | 44.77371 | 43.51948 |
| HMR_5924 | alanine[c] + arginine[s] => alanine[s] + arginine[c] | Transport, extracellular | both | 41.9378 | 40.78877 | 1 | 1 | 505 | 507 | 44.92272 | 43.67843 |
| HMR_5923 | arginine[s] + glycine[c] => arginine[c] + glycine[s] | Transport, extracellular | both | 42.28053 | 41.0971 | 1 | 1 | 501 | 504 | 45.33963 | 44.05298 |
| HMR_5922 | isoleucine[c] + lysine[s] => isoleucine[s] + lysine[c] | Transport, extracellular | both | 42.44266 | 41.24104 | 1 | 1 | 498 | 392 | 45.47244 | 44.16807 |
| HMR_5921 | homoserine[c] + lysine[s] => homoserine[s] + lysine[c] | Transport, extracellular | both | 42.44712 | 41.24809 | 1 | 1 | 496 | 497 | 45.47789 | 44.17669 |
| HMR_5919 | lysine[s] + valine[c] => lysine[c] + valine[s] | Transport, extracellular | un | 0 | 41.15296 | 0 | 1 | null | 502 | 0 | 44.05948 |
| HMR_5917 | lysine[s] + proline[c] => lysine[c] + proline[s] | Transport, extracellular | both | 42.61952 | 41.44274 | 1 | 1 | 492 | 495 | 45.6897 | 44.41765 |
| HMR_5916 | leucine[c] + lysine[s] => leucine[s] + lysine[c] | Transport, extracellular | con | 42.26037 | null | 1 | 0 | 502 | null | 45.25411 | null |
| HMR_5914 | lysine[s] + tyrosine[c] => lysine[c] + tyrosine[s] | Transport, extracellular | both | 42.43575 | 41.23676 | 1 | 1 | 497 | 498 | 45.464 | 44.16287 |
| HMR_5913 | lysine[s] + phenylalanine[c] => lysine[c] + phenylalanine[s] | Transport, extracellular | both | 42.43816 | 41.23908 | 1 | 1 | 499 | 500 | 45.4669 | 44.16565 |
| HMR_5912 | lysine[s] + tryptophan[c] => lysine[c] + tryptophan[s] | Transport, extracellular | un | 0 | 41.24084 | 0 | 1 | null | 499 | 0 | 44.16782 |
| HMR_5911 | lysine[s] + methionine[c] => lysine[c] + methionine[s] | Transport, extracellular | both | 42.36527 | 41.17304 | 1 | 1 | 500 | 501 | 45.38146 | 44.08834 |
| HMR_5907 | glycine[c] + lysine[s] => glycine[s] + lysine[c] | Transport, extracellular | both | 43.11793 | 41.89354 | 1 | 1 | 478 | 481 | 46.30134 | 44.96836 |
| HMR_5584 | arginine[c] + citrulline[s] <=> arginine[s] + citrulline[c] | Transport, extracellular | un | 0 | 389.541 | 0 | 1 | null | 186 | 0 | 430.4238 |
| HMR_5583 | arginine[c] + histidine[s] <=> arginine[s] + histidine[c] | Transport, extracellular | un | 0 | 453.0517 | 0 | 1 | null | 235 | 0 | 416.1791 |
| HMR_5582 | arginine[c] + ornithine[s] <=> arginine[s] + ornithine[c] | Transport, extracellular | un | 0 | 391.2473 | 0 | 1 | null | 383 | 0 | 429.843 |
| HMR_5580 | histidine[c] + lysine[s] <=> histidine[s] + lysine[c] | Transport, extracellular | con | -337.0135 | null | 1 | 0 | 390 | null | 464.5443 | null |
| HMR_5578 | arginine[c] + lysine[s] <=> arginine[s] + lysine[c] | Transport, extracellular | both | 220.4437 | 220.9805 | 1 | 1 | 74 | 394 | 483.2799 | 483.8799 |
| HMR_5570 | asparagine[c] + homoserine[s] <=> asparagine[s] + homoserine[c] | Transport, extracellular | con | 46.78038 | null | 1 | 0 | 205 | null | 561.5343 | null |
| HMR_5562 | isoleucine[c] + leucine[s] <=> isoleucine[s] + leucine[c] | Transport, extracellular | con | 37.00981 | null | 1 | 0 | 219 | null | 561.8308 | null |
| HMR_5560 | leucine[c] + threonine[s] <=> leucine[s] + threonine[c] | Transport, extracellular | un | 0 | -82.55016 | 0 | 1 | null | 46 | 0 | 558.7539 |
| HMR_5554 | cysteine[c] + threonine[s] <=> cysteine[s] + threonine[c] | Transport, extracellular | con | 84.40566 | null | 1 | 0 | 151 | null | 558.2503 | null |
| HMR_5543 | leucine[c] + tyrosine[s] <=> leucine[s] + tyrosine[c] | Transport, extracellular | con | -35.60913 | null | 1 | 0 | 210 | null | 561.8822 | null |
| HMR_5541 | isoleucine[c] + phenylalanine[s] <=> isoleucine[s] + phenylalanine[c] | Transport, extracellular | con | 0.9135497 | null | 1 | 0 | 53 | null | 562.9365 | null |
| HMR_5535 | leucine[c] + phenylalanine[s] <=> leucine[s] + phenylalanine[c] | Transport, extracellular | un | 0 | -36.40831 | 0 | 1 | null | 276 | 0 | 561.8343 |
| HMR_5534 | cysteine[c] + phenylalanine[s] <=> cysteine[s] + phenylalanine[c] | Transport, extracellular | con | 129.2761 | null | 1 | 0 | 182 | null | 553.6195 | null |
| HMR_5529 | tryptophan[c] + valine[s] <=> tryptophan[s] + valine[c] | Transport, extracellular | con | -0.541813 | null | 1 | 0 | 158 | null | 562.9368 | null |
| HMR_5522 | isoleucine[c] + methionine[s] <=> isoleucine[s] + methionine[c] | Transport, extracellular | un | 0 | 14.52603 | 0 | 1 | null | 66 | 0 | 562.4985 |
| HMR_5516 | leucine[c] + methionine[s] <=> leucine[s] + methionine[c] | Transport, extracellular | un | 0 | -22.35993 | 0 | 1 | null | 154 | 0 | 561.9339 |
| HMR_5512 | methionine[c] + tryptophan[s] <=> methionine[s] + tryptophan[c] | Transport, extracellular | con | -15.06535 | null | 1 | 0 | 146 | null | 562.5016 | null |
| HMR_5510 | homoserine[c] + serine[s] <=> homoserine[s] + serine[c] | Transport, extracellular | un | 0 | -40.43595 | 0 | 1 | null | 293 | 0 | 561.8201 |
| HMR_5509 | serine[c] + threonine[s] <=> serine[s] + threonine[c] | Transport, extracellular | both | -2.734026 | -3.922645 | 1 | 1 | 171 | 489 | 562.4021 | 562.3936 |
| HMR_5508 | serine[c] + valine[s] <=> serine[s] + valine[c] | Transport, extracellular | both | 42.27887 | 60.8669 | 1 | 1 | 169 | 489 | 561.7511 | 560.755 |
| HMR_5507 | asparagine[c] + serine[s] <=> asparagine[s] + serine[c] | Transport, extracellular | un | 0 | 6.835033 | 0 | 1 | null | 282 | 0 | 562.3762 |
| HMR_5506 | proline[c] + serine[s] <=> proline[s] + serine[c] | Transport, extracellular | both | -6.534242 | 1.28573 | 1 | 1 | 273 | 158 | 562.3936 | 562.404 |
| HMR_5503 | serine[c] + tyrosine[s] <=> serine[s] + tyrosine[c] | Transport, extracellular | un | 0 | 42.86239 | 0 | 1 | null | 253 | 0 | 561.7161 |
| HMR_5501 | serine[c] + tryptophan[s] <=> serine[s] + tryptophan[c] | Transport, extracellular | un | 0 | 41.98873 | 0 | 1 | null | 287 | 0 | 561.7538 |
| HMR_5499 | glutamine[c] + isoleucine[s] <=> glutamine[s] + isoleucine[c] | Transport, extracellular | un | 0 | 43.13488 | 0 | 1 | null | 216 | 0 | 562.446 |
| HMR_5492 | cysteine[c] + glutamine[s] <=> cysteine[s] + glutamine[c] | Transport, extracellular | con | 88.97525 | null | 1 | 0 | 264 | null | 558.6005 | null |
| HMR_5485 | alanine[c] + homoserine[s] <=> alanine[s] + homoserine[c] | Transport, extracellular | con | 63.30432 | null | 1 | 0 | 255 | null | 560.72 | null |
| HMR_5484 | alanine[c] + threonine[s] <=> alanine[s] + threonine[c] | Transport, extracellular | con | 19.37799 | null | 1 | 0 | 17 | null | 562.3093 | null |
| HMR_5481 | alanine[c] + proline[s] <=> alanine[s] + proline[c] | Transport, extracellular | un | 0 | 27.75344 | 0 | 1 | null | 147 | 0 | 562.0679 |
| HMR_5478 | alanine[c] + tyrosine[s] <=> alanine[s] + tyrosine[c] | Transport, extracellular | con | 65.59202 | null | 1 | 0 | 195 | null | 560.5691 | null |
| HMR_5477 | alanine[c] + phenylalanine[s] <=> alanine[s] + phenylalanine[c] | Transport, extracellular | both | 65.10797 | 71.16915 | 1 | 1 | 376 | 41 | 560.6011 | 560.1318 |
| HMR_5474 | alanine[c] + serine[s] <=> alanine[s] + serine[c] | Transport, extracellular | both | 22.1063 | 29.0342 | 1 | 1 | 199 | 270 | 562.267 | 562.0464 |
| HMR_5471 | glycine[c] + homoserine[s] <=> glycine[s] + homoserine[c] | Transport, extracellular | con | 132.2009 | null | 1 | 0 | 136 | null | 554.1658 | null |
| HMR_5467 | glycine[c] + proline[s] <=> glycine[s] + proline[c] | Transport, extracellular | con | 98.48464 | null | 1 | 0 | 155 | null | 557.8854 | null |

| | | | | | | | | | | | |
|---|---|---|---|---|---|---|---|---|---|---|---|
| HMR_5463 | glycine[c] + phenylalanine[s] <=> glycine[s] + phenylalanine[c] | Transport, extracellular | con | 133.951 | null | 1 | 0 | 223 | null | 553.9263 | null |
| HMR_5460 | glycine[c] + serine[s] <=> glycine[s] + serine[c] | Transport, extracellular | con | 92.08562 | null | 1 | 0 | 251 | null | 558.5111 | null |
| HMR_5459 | glutamine[c] + glycine[s] <=> glutamine[s] + glycine[c] | Transport, extracellular | both | -93.51684 | -93.43708 | 1 | 1 | 405 | 248 | 559.9399 | 559.853 |
| HMR_5458 | alanine[c] + glycine[s] <=> alanine[s] + glycine[c] | Transport, extracellular | con | -70.23188 | null | 1 | 0 | 147 | null | 560.5337 | null |
| HMR_5433 | K+[c] => K+[s] | Transport, extracellular | both | 500.722 | 591.0727 | 1 | 1 | 396 | 395 | 230.0699 | 224.466 |
| HMR_5426 | acetoacetate[c] + pyruvate[m] <=> acetoacetate[m] + pyruvate[c] | Transport, mitochondrial | both | -577.3467 | -518.5006 | 1 | 1 | 338 | 322 | 254.6541 | 252.0767 |
| HMR_5411 | H+[c] + L-lactate[c] <=> H+[m] + L-lactate[m] | Transport, mitochondrial | both | -275.1234 | -419.0729 | 1 | 1 | 133 | 406 | 310.4453 | 283.3715 |
| HMR_5297 | AKG[m] + CoA[m] + NAD+[m] => CO2[m] + H+[m] + NADH[m] + succinyl-CoA[m] | Tricarboxylic acid cycle and glyoxylate/dicarboxylate metabolism | both | 239.2187 | 114.8465 | 1 | 2 | 363 | 379,489 | 140.3659 | 81.61512 |
| HMR_5126 | H+[c] + proline[m] => H+[m] + proline[c] | Transport, mitochondrial | both | 709.675 | 877.4688 | 1 | 1 | 447 | 381 | 90.65889 | 60.48343 |
| HMR_5124 | glycine[c] <=> glycine[m] | Transport, mitochondrial | both | -376.252 | -294.8093 | 1 | 1 | 376 | 455 | 37.86944 | 40.38826 |
| HMR_5122 | glutamate[c] + H+[c] => glutamate[m] + H+[m] | Transport, mitochondrial | both | 159.0627 | 81.4758 | 2 | 2 | 391,376 | 398,489 | 79.42198 | 53.2272 |
| HMR_5087 | proline[c] => proline[s] | Transport, extracellular | both | 589.2851 | 593.5668 | 1 | 1 | 347 | 352 | 277.8997 | 277.0602 |
| HMR_5070 | histidine[c] <=> histidine[s] | Transport, extracellular | both | 957.8291 | 959.1877 | 1 | 1 | 507 | 512 | 45.21624 | 43.70639 |
| HMR_5029 | glucose[c] <=> glucose[s] | Transport, extracellular | con | -177.0513 | null | 2 | 0 | 385,426 | null | 55.75087 | null |
| HMR_5016 | (R)-3-hydroxybutanoate[c] + H+[c] => (R)-3-hydroxybutanoate[m] + H+[m] | Transport, mitochondrial | both | 263.4443 | 147.3204 | 1 | 1 | 350 | 350 | 194.6159 | 123.0397 |
| HMR_4996 | glucosamine[s] => glucosamine[c] | Transport, extracellular | both | 56.85071 | 61.58985 | 1 | 1 | 426 | 415 | 25.32372 | 27.33328 |
| HMR_4990 | H2S[c] <=> H2S[s] | Transport, extracellular | both | -366.2917 | -293.4265 | 1 | 1 | 379 | 384 | 62.67819 | 68.49305 |
| HMR_4989 | nicotinate[c] <=> nicotinate[s] | Transport, extracellular | un | 0 | -958.6225 | 0 | 1 | null | 472 | 0 | 18.87374 |
| HMR_4982 | sarcosine[c] <=> sarcosine[s] | Transport, extracellular | both | -102.4714 | -93.62069 | 1 | 1 | 453 | 385 | 43.41771 | 38.39645 |
| HMR_4977 | acetoacetate[c] + H+[c] => acetoacetate[m] + H+[m] | Transport, mitochondrial | both | 317.0525 | 371.2317 | 2 | 2 | 338,350 | 322,350 | 221.6106 | 240.3783 |
| HMR_4972 | citrate[c] + isocitrate[m] <=> citrate[m] + isocitrate[c] | Transport, mitochondrial | con | -72.2804 | null | 1 | 0 | 118 | null | 526.7408 | null |
| HMR_4931 | cystine[s] + glutamate[c] => cystine[c] + glutamate[s] | Transport, extracellular | both | 65.6381 | 79.64696 | 1 | 1 | 391 | 398 | 45.46773 | 50.16068 |
| HMR_4926 | H+[c] + pyruvate[c] => H+[m] + pyruvate[m] | Transport, mitochondrial | both | 408.1788 | 425.2217 | 2 | 1 | 338,133 | 448 | 258.1049 | 260.4819 |
| HMR_4874 | NH3[c] <=> NH3[m] | Transport, mitochondrial | both | 881.6842 | 958.335 | 1 | 1 | 405 | 503 | 64.51587 | 35.12937 |
| HMR_4873 | NH3[c] <=> NH3[s] | Transport, extracellular | both | 21.81527 | -362.0261 | 1 | 1 | 398 | 403 | 116.9498 | 116.2669 |
| HMR_4867 | malate[c] + sulfate[m] <=> malate[m] + sulfate[c] | Transport, mitochondrial | un | 0 | -31.42297 | 0 | 1 | null | 120 | 0 | 524.8733 |
| HMR_4865 | malate[m] + Pi[c] <=> malate[c] + Pi[m] | Transport, mitochondrial | both | -40.72722 | -21.49642 | 1 | 1 | 104 | 114 | 535.1366 | 535.9307 |
| HMR_4863 | succinate[c] + sulfate[m] <=> succinate[m] + sulfate[c] | Transport, mitochondrial | un | 0 | 21.24562 | 0 | 1 | null | 120 | 0 | 521.5053 |
| HMR_4855 | fumarate[c] + malate[m] <=> fumarate[m] + malate[c] | Transport, mitochondrial | both | 63.98267 | 62.87346 | 1 | 2 | 142 | 503,489 | 541.3968 | 541.5198 |
| HMR_4852 | AKG[c] + malate[m] <=> AKG[m] + malate[c] | Transport, mitochondrial | both | 142.7941 | 133.6729 | 1 | 1 | 98 | 489 | 507.9562 | 509.3626 |
| HMR_4851 | AKG[c] + succinate[m] <=> AKG[m] + succinate[c] | Transport, mitochondrial | un | 0 | 82.13499 | 0 | 1 | null | 120 | 0 | 512.1946 |
| HMR_4733 | (R)-4-phosphopantothenoyl-cysteine[m] <=> CO2[m] + phosphopantetheine[m] | Pantothenate and CoA biosynthesis | both | -762.7353 | -748.6516 | 1 | 1 | 356 | 347 | 107.3392 | 113.4643 |
| HMR_4731 | (R)-4-phosphopantothenoyl-cysteine[m] + ADP[m] <=> ATP[m] + N-pantothenoylcysteine[m] | Pantothenate and CoA biosynthesis | both | 762.7358 | 748.652 | 1 | 1 | 356 | 347 | 107.3392 | 113.4643 |
| HMR_4729 | N-pantothenoylcysteine[c] <=> N-pantothenoylcysteine[m] | Transport, mitochondrial | both | -762.7357 | -748.652 | 1 | 1 | 356 | 347 | 107.3392 | 113.4643 |
| HMR_4720 | pantetheine[c] <=> pantetheine[m] | Transport, mitochondrial | both | 762.7361 | 748.6524 | 1 | 1 | 356 | 347 | 107.3392 | 113.4643 |
| HMR_4716 | N-pantothenoylcysteine[c] => CO2[c] + pantetheine[c] | Pantothenate and CoA biosynthesis | both | 762.7355 | 748.6519 | 1 | 1 | 356 | 347 | 107.3392 | 113.4642 |
| HMR_4715 | ATP[m] + pantetheine[m] => ADP[m] + phosphopantetheine[m] | Pantothenate and CoA biosynthesis | both | 762.7367 | 748.653 | 1 | 1 | 356 | 347 | 107.3392 | 113.4643 |
| HMR_4701 | dimethylglycine[c] + SAH[c] => SAM[c] + sarcosine[c] | Sulfur metabolism | both | 104.8067 | 96.34416 | 1 | 1 | 453 | 385 | 42.92639 | 38.0467 |
| HMR_4700 | formaldehyde[c] + H+[c] + NADPH[c] + sarcosine[c] <=> dimethylglycine[c] + H2O[c] + NADP+[c] | Glycine, serine and threonine metabolism | both | 101.7879 | 93.3976 | 1 | 1 | 453 | 385 | 42.95502 | 38.07699 |
| HMR_4595 | ADP[c] + D-xylulose-5-phosphate[c] <=> ATP[c] + D-xylulose[c] | Pentose and glucuronate interconversions | both | -19.53053 | -14.78347 | 1 | 1 | 413 | 421 | 31.35904 | 28.29092 |
| HMR_4593 | NAD+[c] + xylitol[c] => D-xylulose[c] + H+[c] + NADH[c] | Pentose and glucuronate interconversions | both | 47.85764 | 42.08396 | 1 | 1 | 413 | 421 | 28.10511 | 24.78159 |
| HMR_4589 | cis-aconitate[m] + H2O[m] <=> isocitrate[m] | Tricarboxylic acid cycle and glyoxylate/dicarboxylate metabolism | con | 167.3841 | null | 1 | 0 | 49 | null | 455.8593 | null |
| HMR_4584 | ATP[c] + creatine[c] <=> ADP[c] + creatine-phosphate[c] + 2 H+[c] | Glycine, serine and threonine metabolism | both | -47.82268 | -39.81188 | 1 | 1 | 432 | 489 | 26.1345 | 21.77192 |
| HMR_4533 | CMP-neuNGc[c] + H2O[c] + NAD+[c] => CMP-N-acetylneuraminate[c] + H+[c] + NADH[c] + O2[c] | Amino sugar and nucleotide sugar metabolism | con | 219.7083 | null | 1 | 0 | 354 | null | 176.34 | null |
| HMR_4532 | CMP-neuNGc[c] + PPi[c] <=> CTP[c] + neungc[c] | Amino sugar and nucleotide sugar metabolism | con | -501.9517 | null | 1 | 0 | 354 | null | 188.4044 | null |
| HMR_4531 | H+[c] + N-acetylneuraminate[c] + NADH[c] + O2[c] <=> H2O[c] + NAD+[c] + neungc[c] | Amino sugar and nucleotide sugar metabolism | con | 501.9543 | null | 1 | 0 | 354 | null | 188.4044 | null |
| HMR_4461 | (R)-3-hydroxybutanoate[m] + NAD+[m] <=> acetoacetate[m] + H+[m] + NADH[m] | Butanoate metabolism | both | 232.3723 | 118.8764 | 1 | 1 | 350 | 350 | 195.3622 | 124.6016 |
| HMR_4458 | cis-aconitate[m] + H2O[m] <=> citrate[m] | Tricarboxylic acid cycle and glyoxylate/dicarboxylate metabolism | con | -167.3838 | null | 1 | 0 | 49 | null | 455.8594 | null |
| HMR_4410 | fumarate[m] + H2O[m] <=> malate[m] | Tricarboxylic acid cycle and glyoxylate/dicarboxylate metabolism | un | 0 | 971.125 | 0 | 1 | null | 503 | 0 | 27.53557 |
| HMR_4408 | fumarate[c] + H2O[c] <=> malate[c] | Tricarboxylic acid cycle and glyoxylate/dicarboxylate metabolism | un | 0 | -274.0191 | 0 | 1 | null | 503 | 0 | 41.91592 |
| HMR_4391 | DHAP[c] <=> GAP[c] | Glycolysis / Gluconeogenesis | both | 440.411 | 447.612 | 1 | 1 | 383 | 388 | 29.72294 | 25.03954 |
| HMR_4388 | H+[c] + NADH[c] + pyruvate[c] <=> L-lactate[c] + NAD+[c] | Glycolysis / Gluconeogenesis | both | 64.69543 | 190.4681 | 1 | 2 | 228 | 366,402 | 352.8025 | 304.3996 |
| HMR_4323 | H2S[c] + serine[c] => cysteine[c] + H2O[c] | Cysteine and methionine metabolism | both | 374.7793 | 302.374 | 1 | 1 | 379 | 384 | 62.05165 | 67.82907 |
| HMR_4321 | fructose[c] <=> fructose[s] | Transport, extracellular | both | -121.0896 | -121.9996 | 1 | 1 | 385 | 386 | 50.28198 | 50.44107 |
| HMR_4299 | glucosamine-6-phosphate[c] + H2O[c] => fructose-6-phosphate[c] + NH3[c] | Amino sugar and nucleotide sugar metabolism | both | 58.00473 | 62.91982 | 1 | 1 | 426 | 415 | 25.46895 | 27.47509 |
| HMR_4291 | ADP[c] + ITP[c] <=> ATP[c] + IDP[c] | Purine metabolism | both | -113.6286 | -115.2181 | 1 | 1 | 428 | 426 | 48.86804 | 49.27855 |
| HMR_4280 | H+[m] + NADH[m] + pyruvate[m] <=> L-lactate[m] + NAD+[m] | Pyruvate metabolism | both | 266.2514 | 409.8723 | 1 | 1 | 133 | 406 | 310.4338 | 283.3704 |
| HMR_4262 | nicotinamide D-ribonucleotide[c] + PPi[c] <=> nicotinamide[c] + PRPP[c] | Nicotinate and nicotinamide metabolism | un | 0 | 955.21 | 0 | 1 | null | 472 | 0 | 18.92855 |
| HMR_4259 | H+[c] + H2O[c] + nicotinamide D-ribonucleotide[c] <=> NH3[c] + nicotinate ribonucleotide[c] | Nicotinate and nicotinamide metabolism | un | 0 | -956.5513 | 0 | 1 | null | 472 | 0 | 18.91142 |
| HMR_4254 | H+[c] + nicotinate[c] + PRPP[c] <=> nicotinate ribonucleotide[c] + PPi[c] | Nicotinate and nicotinamide metabolism | un | 0 | 958.6225 | 0 | 1 | null | 472 | 0 | 18.87359 |
| HMR_4210 | PEP[c] + UDP[c] => pyruvate[c] + UTP[c] | Pyrimidine metabolism | both | 282.1577 | 284.3413 | 1 | 1 | 414 | 399 | 225.2533 | 226.3698 |

| | | | | | | | | | | | |
|---|---|---|---|---|---|---|---|---|---|---|---|
| HMR_4171 | GDP[c] + PEP[c] => GTP[c] + pyruvate[c] | Purine metabolism | both | 250.2913 | 252.369 | 1 | 1 | 361 | 365 | 209.0935 | 210.3233 |
| HMR_4145 | citrate[m] + CoA[m] <=> acetyl-CoA[m] + H2O[m] + OAA[m] | Tricarboxylic acid cycle and glyoxylate/dicarboxylate metabolism | un | 0 | -537.2401 | 0 | 1 | null | 489 | 0 | 67.75637 |
| HMR_4141 | H+[m] + NADH[m] + OAA[m] <=> malate[m] + NAD+[m] | Tricarboxylic acid cycle and glyoxylate/dicarboxylate metabolism | both | -256.2166 | -195.1828 | 1 | 2 | 268 | 256,489 | 268.6583 | 269.9385 |
| HMR_4139 | H+[c] + NADH[c] + OAA[c] <=> malate[c] + NAD+[c] | Tricarboxylic acid cycle and glyoxylate/dicarboxylate metabolism | both | -330.2298 | -406.0951 | 2 | 1 | 268,414 | 256 | 266.5152 | 269.2388 |
| HMR_4137 | CoA[m] + NAD+[m] + pyruvate[m] => acetyl-CoA[m] + CO2[m] + H+[m] + NADH[m] | Glycolysis / Gluconeogenesis | both | 546.6356 | 380.6548 | 1 | 1 | 344 | 359 | 161.3396 | 154.1884 |
| HMR_4101 | GTP[c] + OAA[c] => CO2[c] + GDP[c] + PEP[c] | Pyruvate metabolism | con | 60.40496 | null | 1 | 0 | 414 | null | 50.34632 | null |
| HMR_4089 | malate[c] + NADP+[c] => CO2[c] + H+[c] + NADPH[c] + pyruvate[c] | Pyruvate metabolism | both | 55.68775 | 67.76298 | 1 | 1 | 414 | 489 | 47.67091 | 54.25878 |
| HMR_3971 | fumarate[m] + Pi[c] <=> fumarate[c] + Pi[m] | Transport, mitochondrial | un | 0 | -88.07946 | 0 | 1 | null | 489 | 0 | 529.8349 |
| HMR_3958 | isocitrate[m] + NADP+[m] => AKG[m] + CO2[m] + H+[m] + NADPH[m] | Tricarboxylic acid cycle and glyoxylate/dicarboxylate metabolism | both | 199.5893 | 201.4573 | 1 | 1 | 368 | 369 | 152.2816 | 152.7473 |
| HMR_3957 | isocitrate[m] + NAD+[m] => AKG[m] + CO2[m] + H+[m] + NADH[m] | Tricarboxylic acid cycle and glyoxylate/dicarboxylate metabolism | both | 190.8248 | 188.8379 | 1 | 1 | 368 | 369 | 148.8246 | 147.7644 |
| HMR_3952 | THF[c] <=> THF[m] | Transport, mitochondrial | both | -668.9211 | -619.5795 | 1 | 1 | 405 | 429 | 142.1801 | 155.8188 |
| HMR_3950 | 5,10-methylene-THF[c] <=> 5,10-methylene-THF[m] | Transport, mitochondrial | both | 392.9233 | 297.1823 | 1 | 1 | 405 | 429 | 40.10535 | 42.46222 |
| HMR_3949 | malate[m] + OAA[c] => malate[c] + OAA[m] | Transport, mitochondrial | both | 421.4984 | 455.3691 | 2 | 1 | 268,377 | 256 | 259.3567 | 262.8565 |
| HMR_3923 | 5,10-methylene-THF[m] + CO2[m] + H+[m] + NADH[m] + NH3[m] => glycine[m] + NAD+[m] + THF[m] | Folate metabolism | both | 808.6485 | 774.7948 | 1 | 1 | 374 | 370 | 132.0687 | 154.1109 |
| HMR_3899 | AKG[c] + alanine[c] <=> glutamate[c] + pyruvate[c] | Alanine, aspartate and glutamate metabolism | con | -668.7194 | null | 1 | 0 | 376 | null | 73.47571 | null |
| HMR_3867 | formaldehyde[c] <=> formaldehyde[p] | Transport, peroxisomal | both | -124.1606 | -114.2267 | 1 | 1 | 453 | 385 | 43.91719 | 39.51125 |
| HMR_3864 | 3-sulfinoalanine[m] + glutamate[c] + H+[c] => 3-sulfinoalanine[c] + glutamate[m] + H+[m] | Transport, mitochondrial | both | 58.05902 | 48.59677 | 1 | 1 | 421 | 431 | 48.02862 | 39.94689 |
| HMR_3863 | 3-sulfinoalanine[c] + aspartate[m] <=> 3-sulfinoalanine[m] + aspartate[c] | Transport, mitochondrial | both | 92.20881 | 75.36047 | 1 | 1 | 421 | 431 | 50.40377 | 41.76933 |
| HMR_3859 | D-lactate[c] + 2 ferricytochrome C[m] => 2 ferrocytochrome C[m] + 2 H+[c] + pyruvate[c] | Pyruvate metabolism | un | 0 | 57.96077 | 0 | 1 | null | 402 | 0 | 46.25137 |
| HMR_3850 | sarcosine[c] <=> sarcosine[p] | Transport, peroxisomal | both | 124.1594 | 114.2256 | 1 | 1 | 453 | 385 | 43.91733 | 39.51141 |
| HMR_3849 | H2O[p] + O2[p] + sarcosine[p] => formaldehyde[p] + glycine[p] + H2O2[p] | Glycine, serine and threonine metabolism | both | 124.1594 | 114.2256 | 1 | 1 | 453 | 385 | 43.91722 | 39.51129 |
| HMR_3848 | glycine[c] <=> glycine[p] | Transport, peroxisomal | both | -124.1591 | -114.2253 | 1 | 1 | 453 | 385 | 43.91732 | 39.5114 |
| HMR_3845 | serine[c] + THF[c] <=> 5,10-methylene-THF[c] + glycine[c] + H2O[c] | Glycine, serine and threonine metabolism | both | -462.3558 | -403.3471 | 1 | 1 | 405 | 429 | 49.85803 | 51.13618 |
| HMR_3835 | 1-pyrroline-5-carboxylate[c] + H+[c] + NADH[c] => NAD+[c] + proline[c] | Arginine and proline metabolism | both | 89.35603 | 50.50736 | 1 | 1 | 397 | 452 | 71.41064 | 41.4021 |
| HMR_3833 | 1-pyrroline-5-carboxylate[c] + H+[c] + NADPH[c] => NADP+[c] + proline[c] | Arginine and proline metabolism | both | 81.14296 | 47.71352 | 1 | 1 | 397 | 452 | 67.10185 | 39.84461 |
| HMR_3829 | AKG[c] + aspartate[c] <=> glutamate[c] + OAA[c] | Alanine, aspartate and glutamate metabolism | con | 149.8874 | null | 1 | 0 | 377 | null | 77.65659 | null |
| HMR_3827 | AKG[m] + aspartate[m] <=> glutamate[m] + OAA[m] | Alanine, aspartate and glutamate metabolism | con | -147.5874 | null | 1 | 0 | 377 | null | 77.11972 | null |
| HMR_3825 | aspartate[m] + glutamate[c] + H+[c] => aspartate[c] + glutamate[m] + H+[m] | Transport, mitochondrial | both | 61.18769 | 51.29637 | 2 | 1 | 421,377 | 431 | 59.32846 | 49.96697 |
| HMR_3822 | 1-pyrroline-5-carboxylate[m] + 2 H2O[m] + NAD+[m] => glutamate[m] + H+[m] + NADH[m] | Alanine, aspartate and glutamate metabolism | both | 120.5889 | 61.80814 | 1 | 1 | 380 | 408 | 85.33154 | 45.43376 |
| HMR_3820 | 1-pyrroline-5-carboxylate[m] + H2O[m] <=> L-glutamate 5-semialdehyde[m] | Arginine and proline metabolism | both | -783.3651 | -901.6278 | 2 | 2 | 381,380 | 381,408 | 107.3453 | 58.66031 |
| HMR_3819 | 1-pyrroline-5-carboxylate[c] + H2O[c] <=> L-glutamate 5-semialdehyde[c] | Arginine and proline metabolism | both | -153.989 | -83.12415 | 1 | 1 | 381 | 381 | 80.56197 | 48.47073 |
| HMR_3818 | L-glutamate 5-semialdehyde[c] <=> L-glutamate 5-semialdehyde[m] | Transport, mitochondrial | both | -153.9887 | -83.12389 | 1 | 1 | 381 | 381 | 80.56199 | 48.47078 |
| HMR_3806 | glutamate[m] + H+[m] + NADH[m] <=> H2O[m] + L-glutamate 5-semialdehyde[m] + NAD+[m] | Arginine and proline metabolism | both | 870.223 | 926.9322 | 1 | 1 | 380 | 408 | 88.03456 | 48.45246 |
| HMR_3763 | CoA[m] + methylmalonate[m] => (R)-methylmalonyl-CoA[m] + H2O[m] | Valine, leucine and isoleucine degradation | un | 0 | 186.7176 | 0 | 1 | null | 377 | 0 | 94.66132 |
| HMR_3761 | 2-methyl-3-oxopropanoate[m] + CoA[m] + NAD+[m] => (R)-methylmalonyl-CoA[m] + H+[m] + NADH[m] | Valine, leucine, and isoleucine metabolism | un | 0 | 138.8494 | 0 | 2 | null | 377,489 | 0 | 95.58202 |
| HMR_3759 | 2-methyl-3-oxopropanoate[m] + H2O[m] + NAD+[m] => H+[m] + methylmalonate[m] + NADH[m] | Valine, leucine and isoleucine degradation | un | 0 | 149.7057 | 0 | 1 | null | 377 | 0 | 94.88936 |
| HMR_3757 | 3-hydroxyisobutyrate[m] + NAD+[m] => 2-methyl-3-oxopropanoate[m] + H+[m] + NADH[m] | Valine, leucine, and isoleucine metabolism | un | 0 | 330.9161 | 0 | 1 | null | 489 | 0 | 53.1754 |
| HMR_3755 | 3-hydroxyisobutyryl-CoA[m] + H2O[m] => 3-hydroxyisobutyrate[m] + CoA[m] | Valine, leucine and isoleucine degradation | un | 0 | 330.9156 | 0 | 1 | null | 489 | 0 | 53.1754 |
| HMR_3753 | H2O[m] + methacrylyl-CoA[m] => 3-hydroxyisobutyryl-CoA[m] | Valine, leucine and isoleucine degradation | un | 0 | 330.9193 | 0 | 1 | null | 489 | 0 | 53.17541 |
| HMR_3748 | 3-methyl-2-oxobutyrate[m] + CoA[m] + NAD+[m] => CO2[m] + H+[m] + isobutyryl-CoA[m] + NADH[m] | Valine, leucine and isoleucine degradation | un | 0 | 173.2033 | 0 | 2 | null | 372,489 | 0 | 99.66492 |
| HMR_3747 | AKG[c] + valine[c] => 3-methyl-2-oxobutyrate[c] + glutamate[c] | Valine, leucine, and isoleucine metabolism | un | 0 | 317.6482 | 0 | 1 | null | 489 | 0 | 53.59684 |
| HMR_3746 | 3-methyl-2-oxobutyrate[c] <=> 3-methyl-2-oxobutyrate[m] | Transport, mitochondrial | un | 0 | 317.6473 | 0 | 1 | null | 489 | 0 | 53.59676 |
| HMR_3215 | (R)-methylmalonyl-CoA[m] => succinyl-CoA[m] | Valine, leucine, and isoleucine metabolism | un | 0 | 337.834 | 0 | 1 | null | 489 | 0 | 53.28529 |
| HMR_3106 | acetyl-CoA[c] <=> acetyl-CoA[p] | Beta oxidation of even-chain fatty acids (peroxisomal) | both | -196.9886 | -101.876 | 1 | 1 | 22 | 21 | 372.2204 | 399.3254 |
| HMR_3030 | CoA[p] + O-acetylcarnitine[p] <=> acetyl-CoA[p] + L-carnitine[p] | Carnitine shuttle (peroxisomal) | both | 113.8137 | 21.28723 | 1 | 1 | 22 | 21 | 372.2673 | 399.3347 |
| HMR_3029 | L-carnitine[c] + O-acetylcarnitine[p] <=> L-carnitine[p] + O-acetylcarnitine[c] | Carnitine shuttle (peroxisomal) | both | -113.8137 | -21.28712 | 1 | 1 | 22 | 21 | 372.2674 | 399.3348 |
| HMR_2593 | CoA[m] + O-acetylcarnitine[m] <=> acetyl-CoA[m] + L-carnitine[m] | Carnitine shuttle (mitochondrial) | both | -557.3183 | -368.2912 | 1 | 1 | 423 | 390 | 113.4018 | 111.053 |
| HMR_2592 | L-carnitine[c] + O-acetylcarnitine[m] <=> L-carnitine[m] + O-acetylcarnitine[c] | Carnitine shuttle (mitochondrial) | both | 557.3149 | 368.2881 | 1 | 1 | 423 | 390 | 113.4013 | 111.0526 |
| HMR_2591 | CoA[c] + O-acetylcarnitine[c] <=> acetyl-CoA[c] + L-carnitine[c] | Carnitine shuttle (cytosolic) | both | 443.498 | 346.9992 | 2 | 2 | 423,22 | 390,21 | 373.0087 | 402.6875 |
| HMR_1327 | H+[c] + NADPH[c] + prostaglandin E2[c] <=> NADP+[c] + prostaglandin F2alpha[c] | Prostaglandin biosynthesis | un | 0 | 412.6855 | 0 | 1 | null | 331 | 0 | 239.597 |
| HMR_1325 | H+[c] + NADH[c] + prostaglandin H2[c] <=> NAD+[c] + prostaglandin F2alpha[c] | Prostaglandin biosynthesis | un | 0 | 60.11558 | 0 | 1 | null | 331 | 0 | 481.741 |
| HMR_1312 | prostaglandin H2[c] => prostaglandin E2[c] | Prostaglandin biosynthesis | un | 0 | 412.6848 | 0 | 1 | null | 331 | 0 | 239.597 |
| HMR_1284 | 10,11-dihydro-12-epi-LTB4[c] <=> 10,11-dihydro-12-epi-LTB4[m] | Transport, mitochondrial | un | 0 | 491.6877 | 0 | 1 | null | 489 | 0 | 20.1913 |
| HMR_1281 | 10,11-dihydro-12-epi-LTB4[m] + NADP+[m] <=> 10,11-dihydro-12-oxo-LTB4[m] + H+[m] + NADPH[m] | Leukotriene metabolism | un | 0 | 491.6877 | 0 | 1 | null | 489 | 0 | 20.19122 |
| HMR_1267 | 10,11-dihydro-LTB4-CoA[c] <=> 10,11-dihydro-LTB4-CoA[m] | Transport, mitochondrial | un | 0 | -491.6879 | 0 | 1 | null | 489 | 0 | 20.19099 |

| | | | | | | | | | | | |
|---|---|---|---|---|---|---|---|---|---|---|---|
| HMR_1264 | 10,11-dihydro-LTB4-CoA[m] + AMP[m] + PPi[m] <=> 10,11-dihydro-LTB4[m] + ATP[m] + CoA[m] | Leukotriene metabolism | un | 0 | -491.6876 | 0 | 1 | null | 489 | 0 | 20.19093 |
| HMR_1263 | 10,11-dihydro-LTB4-CoA[c] + AMP[c] + PPi[c] <=> 10,11-dihydro-LTB4[c] + ATP[c] + CoA[c] | Leukotriene metabolism | un | 0 | 677.045 | 0 | 1 | null | 489 | 0 | 91.46397 |
| HMR_1232 | 10,11-dihydro-12-oxo-LTB4[m] + H+[m] + NADPH[m] => 10,11-dihydro-LTB4[m] + NADP+[m] | Leukotriene metabolism | un | 0 | 491.6876 | 0 | 1 | null | 489 | 0 | 20.19109 |
| HMR_0454 | ATP[c] + glyceraldehyde[c] => ADP[c] + GAP[c] | Fructose and Mannose metabolism | both | 81.73443 | 65.33471 | 1 | 1 | 383 | 388 | 55.3203 | 44.82028 |
| HMR_0453 | glyceraldehyde[c] + H+[c] + NADPH[c] <=> glycerol[c] + NADP+[c] | Glycerolipid metabolism | both | -85.25277 | -68.96361 | 1 | 1 | 383 | 388 | 55.32292 | 44.82262 |
| HMR_0444 | glycerol[c] <=> glycerol[s] | Transport, extracellular | both | -69.82899 | -53.09124 | 1 | 1 | 383 | 388 | 55.57638 | 45.1565 |
| HMR_0016 | 5.1e-05 (10Z)-heptadecenoic acid[s] + 0.002293 (11Z,14Z)-eicosadienoic acid[s] + 0.000266 (11Z,14Z,17Z)-eicosatrienoic acid[s] + 0.000175 (13Z)-eicosenoic acid[s] + 0.002986 (13Z)-octadecenoic acid[s] + 4e-05 (13Z,16Z)-docosadienoic acid[s] + 8.2e-05 (4Z,7Z,10Z,13Z,16Z)-DPA[s] + 4.9e-05 (6Z,9Z)-octadecadienoic acid[s] + 3.8e-05 (6Z,9Z,12Z,15Z,18Z)-TPA[s] + 3.8e-05 (6Z,9Z,12Z,15Z,18Z,21Z)-THA[s] + 4.8e-05 (7Z)-octadecenoic acid[s] + 0.000361 (7Z) tetradecenoic acid[s] + 0.002284 (9E)-tetradecenoic acid[s] + 3.8e-05 (9Z,12Z,15Z,18Z)-TTA[s] + 3.8e-05 (9Z,12Z,15Z,18Z,21Z)-TPA[s] + 4.1e-05 10,13,16,19-docosatetraenoic acid[s] + 4.1e-05 10,13,16-docosatriynoic acid[s] + 3.8e-05 12,15,18,21-tetracosatetraenoic acid[s] + 4.1e-05 13,16,19-docosatrienoic acid[s] + 0.004492 7-palmitoleic acid[s] + 0.000176 8,11-eicosadienoic acid[s] + 0.000964 9-eicosenoic acid[s] + 0.003649 9-heptadecylenic acid[s] + 0.000818 adrenic acid[s] + 0.005094 arachidonate[s] + 0.00016 behenic acid[s] + 3.4e-05 cerotic acid[s] + 8e-05 cis-cetoleic acid[s] + 0.000241 cis-erucic acid[s] + 0.004644 cis-gondoic acid[s] + 0.02042 cis-vaccenic acid[s] + 0.00116 DHA[s] + 0.002131 dihomo-gamma-linolenate[s] + 0.000906 DPA[s] + 0.000871 eicosanoate[s] + 4.8e-05 elaidate[s] + 0.0009 EPA[s] + 0.000489 gamma-linolenate[s] + 4.2e-05 henicosanoic acid[s] + 0.006112 lauric acid[s] + 3.7e-05 lignocerate[s] + 0.15726 linoleate[s] + 0.007329 linolenate[s] + 5e-05 margaric acid[s] + 4.4e-05 mead acid[s] + 0.035025 myristic acid[s] + 3.7e-05 nervonic acid[s] + 4.6e-05 nonadecylic acid[s] + 0.38306 oleate[s] + 0.000357 omega-3-arachidonic acid[s] + 0.24944 palmitate[s] + 0.039461 palmitolate[s] + 0.003928 pentadecylic acid[s] + 6e-05 physeteric acid[s] + 0.061303 stearate[s] + 4.9e-05 stearidonic acid[s] + 3.8e-05 tricosanoic acid[s] + 6.3e-05 tridecylic acid[s] + 3.4e-05 ximenic acid[s] => NEFA blood pool out[s] | Transport, extracellular | both | 95.28182 | 88.86312 | 1 | 1 | 402 | 409 | 40.72327 | 37.90539 |
| HMR_0015 | NEFA blood pool in[s] => 0.004805 (10Z)-heptadecenoic acid[s] + 0.001642 (11Z,14Z)-eicosadienoic acid[s] + 0.001591 (11Z,14Z,17Z)-eicosatrienoic acid[s] + 9e-06 (13Z)-eicosenoic acid[s] + 9e-06 (13Z)-octadecenoic acid[s] + 5.1e-05 (13Z,16Z)-docosadienoic acid[s] + 9e-06 (4Z,7Z,10Z,13Z,16Z)-DPA[s] + 9e-06 (6Z,9Z)-octadecadienoic acid[s] + 9e-06 (6Z,9Z,12Z,15Z,18Z)-TPA[s] + 9e-06 (6Z,9Z,12Z,15Z,18Z,21Z)-THA[s] + 9e-06 (7Z)-octadecenoic acid[s] + 9e-06 (7Z)-tetradecenoic acid[s] + 9e-06 (9E)-tetradecenoic acid[s] + 9e-06 (9Z,12Z,15Z,18Z)-TTA[s] + 9e-06 (9Z,12Z,15Z,18Z,21Z)-TPA[s] + 9e-06 10,13,16,19-docosatetraenoic acid[s] + 9e-06 10,13,16-docosatriynoic acid[s] + 9e-06 12,15,18,21-tetracosatetraenoic acid[s] + 1.9e-05 13,16,19-docosatrienoic acid[s] + 9e-06 7-palmitoleic acid[s] + 9e-06 8,11-eicosadienoic acid[s] + 9e-06 9-eicosenoic acid[s] + 9e-06 9-heptadecylenic acid[s] + 0.001698 adrenic acid[s] + 0.013716 arachidonate[s] + 0.000746 behenic acid[s] + 0.000513 cerotic acid[s] + 9e-06 cis-cetoleic acid[s] + 0.000131 cis-erucic acid[s] + 9e-06 cis-gondoic acid[s] + 9e-06 cis-vaccenic acid[s] + 0.004619 DHA[s] + 0.002529 dihomo-gamma-linolenate[s] + 0.001866 DPA[s] + 0.00111 eicosanoate[s] + 9e-06 elaidate[s] + 0.002029 EPA[s] + 0.004805 gamma-linolenate[s] + 9e-06 henicosanoic acid[s] + 0.003354 lauric acid[s] + 0.001222 lignocerate[s] + 0.070911 linoleate[s] + 0.000537 linolenate[s] + 0.005598 margaric acid[s] + 0.000443 mead acid[s] + 0.028271 myristic acid[s] + 0.000327 nervonic acid[s] + 9e-06 nonadecylic acid[s] + 0.37462 oleate[s] + 9e-06 omega-3-arachidonic acid[s] + 0.29764 palmitate[s] + 0.068579 palmitolate[s] + 0.003046 pentadecylic acid[s] + 9e-06 physeteric acid[s] + 0.1031 stearate[s] + 7.5e-05 stearidonic acid[s] + 0.000154 tricosanoic acid[s] + 9e-06 tridecylic acid[s] + 9e-06 ximenic acid[s] | Transport, extracellular | both | 97.48426 | 87.989 | 1 | 1 | 402 | 409 | 38.09657 | 35.28272 |
| EXC_OUT_m5559 | valine[s] => | Exchange reactions | un | 0 | 337.2298 | 0 | 1 | null | 343 | 0 | 219.1931 |
| EXC_OUT_m4082 | leucine[s] => | Exchange reactions | un | 0 | 501.3706 | 0 | 1 | null | 323 | 0 | 252.5405 |
| EXC_OUT_m3848 | isoleucine[s] => | Exchange reactions | un | 0 | 495.9645 | 0 | 1 | null | 329 | 0 | 252.524 |
| EXC_OUT_m3413 | glucose[s] => | Exchange reactions | both | 422.4877 | 423.4071 | 1 | 1 | 426 | 332 | 245.7048 | 245.8971 |
| EXC_IN_m4849 | => proline[s] | Exchange reactions | con | 94.94933 | null | 1 | 0 | 392 | null | 72.17909 | null |

**TableS6: Table with processing time for each step in ComMet's pipeline**

| Analysis | Time |
|---|---|
| Preprocessing | ~30 minutes |
| EP sampling | ~12 minutes |
| PCA+basis rotation | ~1 minute |
| Module extraction | ~10 seconds |
| Reaction network construction | 1-2 minutes |
| ICA N optimization (bootstrapping) | ~20 hours |
| Optimized ICA run (9000 iterations with fixed seed) | ~2 hours |
